# Multiplex base editing to convert TAG into TAA codons in the human genome

**DOI:** 10.1101/2021.07.13.452007

**Authors:** Yuting Chen, Eriona Hysolli, Anlu Chen, Stephen Casper, Songlei Liu, Kevin Yang, Chenli Liu, George Church

## Abstract

Large-scale recoding has been shown to enable novel amino acids, biocontainment and viral resistance in bacteria only so far. Here we extend this to human cells demonstrating exceptional base editing to convert TAG to TAA for 33 essential genes via a single transfection, and examine base-editing genome-wide (observing ~ 40 C-to-T off-target events in essential gene exons). We also introduce GRIT, a computational tool for recoding. This demonstrates the feasibility of recoding, and multiplex editing in mammalian cells.

## Main

Human recoding is the pilot project of GP-write aiming to “write” genomes^1^. Recoding confers virus resistance^2^ (Supplementary Fig. 1a), and “blank” codons can be repurposed^3^. The *E. coli* genome has been successfully recoded by our lab and others^4–6^. Building on this previous work, we set out to explore the feasibility of genome-wide TAGs to TAAs replacement in human cells using C-to-T base editor (CBE)^7^. To automate part design, we designed Genome Recoding Informatics Toolbox (GRIT), a python-based platform for genome-scale data analysis tailored to recoding (Fig. 1a). Using GRIT, we determined that of 6888 TAG codons, 6648 are editable across the human haploid genome; 1947 are essential gene TAG codons number (of these, 1937 are editable), and also visualized the distribution of TAG sites throughout the 24 human chromosomes (Fig. 1b, Supplementary Fig. 1b and Supplementary Excel Data 1), based on GRCh38.p13 and the OGEE database^8^.

**Fig. 1.**
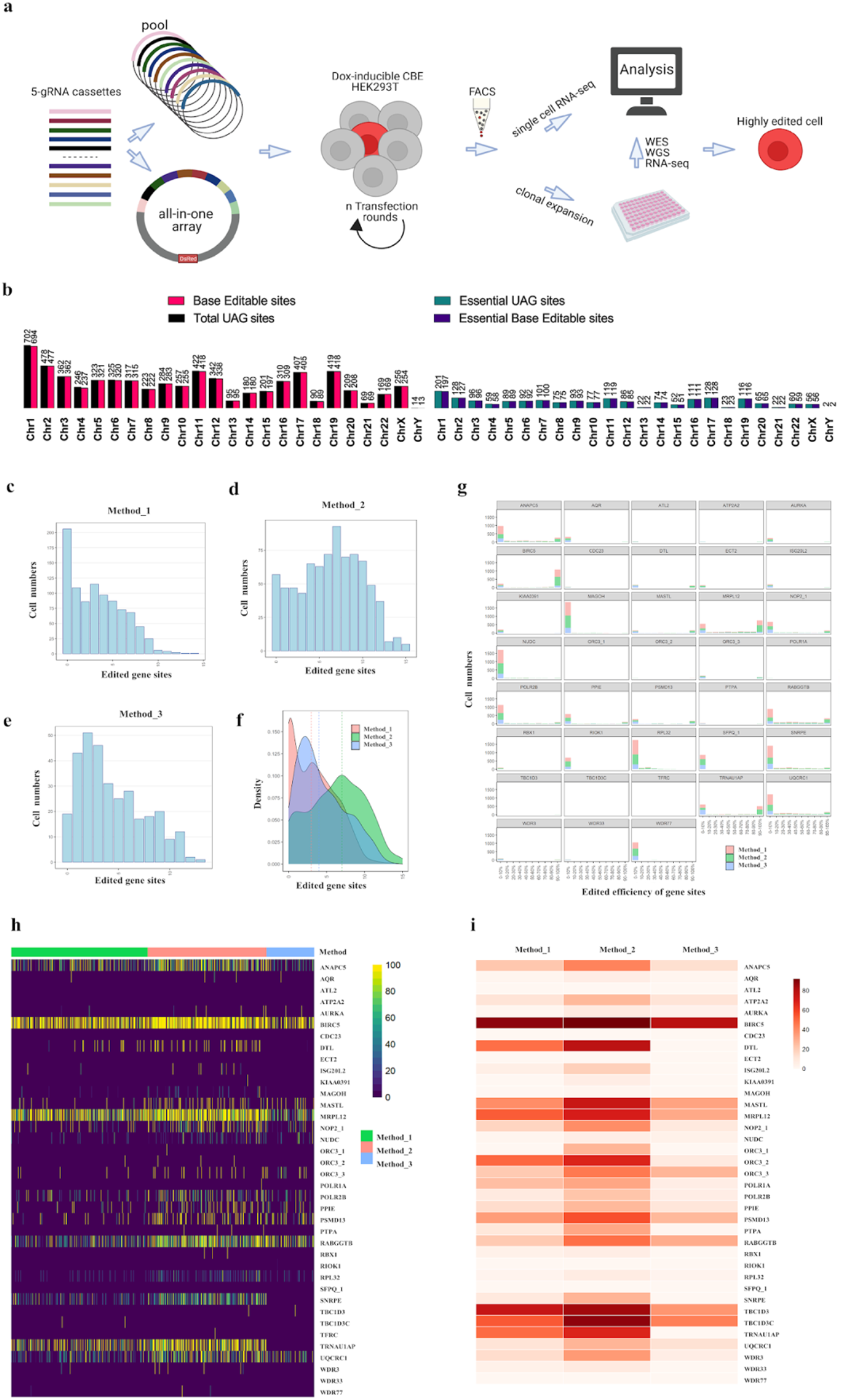
Experimental strategy for converting UAG to UAA using multiplexed base editing. a, Framework for converting TAG codons into TAA in human cells. b, UAGs number and editable UAG sites of all genes and essential genes in each chromosome. c-e, distribution analysis of cells with different number of modified gene targets in populations with different delivery methods based on single cell RNAseq. Method_1, delivery 10 gBlocks with mCherry-inactivated eGFP reporter; Method_2, delivery 10 gBlocks with mCherry-inactivated eGFP reporter and eGFP sgRNA plasmids; Method_3, delivery 43-all-in-one with DsRed. f, Density plot for distribution of number of modified gene targets detected by scRNAseq in 3 populations. Vertical lines indicate the median values of modified gene targets. g, For each gene target, distribution analysis of modified cells with different editing efficiency. Counts from different methods were stacked in the plot. h, Editing efficiency of each sgRNAs in single cells. i, Heat map of target “C” editing efficiency in cell population with three delivery methods based on converting single-cell RNA-Seq into Bulk RNA-Seq. Editing efficiency was indicated with the intensity of red.

CRISPR/Cas9, base editors and prime editing have revolutionized genome editing^9^. However, we need technologies that deliver thousands of gRNAs into a single cell for complete recoding. Firstly, we designed and synthesized gBlock containing 5 individual gRNAs cassettes: 5 previously published sgRNAs^10^ (gBlock-PC) as a control and 5 designed sgRNAs targeting TAG regions of genes (gBlock-YC1), which were transiently co-transfected with evoAPOBEC1-BE4max-NG in HEK293T cells, separately. Sanger sequencing and EditR^11^ analysis showed that the efficiency at each locus of gBlock-PC and gBlock-YC1 is ~40-50% and ~20%-50%, respectively (Supplementary Figs. 2a, b). To further improve editing efficiency from the gRNA array, we generated two doxycycline-inducible stable HEK293T lines with piggybacFNLS-BE3-NG^12^ and evoAPOBEC1-BE4max-NG (Supplementary Fig. 3a) and transiently transfected gBlock-PC or gBlock-YC1 into each of the two inducible CBE lines. The editing efficiency of gBlock-PC is ~60-70% across genes loci in the evoAPOBEC1-BE4max-NG line, which is slightly higher than ~45-65% in the FNLS-BE3-NG line. Similarly, the efficiency from the gBlock-YC1 is above ~30-75% in the evoAPOBEC1-BE4max-NG line, which is significantly higher than ~20-40% in the FNLS-BE3-NG stable line (Supplementary Fig. 3b). Thus, we decided to use the stably expressing evoAPOBEC1-BE4max-NG line in subsequent experiments.

To identify an effective method for converting TAG to TAA, we firstly transfected 10-, 20-, and 30 gBlock pools into evoAPOBEC1-BE4max-NG stable clone 1 with high editing efficiency verified by us (Supplementary Fig. 4a, b) to probe the number of gBlock cassettes that can be delivered at one time with good editing efficiency. We performed Whole Exome Sequencing (WES) and found editing efficiency at most sites of the first 52 gene sites (Supplementary Table 1) is highest when 10 gBlocks are delivered compared to 20 and 30 gBlocks (Supplementary Fig. 5a, b). We also assembled 10 gBlocks into one vector with DsRed by golden gate cloning, and screened a successful 43-gRNA array called 43-all-in-one (Supplementary Fig. 6a, b). Then, we applied the following methods: 1) Method_1: 10 gBlocks + mCherry-inactivated eGFP reporter^13^; 2) Method_2: 10 gBlocks + mCherry-inactivated eGFP reporter and eGFP cognated sgRNA plasmid^13^; 3) Method_3: 43 sgRNAs all-in-one. We sorted ~1000 single cells from each condition and performed single cell RNA-seq to examine the distribution of each targeting locus across three cell populations (Fig. 1a). QC metrics analyses of the samples are shown in Supplementary Figs. 7a, b, c. We mapped a total of 38/52 gene sites, and observed the number of cells decreased as the number of editing sites increased in all three methods and the number of cells with most edited gene sites was the highest in Method_2 (Figs. 1c-e). We plotted the population density of cells (Fig. 1f) and analyzed editing efficiency of each target and targets with editing events exhibited a bimodal distribution (Fig 1g). Editing efficiency of each mapped site in each single cell (Fig. 1h) and total editing efficiency of each target in each sample (Fig. 1i) were also analyzed. Collectively, these data show that Method_2 is the most efficient for TAG to TAA replacement.

To further investigate which method generates highly modified expandable clones, we sorted and cultured 28/96 and 24/96 single clones from populations transfected by Method_2 and Method_3, respectively. For clones from Method_2, we picked 10 well-edited loci (one from each gBlock to validate their delivery), PCR amplified them, followed by Sanger sequencing and EditR analysis, and we found 4 clones without gBlocks and 24 clones with 1-10 gBlocks, of which clone 19 contained all 10 gBlocks (Supplementary Fig. 8a). For clones from Method_3, we used 3 well-edited loci for screening and found 13 clones had no editing, and 11 clones had some edited sites, of which clones 11, 20, 21, and 24 had all 3 sites edited (Supplementary Fig. 8b). We also used Sanger sequencing in all loci in the two highly modified clones (clone 19 and 21). In clone 19, we found TAG to TAA at 33/47 genomic sites, of which 9 sites are homozygous, and 14/47 unedited, while in clone 21, we found 27/40 desired editing sites, of which 10 are homozygous TAA, and 13/40 sites unedited (Supplementary Fig. 8c). To determine whether editing efficiency could increase with subsequent transfection rounds, we transfected gBlocks into the highly modified clone 19 using Method_2 and selected clones 19-1, 19-16, and 19-21 from 22/96 clones due to higher editing (Sanger/EditR) in select loci, as compared to the original clone 19.

Next, we comprehensively assessed on- and off-target efficiencies of CBE genome-wide TAG to TAA conversion. We performed Whole Genome Sequencing (WGS) at 30X on highly modified clones (19, 21, 19-1, 19-16, 19-21) and the negative control. For on-target editing, the heat map showed 39/47 gene sites have been mapped and 28 of them are highly edited in the highly modified clones, and clones 19-1, -16, -21 showed improved editing at select loci compared to clone 19 (Fig. 2a). This result was consistent with our previous finding detected with Sanger sequencing. To find off-target events, we analyzed the single nucleotide variants (SNVs) and insertion/deletions (indels) in highly modified clones (19, 21, 19-1, 19-16, 19-21) compared to the control. After subtracting on-targets, SNVs were 23084, 70356, 35700, 42595 and 31530, respectively (Fig. 2b). Further analysis on these clones revealed 277, 805, 419, 470, 358 SNVs, respectively, were located on exons (Fig 2b), and only 33, 77, 42, 46, 40 SNVs, respectively, were located in exons of essential genes (Fig. 2c and Supplementary Fig. 9). We classified the SNVs into individual mutation types and found that C-to-T (G-to-A) transitions were the most frequent edits as expected (Fig. 2d). SNV mutation ratios were very low as seen in each clone (Fig. 2e), and spread throughout each chromosome (Fig. 2f). In addition to SNVs, the number of indels detected in these clones was 558, 715, 717, 662, 655, respectively, with a small subset located in exons (Fig. 2g) and none in exons of essential genes. Similarly, indel ratios were very low in each clone (Fig. 2h) and chromosome (Fig. 2i).

**Fig. 2.**
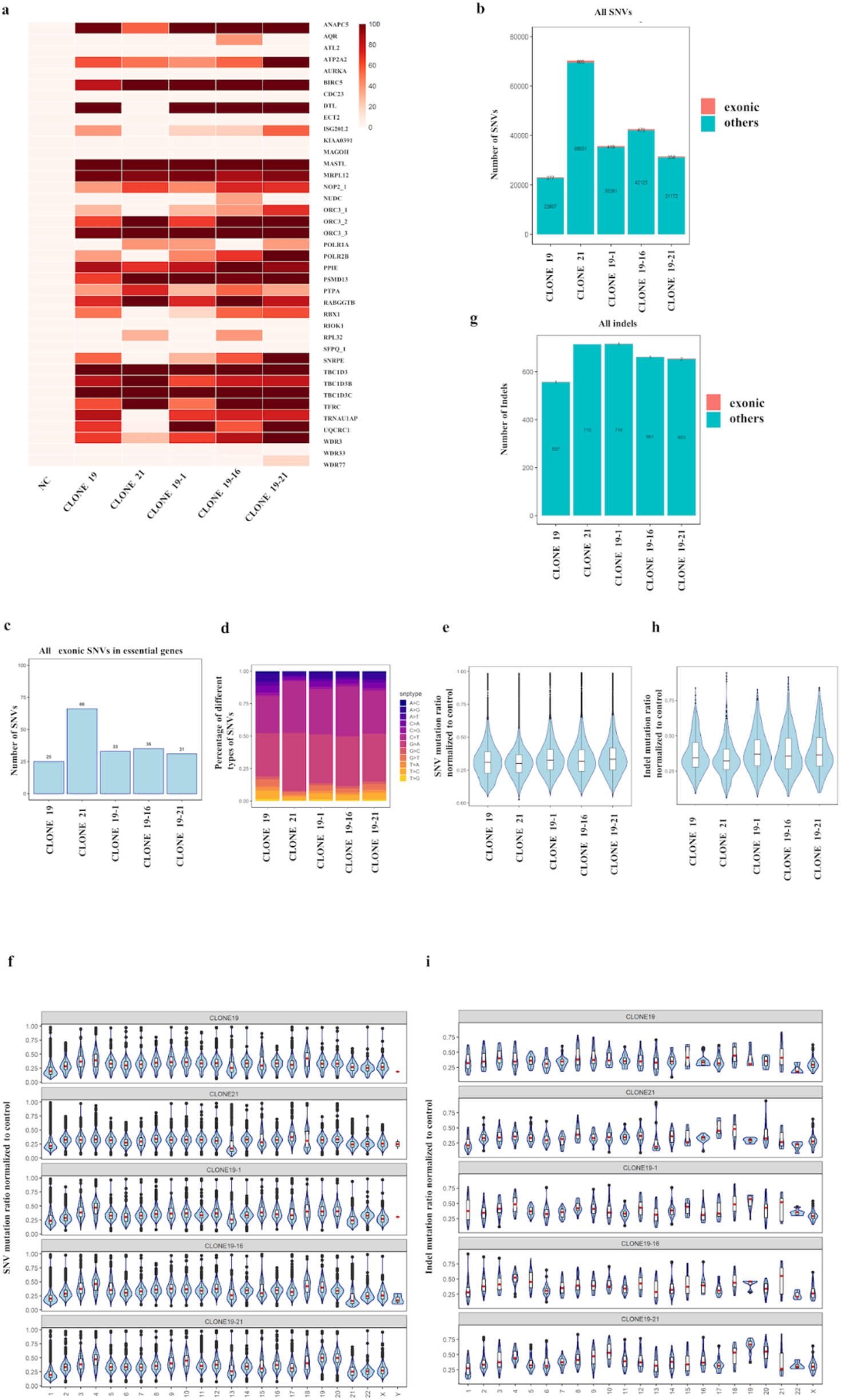
Analysis of the genetic changes of highly modified HEK293T clones identified by WGS. a, Heatmap of target “C” editing efficiency for converting TAGs to TAAs. NC, negative control, HEK293T-BE4max stable cell; clone19 from method_2, clone 21 from method_3; clone19-1, 19-16, 19-21 from second transfection by method_2. b, Number of exonic SNVs (SNVs are located on exons and splicing sites) or other SNVs detected in highly modified clones, as compared to the sequence of the parental HEK293T. The numbers of total SNVs in clone19, clone 21, clone19-1, 19-16, 19-21 were 23084, 70356, 35700, 42595 and 31530, respectively. c, Number of exonic SNVs detected in essential genes. d, distribution of different types of SNV changes. e, mutation ratio of detected C>T or G>T SNVs across samples. f, mutation ratio of detected C>T or G>T SNVs across samples & chromosomes. g, Number of exonic indels or other indels detected in highly modified clones h, mutation ratio of detected indels across samples. i, mutation ratio of detected indels across samples & chromosomes.

Finally, we examined potential gene expression changes before and after editing. We performed UMAP analysis on the single cell RNA-seq data, and did not observe cell clustering driven by a high number of edits, indicating no significant gene expression change as a result of editing (Supplementary Figs. 10, 11). Next, we analyzed the bulk RNA-seq data for highly modified clones (19, 21, and 11), lowly modified clones (5, 16) and the negative control. We performed on-target analysis, and the results were consistent with those of WGS (Supplementary Fig. 12a). Gene expression levels in highly modified clones and lowly modified clones were mostly similar in all genes (Supplementary Figs. 12b-e) and the 43 targeted loci (Supplementary Fig. 12f). A few genes were differentially expressed between highly and wild-type negative control clones (Supplementary Fig.12b), with gene names and gene expression fold change shown in more detail in Supplementary Figure 12e. So, Bulk RNA-seq is an effective method for high throughput screening of single clones with multi-site editing because it is less costly than WES and WGS, and gene expression changes can also be assessed before and after editing. We also examined whether unexpected genomic rearrangements had occurred as a result of the multiplexed genome editing. Karyotyping of individual modified clones (Supplementary Fig.13 and Supplementary Table 3) indicated that there were no observable genomic rearrangements due to multiplex editing.

From a quick calculation, in our best clone 19, in order to get 9 homozygous recoded gene termini on target per cell, we also incurred 24 heterozygote off-target mutations in 4764 (predicted) essential genes. To get 1937 precise edits “on-target”, we tolerate a Poisson average of 5165 hits (mostly C to T) in off-target 4764 essential genes (including ~1550 homozygous). If we reduce the off-target rate by 60X, then we expect less than 1 homozygous off-target gene per cell, and thus, we predict clones with zero homozygous off-targets. Certainly, some genes are more difficult to edit and therefore, CBE-based editing efficiency must significantly improve. Adenine base editors (ABE) have lower off-target burdens, but they are not suitable for recoding TAG. Our group has previously recorded the highest number of CBE-based edits (~6300) in LINE-1 repetitive elements in human cells^14^. However, a single guide RNA was designed for the highest homology among LINE-1 elements, which we predict generates a lower off-target mutation burden compared to multiple gRNAs. In the future, off-target mutations can be ameliorated with DddA-split base editors^15^.

In summary, we developed GRIT, a primary Computer Aided Design (CAD) platform for human recoding. Furthermore, our results demonstrate the feasibility of TAG to TAA in the human genome, but with an observable CBE-mediated off-target burden. We provide a framework for large-scale engineering of mammalian genomes. Once complete, genetically modified human cells will offer a unique chassis with extended functionality that could be broadly applicable for biomedicine, especially for making cell therapies or therapeutic production lines resistant to most natural viruses.

## Acknowledgements

This work (G.M.C, E.H., and Y.C.) was supported by a pilot Harvard Quadrangle Fund for Advancing Seeding Translational Research, Aging and Longevity-Related Research Fund at Harvard Medical School, and U.S Department of Energy grant DE-FG02-02ER63445. C.L. was supported by Strategic Priority Research Program of Chinese Academy of Sciences (Nos. XDPB18, XDB29050501), National Natural Science Foundation of China (Nos. 32025022, 3201101136), Shenzhen Grants (Nos. KQTD2015033117210153). We thank GenScript for gblock DNA synthesis and Chun-Ting Wu for helpful discussions.

## Author contributions

Y.C., E.H., S.C., S.L. and K.Y. performed experiments. Y.C., G.M.C, E.H., A.C., S.L., and C.L. designed the experiments and analyzed the data. Y.C., E.H., A.C., S.L., C.L. and G.M.C wrote the paper with input from all authors.

## Materials and Methods

### Computational design procedure and design rules

The Genome Recoding Informatics Toolbox (GRIT) provides a Python-based platform for working with human genome data, specifically <u>GRCh38.p13</u> (GenBank Assembly Accession GCA_000001405.28). The central functions of GRIT are to parse genome data, find TAG codons, and identify guides for base editors with NG protospacer adjacent motifs (PAMs). Using these data, GRIT identifies 6888 total TAG sites (including ones in alternate isoforms of the same genes). Of these, 6648 are editable across the human haploid genome using base editors with editing window from position 1-13. Additionally, 1947 (1937 of which are editable) of the 6888 TAG codons are in genes that do not have strong evidence of nonessentiality. It is meant for human genome recoding of TAA to TAG, but it can be used for informatics involving coding DNA sequences, chromosomal sequences, gene essentiality, multiple isoforms, multifunctional sites, base editor guides, and primers. It can easily be run from a desktop computer. Though it was only tested on human genome data, it could be repurposed for other eukaryotic species for which high-quality genome data similar to GRCh38.p13.

The key functions for GRIT are in two python files. The main file, \texttt{GRIT.py} contains sample code and functions to replicate results. There are five functions with docstrings provided for replicating results in GRIT.py: \texttt{demo}, \texttt{count_total_sites}, \texttt{count_editing_sites}, \texttt{find_genes_to_recode}, and \texttt{get_all_site_data}. Each can be run from the command line. The second file, \texttt{GRIT_utils.py}, contains a \texttt{Chromosome} class, a \texttt{Gene} class, and helper functions. Additionally, \texttt{plot_tag_sites.ipynb} can be run to reproduce Supplementary Figure1b. Inside of \texttt{GRIT_utils.py}, a chromosome object is instantiated. Sites are found that can be directly edited with a C base editor or edited with a “daisy” chain of A and C editors. See the output of \texttt{demo} to see how these are represented. When a chromosome object is instantiated, GRIT will gather data including chromosome name, wildtype sequence, recoded sequence, indices of sites to recode, base editor sites, gene objects, and edit sites that are part of different genes or different codons read in different frames and which two genes they are part of. Gene objects are generally meant to be instantiated automatically and from within the Chromosome class. When one is instantiated, GRIT gathers data including name, chromosome, strand, wildtype sequence, recoded sequence, active isoform, introns, isoform information, gene essentiality data, and recoding sites.

### Plasmids cloning

FNLS-BE3-NG was generated using the NEBuilder HIFI DNA Assembly kit (New England Biolabs(NEB) cat# E2621L) according to manufacturer’s instructions, by combining a PCR-amplified FNLS-APOBEC1 DNA from pLenti-TRE3G-FNLS-PGK-Puro (Addgene# 110847), PCR-amplified Cas9n-NG DNA from pX330-SpCas9-NG (Addgene# 117919), PCR-amplified UGI DNA from pLenti-TRE3G-FNLS-PGK-Puro (Addgene# 110847) and an NheI/PmeI-digested piggyBac dox inducible expression vector PB-TRE-Cas9^1^ including a puromycin selection marker. The evoAPOBEC1-BE4max-NG DNA from pBT375 (Addgene# 125616) were cloned between the NotI and PmeI sites of the PB-TRE-Cas9 with NotI restriction enzyme site insertion. NEB Stable Competent E. coli (NEB cat# C3040I) was used following the manufacturer’s instructions. Q5 High-Fidelity 2X Master Mix (NEB cat# M0494S) was used for all PCRs. All enzymes and buffers were obtained from New England Biolabs unless otherwise noted. Nuclease-free water (Life Technologies cat# 10977-015) was used for cloning and PCR reactions. All primers and oligos were synthesized by IDT.

### gBlocks synthesis and golden gate assembly

All gBlocks fragment containing 5 sgRNA expression cassettes with high fidelity four-base overhang pair^2^ after cutting with type IIS restriction enzyme BbsI restriction enzyme were designed and directly sent to be synthesized into PUC57 cloning plasmid by GenScript. Two oligos with BbsI cutting sites were annealed and cloned into backbone vector with CMV promoter drive fluorescent protein expression using SpeI-HF. 10 gBlocks and backbone plasmid were cutted by BbsI-HF separately, and then gel extraction using Gel extraction kit (Zymo Research cat# 11-301C). gBlocks fragment and backbone plasmid was ligated by T4 DNA ligase (NEB cat# M0202S) at 16 °C overnight. After the ligation reaction, transform the 2μl reaction mix into a competent E. coli strain NEB Stable, according to the protocol supplied with the cells. Isolate the plasmid DNA from cultures by using a QIAprep spin miniprep kit (cat#27104) according to the manufacturer’s instructions. To validate multiple sgRNAs plasmid, we firstly can roughly check whether the assembly is successful by SpeI cutting. Because There is a SpeI site on either side of the multiple sgRNAs insertion site. When multiple sgRNAs are assembled successfully in the plasmid, two bands will be seen on a gel electrophoresis after the plasmid was cut by SpeI. One band is about 4479 bp and another is about 22140 bp. Then we verified multiple sgRNAs insertion by sanger sequencing.

### Cell culture

HEK293T cells were purchased from ATCC. HEK293T cells were maintained in high-glucose Dulbecco’s Modified Eagle Medium (Gibco cat# 11965092) with 10% (v/v) fetal bovine serum (FBS, Gibco cat# 10082147), at 37 °C with 5% CO_2_ and passaged every 3–4 days, and tested for mycoplasma with Universal Mycoplasma Detection Kit (ATCC® 30-1012K™) every 4-6 weeks.

### Transient transfection

Transfection was conducted using Lipofectamine 3000 (Thermo Fisher Scientific cat# L3000015) using the protocol recommended by the manufacturer with slight modifications outlined below. 24 hours before transfection, 50,000 cells were seeded per well in a 48-well plate along with 250 μl of media. For single gBlock and base editor plasmid, a total of 1ug of DNA (750 ng of base editor plasmid, 250 ng of single gBlock plasmid) and 2 μl of Lipofectamine 3000 were used per well.

### Generation of CBE stable cell lines

24 hours before transfection, 500,000 HEK293T cells were seeded per well in a 6-well plate. A total of 4 μg of piggyBac targeting base editor plasmids were transfected with 1 μg of super transposase plasmid (SBI System Biosciences cat# PB210PA-1) using Lipofectamine 3000 following manufacturer’s instructions. After 48 h, cells were selected for puromycin (2 ug/ml). Cells were grown for 7-10 days under selection for polyclonal pools or sorted by flow cytometry into 96 wells of single cells after 5-7 days of selection for clonal cell lines. Puromycin was periodically confirmed in long-term cultures.

### Transfection of gBlocks pool and multiple sgRNAs plasmid into CBE stable cell

24 hours before transfection,100,000 cells were seeded per well in a 48-well poly-(d-lysine) plates (Corning cat# 354413) along with 300 μl of media with Doxycycline (2 ug/ml), 20 mM cyclic Pifithrin-Alpha (Stem Cell Technologies cat# 72062) and 20 ng/ml human recombinant bFGF (Stem Cell Technologies cat# 78003). For 10 gBlocks pool, 200 ng of each gBlocks and 3ul of Lipofectamine 3000 were used per well and 20 ng of green fluorescent protein as a transfection control. For 20 gBlocks pool, 150 ng of each gBlocks and 3μl of Lipofectamine 3000 were used per well and 20 ng of green fluorescent protein as a transfection control. For 30 gBlocks pool, 100 ng of each gBlocks and 3μl of Lipofectamine 3000 were used per well and 20 ng of green fluorescent protein as a transfection control. After transfection, Doxycycline was added for another 5 days and then cells were harvested for genomic DNA for analysis editing.

### Single cell RNAseq

After transfection, Doxycycline was added for another 5 days and then changed the medium without Doxycycline to continue to culture for 5 days, and then single cell isolation was performed by FACS-sorted with fluorescent protein. Library preparation for 10X Genomics single-cell RNA sequencing was performed following manufacturer’s instructions for Chromium Next GEM Single Cell 3′ Reagent Kits v3.1. Briefly, after single-cell suspension was acquired from flow sorting, cells and reagents were loaded to Chromium Next GEM Chip G with a targeted recovery of 1000 single cells per sample. Droplet generation, reverse transcription, cDNA amplification, fragmentation and adaptor ligation were conducted as protocol instructed. Sequencing of the library was performed on Illumina NovaSeq 6000 S1 flow cell (Read 1: 28 cycles, Read 2: 300 cycles, single i7 Index: 8 cycles), with a targeted depth of 300,000 reads per cell.

Raw sequencing reads were processed with Cell Ranger 5.0.0 to generate gene count matrix. Seurat R package 4.0.1 was used for downstream expression analysis. Due to variance in the sample’s sequencing depth, different cell filters were applied. Sample 1 and 2: gene number > 3000, mitochondrial gene percentage < 7; sample 3: gene number > 5000, mitochondrial gene percentage < 10. Normalization was performed using SCTransform function, with the options to regress out variance from mitochondrial gene ration and cell cycle. Principal component analysis was performed with RunPCA function. Top 40 dimensions were used to generate UMAP embedding with the RunUMAP function.

### Single cell clonal isolation

After transfection, Doxycycline was added for another 5 days and then change the medium without Doxycycline to continue to culture for 5 days, and then single cell isolation was performed by FACS-sorted with fluorescent protein, into flat bottom 96-well plates containing 100 μl of DMEM with 10% FBS and 1% Penicillin/Streptomycin per well. Sorted plates were incubated for 10-14 days until well-characterized colonies were visible, with periodic media changes performed as necessary. And then single cell clones were first detached using 20 μl Accutase (STEMCELL Technologies cat# 07920) and neutralized with 20 μl growth media, and then single cell clones were directly passaged into 24 wells with 800μl media for expansion.

### Genomic DNA extraction

For Sanger sequence, at 5 days post-transfection, cells were washed with PBS, and lysed in containing 200 μl of QuickExtract™ DNA Extraction Solution (Lucigen Cat. # QE09050) per well of 48-well plates, and genomic DNA (gDNA) was extracted using the manufacturer’s protocol. Briefly, the sorted plates were sealed, vortexed and heated at 65°C for 6 min then 98°C for 2 min. All primers for Sanger sequencing are listed in Supplementary Table 2. For whole exome sequence and whole genome sequence, DNA was extracted using the PureLink™ Genomic Plant DNA Purification Kit (Thermo Fisher cat# K183001) according to the manufacturer’s protocol.

### Western blotting

For western blot, HEK293T clone cells were lysed 5 d after Doxycycline was added using RIPA buffer supplemented with proteinase and phosphatase inhibitors. Total protein was quantified using the BCA kit (Beyotime cat# P0012). 20 μg per well of total protein was separated by electrophoresis using a 15-well 4 %-12 % Tris-Gly and transferred to a PVDF membrane for 120 min at 300 mA before blocking with 10% skimmed milk powder for 2 h at 4°C. PVDF membranes were incubated with a 1:1,000 dilution of anti-GAPDH (ABclonal, A19056) and a 1:1,000 dilution of anti-Cas9 (ABclonal, A14997)) overnight. Then, membranes were incubated with a 1:1,000 dilution of HRP Goat Anti-Rabbit IgG (H+L) (ABclonal, AS014) for 2 h and visualized using Tanon imager (Supplementary Fig. 4b).

### Whole exome sequencing and whole genome sequencing

For whole exome sequence, 1.5-5ug DNA processed with Exome Kit Agilent SureSelect XT Human All Exon V5, and sequenced with Illumina NovaSeq6000 S4 (2×150bp) at 50× coverage. Processing, sequencing and preliminary analysis conducted by Psomagen (South Korea). For whole genome sequencing, Library generation and sequencing were carried out using the Illumina Truseq Kit with 30X coverage at Beijing Genomics Institute (BGI) Hong Kong.

### Preparation of RNA libraries for bulk RNA sequencing

HEK293T cells cultured in 6-well plates were washed with PBS and harvested by adding 600 μl TRIzol (Life Technologies cat#15596026) directly to the cells. Total RNA were extracted with Direct-zol RNA Miniprep Plus kit (Zymo Research cat# R2070) following manufacturer’s protocol. RNA integrity was confirmed by the presence of 18S and 28S bands on a 2% E-Gel EX (Invitrogen cat# G402002). RNA libraries were prepared with NEBNext Poly(A) mRNA Magnetic Isolation Module (NEB cat# E7490L) and NEBNext Ultra II Directional RNA Library Prep Kit for Illumina (NEB cat# E7760L). 500 ng RNA was used at input, and the quality of final libraries were confirmed by qPCR and TapeStation. Sequencing of the libraries was performed by the Biopolymers Facility at Harvard Medical School using NovaSeq 6000.

### On-target analysis for whole exome sequencing, bulk RNA sequencing and whole genome sequencing

BWA was used to map sequencing reads to the reference human genome (hg38). Bam files were further analyzed with CRISPResso2 version 2.0.31^3^. An input gene list with 20bp gRNA sequence (no PAM included) for each target and chromosome coordinates for a 41bp mapping region with the target edits centered were generated. CRISPRessoWGS mode was applied to detect genomic changes in 52 selected regions among samples with customized usages -wc -15 -w 10 -p 5 and base edit-related usages -base_-edit --conversion_nuc_from C (--conversion_nuc_from G) --base_editor_output. “SAMPLES_QUANTIFICATION_SUMMARY.txt” was used to quantify the percentage of modified reads and “Selected_nucleotide_percentage_table_around_sgRNA.txt” was used to quantify the desired base edits (C>T or G>A) for each target. Subsequently, heat maps for percentage of modified reads and desired base edits were plotted respectively.

### Off-target analysis by whole genome sequencing

We called SNPs and indels using somatic tumor-normal approach (using a control sample as a normal, and edited samples as ‘tumor’), and two variant callers (mutect2 followed by FilterMutectCalls (from gatk package v4.2.0.0) and strelka2 v2.9.10) were applied and only variants passed filters were selected. For mutect2-called variants, reference counts and alternative counts were calculated based on tier 1 A/C/G/T counts while those for strelka2-called variants were pre-calculated. Shared variants from vcf files were selected by bedtools v2.29.2 to confirm a variant to be called (a similar approach was taken by Zuo et al.^4^ SNVs and indels were separated based on the length of reference allele and alternative allele. Annovar ^5^was used to further annotate the SNVs and indels using refGene, a gene-based annotation, to illustrate the distribution of different variant types. Whether detected variants were in essential genes were also examined.

### On-target analysis for scRNAseq

BAM files were generated from fastq files using Cell Ranger 5.0.0. BAM files were filtered for cell barcodes passed quality control and variants were called using CRISPResso2^3^ as described in “On-target analysis for Whole exome sequencing, bulk RNA sequencing and whole genome sequencing”. Edited targets were defined as targets with mapped and at least 2 reads of desired C or G. Individual cells with different numbers of edited targets were quantified and plotted to demonstrate the distribution among different delivery and enrichment methods as shown in Figure 1b-d and overlapped density plot was shown in Figure 1e. For each target, on-target editing efficiency was also plotted as shown in Figure 1f.

### Evaluate gene expression levels by RNA sequencing data analysis

STAR 2.5.2b was used for alignment of reads and quantification of gene expression. Briefly, a human genome reference index was built using genome primary assembly: (ftp://ftp.ebi.ac.uk/pub/databases/gencode/Gencode_human/release_27/GRCh38.primary_assembly.genome.fa.gz) and annotation file: (ftp://ftp.ebi.ac.uk/pub/databases/gencode/Gencode_human/release_27/gencode.v27.primary_assembly.annotation.gtf.gz) from GENCODE. Per gene counts were generated using STAR-quantMode GeneCounts. Differential gene expression analysis was performed using DESeq2 with raw counts from STAR. Genes with an adjusted p-value less than 0.05 were called differentially expressed. For figures that used transcripts per million (TPM) values, TPM counts were generated using Salmon. TPM for each gene was produced by aggregating the TPM value from all transcripts from the same gene.

### Karyotype analysis of highly modified single cell clones

Highly modified HEK293T clones (clone 19, clone 21) were expanded and karyotypically compared with the control groups and the wildtype HEK 293T. Actively growing cells were passaged 1–2 days prior to sending to BWH CytoGenomics Core Laboratory. The cells were received by the core at 60–80% confluency. Chromosomal count, variances and abnormalities were investigated.

### Statistics and reproducibility

All statistical analyses were performed on at least three biologically independent experiments using GraphPad prism9. Detailed information on exact sample sizes and experimental replicates can be found in the individual figure legends. Tests for statistically significant differences between groups were performed using a two-tailed Student’s *t*-test and all *P*< 0.05 were considered significant.

### Reporting Summary

Further information on experimental design is available in the Nature Research Reporting Summary linked to this article.

### Data availability

Sequencing data have been deposited in the NCBI Sequence Read Archive database with accession code PRJNA730314. All plasmids in this study will be available upon reasonable request.

### Code availability

Codes have been uploaded to the Github repository github.com/thestephencasper/GRIT, including code and files for reproducibility.

---

**Supplementary Figure 1.**
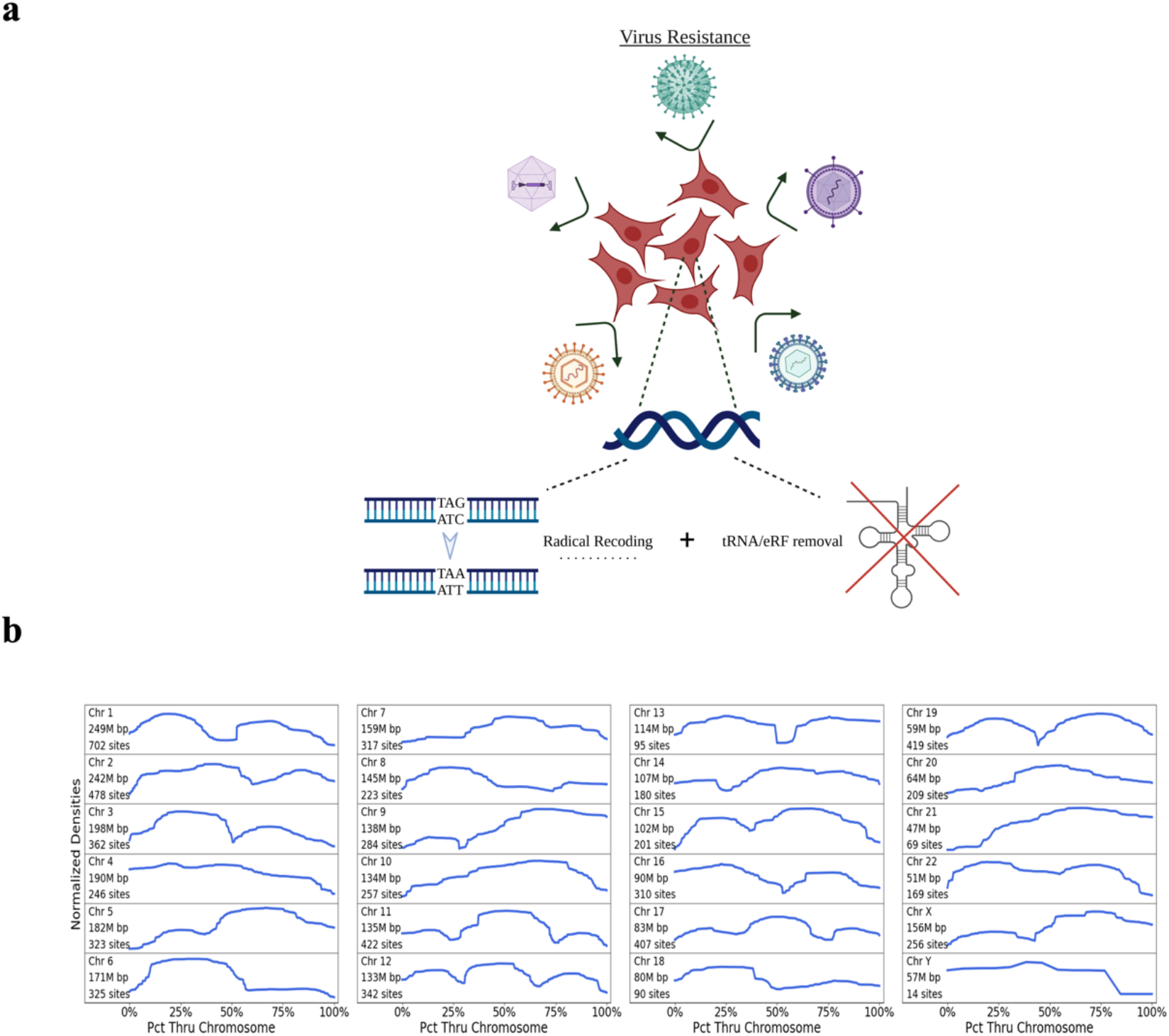
Virus resistance schematic and TAGs number count across human genome by software GRIT. (a) Virus resistance schematic: removal of redundant codons and their respective eukaryotic release factor (eRF) and/or tRNA genes in the genome. (b) Kernel density curves for the densities of TAG codons in the GRCh38.p13 build of the human genome obtained using GRIT. The density curves for the chromosomes are normalized to have uniform height and width. Chromosome lengths and total TAG counts are given on the left-hand side.

**Supplementary Figure 2.**
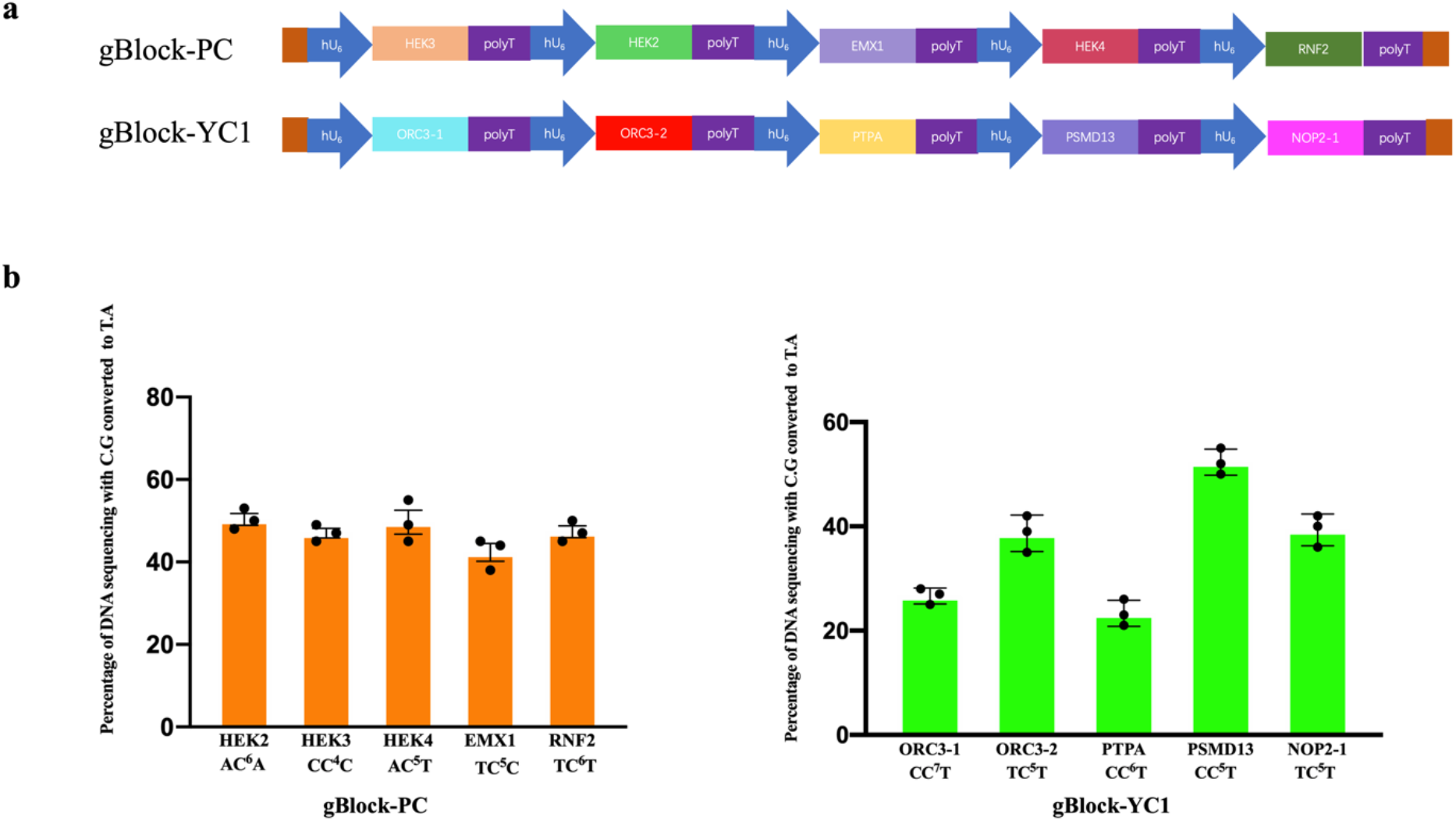
gBlocks design and editing efficiency validation. (a)Schematic diagram of gBlock-PC and gBlock-YC1. gBlock-PC carries published 5 sgRNA targeting at 5 endogenous loci (HEK2, HEK3, HEK4, EMX1, RNF2) and gBlock-YC1 carries5 sgRNA targeting at TAG of 5 genes loci (ORC3-1, ORC3-2, PTPA, PMSD13, NOP2-1). (b)Co-transfected with gblock-PC and gBlock-YC1 with evoAPOBEC-BE4max-NG into HEK293T cells separately. Frequency (%) of C-to-T conversion was obtained by Sanger sequence and editR analysis. Dots represent individual biological replicates and bars represent mean values.

**Supplementary Figure 3.**
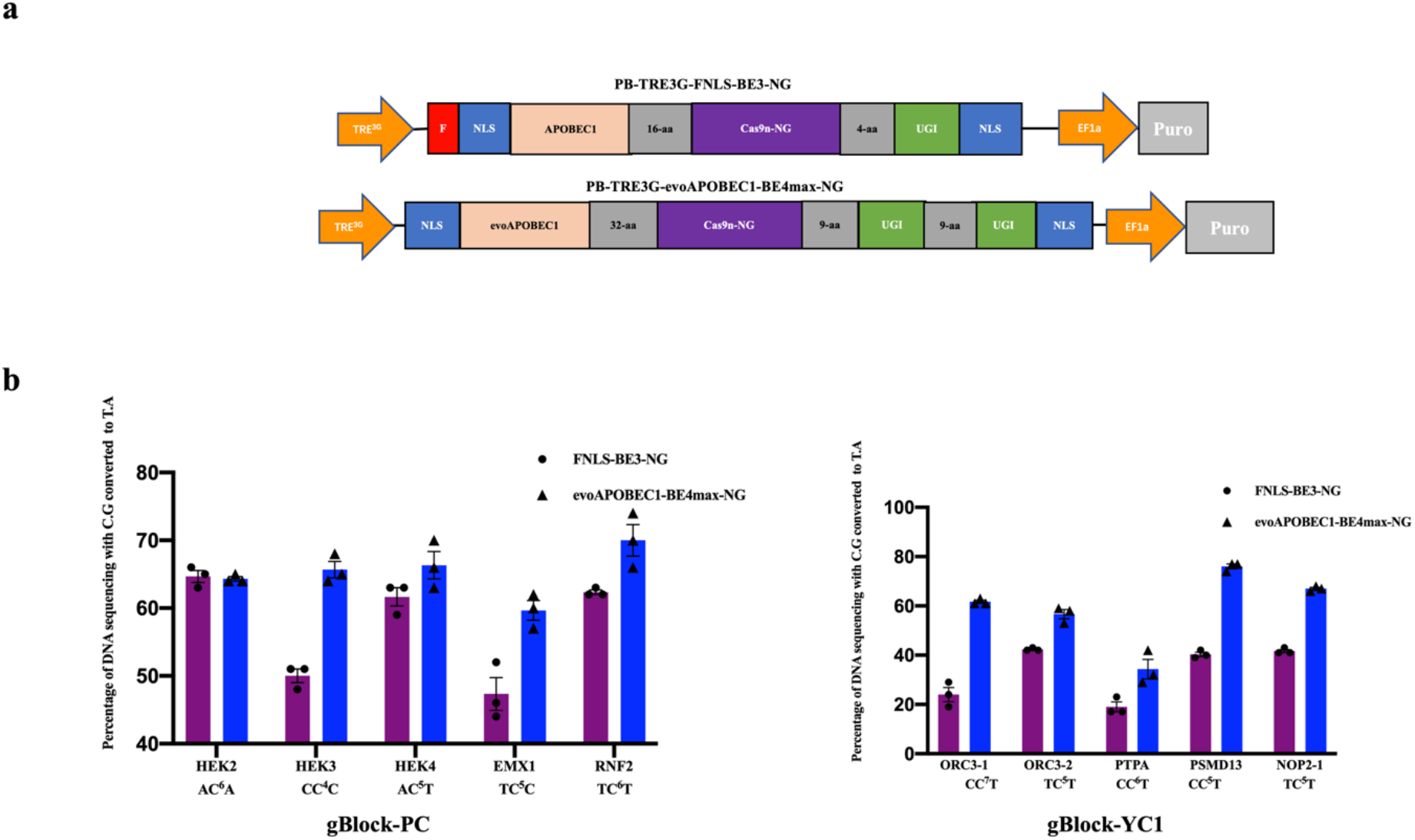
Generation of doxycycline inducible cytosine base editor stable HEK293T cell line and editing efficiency validation. (a)Schematic diagram of dox-inducible cytidine deaminase piggyBac construct. F, Flag tag; NLS, Nuclear localization signal, cas9n-NG, Cas9D10A with recognizing NG PAM. APOBAEC1, rat APOBEC1; evoAPOBAEC1, evolved rat APOBEC1. (b)Frequency (%) of C-to-T conversion in two stable HEK293T cell lines (FNLS-BE3-NG and evoAPOBEC1-BE4max-NG) transduced with gBlock-PC and gBlock-YC1 separately. Dots and triangle represent individual biological replicates and bars represent mean values.

**Supplementary Figure 4.**
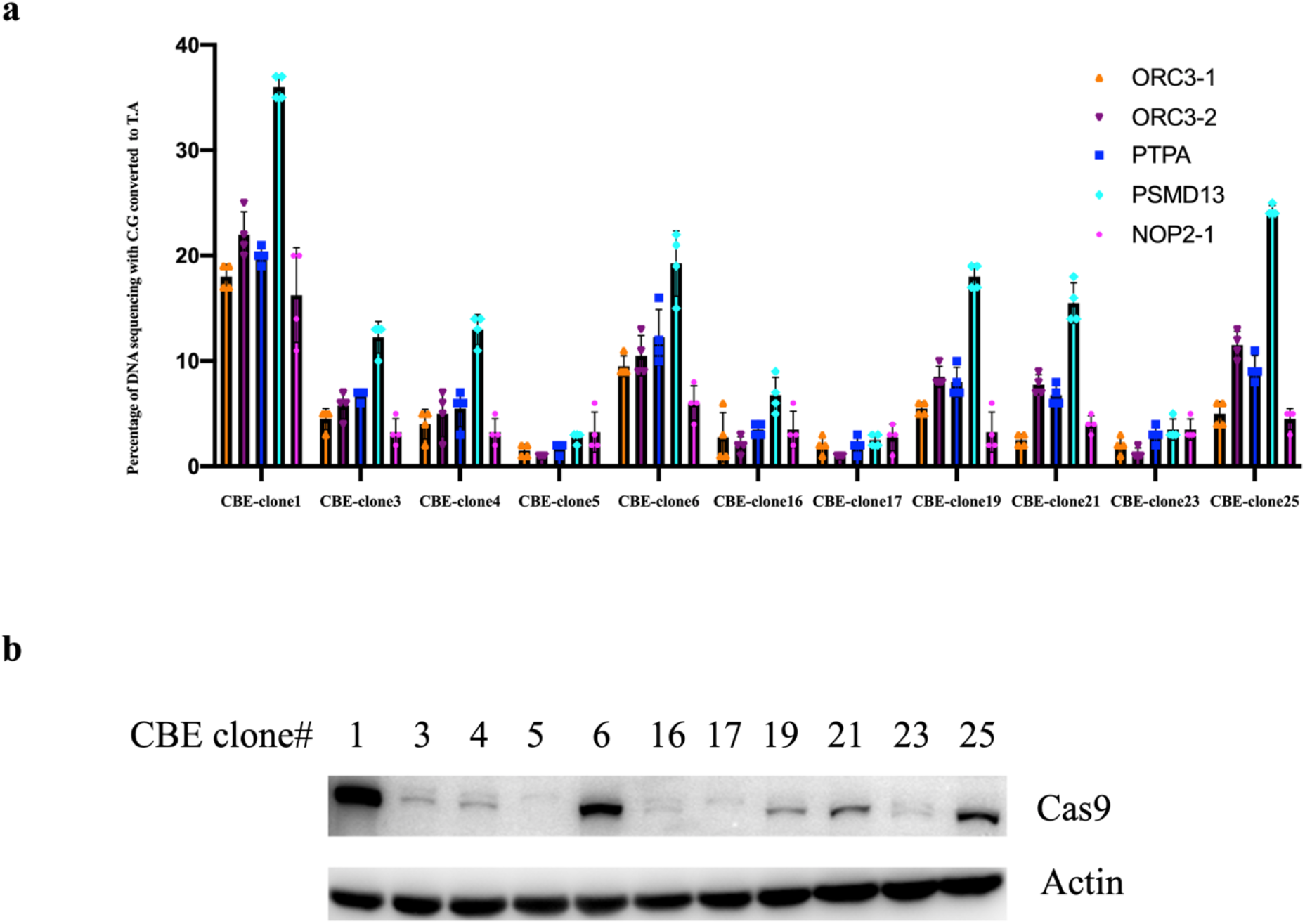
Inducible cytosine base editor (CBE) single clone screening. (a) 11 single clones from the drug resistant CBE stable cell population and transfected them with gBlock-YC1. The editing efficiency of all five sites in clone 1 was higher than that of other clones.values and error bars reflect the means and s.d. of four independent experiments. (b)The protein levels of Cytosine base editor in each clone 5 days after Doxycycline inducible. Anti-Cas9(top) and anti-Actin(bottom) were used. Western blotting images are representative of three independent experiments.

**Supplementary Figure 5.**
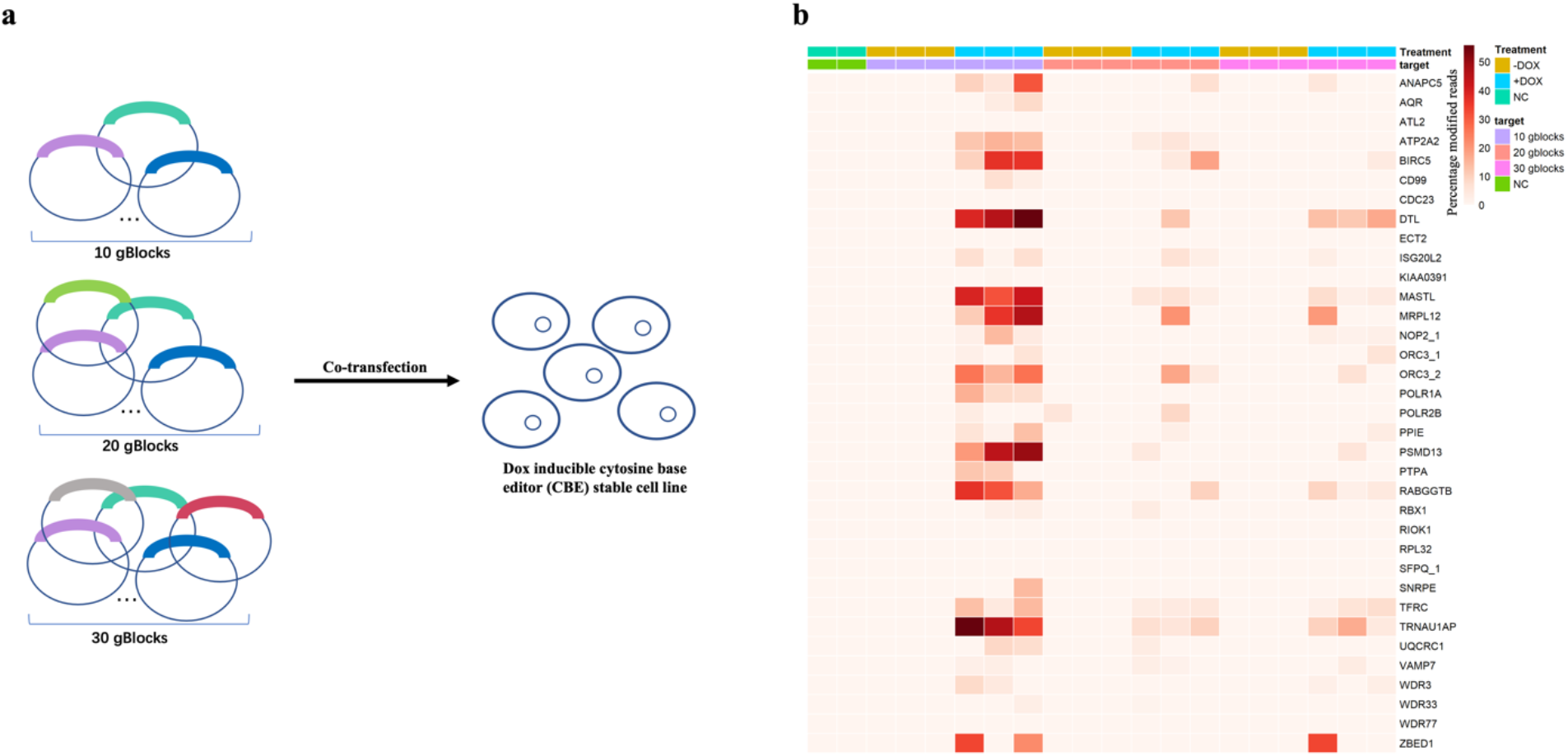
Delivery of different number of gBlocks pools into HEK293T CBE stable cell line. (a) Co-transfection of 10, 20, 30 gBlocks into HEK293T evoAPOBEC1-BE4max-NG stable cell line by lipofectamine 3000 separately. (b) Heatmap of mutation frequency (%) of target “C” in HEK293T cells based on whole exon sequencing under different gBlocks pools and mediums with or without Doxycycline.

**Supplementary Figure 6.**
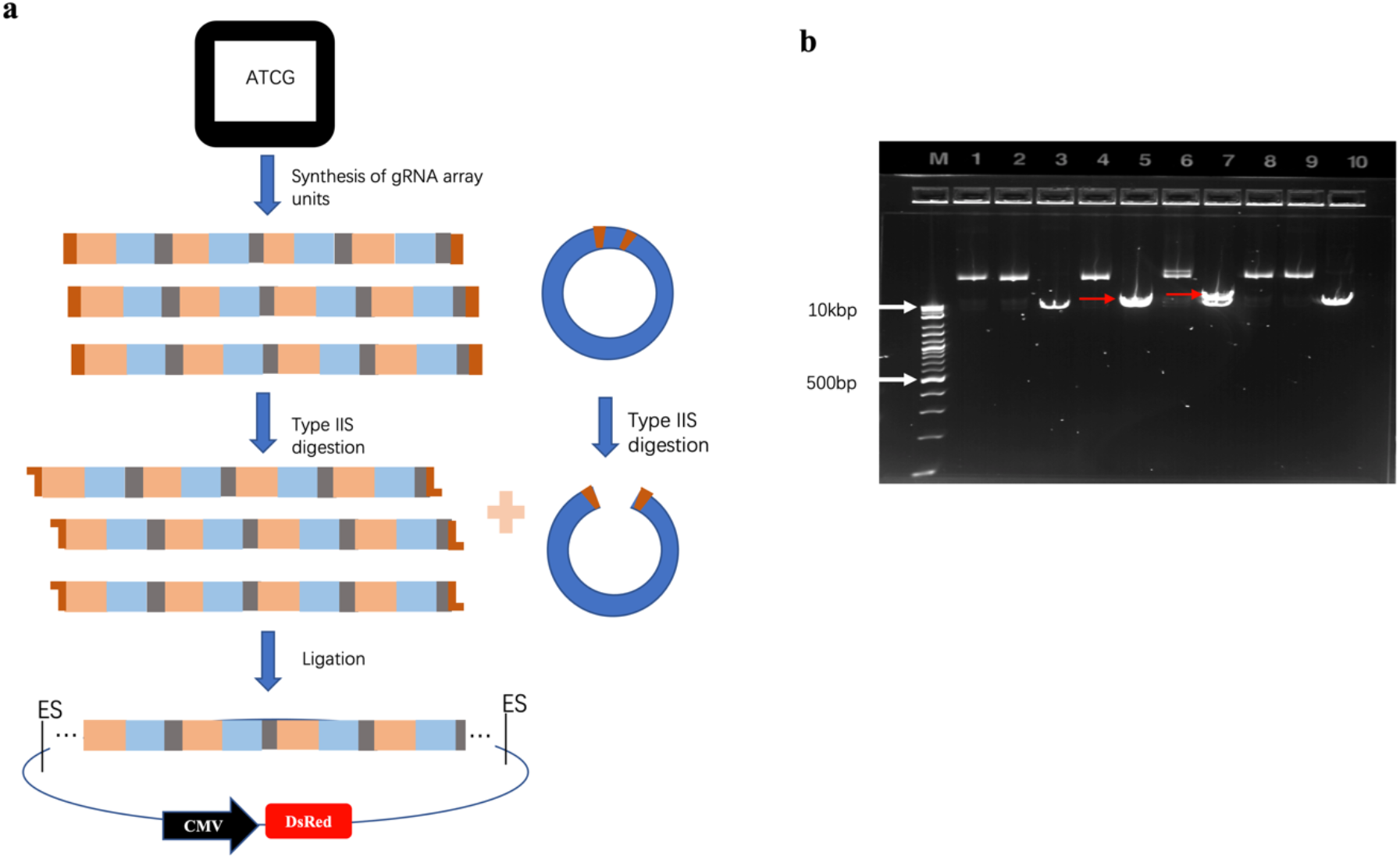
Synthesis and golden gate assembly strategy for 43 sgRNAs all-in-one plasmid. (a) workflow of gBlocks assembly. sgRNAs design by software; multiple gRNA array units are synthesized in tandem, and the ends of each synthesized piece of DNA contain overhangs for specified Type IIs restriction sites for Golden Gate; destination plasmid containing Bbs1 site and two spe1 restrictive endonuclease site on both sides of the Bbs1 site, ES, endonuclease site. (b) Agarose gel electrophoresis analysis of the final all-in-one plasmid. DNA ladder was shown on the left. All the plasmids were linearized by endonuclease enzyme spe1. The empty vector on the far right was also shown as a control. Two out of the nine tested plasmids have the right insertion size. Red arrow is 22Kb.

**Supplementary Figure 7.**
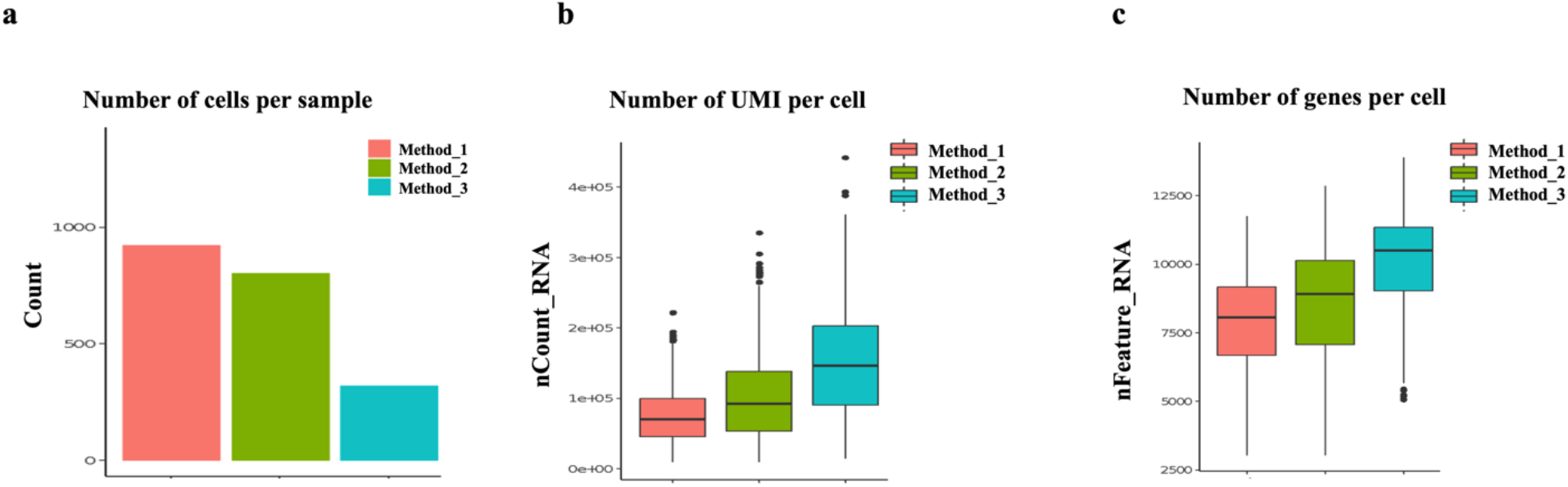
Basic quality metrics of single-cell RNA-seq of 3 different delivery methods. (a) Number of cells that were captured. (b) Number of UMI per cell. c. Number of detected genes per cell.

**Supplementary Figure 8.**
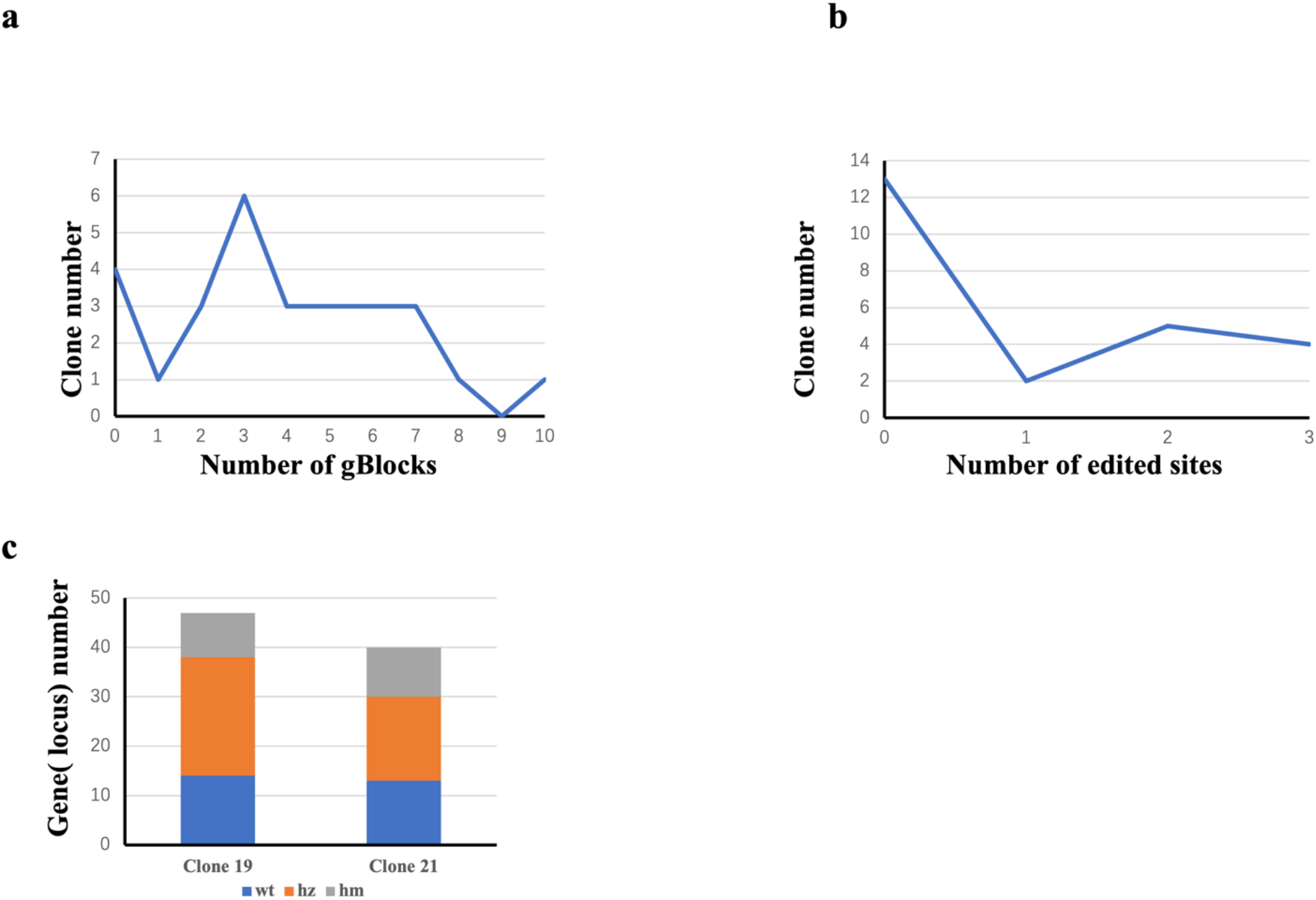
Single clone screening by Sanger sequencing. (a) Picked 10 well edited loci (one from each gBlock to validate their delivery) the peak number of gBlocks is 3, and only one clone have all 10 gBlocks. (b) 3 well edited loci for screening and half of clones without any editing and 4 clones have all 3 editing sites. (c) Allele editing of all target sites in each clone by Sanger sequencing and EditR. WT (wildtype) - no allele editing; HZ (heterozygous) – partial allele editing; HM (homozygous) - all allele editing.

**Supplementary Figure 9.**
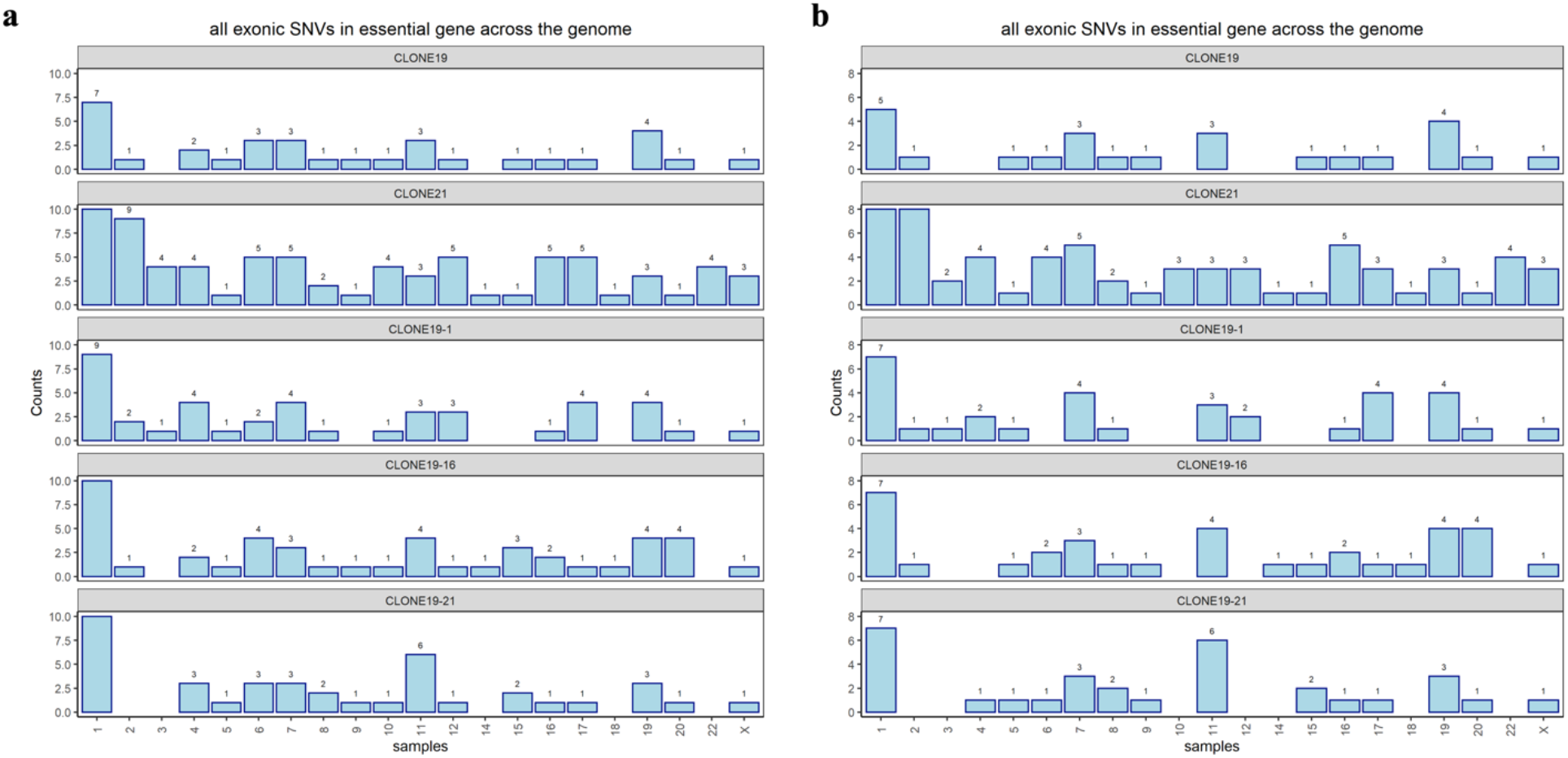
Chromosome distribution of exonic SNVs in essential genes. (a) With and (b) without the ones in selected 50 essential gene targets. X axis indicates each chromosome, and y axis indicates the count on that chromosome. Number of exonic SNVs in essential genes on each chromosome was marked on top of each bar for better demonstration.

**Supplementary Figure 10.**
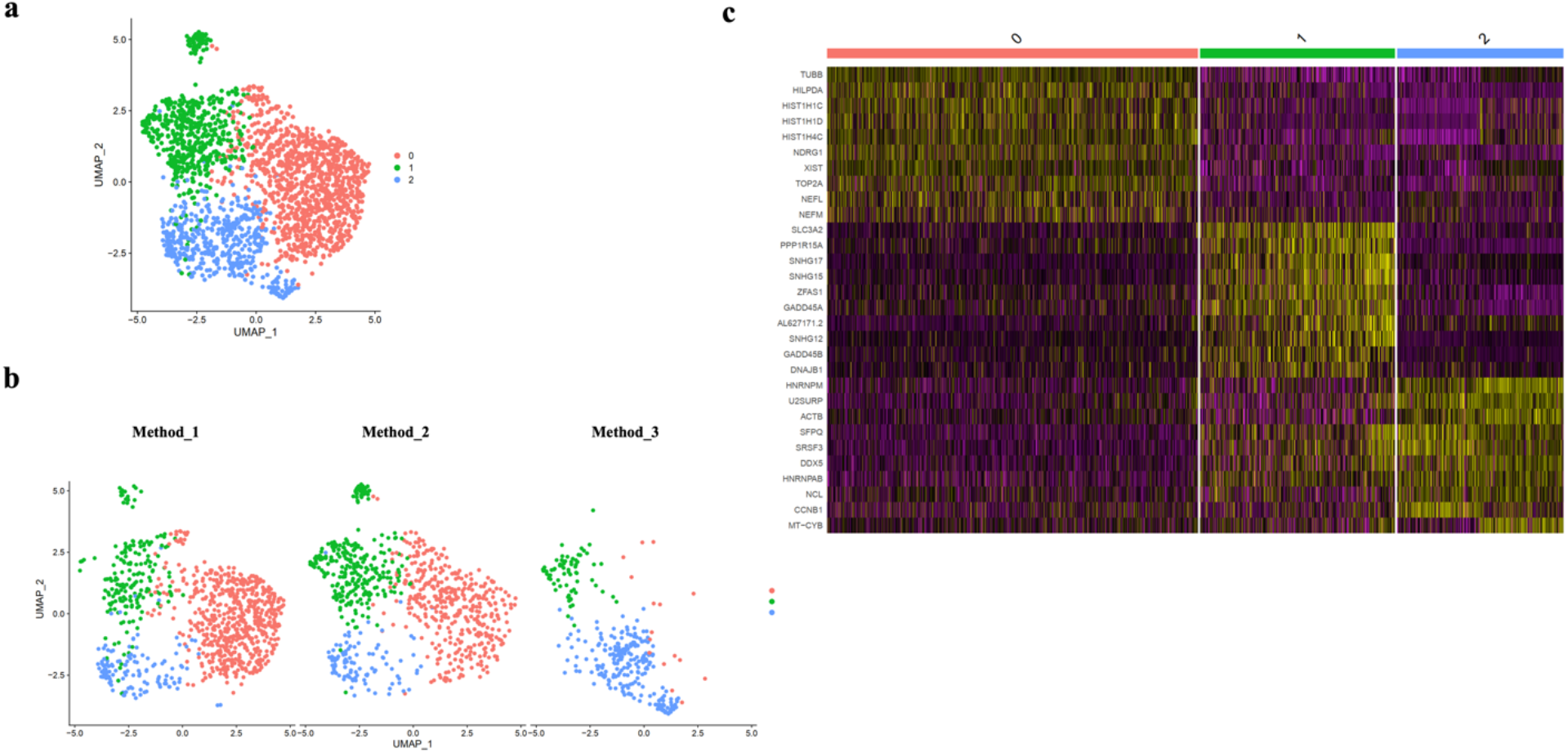
Clustering analysis of single cells from three different delivery methods. (a) UMAP of all single cells from three samples, clustering with 0.3 resolution showed three clusters. (b) Distribution of single cells from three samples in the three clusters. (c)Top 10 enriched genes in each of the three clusters.

**Supplementary Figure 11.**
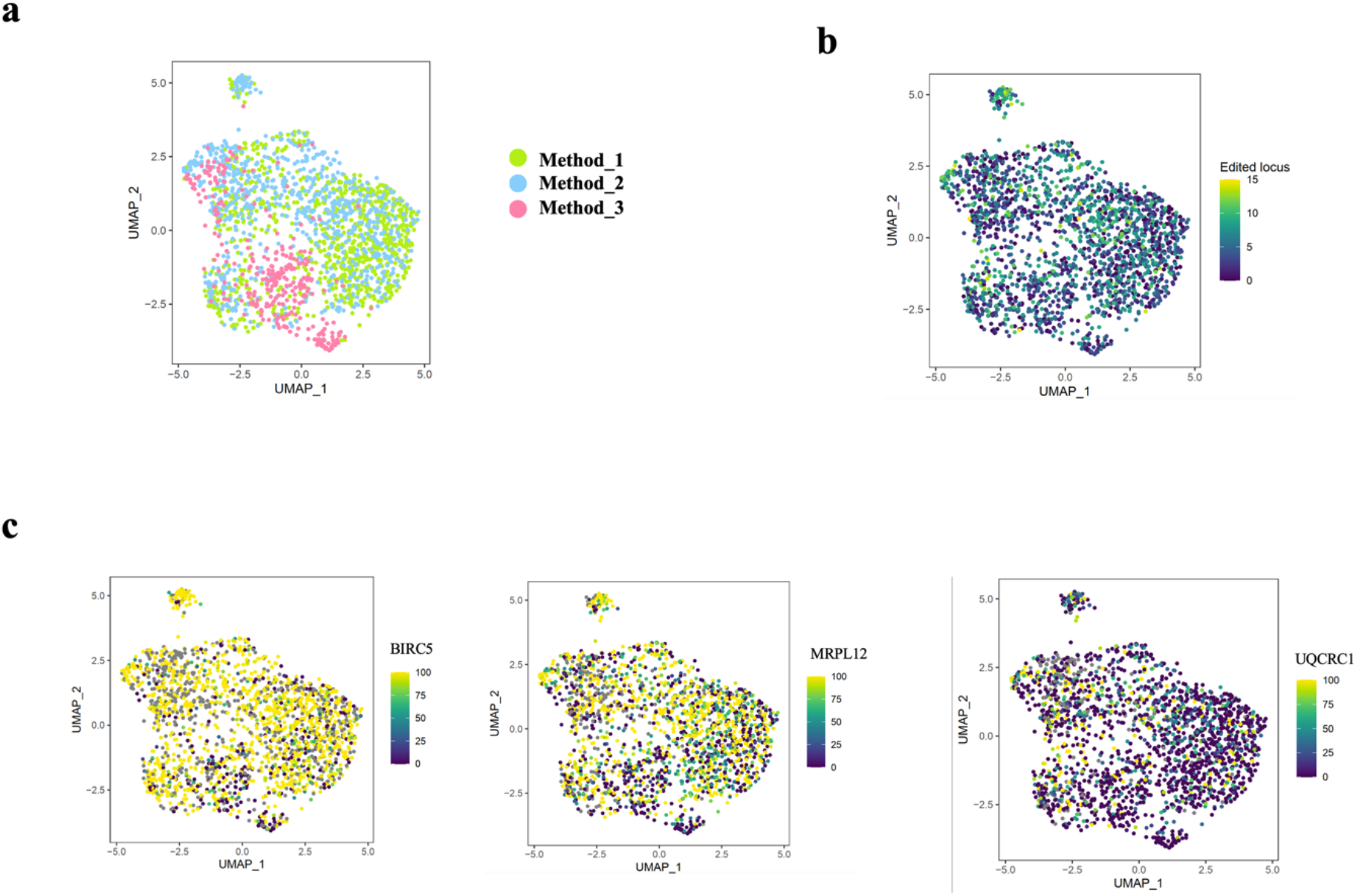
Analysis of on-target editing efficiency and single cell clusters. (a) Single-cell UMAP colored by three different delivery methods. (b) Distribution of number of edited loci in single cells on UMAP. (c) Distribution of editing efficiency for representative loci (BIRC5, MRPL12, UQCRC1) on UMAP.

**Supplementary Figure12.**
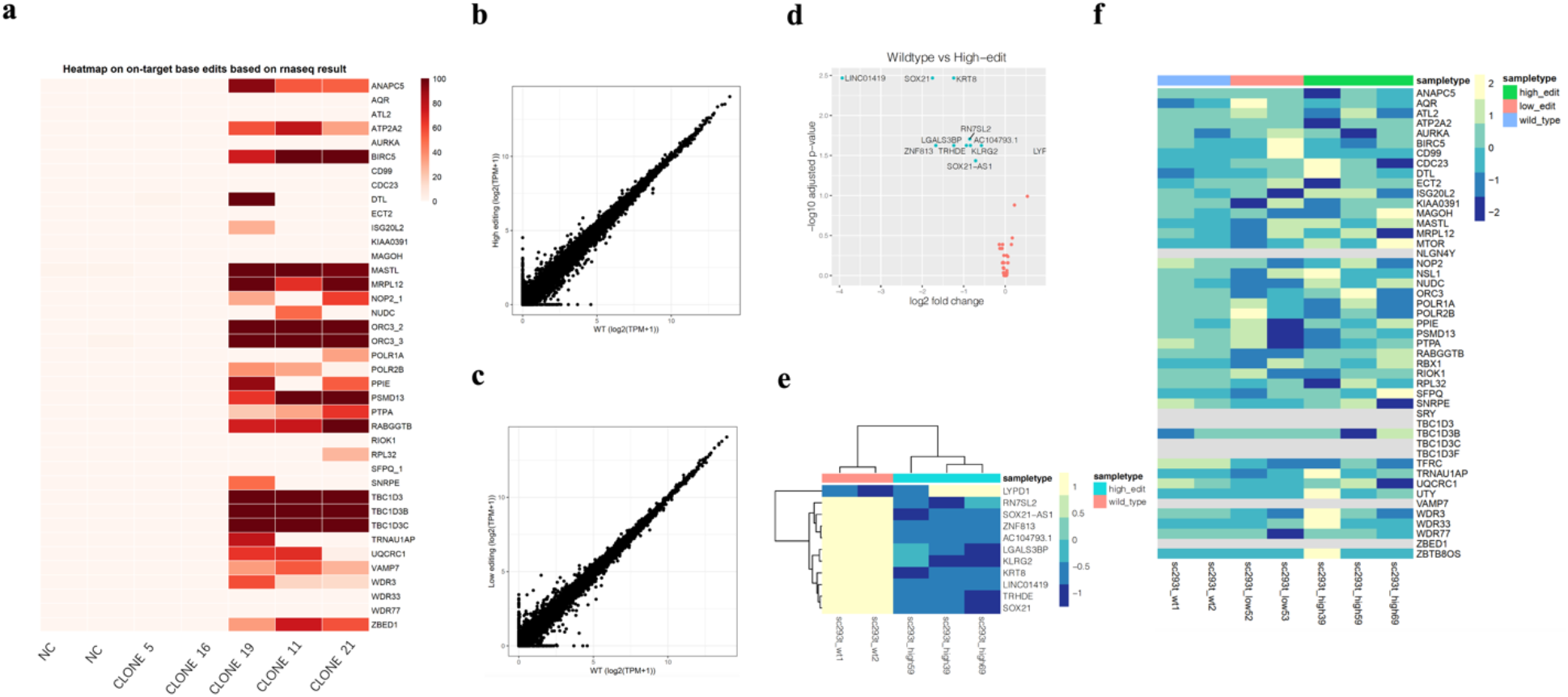
On-target and Gene expression analysis in highly modified HEK293T clones and lowly modified clones by bulk RNAseq. (a) On-target editing efficiency in two negative control (NC) clones, two lowly modified clones, and three highly modified clones. (b)Transcriptional correlation of wild-type negative control clones and highly modified clones. (c) Transcriptional correlation of wild-type negative control clones and lowly modified clones. (d)Volcano plot for differentially expressed genes between wild-type negative control clones and highly modified clones. (e) Heatmap for differentially expressed genes between wild-type negative control clones and highly modified clones. (f) Expression level of targeted loci in three conditions.

**Supplementary Figure 13.**
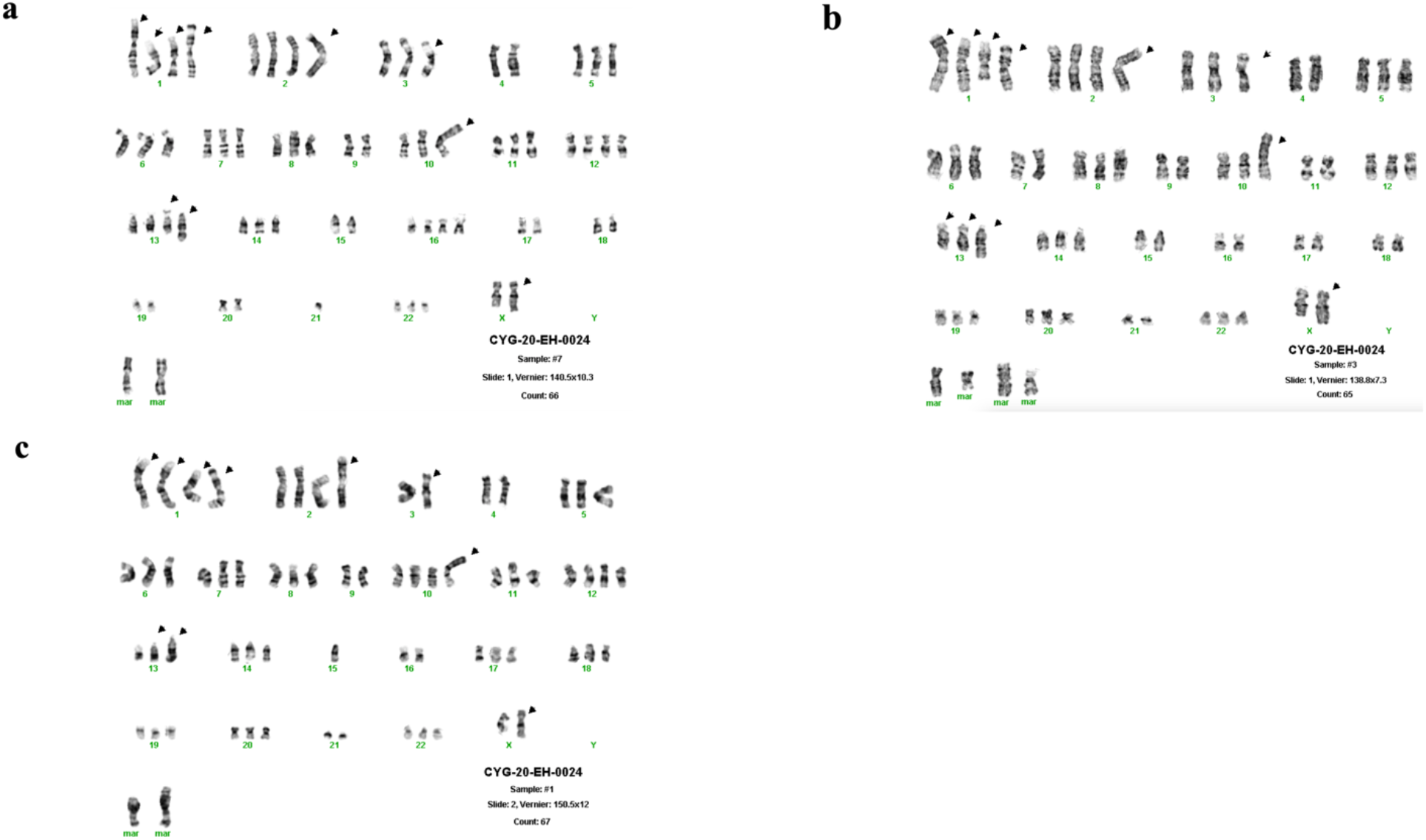
Karyotype analysis of the highly and lowly modified HEK293T clones. The chromosomal arrangement of one of highly modified HEK293T clone(a) and one lowly modified HEK293T clone(b) and Wildtype HEK 293T(c) were determined using Karyotype analysis. The black arrows indicated clonal abnormalities.

**Table S1.**
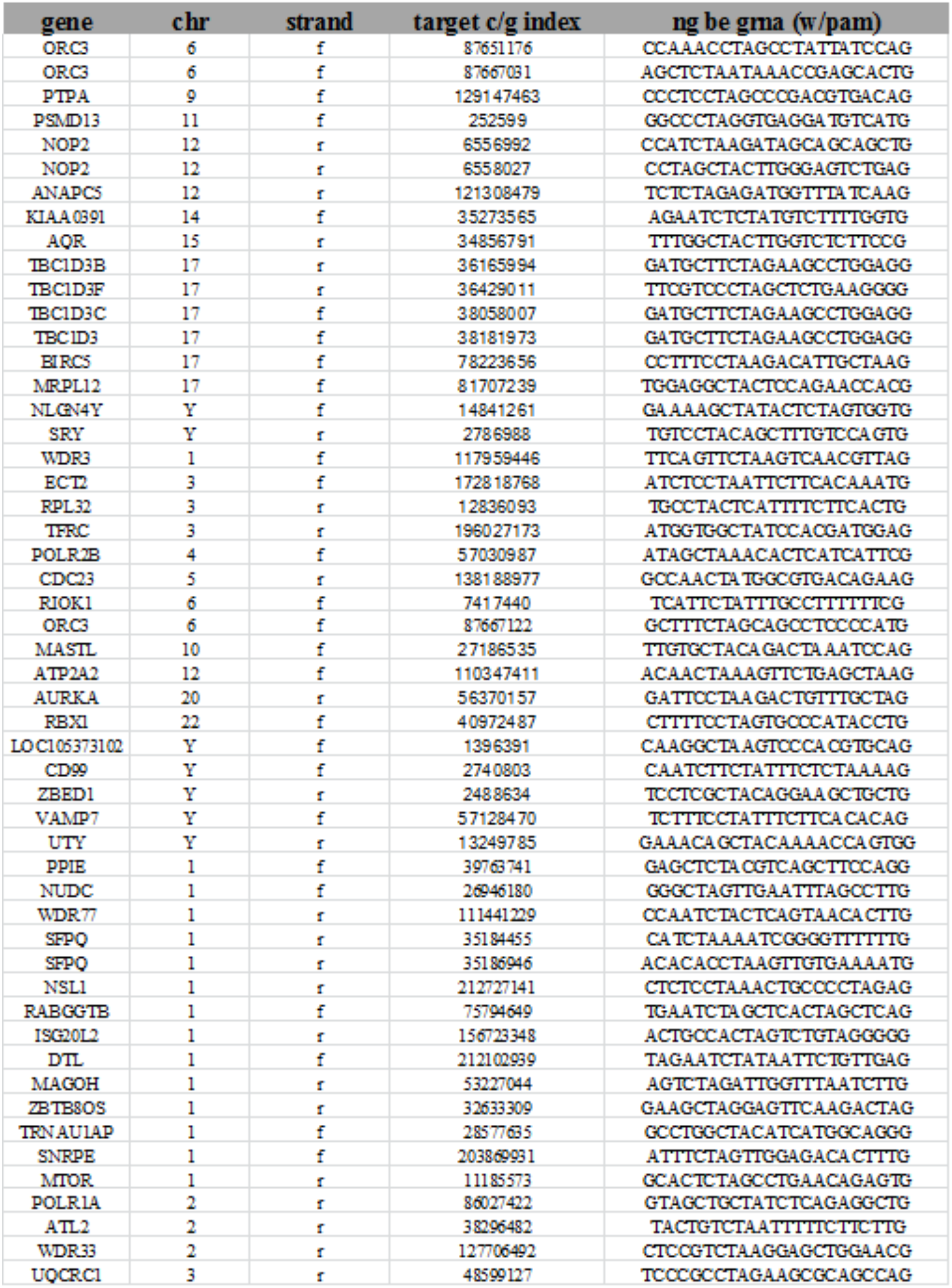
50 sgRNAs sequences targeting 52 gene sites.

**Table S2.**
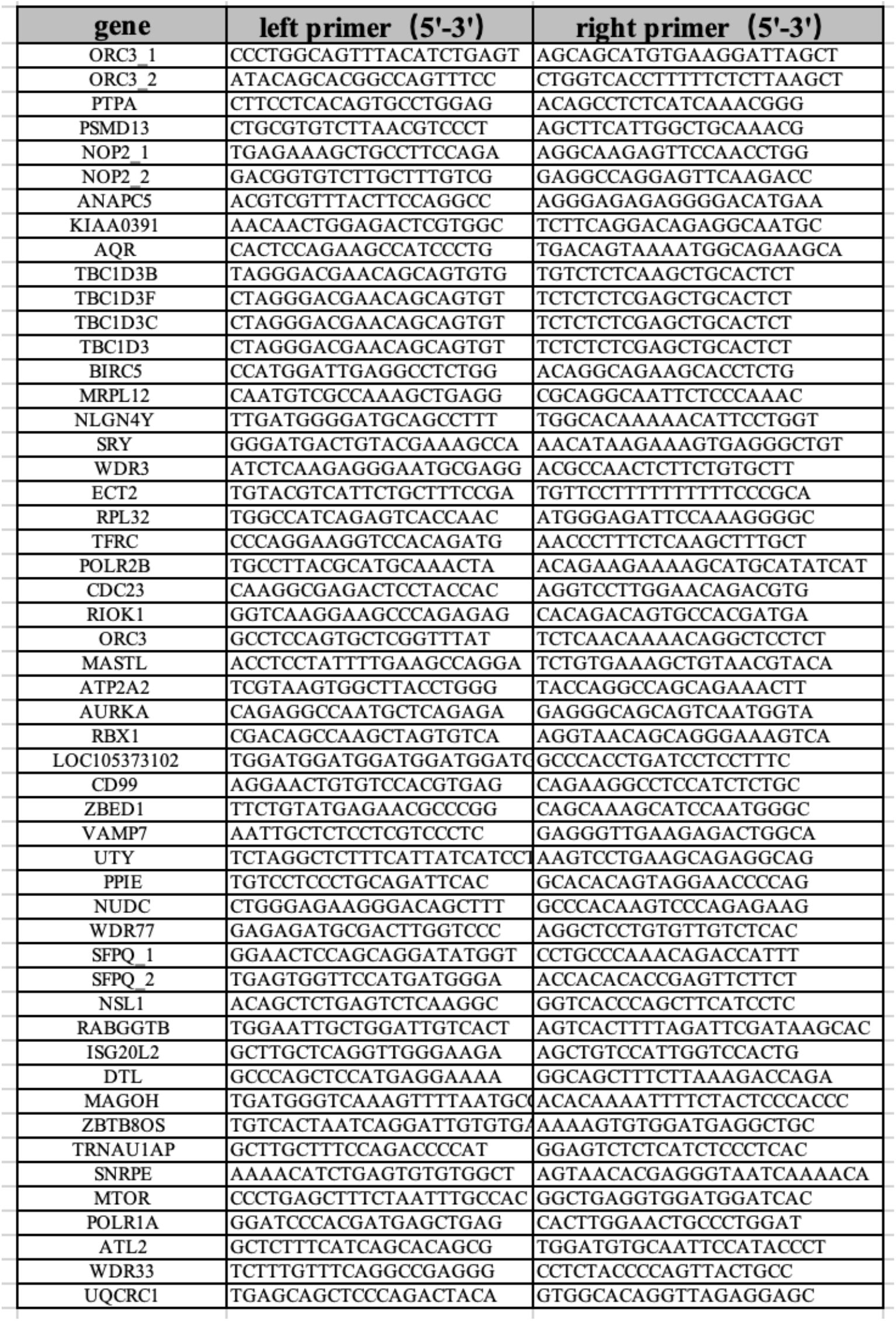
Sanger sequence primers used in this study.

**Table S3.** Summary of the karyotype analysis of HEK293T highly modified clones (19,21) and lowly modified clones (11,16).

|  | HEK293T | Clone11 | Clone16 | Clone19 | Clone21 | 293T9 (CYG-18-PK-0040) | 293T (CYG-18-PK-0021) |
| --- | --- | --- | --- | --- | --- | --- | --- |
| -X |  | x | x |  |  |  |  |
| add(X) (q28) | x | x | x | x | x | x |  |
| der(X) add(X) (p11.2) add(X) (q28) |  |  |  |  |  |  |  |
| add(1) (p36.1) | x | x | x | xx | x | x |  |
| add(1) (q42) | xx | xx | xx | xx | xx | xx |  |
| del(1) (q31) | x | x | x | xx | xx | x | x |
| -2 |  |  |  |  |  |  | x |
| i(2) (q10) | x | x | x | x | x |  |  |
| -3 |  |  |  | x |  |  |  |
| add(3) (p24) | x | x | x |  | x | x |  |
| del(3) (p22) |  |  |  |  |  |  | x |
| add(3) (q12) |  |  |  |  |  | x |  |
| -4 | xx | x | x | xx | x | x | x |
| add(4) (p16) |  | x |  |  |  |  |  |
| add(7) (q36) |  |  |  |  |  |  |  |
| -7 |  |  | x |  |  | x |  |
| -8 | x | x | x | x |  | x |  |
| -9 | x | x |  | x | x |  |  |
| add(9) (q34) |  | x |  |  |  |  | x |
| add(10) (p13) | x |  | x | x | x | x |  |
| -10 |  | x |  |  |  |  |  |
| add(11) (q23) |  |  |  | x |  |  |  |
| -11 |  |  | x | x |  |  |  |
| -12 |  |  |  | x |  |  |  |
| add(13) (p11) | x | x | x | x | x | x | x |
| add(13) (q34) | x | x | x | x | x | x |  |
| -13 |  |  | x |  |  |  | x |
| -14 |  | x |  |  |  |  |  |
| -15 | x | x | x | x | x | x | x |
| -16 |  |  |  | x |  |  |  |
| -17 |  |  | x | x | x |  |  |
| -18 | x | x | x | x | x | x | x |
| -21 | x | x | x | x | x | x | x |
| -22 |  | x |  |  |  |  |  |
| mar1 | x | x |  |  |  |  |  |
| mar | 1-4 | 1-6 | 3-5 | 4-5 | 1-4 | 1-4 | 3-5 |

