## Supplementary Excel Data 1 for "Multiplex base editing to convert TAG into TAA codons in the human genome"

uagzuaa_1_13_window

| | | gene | gene_ogee | gene_essentiality | site_inde | cbe_targets | abe_targets |
|---|---|---|---|---|---|---|---|
| 0 | 1 | OR4F5 | {'ess': 0, 'nonest': 3} | True | 70007 | AAGATCTAAAACTTTACACTAG | [] |
| 1 | 1 | PERM1 | unavailable | False | 976171 | TGGCTCCTAGGAGCTGGGGCT | [] |
| 2 | 1 | RNF223 | unavailable | False | 1071816 | TGGCCCTAATTATCAGTCAGAG | [] |
| 3 | 1 | TTLL10 | {'ess': 0, 'nonest': 7} | True | 1185524 | CCTTAGCTAGGGAGAGTCCGG | [] |
| 4 | 1 | TTLL10 | {'ess': 0, 'nonest': 7} | True | 1197846 | TGGCTGCCCCTAGGGCCTCGC | [] |
| 5 | 1 | TNFRSF1 | {'ess': 0, 'nonest': 7} | True | 1203590 | TCCTAAGACCCCACCCCATCA | [] |
| 6 | 1 | ACAP3 | {'ess': 0, 'nonest': 7} | True | 1293563 | CCGGCCCTAGCTCTCTTCCAG | [] |
| 7 | 1 | CPTP | unavailable | False | 1327762 | CCCGCCCCTAGGGCAGGTCC | [] |
| 8 | 1 | MXRA8 | {'ess': 0, 'nonest': 7} | True | 1353603 | CCTATTTGCAGTTCTCCTTCCG | [] |
| 9 | 1 | MRPL20 | {'ess': 6, 'nonest': 2} | False | 1402568 | TGGCCTAGGGTAACCCTTTAAG | [] |
| 10 | 1 | ATAD3B | {'ess': 0, 'nonest': 7} | True | 1495816 | CCAGTGCCTACAACAGGGGGT | [] |
| 11 | 1 | MMP23B | {'ess': 0, 'nonest': 7} | True | 1634328 | TCACTACACTTTCCCTTTCTTG | [] |
| 12 | 1 | NADK | {'ess': 4, 'nonest': 5} | False | 1752903 | -1 | -1 |
| 13 | 1 | CALML6 | {'ess': 1, 'nonest': 7} | True | 1917192 | CTGCACCTACTGGATCAGCTT | [] |
| 14 | 1 | PRKCZ | {'ess': 1, 'nonest': 7} | True | 2157294 | GGGACGCCCTAGTAGCTGGCT | [] |
| 15 | 1 | SKI | {'ess': 2, 'nonest': 6} | False | 2306764 | GGAATCTACGGCTCCAGCTCC | [] |
| 16 | 1 | MORN1 | {'ess': 0, 'nonest': 7} | True | 2355447 | CGTCTGATCCTAGAAGAGTGAG | [] |
| 17 | 1 | RER1 | {'ess': 1, 'nonest': 5} | False | 2403123 | TCCCGCTTCTAGCTGGCGAAG | [] |
| 18 | 1 | PRXL2B | unavailable | False | 2589478 | AGCACACCTAGGGCAGCAGAC | [] |
| 19 | 1 | MMEL1 | {'ess': 1, 'nonest': 7} | True | 2590989 | TTGGCTACCACACGCGGCATC | [] |
| 20 | 1 | ARHGEF1 | {'ess': 0, 'nonest': 7} | True | 3480586 | GCTACACGTCCGTCTCCACCC | [] |
| 21 | 1 | MEGF6 | {'ess': 1, 'nonest': 7} | True | 3490527 | GCCTCTACTAGTGCCTCGCTG | [] |
| 22 | 1 | TPRG1L | {'ess': 0, 'nonest': 7} | True | 3628602 | CAGCGGCTAAAAGCCTATTTTG | [] |
| 23 | 1 | WRAP73 | unavailable | False | 3630974 | GCTGCTACGTGTGGCCGCCCA | [] |
| 24 | 1 | CCDC27 | {'ess': 0, 'nonest': 7} | True | 3768596 | ACTACGGCTGGATCCCAAACT | [] |
| 25 | 1 | CCDC27 | {'ess': 0, 'nonest': 7} | True | 3771522 | CCTGGCTGGGCCTACTTGGAG | [] |
| 26 | 1 | C1orf174 | {'ess': 0, 'nonest': 7} | True | 3889959 | CCCCTACATTTCTGCATCATCG | [] |
| 27 | 1 | CHD5 | {'ess': 1, 'nonest': 6} | True | 6106279 | GCTAGATATCTGTCAAAAAAAG | [] |
| 28 | 1 | RNF207 | {'ess': 0, 'nonest': 7} | True | 6219406 | GTCCCATTTGCTAAGTCGGGT | [] |
| 29 | 1 | GPR153 | {'ess': 0, 'nonest': 7} | True | 6249327 | GGTCCTAGGACGCGGAGCCCA | [] |
| 30 | 1 | ACOT7 | {'ess': 0, 'nonest': 7} | True | 6264596 | GGAGTCTAGGGCTGAGGCTCC | [] |
| 31 | 1 | HES2 | {'ess': 0, 'nonest': 7} | True | 6418872 | GAGGGGCTACCACGGCCGCC | [] |
| 32 | 1 | ESPN | {'ess': 1, 'nonest': 6} | True | 6446423 | CAGCCCTGCTAGTGCCTGCTC | ['CTTCCCCCCACCCTCCTTCCAG', 'GTGCCTGCTCACCTTCCCCCCG'] |
| 33 | 1 | ESPN | {'ess': 1, 'nonest': 6} | True | 6450145 | CGGCCTCTAGTACTTAGCGAT | [] |
| 34 | 1 | ZBTB48 | {'ess': 0, 'nonest': 7} | True | 6588211 | GGGCTATGTGTCACAGTCCTC | [] |
| 35 | 1 | KLHL21 | {'ess': 0, 'nonest': 7} | True | 6593364 | CTGGGGCTAGTGCAGCTCATC | [] |
| 36 | 1 | PARK7 | {'ess': 0, 'nonest': 7} | True | 7985053 | TGCTCTAGTCTTTAAGAACAAG | [] |
| 37 | 1 | ERRFI1 | {'ess': 0, 'nonest': 7} | True | 8013209 | GGTCTAAGGAGAAACCACATAG | [] |
| 38 | 1 | SLC2A7 | {'ess': 0, 'nonest': 7} | True | 9003299 | CCACTAAAAGGAAGTTTCCTTG | [] |
| 39 | 1 | SLC2A7 | {'ess': 0, 'nonest': 7} | True | 9007486 | GAGCTGCACCTAGATGTCTGG | [] |
| 40 | 1 | SLC2A5 | {'ess': 0, 'nonest': 7} | True | 9041620 | TGGCCTGCTATGTTGGCTCGG | [] |
| 41 | 1 | PIK3CD | {'ess': 4, 'nonest': 3} | False | 9727045 | CCACTACTGCCTGTTGTCTTTG | [] |
| 42 | 1 | CTNNBIP1 | {'ess': 0, 'nonest': 7} | True | 9850717 | GCTTTGCAGCTACTGCCTCCG | [] |
| 43 | 1 | NMNAT1 | {'ess': 0, 'nonest': 1} | False | 9982700 | AATTCCTATGTCTTAGCTTCTG | [] |
| 44 | 1 | UBE4B | {'ess': 4, 'nonest': 4} | False | 10171136 | CAAGCTCCTAGGAAATCACAGG | [] |
| 45 | 1 | DFFA | {'ess': 0, 'nonest': 7} | True | 10461489 | GCTGCTGCTATGTGGGATCCT | [] |
| 46 | 1 | PEX14 | {'ess': 2, 'nonest': 5} | False | 10629886 | GCCCTAGTCCCGCTCACTCTC | [] |
| 47 | 1 | CASZ1 | {'ess': 0, 'nonest': 7} | True | 10638941 | CGCCGCTAGGGCGAGGAGGCT | [] |
| 48 | 1 | TARDBP | {'ess': 2, 'nonest': 5} | False | 11022653 | ACTGTCTACATTCCCCAGCCA | [] |
| 49 | 1 | MASP2 | {'ess': 0, 'nonest': 7} | True | 11044939 | GGAGGCTAGAGGCTCTGCTCT | [] |
| 50 | 1 | EXOSC10 | {'ess': 9, 'nonest': 2} | False | 11066717 | CCAGGACTATCTCTGTGGCCA | [] |
| 51 | 1 | MTOR | {'ess': 8, 'nonest': 1} | False | 11185573 | GCACTCTAGCCTGAACAGAGT | [] |
| 52 | 1 | DISP3 | unavailable | False | 11536885 | CCGGGCTATAGGGAGGCCCCT | [] |
| 53 | 1 | DRAXIN | unavailable | False | 11719895 | GGGCCGCTAGACGTTGATGAA | [] |
| 54 | 1 | AGTRAP | {'ess': 0, 'nonest': 7} | True | 11750103 | TAGGCACTACGGTCCTGAGAA | [] |
| 55 | 1 | C1orf167 | {'ess': 0, 'nonest': 1} | False | 11789458 | ATTCTATGAGAACAAGTCACAG | [] |
| 56 | 1 | MTHFR | {'ess': 0, 'nonest': 7} | True | 11790539 | CCAGTCTAGCTGCCATTGTCA | [] |
| 57 | 1 | MFN2 | {'ess': 6, 'nonest': 2} | False | 12011564 | GTGCCCACTATCTGCTGGGCT | [] |
| 58 | 1 | TNFRSF1 | {'ess': 2, 'nonest': 6} | False | 12195193 | CAGGCCCTACTTTATAAAACAG | [] |
| 59 | 1 | DHRS3 | {'ess': 0, 'nonest': 7} | True | 12568339 | CTGTCTCTATGTCCGCCCTTT | [] |
| 60 | 1 | PRAMEF1 | {'ess': 0, 'nonest': 7} | True | 12795895 | CCTTCCCTAGCAGCAAAGATG | [] |
| 61 | 1 | PRAMEF2 | {'ess': 0, 'nonest': 7} | True | 12861778 | CCTTCCCTAGCAGCAAAGATG | [] |
| 62 | 1 | PRAMEF1 | {'ess': 0, 'nonest': 7} | True | 12892915 | CCTTCCCTAGGAGCAAAGATG | [] |
| 63 | 1 | PRAMEF1 | {'ess': 0, 'nonest': 3} | True | 13196822 | CCTTCCCTAGCAGCAAAGATG | [] |
| 64 | 1 | PRAMEF1 | {'ess': 0, 'nonest': 4} | True | 13223331 | CTAGGCAGGCCTTCCCCACAA | [] |
| 65 | 1 | PRAMEF2 | unavailable | False | 13308886 | CCTTCCCTAGGAGCAAAGATG | [] |
| 66 | 1 | PRAMEF1 | {'ess': 0, 'nonest': 3} | True | 13342527 | CCTTCCCTAGCAGCAAAGATG | [] |
| 67 | 1 | PRAMEF2 | unavailable | False | 13369066 | CCCACTAGGCAGGCCTTCCCC | [] |
| 68 | 1 | PRAMEF1 | {'ess': 1, 'nonest': 6} | True | 13392501 | CCTTCACTAAGAGCAAAGATGG | [] |
| 69 | 1 | PRDM2 | {'ess': 0, 'nonest': 7} | True | 13780526 | CTTTTCTACAGGAAGTTCCTAG | [] |
| 70 | 1 | PRDM2 | {'ess': 0, 'nonest': 7} | True | 13816546 | AGTGTCTACTCTTTGAAGAATG | [] |
| 71 | 1 | FHAD1 | {'ess': 0, 'nonest': 3} | True | 15383157 | TCCCATCTACTAATGAGGAAAG | [] |
| 72 | 1 | FHAD1 | {'ess': 0, 'nonest': 3} | True | 15397412 | GTTTCTCTAGTACTTTCCACTG | [] |
| 73 | 1 | EFHD2 | {'ess': 0, 'nonest': 7} | True | 15426723 | CCCGCTACTTAAAGGTGGACT | [] |
| 74 | 1 | CTRC | {'ess': 1, 'nonest': 7} | True | 15446770 | ACCTGTCTAATGCAGGGTGGG | [] |
| 75 | 1 | SLC25A34 | {'ess': 0, 'nonest': 7} | True | 15739405 | CCGTCGTCTAGGTGCCCTTGT | [] |
| 76 | 1 | SRARP | unavailable | False | 16006345 | CAGCTAATCGGCCTGCCCACA | [] |
| 77 | 1 | MST1L | unavailable | False | 16757231 | CTGGGCCTAACCCAGTCTCAT | [] |
| 78 | 1 | MFAP2 | {'ess': 0, 'nonest': 7} | True | 16974919 | ACCACCCTAGCAGCTCCCACA | [] |
| 79 | 1 | ATP13A2 | {'ess': 0, 'nonest': 7} | True | 16985984 | TTATTTGCTACACTGACAGCAG | [] |
| 80 | 1 | ATP13A2 | {'ess': 0, 'nonest': 7} | True | 16986220 | CCTGCACTACCTCAGGGGGCC | [] |
| 81 | 1 | PADI6 | unavailable | False | 17401437 | GGGTCTAAGGTACCATCTTCCC | ['CTTCCACCATTTGAAGGCAAAG'] |
| 82 | 1 | ARHGEF1 | {'ess': 0, 'nonest': 7} | True | 17697379 | GGAGGCGCTATAGCATCAAGG | [] |
| 83 | 1 | KLHDC7A | {'ess': 0, 'nonest': 7} | True | 18482314 | GACTGCTAGACCCTGGTCTGA | [] |
| 84 | 1 | PAX7 | {'ess': 0, 'nonest': 7} | True | 18744928 | GGGCCCTAGTAGGCCTGGCCA | [] |
| 85 | 1 | TAS1R2 | {'ess': 0, 'nonest': 7} | True | 18839598 | ATGGCACTAGTCCCTCCTCAT | [] |
| 86 | 1 | AKR7A3 | {'ess': 0, 'nonest': 7} | True | 19282730 | ATGGGCCTAGCGGAAGTAGTT | [] |
| 87 | 1 | AKR7A2 | {'ess': 0, 'nonest': 7} | True | 19304224 | ATGGGCCTAGCGGAAGTAGTT | [] |
| 88 | 1 | TMCO4 | {'ess': 0, 'nonest': 7} | True | 19745453 | AAAAACCTAGCCCAGGTCTGC | [] |
| 89 | 1 | OTUD3 | {'ess': 0, 'nonest': 7} | True | 19907563 | TGTGGCTACCTCTGCTCTCCA | [] |
| 90 | 1 | PLA2G5 | {'ess': 0, 'nonest': 7} | True | 20090691 | GGAGGCCTAGGAGCAGAGGAT | [] |
| 91 | 1 | PLA2G2D | {'ess': 0, 'nonest': 7} | True | 20114113 | GCTTCTAGCACCCAGGGGTCT | [] |
| 92 | 1 | PLA2G2F | {'ess': 0, 'nonest': 7} | True | 20149400 | GAGGCTCTAGGGAGGGGCGGG | [] |
| 93 | 1 | PLA2G2C | {'ess': 0, 'nonest': 3} | True | 20163259 | CAGGCTAGCCCAGGCTTTTTTC | [] |
| 94 | 1 | PLA2G2C | {'ess': 0, 'nonest': 3} | True | 20163990 | GTCCCTAGCACCAGGGCTTAT | [] |
| 95 | 1 | VWA5B1 | {'ess': 0, 'nonest': 3} | True | 20354262 | ACTCAACTACACATAATTCGGG | [] |
| 96 | 1 | RAP1GAP | {'ess': 1, 'nonest': 6} | True | 21597719 | GGCCCGGCTAACAGCCCTGCA | [] |
| 97 | 1 | USP48 | {'ess': 3, 'nonest': 5} | False | 21705869 | TATACCTAATTACTGCATGACG | [] |
| 98 | 1 | HSPG2 | {'ess': 0, 'nonest': 5} | True | 21823315 | AGGTGCCTACGAGGGGCAGGG | [] |
| 99 | 1 | CELA3B | {'ess': 0, 'nonest': 7} | True | 21989278 | TTGGTTCTAGTGGCTTGCTATG | [] |
| 100 | 1 | CELA3A | {'ess': 0, 'nonest': 7} | True | 22012456 | TTGGTTCTAGTGGCTTGCTATG | [] |
| 101 | 1 | EPHA8 | {'ess': 0, 'nonest': 7} | True | 22589378 | CTTTTCTAAAGGCAATGAAGAG | [] |
| 102 | 1 | C1QC | {'ess': 0, 'nonest': 7} | True | 22647782 | GCCCGCCCTAGTCGGGGAAGA | [] |
| 103 | 1 | HTR1D | {'ess': 1, 'nonest': 5} | False | 23193085 | ATAAGACTAGGAGGCCTTCCG | [] |
| 104 | 1 | HNRNPR | {'ess': 0, 'nonest': 7} | True | 23310453 | TACTTGTCTACTTCCACTGTTG | [] |
| 105 | 1 | TCEA3 | {'ess': 0, 'nonest': 7} | True | 23381474 | GCAGAACTACAAAACCAGAAAG | [] |
| 106 | 1 | ASAP3 | {'ess': 0, 'nonest': 7} | True | 23429855 | CAAGGAGCTAGTCTTGCAAAAG | [] |
| 107 | 1 | STPG1 | unavailable | False | 24358542 | GTGACATCCCTACAGAACCGG | [] |
| 108 | 1 | NIPAL3 | {'ess': 0, 'nonest': 7} | True | 24454096 | GAGAGCTAAGATTGTACCACTG | [] |
| 109 | 1 | SRRM1 | {'ess': 7, 'nonest': 1} | False | 24672285 | ATTTCCCCCTAAGACTGTGGG | [] |
| 110 | 1 | MACO1 | unavailable | False | 25491356 | TCGATGTCTATAAATTAGCCTG | [] |
| 111 | 1 | LDLRAP1 | {'ess': 0, 'nonest': 7} | True | 25565415 | TTTCTAACAGCTCTGGACAACG | [] |
| 112 | 1 | SELENON | unavailable | False | 25815717 | GGCACTCTAGGGCTGGAGGAG | [] |
| 113 | 1 | STMN1 | {'ess': 1, 'nonest': 6} | True | 25885722 | AAGTACCTAGTGATGGTGAGAG | [] |
| 114 | 1 | PAFAH2 | {'ess': 0, 'nonest': 7} | True | 25961988 | GTTGTGCCTACAGGCTGGACA | [] |
| 115 | 1 | EXTL1 | {'ess': 2, 'nonest': 5} | False | 26035346 | GCCCCCCTAGGGCTTCTCCAG | [] |
| 116 | 1 | TRIM63 | {'ess': 0, 'nonest': 7} | True | 26053803 | ATTGCCAATGTCTAACACAGTG | [] |
| 117 | 1 | C1orf232 | unavailable | False | 26164160 | GCCTGCTAGTCACCCTTGGGC | [] |
| 118 | 1 | GPN2 | {'ess': 7, 'nonest': 1} | False | 26879676 | CCTTGTTGCTACAGCTGCATGC | [] |
| 119 | 1 | NUDC | {'ess': 7, 'nonest': 1} | False | 26946180 | AACAGGGGCTAGTTGAATTTAG | [] |
| 120 | 1 | WDTC1 | {'ess': 1, 'nonest': 7} | True | 27306382 | GGGTCTAGCTGGGCCGGCACT | [] |
| 121 | 1 | SYTL1 | {'ess': 0, 'nonest': 6} | True | 27353851 | GGGGCTACGTCCTGGGGGCC | [] |
| 122 | 1 | FCN3 | {'ess': 0, 'nonest': 7} | True | 27369235 | TGCCCTATCGAAGCATCATCC | [] |
| 123 | 1 | GPR3 | {'ess': 0, 'nonest': 7} | True | 27394790 | ACTCAGCTAGACATCACTGGG | [] |
| 124 | 1 | AHDC1 | {'ess': 0, 'nonest': 5} | True | 27547303 | TTGGCACTACAGGGATGTGAC | [] |
| 125 | 1 | FGR | {'ess': 1, 'nonest': 6} | True | 27612913 | GACAGGCTATGTCTGATCCCC | [] |
| 126 | 1 | IFI6 | {'ess': 1, 'nonest': 7} | True | 27666380 | TGGCTACTCCTCATCCTCCTC | ['CTCCTCACTATCGAGATACTTG'] |
| 127 | 1 | FAM76A | {'ess': 0, 'nonest': 7} | True | 27750221 | CCCTCTAAATAAACAAAATCAG | [] |
| 128 | 1 | XKR8 | {'ess': 1, 'nonest': 7} | True | 27967199 | TTCACCTATCCCTTCACTGGC | [] |
| 129 | 1 | PTAFR | {'ess': 0, 'nonest': 7} | True | 28149992 | AGCAGGGACTAATTTTTGAGGG | [] |
| 130 | 1 | TRNAU1A | {'ess': 7, 'nonest': 1} | False | 28577535 | GCCTGGCTACATCATGGCAGG | [] |
| 131 | 1 | EPB41 | {'ess': 0, 'nonest': 7} | True | 29069293 | TATCTATGAAAGGAAAAACTTG | [] |
| 132 | 1 | EPB41 | {'ess': 0, 'nonest': 7} | True | 29072660 | TACCCTATTGATAGAAGAAATG | [] |
| 133 | 1 | PTPRU | {'ess': 0, 'nonest': 7} | True | 29325660 | GGCCCCGCTATCTTGACTCCA | [] |
| 134 | 1 | SDC3 | {'ess': 0, 'nonest': 7} | True | 30873210 | ACTAGGCATAGAACTCCTCCTG | [] |
| 135 | 1 | NKAIN1 | {'ess': 0, 'nonest': 7} | True | 31181702 | CAGAGGCTACCCCGACCTGCG | [] |
| 136 | 1 | COL16A1 | {'ess': 0, 'nonest': 7} | True | 31670589 | AGCAGCGCTAGGTTCTTACAG | [] |
| 137 | 1 | PTP4A2 | {'ess': 2, 'nonest': 6} | False | 31908851 | CCTTCTACTGAACACAGCAATG | [] |
| 138 | 1 | TXLNA | {'ess': 1, 'nonest': 7} | True | 32195194 | GGCTCTCTAGGCCCTGGCGGA | [] |
| 139 | 1 | CCDC28B | {'ess': 0, 'nonest': 7} | True | 32204525 | TGGGGGCTATCTTCTGGGGAG | [] |
| 140 | 1 | CCDC28B | {'ess': 0, 'nonest': 7} | True | 32204797 | CACTTTTCTTCTAATTTTGGGG | [] |
| 141 | 1 | CCDC28B | {'ess': 0, 'nonest': 7} | True | 32205247 | GGGACGCCTACGCAGCGGACT | [] |
| 142 | 1 | IQCC | {'ess': 1, 'nonest': 6} | True | 32208081 | AGGGTCTAGCCAGGTGGTTCT | [] |
| 143 | 1 | MARCKSL | {'ess': 0, 'nonest': 7} | True | 32234596 | CCTAGCTACTCATTCTGCTCAG | [] |
| 144 | 1 | ZBTB8B | {'ess': 1, 'nonest': 4} | False | 32485417 | ATTGCTACGTAAGTGTCTCTCG | [] |
| 145 | 1 | ZBTB8A | {'ess': 0, 'nonest': 7} | True | 32600418 | ATTACAGCTACCTAATGTTGGG | [] |
| 146 | 1 | ZBTB8OS | {'ess': 8, 'nonest': 1} | False | 32633309 | GAAGCTAGGAGTTCAAGACTAG | [] |
| 147 | 1 | RBBP4 | {'ess': 7, 'nonest': 1} | False | 32679704 | ATCTAGGACCCTTGTCCTTCTG | [] |
| 148 | 1 | SYNC | {'ess': 0, 'nonest': 7} | True | 32681949 | ATATATTTCTAAACAGCCCCTG | [] |
| 149 | 1 | SYNC | {'ess': 0, 'nonest': 7} | True | 32683785 | CACTATGCCCGGCTAATTTTTG | [] |
| 150 | 1 | YARS1 | unavailable | False | 32775980 | -1 | -1 |
| 151 | 1 | AK2 | {'ess': 5, 'nonest': 2} | False | 33010828 | CTGATACTAGGCTGAAATAGAG | [] |
| 152 | 1 | AZIN2 | unavailable | False | 33101865 | GGACTCTAGAGACATGTAGCAG | [] |
| 153 | 1 | AZIN2 | unavailable | False | 33101891 | AGCGCTACACGGCAATGAAATG | [] |
| 154 | 1 | AZIN2 | unavailable | False | 33160887 | TGTCTATAAACACCAGTCCTTG | [] |
| 155 | 1 | TRIM62 | {'ess': 0, 'nonest': 7} | True | 33147176 | CCTGGACTAGATGCGGACGGT | [] |
| 156 | 1 | A3GALT2 | unavailable | False | 33306765 | GGCGGCGCTAGTTCCGCAGCA | [] |
| 157 | 1 | PHC2 | {'ess': 0, 'nonest': 7} | True | 33324864 | ACCAGCCCTAGGAGTCCTTGA | [] |
| 158 | 1 | ZSCAN20 | {'ess': 0, 'nonest': 7} | True | 33495475 | CCCTACGACGCCTTCCCTCCT | [] |
| 159 | 1 | CSMD2 | {'ess': 0, 'nonest': 7} | True | 33519517 | GCCGGGTGGCTATACTGCTGT | [] |
| 160 | 1 | C1orf94 | {'ess': 0, 'nonest': 7} | True | 34216760 | AGAGGAGATCTAAAAGTTTATG | [] |
| 161 | 1 | GJA4 | {'ess': 0, 'nonest': 7} | True | 34795214 | CTCTATACATACTGCTTCTTAG | [] |
| 162 | 1 | TMEM35B | unavailable | False | 34981943 | GCTCTACTTCCAGGATTCCTTG | [] |
| 163 | 1 | ZMYM6 | {'ess': 0, 'nonest': 7} | True | 34987103 | TATTGCTACTCTTTCTCCTTCG | ['TTCACTAATTTTTCTATCCTTG'] |
| 164 | 1 | SFPQ | {'ess': 8, 'nonest': 1} | False | 35184455 | CATCTAAAATCGGGGTTTTTTG | [] |
| 165 | 1 | SFPQ | {'ess': 8, 'nonest': 1} | False | 35186945 | ACACACCTAAGTTGTGAAAATG | [] |
| 166 | 1 | KIAA0319L | {'ess': 1, 'nonest': 7} | True | 35434893 | GTGGCTACAGGATCTCCTCCC | [] |
| 167 | 1 | KIAA0319L | {'ess': 1, 'nonest': 7} | True | 35447199 | GTCTCTCCCTATTTTAGACTAG | [] |
| 168 | 1 | AGO1 | unavailable | False | 35919708 | -1 | -1 |
| 169 | 1 | TEKT2 | {'ess': 0, 'nonest': 7} | True | 36086185 | TCCAAGGCTAGGCCAGCTCCA | [] |
| 170 | 1 | THRAP3 | {'ess': 2, 'nonest': 5} | False | 36304016 | GTGGCCCCTACTCGGTTGTGG | [] |
| 171 | 1 | SH3D21 | unavailable | False | 36327879 | AAGCAGCTAGGCAGGGGGTGG | [] |
| 172 | 1 | OSCP1 | {'ess': 0, 'nonest': 7} | True | 36418138 | CAGCTATAACTCATCCATCATG | [] |
| 173 | 1 | CSF3R | {'ess': 0, 'nonest': 5} | True | 36466356 | AAGCCCTAGAAGCTCCCCAGC | [] |
| 174 | 1 | GRIK3 | {'ess': 0, 'nonest': 7} | True | 36801850 | CTGTGCCTAGGGGAACACAGG | [] |
| 175 | 1 | MEAF6 | {'ess': 2, 'nonest': 6} | False | 37494098 | GTGTCTTCTAATAGTCCTAAAG | [] |
| 176 | 1 | SNIP1 | {'ess': 7, 'nonest': 1} | False | 37537747 | TTCTTAGTTTGCTAGCTGTCAG | [] |
| 177 | 1 | RSPO1 | {'ess': 0, 'nonest': 5} | True | 37612754 | TGTCCCTAGGCAGGCCCTGCA | [] |
| 178 | 1 | EPHA10 | {'ess': 0, 'nonest': 7} | True | 37761366 | CGCTCTACACTCCCACCCCAC | [] |
| 179 | 1 | SF3A3 | {'ess': 6, 'nonest': 2} | False | 37958185 | CTGAACACTAGAGCAGTCCTTG | [] |
| 180 | 1 | LOC10798 | unavailable | False | 38055586 | GAGTACTATAGACCGGGAATTG | [] |
| 181 | 1 | RRAGC | {'ess': 2, 'nonest': 5} | False | 38839552 | TCAGACTAGATGGCGTTTCGTG | [] |
| 182 | 1 | MYCBP | {'ess': 0, 'nonest': 7} | True | 38864959 | GAGAAGAATCCTATTCAGCACG | [] |
| 183 | 1 | LOC10537 | unavailable | False | 39490021 | AGCCCCGGCTTCTAGGCGGTG | [] |
| 184 | 1 | BMP8A | {'ess': 0, 'nonest': 7} | True | 39512345 | TTCCTACTGCCTGGCATAAAAG | [] |
| 185 | 1 | PABPC4 | {'ess': 1, 'nonest': 7} | True | 39561697 | CTTGTCTAAGAGGTAGCAGCAG | [] |
| 186 | 1 | NT5C1A | {'ess': 0, 'nonest': 7} | True | 39659120 | GTTCAGCTACTGTGCAGATGG | [] |
| 187 | 1 | HPCAL4 | {'ess': 0, 'nonest': 7} | True | 39682535 | CACCAGCTTCTACTTCTGCATG | [] |
| 188 | 1 | PPIE | {'ess': 7, 'nonest': 1} | False | 39762623 | AGGAAGCTTCTAGATGGTGTGG | [] |
| 189 | 1 | PPIE | {'ess': 7, 'nonest': 1} | False | 39763741 | CGAGCTCTACGTCAGCTTCCA | [] |
| 190 | 1 | BMP8B | {'ess': 0, 'nonest': 7} | True | 39773865 | TTCCTACTGCCTGGCATAAAAG | [] |
| 191 | 1 | MFSD2A | {'ess': 0, 'nonest': 7} | True | 39966567 | GGCCCTAGAGGATGCTAGCCA | [] |
| 192 | 1 | CTPS1 | unavailable | False | 40999831 | TGGTTTCTACTATATAAATAAG | ['ATAAATAAACCATTACAAATTG'] |
| 193 | 1 | EDN2 | {'ess': 0, 'nonest': 7} | True | 41479408 | ACACTATCTCTTCCTCCACCTG | [] |
| 194 | 1 | HIVEP3 | {'ess': 0, 'nonest': 5} | True | 41510450 | AGAGAGGCTAAGCGTTGGGGG | [] |
| 195 | 1 | GUCA2A | {'ess': 0, 'nonest': 7} | True | 42162905 | GCCCCCCTAGCATCGGGTACA | [] |
| 196 | 1 | FOXJ3 | {'ess': 0, 'nonest': 5} | True | 42179709 | AGAAACAAGCCCTACACAATTG | [] |
| 197 | 1 | ZMYND12 | {'ess': 0, 'nonest': 7} | True | 42430735 | GGGTCACTAAGTAATGGGATGG | [] |
| 198 | 1 | PPIH | {'ess': 2, 'nonest': 6} | False | 42667418 | TGGACTACATCTCCCCACACT | [] |
| 199 | 1 | CLDN19 | {'ess': 0, 'nonest': 7} | True | 42735761 | GAGCACCTATGCCCATCCTAT | [] |
| 200 | 1 | ERMAP | {'ess': 0, 'nonest': 7} | True | 42843231 | GCATATCCCTAAAAAGAAGGAG | [] |
| 201 | 1 | FAM183A | {'ess': 1, 'nonest': 7} | True | 43156312 | CGCTGGGATGCTACTTGTGGT | [] |
| 202 | 1 | TIE1 | {'ess': 1, 'nonest': 6} | True | 43314665 | TGACTGGCCTACTGGCAGGTG | [] |
| 203 | 1 | KDM4A | {'ess': 2, 'nonest': 5} | False | 43704369 | GCACCTACTCCATGATGGCCC | [] |
| 204 | 1 | ST3GAL3 | {'ess': 1, 'nonest': 7} | True | 43800792 | GCTTGCTACCTACTTGACACTG | [] |
| 205 | 1 | ST3GAL3 | {'ess': 1, 'nonest': 7} | True | 43920350 | ATGAGCCTAGGGGAGTGGCAG | [] |
| 206 | 1 | ATP6V0B | {'ess': 7, 'nonest': 1} | False | 43978006 | CTAGTCACCCATCTTCACTCTG | [] |
| 207 | 1 | CCDC24 | {'ess': 0, 'nonest': 7} | True | 43996178 | CCAGAGCCTACTCGGTGACCA | [] |
| 208 | 1 | SLC6A9 | {'ess': 1, 'nonest': 6} | True | 43998123 | CCAAACTAGAAAGGGCATTTG | [] |
| 209 | 1 | KLF17 | {'ess': 0, 'nonest': 7} | True | 44130755 | CTGACCTAAGGACCAGCAGCA | [] |
| 210 | 1 | ARMH1 | unavailable | False | 44708780 | ACGTGGCTAACCTGACTCACT | [] |
| 211 | 1 | ARMH1 | unavailable | False | 44722246 | ATGTGGCTACATGGAAGACAGG | [] |
| 212 | 1 | PLK3 | {'ess': 1, 'nonest': 6} | True | 44805677 | TGGGTCCTAGGCTGGGCTGCG | [] |
| 213 | 1 | PTCH2 | {'ess': 0, 'nonest': 7} | True | 44820688 | GCAGAGGCTAGATCTCCTCGG | [] |
| 214 | 1 | GPBP1L1 | {'ess': 1, 'nonest': 5} | False | 45626255 | ATATGCCTACTTCCAGGCATCG | [] |
| 215 | 1 | MAST2 | {'ess': 1, 'nonest': 6} | True | 46030065 | ACTGCTATGTTTGCTTTAAAAG | [] |
| 216 | 1 | PIK3R3 | unavailable | False | 46175585 | CTGGGCACTTCTAGAGGATCA | [] |
| 217 | 1 | NSUN4 | {'ess': 5, 'nonest': 3} | False | 46361845 | GGTGATACTATGTCAGCCTACG | [] |
| 218 | 1 | TMEM275 | unavailable | False | 46532993 | GTCCGCTAGGGGGCGGCGCG | [] |
| 219 | 1 | ATPAF1 | {'ess': 0, 'nonest': 5} | True | 46635775 | ACAGTTCTAAGTCTTATTTTGG | [] |
| 220 | 1 | TEX38 | {'ess': 0, 'nonest': 4} | True | 46673455 | GAGGAGGCTACAGTTCTGAAG | [] |
| 221 | 1 | EFCAB14 | unavailable | False | 46679850 | TGATGAAGCTAGATACCTAAAG | [] |
| 222 | 1 | CYP4B1 | {'ess': 0, 'nonest': 7} | True | 46818813 | CAGAGCTACTTCCCAGACCCA | [] |
| 223 | 1 | CYP4X1 | {'ess': 0, 'nonest': 7} | True | 47050173 | CTGAGATCTAACATTCAGAGAG | [] |
| 224 | 1 | TRABD2B | {'ess': 0, 'nonest': 4} | True | 47765774 | AGTGTCTAGCCAATCCCATGTG | [] |
| 225 | 1 | SPATA6 | {'ess': 0, 'nonest': 7} | True | 48351727 | CATTGGTCTACATTTTTCTTTG | [] |
| 226 | 1 | DMRTA2 | {'ess': 0, 'nonest': 7} | True | 50418664 | TCGCCCTACTGCTTCTCCGGA | [] |
| 227 | 1 | TTC39A | {'ess': 0, 'nonest': 7} | True | 51288156 | CAAAGCTACAAGGACACTGATG | [] |
| 228 | 1 | TTC39A | {'ess': 0, 'nonest': 7} | True | 51290347 | TGGCCTACACAGTGGGAGCAC | [] |
| 229 | 1 | OSBPL9 | {'ess': 2, 'nonest': 7} | False | 51787788 | CTTCCAACCTAATGCTTGGCAG | [] |
| 230 | 1 | RAB3B | {'ess': 0, 'nonest': 6} | True | 51919926 | TTGCCTAGCATGAGCAGTTCTG | [] |
| 231 | 1 | BTF3L4 | {'ess': 1, 'nonest': 6} | True | 52086719 | TTTTCTACAAGATCTGAAAAAG | [] |
| 232 | 1 | TUT4 | unavailable | False | 52463562 | ACACCACTACAGATGCCAGAAG | [] |
| 233 | 1 | PODN | {'ess': 0, 'nonest': 7} | True | 53082160 | -1 | -1 |
| 234 | 1 | MAGOH | {'ess': 8, 'nonest': 1} | False | 53227044 | AGTCTAGATTGGTTTAATCTTG | [] |
| 235 | 1 | DMRTB1 | {'ess': 0, 'nonest': 7} | True | 53466661 | GGGCCTAGTCGGACTGCTCCT | [] |
| 236 | 1 | NDC1 | unavailable | False | 53767969 | TAACTATTCTTTGAACTCCAAG | [] |
| 237 | 1 | CYB5RL | {'ess': 0, 'nonest': 1} | False | 54174618 | CCAGGGCTAGAAGAGGAAATAG | [] |
| 238 | 1 | ACOT11 | {'ess': 1, 'nonest': 7} | True | 54609611 | GGGCATCTAGAGGGTCTGGAG | [] |
| 239 | 1 | TTC22 | {'ess': 0, 'nonest': 7} | True | 54781242 | AGCCCTAGAATGAGACAGCCC | [] |
| 240 | 1 | LEXM | unavailable | False | 54824990 | TTATACTAGATGCACACATATG | [] |
| 241 | 1 | LEXM | unavailable | False | 54837248 | CCAGGCTACCTATGGGGACAT | [] |
| 242 | 1 | LEXM | unavailable | False | 54841575 | GAGATCCTACATTGGTGAGAGG | [] |
| 243 | 1 | USP24 | {'ess': 0, 'nonest': 7} | True | 55069044 | TCTAGGGATCAACATCATCAAG | [] |
| 244 | 1 | PLPP3 | unavailable | False | 56496550 | TGGCACCTACATCATGTTGTGG | [] |
| 245 | 1 | FYB2 | unavailable | False | 56719670 | TTTTCTAAGGTGACCAACTTTG | [] |
| 246 | 1 | FYB2 | unavailable | False | 56731891 | CATCTTACCTAACAGGGAAAAG | [] |
| 247 | 1 | C8B | {'ess': 0, 'nonest': 7} | True | 56829403 | ATCTGCTAGGAGCAGTCAAGTG | [] |
| 248 | 1 | DAB1 | {'ess': 0, 'nonest': 5} | True | 57010694 | GCTATCTAGCTACCGGCCTGT | [] |
| 249 | 1 | TACSTD2 | {'ess': 1, 'nonest': 6} | True | 58576184 | CGCCGGGTACCTACAAGCTCG | [] |
| 250 | 1 | PATJ | unavailable | False | 61991067 | GGTCCTAATAGACACTCAAAAG | [] |
| 251 | 1 | L1TD1 | {'ess': 0, 'nonest': 7} | True | 62211371 | GCTAAGGTATATTATTCCCCAG | [] |
| 252 | 1 | KANK4 | {'ess': 0, 'nonest': 7} | True | 62238276 | TGCAGCCCCTACAGCCCCAGG | [] |
| 253 | 1 | USP1 | {'ess': 2, 'nonest': 6} | False | 62450980 | TCTATAATTTCTTATAAAATAG | [] |
| 254 | 1 | FOXD3 | {'ess': 1, 'nonest': 7} | True | 63324694 | TCCCTATTGCGCCGGCCATTT | [] |
| 255 | 1 | ALG6 | {'ess': 2, 'nonest': 6} | False | 63437019 | ACAGCTAGCTGATTTTCTTCTG | [] |
| 256 | 1 | ITGB3BP | {'ess': 0, 'nonest': 7} | True | 63449777 | ACGTCTAAACTTCAAACCTCAG | [] |
| 257 | 1 | EFCAB7 | {'ess': 0, 'nonest': 7} | True | 63552940 | ACAGCTACTGTTTTAGACAATG | [] |
| 258 | 1 | ROR1 | {'ess': 0, 'nonest': 6} | True | 64142657 | GACTTCTATTTACCGCACGCTG | [] |
| 259 | 1 | UBE2U | {'ess': 1, 'nonest': 7} | True | 64244158 | TTCCCCTATGCTGCTTCCAAAG | [] |
| 260 | 1 | LEPROT | {'ess': 0, 'nonest': 7} | True | 65431918 | AAAGTGCTACCACTGCTCCCA | [] |
| 261 | 1 | LEPR | {'ess': 0, 'nonest': 7} | True | 65626169 | TTCCTAAGTCAACCAACCTCCG | ['CCTCCACCCAGTAGTTCCTTTG'] |
| 262 | 1 | MIER1 | {'ess': 0, 'nonest': 7} | True | 66946309 | CTCTACATGTTATTTACCCTTG | [] |
| 263 | 1 | C1orf141 | {'ess': 0, 'nonest': 5} | True | 67109158 | CCGTCTCTACTAAAAATACAAG | ['TACAAAAAAATTAGCCAGCATG'] |
| 264 | 1 | IL23R | {'ess': 0, 'nonest': 7} | True | 67259127 | CAGCTCTACTTTTCCAAGAGTG | [] |
| 265 | 1 | GNG12 | {'ess': 2, 'nonest': 7} | False | 67705450 | TTCCACTATAAGATGATGCAAG | [] |
| 266 | 1 | WLS | {'ess': 0, 'nonest': 6} | True | 68126225 | CTCCTACTCCTGGGCCTCCTT | [] |
| 267 | 1 | LRRC7 | {'ess': 0, 'nonest': 7} | True | 70128553 | AGCTACAGTAAATGTTTTCCTG | [] |
| 268 | 1 | HHLA3 | {'ess': 0, 'nonest': 7} | True | 70366530 | TTTACCTTACTCTATGGATTCG | [] |
| 269 | 1 | CTH | {'ess': 0, 'nonest': 7} | True | 70439126 | TACTAGCTGTGACTTCCACTTG | [] |
| 270 | 1 | NEGR1 | {'ess': 0, 'nonest': 7} | True | 71587377 | CCTTTTGGATACTAGCTACTGG | ['CTACTGACTTCTTTTTTATCTG'] |
| 271 | 1 | LRRIQ3 | {'ess': 0, 'nonest': 7} | True | 74095060 | GCTTCTAGGAACTTTCCAAAAG | [] |
| 272 | 1 | FPGT | {'ess': 0, 'nonest': 7} | True | 74205831 | AAAATATCTCTACATCAAACTG | [] |
| 273 | 1 | ERICH3 | unavailable | False | 74571116 | AGTCTCCTAGACCTGCACGTT | [] |
| 274 | 1 | SLC44A5 | {'ess': 0, 'nonest': 6} | True | 75203726 | TCTTCTACTGCTTCCTAGTTTG | [] |
| 275 | 1 | SLC44A5 | {'ess': 0, 'nonest': 6} | True | 75206606 | TTTTCTACTCTTTCTGCTTCTG | [] |
| 276 | 1 | RABGGTB | {'ess': 7, 'nonest': 1} | False | 75794549 | TGAATCTAGCTCACTAGCTCAG | [] |
| 277 | 1 | PIGK | {'ess': 2, 'nonest': 6} | False | 77092373 | TCTAAAAAATGAACTTCATATG | [] |
| 278 | 1 | AK5 | {'ess': 0, 'nonest': 7} | True | 77502677 | TTCTTCCTATTGTTTTGGTGAG | [] |
| 279 | 1 | NEXN | {'ess': 0, 'nonest': 7} | True | 77942912 | GAGGGGAGCTAGTAGTCATCA | [] |
| 280 | 1 | FUBP1 | {'ess': 3, 'nonest': 4} | False | 77945611 | ATGAGCTATGATTTGGTGCTTG | [] |
| 281 | 1 | FUBP1 | {'ess': 3, 'nonest': 4} | False | 77947513 | CTCCTCACCCCTACAGAGCTA | [] |
| 282 | 1 | DNAJB4 | {'ess': 0, 'nonest': 7} | True | 78016246 | CTTCATTCTATGAGGCAGGAAG | [] |
| 283 | 1 | PTGFR | {'ess': 1, 'nonest': 6} | True | 78503941 | CCTTCTATTTTCTGGTCCACAG | [] |
| 284 | 1 | PTGFR | {'ess': 1, 'nonest': 6} | True | 78536686 | CTATTAAGCTAGGTGCTTGCTG | [] |
| 285 | 1 | IFI44 | {'ess': 0, 'nonest': 7} | True | 78661390 | AACTAAGGCTCAAAAAAAATTG | [] |
| 286 | 1 | IFI44 | {'ess': 0, 'nonest': 7} | True | 78663810 | CATATCTATTTTTTTCCTTGTG | [] |
| 287 | 1 | TTLL7 | {'ess': 0, 'nonest': 7} | True | 83912488 | TATACTAGTTGCTTGTTTTCTG | [] |
| 288 | 1 | SAMD13 | {'ess': 0, 'nonest': 7} | True | 84349773 | TTGACTGTACTATGAAGAGTTG | [] |
| 289 | 1 | GNG5 | {'ess': 0, 'nonest': 7} | True | 84501844 | ATTTTACTACAAAAAGGAACAG | [] |
| 290 | 1 | SPATA1 | {'ess': 0, 'nonest': 1} | False | 84552123 | ACATACCTAATTGAGATTGTAG | [] |
| 291 | 1 | SSX2IP | {'ess': 0, 'nonest': 7} | True | 84647432 | TCCACATGTCTAAGGTAAGTCG | ['AAGTCATCTTTTTCTACATGAG'] |
| 292 | 1 | MCOLN3 | {'ess': 0, 'nonest': 7} | True | 85019122 | TGATAGCTACTTTTTACAACAG | [] |
| 293 | 1 | C1orf52 | {'ess': 0, 'nonest': 7} | True | 85252628 | TGTTTCTACTTTTTCTTCTTTG | [] |
| 294 | 1 | COL24A1 | {'ess': 1, 'nonest': 7} | True | 85960769 | TTCCAGCTATCCCGGGAGGCT | [] |
| 295 | 1 | CLCA1 | {'ess': 0, 'nonest': 5} | True | 86500044 | TTCAGCCCTAGGCTATTGACAG | [] |
| 296 | 1 | HS2ST1 | {'ess': 0, 'nonest': 7} | True | 87097938 | TCTCCCCTACCAGCATTCGGA | [] |
| 297 | 1 | RBMXL1 | {'ess': 0, 'nonest': 7} | True | 88992653 | TTGTTTCTAGTATCTGCTTCTG | [] |
| 298 | 1 | ZNF644 | {'ess': 0, 'nonest': 7} | True | 90916797 | CTGGTTCTATGAAGCTGCTTCG | [] |
| 299 | 1 | TGFBR3 | {'ess': 1, 'nonest': 4} | False | 91693738 | GTTGGGCTAGGCCGTGCTGCT | [] |
| 300 | 1 | C1orf146 | {'ess': 0, 'nonest': 7} | True | 92245673 | GTACTCTAATTTGGGTTAACTG | [] |
| 301 | 1 | DIPK1A | unavailable | False | 92843382 | AATGAACTAAGAGTCATTAGTG | [] |
| 302 | 1 | TMED5 | {'ess': 0, 'nonest': 7} | True | 93154826 | TGCTCTAAGCAGAGTTTGTATG | [] |
| 303 | 1 | CCDC18 | {'ess': 0, 'nonest': 1} | False | 93245492 | TGAAACTCTAAGTTCTGATAAG | [] |
| 304 | 1 | ARHGAP2 | {'ess': 0, 'nonest': 7} | True | 94173868 | GACATCCCTACACAAATTGTGG | [] |
| 305 | 1 | ABCD3 | {'ess': 0, 'nonest': 7} | True | 94517128 | CTCTAAGAGCCAAACTCAACTG | [] |
| 306 | 1 | SLC44A3 | {'ess': 0, 'nonest': 7} | True | 94894921 | GTATCTATCTCACAATGGCCTG | [] |
| 307 | 1 | CNN3 | {'ess': 0, 'nonest': 7} | True | 94897741 | GGATCTAATAATCAATGCCTTG | [] |
| 308 | 1 | RWDD3 | {'ess': 0, 'nonest': 7} | True | 95246517 | TACACTAATCATTTTCTCTTTG | [] |
| 309 | 1 | FRRS1 | {'ess': 0, 'nonest': 7} | True | 99708897 | TCCTCTACAGACATGACCCCA | [] |
| 310 | 1 | AGL | {'ess': 0, 'nonest': 7} | True | 99921650 | ATAAACTATAAATCATAAAGTG | [] |
| 311 | 1 | SLC35A3 | {'ess': 0, 'nonest': 7} | True | 1E+08 | TATACAACTATGCTTTAGTGGG | [] |
| 312 | 1 | RTCA | unavailable | False | 1E+08 | ATACTCTATAGATTTGGATTTG | [] |
| 313 | 1 | CDC14A | {'ess': 1, 'nonest': 6} | True | 1E+08 | AGAATCTCCTAGGGAAAACTTG | ['GAAAACTTACCTCTCCAAATCG'] |
| 314 | 1 | VCAM1 | {'ess': 1, 'nonest': 6} | True | 1.01E+08 | TTAGCTACACTTTTGACTTCTG | [] |
| 315 | 1 | SLC30A7 | {'ess': 1, 'nonest': 7} | True | 1.01E+08 | AACCGTTCTAAAACTTTTGTGG | [] |
| 316 | 1 | SLC30A7 | {'ess': 1, 'nonest': 7} | True | 1.01E+08 | CATTCACTACATGGCTGCAAAG | [] |
| 317 | 1 | DPH5 | {'ess': 4, 'nonest': 4} | False | 1.01E+08 | TCTGAGCTATTTTCTGGTATGG | [] |
| 318 | 1 | S1PR1 | {'ess': 2, 'nonest': 5} | False | 1.01E+08 | CCAGTTCTAGGAAGAAGAGTTG | [] |
| 319 | 1 | OLFM3 | {'ess': 0, 'nonest': 7} | True | 1.02E+08 | TTTGCCTATGTGTCATCCTCTG | [] |
| 320 | 1 | COL11A1 | {'ess': 2, 'nonest': 6} | False | 1.03E+08 | ACAGCTAGGTTGTATTTAAAAG | [] |
| 321 | 1 | NTNG1 | {'ess': 0, 'nonest': 7} | True | 1.07E+08 | TGACACCTAGAACACCAGGGG | [] |
| 322 | 1 | FAM102B | {'ess': 0, 'nonest': 7} | True | 1.09E+08 | TCTAGCGTTTGATCACAACTTG | [] |
| 323 | 1 | STXBP3 | {'ess': 1, 'nonest': 6} | True | 1.09E+08 | ATGCTATTCATCTTTAATTAAG | [] |
| 324 | 1 | AKNAD1 | {'ess': 1, 'nonest': 6} | True | 1.09E+08 | CGTTGAACTAGTATTTCAGTCG | [] |
| 325 | 1 | GPSM2 | {'ess': 1, 'nonest': 5} | False | 1.09E+08 | CATAGTAACTAATGGTCTGCCG | [] |
| 326 | 1 | CLCC1 | {'ess': 1, 'nonest': 5} | False | 1.09E+08 | GTTCCTCTAGCCACAGGGGCT | [] |
| 327 | 1 | WDR47 | {'ess': 0, 'nonest': 7} | True | 1.09E+08 | GGTGTGCTCTACCCATTGTAAG | [] |
| 328 | 1 | TMEM167 | {'ess': 0, 'nonest': 7} | True | 1.09E+08 | ACTCTCCTACACTGGGTTCAG | [] |
| 329 | 1 | C1orf194 | {'ess': 0, 'nonest': 7} | True | 1.09E+08 | TCAACACTAGGTGGAGAAGAAG | [] |
| 330 | 1 | KIAA1324 | {'ess': 0, 'nonest': 7} | True | 1.09E+08 | AACTATGATTCAGCTCCCAAAG | [] |
| 331 | 1 | PSRC1 | {'ess': 2, 'nonest': 6} | False | 1.09E+08 | TTTGCTACTGAAGTCTTGCTTG | [] |
| 332 | 1 | SORT1 | {'ess': 1, 'nonest': 6} | True | 1.09E+08 | AGAGCTATTCCAAGAGGTCCTG | [] |
| 333 | 1 | GSTM2 | {'ess': 0, 'nonest': 7} | True | 1.1E+08 | AAGGCCCTACTTGTTGCCCCA | [] |

| | | | | | | | |
|---|---|---|---|---|---|---|---|
| 334 | 1 | GSTM1 | {'exs': 0, 'nonexs': 7} | True | 1.1E+08 | AAGGCCCTACTTGTTGCCCCA | [] |
| 335 | 1 | GSTM5 | {'exs': 0, 'nonexs': 7} | True | 1.1E+08 | GGGCCCTATTTGCTGTTCCAT | [] |
| 336 | 1 | EPS8L3 | {'exs': 0, 'nonexs': 7} | True | 1.1E+08 | GGTGCCTAAGGGCTTATCTGA | [] |
| 337 | 1 | CSF1 | {'exs': 2, 'nonexs': 5} | False | 1.1E+08 | AGAATTCCCTCTACACTGGCAG | [] |
| 338 | 1 | KCNC4 | {'exs': 0, 'nonexs': 7} | True | 1.1E+08 | GGTGCAGCTAAGGGGCAGAGT | [] |
| 339 | 1 | KCNC4 | {'exs': 0, 'nonexs': 7} | True | 1.1E+08 | ACAGAGCTAGATCTTGCCTAAG | ['TTGCCTAAAAAAAAAAAAAAAG'] |
| 340 | 1 | RBM15 | {'exs': 1, 'nonexs': 7} | True | 1.1E+08 | AACCACTATAACAGGGTCAGC | [] |
| 341 | 1 | PROK1 | {'exs': 1, 'nonexs': 7} | True | 1.1E+08 | GGCAAGCGCCTAAAAATTGATG | [] |
| 342 | 1 | WDR77 | {'exs': 7, 'nonexs': 1} | False | 1.11E+08 | CCAATCTACTCAGTAACACTTG | [] |
| 343 | 1 | ADORA3 | {'exs': 0, 'nonexs': 4} | True | 1.11E+08 | CTACTCAGAATTCTTCTCAATG | [] |
| 344 | 1 | ADORA3 | {'exs': 0, 'nonexs': 4} | True | 1.12E+08 | TAATTGCTAAAGGGTAGGGGAG | [] |
| 345 | 1 | RAP1A | {'exs': 0, 'nonexs': 7} | True | 1.12E+08 | TGGGCCTAGAGCAGCAGACAT | [] |
| 346 | 1 | CTTNBP2 | {'exs': 1, 'nonexs': 7} | True | 1.12E+08 | CTAGGGACTAGCTGCTGGTAG | [] |
| 347 | 1 | ST7L | {'exs': 0, 'nonexs': 7} | True | 1.13E+08 | CCCAGCTACTTGGGAGGCTGA | [] |
| 348 | 1 | CAPZA1 | {'exs': 0, 'nonexs': 7} | True | 1.13E+08 | GGCATGCTCCTATGGCCAGGA | [] |
| 349 | 1 | MOV10 | {'exs': 0, 'nonexs': 7} | True | 1.13E+08 | ATTAACCTAATGCAAGAGGGAG | [] |
| 350 | 1 | MOV10 | {'exs': 0, 'nonexs': 7} | True | 1.13E+08 | CTCAGTCTAAGAATGACAGGAG | [] |
| 351 | 1 | TAFA3 | unavailable | False | 1.13E+08 | CAAGAGCTATCGTGTGACCTG | [] |
| 352 | 1 | RSBN1 | {'exs': 1, 'nonexs': 7} | True | 1.14E+08 | CTGCTCTCTCTACTGCAGAGA | [] |
| 353 | 1 | HIPK1 | {'exs': 1, 'nonexs': 6} | True | 1.14E+08 | CCAACTATAAGTAGGAATACTG | [] |
| 354 | 1 | TRIM33 | {'exs': 1, 'nonexs': 6} | True | 1.14E+08 | GATACTACGGGAAAGGGTCCA | [] |
| 355 | 1 | CASQ2 | {'exs': 0, 'nonexs': 7} | True | 1.16E+08 | GTTGGGCTATTCATCATCATCG | [] |
| 356 | 1 | NHLH2 | {'exs': 0, 'nonexs': 7} | True | 1.16E+08 | CCCGCCCTACACGTCCAGGAC | [] |
| 357 | 1 | SLC22A15 | {'exs': 0, 'nonexs': 7} | True | 1.16E+08 | ACCAGGCGCTAAGGATCAGGA | [] |
| 358 | 1 | ATP1A1 | {'exs': 6, 'nonexs': 2} | False | 1.16E+08 | AGGACGGGGGGCTAATAGTAG | [] |
| 359 | 1 | PTGFRN | {'exs': 0, 'nonexs': 7} | True | 1.17E+08 | CCAGCCTAGTCCATCTCCATC | [] |
| 360 | 1 | MAN1A2 | {'exs': 0, 'nonexs': 7} | True | 1.18E+08 | GGAGGCTAACTTCAAGATCCAG | [] |
| 361 | 1 | WDR3 | {'exs': 9, 'nonexs': 0} | False | 1.18E+08 | TTCAGTTCTAAGTCAACGTTAG | [] |
| 362 | 1 | SPAG17 | {'exs': 0, 'nonexs': 7} | True | 1.18E+08 | CCTAATGTGGAGAAACTTTACG | [] |
| 363 | 1 | WARS2 | {'exs': 7, 'nonexs': 1} | False | 1.19E+08 | ACTTCCTATAGAAAACCCACCG | ['CCACCAATTTCTTCACCTCCTG'] |
| 364 | 1 | ZNF697 | {'exs': 0, 'nonexs': 7} | True | 1.2E+08 | CCAGCCCCTAACACAGGTGCA | [] |
| 365 | 1 | REG4 | {'exs': 0, 'nonexs': 7} | True | 1.2E+08 | CTTGCTCTATGGTCGGTACTTG | [] |
| 366 | 1 | REG4 | {'exs': 0, 'nonexs': 7} | True | 1.2E+08 | CTTTCCTATGTATCCGTATGTG | [] |
| 367 | 1 | NBPF7 | unavailable | False | 1.2E+08 | TTCAACTAGTGCCAAGAAAGTG | [] |
| 368 | 1 | LOC10192 | unavailable | False | 1.45E+08 | CCACTGCTACCGTGTTCTAGT | [] |
| 369 | 1 | LIX1L | {'exs': 1, 'nonexs': 4} | False | 1.46E+08 | ATGCCTAGCAGTTGGAAGAATG | [] |
| 370 | 1 | NOTCH2N | unavailable | False | 1.46E+08 | CCAGTGTCTAATTCTCATCGTG | [] |
| 371 | 1 | PRKAB2 | {'exs': 1, 'nonexs': 7} | True | 1.47E+08 | TGCGCTAAGGACCATCACACT | [] |
| 372 | 1 | FMO5 | {'exs': 0, 'nonexs': 7} | True | 1.47E+08 | CAATAGGACAACTAGAAGTAAG | [] |
| 373 | 1 | FMO5 | {'exs': 0, 'nonexs': 7} | True | 1.47E+08 | TTTCTATACACAGCCTCTTCTG | ['CCTCTTCTAATTCTGAAAAGTG'] |
| 374 | 1 | ACP6 | {'exs': 2, 'nonexs': 7} | False | 1.48E+08 | CAGGACTACTTGGCATACTCC | ['CTCCATCTTCAACGATAGTAAG'] |
| 375 | 1 | ACP6 | {'exs': 2, 'nonexs': 7} | False | 1.48E+08 | TTGGTGCTAACTCAGGTAGGTG | [] |
| 376 | 1 | NBPF11 | {'exs': 0, 'nonexs': 1} | False | 1.48E+08 | TACAACCTATATGCACCCATAG | [] |
| 377 | 1 | NOTCH2N | unavailable | False | 1.49E+08 | CCAGTGTCTAATTCTCATCGTG | [] |
| 378 | 1 | NUDT4B | unavailable | False | 1.49E+08 | TCTCTCTATCTTACACTAGATG | [] |
| 379 | 1 | PDE4DIP | {'exs': 5, 'nonexs': 4} | False | 1.49E+08 | AGGCACCTACTTGTCCCTCTG | [] |
| 380 | 1 | PDE4DIP | {'exs': 5, 'nonexs': 4} | False | 1.49E+08 | TTCTGAAGCTATAGCTACAGAG | [] |
| 381 | 1 | NOTCH2N | unavailable | False | 1.49E+08 | CCAGTGTCTAATTCTCATCGTG | [] |
| 382 | 1 | NBPF19 | unavailable | False | 1.5E+08 | GGCTGCCTATTGTGGGAATATG | [] |
| 383 | 1 | FCGR1A | {'exs': 3, 'nonexs': 5} | False | 1.5E+08 | CCGCTGCTACGTGGCCCCCTG | [] |
| 384 | 1 | LOC11226 | unavailable | False | 1.5E+08 | TGATTCTAGATTATTTGGTTTG | [] |
| 385 | 1 | H4C14 | unavailable | False | 1.5E+08 | GGCGGCCTAGCCTCCGAAGCC | [] |
| 386 | 1 | LOC11226 | unavailable | False | 1.5E+08 | ATTGGCCTATCCGCACTGGCC | [] |
| 387 | 1 | H4C15 | unavailable | False | 1.5E+08 | GGCGGCCTAGCCTCCGAAGCC | [] |
| 388 | 1 | MTMR11 | {'exs': 0, 'nonexs': 7} | True | 1.5E+08 | CTCTACCCCAGATCCCCCTCT | [] |
| 389 | 1 | VPS45 | {'exs': 4, 'nonexs': 4} | False | 1.5E+08 | CCGTGGGCTACCTGGAGATGA | [] |
| 390 | 1 | ANP32E | {'exs': 1, 'nonexs': 7} | True | 1.5E+08 | CTAGTCATCTTCTTCCTCTCCG | ['CTCCATCGTCTTCAGCATCTCG'] |
| 391 | 1 | C1orf54 | {'exs': 0, 'nonexs': 7} | True | 1.5E+08 | CTTACCTTCTACATGAAATACG | ['ATACATCCCCACCTGAACAAAG'] |
| 392 | 1 | CIART | unavailable | False | 1.5E+08 | CCCTGGGCTAGAGGTTGAGAA | [] |
| 393 | 1 | ADAMTSL | {'exs': 2, 'nonexs': 6} | False | 1.51E+08 | TTCTATATTGGGGGTCCATATG | [] |
| 394 | 1 | ADAMTSL | {'exs': 2, 'nonexs': 6} | False | 1.51E+08 | TTGAGCTATGCTGGAAAGGCTG | [] |
| 395 | 1 | MCL1 | {'exs': 5, 'nonexs': 3} | False | 1.51E+08 | TAAGGCTATCTTATTAGATATG | [] |
| 396 | 1 | CTSS | {'exs': 1, 'nonexs': 6} | True | 1.51E+08 | ATCCTCTAGATTTCTGGGTAAG | [] |
| 397 | 1 | ARNT | {'exs': 0, 'nonexs': 7} | True | 1.51E+08 | ATAGTTCTATTCTGAAAAGGGG | [] |
| 398 | 1 | SETDB1 | {'exs': 2, 'nonexs': 6} | False | 1.51E+08 | CCTCTAAAGAAGACGTCCTCTG | [] |
| 399 | 1 | MINDY1 | unavailable | False | 1.51E+08 | GCAGAGCTACAGCAGAATGCA | [] |
| 400 | 1 | BNIPL | {'exs': 0, 'nonexs': 7} | True | 1.51E+08 | CCTGTGCTATGTCCCTCCTGA | [] |
| 401 | 1 | CDC42SE | {'exs': 2, 'nonexs': 5} | False | 1.51E+08 | TATTGGAGCTATAAGCCCCTAG | [] |
| 402 | 1 | ZNF687 | {'exs': 1, 'nonexs': 6} | True | 1.51E+08 | GGAGACTAGTTGTCCCCAACA | [] |
| 403 | 1 | CGN | {'exs': 0, 'nonexs': 7} | True | 1.52E+08 | CGAGCTAACAGGAGCTGGTCT | [] |
| 404 | 1 | SNX27 | {'exs': 1, 'nonexs': 7} | True | 1.52E+08 | TCTAACTAATATTCCTAGAAAG | [] |
| 405 | 1 | SNX27 | {'exs': 1, 'nonexs': 7} | True | 1.52E+08 | GCTAGGTGGCCACATCTCTCT | [] |
| 406 | 1 | C2CD4D | {'exs': 0, 'nonexs': 7} | True | 1.52E+08 | CTCAGGCTACAGGCTGAGATG | [] |
| 407 | 1 | S100A10 | {'exs': 1, 'nonexs': 6} | True | 1.52E+08 | CTGCCTACTTCTTTCCCTTCTG | [] |
| 408 | 1 | PRR9 | {'exs': 0, 'nonexs': 7} | True | 1.53E+08 | AGCAGCTACTTTCCCTTGCCA | [] |
| 409 | 1 | LORICRIN | unavailable | False | 1.53E+08 | GGGGATCTATTTGGACGGCCA | [] |
| 410 | 1 | S100A12 | {'exs': 1, 'nonexs': 6} | True | 1.53E+08 | AGCTACCTACTCTTTGTGGGTG | [] |
| 411 | 1 | S100A8 | {'exs': 1, 'nonexs': 6} | True | 1.53E+08 | GTAACTCAGCTACTCTTTGTGG | [] |
| 412 | 1 | S100A3 | {'exs': 0, 'nonexs': 7} | True | 1.54E+08 | CAGAGGCTACTGGGAGCAGGG | [] |
| 413 | 1 | S100A16 | {'exs': 0, 'nonexs': 7} | True | 1.54E+08 | GGGGTCTCTAGCTGCTGCTCT | [] |
| 414 | 1 | DENND4B | {'exs': 0, 'nonexs': 7} | True | 1.54E+08 | GGTCTCTAGCATTCTGGAGGT | [] |
| 415 | 1 | SLC39A1 | {'exs': 1, 'nonexs': 7} | True | 1.54E+08 | AGCCCCCTAGATTTGGATGAAG | [] |
| 416 | 1 | JTB | {'exs': 2, 'nonexs': 5} | False | 1.54E+08 | GTAGCTATATGGACTCGATTTG | [] |
| 417 | 1 | NUP210L | {'exs': 0, 'nonexs': 7} | True | 1.54E+08 | GGCTATGGCCAGCGACCACTA | [] |
| 418 | 1 | TPM3 | {'exs': 1, 'nonexs': 6} | True | 1.54E+08 | GGCGTTCTACATCTCATTCAGG | [] |
| 419 | 1 | HAX1 | {'exs': 0, 'nonexs': 7} | True | 1.54E+08 | TTAACAAGGCTACCGGGACCG | [] |
| 420 | 1 | IL6R | {'exs': 1, 'nonexs': 6} | True | 1.54E+08 | CCCAGCTACTTAGGAGGCTGA | [] |
| 421 | 1 | IL6R | {'exs': 1, 'nonexs': 6} | True | 1.54E+08 | AGCCAGCTATCTGGGGAAGAA | [] |
| 422 | 1 | TDRD10 | {'exs': 0, 'nonexs': 7} | True | 1.55E+08 | GACCACAAACTAGAACTGGCAG | [] |
| 423 | 1 | TDRD10 | {'exs': 0, 'nonexs': 7} | True | 1.55E+08 | TCCAGGCCTAGGCAGCAGCCT | [] |
| 424 | 1 | ADAR | {'exs': 2, 'nonexs': 5} | False | 1.55E+08 | GAGCATACTATACTGGGCAGAG | [] |
| 425 | 1 | PMVK | {'exs': 6, 'nonexs': 2} | False | 1.55E+08 | TAGTGACTAAAGTCTGGAGCGG | [] |
| 426 | 1 | LOC10192 | unavailable | False | 1.55E+08 | GAAGCCTATTGTGTACGCCGC | [] |
| 427 | 1 | SHC1 | {'exs': 4, 'nonexs': 5} | False | 1.55E+08 | CAGGCTAAGGGATGAGCCTCT | [] |
| 428 | 1 | ADAM15 | {'exs': 0, 'nonexs': 7} | True | 1.55E+08 | ACCACTAGGGGCGGGATTCCA | [] |
| 429 | 1 | EFNA3 | {'exs': 1, 'nonexs': 7} | True | 1.55E+08 | CAGAGCTAGGAGGCCAAGAAC | [] |
| 430 | 1 | TRIM46 | {'exs': 2, 'nonexs': 6} | False | 1.55E+08 | ATTCTCCTACCTCAGCCTCCC | [] |
| 431 | 1 | MUC1 | {'exs': 2, 'nonexs': 6} | False | 1.55E+08 | TGCCCCTACAAGTTGGCAGAA | [] |
| 432 | 1 | MUC1 | {'exs': 2, 'nonexs': 6} | False | 1.55E+08 | TACTGCTAGGGGGCACATAGC | [] |
| 433 | 1 | RUSC1 | {'exs': 1, 'nonexs': 7} | True | 1.55E+08 | CCAGGGCTACAGAACAAGGGA | [] |
| 434 | 1 | ASH1L | {'exs': 0, 'nonexs': 7} | True | 1.55E+08 | GGCTAGAAGCATGGCATCATAG | [] |
| 435 | 1 | MSTO1 | {'exs': 3, 'nonexs': 4} | False | 1.56E+08 | GCAGCTCTGCTACGTCATCGT | [] |
| 436 | 1 | YY1AP1 | {'exs': 1, 'nonexs': 7} | True | 1.56E+08 | GTCATATTTCCTACCTCTCTAG | [] |
| 437 | 1 | KHDC4 | unavailable | False | 1.56E+08 | TGTTTCCTAGGGAGCCATCCA | [] |
| 438 | 1 | UBQLN4 | {'exs': 0, 'nonexs': 7} | True | 1.56E+08 | GAGCTTCACTAATGGGTGATGG | [] |
| 439 | 1 | LMNA | {'exs': 1, 'nonexs': 7} | True | 1.56E+08 | GGTCTAAAGCTAGGGAAGGCA | [] |
| 440 | 1 | PMF1-BGL | unavailable | False | 1.56E+08 | TCACTACCTCGCTGCCCTCCT | [] |
| 441 | 1 | BGLAP | {'exs': 0, 'nonexs': 7} | True | 1.56E+08 | GACACCCTAGACCGGGCCGTA | [] |
| 442 | 1 | RHBG | {'exs': 1, 'nonexs': 1} | False | 1.56E+08 | CTTGACTAGTCTTGCTTTCCTG | [] |
| 443 | 1 | C1orf61 | {'exs': 0, 'nonexs': 7} | True | 1.56E+08 | TGGCTCTATCGTCCACACGGA | [] |
| 444 | 1 | HAPLN2 | {'exs': 1, 'nonexs': 7} | True | 1.57E+08 | GGGCGCCTAATTCTCGGCGTA | [] |
| 445 | 1 | BCAN | {'exs': 0, 'nonexs': 7} | True | 1.57E+08 | TGCCCCCTAGGGACCTGGCAT | [] |
| 446 | 1 | NES | {'exs': 0, 'nonexs': 7} | True | 1.57E+08 | CTTTTCCTAGTCCTCCCCTGA | [] |
| 447 | 1 | ISG20L2 | {'exs': 8, 'nonexs': 1} | False | 1.57E+08 | ACTGCCACTAGTCTGTAGGGG | [] |
| 448 | 1 | HDGF | {'exs': 0, 'nonexs': 6} | True | 1.57E+08 | TGGTGGCTACAGGCTGTGAGG | [] |
| 449 | 1 | PRCC | {'exs': 4, 'nonexs': 4} | False | 1.57E+08 | AATGAGCTACGATGCGAGCAG | [] |
| 450 | 1 | PRCC | {'exs': 4, 'nonexs': 4} | False | 1.57E+08 | AGAGCCCTAGAATCCATATTTG | [] |
| 451 | 1 | SH2D2A | {'exs': 0, 'nonexs': 7} | True | 1.57E+08 | CCGCCTACTGAGGAGGCCCAA | [] |
| 452 | 1 | NTRK1 | {'exs': 0, 'nonexs': 7} | True | 1.57E+08 | CCCCCTAGCCCAGGACATCCA | [] |
| 453 | 1 | LRRC71 | unavailable | False | 1.57E+08 | GGAGGGGGCTAGGGGAAGAAT | [] |
| 454 | 1 | ETV3 | {'exs': 1, 'nonexs': 7} | True | 1.57E+08 | CAGTTCTAAGCATCAGCAGCTG | [] |
| 455 | 1 | FCRL3 | {'exs': 1, 'nonexs': 7} | True | 1.58E+08 | CACAGGGCTAAGTGTTACAAAG | [] |
| 456 | 1 | FCRL3 | {'exs': 1, 'nonexs': 7} | True | 1.58E+08 | GGGGCTAGTGGTCTGAGGCCA | [] |
| 457 | 1 | FCRL1 | {'exs': 1, 'nonexs': 7} | True | 1.58E+08 | GATGTCTAAGGAAACCTGGAAG | [] |
| 458 | 1 | FCRL1 | {'exs': 1, 'nonexs': 7} | True | 1.58E+08 | AGGCACTAGAGGGAGAGACCT | [] |
| 459 | 1 | CD5L | {'exs': 0, 'nonexs': 7} | True | 1.58E+08 | TGCCCACCTACACTGGGAAAT | [] |
| 460 | 1 | CD5L | {'exs': 0, 'nonexs': 7} | True | 1.58E+08 | CAGGATACTATCCTATAAAGAG | [] |
| 461 | 1 | KIRREL1 | unavailable | False | 1.58E+08 | CTGGCCCCTACACGTGAGTCT | [] |
| 462 | 1 | OR6K2 | {'exs': 0, 'nonexs': 3} | True | 1.59E+08 | CTCAACTAAAATATCTTACTTG | [] |
| 463 | 1 | OR6K6 | {'exs': 0, 'nonexs': 3} | True | 1.59E+08 | ATCTATTTCCCAGCCCAACCAG | [] |
| 464 | 1 | AIM2 | {'exs': 0, 'nonexs': 7} | True | 1.59E+08 | ACTTCTCTATGTTTTTTTTTTG | [] |
| 465 | 1 | CADM3 | {'exs': 0, 'nonexs': 7} | True | 1.59E+08 | CTAGATGAAATATTCCTTCTTG | [] |
| 466 | 1 | ACKR1 | unavailable | False | 1.59E+08 | AGAACTAGGATTTGCTTCCAAG | [] |
| 467 | 1 | OR10J4 | unavailable | False | 1.59E+08 | CACCTAAGAGAGAGGCTTTCTG | [] |
| 468 | 1 | FCRL6 | {'exs': 0, 'nonexs': 7} | True | 1.6E+08 | CTAGCAGAGAACCTCCTCACA | [] |
| 469 | 1 | PIGM | {'exs': 1, 'nonexs': 7} | True | 1.6E+08 | GAACATACACTAGTCATATTTG | [] |
| 470 | 1 | CASQ1 | {'exs': 1, 'nonexs': 7} | True | 1.6E+08 | CAACTAGTCATCATCATCATCG | [] |
| 471 | 1 | CD84 | {'exs': 0, 'nonexs': 7} | True | 1.61E+08 | AGCCTAGATCACAATTTCATAG | [] |
| 472 | 1 | SLAMF7 | {'exs': 0, 'nonexs': 7} | True | 1.61E+08 | TGTCTAGATAACATTCTCATAG | [] |
| 473 | 1 | LY9 | {'exs': 0, 'nonexs': 7} | True | 1.61E+08 | GAGGGCCTAGCCCTCCCGCAC | [] |
| 474 | 1 | CD244 | {'exs': 0, 'nonexs': 7} | True | 1.61E+08 | CAACTAGGAATAAACATCAAAG | [] |
| 475 | 1 | TSTD1 | {'exs': 0, 'nonexs': 7} | True | 1.61E+08 | CCTGCCTAACTCTCTTTCTCC | ['TTCTCCAACCATTCTCTATAGG'] |
| 476 | 1 | KLHDC9 | {'exs': 0, 'nonexs': 7} | True | 1.61E+08 | GTGTCTCTAGCTCTGGTCAGA | [] |
| 477 | 1 | PFDN2 | {'exs': 7, 'nonexs': 1} | False | 1.61E+08 | CCCTAGGAGACCAACACTCCA | [] |
| 478 | 1 | PPOX | {'exs': 0, 'nonexs': 7} | True | 1.61E+08 | CCCCCGCTAGAAGAGGGGAAA | [] |
| 479 | 1 | PPOX | {'exs': 0, 'nonexs': 7} | True | 1.61E+08 | GGGTGCCTAGTCAGCCCGGGT | [] |
| 480 | 1 | NDUFS2 | {'exs': 3, 'nonexs': 5} | False | 1.61E+08 | CTCTACTACACAATAGGCCTCG | [] |
| 481 | 1 | FCER1G | {'exs': 0, 'nonexs': 7} | True | 1.61E+08 | TATTCTAAAGCTACTGTGGTGG | [] |
| 482 | 1 | SDHC | {'exs': 5, 'nonexs': 2} | False | 1.61E+08 | AAGGATCTAAACAAATAAGGAG | [] |
| 483 | 1 | FCGR2B | {'exs': 1, 'nonexs': 6} | True | 1.62E+08 | GTTAGACTAAGAGCTGACCAAG | [] |
| 484 | 1 | FCGR2B | {'exs': 1, 'nonexs': 6} | True | 1.62E+08 | TGGAGACTAAATACGGTTCTGG | [] |
| 485 | 1 | FCRLA | {'exs': 0, 'nonexs': 7} | True | 1.62E+08 | GTTTACTTCTATTCAGCAGTAG | [] |
| 486 | 1 | ATF6 | {'exs': 0, 'nonexs': 7} | True | 1.62E+08 | GGGTGCTATTGTAATGACTCAG | [] |
| 487 | 1 | NOS1AP | {'exs': 0, 'nonexs': 7} | True | 1.62E+08 | CGGCACCTACACGGCGATCTC | ['GATCTCATCATCCAGGCCGTCG'] |
| 488 | 1 | SPATA46 | unavailable | False | 1.62E+08 | ATCTCTCTAGAGACATAAGTAG | [] |
| 489 | 1 | C1orf226 | {'exs': 0, 'nonexs': 7} | True | 1.62E+08 | AGAGGCTCTATTCAAAGGACAG | [] |
| 490 | 1 | DDR2 | {'exs': 2, 'nonexs': 6} | False | 1.63E+08 | GTGTCAACACCTAATTCACATG | [] |
| 491 | 1 | RGS5 | {'exs': 1, 'nonexs': 7} | True | 1.63E+08 | ATTACTACTTGATTAACTCCTG | [] |
| 492 | 1 | LRRC52 | {'exs': 0, 'nonexs': 7} | True | 1.66E+08 | TGGCAACTAAATAAGCTGAGGG | [] |
| 493 | 1 | FAM78B | {'exs': 0, 'nonexs': 7} | True | 1.66E+08 | AGTCTGCTTCTACTTAGGAGGG | [] |
| 494 | 1 | LOC10537 | unavailable | False | 1.67E+08 | GGACAGTCCTAGGAAAAGTGAG | [] |
| 495 | 1 | MPZL1 | {'exs': 1, 'nonexs': 7} | True | 1.68E+08 | GTGGTCTAACTGTGCATATATG | [] |
| 496 | 1 | ADCY10 | {'exs': 0, 'nonexs': 7} | True | 1.68E+08 | ATAGTCTAGGTAAGCCTTAATG | [] |
| 497 | 1 | ADCY10 | {'exs': 0, 'nonexs': 7} | True | 1.68E+08 | TGTAGCTATAGATGCGCCACAG | [] |
| 498 | 1 | TIPRL | {'exs': 4, 'nonexs': 4} | False | 1.68E+08 | TAGCTGGGACTACAGGTGCCC | [] |
| 499 | 1 | XCL2 | {'exs': 0, 'nonexs': 7} | True | 1.69E+08 | AGAGACTACTAGCCAGTCAGG | [] |
| 500 | 1 | XCL1 | {'exs': 0, 'nonexs': 7} | True | 1.69E+08 | AGAGACTACTAGCCAGTCAGAG | [] |
| 501 | 1 | DPT | {'exs': 0, 'nonexs': 7} | True | 1.69E+08 | ATGTGGCAAATCTAAACATTTG | [] |
| 502 | 1 | NME7 | {'exs': 1, 'nonexs': 6} | True | 1.69E+08 | CCACTAATTATCCAAGATCTTG | [] |
| 503 | 1 | F5 | {'exs': 0, 'nonexs': 7} | True | 1.7E+08 | TCAATTCTAGTAAATATCACAG | [] |
| 504 | 1 | C1orf112 | {'exs': 0, 'nonexs': 7} | True | 1.7E+08 | ATCTATTTGATTTGAAATTCAG | [] |
| 505 | 1 | MROH9 | unavailable | False | 1.71E+08 | CTATAAGGCCTTATCATTCTTG | [] |
| 506 | 1 | FMO2 | {'exs': 0, 'nonexs': 7} | True | 1.71E+08 | ATGCTGACTAGGACCATTGAAG | [] |
| 507 | 1 | FMO1 | {'exs': 0, 'nonexs': 7} | True | 1.71E+08 | TCTATGTATGTGATATAATCTG | [] |
| 508 | 1 | PRRC2C | unavailable | False | 1.72E+08 | CTCCTCCTATCTGTTTGCTCTG | [] |
| 509 | 1 | DNM3 | {'exs': 1, 'nonexs': 7} | True | 1.72E+08 | CTGTTCCTATTTGGCCATCTTG | [] |
| 510 | 1 | DNM3 | {'exs': 1, 'nonexs': 7} | True | 1.72E+08 | AGACTAAATGTTCTGCAATAAG | [] |
| 511 | 1 | DNM3 | {'exs': 1, 'nonexs': 7} | True | 1.72E+08 | TCCTCTCTTCTAAATCTTGCGG | [] |
| 512 | 1 | TNFSF18 | {'exs': 0, 'nonexs': 7} | True | 1.73E+08 | AAGTCTCTAGGAGATGAATTGG | [] |
| 513 | 1 | ZBTB37 | {'exs': 0, 'nonexs': 5} | True | 1.74E+08 | AACCTACCTGTCAACTTCACTG | [] |
| 514 | 1 | ZBTB37 | {'exs': 0, 'nonexs': 5} | True | 1.74E+08 | ATTTTCTATTCATACACAGTAG | [] |
| 515 | 1 | RABGAP1I | {'exs': 0, 'nonexs': 7} | True | 1.75E+08 | TTACCTACAAATATACAAACTG | [] |
| 516 | 1 | RABGAP1I | {'exs': 0, 'nonexs': 7} | True | 1.75E+08 | CTGGAACTATGTGCTCTCCTTG | [] |
| 517 | 1 | KIAA0040 | {'exs': 0, 'nonexs': 2} | False | 1.75E+08 | CCTAACTAAACAGGCAGTGATG | [] |
| 518 | 1 | COP1 | unavailable | False | 1.76E+08 | GTACCTAATCACTCCTGTAAAG | [] |
| 519 | 1 | ASTN1 | {'exs': 0, 'nonexs': 7} | True | 1.77E+08 | ATGGTGCTAGATCTCTTTGCTG | [] |
| 520 | 1 | TEX35 | unavailable | False | 1.79E+08 | AATTGAGCTACCATAGATCCAG | [] |
| 521 | 1 | RALGPS2 | {'exs': 0, 'nonexs': 7} | True | 1.79E+08 | CTTCTACTCAAAAGTCATCAAG | [] |
| 522 | 1 | ABL2 | {'exs': 1, 'nonexs': 6} | True | 1.79E+08 | CTACCTCTGCACCACATCACT | [] |
| 523 | 1 | SOAT1 | {'exs': 0, 'nonexs': 7} | True | 1.79E+08 | CAAGCTTCTAAAACACGTAACG | [] |
| 524 | 1 | AXDND1 | unavailable | False | 1.8E+08 | TGCTATCTCTGGTGACATAAAG | [] |
| 525 | 1 | NPHS2 | {'exs': 1, 'nonexs': 7} | True | 1.8E+08 | TCCTTCCTATAACATGGGAGAG | [] |
| 526 | 1 | TOR1AIP2 | {'exs': 0, 'nonexs': 1} | False | 1.8E+08 | GAAGCTATTTGTTTTGGCCCAG | [] |
| 527 | 1 | CACNA1E | {'exs': 0, 'nonexs': 5} | True | 1.82E+08 | GAGCAGCCTCTAGCATTTGTC | ['GCATTTGTCATCTTCTTCCGTG'] |
| 528 | 1 | RGS8 | {'exs': 0, 'nonexs': 7} | True | 1.83E+08 | AGGTCTAACTGAGCCTCCTCT | [] |
| 529 | 1 | NPL | {'exs': 2, 'nonexs': 5} | False | 1.83E+08 | GCACTAGCTACCAGCTTCCAA | [] |
| 530 | 1 | LAMC1 | {'exs': 0, 'nonexs': 7} | True | 1.83E+08 | AAGACACTAGGGCTTTTCAATG | [] |
| 531 | 1 | NMNAT2 | {'exs': 0, 'nonexs': 7} | True | 1.83E+08 | GGGCTGCTAGCCGGAGGCATT | [] |
| 532 | 1 | NCF2 | {'exs': 0, 'nonexs': 7} | True | 1.84E+08 | ACATCCTAGACTTCTCTCCGAG | [] |
| 533 | 1 | ARPC5 | {'exs': 0, 'nonexs': 7} | True | 1.84E+08 | CCAGACTACACAGTTTTTCTTG | [] |
| 534 | 1 | TSEN15 | {'exs': 2, 'nonexs': 5} | False | 1.84E+08 | ATCTTCTAAGAGAAATATTCTG | [] |
| 535 | 1 | TSEN15 | {'exs': 2, 'nonexs': 5} | False | 1.84E+08 | TGGACACTAGAAATCCAGACCG | [] |
| 536 | 1 | C1orf21 | {'exs': 0, 'nonexs': 7} | True | 1.85E+08 | -1 | -1 |
| 537 | 1 | TRMT1L | unavailable | False | 1.85E+08 | TTATTTCTAGATGTTTCGATAG | [] |
| 538 | 1 | HMCN1 | {'exs': 0, 'nonexs': 7} | True | 1.86E+08 | GTGGAACTACCTTTGAGAACTG | [] |
| 539 | 1 | PRG4 | {'exs': 0, 'nonexs': 7} | True | 1.86E+08 | AGTCTAAGGACAGTTGTACCAG | [] |
| 540 | 1 | ODR4 | unavailable | False | 1.86E+08 | CATCTCTATACTAGACTGTCAG | [] |
| 541 | 1 | ODR4 | unavailable | False | 1.86E+08 | CACCCTAATCACTGAAGTAATG | [] |
| 542 | 1 | PTGS2 | {'exs': 0, 'nonexs': 7} | True | 1.87E+08 | ACTTCTACAGTTCAGTCGAACG | [] |
| 543 | 1 | PLA2G4A | {'exs': 0, 'nonexs': 7} | True | 1.87E+08 | AGTACATGAACTATGCTTTGGG | [] |
| 544 | 1 | UCHL5 | {'exs': 4, 'nonexs': 5} | False | 1.93E+08 | TTCTTTTATCCTATTTTCCCTG | [] |
| 545 | 1 | RO60 | unavailable | False | 1.93E+08 | CAAAACTATTTTACAGTGTCTG | [] |
| 546 | 1 | RO60 | unavailable | False | 1.93E+08 | ATGTCTAAAATGATTTATTTAG | [] |
| 547 | 1 | B3GALT2 | {'exs': 1, 'nonexs': 7} | True | 1.93E+08 | TCTTTTCTAATGTAGTTTACGG | [] |
| 548 | 1 | KCNT2 | {'exs': 1, 'nonexs': 7} | True | 1.96E+08 | CTACTAATGGCAAGAAATCACG | [] |
| 549 | 1 | KCNT2 | {'exs': 1, 'nonexs': 7} | True | 1.96E+08 | TCCTAACAAATACTTATATAAG | [] |
| 550 | 1 | CFH | {'exs': 0, 'nonexs': 7} | True | 1.97E+08 | TGATTCTATCTTTTTGCACAAG | [] |
| 551 | 1 | CFHR1 | {'exs': 0, 'nonexs': 7} | True | 1.97E+08 | TGATTCTATCTTTTTGCACAAG | [] |
| 552 | 1 | CFHR4 | {'exs': 1, 'nonexs': 7} | True | 1.97E+08 | CCATTGCTAATAGGTGGAGGAG | [] |
| 553 | 1 | CFHR2 | {'exs': 0, 'nonexs': 7} | True | 1.97E+08 | ATTCTATTTTTCTTCACAACTG | [] |
| 554 | 1 | F13B | {'exs': 0, 'nonexs': 7} | True | 1.97E+08 | TTCATTTCTATGTTCTTAAGGG | [] |
| 555 | 1 | ASPM | {'exs': 1, 'nonexs': 6} | True | 1.97E+08 | TTTACTAATAAGGAATGCCAAG | [] |
| 556 | 1 | CRB1 | {'exs': 0, 'nonexs': 7} | True | 1.97E+08 | TCCTAAATCAGTCTCTCCATTG | [] |
| 557 | 1 | DENND1B | {'exs': 0, 'nonexs': 6} | True | 1.98E+08 | TCGAGCTATGCGGTTTCAGCC | [] |
| 558 | 1 | PTPRC | {'exs': 2, 'nonexs': 5} | False | 1.99E+08 | TCCTATGAACCTTGATTTAAAG | [] |
| 559 | 1 | CAMSAP2 | unavailable | False | 2.01E+08 | CCAACTTCTATGCCTTAGTGGG | [] |
| 560 | 1 | GPR25 | {'exs': 0, 'nonexs': 7} | True | 2.01E+08 | GGGCAGCTACCAGGAGGCCGA | [] |
| 561 | 1 | KIF21B | {'exs': 0, 'nonexs': 6} | True | 2.01E+08 | TCACTCCTAGGGTGGGCCGCT | [] |
| 562 | 1 | TMEM9 | {'exs': 0, 'nonexs': 7} | True | 2.01E+08 | GCCCATCTAGCTGAGCATCTT | [] |
| 563 | 1 | IGFN1 | {'exs': 0, 'nonexs': 5} | True | 2.01E+08 | TGAGGCTAGGTGCTGGGTTCT | [] |
| 564 | 1 | TNNT2 | {'exs': 0, 'nonexs': 5} | True | 2.01E+08 | CAGGCTCTATTTCCAGCGCCC | [] |
| 565 | 1 | TNNI1 | {'exs': 0, 'nonexs': 7} | True | 2.01E+08 | TACCTCTATTGTGAGGTCGGAG | [] |
| 566 | 1 | NAV1 | {'exs': 0, 'nonexs': 7} | True | 2.02E+08 | AGCTCCCTTGTCTATTCTTCTG | [] |
| 567 | 1 | NAV1 | {'exs': 0, 'nonexs': 7} | True | 2.02E+08 | -1 | -1 |
| 568 | 1 | LMOD1 | {'exs': 0, 'nonexs': 7} | True | 2.02E+08 | CTGGTCCTACTGAAGCAGTTTG | [] |
| 569 | 1 | TIMM17A | {'exs': 1, 'nonexs': 7} | True | 2.02E+08 | AGAAGTCCTACTGATATTGTCG | [] |
| 570 | 1 | RNPEP | {'exs': 1, 'nonexs': 7} | True | 2.02E+08 | AGCCTCTAACTGCCCTTGGGT | [] |
| 571 | 1 | PTPN7 | {'exs': 1, 'nonexs': 6} | True | 2.02E+08 | CACATTTCTAGGAGGGAGGGA | [] |
| 572 | 1 | UBE2T | {'exs': 3, 'nonexs': 5} | False | 2.02E+08 | AAGTCCCCTAAACATCAGGATG | [] |
| 573 | 1 | PPP1R12B | {'exs': 1, 'nonexs': 6} | True | 2.02E+08 | GACTAGAAAATATTTTCATTTG | [] |
| 574 | 1 | PPP1R12B | {'exs': 1, 'nonexs': 6} | True | 2.03E+08 | CAAGAACTAGAGCTCCTGGGT | [] |
| 575 | 1 | PPP1R12B | {'exs': 1, 'nonexs': 6} | True | 2.03E+08 | CCTAGCCTACTTGGACAGTTTG | [] |
| 576 | 1 | SYT2 | {'exs': 0, 'nonexs': 7} | True | 2.03E+08 | CTGTCTACTTGTTCTTGCCCAG | [] |
| 577 | 1 | TMEM183 | {'exs': 1, 'nonexs': 6} | True | 2.03E+08 | CAGTAACTACGCTCTCAGGGA | [] |
| 578 | 1 | PPFIA4 | {'exs': 0, 'nonexs': 7} | True | 2.03E+08 | CCATGGCTAGTCCCGGAAGGC | [] |
| 579 | 1 | PPFIA4 | {'exs': 0, 'nonexs': 7} | True | 2.03E+08 | TAGACACCCTAAGATTGAATAG | [] |
| 580 | 1 | ADORA1 | {'exs': 0, 'nonexs': 7} | True | 2.03E+08 | AAGGCGGGGTCTAGTCATCAG | [] |
| 581 | 1 | CHI3L1 | {'exs': 0, 'nonexs': 7} | True | 2.03E+08 | GAGGGCTACGTTGCAGCGAGT | [] |
| 582 | 1 | BTG2 | {'exs': 1, 'nonexs': 6} | True | 2.03E+08 | GGAAGGGCCTAGCTGGAGACT | [] |
| 583 | 1 | PRELP | {'exs': 0, 'nonexs': 7} | True | 2.03E+08 | GGGCCTAGATGACCACGGACT | [] |
| 584 | 1 | OPTC | {'exs': 1, 'nonexs': 7} | True | 2.04E+08 | TCCGAGCTACGTGAAGCGGCC | [] |
| 585 | 1 | ATP2B4 | {'exs': 0, 'nonexs': 7} | True | 2.04E+08 | GAGGGAACTATGGAACGCTTTG | [] |
| 586 | 1 | SNRPE | {'exs': 7, 'nonexs': 1} | False | 2.04E+08 | ATTTCTAGTTGGAGACACTTTG | [] |
| 587 | 1 | KLHDC8A | {'exs': 1, 'nonexs': 7} | True | 2.05E+08 | CAGAGACAGCTAGGAGTCAGA | [] |
| 588 | 1 | LEMD1 | {'exs': 0, 'nonexs': 7} | True | 2.05E+08 | GTATCTACAGCTTTGCGTTTAG | [] |
| 589 | 1 | SLC45A3 | {'exs': 0, 'nonexs': 7} | True | 2.06E+08 | AGTTTTCTACGCTGAGTATTTG | [] |
| 590 | 1 | RAB29 | unavailable | False | 2.06E+08 | ACACTACTAGCAGCAGGACCA | [] |
| 591 | 1 | SLC41A1 | {'exs': 0, 'nonexs': 7} | True | 2.06E+08 | CCAAGCTAGTCCCCGACATCC | [] |
| 592 | 1 | SLC26A9 | {'exs': 0, 'nonexs': 7} | True | 2.06E+08 | TTTATCCCTATGTCCGTGACAG | [] |
| 593 | 1 | RAB7B | unavailable | False | 2.06E+08 | TTGACCAAGCACTAAGAGAGGG | [] |
| 594 | 1 | AVPR1B | {'exs': 0, 'nonexs': 7} | True | 2.06E+08 | AGTCTTTCCTAAAAGATGATGG | [] |
| 595 | 1 | SRGAP2 | {'exs': 0, 'nonexs': 1} | False | 2.06E+08 | GATGTTGCTAACAGCCACAATG | [] |
| 596 | 1 | SRGAP2 | {'exs': 0, 'nonexs': 1} | False | 2.06E+08 | TGGTCTAATGAAAATTATATAG | [] |
| 597 | 1 | DYRK3 | {'exs': 1, 'nonexs': 6} | True | 2.07E+08 | GTCCACTAGCTAATCAGTTTTG | [] |
| 598 | 1 | IL19 | {'exs': 1, 'nonexs': 6} | True | 2.07E+08 | CCTCTTACTAAGGAATATGTTG | [] |
| 599 | 1 | IL20 | {'exs': 1, 'nonexs': 6} | True | 2.07E+08 | TCCTCCTATTCTGTCTCCTCC | ['CTCCATCCATTGCAGAAGAATG'] |
| 600 | 1 | PIGR | {'exs': 0, 'nonexs': 7} | True | 2.07E+08 | CACCGTCTAGGCTTCCTGGGG | [] |
| 601 | 1 | C1orf116 | {'exs': 0, 'nonexs': 7} | True | 2.07E+08 | GCAGAGTCTACTCCTTCAACAG | [] |
| 602 | 1 | PFKFB2 | {'exs': 1, 'nonexs': 6} | True | 2.07E+08 | TCGGCTAGTCGGCCCCTTCTT | [] |
| 603 | 1 | CD55 | {'exs': 0, 'nonexs': 7} | True | 2.07E+08 | ATGTCTGACTAGATGCCTCTGG | [] |
| 604 | 1 | CD55 | {'exs': 0, 'nonexs': 7} | True | 2.07E+08 | GTGACCTAGAGGATGCACATC | ['CATCATCTTTTTTGAAGAAATG'] |
| 605 | 1 | CD55 | {'exs': 0, 'nonexs': 7} | True | 2.07E+08 | AAGTCTAAGAAACTAGGAACAG | [] |
| 606 | 1 | CD46 | {'exs': 0, 'nonexs': 7} | True | 2.08E+08 | TGGAATCTATTCAGCCTCTCTG | [] |
| 607 | 1 | G0S2 | {'exs': 0, 'nonexs': 7} | True | 2.1E+08 | ACAGTTCCTAGGAGGCGTGCT | [] |
| 608 | 1 | HSD11B1 | {'exs': 1, 'nonexs': 7} | True | 2.1E+08 | TTCCTACTTGTTTATGAATCTG | [] |
| 609 | 1 | TRAF3IP3 | {'exs': 0, 'nonexs': 7} | True | 2.1E+08 | CCTGCTATCTGCTCCCTGTAA | [] |
| 610 | 1 | TRAF5 | {'exs': 0, 'nonexs': 7} | True | 2.11E+08 | CAGTGACTAGAGATCCTCCAG | [] |
| 611 | 1 | NEK2 | {'exs': 2, 'nonexs': 6} | False | 2.12E+08 | GCCAAGGCTACATTGAGCTGT | [] |
| 612 | 1 | NEK2 | {'exs': 2, 'nonexs': 6} | False | 2.12E+08 | ACCTGGCTAGCGCATGCCCAG | [] |
| 613 | 1 | LPGAT1 | {'exs': 0, 'nonexs': 7} | True | 2.12E+08 | CAATTCCTAAAACAGGCAATGG | [] |
| 614 | 1 | DTL | {'exs': 7, 'nonexs': 1} | False | 2.12E+08 | TAGAATCTATAATTCTGTTGAG | [] |
| 615 | 1 | PACC1 | unavailable | False | 2.12E+08 | TGACTATCTGTGAAGCAAACAG | [] |
| 616 | 1 | NSL1 | {'exs': 7, 'nonexs': 1} | False | 2.13E+08 | CTCTCCTAAACTGCCCCTAGA | [] |
| 617 | 1 | TATDN3 | {'exs': 0, 'nonexs': 7} | True | 2.13E+08 | TTGAAGCTATTTCTGGAGCAAG | [] |
| 618 | 1 | VASH2 | {'exs': 0, 'nonexs': 7} | True | 2.13E+08 | GGCTTGGCTAAATTCGGATTTG | [] |
| 619 | 1 | PROX1 | {'exs': 0, 'nonexs': 5} | True | 2.14E+08 | GAAATTTCTACTCATGAAGCAG | [] |
| 620 | 1 | KCNK2 | {'exs': 0, 'nonexs': 7} | True | 2.15E+08 | GAGGGCTATTTGATGTTCTCAG | ['ATCACAGCAATCTCTTCACCAG'] |
| 621 | 1 | GPATCH2 | {'exs': 1, 'nonexs': 7} | True | 2.17E+08 | TTCACCTACAAATCCTGAAATG | [] |
| 622 | 1 | GPATCH2 | {'exs': 1, 'nonexs': 7} | True | 2.18E+08 | AGACTATGTAATTGTTTTTCTG | [] |
| 623 | 1 | RRP15 | {'exs': 4, 'nonexs': 4} | False | 2.18E+08 | CCAGACCTAGGAGCTCCCTGA | [] |
| 624 | 1 | IARS2 | {'exs': 6, 'nonexs': 1} | False | 2.2E+08 | CTGTTAATACTATTTTCCACTG | [] |
| 625 | 1 | MTARC2 | unavailable | False | 2.21E+08 | CACTACACCATCCGATACACAG | [] |
| 626 | 1 | HLX | {'exs': 0, 'nonexs': 7} | True | 2.21E+08 | ACAGTCTATAAGCAGCCAAGCG | [] |
| 627 | 1 | CAPN8 | {'exs': 0, 'nonexs': 1} | False | 2.24E+08 | AACCTAGGAGGAGCCTACACA | [] |
| 628 | 1 | ENAH | {'exs': 0, 'nonexs': 7} | True | 2.25E+08 | CTGTTCCTCTATGCAGTATTTG | [] |
| 629 | 1 | LEFTY1 | {'exs': 0, 'nonexs': 7} | True | 2.26E+08 | GCCTATGGCTGGAGCCTCCTT | [] |
| 630 | 1 | LEFTY2 | {'exs': 0, 'nonexs': 7} | True | 2.26E+08 | GCCTATGGCTGGAGCCTCCTT | [] |
| 631 | 1 | LEFTY2 | {'exs': 0, 'nonexs': 7} | True | 2.26E+08 | TCTCACTATGTTCTAAAAGCTG | [] |
| 632 | 1 | COQ8A | unavailable | False | 2.27E+08 | CGCAGCCCTACTGCTGGGCCT | [] |
| 633 | 1 | JMJD4 | {'exs': 0, 'nonexs': 7} | True | 2.28E+08 | AGGTGCTATGGGGCCGCAGCA | [] |
| 634 | 1 | SNAP47 | {'exs': 0, 'nonexs': 7} | True | 2.28E+08 | TGCCCTGCTAGGAAAGTCTCT | [] |
| 635 | 1 | SNAP47 | {'exs': 0, 'nonexs': 7} | True | 2.28E+08 | CCTAGGTCAGCCTCTTCATCC | [] |
| 636 | 1 | WNT3A | {'exs': 0, 'nonexs': 7} | True | 2.28E+08 | CCGGTGCCTACTTGCAGGTGT | [] |
| 637 | 1 | WNT3A | {'exs': 0, 'nonexs': 7} | True | 2.28E+08 | AGCCCTAGGAAGTGTCCCCAT | [] |
| 638 | 1 | IBA57 | unavailable | False | 2.28E+08 | TCGGACTACTTGGAGACTGTAG | [] |
| 639 | 1 | TRIM17 | {'exs': 0, 'nonexs': 7} | True | 2.28E+08 | TGTCTATCCTTTCACCCACATG | [] |
| 640 | 1 | TRIM17 | {'exs': 0, 'nonexs': 7} | True | 2.28E+08 | GCCGCCCTACGAAGGCACAGT | [] |
| 641 | 1 | H3-4 | unavailable | False | 2.28E+08 | GCCCTCCTAGGCCCGCTCCCC | [] |
| 642 | 1 | RAB4A | {'exs': 0, 'nonexs': 7} | True | 2.29E+08 | TCTCCTAACAACCACACTCCTG | [] |
| 643 | 1 | ACTA1 | {'exs': 0, 'nonexs': 7} | True | 2.29E+08 | GTGTGTCTAGAAGCATTTGCGG | [] |
| 644 | 1 | GALNT2 | {'exs': 0, 'nonexs': 7} | True | 2.3E+08 | TATCTTCTATCTTAAGCTATTG | ['GCTATTATCTAACAGGCAGGGG'] |
| 645 | 1 | GALNT2 | {'exs': 0, 'nonexs': 7} | True | 2.3E+08 | ACCCTCCTACTGCTGCAGGTT | [] |
| 646 | 1 | TTC13 | {'exs': 0, 'nonexs': 7} | True | 2.31E+08 | AGCAGAACTAGAGTTTCTTAAG | [] |
| 647 | 1 | FAM89A | {'exs': 0, 'nonexs': 7} | True | 2.31E+08 | AGACCCTCTAGATGGACTCCA | [] |
| 648 | 1 | TRIM67 | {'exs': 0, 'nonexs': 7} | True | 2.31E+08 | AGCGGGGCTAATTGCCTGACA | [] |
| 649 | 1 | EGLN1 | {'exs': 1, 'nonexs': 7} | True | 2.31E+08 | GCTCTAGAAGACGTCTTTACCG | [] |
| 650 | 1 | TSNAX | {'exs': 0, 'nonexs': 7} | True | 2.32E+08 | TCTAAGAAATGCCCTCTTCTTG | [] |
| 651 | 1 | DISC1 | {'exs': 0, 'nonexs': 7} | True | 2.32E+08 | TCAGAACTAGGTATGCAAAGTG | [] |
| 652 | 1 | DISC1 | {'exs': 0, 'nonexs': 7} | True | 2.32E+08 | TCTTCTAATCTTTGTTGTAACG | [] |
| 653 | 1 | DISC1 | {'exs': 0, 'nonexs': 7} | True | 2.32E+08 | GTGCACTACCTACCATAGGCA | [] |
| 654 | 1 | DISC1 | {'exs': 0, 'nonexs': 7} | True | 2.32E+08 | TTCTACAGACAGGCTTCCCCT | [] |
| 655 | 1 | SIPA1L2 | {'exs': 0, 'nonexs': 7} | True | 2.32E+08 | GATTGGCTAAGATTTTTTGTCG | [] |
| 656 | 1 | COA6 | unavailable | False | 2.34E+08 | ACAGCCTAGGATTTTGCAGTTG | [] |
| 657 | 1 | TARBP1 | {'exs': 0, 'nonexs': 7} | True | 2.34E+08 | CTGGACTCTATGAGGGCCCAG | [] |
| 658 | 1 | LYST | {'exs': 1, 'nonexs': 5} | False | 2.36E+08 | TGGCCACTAAAAAAAATCTGGG | [] |
| 659 | 1 | LYST | {'exs': 1, 'nonexs': 5} | False | 2.36E+08 | ACTCTAAGCTGTAGTGCAACAG | [] |
| 660 | 1 | GPR137B | {'exs': 0, 'nonexs': 7} | True | 2.36E+08 | ACTGATGCTACCCAAGGCTTG | [] |
| 661 | 1 | ERO1B | unavailable | False | 2.36E+08 | -1 | -1 |
| 662 | 1 | EDARADD | {'exs': 0, 'nonexs': 7} | True | 2.36E+08 | AGAGCTCTAGAAGTGCCTGGA | [] |
| 663 | 1 | LGALS8 | {'exs': 0, 'nonexs': 7} | True | 2.37E+08 | GCTACCAGCTCCTTACTTCCA | [] |
| 664 | 1 | CHRM3 | {'exs': 0, 'nonexs': 7} | True | 2.4E+08 | CTCATTCTACAAGGCCTGCTC | [] |
| 665 | 1 | OPN3 | {'exs': 0, 'nonexs': 7} | True | 2.42E+08 | ATTCCTACAAAGGACGAACTTG | [] |
| 666 | 1 | CHML | {'exs': 0, 'nonexs': 7} | True | 2.42E+08 | CTCTTTTCTAATTTTGAAGGTG | [] |
| 667 | 1 | BECN2 | unavailable | False | 2.42E+08 | GAAAGCTACTTTTGATACCTTG | [] |
| 668 | 1 | MAP1LC3C | {'exs': 0, 'nonexs': 7} | True | 2.42E+08 | CATGGGCTAGAGAGGATTGCA | [] |
| 669 | 1 | SDCCAG8 | {'exs': 1, 'nonexs': 7} | True | 2.43E+08 | ACGTTTGCTATTGTGAAGGGTG | [] |
| 670 | 1 | AKT3 | {'exs': 2, 'nonexs': 5} | False | 2.44E+08 | TAGTGAAGTCTAAAAGGCATTG | [] |

| | | | | | | | |
|---|---|---|---|---|---|---|---|
| 671 | 1 | AKT3 | {'ess': 2, 'nonest': 5} | False | 2.44E+08 | ATCAAACCTAAATCTTATTTTG | [] |
| 672 | 1 | CATSPERI | unavailable | False | 2.45E+08 | TTGTCCTAGTTGAAAAAAATCG | ['AAAAATCACCCCAAGCCTCAGG'] |
| 673 | 1 | CATSPERI | unavailable | False | 2.45E+08 | AGTTTTCCTAATTCTTCTTACG | [] |
| 674 | 1 | EFCAB2 | {'ess': 0, 'nonest': 7} | True | 2.45E+08 | CTCTACCTAAACATTCAATAAG | [] |
| 675 | 1 | KIF26B | {'ess': 0, 'nonest': 7} | True | 2.46E+08 | GCTCATCTACCGGCGCCTGGA | [] |
| 676 | 1 | TFB2M | {'ess': 5, 'nonest': 4} | False | 2.47E+08 | TAGTTGCTACCTATCTTCCAGG | [] |
| 677 | 1 | CNST | {'ess': 0, 'nonest': 7} | True | 2.47E+08 | CTGCTAGGAGAGGTAGATCCAC | [] |
| 678 | 1 | AHCTF1 | {'ess': 7, 'nonest': 2} | False | 2.47E+08 | CGAATTTCTATGGGAGGAGGTG | [] |
| 679 | 1 | NLRP3 | {'ess': 0, 'nonest': 7} | True | 2.47E+08 | CTCCTACCAAGAAGGCTCAAAG | [] |
| 680 | 1 | OR2W5 | unavailable | False | 2.47E+08 | CATTACTAGAGTGTTCCCCTCG | [] |
| 681 | 1 | OR2C3 | {'ess': 0, 'nonest': 3} | True | 2.48E+08 | GGAGTCTCTAAATTTGCGCCAG | [] |
| 682 | 1 | OR13G1 | {'ess': 0, 'nonest': 3} | True | 2.48E+08 | AACTACTAGTGTTTCAGAAATG | [] |
| 683 | 1 | OR11L1 | {'ess': 0, 'nonest': 3} | True | 2.48E+08 | TTTTCTTTACCTAATAAAGGAG | ['TAAAGGAACTTCCTTTTACTTG'] |
| 684 | 1 | OR2L8 | {'ess': 0, 'nonest': 3} | True | 2.48E+08 | GTGTTTCTACATTTTCACAGAG | [] |
| 685 | 1 | OR2L5 | {'ess': 0, 'nonest': 1} | False | 2.48E+08 | GTATGTCTACATTTTCACCGAG | [] |
| 686 | 1 | OR2L2 | {'ess': 0, 'nonest': 3} | True | 2.48E+08 | GTATGTCTACATTTTCACTGAG | [] |
| 687 | 1 | OR2L3 | {'ess': 0, 'nonest': 3} | True | 2.48E+08 | ATGTTTCTACATTTTCCCAGAG | [] |
| 688 | 1 | OR2T4 | {'ess': 0, 'nonest': 3} | True | 2.48E+08 | TGGTCTACTCCATAGCTTTTTG | [] |
| 689 | 1 | OR2T6 | {'ess': 1, 'nonest': 3} | False | 2.48E+08 | CCACATGTCCCCTAACATCTTG | [] |
| 690 | 1 | OR2T2 | {'ess': 0, 'nonest': 3} | True | 2.48E+08 | GAGTCCCTGCTAGCCCTTCCT | [] |
| 691 | 1 | OR2T3 | {'ess': 0, 'nonest': 3} | True | 2.48E+08 | TGCCCTTTACTACTTTTCTTGG | [] |
| 692 | 1 | OR2T5 | {'ess': 0, 'nonest': 3} | True | 2.48E+08 | TCCTAAAGGACGAATCTCACAG | [] |
| 693 | 1 | OR2G6 | {'ess': 0, 'nonest': 3} | True | 2.49E+08 | TTTCCTAGTCCTTGTGGCTTTG | [] |
| 694 | 1 | OR2T29 | {'ess': 0, 'nonest': 3} | True | 2.49E+08 | TCCTAAAGGACGAATCTCACAG | [] |
| 695 | 1 | OR2T34 | {'ess': 0, 'nonest': 3} | True | 2.49E+08 | TGTCCTTTACTACTTTTCTTGG | [] |
| 696 | 1 | OR2T11 | {'ess': 1, 'nonest': 3} | False | 2.49E+08 | ACTCTCTAAGCATCACTTGTTG | [] |
| 697 | 1 | OR2T35 | {'ess': 0, 'nonest': 3} | True | 2.49E+08 | GAGTCCCTGCTAGCCCTTCCT | [] |
| 698 | 1 | OR14I1 | {'ess': 0, 'nonest': 3} | True | 2.49E+08 | CCAGGATGTTCTACTTTTGCAG | [] |
| 699 | 1 | SH3BP5L | {'ess': 0, 'nonest': 7} | True | 2.49E+08 | CCTCGGCTACAGGCTGACGCT | [] |
| 700 | 1 | ZNF672 | {'ess': 1, 'nonest': 7} | True | 2.49E+08 | CTCCCTCAACTAGGACACAGA | [] |
| 701 | 2 | FAM110C | {'ess': 0, 'nonest': 7} | True | 42847 | GAAATCCTAAAATGAATCCTTG | ['GAATCCTTATAAATTTCAGCAG'] |
| 702 | 2 | TPO | {'ess': 0, 'nonest': 7} | True | 1540914 | CACTAAGCAGGCTCACATTCTG | [] |
| 703 | 2 | PXDN | {'ess': 0, 'nonest': 7} | True | 1634203 | GGAGCCTAGGGCTTTTCCTCC | [] |
| 704 | 2 | MYT1L | {'ess': 1, 'nonest': 6} | True | 1885430 | GAGCTAGTTCAGGAAGTCACTG | [] |
| 705 | 2 | TRAPPC12 | unavailable | False | 3457029 | CAAGTCCCTATGGCGTGGGGC | [] |
| 706 | 2 | TRAPPC12 | unavailable | False | 3479460 | AGGCAGCTAGGCCAGCTTGAG | [] |
| 707 | 2 | ADI1 | {'ess': 0, 'nonest': 7} | True | 3498962 | ACTGCTAGGCGGTCTGTGCCA | [] |
| 708 | 2 | LOC10537 | unavailable | False | 4024957 | TTTTGACTAAAAACTAAATGGG | [] |
| 709 | 2 | CMPK2 | {'ess': 0, 'nonest': 7} | True | 6849849 | GAGTAACTACGGTTCACTAAAG | ['ACTAAAACTATTCTGGATTAGG'] |
| 710 | 2 | CMPK2 | {'ess': 0, 'nonest': 7} | True | 6851445 | ACTGGGACACCTACTTTTGACG | [] |
| 711 | 2 | RSAD2 | {'ess': 0, 'nonest': 7} | True | 6895941 | CGCTCTACCAATCCAGCTTCAC | [] |
| 712 | 2 | RNF144A | {'ess': 0, 'nonest': 7} | True | 7039759 | CTTCCTCTAGGTGGGTAACGG | [] |
| 713 | 2 | ASAP2 | {'ess': 0, 'nonest': 7} | True | 9390178 | GTATGACCTACATTACTGGAAG | [] |
| 714 | 2 | IAH1 | {'ess': 0, 'nonest': 7} | True | 9488328 | GCTAATGGTCTCCATCTCCCAC | [] |
| 715 | 2 | HPCAL1 | {'ess': 0, 'nonest': 7} | True | 10424612 | CCAACCTAGAAGATGCCCTCT | [] |
| 716 | 2 | ODC1 | {'ess': 2, 'nonest': 6} | False | 10440723 | GCTATCTACACATTAATACTAG | [] |
| 717 | 2 | ATP6V1C2 | {'ess': 0, 'nonest': 7} | True | 10783262 | CTTTCTAGTCAAGAAGACTAAG | [] |
| 718 | 2 | C2orf50 | {'ess': 2, 'nonest': 7} | False | 11144110 | GCTCCTAGATGGGCTGCATCT | [] |
| 719 | 2 | LRATD1 | unavailable | False | 14634857 | CGGCTACTCCTTGTCGTCCAC | [] |
| 720 | 2 | MYCN | {'ess': 0, 'nonest': 7} | True | 15940096 | AAGCGTCTAGCAAGTCCGAGC | [] |
| 721 | 2 | CYRIA | unavailable | False | 16552935 | AGCTCTACTGAAGCATTGCTCG | [] |
| 722 | 2 | VSNL1 | {'ess': 1, 'nonest': 7} | True | 17655214 | GTGCCTACCATTTTGTAGATAG | [] |
| 723 | 2 | RDH14 | {'ess': 0, 'nonest': 7} | True | 18555190 | TCCTTGTTCCTATTTTAGCAGG | [] |
| 724 | 2 | NT5C1B | {'ess': 1, 'nonest': 7} | True | 18563795 | CCCCTAACTACTGAATTTTTTG | ['TTTTTATTAAAGCCATAAGCTG'] |
| 725 | 2 | PUM2 | {'ess': 1, 'nonest': 6} | True | 20278206 | TTCAGTAGCTATGTAACTAATG | [] |
| 726 | 2 | HS1BP3 | {'ess': 0, 'nonest': 7} | True | 20557561 | ACATTTCTATAGCATTTGGATG | [] |
| 727 | 2 | GDF7 | {'ess': 1, 'nonest': 7} | True | 20671424 | CTCGCGCTACCTGCAGCCGCA | [] |
| 728 | 2 | ATAD2B | {'ess': 1, 'nonest': 7} | True | 23773858 | TCTCCTCATCTATAAAATAAAG | [] |
| 729 | 2 | ATAD2B | {'ess': 1, 'nonest': 7} | True | 23801443 | ACTCACCTATAGGGTGTCTTGG | [] |
| 730 | 2 | FKBP1B | {'ess': 0, 'nonest': 7} | True | 24063062 | GCTCATCTAGCAGGGATGGGG | [] |
| 731 | 2 | ITSN2 | {'ess': 0, 'nonest': 7} | True | 24203625 | ACCCCTACAGGAGAGTTTTTTG | [] |
| 732 | 2 | NCOA1 | {'ess': 3, 'nonest': 4} | False | 24768122 | CTCTAGTCTGTAGTCACCACAG | [] |
| 733 | 2 | ADCY3 | {'ess': 0, 'nonest': 7} | True | 24820107 | TTTCTTCTACCACCTGGAGAGG | [] |
| 734 | 2 | DNAJC27 | {'ess': 0, 'nonest': 7} | True | 24947615 | CTTTCTACTTGATGTTTTTCAG | [] |
| 735 | 2 | DNMT3A | {'ess': 1, 'nonest': 7} | True | 25239489 | ACTAGCCCACTAGTACAGGTG | [] |
| 736 | 2 | DTNB | {'ess': 0, 'nonest': 7} | True | 25377329 | GAAGGTGCTAGTACATGCAGG | [] |
| 737 | 2 | DTNB | {'ess': 0, 'nonest': 7} | True | 25379318 | TCTGCTAACCTGTGGCAGCAT | [] |
| 738 | 2 | DTNB | {'ess': 0, 'nonest': 7} | True | 25387098 | CGCCCCTACGCGATCCTGTGT | [] |
| 739 | 2 | DTNB | {'ess': 0, 'nonest': 7} | True | 25558326 | TTCACACTACTGGGTCACAGAG | [] |
| 740 | 2 | CIB4 | {'ess': 0, 'nonest': 7} | True | 26629487 | TGTGTTCTAGGTGGTTGGAAGG | [] |
| 741 | 2 | TMEM214 | {'ess': 1, 'nonest': 7} | True | 27040836 | GGGCCTACTGCTGGGATATCA | [] |
| 742 | 2 | EMILIN1 | {'ess': 0, 'nonest': 7} | True | 27080014 | CCCCAGTCTACGCGTGTTCAA | [] |
| 743 | 2 | CGREF1 | {'ess': 1, 'nonest': 6} | True | 27099783 | ATGGAGACTACCATTGCGGCT | [] |
| 744 | 2 | CGREF1 | {'ess': 1, 'nonest': 6} | True | 27101273 | CTTCAAGATCTAGATCTCATCG | ['ATCATTCTCCACTTGAACAATG'] |
| 745 | 2 | ABHD1 | {'ess': 0, 'nonest': 7} | True | 27129552 | TGTCTCTAGATCTTCAGTATTG | [] |
| 746 | 2 | ABHD1 | {'ess': 0, 'nonest': 7} | True | 27130352 | TGGCATCTATCTTGGTTCTAGG | [] |
| 747 | 2 | PREB | {'ess': 6, 'nonest': 2} | False | 27131413 | AGGGAAGCTAAAGGAAACCTGG | [] |
| 748 | 2 | PRR30 | unavailable | False | 27137090 | GGAACTAGACTGATGACTTTTG | [] |
| 749 | 2 | SLC5A6 | {'ess': 0, 'nonest': 7} | True | 27200389 | CTCTGGCTATGGCCTGGCACA | [] |
| 750 | 2 | CAD | {'ess': 7, 'nonest': 2} | False | 27243517 | CCAGGCCCTAGAAACGGCCCA | [] |
| 751 | 2 | SLC30A3 | {'ess': 1, 'nonest': 7} | True | 27255085 | ACTAGTCACATCTATTCCCCTG | [] |
| 752 | 2 | GTF3C2 | {'ess': 5, 'nonest': 1} | False | 27326674 | GCCAAGGCTAGGGAGTGGGCA | [] |
| 753 | 2 | PPM1G | {'ess': 2, 'nonest': 6} | False | 27381598 | GATGACTGCTAGTCTCGCTTGG | [] |
| 754 | 2 | NRBP1 | {'ess': 6, 'nonest': 2} | False | 27441811 | TGAGCTCTAAGAGGAGACGGT | [] |
| 755 | 2 | KRTCAP3 | {'ess': 0, 'nonest': 7} | True | 27444055 | TGTCCTAAACCCAACTTCTCTG | [] |
| 756 | 2 | IFT172 | {'ess': 0, 'nonest': 7} | True | 27444631 | TACCAACTACTGAAAGGAAAAG | [] |
| 757 | 2 | FNDC4 | {'ess': 0, 'nonest': 7} | True | 27492395 | CATCCCTATCCCCAGTTGTTAG | [] |
| 758 | 2 | ZNF512 | {'ess': 1, 'nonest': 7} | True | 27621460 | TGACTGCCTATTTCCTTCCTCG | [] |
| 759 | 2 | GPN1 | {'ess': 7, 'nonest': 1} | False | 27650199 | CTATTTATTGTTTCTCTTCCAG | [] |
| 760 | 2 | MRPL33 | {'ess': 7, 'nonest': 1} | False | 27779555 | ATTACACGCTACAGGATTTCTG | [] |
| 761 | 2 | BABAM2 | unavailable | False | 28338512 | TGTTTCCTAGAGCTTTCCATTG | [] |
| 762 | 2 | PLB1 | {'ess': 1, 'nonest': 7} | True | 28633320 | GCCAACCTATCTGATTACAAAG | [] |
| 763 | 2 | PLB1 | {'ess': 1, 'nonest': 7} | True | 28633469 | TGCTAGGTGCCATGTACACATG | [] |
| 764 | 2 | PLB1 | {'ess': 1, 'nonest': 7} | True | 28643060 | CCGGGCCTAGAGGGCCACAGT | [] |
| 765 | 2 | SPDYA | {'ess': 0, 'nonest': 7} | True | 28840479 | TATTACTAAATCATACCTTCAG | [] |
| 766 | 2 | TOGARAM | unavailable | False | 29027014 | CCTGCCCTACCTGGTGTCCTG | [] |
| 767 | 2 | CLIP4 | {'ess': 0, 'nonest': 7} | True | 29179632 | GGTACCTAACACTTCACATACG | [] |
| 768 | 2 | LBH | {'ess': 1, 'nonest': 7} | True | 30257620 | ACAGGGACTCTACTGCTCTTTG | ['TGCTCTTTATTTTCTTTCGCTG'] |
| 769 | 2 | CAPN13 | {'ess': 0, 'nonest': 7} | True | 30748354 | TGGAAACTATCTCAGGCTGGG | [] |
| 770 | 2 | GALNT14 | {'ess': 1, 'nonest': 7} | True | 30911022 | GGGATGCTATGTGCTCGATGT | [] |
| 771 | 2 | FEZ2 | {'ess': 1, 'nonest': 7} | True | 36553162 | TGCTGCTCTATGTAGGACACAG | [] |
| 772 | 2 | EIF2AK2 | {'ess': 3, 'nonest': 6} | False | 37107272 | AAGGGCTCTAACATGTGTGTCG | [] |
| 773 | 2 | CEBPZOS | unavailable | False | 37201723 | TACTTACATCTAATTTTTGCTG | [] |
| 774 | 2 | RMDN2 | unavailable | False | 38054876 | CCTTACTATGGAGCATGTCCTG | [] |
| 775 | 2 | RMDN2 | unavailable | False | 38060889 | CAGACTACAGCTCCTATTAATG | [] |
| 776 | 2 | ATL2 | {'ess': 8, 'nonest': 1} | False | 38296492 | TACTGTCTAATTTTTCTTCTTG | [] |
| 777 | 2 | HNRNPLL | unavailable | False | 38564853 | ACTCTCTATAAAAGCCAATTCG | ['AATTCAAATACAGCTTGGGAAG'] |
| 778 | 2 | DHX57 | {'ess': 0, 'nonest': 7} | True | 38798442 | GTTCAGCCACCTAAAATGAAAG | [] |
| 779 | 2 | MORN2 | {'ess': 0, 'nonest': 7} | True | 38882514 | AACATCACATCTACATGTGAAG | [] |
| 780 | 2 | CDKL4 | {'ess': 0, 'nonest': 7} | True | 39175994 | ATTTTCCTAAATGTTTGGAAGG | [] |
| 781 | 2 | THUMPD2 | {'ess': 0, 'nonest': 7} | True | 39736734 | AGCCTGCTACAGTCCAGAAGA | [] |
| 782 | 2 | THUMPD2 | {'ess': 0, 'nonest': 7} | True | 39744606 | TCCTAACTTAAACTTTTTCCCG | ['TTTTTCCCAAATGGAATGTCAG'] |
| 783 | 2 | THUMPD2 | {'ess': 0, 'nonest': 7} | True | 39768850 | TCAGCTACTATAAGCCTATGTG | [] |
| 784 | 2 | DYNC2LI1 | {'ess': 0, 'nonest': 7} | True | 43810508 | ACTTGGCTAGCAGGTACACAC | ['CACACATCTACAGATTGGAATG'] |
| 785 | 2 | ABCG5 | {'ess': 1, 'nonest': 7} | True | 43813115 | GCTTTCACTACCTGCTAATGAG | [] |
| 786 | 2 | PPM1B | {'ess': 0, 'nonest': 7} | True | 44232344 | AAGGCTACCATGGGTCTTCTAG | [] |
| 787 | 2 | SLC3A1 | {'ess': 0, 'nonest': 7} | True | 44320638 | GTGCCTAACACGAGGTATACAG | [] |
| 788 | 2 | CAMKMT | unavailable | False | 44475793 | TGCTTCTAGTAAATCAACACTG | [] |
| 789 | 2 | CAMKMT | unavailable | False | 44505663 | CCTCAGCCTATTCAATGGTAAG | [] |
| 790 | 2 | CAMKMT | unavailable | False | 44772112 | TTAATCTTCTATCCATGTTTGG | [] |
| 791 | 2 | SIX2 | {'ess': 0, 'nonest': 7} | True | 45006169 | GTTCTAGGAGCCCAGGTCCAC | [] |
| 792 | 2 | PRKCE | {'ess': 3, 'nonest': 6} | False | 46010736 | AAAGAATTCTACAGCTCTCTCG | ['AGCTCTCTCATCCCTTCACAAG'] |
| 793 | 2 | PRKCE | {'ess': 3, 'nonest': 6} | False | 46024334 | CTTCAACTTCCTATTGCCTCTG | [] |
| 794 | 2 | RHOQ | {'ess': 0, 'nonest': 7} | True | 46576804 | GACCTACCTCACTGAGTTCATG | [] |
| 795 | 2 | PIGF | {'ess': 0, 'nonest': 7} | True | 46581505 | CTTTCTATTCCAGTATATCCAG | [] |
| 796 | 2 | CRIPT | {'ess': 1, 'nonest': 7} | True | 46624226 | AATACATCTAGACAGATGTTTG | [] |
| 797 | 2 | MCFD2 | {'ess': 1, 'nonest': 7} | True | 46905462 | AACATCTACTGCAGTGATTTTG | [] |
| 798 | 2 | STPG4 | unavailable | False | 47129825 | AACTGTCTACATTATACTTACG | [] |
| 799 | 2 | MSH2 | {'ess': 1, 'nonest': 6} | True | 47562110 | AATGCCTATGCTGATGCACTTG | [] |
| 800 | 2 | KCNK12 | {'ess': 0, 'nonest': 7} | True | 47520906 | GGCGGTCTACCTGGAGGCGCT | [] |
| 801 | 2 | MSH6 | {'ess': 0, 'nonest': 7} | True | 47806859 | GTCTATAATTCCTTAATCAAAG | [] |
| 802 | 2 | FOXN2 | {'ess': 0, 'nonest': 7} | True | 48375442 | AGTATTTCTATTTTTTCCGTTG | [] |
| 803 | 2 | PPP1R21 | unavailable | False | 48514743 | ATTTCAAAACTATCGACTCTTG | [] |
| 804 | 2 | STON1 | {'ess': 0, 'nonest': 7} | True | 48595301 | CTACTCCTACTGAGTTATGCAG | [] |
| 805 | 2 | NRXN1 | {'ess': 1, 'nonest': 6} | True | 50922504 | TTGAAAGCAGCTACAGAACAAG | [] |
| 806 | 2 | ERLEC1 | {'ess': 0, 'nonest': 7} | True | 53821343 | CGAGACTAGTCTGGGCAATATG | [] |
| 807 | 2 | PSME4 | {'ess': 1, 'nonest': 7} | True | 53866088 | CCTTTCTATGCATAATAGCATG | [] |
| 808 | 2 | ACYP2 | {'ess': 0, 'nonest': 7} | True | 54139188 | CATCTCCCTAACGCTGGAGGA | [] |
| 809 | 2 | ACYP2 | {'ess': 0, 'nonest': 7} | True | 54141906 | GTGAGTCTAGATCAGACCACTG | [] |
| 810 | 2 | EML6 | {'ess': 0, 'nonest': 7} | True | 54966853 | CAAAGCCTATGGCCAGCCCAT | [] |
| 811 | 2 | CLHC1 | unavailable | False | 55175789 | TATAGAACTACCAAAACACATG | [] |
| 812 | 2 | EFEMP1 | {'ess': 0, 'nonest': 7} | True | 55867072 | GAAAAGACTAAAATGAAAATGG | [] |
| 813 | 2 | LOC10798 | unavailable | False | 56184108 | CTGCTCTAGCGACGCCGCCGC | [] |
| 814 | 2 | VRK2 | {'ess': 1, 'nonest': 6} | True | 58154761 | CCTAAGCTTCTACCTAAACTGG | [] |
| 815 | 2 | VRK2 | {'ess': 1, 'nonest': 6} | True | 58154769 | GCCTAAGCTTCTACCTAAACTG | [] |
| 816 | 2 | FANCL | {'ess': 3, 'nonest': 5} | False | 58160047 | TATCTTCTAGAACATATTACTG | [] |
| 817 | 2 | BCL11A | {'ess': 1, 'nonest': 6} | True | 60460403 | ATACCTCTATTCAGTTTTTATG | ['TTATATCATTATTCAACACTCG'] |
| 818 | 2 | PUS10 | {'ess': 0, 'nonest': 7} | True | 60942394 | GAAAGCTAGTCATCCAGAGCAG | [] |
| 819 | 2 | COMMD1 | {'ess': 1, 'nonest': 7} | True | 62012723 | TCTTGCTACTTCTTACTGACAG | [] |
| 820 | 2 | EHBP1 | {'ess': 0, 'nonest': 7} | True | 63045499 | CTGATGGCTACTGAAGAACACG | ['AACACATTTCTCCTCTTTCTTG'] |
| 821 | 2 | WDPCP | unavailable | False | 63258420 | TAGATCTATGTGGTGGCATCAG | [] |
| 822 | 2 | WDPCP | unavailable | False | 63308836 | GTATACCTATGTGACAAACCTG | [] |
| 823 | 2 | WDR92 | {'ess': 7, 'nonest': 2} | False | 68134700 | AGACTTACTACTTCCAGAGGTG | [] |
| 824 | 2 | PNO1 | {'ess': 6, 'nonest': 3} | False | 68174779 | CTGATCTGCTAGCCACAGCTC | [] |
| 825 | 2 | APLF | {'ess': 0, 'nonest': 7} | True | 68578021 | TAGTTACTATTTTCTTTTCATG | ['TTTCTTTTCATAAACCTTTTTG'] |
| 826 | 2 | BMP10 | {'ess': 1, 'nonest': 6} | True | 68865630 | CTTCTATCTACAGCCACATTCG | [] |
| 827 | 2 | GKN2 | {'ess': 0, 'nonest': 7} | True | 68945367 | TCATCCTAAACATGAATGTCTG | [] |
| 828 | 2 | ANTXR1 | {'ess': 0, 'nonest': 7} | True | 69145375 | CTCTGCTATTCCATGCAAGCAG | [] |
| 829 | 2 | ANTXR1 | {'ess': 0, 'nonest': 7} | True | 69245494 | GGGCTCTAGACAGAAGGCCTT | [] |
| 830 | 2 | AAK1 | {'ess': 2, 'nonest': 6} | False | 69475868 | GTCACGGCTACAGGTCTATGA | [] |
| 831 | 2 | LOC11226 | unavailable | False | 69465424 | TGCTCCTAGGGTTTCTTGCCT | [] |
| 832 | 2 | PCBP1 | {'ess': 4, 'nonest': 4} | False | 70088813 | ACTGTTCTAGCTGCACCCCAT | [] |
| 833 | 2 | TIA1 | {'ess': 0, 'nonest': 6} | True | 70216203 | AACCATCCCTACCGAACAAATG | [] |
| 834 | 2 | ADD2 | {'ess': 0, 'nonest': 7} | True | 70672901 | GGAGAGACTAAAGGGCTCTTTG | [] |
| 835 | 2 | CD207 | {'ess': 0, 'nonest': 3} | True | 70825296 | ATGATGGCTAAATGTAACGTGG | [] |
| 836 | 2 | CD207 | {'ess': 0, 'nonest': 3} | True | 70830808 | ACTTCTAAATCCTCTCTACCCG | ['CCACCCCTCCCACTTTAACCTG'] |
| 837 | 2 | ATP6V1B1 | {'ess': 1, 'nonest': 7} | True | 70965120 | CGGGGCTAGAGCGCAGTGTCA | [] |
| 838 | 2 | NAGK | {'ess': 0, 'nonest': 7} | True | 71076677 | CAACACTGCTACGATGTGTCTG | [] |
| 839 | 2 | NAGK | {'ess': 0, 'nonest': 7} | True | 71078507 | AGCCCCCTAGGAAAAGGTGTA | [] |
| 840 | 2 | LOC10798 | unavailable | False | 71932887 | ACAGACTACCAGGAGCTCTGT | [] |
| 841 | 2 | EXOC6B | {'ess': 0, 'nonest': 7} | True | 72454322 | GAATTTCTATAGTCCTCTTTGG | [] |
| 842 | 2 | EMX1 | {'ess': 0, 'nonest': 7} | True | 72933953 | CCCACCCTAGTCATTGGAGGT | [] |
| 843 | 2 | SFXN5 | {'ess': 0, 'nonest': 7} | True | 72945080 | ATCTCCGGCTCTAATTGGGATG | [] |
| 844 | 2 | RAB11FIP5 | {'ess': 0, 'nonest': 7} | True | 73075520 | GAAGGCTATTTGGGGGGGCCC | [] |
| 845 | 2 | PRADC1 | unavailable | False | 73229453 | ACTCTTCTACCAGAAGGTCCAG | [] |
| 846 | 2 | TET3 | {'ess': 1, 'nonest': 6} | True | 74102175 | TGGCACCTAGATCCAGCGGCT | [] |
| 847 | 2 | PCGF1 | {'ess': 0, 'nonest': 7} | True | 74505242 | CCCTACCTCCTCTTCTCTTTCC | ['CTTTCACACTGTATTGTAAAAG'] |
| 848 | 2 | SEMA4F | {'ess': 1, 'nonest': 7} | True | 74680208 | TTTGCCCAGCTCTAGATGGATG | [] |
| 849 | 2 | HK2 | {'ess': 7, 'nonest': 1} | False | 74890940 | CAGGGGTTCTATCGCTGTCCA | [] |
| 850 | 2 | TACR1 | {'ess': 0, 'nonest': 7} | True | 75049431 | TGTGGCCTAGGAGAGCACATT | [] |
| 851 | 2 | EVA1A | unavailable | False | 75493235 | CTCCCTAATAGTAGCGATTCAG | [] |
| 852 | 2 | GCFC2 | unavailable | False | 75702169 | ATATCCTAATCTTCATATATTG | [] |
| 853 | 2 | REG3G | {'ess': 0, 'nonest': 7} | True | 79029275 | CCTGCCCTAGTCCTTGAACTT | [] |
| 854 | 2 | REG1B | {'ess': 0, 'nonest': 7} | True | 79085215 | CTTCCTCTAGTTTTTGAACTTG | [] |
| 855 | 2 | REG1A | {'ess': 1, 'nonest': 7} | True | 79123214 | TTGCCTCTAGTTTTTGAACTTG | [] |
| 856 | 2 | REG3A | {'ess': 0, 'nonest': 7} | True | 79157225 | CCTGCACTAGTCAGTGAACTTG | [] |
| 857 | 2 | CTNNA2 | {'ess': 0, 'nonest': 7} | True | 80647871 | TCGTCCTAGAAGGAATCCATTG | [] |
| 858 | 2 | TRABD2A | unavailable | False | 84838230 | TCAAGGCTACTGTGAAGCTGG | [] |
| 859 | 2 | RETSAT | {'ess': 0, 'nonest': 7} | True | 85343241 | GGAACTAATTCTTTTTCTTCTG | [] |
| 860 | 2 | SFTPB | {'ess': 0, 'nonest': 5} | True | 85659780 | GCCTCCTAAGGTGCTGGGATT | [] |
| 861 | 2 | POLR1A | {'ess': 7, 'nonest': 1} | False | 86027422 | GTAGCTGCTATCTCAGAGGCT | [] |
| 862 | 2 | MRPL35 | {'ess': 6, 'nonest': 2} | False | 86210667 | CTGATCTATACTTTCAGGTTTG | [] |
| 863 | 2 | REEP1 | {'ess': 0, 'nonest': 6} | True | 86217099 | AAGGATTCTAGGCGGTGCCTG | [] |
| 864 | 2 | RNF103-C | unavailable | False | 86505803 | CAGCCCCTAGCTGCGGAGTGT | [] |
| 865 | 2 | CHMP3 | unavailable | False | 86505803 | CAGCCCCTAGCTGCGGAGTGT | [] |
| 866 | 2 | CD8B | {'ess': 0, 'nonest': 7} | True | 86815606 | GCTTCTTGCCTATGTTTTCAGG | [] |
| 867 | 2 | RGPD1 | {'ess': 0, 'nonest': 5} | True | 86993646 | GACTCCTAGTTCTTCAGTTTTG | [] |
| 868 | 2 | RGPD1 | {'ess': 0, 'nonest': 5} | True | 87012546 | TTGCTTTCTATTTTTTCTTGGG | [] |
| 869 | 2 | RGPD2 | {'ess': 0, 'nonest': 7} | True | 87757391 | TTGCTTTCTATTTTTTCTTGGG | [] |
| 870 | 2 | RGPD2 | {'ess': 0, 'nonest': 7} | True | 87776276 | GACTCCTAGTTCTTCAGTTTTG | [] |
| 871 | 2 | THNSL2 | {'ess': 0, 'nonest': 5} | True | 88180122 | GCCAGGCTACTGGGAGGTGTT | [] |
| 872 | 2 | FOXI3 | unavailable | False | 88448206 | GAGTCTGCTCTACACCTCGGA | [] |
| 873 | 2 | TEX37 | unavailable | False | 88529473 | ATCCTTCCTACTGGTAAAATGG | [] |
| 874 | 2 | EIF2AK3 | {'ess': 0, 'nonest': 7} | True | 88557735 | TAAGGCTAATTGCTTGGCAAAG | [] |
| 875 | 2 | LOC10106 | unavailable | False | 90362906 | ACGCTACCCTAGCTCAAACAAG | [] |
| 876 | 2 | PROM2 | {'ess': 0, 'nonest': 7} | True | 95288995 | AAGGCCCTACAGCTTCAGGGA | [] |
| 877 | 2 | KCNIP3 | {'ess': 1, 'nonest': 6} | True | 95384048 | CCTAGATGACATTCTCAAACAG | [] |
| 878 | 2 | FAHD2A | {'ess': 0, 'nonest': 7} | True | 95414338 | CCAGCACTAGATTTTCTTTTTG | [] |
| 879 | 2 | ANKRD36 | unavailable | False | 95851171 | TCATCTAGCTCTCAAACACCTG | [] |
| 880 | 2 | GPAT2 | {'ess': 0, 'nonest': 7} | True | 96022158 | AGTTCTAGCTACAAATGAACTG | [] |
| 881 | 2 | GPAT2 | {'ess': 0, 'nonest': 7} | True | 96022669 | CTCACCCCTAGGTCTCTGAAG | [] |
| 882 | 2 | ASTL | {'ess': 1, 'nonest': 7} | True | 96124246 | GGGCTCCTACCAGTGCTGTGG | [] |
| 883 | 2 | ARID5A | {'ess': 1, 'nonest': 7} | True | 96552312 | GCTGGCCTACAGCTTGGTGTT | [] |
| 884 | 2 | KANSL3 | unavailable | False | 96600065 | ATCTACCTAACAGAAGGTGGTG | [] |
| 885 | 2 | CNNM4 | {'ess': 1, 'nonest': 6} | True | 96802836 | CCCTATTCCCACCGACTCCCC | [] |
| 886 | 2 | FAHD2B | {'ess': 0, 'nonest': 7} | True | 97082405 | GTCGGGCTACACAGACCAGCT | [] |
| 887 | 2 | ANKRD36 | unavailable | False | 97118522 | TTGGTTGACCTATACCTTGCTG | [] |
| 888 | 2 | ANKRD36 | unavailable | False | 97250166 | TCATCTAGCTCTCAAACACCTG | [] |
| 889 | 2 | ANKRD36 | unavailable | False | 97507041 | TCATCTAGCTCTCAAACACCTG | [] |
| 890 | 2 | ANKRD36 | unavailable | False | 97522061 | TCCAAATCTATAAAATTTCTAG | [] |
| 891 | 2 | ANKRD36 | unavailable | False | 97584937 | TTGGTTGACCTATACCTTGCTG | [] |
| 892 | 2 | ACTR1B | {'ess': 0, 'nonest': 7} | True | 97656857 | TTGGGCACTAGAAAGTTTTGCG | [] |
| 893 | 2 | ZAP70 | {'ess': 1, 'nonest': 6} | True | 97738231 | AAAGGCTATGAACTGGGTGAAG | [] |
| 894 | 2 | VWA3B | {'ess': 0, 'nonest': 7} | True | 98312235 | GAGTCCTAGACGCCCGGCCGC | [] |
| 895 | 2 | INPP4A | {'ess': 0, 'nonest': 7} | True | 98582395 | CTCTTGCTACAGTGTACTACTG | [] |
| 896 | 2 | KIAA1211L | unavailable | False | 98794531 | GCCCACCTATTTTATAATCTGG | [] |
| 897 | 2 | TSGA10 | {'ess': 1, 'nonest': 7} | True | 99018525 | ATCTCTACTCTGGATGGCCCT | [] |
| 898 | 2 | TSGA10 | {'ess': 1, 'nonest': 7} | True | 99027208 | CTACTATACGGCCTTTAAAAAG | [] |
| 899 | 2 | TSGA10 | {'ess': 1, 'nonest': 7} | True | 99028755 | TTCTCTATTCAACTTCTAACTG | [] |
| 900 | 2 | LYG2 | {'ess': 0, 'nonest': 5} | True | 99242363 | AGAGCTTTGCCTAGAAGCTTTG | [] |
| 901 | 2 | TXNDC9 | {'ess': 0, 'nonest': 7} | True | 99306772 | ACGGCCCTAGGGCCACCACCT | [] |
| 902 | 2 | TXNDC9 | {'ess': 0, 'nonest': 7} | True | 99319581 | TGAGCTCTAATCATCATCAGAG | [] |
| 903 | 2 | AFF3 | {'ess': 0, 'nonest': 5} | True | 99551473 | AGGTCCCTATGACAGGTGGGC | [] |
| 904 | 2 | NPAS2 | {'ess': 0, 'nonest': 6} | True | 1.01E+08 | GAGGTACTACCCAGGACCGAG | [] |
| 905 | 2 | NPAS2 | {'ess': 0, 'nonest': 6} | True | 1.01E+08 | ATGGCTATAGAGTGTGCCCAAG | [] |
| 906 | 2 | TBC1D8 | {'ess': 0, 'nonest': 7} | True | 1.01E+08 | TGTCTTGCTACAAGTTACTCAG | [] |
| 907 | 2 | TBC1D8 | {'ess': 0, 'nonest': 7} | True | 1.01E+08 | ATCTAGAAACCTGAACACACCG | [] |
| 908 | 2 | RNF149 | {'ess': 0, 'nonest': 7} | True | 1.01E+08 | GGCACGTGTGCTAGGAGATGG | [] |
| 909 | 2 | MAP4K4 | {'ess': 1, 'nonest': 6} | True | 1.02E+08 | TGCTTCTACCAGCTCAGAAGAG | [] |
| 910 | 2 | IL1R1 | {'ess': 0, 'nonest': 5} | True | 1.02E+08 | TCCATGCTACCCGAGAGGCAC | [] |
| 911 | 2 | IL1RL2 | {'ess': 1, 'nonest': 7} | True | 1.02E+08 | CTTTCCTATAAGAGCCCCAGAG | [] |
| 912 | 2 | IL1RL1 | {'ess': 0, 'nonest': 7} | True | 1.02E+08 | AGGCACTATTGCTTCTGGGCA | [] |
| 913 | 2 | SLC9A4 | {'ess': 0, 'nonest': 7} | True | 1.03E+08 | AATAACACTATTTTTTTTGGAG | [] |
| 914 | 2 | MFSD9 | {'ess': 0, 'nonest': 7} | True | 1.03E+08 | CATCTACTCACTTTTTAATTTG | ['TTACTATTCCCATCACCACTAG'] |
| 915 | 2 | TMEM182 | {'ess': 0, 'nonest': 7} | True | 1.03E+08 | AAGTGCCTACCTCAGAAAGTAG | [] |
| 916 | 2 | GPR45 | {'ess': 0, 'nonest': 7} | True | 1.05E+08 | GACCCCCTAAACCGCAGACTG | [] |
| 917 | 2 | TGFBRAP1 | {'ess': 1, 'nonest': 4} | False | 1.05E+08 | GGGTCTCTATGTTCCATCTCCC | [] |
| 918 | 2 | CD8B2 | unavailable | False | 1.07E+08 | CCCTTGCTAGGCATCCATGGG | [] |
| 919 | 2 | ST6GAL2 | {'ess': 0, 'nonest': 7} | True | 1.07E+08 | CCCAGCTACTTGGGAGGCTGA | [] |
| 920 | 2 | RGPD4 | {'ess': 0, 'nonest': 7} | True | 1.08E+08 | GACTCCTAGTTCTTCAGTTTTG | [] |
| 921 | 2 | SULT1C4 | {'ess': 0, 'nonest': 7} | True | 1.08E+08 | TCCTTACTAGAACTGGAAGTGG | [] |
| 922 | 2 | GCC2 | {'ess': 0, 'nonest': 7} | True | 1.09E+08 | CCATCAACCTATCGAAGTCCAG | [] |
| 923 | 2 | RANBP2 | {'ess': 3, 'nonest': 5} | False | 1.09E+08 | TGGAAGCCTATTCATGCCATAG | [] |
| 924 | 2 | CCDC138 | {'ess': 0, 'nonest': 7} | True | 1.09E+08 | TTTCTAGTATTTGAAGATCAAG | [] |
| 925 | 2 | CCDC138 | {'ess': 0, 'nonest': 7} | True | 1.09E+08 | GCCAGTCTAAGGGCTTGCACT | [] |
| 926 | 2 | LOC10798 | unavailable | False | 1.09E+08 | TAGAAGCTAAGGAATTAGGAAG | [] |
| 927 | 2 | NPHP1 | {'ess': 1, 'nonest': 7} | True | 1.1E+08 | AACCCACCTAATGCCATTCCAG | [] |
| 928 | 2 | LIMS4 | unavailable | False | 1.1E+08 | GGGTGACTAAGGGTAGGATGG | [] |
| 929 | 2 | RGPD5 | {'ess': 2, 'nonest': 5} | False | 1.11E+08 | GATGGCCTAGTGATCGTGTAAG | [] |
| 930 | 2 | ACOXL | {'ess': 0, 'nonest': 7} | True | 1.11E+08 | GTTACTACACATGAAATCTTAG | [] |
| 931 | 2 | ACOXL | {'ess': 0, 'nonest': 7} | True | 1.11E+08 | GGCTACAGGAAACACTTCCAAG | [] |
| 932 | 2 | ACOXL | {'ess': 0, 'nonest': 7} | True | 1.11E+08 | TCTAAACTTCCATTTTCACATG | [] |
| 933 | 2 | ACOXL | {'ess': 0, 'nonest': 7} | True | 1.11E+08 | AATGCTAGGCATCTTCCGCCT | [] |
| 934 | 2 | BCL2L11 | {'ess': 1, 'nonest': 7} | True | 1.11E+08 | CCCAGCTACTTGGGAGGCTGA | [] |
| 935 | 2 | BCL2L11 | {'ess': 1, 'nonest': 7} | True | 1.11E+08 | AGCTGTTTCTAGTCCCAGTTAG | [] |
| 936 | 2 | BCL2L11 | {'ess': 1, 'nonest': 7} | True | 1.11E+08 | AAAGGTCACAACTAGGCAACAG | [] |
| 937 | 2 | BCL2L11 | {'ess': 1, 'nonest': 7} | True | 1.11E+08 | CTTCCTCTATTTCTCTAACTGG | [] |
| 938 | 2 | BCL2L11 | {'ess': 1, 'nonest': 7} | True | 1.11E+08 | AACTACACGACAACTTCCTCTG | ['CTTCCTCTATTTCTCTAACTGG'] |
| 939 | 2 | FBLN7 | {'ess': 1, 'nonest': 7} | True | 1.12E+08 | CGCTGCCCTAGAAGCCTGGGT | [] |
| 940 | 2 | ZC3H6 | {'ess': 0, 'nonest': 7} | True | 1.12E+08 | AATAGCTAACAAAATGGTGAAG | [] |
| 941 | 2 | TTL | {'ess': 0, 'nonest': 7} | True | 1.13E+08 | TGCTACAACCAGCCTTCTCCC | [] |
| 942 | 2 | CHCHD5 | {'ess': 0, 'nonest': 7} | True | 1.13E+08 | AGGGGCTATATGGGGCAAGAA | [] |
| 943 | 2 | NT5DC4 | {'ess': 0, 'nonest': 3} | True | 1.13E+08 | GGCACTAGGAAGTGCTCAAAAG | [] |
| 944 | 2 | NT5DC4 | {'ess': 0, 'nonest': 3} | True | 1.13E+08 | TCTTTGCCCTAACCCTTATAAG | [] |
| 945 | 2 | NT5DC4 | {'ess': 0, 'nonest': 3} | True | 1.13E+08 | TTGGAATCCTAGATGGCAGAAG | [] |
| 946 | 2 | IL1A | {'ess': 2, 'nonest': 6} | False | 1.13E+08 | AGACCTACGCCTGGTTTTCCAC | [] |
| 947 | 2 | IL37 | unavailable | False | 1.13E+08 | TTCCTAATCGCTGACCTCACTG | [] |
| 948 | 2 | IL36B | unavailable | False | 1.13E+08 | CTTATTTCTACATCCTTCCTGG | [] |
| 949 | 2 | IL36RN | unavailable | False | 1.13E+08 | GTTGCCCTAGTCACACTGCTG | [] |
| 950 | 2 | IL1F10 | {'ess': 1, 'nonest': 7} | True | 1.13E+08 | TCCCTACCAGCTCTGTTCAAAG | [] |
| 951 | 2 | IL1RN | {'ess': 0, 'nonest': 7} | True | 1.13E+08 | GCAGTACTACTCGTCCTCCTG | [] |
| 952 | 2 | PAX8 | {'ess': 1, 'nonest': 7} | True | 1.13E+08 | TGGCAACTACAGATGGTCAAAG | [] |
| 953 | 2 | CBWD2 | {'ess': 0, 'nonest': 7} | True | 1.13E+08 | TCCTTATCTAAATTTCTGCCTG | [] |
| 954 | 2 | FOXD4L1 | {'ess': 2, 'nonest': 5} | False | 1.14E+08 | AGCCTAGAGCCCTCTGCCACC | [] |
| 955 | 2 | RABL2A | {'ess': 0, 'nonest': 7} | True | 1.14E+08 | CAAACTATTCCTAAAAAAAAAG | [] |
| 956 | 2 | SLC35F5 | {'ess': 0, 'nonest': 7} | True | 1.14E+08 | TTTTTCTACAAAAACTTGCCAG | [] |
| 957 | 2 | SLC35F5 | {'ess': 0, 'nonest': 7} | True | 1.14E+08 | GCTAACTAGCTCCATCCTCCT | [] |
| 958 | 2 | EN1 | {'ess': 0, 'nonest': 7} | True | 1.19E+08 | CGGCTACTCGCTCTCGTCTTT | [] |
| 959 | 2 | C1QL2 | {'ess': 0, 'nonest': 7} | True | 1.19E+08 | CGCGCCCCTAATCCGGGTACA | [] |
| 960 | 2 | GLI2 | {'ess': 1, 'nonest': 4} | False | 1.21E+08 | GCCTCTAGGTCATCATGTTCAG | [] |
| 961 | 2 | TSN | {'ess': 0, 'nonest': 7} | True | 1.22E+08 | GCCTCCTATTTTTCAACACAAG | [] |
| 962 | 2 | TEX51 | unavailable | False | 1.27E+08 | TCTCCTAGGCCTCAGAGCTTT | [] |
| 963 | 2 | MAP3K2 | {'ess': 2, 'nonest': 6} | False | 1.27E+08 | TTACTGGCTGCTAGTGATAATG | [] |
| 964 | 2 | PROC | {'ess': 0, 'nonest': 5} | True | 1.27E+08 | GGAGGGTCGCTAAGGTGCCCA | [] |
| 965 | 2 | IWS1 | {'ess': 4, 'nonest': 4} | False | 1.27E+08 | CCAGGCCTATAGGGGCATCCT | [] |
| 966 | 2 | MYO7B | {'ess': 0, 'nonest': 5} | True | 1.28E+08 | CCGCTGCTAGGTGCTGGCCAG | [] |
| 967 | 2 | WDR33 | {'ess': 7, 'nonest': 1} | False | 1.28E+08 | CAGCTTCTACCGACCCCTTCC | ['CCCCTTCCACCACCCCGTGAAG'] |
| 968 | 2 | WDR33 | {'ess': 7, 'nonest': 1} | False | 1.28E+08 | CTCCGTCTAAGGAGCTGGAAC | [] |
| 969 | 2 | WDR33 | {'ess': 7, 'nonest': 1} | False | 1.28E+08 | TGTTCTGGCTACAATGTCCTTG | [] |
| 970 | 2 | SAP130 | {'ess': 5, 'nonest': 4} | False | 1.28E+08 | GGGTCTAGACTTTTTCCTTTCG | [] |
| 971 | 2 | HS6ST1 | {'ess': 0, 'nonest': 7} | True | 1.28E+08 | CCGCCACTACCACTTCTCAAT | [] |
| 972 | 2 | POTEF | {'ess': 0, 'nonest': 5} | True | 1.3E+08 | GTCCACCTACAAGCATTTGCG | [] |
| 973 | 2 | MZT2B | {'ess': 0, 'nonest': 7} | True | 1.3E+08 | CCATCCTAGGTGCTGCCCCGT | [] |
| 974 | 2 | POTEI | {'ess': 0, 'nonest': 5} | True | 1.3E+08 | GTCCACCTAGAAGCATTTGCG | [] |
| 975 | 2 | POTEJ | {'ess': 0, 'nonest': 5} | True | 1.31E+08 | GTCCACCTAGAAGCATTTGCG | [] |
| 976 | 2 | GPR148 | {'ess': 0, 'nonest': 7} | True | 1.31E+08 | AGAACTCTAGGAAATGGTAAAG | [] |
| 977 | 2 | AMER3 | unavailable | False | 1.31E+08 | CTGTCCTACAGATGGGGCTCA | [] |
| 978 | 2 | PLEKHB2 | {'ess': 0, 'nonest': 7} | True | 1.31E+08 | GCCCCTAGAAGACCCAAAATAG | [] |
| 979 | 2 | POTEE | {'ess': 0, 'nonest': 4} | True | 1.31E+08 | GTCCACCTAGAAGCATTTGCG | [] |
| 980 | 2 | MZT2A | {'ess': 0, 'nonest': 7} | True | 1.31E+08 | CCATCCTAGGTGCTGCCCTGC | [] |
| 981 | 2 | ANKRD30 | {'ess': 0, 'nonest': 1} | False | 1.32E+08 | ACTGTATCCTATTCATCAGGTG | [] |
| 982 | 2 | MGAT5 | {'ess': 0, 'nonest': 5} | True | 1.34E+08 | TAGCTGCTATAGGCAGTCTTTG | [] |
| 983 | 2 | MAP3K19 | unavailable | False | 1.35E+08 | TCCTTGACTAGTGAGACCACAG | [] |
| 984 | 2 | RAB3GAP1 | {'ess': 0, 'nonest': 7} | True | 1.35E+08 | GGGGGGCTACACATCCCCATC | [] |
| 985 | 2 | UBXN4 | {'ess': 0, 'nonest': 7} | True | 1.36E+08 | GTCACACTACATCTGTTGAGTG | [] |
| 986 | 2 | KYNU | {'ess': 0, 'nonest': 7} | True | 1.43E+08 | CATTCCTAATTAAAGAACTCCG | [] |
| 987 | 2 | KYNU | {'ess': 0, 'nonest': 7} | True | 1.43E+08 | CACTGCTAATTTTTTTGTTTCTG | [] |
| 988 | 2 | LYPD6 | {'ess': 0, 'nonest': 7} | True | 1.49E+08 | CACTGAGCCACTATAACATGAG | [] |
| 989 | 2 | MMADHC | {'ess': 0, 'nonest': 7} | True | 1.5E+08 | ATTTCTGCTAATTTCCACTTAG | ['CACTTAATTTCTTCATAATATG'] |
| 990 | 2 | NMI | {'ess': 1, 'nonest': 7} | True | 1.51E+08 | TAAGTCTATTCTTCAAAGTATG | [] |
| 991 | 2 | RIF1 | {'ess': 4, 'nonest': 4} | False | 1.51E+08 | AATACTAAATAGAATTTTCATG | [] |
| 992 | 2 | NEB | {'ess': 0, 'nonest': 7} | True | 1.51E+08 | AAATGCCTAAATAGCTTCAACG | [] |
| 993 | 2 | STAM2 | {'ess': 0, 'nonest': 7} | True | 1.52E+08 | TTTGTTTCTAAAGGAGAGGCTG | [] |
| 994 | 2 | GALNT5 | {'ess': 1, 'nonest': 7} | True | 1.57E+08 | AACTAGCTGCTCTGCACATCTG | [] |
| 995 | 2 | UPP2 | {'ess': 0, 'nonest': 7} | True | 1.58E+08 | GATGCTGTCCCTAGCTGAGGA | [] |
| 996 | 2 | UPP2 | {'ess': 0, 'nonest': 7} | True | 1.58E+08 | ACGTCTAGTCACAAAGTCCAAG | [] |
| 997 | 2 | CCDC148 | {'ess': 0, 'nonest': 7} | True | 1.58E+08 | GAGCTCTATATTTTAAATACAG | [] |
| 998 | 2 | PKP4 | {'ess': 0, 'nonest': 7} | True | 1.59E+08 | AGCCGTCTATTTTGTTCTCTTG | [] |
| 999 | 2 | PKP4 | {'ess': 0, 'nonest': 7} | True | 1.59E+08 | CTTGACTATAATACAGCTGTTG | [] |
| 1000 | 2 | PKP4 | {'ess': 0, 'nonest': 7} | True | 1.59E+08 | TGATGCTACACCCATGAGTCTG | [] |
| 1001 | 2 | BAZ2B | {'ess': 0, 'nonest': 7} | True | 1.59E+08 | TCCACTTCTAAGATGCCTCAGG | [] |
| 1002 | 2 | ITGB6 | {'ess': 0, 'nonest': 7} | True | 1.6E+08 | AGTTCTAGCAATCTGTGGAAAG | [] |
| 1003 | 2 | TBR1 | {'ess': 1, 'nonest': 7} | True | 1.61E+08 | GGCGGCCTAGCTGTGCGAGTA | [] |
| 1004 | 2 | DPP4 | {'ess': 0, 'nonest': 7} | True | 1.62E+08 | TTTGAGGTGCTAAGGTAAAGAG | [] |
| 1005 | 2 | IFIH1 | {'ess': 0, 'nonest': 5} | True | 1.62E+08 | CAAGTGCTAATCCTCATCACTG | ['TCACTAAATAAACAGCATTCTG'] |
| 1006 | 2 | KCNH7 | {'ess': 1, 'nonest': 7} | True | 1.62E+08 | CTACTTACAGCCAATCAACATG | [] |
| 1007 | 2 | GRB14 | {'ess': 1, 'nonest': 6} | True | 1.64E+08 | CTTGTCTAGAGAGCAATCCTAG | [] |

| | | | | | | | |
|---|---|---|---|---|---|---|---|
| 1008 | 2 | SLC38A11 | {'exs': 0, 'nonexs': 7} | True | 1.65E+08 | CAGACTATAAAGAAAGATCTG | [] |
| 1009 | 2 | CSRNP3 | {'exs': 0, 'nonexs': 7} | True | 1.66E+08 | TTAAGCTACTAAACAGGGACTG | [] |
| 1010 | 2 | SCN9A | {'exs': 0, 'nonexs': 7} | True | 1.66E+08 | GCTCTATTTTTTGCTTTCCTTG | [] |
| 1011 | 2 | SCN9A | {'exs': 0, 'nonexs': 7} | True | 1.66E+08 | AAATCAGCTAGAAACATACCTG | [] |
| 1012 | 2 | B3GALT1 | {'exs': 0, 'nonexs': 5} | True | 1.68E+08 | AAAAATCCTAACATCTGAGATG | [] |
| 1013 | 2 | CERS6 | unavailable | False | 1.69E+08 | CAGCCCTCTAGTTAGCTCTTTG | [] |
| 1014 | 2 | NOSTRIN | {'exs': 1, 'nonexs': 7} | True | 1.69E+08 | CCAAGAGCTACTTTTTATTTGG | [] |
| 1015 | 2 | G6PC2 | {'exs': 1, 'nonexs': 7} | True | 1.69E+08 | ACTCTACTGACTCTTCTTTCCG | [] |
| 1016 | 2 | LRP2 | {'exs': 0, 'nonexs': 7} | True | 1.69E+08 | CTGGTATAGCTATACTTCAGAG | [] |
| 1017 | 2 | FASTKD1 | {'exs': 0, 'nonexs': 7} | True | 1.7E+08 | TAAAAACTACAAACATGACTTG | [] |
| 1018 | 2 | PPIG | {'exs': 0, 'nonexs': 7} | True | 1.7E+08 | TCTCACTAGATGCACTTAACAG | [] |
| 1019 | 2 | CCDC173 | unavailable | False | 1.7E+08 | TTTTCTACCATGTAAAACCTAG | [] |
| 1020 | 2 | SSB | {'exs': 3, 'nonexs': 5} | False | 1.7E+08 | TTACTAAACTACTGGTCTCCAG | [] |
| 1021 | 2 | DCAF17 | {'exs': 0, 'nonexs': 7} | True | 1.71E+08 | TTACCTACACTGTGACCACCAG | [] |
| 1022 | 2 | DYNC1I2 | {'exs': 5, 'nonexs': 3} | False | 1.72E+08 | CAGGAACTAAGCAGGTATTCGG | [] |
| 1023 | 2 | ITGA6 | {'exs': 1, 'nonexs': 6} | True | 1.73E+08 | AATACTATGCATCAGAAGTAAG | [] |
| 1024 | 2 | ITGA6 | {'exs': 1, 'nonexs': 6} | True | 1.73E+08 | CTATGAGTAGCTTTCATTTTCG | [] |
| 1025 | 2 | PDK1 | {'exs': 1, 'nonexs': 6} | True | 1.73E+08 | TGTGTCTAGGCACTGCGGAAC | [] |
| 1026 | 2 | RAPGEF4 | {'exs': 0, 'nonexs': 7} | True | 1.73E+08 | GAAATGTCTATGGTCGACGAGG | [] |
| 1027 | 2 | SP9 | {'exs': 1, 'nonexs': 6} | True | 1.74E+08 | CGGCCTCTAAGAGTCGTTGGG | [] |
| 1028 | 2 | SCRN3 | {'exs': 0, 'nonexs': 7} | True | 1.74E+08 | TGATCACTAAGAACTAACTTTG | [] |
| 1029 | 2 | LNPK | {'exs': 0, 'nonexs': 6} | True | 1.76E+08 | ATTTACTACTCTGCCGTCAAAG | [] |
| 1030 | 2 | LNPK | {'exs': 0, 'nonexs': 6} | True | 1.76E+08 | GTGTTCTAAAGATTCTGAAAAG | [] |
| 1031 | 2 | HOXD12 | {'exs': 0, 'nonexs': 7} | True | 1.76E+08 | CGCGGCTAGTAGAGCGCCAGC | [] |
| 1032 | 2 | HOXD10 | {'exs': 0, 'nonexs': 7} | True | 1.76E+08 | TCAGACCTAAGAAAACGTGAGG | [] |
| 1033 | 2 | HOXD4 | {'exs': 0, 'nonexs': 7} | True | 1.76E+08 | CCACTTCTATAAGGTCGTCAGG | [] |
| 1034 | 2 | HOXD3 | {'exs': 0, 'nonexs': 7} | True | 1.76E+08 | GCGGCCGCTACAGATGCGTCA | [] |
| 1035 | 2 | MTX2 | {'exs': 3, 'nonexs': 6} | False | 1.76E+08 | CACATAACTCTATGACAGCCTG | [] |
| 1036 | 2 | NFE2L2 | {'exs': 1, 'nonexs': 6} | True | 1.77E+08 | ATCTAGTTTTTCTTAACATCTG | [] |
| 1037 | 2 | TTC30B | {'exs': 1, 'nonexs': 7} | True | 1.78E+08 | TATTACTATATATTCCATCCTG | ['CCTATAATCTCATAAATCAAAG'] |
| 1038 | 2 | TTC30A | {'exs': 0, 'nonexs': 7} | True | 1.78E+08 | CATAACTACTTATTCCATCCTG | ['CCTATAATCTCATAAATCAAAG'] |
| 1039 | 2 | OSBPL6 | {'exs': 0, 'nonexs': 7} | True | 1.78E+08 | CCCAGTCTACCAAAGAACAGG | [] |
| 1040 | 2 | PJVK | unavailable | False | 1.78E+08 | CATCACTCTTCTATCTAAAAGG | [] |
| 1041 | 2 | FKBP7 | {'exs': 0, 'nonexs': 7} | True | 1.78E+08 | AAATATGCTATAGTTCATCGTG | [] |
| 1042 | 2 | CCDC141 | {'exs': 0, 'nonexs': 1} | False | 1.79E+08 | CGCCGCTAGAGTTTTGGGCCC | [] |
| 1043 | 2 | NEUROD1 | {'exs': 1, 'nonexs': 6} | True | 1.82E+08 | GTGCCTCTAATCATGAAATATG | [] |
| 1044 | 2 | PDE1A | {'exs': 0, 'nonexs': 5} | True | 1.82E+08 | CCTATGAATGTGTCTCATCATG | [] |
| 1045 | 2 | NUP35 | {'exs': 1, 'nonexs': 6} | True | 1.83E+08 | GTTCTACTACCAGCCAAACATG | [] |
| 1046 | 2 | ZNF804A | {'exs': 0, 'nonexs': 7} | True | 1.85E+08 | GTGATGACTAGAAGAGAGGTTG | [] |
| 1047 | 2 | FSIP2 | {'exs': 0, 'nonexs': 5} | True | 1.86E+08 | ATGAATTTCTAGCAACGGCCTG | [] |
| 1048 | 2 | ASDURF | unavailable | False | 1.9E+08 | GGTTGACTATTTCTTGATTTTG | [] |
| 1049 | 2 | ANKAR | {'exs': 0, 'nonexs': 5} | True | 1.9E+08 | CATTCCCACTATCGCCGCCCA | [] |
| 1050 | 2 | ANKAR | {'exs': 0, 'nonexs': 5} | True | 1.9E+08 | CCGCTCGCTAGCGGAACAGCA | [] |
| 1051 | 2 | ORMDL1 | {'exs': 0, 'nonexs': 7} | True | 1.9E+08 | CAATAACTAATAAATCATCTGG | [] |
| 1052 | 2 | INPP1 | {'exs': 1, 'nonexs': 6} | True | 1.9E+08 | TTCCTCTAGGTATGCGTCTCTG | [] |
| 1053 | 2 | NAB1 | {'exs': 0, 'nonexs': 7} | True | 1.91E+08 | CACAGCTATCTTGAATCTTCAG | [] |
| 1054 | 2 | STAT1 | {'exs': 2, 'nonexs': 6} | False | 1.91E+08 | TGCTCTATACTGTGTTCATCTG | [] |
| 1055 | 2 | TMEFF2 | {'exs': 0, 'nonexs': 7} | True | 1.92E+08 | AATGCTATTACCTAAAGTTTGG | [] |
| 1056 | 2 | TMEFF2 | {'exs': 0, 'nonexs': 7} | True | 1.92E+08 | TGCTCTAGTTAGGCTTACCAAG | [] |
| 1057 | 2 | STK17B | {'exs': 1, 'nonexs': 6} | True | 1.96E+08 | AAGTGCTAACAGAGCAAATCTG | [] |
| 1058 | 2 | CCDC150 | {'exs': 0, 'nonexs': 7} | True | 1.97E+08 | TGTTCAACTTCTACCTGAAATG | [] |
| 1059 | 2 | ANKRD44 | {'exs': 0, 'nonexs': 5} | True | 1.97E+08 | GTAACCTACTTGGCTGTGTCAG | [] |
| 1060 | 2 | MARS2 | {'exs': 5, 'nonexs': 3} | False | 1.98E+08 | CTGAGTTTCTAGGTCCGGTGG | [] |
| 1061 | 2 | BOLL | {'exs': 0, 'nonexs': 7} | True | 1.98E+08 | GCAGCTTCTAGCTGGTTCGTT | [] |
| 1062 | 2 | FTCDNL1 | unavailable | False | 2E+08 | TGGTCTAAAGCCATGCAGCCA | [] |
| 1063 | 2 | FTCDNL1 | unavailable | False | 2E+08 | CACTCCACACTATTCCACTGTG | [] |
| 1064 | 2 | CLK1 | {'exs': 2, 'nonexs': 5} | False | 2.01E+08 | AGATCTATATACTTTTCTTCAG | [] |
| 1065 | 2 | PPIL3 | {'exs': 0, 'nonexs': 7} | True | 2.01E+08 | TCATAGCTACTGAGCAAATGGG | [] |
| 1066 | 2 | FAM126B | {'exs': 0, 'nonexs': 7} | True | 2.01E+08 | AAAACTACACCTGGGATATTAG | [] |
| 1067 | 2 | CFLAR | {'exs': 2, 'nonexs': 6} | False | 2.01E+08 | GACATCTAGTAAGATGCATCCG | [] |
| 1068 | 2 | CFLAR | {'exs': 2, 'nonexs': 6} | False | 2.01E+08 | GACCATCCTATAGGACATCTAG | [] |
| 1069 | 2 | CASP10 | {'exs': 0, 'nonexs': 7} | True | 2.01E+08 | AGGAGCTACAGGTCATCTGGC | [] |
| 1070 | 2 | CASP10 | {'exs': 0, 'nonexs': 7} | True | 2.01E+08 | ACTCTCTGCTATAATGAAAGTG | [] |
| 1071 | 2 | CASP10 | {'exs': 0, 'nonexs': 7} | True | 2.01E+08 | TGGAAAGAACTAGGAAACGCTG | [] |
| 1072 | 2 | STRADB | {'exs': 1, 'nonexs': 6} | True | 2.01E+08 | CAGCCCTAGAATTCCCAGTATG | [] |
| 1073 | 2 | C2CD6 | unavailable | False | 2.02E+08 | CTACCTTTTAATTTTTAATCTG | [] |
| 1074 | 2 | ALS2 | {'exs': 0, 'nonexs': 7} | True | 2.02E+08 | CAGCCTAGTTAAGCTTCTCACG | [] |
| 1075 | 2 | KIAA2012 | unavailable | False | 2.02E+08 | TCACTCTATAAAAGCCTATACG | [] |
| 1076 | 2 | SUMO1 | {'exs': 0, 'nonexs': 7} | True | 2.02E+08 | AGAATATCTAAACTGTTGAATG | [] |
| 1077 | 2 | FAM117B | {'exs': 0, 'nonexs': 7} | True | 2.03E+08 | TGCACTGTGACCTATCCTTCAG | [] |
| 1078 | 2 | RAPH1 | {'exs': 0, 'nonexs': 7} | True | 2.03E+08 | TGGTGGCTACCAGTCTCTGGA | [] |
| 1079 | 2 | PARD3B | {'exs': 0, 'nonexs': 7} | True | 2.06E+08 | GCACTCAGCTATACGGCTGCA | [] |
| 1080 | 2 | ZDBF2 | {'exs': 0, 'nonexs': 7} | True | 2.06E+08 | CAACTTCTATTTTACCTCATTG | ['TCATTACCTCTCAGTTTGTTTG'] |
| 1081 | 2 | FZD5 | {'exs': 1, 'nonexs': 6} | True | 2.08E+08 | AGCCTCCTACACGTGCGACAG | [] |
| 1082 | 2 | C2orf80 | {'exs': 0, 'nonexs': 7} | True | 2.08E+08 | CAATTTTCTAAGTGACCTGCAG | [] |
| 1083 | 2 | PIKFYVE | {'exs': 2, 'nonexs': 5} | False | 2.08E+08 | GTGACTACCTCCTACCTTTTTG | [] |
| 1084 | 2 | UNC80 | {'exs': 0, 'nonexs': 5} | True | 2.1E+08 | CTTTAACTAACTGTGGGTAAGG | [] |
| 1085 | 2 | ACADL | {'exs': 0, 'nonexs': 7} | True | 2.1E+08 | GGCAGATGTCTACTTGTCAAAG | [] |
| 1086 | 2 | CPS1 | {'exs': 0, 'nonexs': 6} | True | 2.11E+08 | CATCTCTATGCTGCTTTTCCAG | [] |
| 1087 | 2 | IKZF2 | {'exs': 3, 'nonexs': 5} | False | 2.13E+08 | AAAAGGCCTAGTGGAATGTGTG | [] |
| 1088 | 2 | SPAG16 | {'exs': 0, 'nonexs': 7} | True | 2.13E+08 | TTTCCTATGAGAAATCAGCCTG | [] |
| 1089 | 2 | SPAG16 | {'exs': 0, 'nonexs': 7} | True | 2.14E+08 | AGATATCTAAAAGGAAGCACTG | [] |
| 1090 | 2 | VWC2L | {'exs': 0, 'nonexs': 7} | True | 2.15E+08 | TTTGTCCTACACAGTCTGCTTG | [] |
| 1091 | 2 | BARD1 | {'exs': 4, 'nonexs': 4} | False | 2.15E+08 | AAGAGCTATGATTGCACCACTG | [] |
| 1092 | 2 | MREG | {'exs': 0, 'nonexs': 7} | True | 2.16E+08 | TGCTTCTACTTAAAATGAAGAG | [] |
| 1093 | 2 | XRCC5 | {'exs': 5, 'nonexs': 1} | False | 2.16E+08 | CGACCTATATCATGTCCAACTG | [] |
| 1094 | 2 | RPL37A | {'exs': 6, 'nonexs': 1} | False | 2.17E+08 | GGAGCGTCTACTGGTCTTTCA | ['TTTCAACTCCTTCAGTCTTCTG'] |
| 1095 | 2 | IGFBP2 | {'exs': 0, 'nonexs': 7} | True | 2.17E+08 | TGCGGTCTACTGCATCCGCTG | [] |
| 1096 | 2 | SLC11A1 | {'exs': 1, 'nonexs': 7} | True | 2.18E+08 | CTGGTGTGGGCCTAGCCAGAG | [] |
| 1097 | 2 | CTDSP1 | {'exs': 1, 'nonexs': 7} | True | 2.18E+08 | CATATGGCTACAGGGCAGGTG | [] |
| 1098 | 2 | CTDSP1 | {'exs': 1, 'nonexs': 7} | True | 2.18E+08 | CCTCACTAGCTCCCTGGCCGT | [] |
| 1099 | 2 | RNF25 | {'exs': 0, 'nonexs': 7} | True | 2.19E+08 | TCCTGCTAGGAACCATCCTTAG | [] |
| 1100 | 2 | CDK5R2 | {'exs': 0, 'nonexs': 7} | True | 2.19E+08 | TGGGTATCCCTAGCGGTCCAG | [] |
| 1101 | 2 | FEV | {'exs': 0, 'nonexs': 7} | True | 2.19E+08 | CCGCCCCGTCTAGTGGTAATG | [] |
| 1102 | 2 | CRYBA2 | {'exs': 1, 'nonexs': 7} | True | 2.19E+08 | GAGCCTAGTGCTGGACTCTCC | [] |
| 1103 | 2 | CFAP65 | unavailable | False | 2.19E+08 | CCGAGCTAGGCCAGGAGCTTT | [] |
| 1104 | 2 | CNPPD1 | unavailable | False | 2.19E+08 | CCCTACCTAGCCTGGGAAAAC | [] |
| 1105 | 2 | ZFAND2B | {'exs': 0, 'nonexs': 7} | True | 2.19E+08 | AGGGCCCTAGCACAGGCTGCA | [] |
| 1106 | 2 | ATG9A | {'exs': 2, 'nonexs': 6} | False | 2.19E+08 | CCTTGTCTATACCTGTGGGGA | [] |
| 1107 | 2 | SPEG | {'exs': 0, 'nonexs': 7} | True | 2.19E+08 | GTGCCTCTAGGGGCCGCCAGG | [] |
| 1108 | 2 | ASIC4 | unavailable | False | 2.2E+08 | CTAGCAAGCAAAATCTTCAAAG | [] |
| 1109 | 2 | TMEM198 | {'exs': 0, 'nonexs': 7} | True | 2.2E+08 | ATATGGCTATACCCGCACTGG | [] |
| 1110 | 2 | OBSL1 | {'exs': 0, 'nonexs': 7} | True | 2.2E+08 | TGCTATCTAGAAGGCCTGTTTG | [] |
| 1111 | 2 | OBSL1 | {'exs': 0, 'nonexs': 7} | True | 2.2E+08 | CCGCGGGCTAGGCGTAAGCGC | [] |
| 1112 | 2 | OBSL1 | {'exs': 0, 'nonexs': 7} | True | 2.2E+08 | AGCTGCTACTCAGTGCCAGCT | [] |
| 1113 | 2 | PAX3 | {'exs': 0, 'nonexs': 7} | True | 2.22E+08 | TTGCCCCCTAAAAAGTCCAAGG | [] |
| 1114 | 2 | KCNE4 | {'exs': 0, 'nonexs': 5} | True | 2.23E+08 | GGGGTGCTAGGAATTCTGATG | [] |
| 1115 | 2 | MRPL44 | {'exs': 6, 'nonexs': 2} | False | 2.24E+08 | TGGCGGCTAGCTGGCAGTGAT | [] |
| 1116 | 2 | FAM124B | {'exs': 0, 'nonexs': 7} | True | 2.24E+08 | CTATATAAAGAATTCTTCTTCG | ['TTTCTTCACCCTGAACTTGCAG', 'TTCATCTTCTTCTTTTTCTTCG'] |
| 1117 | 2 | DOCK10 | {'exs': 2, 'nonexs': 7} | False | 2.25E+08 | TGCTTTGCTAATTACTCGAGTG | [] |
| 1118 | 2 | DOCK10 | {'exs': 2, 'nonexs': 7} | False | 2.25E+08 | ACATAGTCTAGAAAACCCAGGG | [] |
| 1119 | 2 | IRS1 | {'exs': 1, 'nonexs': 6} | True | 2.27E+08 | TTGAGCTACTGACGGTCCTCT | [] |
| 1120 | 2 | COL4A4 | {'exs': 0, 'nonexs': 7} | True | 2.27E+08 | TCTAATTGCAGGTTTTTCTCAG | [] |
| 1121 | 2 | COL4A4 | {'exs': 0, 'nonexs': 7} | True | 2.27E+08 | CATTCTCTAGCTATACTTCACG | [] |
| 1122 | 2 | MFF | {'exs': 2, 'nonexs': 6} | False | 2.27E+08 | TTACCTCTAGCGGCGAAACCA | [] |
| 1123 | 2 | TM4SF20 | {'exs': 0, 'nonexs': 7} | True | 2.27E+08 | TCCCATTAAACTACACAATTTG | [] |
| 1124 | 2 | AGFG1 | {'exs': 0, 'nonexs': 7} | True | 2.28E+08 | ATAAGGCTATAAGAAAGGATTG | [] |
| 1125 | 2 | C2orf83 | {'exs': 0, 'nonexs': 7} | True | 2.28E+08 | AGGTGGGAAGCTAGTTCCCCT | [] |
| 1126 | 2 | FBXO36 | {'exs': 2, 'nonexs': 7} | False | 2.3E+08 | CCTAAGGTTGCTTTTCTCTCAG | [] |
| 1127 | 2 | SLC16A14 | {'exs': 1, 'nonexs': 7} | True | 2.3E+08 | ATTACATGATACTAAACATGTG | [] |
| 1128 | 2 | SP140 | {'exs': 0, 'nonexs': 7} | True | 2.3E+08 | CAGTCCTACAGCTCCTTCGGA | [] |
| 1129 | 2 | SP100 | {'exs': 1, 'nonexs': 6} | True | 2.3E+08 | GCATCCTAATCTTCTTTACCTG | [] |
| 1130 | 2 | SP100 | {'exs': 1, 'nonexs': 6} | True | 2.31E+08 | TGGCTAAATAAACATTATAATG | [] |
| 1131 | 2 | CAB39 | {'exs': 5, 'nonexs': 4} | False | 2.31E+08 | TCTCTCTATTTGTATGCAGCAG | [] |
| 1132 | 2 | PSMD1 | {'exs': 7, 'nonexs': 1} | False | 2.31E+08 | CCCCTAGCCTGCTCTTTCACT | [] |
| 1133 | 2 | HTR2B | {'exs': 0, 'nonexs': 7} | True | 2.31E+08 | TCTGCTATACATAACTAACTTG | [] |
| 1134 | 2 | ARMC9 | {'exs': 0, 'nonexs': 7} | True | 2.31E+08 | CTTCGCTAGAGAAACACACACG | [] |
| 1135 | 2 | NCL | {'exs': 6, 'nonexs': 3} | False | 2.31E+08 | GACAGAAGCTATTCAAACTTCG | [] |
| 1136 | 2 | NMUR1 | {'exs': 0, 'nonexs': 7} | True | 2.32E+08 | AGATGCCTAACAGAAGGAAGGG | [] |
| 1137 | 2 | TEX44 | unavailable | False | 2.32E+08 | AGGGCTCTAATCATAGTTGGGG | [] |
| 1138 | 2 | PTMA | {'exs': 2, 'nonexs': 5} | False | 2.32E+08 | TGCTGTCTAGTCATCCTCGTC | [] |
| 1139 | 2 | PDE6D | {'exs': 0, 'nonexs': 7} | True | 2.32E+08 | TTCTATGATAACGTTCCCACTG | [] |
| 1140 | 2 | COPS7B | {'exs': 0, 'nonexs': 7} | True | 2.32E+08 | CCGGCCCTAGTGGCGGCTGG | [] |
| 1141 | 2 | NPPC | {'exs': 0, 'nonexs': 7} | True | 2.32E+08 | GCCGCACTAACATCCCAGGCC | [] |
| 1142 | 2 | CHRND | {'exs': 0, 'nonexs': 7} | True | 2.33E+08 | CCCACCCTAGATGAAGCGCTT | [] |
| 1143 | 2 | EFHD1 | {'exs': 0, 'nonexs': 7} | True | 2.33E+08 | GCAGGACTATGTATTGAAGTTG | [] |
| 1144 | 2 | SAG | {'exs': 0, 'nonexs': 7} | True | 2.33E+08 | CAAAAACTAAATTTGCATCCTG | [] |
| 1145 | 2 | DGKD | {'exs': 1, 'nonexs': 6} | True | 2.33E+08 | CACTTTCTAAGTAGTGCTGGTG | [] |
| 1146 | 2 | DGKD | {'exs': 1, 'nonexs': 6} | True | 2.33E+08 | CAGAGGCTAGGCCTCGACGGC | [] |
| 1147 | 2 | DGKD | {'exs': 1, 'nonexs': 6} | True | 2.33E+08 | CGGAGGCTAGGCCTCGGCGGC | [] |
| 1148 | 2 | MROH2A | unavailable | False | 2.34E+08 | GACCACCTAGGACATTCCTTG | [] |
| 1149 | 2 | HJURP | {'exs': 5, 'nonexs': 3} | False | 2.34E+08 | TAACCTAGCTACACACTTTTAG | [] |
| 1150 | 2 | AGAP1 | {'exs': 1, 'nonexs': 6} | True | 2.36E+08 | TCCCACCGCTAATTCATCATGG | [] |
| 1151 | 2 | AGAP1 | {'exs': 1, 'nonexs': 6} | True | 2.36E+08 | AGGCCTAGATGCATGGTTTTTG | [] |
| 1152 | 2 | MLPH | {'exs': 0, 'nonexs': 7} | True | 2.38E+08 | GCGGACTAAGGATTTCCACATG | [] |
| 1153 | 2 | RAB17 | {'exs': 0, 'nonexs': 7} | True | 2.38E+08 | CACCTAGTGGGCGCAGCATTT | [] |
| 1154 | 2 | RBM44 | {'exs': 0, 'nonexs': 5} | True | 2.38E+08 | GAATTCCTAACTGGAAGCAGAG | [] |
| 1155 | 2 | RAMP1 | {'exs': 0, 'nonexs': 7} | True | 2.38E+08 | CCCCGCTACTCACCCACTGCA | [] |
| 1156 | 2 | RAMP1 | {'exs': 0, 'nonexs': 7} | True | 2.38E+08 | CCCGCCTACACAATGCCCTCA | [] |
| 1157 | 2 | SCLY | {'exs': 0, 'nonexs': 5} | True | 2.38E+08 | AGTGCTAGGCCTGGTCCTCCA | [] |
| 1158 | 2 | ASB1 | {'exs': 0, 'nonexs': 7} | True | 2.38E+08 | TTGGAGTCTACTCATGGAGTAG | [] |
| 1159 | 2 | TWIST2 | {'exs': 1, 'nonexs': 7} | True | 2.39E+08 | GCGCGGCGCTAGTGGGAGGC | [] |
| 1160 | 2 | HDAC4 | {'exs': 1, 'nonexs': 6} | True | 2.39E+08 | CGAGGGAGTGCTACAGGGGCG | [] |
| 1161 | 2 | NDUFA10 | {'exs': 3, 'nonexs': 5} | False | 2.4E+08 | GTTGTCCTAGCAACTATCTTGG | [] |
| 1162 | 2 | COPS9 | unavailable | False | 2.4E+08 | TGCATCCTACCTCACATCTATG | [] |
| 1163 | 2 | ANKMY1 | {'exs': 0, 'nonexs': 7} | True | 2.41E+08 | GGACCTACTCGATCATGGACA | [] |
| 1164 | 2 | RNPEPL1 | {'exs': 0, 'nonexs': 3} | True | 2.41E+08 | ACAGGGCTAGGCAGACACATT | [] |
| 1165 | 2 | MTERF4 | unavailable | False | 2.41E+08 | CAGGTCTATGTTTTGTTACGTG | [] |
| 1166 | 2 | MTERF4 | unavailable | False | 2.41E+08 | TAGGTTCTAGGGAGGGTGGCA | [] |
| 1167 | 2 | MTERF4 | unavailable | False | 2.41E+08 | TCACAGCTATTCCTCCTCGTC | [] |
| 1168 | 2 | ANO7 | {'exs': 0, 'nonexs': 7} | True | 2.41E+08 | CTCCCCTAAGGACCACCAGAT | [] |
| 1169 | 2 | FARP2 | {'exs': 0, 'nonexs': 7} | True | 2.41E+08 | AAGGGTCTATGTGCCCATGCA | [] |
| 1170 | 2 | FARP2 | {'exs': 0, 'nonexs': 7} | True | 2.41E+08 | AATGTCCTAACTCTGAATGTTG | [] |
| 1171 | 2 | FARP2 | {'exs': 0, 'nonexs': 7} | True | 2.41E+08 | AAAAGTCTACTCCATTTTTGTG | [] |
| 1172 | 2 | STK25 | {'exs': 1, 'nonexs': 6} | True | 2.41E+08 | TTCTCTATAGAAGGATCATCTG | [] |
| 1173 | 2 | ATG4B | {'exs': 0, 'nonexs': 7} | True | 2.42E+08 | GCTCTACATCAGAAGAATCTAG | [] |
| 1174 | 2 | ING5 | {'exs': 2, 'nonexs': 6} | False | 2.42E+08 | TTATGCCTAGCATACCCTTGAG | [] |
| 1175 | 2 | ING5 | {'exs': 2, 'nonexs': 6} | False | 2.42E+08 | -1 | -1 |
| 1176 | 2 | D2HGDH | {'exs': 0, 'nonexs': 7} | True | 2.42E+08 | CAGCTACTAGGGAGGCAGGAG | [] |
| 1177 | 2 | D2HGDH | {'exs': 0, 'nonexs': 7} | True | 2.42E+08 | CCAAATCCTAGGGCTCAGACA | [] |
| 1178 | 2 | GAL3ST2 | {'exs': 0, 'nonexs': 7} | True | 2.42E+08 | CCCCTCTACGCCCCCAGGAAC | [] |
| 1179 | 3 | IL5RA | {'exs': 1, 'nonexs': 7} | True | 3092215 | AGCTACTTACCCACATAAATAG | [] |
| 1180 | 3 | ITPR1 | {'exs': 0, 'nonexs': 7} | True | 4737285 | AGGGCACTAAATGTCATCCTTG | [] |
| 1181 | 3 | GRM7 | {'exs': 0, 'nonexs': 7} | True | 7686426 | CAAAAGCTATACTGTTGGTGGG | [] |
| 1182 | 3 | SRGAP3 | {'exs': 0, 'nonexs': 7} | True | 8992901 | CGCATATCCTACCGGCCCATC | ['GCCCATCACCCCCCCAAAGCCG'] |
| 1183 | 3 | THUMPD3 | {'exs': 0, 'nonexs': 7} | True | 9384731 | TTTATGCTAACATTTCCAGTGG | [] |
| 1184 | 3 | SETD5 | {'exs': 0, 'nonexs': 5} | True | 9469796 | AGACCACTAATGCTCATCACTG | [] |
| 1185 | 3 | SETD5 | {'exs': 0, 'nonexs': 5} | True | 9476090 | AGCCCTAGGAAAGTCCCGTCT | [] |
| 1186 | 3 | CPNE9 | {'exs': 0, 'nonexs': 7} | True | 9727381 | CCCGACCTACACCACGGATGT | [] |
| 1187 | 3 | OGG1 | {'exs': 0, 'nonexs': 7} | True | 9756842 | GAAACCTAGGGAGGACACCTA | [] |
| 1188 | 3 | OGG1 | {'exs': 0, 'nonexs': 7} | True | 9757149 | CATCTAGCCTTCCGGCCCTTT | [] |
| 1189 | 3 | OGG1 | {'exs': 0, 'nonexs': 7} | True | 9781578 | AGGCTGCCTAGACCTTAGGGG | [] |
| 1190 | 3 | OGG1 | {'exs': 0, 'nonexs': 7} | True | 9781681 | CAATCACTAAGTGACCTTCCAG | [] |
| 1191 | 3 | CAMK1 | {'exs': 2, 'nonexs': 6} | False | 9757538 | CAGGGCCCTAGAGCTGGTGGG | [] |
| 1192 | 3 | TADA3 | {'exs': 4, 'nonexs': 4} | False | 9780356 | GTGAGGGCTACCCATCCAGCA | [] |
| 1193 | 3 | RPUSD3 | {'exs': 2, 'nonexs': 6} | False | 9838015 | GGGAGGACTATTGTAAGCGGA | [] |
| 1194 | 3 | EMC3 | unavailable | False | 9969586 | TGTAAACTATGTTTTTACTAAG | [] |
| 1195 | 3 | FANCD2 | {'exs': 4, 'nonexs': 5} | False | 10101261 | TGGGGTCTAATCAGAGTCATCG | ['CATAACTCTCATCACTATCTTG'] |
| 1196 | 3 | BRK1 | {'exs': 1, 'nonexs': 4} | False | 10126294 | ACGGCACTGTTCTAGGTGAGT | [] |
| 1197 | 3 | VHL | {'exs': 4, 'nonexs': 5} | False | 10143003 | GGCTAGTTCCTACTCCATCCT | [] |
| 1198 | 3 | ATP2B2 | {'exs': 0, 'nonexs': 7} | True | 10328813 | TCAGCTAAAGCGACGTCTCCA | [] |
| 1199 | 3 | ATP2B2 | {'exs': 0, 'nonexs': 7} | True | 10329099 | AGAGCTACGGAACGCCTTCAC | [] |
| 1200 | 3 | ATP2B2 | {'exs': 0, 'nonexs': 7} | True | 10332029 | AGGGTTCTCTAGCCCTCTAGG | [] |
| 1201 | 3 | SLC6A1 | {'exs': 1, 'nonexs': 7} | True | 11036965 | ACCCTAGATGTAGGCCTCCTT | [] |
| 1202 | 3 | ATG7 | {'exs': 1, 'nonexs': 7} | True | 11564284 | GAAGCCTATGATTGGTACATAG | [] |
| 1203 | 3 | TAMM41 | unavailable | False | 11817170 | ACAAGCTATTTTCCACATACAG | [] |
| 1204 | 3 | SYN2 | {'exs': 0, 'nonexs': 1} | False | 12190524 | GAAGAGCTAATCTGAAAAGAGG | [] |
| 1205 | 3 | TIMP4 | {'exs': 0, 'nonexs': 7} | True | 12153514 | TCCCTACTAGGGCTGAACGAT | [] |
| 1206 | 3 | PPARG | {'exs': 2, 'nonexs': 5} | False | 12434144 | TCTCTGCTAGTACAAGTCCTTG | [] |
| 1207 | 3 | RAF1 | {'exs': 5, 'nonexs': 3} | False | 12584513 | AAGTCAACTAGAAGACAGGCAG | [] |
| 1208 | 3 | CAND2 | {'exs': 2, 'nonexs': 6} | False | 12833981 | GGGGACTAGCTGAGCTCCATT | [] |
| 1209 | 3 | RPL32 | {'exs': 7, 'nonexs': 0} | False | 12836093 | TGCCTACTCATTTTCTTCACTG | [] |
| 1210 | 3 | IQSEC1 | {'exs': 0, 'nonexs': 7} | True | 12900982 | GGCTGTCTACACAATTGTGCTG | [] |
| 1211 | 3 | NUP210 | {'exs': 0, 'nonexs': 7} | True | 13317680 | CGCGGCCTAGTGGGAGGCATA | [] |
| 1212 | 3 | NUP210 | {'exs': 0, 'nonexs': 7} | True | 13347704 | CTTGTCTATCTCTGTGCTGCTG | [] |
| 1213 | 3 | C3orf20 | {'exs': 0, 'nonexs': 7} | True | 14772874 | GGCGCTACTTCTTGGAGGCCT | [] |
| 1214 | 3 | FGD5 | {'exs': 1, 'nonexs': 6} | True | 14933166 | GATAACTGCTATAACACACTCG | [] |
| 1215 | 3 | NR2C2 | {'exs': 1, 'nonexs': 6} | True | 15043007 | TTTGCGCTATAGACTGGCTCC | [] |
| 1216 | 3 | METTL6 | {'exs': 0, 'nonexs': 7} | True | 15414808 | CCCAACTACTTCAGAGGCTGA | [] |
| 1217 | 3 | METTL6 | {'exs': 0, 'nonexs': 7} | True | 15415633 | TTTCTATACACTCACTACACAG | [] |
| 1218 | 3 | EAF1 | {'exs': 3, 'nonexs': 6} | False | 15439154 | GCACTAGTCATCACTGTCACTG | [] |
| 1219 | 3 | BTD | {'exs': 0, 'nonexs': 7} | True | 15645487 | TTCCTAGTCCCTCTCATACAAG | [] |
| 1220 | 3 | ANKRD28 | {'exs': 0, 'nonexs': 7} | True | 15781622 | TGTTCGGCAGGCTAATGTTTAG | [] |
| 1221 | 3 | GALNT15 | unavailable | False | 16231872 | CCTACCCATTGTTCACAATCTG | [] |
| 1222 | 3 | OXNAD1 | {'exs': 0, 'nonexs': 6} | True | 16303561 | CTCCTACCACCACTTCTCAAAG | [] |
| 1223 | 3 | EFHB | {'exs': 0, 'nonexs': 7} | True | 19879677 | CTCATCTAGAACATTTCTGATG | [] |
| 1224 | 3 | ZNF385D | {'exs': 0, 'nonexs': 7} | True | 21459363 | AAGGGCACTATCTGGAATATGG | [] |
| 1225 | 3 | UBE2E2 | {'exs': 0, 'nonexs': 7} | True | 23345869 | GCTAACCTTAAATACAAATTTG | [] |
| 1226 | 3 | UBE2E2 | {'exs': 0, 'nonexs': 7} | True | 23589830 | CAGGCCCCTATGTGGCGTACC | [] |
| 1227 | 3 | THRB | {'exs': 0, 'nonexs': 7} | True | 24122883 | CTAATCCTCGAACACTTCCAAG | [] |
| 1228 | 3 | NGLY1 | {'exs': 2, 'nonexs': 7} | False | 25722734 | CTACTGCATGCTCCTCTTACAG | [] |
| 1229 | 3 | OXSM | {'exs': 2, 'nonexs': 6} | False | 25794493 | TCTACAGTCCAGCAATACAAAG | [] |
| 1230 | 3 | LRRC3B | {'exs': 0, 'nonexs': 7} | True | 26710651 | GTTTGGACACTATACCACAGTG | [] |
| 1231 | 3 | NEK10 | {'exs': 0, 'nonexs': 7} | True | 27114279 | TTTTCCTAATGCATATACTCAG | [] |
| 1232 | 3 | NEK10 | {'exs': 0, 'nonexs': 7} | True | 27171468 | AAACCTATACCCTGGGGATATG | [] |
| 1233 | 3 | SLC4A7 | {'exs': 4, 'nonexs': 4} | False | 27376763 | CAATTCTATAATGAAGTTTCAG | [] |
| 1234 | 3 | CMC1 | {'exs': 0, 'nonexs': 7} | True | 28319628 | TATCTGCCTACATGCTTGTTGG | [] |
| 1235 | 3 | ZCWPW2 | {'exs': 0, 'nonexs': 7} | True | 28524687 | TTCTAAAACTCAGACATCAAAG | [] |
| 1236 | 3 | TGFBR2 | {'exs': 2, 'nonexs': 6} | False | 30691598 | AGAGCTATTTGGTAGTGTTTAG | [] |
| 1237 | 3 | GADL1 | {'exs': 0, 'nonexs': 7} | True | 30728241 | ACAGCTACATGTCTTTACCCAG | [] |
| 1238 | 3 | OSBPL10 | {'exs': 0, 'nonexs': 7} | True | 31664018 | CCATCTACCCTCTCAGGACAA | [] |
| 1239 | 3 | ZNF860 | {'exs': 0, 'nonexs': 7} | True | 31990977 | TGACCTCTAACTGAAGACCTTG | [] |
| 1240 | 3 | CMTM7 | {'exs': 0, 'nonexs': 7} | True | 32449659 | TATTGATTCTATGAAGGGAATG | [] |
| 1241 | 3 | CMTM7 | {'exs': 0, 'nonexs': 7} | True | 32452088 | GGGGCTAAGCCAATGGACTTT | [] |
| 1242 | 3 | CCR4 | {'exs': 0, 'nonexs': 7} | True | 32954504 | ATTTTTCTACAGAGCATCATGG | [] |
| 1243 | 3 | CRTAP | {'exs': 0, 'nonexs': 7} | True | 33143447 | TGGGCTAGCTGGTCTCCTCCA | [] |
| 1244 | 3 | SUSD5 | {'exs': 0, 'nonexs': 7} | True | 33152741 | TGCCTAGACCTTCTCCATCTC | [] |
| 1245 | 3 | CLASP2 | {'exs': 2, 'nonexs': 1} | False | 33498630 | GAGCTTCACTAACTTTGTCCAG | [] |
| 1246 | 3 | DCLK3 | {'exs': 2, 'nonexs': 6} | False | 36715327 | AAGGTGGTGACTATGATACCTG | [] |
| 1247 | 3 | LRRFIP2 | {'exs': 1, 'nonexs': 6} | True | 37053850 | GGTTTTCCTACTGCTGGGCCA | [] |
| 1248 | 3 | C3orf35 | {'exs': 0, 'nonexs': 3} | True | 37435129 | CATTTCTATCCCAGGAACCAAG | [] |
| 1249 | 3 | CTDSPL | {'exs': 0, 'nonexs': 7} | True | 37980866 | GCCAGGGCTACCTATTGCAGA | [] |
| 1250 | 3 | PLCD1 | {'exs': 0, 'nonexs': 7} | True | 38007772 | TCCAGCCTAGTCCTGGAGGGA | [] |
| 1251 | 3 | DLEC1 | {'exs': 0, 'nonexs': 7} | True | 38117206 | CCACCTACTGAGCCCAAGTCT | [] |
| 1252 | 3 | DLEC1 | {'exs': 0, 'nonexs': 7} | True | 38119583 | CCGGCCTAGGTGACACAGCAA | [] |
| 1253 | 3 | EXOG | {'exs': 1, 'nonexs': 6} | True | 38524361 | ACTAGGATGGCTTTCTTATCTG | [] |
| 1254 | 3 | SCN10A | {'exs': 0, 'nonexs': 7} | True | 38697348 | TGTTCTCACTAGGGCCCAGGG | [] |
| 1255 | 3 | WDR48 | {'exs': 2, 'nonexs': 6} | False | 39094251 | TTGTCTATGTATCCCTTCTCTG | [] |
| 1256 | 3 | TTC21A | {'exs': 0, 'nonexs': 7} | True | 39131610 | GCCTCTAGAACTGTAGAAAATG | [] |
| 1257 | 3 | TTC21A | {'exs': 0, 'nonexs': 7} | True | 39138787 | GACCCCAGCTAGGGCCTCAGG | [] |
| 1258 | 3 | ENTPD3 | {'exs': 0, 'nonexs': 7} | True | 40424771 | TATCAGCTATTCCTGAAATGAG | [] |
| 1259 | 3 | ZNF620 | {'exs': 0, 'nonexs': 7} | True | 40516862 | TAGCCCTATGCTTGGACAGGT | [] |
| 1260 | 3 | ULK4 | {'exs': 0, 'nonexs': 7} | True | 41249928 | TTGCCTAGTGCCCAACGGCTT | [] |
| 1261 | 3 | ULK4 | {'exs': 0, 'nonexs': 7} | True | 41391663 | GCTATATTACACTCATTTTCAG | [] |
| 1262 | 3 | TRAK1 | {'exs': 0, 'nonexs': 7} | True | 42217150 | CTGATTCCTAAAGCCCTGGGT | [] |
| 1263 | 3 | CCK | {'exs': 0, 'nonexs': 7} | True | 42258097 | GGTCCTCTAGGAGGGGTACTC | ['GTACTCATACTCCTCGGCACTG'] |
| 1264 | 3 | LYZL4 | {'exs': 0, 'nonexs': 7} | True | 42397264 | GCTACAGCTTGCAACCATCCA | [] |
| 1265 | 3 | ZBTB47 | {'exs': 1, 'nonexs': 6} | True | 42664597 | GCTCGGCAGCTAGTTGTTGGC | [] |
| 1266 | 3 | HHATL | {'exs': 1, 'nonexs': 7} | True | 42692750 | CTCCTACTCCGGCTTCTCTTT | [] |
| 1267 | 3 | CCDC13 | {'exs': 0, 'nonexs': 7} | True | 42708979 | CTGTCATCCTATTGCTTGCCTG | [] |
| 1268 | 3 | CCDC13 | {'exs': 0, 'nonexs': 7} | True | 42735388 | TGGAGCTACTGCCACCATCTT | [] |
| 1269 | 3 | HIGD1A | {'exs': 0, 'nonexs': 7} | True | 42785270 | TTCTTCTAAGGCTTAGGTTTTG | [] |
| 1270 | 3 | ZNF662 | {'exs': 1, 'nonexs': 7} | True | 42915353 | CACTCTCTAATGTTCAAGAAGG | [] |
| 1271 | 3 | POMGNT2 | unavailable | False | 43079688 | CTCGCTACGTGTTGCACACCA | [] |
| 1272 | 3 | ANO10 | {'exs': 0, 'nonexs': 7} | True | 43372760 | TGACTAGACCCCAGGACCTAT | [] |
| 1273 | 3 | ANO10 | {'exs': 0, 'nonexs': 7} | True | 43457457 | ATTATGCTAAATAAAAAGCCAG | [] |
| 1274 | 3 | TOPAZ1 | unavailable | False | 44332010 | TTAGCTAATGGTTACTGAAAAG | [] |
| 1275 | 3 | ZNF445 | {'exs': 1, 'nonexs': 5} | False | 44449574 | CCCAGGCTAATCTCTAATATGG | [] |
| 1276 | 3 | ZNF197 | {'exs': 0, 'nonexs': 7} | True | 44644219 | ACATATGACACTAGATTTTCTG | [] |
| 1277 | 3 | ZNF660-ZNF197 | unavailable | False | 44644219 | ACATATGACACTAGATTTTCTG | [] |
| 1278 | 3 | TMEM42 | {'exs': 1, 'nonexs': 7} | True | 44865179 | TGGTGCTACTGCTGCTTGTGT | [] |
| 1279 | 3 | TGM4 | {'exs': 0, 'nonexs': 7} | True | 44913724 | GACAAGGCTACTTGGTGATGAG | [] |
| 1280 | 3 | ZDHHC3 | {'exs': 0, 'nonexs': 7} | True | 44917894 | CACAGCAGCATCTATTCATCTG | [] |
| 1281 | 3 | CLEC3B | {'exs': 1, 'nonexs': 7} | True | 45035923 | CCGGCTACACGATCCCGAACT | [] |
| 1282 | 3 | LIMD1 | {'exs': 0, 'nonexs': 7} | True | 45677058 | CTCTGGCTAGAAGTGGTGCTG | [] |
| 1283 | 3 | FYCO1 | {'exs': 0, 'nonexs': 6} | True | 45921764 | CTACAGGAAATCACTTCATCG | [] |
| 1284 | 3 | CXCR6 | {'exs': 0, 'nonexs': 7} | True | 45947509 | CAAGGCCTATAACTGGAACATG | [] |
| 1285 | 3 | CCR3 | {'exs': 0, 'nonexs': 6} | True | 46266225 | TCTGACCTAAAACACAATAGAG | [] |
| 1286 | 3 | CCR2 | {'exs': 0, 'nonexs': 7} | True | 46359859 | CTGTCTCTAGGCTCCTTCTTTG | [] |
| 1287 | 3 | LRRC2 | {'exs': 0, 'nonexs': 7} | True | 46529900 | GTAGCTACCTGTCTATATCTTG | [] |
| 1288 | 3 | ALS2CL | {'exs': 0, 'nonexs': 7} | True | 46670983 | GCCAGGCTACCAGAGCTCCCT | [] |
| 1289 | 3 | ALS2CL | {'exs': 0, 'nonexs': 7} | True | 46675900 | CTTCTATTTGCAGTGTTTCATG | [] |
| 1290 | 3 | KIF9 | {'exs': 1, 'nonexs': 7} | True | 47228651 | GGTTCTATTTTCTATGTGCCTG | [] |
| 1291 | 3 | SMARCC1 | {'exs': 3, 'nonexs': 4} | False | 47588208 | TCCAGGCTAAGGAGCAGCTGA | [] |
| 1292 | 3 | MAP4 | {'exs': 1, 'nonexs': 7} | True | 47852791 | ACTGGTCTAGTGGACAGCCCG | [] |
| 1293 | 3 | MAP4 | {'exs': 1, 'nonexs': 7} | True | 47975457 | GACACGCTAGGCTTCTAGGAG | [] |
| 1294 | 3 | CAMP | {'exs': 1, 'nonexs': 5} | False | 48225423 | CACACACTAGGACTCTGTCCT | [] |
| 1295 | 3 | NME6 | {'exs': 1, 'nonexs': 7} | True | 48294665 | TCCTGTTCCAGCTACATAGTGG | [] |
| 1296 | 3 | PLXNB1 | {'exs': 0, 'nonexs': 7} | True | 48404485 | TTCCTATAGATCTGTGACCTTG | [] |
| 1297 | 3 | TREX1 | {'exs': 0, 'nonexs': 5} | True | 48467599 | CTTGGCCTACTCCCCAGGTGT | [] |
| 1298 | 3 | SHISA5 | {'exs': 0, 'nonexs': 7} | True | 48469337 | GCAGGGCTAGAGTTGGCAGGG | [] |
| 1299 | 3 | UQCRC1 | {'exs': 7, 'nonexs': 1} | False | 48599127 | TCCCGCCTAGAAGCGCAGCCA | [] |
| 1300 | 3 | NCKIPSD | {'exs': 1, 'nonexs': 7} | True | 48674543 | GCTAGCTGGGAGCCTCCCCCA | [] |
| 1301 | 3 | PRKAR2A | {'exs': 0, 'nonexs': 7} | True | 48751584 | GCACACCTACTGCCCGAGGTT | [] |
| 1302 | 3 | ARIH2 | {'exs': 0, 'nonexs': 7} | True | 48983222 | CTGCTCCGCTATATGCATCTG | [] |
| 1303 | 3 | USP19 | {'exs': 0, 'nonexs': 7} | True | 49108411 | CTGCTAATCCACCTCCTCCAT | [] |
| 1304 | 3 | GPX1 | {'exs': 0, 'nonexs': 7} | True | 49357387 | GGGCGCCCTAGGCACAGCTGG | [] |
| 1305 | 3 | GPX1 | {'exs': 0, 'nonexs': 7} | True | 49357825 | AGATTTTCTATGAGTCACCGGG | [] |
| 1306 | 3 | GPX1 | {'exs': 0, 'nonexs': 7} | True | 49357981 | CCCAGCCACTACTGCACGTCC | [] |
| 1307 | 3 | MST1 | {'exs': 1, 'nonexs': 6} | True | 49684027 | CTGGGCCTAACCCAGTCTCAT | [] |
| 1308 | 3 | RNF123 | {'exs': 0, 'nonexs': 7} | True | 49721304 | TGAGGGCTAGGCAGCTGAGGA | [] |
| 1309 | 3 | AMIGO3 | {'exs': 0, 'nonexs': 5} | True | 49717950 | GGGCAGTCTAGGTTGTCATGG | [] |
| 1310 | 3 | IP6K1 | {'exs': 0, 'nonexs': 7} | True | 49727121 | GGGCCTACTGGTTCTCGTCCC | [] |
| 1311 | 3 | CDHR4 | {'exs': 0, 'nonexs': 7} | True | 49793401 | GTAGCCTAGGCATCAAGTGATG | [] |
| 1312 | 3 | RBM6 | {'exs': 1, 'nonexs': 7} | True | 50060881 | GTTCCTATCGAGAAATCCTCTG | [] |
| 1313 | 3 | SLC38A3 | unavailable | False | 50220176 | CACCCTAGTGGTTTCCTCCAT | [] |
| 1314 | 3 | HYAL2 | {'exs': 0, 'nonexs': 7} | True | 50318128 | GGAGACCCCTACAAGGTCCAG | [] |
| 1315 | 3 | HEMK1 | {'exs': 2, 'nonexs': 6} | False | 50580416 | CATGCTATGGCCCAGACCTCC | [] |
| 1316 | 3 | HEMK1 | {'exs': 2, 'nonexs': 6} | False | 50582092 | GGGCAGCTACCCTGAGATGCT | [] |
| 1317 | 3 | MAPKAPK3 | {'exs': 1, 'nonexs': 6} | True | 50649045 | CATGAGCTACTGGTTGTTGCAG | [] |
| 1318 | 3 | MANF | {'exs': 0, 'nonexs': 1} | False | 51389088 | GCAGACTACAAATCGGTCCGT | [] |
| 1319 | 3 | RBM15B | {'exs': 0, 'nonexs': 4} | True | 51394071 | CTTGGGCTAGGCAGTGTCTCT | [] |
| 1320 | 3 | RAD54L2 | {'exs': 2, 'nonexs': 7} | False | 51663419 | GCTCCCTAGCTATTTCCCAGT | [] |
| 1321 | 3 | IQCF6 | {'exs': 0, 'nonexs': 5} | True | 51778622 | GCCCTAGGTCATGAGGATTTC | [] |
| 1322 | 3 | RPL29 | {'exs': 5, 'nonexs': 3} | False | 51993748 | GATATCTACTCTGAAGCCTTTG | [] |
| 1323 | 3 | POC1A | {'exs': 1, 'nonexs': 6} | True | 52123578 | ATGGATGGCGTCTAGGGTCGT | [] |
| 1324 | 3 | ALAS1 | {'exs': 2, 'nonexs': 6} | False | 52214030 | TGTGACTAGCAGATTCTCTGAG | [] |
| 1325 | 3 | TLR9 | {'exs': 0, 'nonexs': 7} | True | 52221216 | CTCACGGCTATTCGGCCGTGG | [] |
| 1326 | 3 | TWF2 | {'exs': 0, 'nonexs': 7} | True | 52229033 | AGCCTCCTAGCTGTCATCCCC | ['ATCCCCATTTTCACCCGGGCCG'] |
| 1327 | 3 | DNAH1 | {'exs': 0, 'nonexs': 7} | True | 52392236 | TGGAATCCTAGAAGGGGCTGA | [] |
| 1328 | 3 | DNAH1 | {'exs': 0, 'nonexs': 7} | True | 52400445 | GTCTGAGTCTAGTAGTCCAGG | [] |
| 1329 | 3 | SEMA3G | {'exs': 0, 'nonexs': 7} | True | 52435602 | TCTACGTGGCCTCCACCTCCC | [] |
| 1330 | 3 | TNNC1 | {'exs': 0, 'nonexs': 7} | True | 52451274 | CATCTACTCCACACCCTTCATG | [] |
| 1331 | 3 | NISCH | {'exs': 0, 'nonexs': 5} | True | 52492481 | CCTGGGCTAGCCGGTGAGCTC | [] |
| 1332 | 3 | STAB1 | {'exs': 0, 'nonexs': 7} | True | 52519896 | AGAGACTAGTTAGCAAAGGCAG | [] |
| 1333 | 3 | GLT8D1 | {'exs': 0, 'nonexs': 7} | True | 52696147 | AAACCACTACAGCCCACTGTAG | [] |
| 1334 | 3 | ITIH4 | {'exs': 0, 'nonexs': 7} | True | 52813420 | CATCAGAACTACAGCTCCACAG | [] |
| 1335 | 3 | TKT | {'exs': 6, 'nonexs': 2} | False | 53225755 | CCCGCCCTAGGCCTTGGTGAT | [] |
| 1336 | 3 | CACNA1D | {'exs': 0, 'nonexs': 7} | True | 53811405 | TGGGGGCTACAAGGTGGTGAT | [] |
| 1337 | 3 | IL17RB | {'exs': 0, 'nonexs': 7} | True | 53865307 | GGTGGGCTACAAGGAGCAGCA | [] |
| 1338 | 3 | WNT5A | {'exs': 1, 'nonexs': 4} | False | 55470091 | CCCACTACTTGCACACAAACTG | [] |
| 1339 | 3 | ERC2 | {'exs': 0, 'nonexs': 7} | True | 55683822 | CAGGCCCGGCTATGCCCATAT | [] |
| 1340 | 3 | ERC2 | {'exs': 0, 'nonexs': 7} | True | 55943493 | TTGGTCTAATCTTCACAGGAAG | [] |
| 1341 | 3 | HESX1 | {'exs': 1, 'nonexs': 7} | True | 57198196 | TCTATCTATTCCAGCAGATTTG | [] |
| 1342 | 3 | PDE12 | {'exs': 1, 'nonexs': 7} | True | 57560003 | ACACATCTATTTCCATTTTAAG | ['CATTTTAAATCACATACAAGTG'] |
| 1343 | 3 | PDE12 | {'exs': 1, 'nonexs': 7} | True | 57654792 | AAGAGACCTAGAAAGGCATTTG | [] |
| 1344 | 3 | SLMAP | {'exs': 3, 'nonexs': 5} | False | 57917154 | CCTTTATTTCTAACTAGCATGG | [] |

| | | | | | | | |
|---|---|---|---|---|---|---|---|
| 1345 | 3 | SLMAP | {'exs': 3, 'nonexs': 5} | False | 57925927 | CTTTCTCTACCACAATGGACCG | [] |
| 1346 | 3 | LOC10537 | unavailable | False | 58180501 | TACTCCTAGGATGACATGACTG | [] |
| 1347 | 3 | DNASE1L3 | {'exs': 0, 'nonexs': 7} | True | 58192586 | CCCTTGGGTCTAGGAGCGTTT | [] |
| 1348 | 3 | PDHB | {'exs': 1, 'nonexs': 7} | True | 58428033 | AGTCCAAACTAAATATTTAATG | [] |
| 1349 | 3 | FAM107A | {'exs': 0, 'nonexs': 7} | True | 58566587 | GCCCTACAGCTCTCTCTCTTC | [] |
| 1350 | 3 | FAM3D | {'exs': 0, 'nonexs': 7} | True | 58634278 | GCCACCCTAAAATGGCTTCGG | [] |
| 1351 | 3 | CADPS | {'exs': 0, 'nonexs': 7} | True | 62399405 | AATGGTCTAATCGTCTTCTTCG | [] |
| 1352 | 3 | THOC7 | {'exs': 8, 'nonexs': 1} | False | 63834131 | CCTGTCTATGGCTTAGGATCTG | [] |
| 1353 | 3 | PSMD6 | {'exs': 8, 'nonexs': 1} | False | 64010908 | CTTTATCTATTTTGCAGTGTAG | [] |
| 1354 | 3 | ADAMTS9 | {'exs': 0, 'nonexs': 7} | True | 64522170 | CTTACCTTAGCTATAAAACTCG | [] |
| 1355 | 3 | LRIG1 | {'exs': 1, 'nonexs': 7} | True | 66380262 | GACAAAACCTAGCTTTTTGGTG | [] |
| 1356 | 3 | TAFA4 | unavailable | False | 68733141 | TCTTCGCTACCGCGTTACCTAC | ['ACCTAAAACAAATCAAAATAAG'] |
| 1357 | 3 | FRMD4B | {'exs': 0, 'nonexs': 7} | True | 69171865 | CAGACTAATGTTCCAGGCTTTG | [] |
| 1358 | 3 | MITF | {'exs': 0, 'nonexs': 7} | True | 69965247 | ATTCGCTAACAAGTGTGCTCCG | [] |
| 1359 | 3 | MDFIC2 | unavailable | False | 70196925 | TTCACTGTGCTAGCGGTAACAG | [] |
| 1360 | 3 | EBLN2 | unavailable | False | 73062899 | TTGGCAACTAACTTGACTTGCG | [] |
| 1361 | 3 | ZNF717 | {'exs': 1, 'nonexs': 1} | False | 75730449 | CACAGCTAGTTTGTTACTGGTG | [] |
| 1362 | 3 | CHMP2B | {'exs': 1, 'nonexs': 7} | True | 87253821 | TTTGACTAATCTACTCCTAAAG | [] |
| 1363 | 3 | HTR1F | {'exs': 0, 'nonexs': 7} | True | 87991849 | TTTAAAACTAACATCGACATCG | [] |
| 1364 | 3 | EPHA3 | {'exs': 1, 'nonexs': 6} | True | 89399505 | CCCCCTCCTCTAGACTGCATT | [] |
| 1365 | 3 | LOC10192 | unavailable | False | 91374235 | TGCCTAAGAATCTTCACTAAAG | [] |
| 1366 | 3 | EPHA6 | {'exs': 1, 'nonexs': 6} | True | 97010028 | CAACTAGAAGCAGCGGCCTTT | [] |
| 1367 | 3 | RIOX2 | unavailable | False | 97945183 | TGCAAAGGCACTAGACTACTTG | [] |
| 1368 | 3 | OR5H14 | {'exs': 0, 'nonexs': 3} | True | 98150317 | GATCTAAACATCATTTCTTTTG | [] |
| 1369 | 3 | OR5H6 | {'exs': 0, 'nonexs': 3} | True | 98265261 | TGAGATCTAAACATTGCTTTTG | [] |
| 1370 | 3 | OR5H2 | {'exs': 0, 'nonexs': 3} | True | 98283831 | GAAATCTAAACATTTCTTTTTG | ['CTTTTTACCATTTTTGTGAATG'] |
| 1371 | 3 | ADGRG7 | unavailable | False | 1.01E+08 | ACTGTCTAGATGCTTTCCTTTG | [] |
| 1372 | 3 | ABI3BP | {'exs': 0, 'nonexs': 7} | True | 1.01E+08 | GCAGCATCTACCATTTTCCAGG | [] |
| 1373 | 3 | ABI3BP | {'exs': 0, 'nonexs': 7} | True | 1.01E+08 | TCCTTGGCTACTTAAGCCAAGG | [] |
| 1374 | 3 | SENP7 | {'exs': 0, 'nonexs': 7} | True | 1.01E+08 | ACAGATTAACTAGCTACTGCTG | [] |
| 1375 | 3 | LOC10537 | unavailable | False | 1.02E+08 | AAATCTACTGTGGCGCTGCCTC | [] |
| 1376 | 3 | CEP97 | {'exs': 3, 'nonexs': 5} | False | 1.02E+08 | GCTACACAGTAACACCAACATG | [] |
| 1377 | 3 | NXPE3 | unavailable | False | 1.02E+08 | GACAGGCTAGGTTTCCAAGGG | [] |
| 1378 | 3 | NFKBIZ | {'exs': 0, 'nonexs': 7} | True | 1.02E+08 | TAATGGAGCTAATACGGTGGAG | [] |
| 1379 | 3 | ALCAM | {'exs': 0, 'nonexs': 7} | True | 1.06E+08 | AGACTACTTACTGAACACCTTG | [] |
| 1380 | 3 | CBLB | {'exs': 0, 'nonexs': 7} | True | 1.06E+08 | GGCTGCTATAGATTTAGACGTG | [] |
| 1381 | 3 | CBLB | {'exs': 0, 'nonexs': 7} | True | 1.06E+08 | GTTTGCTATTTGTCAGCCATAG | [] |
| 1382 | 3 | CCDC54 | {'exs': 1, 'nonexs': 7} | True | 1.07E+08 | AGAACATAACTCTATAGTCGAG | [] |
| 1383 | 3 | BBX | {'exs': 0, 'nonexs': 7} | True | 1.08E+08 | TGGACTCACTAGCATCCCTCGG | [] |
| 1384 | 3 | HHLA2 | {'exs': 1, 'nonexs': 7} | True | 1.08E+08 | TTTCCTATACTTTTCCTGAAAG | [] |
| 1385 | 3 | MYH15 | {'exs': 1, 'nonexs': 7} | True | 1.08E+08 | GATGCTATTCTTCTTGAACCTG | [] |
| 1386 | 3 | CIP2A | unavailable | False | 1.09E+08 | GTCTATATACTGAGATTCACAG | [] |
| 1387 | 3 | TRAT1 | {'exs': 0, 'nonexs': 7} | True | 1.09E+08 | CATGGTCCAGCTAGTTTATAGG | [] |
| 1388 | 3 | GUCA1C | {'exs': 0, 'nonexs': 7} | True | 1.09E+08 | TCTCACAGCTACTTCATTTTCG | ['ATTTTCACCTTCCCTAGACCAG'] |
| 1389 | 3 | DPPA2 | {'exs': 0, 'nonexs': 7} | True | 1.09E+08 | GGTTGCTGCTACTTCTCTACTG | [] |
| 1390 | 3 | DPPA4 | {'exs': 0, 'nonexs': 7} | True | 1.09E+08 | GATATTCTATTCCCATTGGAGG | [] |
| 1391 | 3 | NECTIN3 | unavailable | False | 1.11E+08 | CTAAACATACCACTCCCTCCTG | [] |
| 1392 | 3 | CD96 | {'exs': 1, 'nonexs': 7} | True | 1.12E+08 | ACTAGAGGGTCTCCATCTCATG | [] |
| 1393 | 3 | PHLDB2 | {'exs': 0, 'nonexs': 7} | True | 1.12E+08 | GTTCACTACAACAAGAAGTGAG | [] |
| 1394 | 3 | ABHD10 | {'exs': 1, 'nonexs': 7} | True | 1.12E+08 | TGATACTAGTTAACTATAGTTG | [] |
| 1395 | 3 | TAGLN3 | {'exs': 0, 'nonexs': 6} | True | 1.12E+08 | CGCGTCCTACATGATCTGCCT | [] |
| 1396 | 3 | C3orf52 | {'exs': 0, 'nonexs': 7} | True | 1.12E+08 | TCCAGCTAAGCGACAGAACGA | [] |
| 1397 | 3 | GCSAM | unavailable | False | 1.12E+08 | TTCACTATAAATGGGAAAACTG | [] |
| 1398 | 3 | NEPRO | unavailable | False | 1.13E+08 | AACATCTAAACTCCCATTAAAG | [] |
| 1399 | 3 | NEPRO | unavailable | False | 1.13E+08 | TTATGCACTCTAGTTTCTTTGG | [] |
| 1400 | 3 | BOC | {'exs': 0, 'nonexs': 7} | True | 1.13E+08 | TTTATGCTAAGAGTAACAAGAG | [] |
| 1401 | 3 | BOC | {'exs': 0, 'nonexs': 7} | True | 1.13E+08 | TCTGCCTAAATTGTGAGAGGTG | [] |
| 1402 | 3 | ATP6V1A | {'exs': 7, 'nonexs': 1} | False | 1.14E+08 | GCTTCTAATCTTCAAGGCTACG | [] |
| 1403 | 3 | GRAMD1C | {'exs': 0, 'nonexs': 7} | True | 1.14E+08 | TCAGATCACTAGCTTTCAACAG | [] |
| 1404 | 3 | TIGIT | {'exs': 0, 'nonexs': 5} | True | 1.14E+08 | TGGTTGCTAACCAGTCTCTGTG | [] |
| 1405 | 3 | IGSF11 | {'exs': 0, 'nonexs': 7} | True | 1.19E+08 | CATGTCCTATACCAAGGACCC | [] |
| 1406 | 3 | IGSF11 | {'exs': 0, 'nonexs': 7} | True | 1.19E+08 | TATCCTCCTAGTTGGGCACATG | [] |
| 1407 | 3 | TEX55 | unavailable | False | 1.19E+08 | TCCAATTCTATACCTGGCACAG | [] |
| 1408 | 3 | POGLUT1 | {'exs': 0, 'nonexs': 7} | True | 1.19E+08 | TATGATGACTACTATAGTTCAG | [] |
| 1409 | 3 | ADPRH | {'exs': 1, 'nonexs': 7} | True | 1.2E+08 | TCTCCCTAAAGGGAAATTACAG | [] |
| 1410 | 3 | PLA1A | {'exs': 0, 'nonexs': 7} | True | 1.2E+08 | ACTACACACAGGCTATCTTCAG | [] |
| 1411 | 3 | HGD | {'exs': 0, 'nonexs': 7} | True | 1.21E+08 | GGCTAGATCTAAAGAAAATCAG | [] |
| 1412 | 3 | STXBP5L | {'exs': 0, 'nonexs': 7} | True | 1.21E+08 | AGGGCTACAACCTAGAAATAAG | [] |
| 1413 | 3 | GOLGB1 | {'exs': 1, 'nonexs': 7} | True | 1.22E+08 | AACTAAGTCTATAGATGGCCCG | [] |
| 1414 | 3 | IQCB1 | {'exs': 0, 'nonexs': 7} | True | 1.22E+08 | GTTACTCACTAAGGTGGTTTGG | [] |
| 1415 | 3 | ILDR1 | {'exs': 0, 'nonexs': 7} | True | 1.22E+08 | GCTTGGTGACTAAATGACCACG | ['ACCACACTCCTTCCACTATGAG'] |
| 1416 | 3 | ILDR1 | {'exs': 0, 'nonexs': 7} | True | 1.22E+08 | ACTCCTTCCACTATGAGAGCTG | [] |
| 1417 | 3 | CSTA | {'exs': 0, 'nonexs': 7} | True | 1.22E+08 | CATGCTGCTAAAAGCCCGTCAC | [] |
| 1418 | 3 | CCDC58 | {'exs': 1, 'nonexs': 6} | True | 1.22E+08 | GGTTTCTAAACCCTCCTTGCAG | [] |
| 1419 | 3 | FAM162A | {'exs': 0, 'nonexs': 7} | True | 1.22E+08 | TACCTCTGCTACTCTGTTTTGG | [] |
| 1420 | 3 | PARP15 | {'exs': 0, 'nonexs': 7} | True | 1.23E+08 | TGTCTACCCTACTCTGCAAGGG | [] |
| 1421 | 3 | HSPBAP1 | {'exs': 0, 'nonexs': 7} | True | 1.23E+08 | TCTCCTGACCTAAATGAAAATG | [] |
| 1422 | 3 | ADCY5 | {'exs': 0, 'nonexs': 7} | True | 1.23E+08 | ACAGCTGCTAACTGAGCGGGG | [] |
| 1423 | 3 | HACD2 | unavailable | False | 1.23E+08 | CAGGAACTATTCAAATTTCTTG | [] |
| 1424 | 3 | KALRN | {'exs': 2, 'nonexs': 6} | False | 1.25E+08 | TGCGCGCTAAACGTAAGTGGAC | [] |
| 1425 | 3 | KALRN | {'exs': 2, 'nonexs': 6} | False | 1.25E+08 | GGGAGATGGCTACGTCCCTTG | [] |
| 1426 | 3 | MUC13 | {'exs': 0, 'nonexs': 3} | True | 1.25E+08 | ACTCACCTAATAGTCAGGGCGG | [] |
| 1427 | 3 | SLC12A8 | {'exs': 0, 'nonexs': 7} | True | 1.25E+08 | GCATCTAGTAGTGAGGCATCAG | [] |
| 1428 | 3 | SLC41A3 | {'exs': 0, 'nonexs': 7} | True | 1.26E+08 | GAAATCTAAATAGAGGTTTTTG | [] |
| 1429 | 3 | CHST13 | {'exs': 0, 'nonexs': 7} | True | 1.27E+08 | GGACCGCTAGAGCAGCCGCAG | [] |
| 1430 | 3 | C3orf22 | {'exs': 0, 'nonexs': 7} | True | 1.27E+08 | CAGTCTCTAGGAGAGGCCCCT | [] |
| 1431 | 3 | TXNRD3 | {'exs': 0, 'nonexs': 1} | False | 1.27E+08 | GCAGCAGGCCTAGCCTCAGCA | [] |
| 1432 | 3 | RUVBL1 | {'exs': 7, 'nonexs': 1} | False | 1.28E+08 | TGGTCATCTATTTCCAGGTGTG | [] |
| 1433 | 3 | DNAJB8 | {'exs': 1, 'nonexs': 7} | True | 1.28E+08 | CTACTTGCTGTCCATCCATTTG | [] |
| 1434 | 3 | GATA2 | {'exs': 3, 'nonexs': 5} | False | 1.28E+08 | TCTGTTCCCTAGCCCATGGCG | [] |
| 1435 | 3 | RPN1 | {'exs': 7, 'nonexs': 1} | False | 1.29E+08 | CAGGGGCTACAGGGCATCCAG | [] |
| 1436 | 3 | ISY1 | {'exs': 5, 'nonexs': 4} | False | 1.29E+08 | CTGGGTCCTAATACCCCAGGA | [] |
| 1437 | 3 | COPG1 | unavailable | False | 1.29E+08 | AGTAGACTAGTGGTGGCCCCA | [] |
| 1438 | 3 | EFCAB12 | unavailable | False | 1.29E+08 | TGGCTAGCTCTAGTTGATGTAG | [] |
| 1439 | 3 | H1-8 | unavailable | False | 1.3E+08 | GGCCCTAAGCTTCAGCCCTCT | [] |
| 1440 | 3 | COL6A5 | {'exs': 1, 'nonexs': 0} | False | 1.3E+08 | TTCTGGCTAATACACAGAGTAG | [] |
| 1441 | 3 | COL6A5 | {'exs': 1, 'nonexs': 0} | False | 1.3E+08 | CTTTTCTCCTATGATGACCTTG | [] |
| 1442 | 3 | COL6A6 | {'exs': 0, 'nonexs': 7} | True | 1.31E+08 | TAATCCCTACTGGATAAAAAAG | [] |
| 1443 | 3 | ASTE1 | {'exs': 0, 'nonexs': 7} | True | 1.31E+08 | TTACCAACTAGCGAGGAGCAAG | [] |
| 1444 | 3 | NEK11 | {'exs': 1, 'nonexs': 6} | True | 1.31E+08 | AGTTGCCTAGAGATGGTTCTGG | [] |
| 1445 | 3 | NUDT16 | {'exs': 0, 'nonexs': 7} | True | 1.31E+08 | GCTGCCTCTAGTGATGAGCTGG | [] |
| 1446 | 3 | ACP3 | unavailable | False | 1.32E+08 | CTGTGCACACTAATCTGTACTG | [] |
| 1447 | 3 | ACP3 | unavailable | False | 1.32E+08 | TACTGGCTAATACTGAATGTGG | [] |
| 1448 | 3 | UBA5 | {'exs': 1, 'nonexs': 7} | True | 1.33E+08 | TATCTACATATTCTTCATTTTG | [] |
| 1449 | 3 | CEP63 | {'exs': 2, 'nonexs': 5} | False | 1.35E+08 | AAGAGGCTACTTTAAGGCTGTG | [] |
| 1450 | 3 | PPP2R3A | {'exs': 1, 'nonexs': 6} | True | 1.36E+08 | GGCAGCTATTCTTCATCCACTG | [] |
| 1451 | 3 | PCCB | {'exs': 0, 'nonexs': 7} | True | 1.36E+08 | TTGACCTAAGGTTCCTTACCTG | [] |
| 1452 | 3 | IL20RB | {'exs': 0, 'nonexs': 7} | True | 1.37E+08 | CCGCAAACCTATGAGATCCAGG | [] |
| 1453 | 3 | DZIP1L | {'exs': 0, 'nonexs': 7} | True | 1.38E+08 | TCCAAATCTACTGCATCAAAAG | ['GCATCAAAAATTCTACAGGTGG'] |
| 1454 | 3 | CEP70 | {'exs': 0, 'nonexs': 7} | True | 1.38E+08 | TCTATGGACTAATCCAAACAAG | ['ACAAACTCTAACTGCTTGTAAG'] |
| 1455 | 3 | FOXL2NB | unavailable | False | 1.39E+08 | GGATGTACTAATAGACACAGTG | [] |
| 1456 | 3 | PRR23B | {'exs': 0, 'nonexs': 7} | True | 1.39E+08 | GGGGGTCTATGCCTGGAACAG | [] |
| 1457 | 3 | NMNAT3 | {'exs': 0, 'nonexs': 7} | True | 1.4E+08 | CCCCCCTCCCTAGCTTGTCTT | [] |
| 1458 | 3 | CLSTN2 | {'exs': 0, 'nonexs': 7} | True | 1.41E+08 | TGGGCACTAGTAGGGGAGGGT | [] |
| 1459 | 3 | SLC25A36 | {'exs': 0, 'nonexs': 7} | True | 1.41E+08 | GTGCTGCTATCCATTGAGTAGG | [] |
| 1460 | 3 | RASA2 | {'exs': 0, 'nonexs': 7} | True | 1.42E+08 | TGTTGCTACCACGCTTTCACTG | [] |
| 1461 | 3 | RASA2 | {'exs': 0, 'nonexs': 7} | True | 1.42E+08 | TAAACTCTAAGATGCTTTCCCG | ['CCAACAATTGGATTTTCCCTAG'] |
| 1462 | 3 | ATP1B3 | {'exs': 3, 'nonexs': 5} | False | 1.42E+08 | CTCATACTATGCACGTGCTGTG | [] |
| 1463 | 3 | XRN1 | {'exs': 3, 'nonexs': 5} | False | 1.42E+08 | TTTTTCTATTTTAAAATAAAAG | [] |
| 1464 | 3 | DIPK2A | unavailable | False | 1.44E+08 | ATTGTGCTAGGTATTCACGCAG | [] |
| 1465 | 3 | DIPK2A | unavailable | False | 1.44E+08 | CACCATAGACTACCTCACGTTG | [] |
| 1466 | 3 | PLSCR4 | {'exs': 0, 'nonexs': 7} | True | 1.46E+08 | TCTCTCTATCTTGAACGTTGTG | [] |
| 1467 | 3 | PLSCR1 | {'exs': 0, 'nonexs': 7} | True | 1.47E+08 | CTAATCCACTACCACACTCCTG | [] |
| 1468 | 3 | PLSCR5 | {'exs': 0, 'nonexs': 7} | True | 1.47E+08 | GGTCCTAAGCTATCCTTCTATG | [] |
| 1469 | 3 | CPB1 | {'exs': 0, 'nonexs': 5} | True | 1.49E+08 | TCAACTAGTACAGGTGTTCCAG | [] |
| 1470 | 3 | GYG1 | {'exs': 0, 'nonexs': 7} | True | 1.49E+08 | GTGTTTCTACTGGAGGTAAGTG | [] |
| 1471 | 3 | GYG1 | {'exs': 0, 'nonexs': 7} | True | 1.49E+08 | GATACTAAGTATCAGAAACAAG | [] |
| 1472 | 3 | TM4SF1 | {'exs': 1, 'nonexs': 7} | True | 1.49E+08 | AGTATGTTACACTAATACAAAG | [] |
| 1473 | 3 | COMMD2 | {'exs': 0, 'nonexs': 7} | True | 1.5E+08 | ACTGGTACTACTTGATGTTGCG | [] |
| 1474 | 3 | PFN2 | {'exs': 0, 'nonexs': 7} | True | 1.5E+08 | GCCTAGCAGCTAGAACCCAGA | [] |
| 1475 | 3 | TMEM183 | unavailable | False | 1.5E+08 | CAGTAACTACGCTCTCAGGGA | [] |
| 1476 | 3 | SELENOT | unavailable | False | 1.51E+08 | TGGTGCTATGATCGATGGTGTG | [] |
| 1477 | 3 | GPR87 | {'exs': 0, 'nonexs': 7} | True | 1.51E+08 | ATAAAAGGCCTACACATCAGTG | [] |
| 1478 | 3 | AADAC | {'exs': 0, 'nonexs': 7} | True | 1.52E+08 | TTTACTATAGATTTTCCTTTAG | [] |
| 1479 | 3 | MBNL1 | {'exs': 6, 'nonexs': 2} | False | 1.52E+08 | TTGTGGCTAGTCAGATGTTCGG | [] |
| 1480 | 3 | MBNL1 | {'exs': 6, 'nonexs': 2} | False | 1.52E+08 | TGATGAAAATTCTACATCTGGG | [] |
| 1481 | 3 | ARHGEF26 | {'exs': 1, 'nonexs': 6} | True | 1.54E+08 | GAGACTACACGTTGGTCTCCAC | [] |
| 1482 | 3 | GPR149 | {'exs': 0, 'nonexs': 7} | True | 1.54E+08 | AAATACCCACTAACTACCCTTG | [] |
| 1483 | 3 | STRIT1 | unavailable | False | 1.55E+08 | ACCTTTCTAAGAGAAGACAACG | ['GACAACATAAATCATTATGATG'] |
| 1484 | 3 | PLCH1 | {'exs': 0, 'nonexs': 7} | True | 1.55E+08 | AGGGACCACTACCTCCATACAC | [] |
| 1485 | 3 | SSR3 | {'exs': 1, 'nonexs': 7} | True | 1.57E+08 | ATGGTCTATTTGGAGCCAGTAG | [] |
| 1486 | 3 | LEKR1 | {'exs': 0, 'nonexs': 1} | False | 1.57E+08 | TGTGAAGCTAGGAGGTGAGAAG | [] |
| 1487 | 3 | LEKR1 | {'exs': 0, 'nonexs': 1} | False | 1.57E+08 | GCAAAATCTAACATTTTTTAGG | [] |
| 1488 | 3 | SLC66A1L | unavailable | False | 1.58E+08 | CAACTATGCCAACAGCATCCTG | [] |
| 1489 | 3 | KPNA4 | {'exs': 4, 'nonexs': 4} | False | 1.61E+08 | TTCTAAAACTGGAACCCTTCTG | [] |
| 1490 | 3 | SPTSSB | unavailable | False | 1.61E+08 | CCGACTCTACTGAAGGGGCAC | [] |
| 1491 | 3 | ZBBX | {'exs': 1, 'nonexs': 5} | False | 1.67E+08 | TACCTCAGCTAGGAGACATCCC | ['TCTTCCCACTTCTTCCTTCTTG', 'GACATCCATTTTACTCTTCCCG'] |
| 1492 | 3 | GOLIM4 | {'exs': 1, 'nonexs': 6} | True | 1.68E+08 | GCCGCTACATTTCAGCTCTTCC | [] |
| 1493 | 3 | PHC3 | {'exs': 1, 'nonexs': 7} | True | 1.7E+08 | CCCTTTCCTAAATAAACTGAGG | [] |
| 1494 | 3 | SKIL | {'exs': 0, 'nonexs': 7} | True | 1.7E+08 | AACAGTTTCTATTCTTTAGCAG | [] |
| 1495 | 3 | SLC7A14 | {'exs': 0, 'nonexs': 7} | True | 1.7E+08 | GTGTTTCTCCTACTCTGGAGAG | [] |
| 1496 | 3 | RPL22L1 | {'exs': 3, 'nonexs': 5} | False | 1.71E+08 | TTTGCCTAGTCCTCCGACTCTC | [] |
| 1497 | 3 | ECT2 | {'exs': 8, 'nonexs': 0} | False | 1.73E+08 | ATCTCCTAATTCTTCACAAATG | [] |
| 1498 | 3 | SPATA16 | {'exs': 0, 'nonexs': 7} | True | 1.73E+08 | AATGACTACCTTTGCTGAACAG | [] |
| 1499 | 3 | SPATA16 | {'exs': 0, 'nonexs': 7} | True | 1.73E+08 | AAACCTAGTACCTGATATTTAG | [] |
| 1500 | 3 | NLGN1 | {'exs': 0, 'nonexs': 5} | True | 1.74E+08 | TCTGGCTATACCCTGGTTGTTG | [] |
| 1501 | 3 | NAALADL2 | {'exs': 0, 'nonexs': 7} | True | 1.76E+08 | GCCTATTCAGCCTTTTTTAAAG | [] |
| 1502 | 3 | TBL1XR1 | {'exs': 1, 'nonexs': 7} | True | 1.77E+08 | CTAGTAGCGCTATTTCCGAAGG | [] |
| 1503 | 3 | ZMAT3 | {'exs': 0, 'nonexs': 7} | True | 1.79E+08 | ATAATCACTATACATATCCCAG | [] |
| 1504 | 3 | TTC14 | {'exs': 0, 'nonexs': 7} | True | 1.81E+08 | ATATCACTAAATTTCTAAGAGG | [] |
| 1505 | 3 | TTC14 | {'exs': 0, 'nonexs': 7} | True | 1.81E+08 | CCTAAGCCTACCCGTGGTCAG | [] |
| 1506 | 3 | ATP11B | {'exs': 0, 'nonexs': 7} | True | 1.83E+08 | CTCAGCTAGCAGGTGCCTACT | [] |
| 1507 | 3 | ATP11B | {'exs': 0, 'nonexs': 7} | True | 1.83E+08 | TACCTAGTGCACGCAACAACTG | [] |
| 1508 | 3 | ATP11B | {'exs': 0, 'nonexs': 7} | True | 1.83E+08 | ATCTTCCTATAACTCGTTCCAG | [] |
| 1509 | 3 | DCUN1D1 | {'exs': 0, 'nonexs': 7} | True | 1.83E+08 | TGCTACACTGTTGTACTTTTTG | [] |
| 1510 | 3 | MCF2L2 | {'exs': 0, 'nonexs': 7} | True | 1.83E+08 | CTCCTACCTCAGCCTCCCAAA | [] |
| 1511 | 3 | HTR3D | {'exs': 0, 'nonexs': 7} | True | 1.84E+08 | ACCTGCCTAGGTGTTCCAGAG | [] |
| 1512 | 3 | HTR3C | {'exs': 0, 'nonexs': 7} | True | 1.84E+08 | GTCTGCCTAGGTGTTCCAGAG | [] |
| 1513 | 3 | HTR3E | {'exs': 0, 'nonexs': 7} | True | 1.84E+08 | ACCTGCCTAGGTGTTCCAGAG | [] |
| 1514 | 3 | AP2M1 | {'exs': 4, 'nonexs': 4} | False | 1.84E+08 | CCTAGTGGCAGCTAGCAGCGA | [] |
| 1515 | 3 | ABCF3 | {'exs': 0, 'nonexs': 7} | True | 1.84E+08 | GTGGCCCTAGAGGAAGCCTTC | [] |
| 1516 | 3 | EEF1AKMT | unavailable | False | 1.84E+08 | AGGTCTACCACACCTCACACA | [] |
| 1517 | 3 | CAMK2N2 | {'exs': 0, 'nonexs': 7} | True | 1.84E+08 | CGCGTCTACACTCCGGACGGC | [] |
| 1518 | 3 | ECE2 | {'exs': 0, 'nonexs': 7} | True | 1.84E+08 | AGGTCTACCACACCTCACACA | [] |
| 1519 | 3 | CLCN2 | {'exs': 0, 'nonexs': 7} | True | 1.84E+08 | TCCCGCCTAGCTCTGACCCTG | [] |
| 1520 | 3 | CHRD | {'exs': 0, 'nonexs': 7} | True | 1.84E+08 | GCTGCTCCCTAAGAGCCTTCG | [] |
| 1521 | 3 | VPS8 | {'exs': 0, 'nonexs': 7} | True | 1.85E+08 | TTCTAGGCATGCTAAAACAATG | [] |
| 1522 | 3 | AHSG | {'exs': 0, 'nonexs': 6} | True | 1.87E+08 | GTCTAGCCTAGACCTTGAAGTG | [] |
| 1523 | 3 | SST | {'exs': 1, 'nonexs': 6} | True | 1.88E+08 | GAAAGCTAACAGGATGTGAAAG | [] |
| 1524 | 3 | RTP2 | {'exs': 0, 'nonexs': 7} | True | 1.88E+08 | AGCTCCACTAAAAGAAGGCAGG | [] |
| 1525 | 3 | LPP | {'exs': 0, 'nonexs': 7} | True | 1.89E+08 | TGAATCTAAAGGTCAGTGCTCG | [] |
| 1526 | 3 | TPRG1 | {'exs': 0, 'nonexs': 7} | True | 1.89E+08 | TACAGCTCCTATAATTCATAAG | [] |
| 1527 | 3 | TP63 | {'exs': 0, 'nonexs': 7} | True | 1.9E+08 | GGCTCTATGGGTACACTGATCC | [] |
| 1528 | 3 | IL1RAP | {'exs': 1, 'nonexs': 7} | True | 1.91E+08 | CCCAGCTACTCAGGAGGCTGA | [] |
| 1529 | 3 | GMNC | unavailable | False | 1.91E+08 | GGGTCACTAAGACTGCTTAGGG | [] |
| 1530 | 3 | FGF12 | {'exs': 0, 'nonexs': 7} | True | 1.92E+08 | TTCTCAGCTATGTTGAATCTTG | [] |
| 1531 | 3 | PLAAT1 | unavailable | False | 1.93E+08 | ACACAGCTATTTCCTGAAATCG | [] |
| 1532 | 3 | CPN2 | {'exs': 0, 'nonexs': 7} | True | 1.94E+08 | GCTACTAGGGCCCTGCTGCCC | [] |
| 1533 | 3 | ATP13A3 | {'exs': 0, 'nonexs': 5} | True | 1.94E+08 | TCACTGCTATGTTATGGTGATG | [] |
| 1534 | 3 | XXYLT1 | unavailable | False | 1.95E+08 | GAGCGCCTAGTCCTCCGGGAT | [] |
| 1535 | 3 | PPP1R2 | {'exs': 5, 'nonexs': 3} | False | 1.96E+08 | CGTCTATGAACTTCGTAATTTG | [] |
| 1536 | 3 | TFRC | {'exs': 8, 'nonexs': 0} | False | 1.96E+08 | GGTGGCTATCCACGATGGAGA | [] |
| 1537 | 3 | TM4SF19 | {'exs': 0, 'nonexs': 7} | True | 1.96E+08 | GGATACCTAAAGGGGAAGTTTG | [] |
| 1538 | 3 | BDH1 | {'exs': 0, 'nonexs': 7} | True | 1.98E+08 | CTTTGACTACGACGTGACCTTG | [] |
| 1539 | 3 | FYTTD1 | {'exs': 0, 'nonexs': 7} | True | 1.98E+08 | GACCTATCCCACGGTGACAAAC | [] |
| 1540 | 3 | LRCH3 | {'exs': 0, 'nonexs': 7} | True | 1.98E+08 | AGTTTCTAGTAGTCCTGTGAAG | [] |
| 1541 | 3 | LMLN | {'exs': 0, 'nonexs': 7} | True | 1.98E+08 | ATTCCTAGTGCCATATACACAG | [] |
| 1542 | 4 | ZNF718 | unavailable | False | 162121 | CATTTCTAAAATTTTCCTCCAG | [] |
| 1543 | 4 | ZNF732 | {'exs': 0, 'nonexs': 7} | True | 271098 | TATTTCTAGAGTTTCTCTCCAG | [] |
| 1544 | 4 | ZNF141 | {'exs': 1, 'nonexs': 7} | True | 383129 | GAAAGCTAAAATGGGCTTGTTG | [] |
| 1545 | 4 | PIGG | {'exs': 0, 'nonexs': 7} | True | 539368 | GCTTAGTCTAAGACTGTGTGAG | [] |
| 1546 | 4 | SLC49A3 | unavailable | False | 681957 | CTCAGCTACGTGATCACCCAC | [] |
| 1547 | 4 | CPLX1 | {'exs': 0, 'nonexs': 7} | True | 788500 | GCGCGGCTACTTCTTGAGCAT | [] |
| 1548 | 4 | TMEM175 | {'exs': 0, 'nonexs': 7} | True | 950397 | CGCGCTCTAACAGACGAAGAG | [] |
| 1549 | 4 | TMEM175 | {'exs': 0, 'nonexs': 7} | True | 959495 | TGGCTGCTAGCAGGGGGCAGG | [] |
| 1550 | 4 | DGKQ | {'exs': 0, 'nonexs': 7} | True | 960519 | CAGCCACCCCTACCTAGGATC | [] |
| 1551 | 4 | SLC26A1 | {'exs': 0, 'nonexs': 7} | True | 988832 | GCCCTGCTACAGATGGGCATC | [] |
| 1552 | 4 | FGFRL1 | {'exs': 0, 'nonexs': 5} | True | 1025346 | TGCCGTCTAGCACTGATAGTGC | [] |
| 1553 | 4 | LOC10192 | unavailable | False | 1200642 | ATCTCCCTAGCAGGAAGCTGG | [] |
| 1554 | 4 | CTBP1 | {'exs': 0, 'nonexs': 7} | True | 1212229 | CCCGGGCTACAACTGGTCACT | [] |
| 1555 | 4 | MAEA | {'exs': 1, 'nonexs': 6} | True | 1328874 | ACGAAGCTAAGGCAGGGTGGG | [] |
| 1556 | 4 | MAEA | {'exs': 1, 'nonexs': 6} | True | 1329168 | CTACATGATGTACACCTTCTCG | [] |
| 1557 | 4 | UVSSA | unavailable | False | 1350028 | CCAGTCCTAAGGCCCGACTGT | [] |
| 1558 | 4 | UVSSA | unavailable | False | 1371980 | CTGAGGGCTACGGGTGCAGAG | [] |
| 1559 | 4 | UVSSA | unavailable | False | 1385960 | CGCTCTCTAGTTCAGTGCGTAC | [] |
| 1560 | 4 | FAM53A | {'exs': 0, 'nonexs': 7} | True | 1619067 | GGAGGGCCGTGCTATGTCATG | [] |
| 1561 | 4 | TACC3 | {'exs': 7, 'nonexs': 2} | False | 1740560 | ACACCCTAGAGCCTAGGGTGA | [] |
| 1562 | 4 | LETM1 | {'exs': 6, 'nonexs': 2} | False | 1814423 | TGGTTCTAGCTCTTCACCTCTG | [] |
| 1563 | 4 | NSD2 | unavailable | False | 1942400 | TCATTCTAAGTGCAGTACAGAG | [] |
| 1564 | 4 | NSD2 | unavailable | False | 1978808 | CTGGCGCTATTTGCCCTCTGT | [] |
| 1565 | 4 | NELFA | unavailable | False | 1983318 | AGGTGGTTCTAGGACACATTGG | [] |
| 1566 | 4 | C4orf48 | {'exs': 0, 'nonexs': 5} | True | 2041905 | GAGGAACCTAGGACGCTCCAG | [] |
| 1567 | 4 | POLN | {'exs': 0, 'nonexs': 5} | True | 2072113 | TGGGGCTACAGACAAAATGAAG | [] |
| 1568 | 4 | MXD4 | {'exs': 0, 'nonexs': 7} | True | 2250543 | ACGGGCCTACGAGAGGGCGGG | [] |
| 1569 | 4 | ZFYVE28 | {'exs': 0, 'nonexs': 7} | True | 2270563 | GAGGCTAGCGTGGGCAGGACA | [] |
| 1570 | 4 | ZFYVE28 | {'exs': 0, 'nonexs': 7} | True | 2274002 | CTCCCCTAAAGCGCAGCCCGT | [] |
| 1571 | 4 | MFSD10 | {'exs': 0, 'nonexs': 7} | True | 2930837 | CTCAGCTACTCAGCCTTGAGC | [] |
| 1572 | 4 | MFSD10 | {'exs': 0, 'nonexs': 7} | True | 2931102 | AGAGCACCTAGGCTGCGCAGT | [] |
| 1573 | 4 | GRK4 | {'exs': 1, 'nonexs': 6} | True | 3038658 | GCCTCCACCTAGCACCATGTG | [] |
| 1574 | 4 | MSANTD1 | unavailable | False | 3255964 | GCTGGCCTAGACGCTGGACTT | [] |
| 1575 | 4 | OTOP1 | {'exs': 0, 'nonexs': 7} | True | 4188802 | CCCAGACTATATCTTACAATAG | [] |
| 1576 | 4 | NSG1 | unavailable | False | 4417434 | TCCCGCTAAGCTGACTTCTCAC | [] |
| 1577 | 4 | STX18 | {'exs': 4, 'nonexs': 4} | False | 4420033 | CCCTGGCTAGCTGTCGTACCA | [] |
| 1578 | 4 | MSX1 | {'exs': 0, 'nonexs': 7} | True | 4863142 | CCTCTATGTCAGGTGGTACATG | [] |
| 1579 | 4 | EVC2 | {'exs': 1, 'nonexs': 7} | True | 5562847 | CTTGGGCTAGTCCATGCCCAA | [] |
| 1580 | 4 | EVC | {'exs': 0, 'nonexs': 7} | True | 5811036 | -1 | -1 |
| 1581 | 4 | PPP2R2C | {'exs': 0, 'nonexs': 7} | True | 6323301 | ACATACCTAGTGCATGTCAGAG | [] |
| 1582 | 4 | GRPEL1 | {'exs': 8, 'nonexs': 1} | False | 7060861 | AGCAGCTAAGCTTCCTTCACCC | ['TTCACCACCCCCACCAGGGCGG'] |
| 1583 | 4 | AFAP1 | {'exs': 1, 'nonexs': 5} | False | 7763764 | GTCCCCTAGGTCCCGTTCTTC | ['TCTTCAATTCCCATTCCTAGAG'] |
| 1584 | 4 | ACOX3 | {'exs': 0, 'nonexs': 7} | True | 8368960 | CCCACTAGAGCTTCGATTTCAG | [] |
| 1585 | 4 | ACOX3 | {'exs': 0, 'nonexs': 7} | True | 8388307 | TCTGCTGGTCTAGACTGTAAAG | [] |
| 1586 | 4 | CPZ | {'exs': 0, 'nonexs': 7} | True | 8619616 | GCCGGGGCTAGTACTTGAGCA | [] |
| 1587 | 4 | CLNK | {'exs': 0, 'nonexs': 7} | True | 10490466 | ACCAGGCTACAGAGGCAAGAG | [] |
| 1588 | 4 | RAB28 | {'exs': 0, 'nonexs': 7} | True | 13369875 | CCTGCACTACTGTACTGAACAG | [] |
| 1589 | 4 | CC2D2A | {'exs': 0, 'nonexs': 5} | True | 15563353 | AATGCTAATGATTACAGAACAG | [] |
| 1590 | 4 | FAM200B | {'exs': 0, 'nonexs': 7} | True | 15688950 | TTTACTATGATGGGTGTGCTTG | [] |
| 1591 | 4 | BST1 | {'exs': 0, 'nonexs': 7} | True | 15723455 | TGGCTAACACGGTGAAACCCC | [] |
| 1592 | 4 | BST1 | {'exs': 0, 'nonexs': 7} | True | 15736118 | CAAAGGCTAGAACCGGCGGTT | [] |
| 1593 | 4 | BST1 | {'exs': 0, 'nonexs': 7} | True | 15737743 | TATCGTGCTAGAGGGAATACAG | [] |
| 1594 | 4 | QDPR | {'exs': 0, 'nonexs': 7} | True | 17487130 | TGAGGCCTAAAAATATGCTGGG | [] |
| 1595 | 4 | LAP3 | {'exs': 0, 'nonexs': 7} | True | 17607588 | ATCTGAACTAAGCATTGTCTTG | [] |
| 1596 | 4 | NCAPG | {'exs': 6, 'nonexs': 3} | False | 17843424 | TTCCTAACTTAGATCTTCATTG | [] |
| 1597 | 4 | LCORL | {'exs': 0, 'nonexs': 7} | True | 17845720 | AATCTATTTCTCTTCGCTTTTG | [] |
| 1598 | 4 | LCORL | {'exs': 0, 'nonexs': 7} | True | 17883719 | AGTAATCTACACAGGCTTGCTG | [] |
| 1599 | 4 | PACRGL | {'exs': 0, 'nonexs': 7} | True | 20727294 | TGATGATGCTAAGGCTCCCCTC | [] |
| 1600 | 4 | KCNIP4 | {'exs': 0, 'nonexs': 7} | True | 20731936 | GTAGCTAGCTGCTAATACTCAG | [] |
| 1601 | 4 | PI4K2B | {'exs': 1, 'nonexs': 6} | True | 25277186 | TACTACCAGGAGGAAAAAAATG | [] |
| 1602 | 4 | ZCCHC4 | {'exs': 0, 'nonexs': 7} | True | 25352081 | CTTTTCTATATGCCCTGAAAAG | [] |
| 1603 | 4 | SLC34A2 | {'exs': 0, 'nonexs': 7} | True | 25676748 | CCCCTACAAGGCCGTGCATTC | [] |
| 1604 | 4 | TBC1D19 | {'exs': 0, 'nonexs': 7} | True | 26722869 | AGGTTAACACTCTATGCCTCAG | [] |
| 1605 | 4 | PCDH7 | {'exs': 0, 'nonexs': 6} | True | 31142495 | TCAGGGCTACATCTGGAAGAG | [] |
| 1606 | 4 | PCDH7 | {'exs': 0, 'nonexs': 6} | True | 31142848 | TCCTTTATAGCCTACAGGTAAG | ['GGTAAACTTCTCTTCTAGTGAG'] |
| 1607 | 4 | ARAP2 | {'exs': 0, 'nonexs': 7} | True | 36067906 | TTATTTCCTACTTCAAAATCTG | [] |
| 1608 | 4 | DTHD1 | {'exs': 0, 'nonexs': 7} | True | 36343812 | CAGGGGCTACATCAAAACACTG | [] |
| 1609 | 4 | C4orf19 | {'exs': 0, 'nonexs': 7} | True | 37590999 | CTCCCCTAATCAGTCTCATCCC | ['TCATCCAAATCTTCTCCTGCAG'] |
| 1610 | 4 | RELL1 | {'exs': 0, 'nonexs': 7} | True | 37631387 | CACCTGCTACTCTGTGCCACT | [] |
| 1611 | 4 | TLR1 | {'exs': 0, 'nonexs': 5} | True | 38796470 | GTGTAATCTATTTCTTTGCTTG | [] |
| 1612 | 4 | FAM114A1 | {'exs': 1, 'nonexs': 7} | True | 38927175 | GTAATGTGCTAGAGGAGGAGAG | [] |
| 1613 | 4 | KLHL5 | {'exs': 0, 'nonexs': 6} | True | 39126794 | TAGGGCAGTCTAATGACTTCTG | [] |
| 1614 | 4 | UAS | {'exs': 5, 'nonexs': 4} | False | 39463640 | AGTTTCTAGACTAAAGCCATAG | [] |
| 1615 | 4 | UGDH | {'exs': 0, 'nonexs': 7} | True | 39500142 | TCTCTACACTTTAGGTTTCTTG | [] |
| 1616 | 4 | PDS5A | {'exs': 5, 'nonexs': 3} | False | 39898255 | GTAAAGCTAGAGAAAAGTGTG | [] |
| 1617 | 4 | CHRNA9 | {'exs': 0, 'nonexs': 7} | True | 40354519 | ATCTGTGACTAATCCGCTCTTG | [] |
| 1618 | 4 | NSUN7 | {'exs': 1, 'nonexs': 7} | True | 40786773 | CTCTTCTATTTGTCAAAATCAG | [] |
| 1619 | 4 | APBB2 | {'exs': 0, 'nonexs': 7} | True | 40816091 | GTGCAGCTATGGCATTTCGGT | [] |
| 1620 | 4 | APBB2 | {'exs': 0, 'nonexs': 7} | True | 40881405 | CCCCACCTAATCACTCCTAGG | [] |
| 1621 | 4 | TMEM33 | {'exs': 1, 'nonexs': 6} | True | 41954198 | TTAAACTATGGAACTGTTGGTG | [] |
| 1622 | 4 | BEND4 | {'exs': 0, 'nonexs': 7} | True | 42117517 | ACTAATCCCCAGATCCATCCTG | [] |
| 1623 | 4 | BEND4 | {'exs': 0, 'nonexs': 7} | True | 42117725 | TGCATCCTAATGAATTCTGCAG | [] |
| 1624 | 4 | KCTD8 | {'exs': 0, 'nonexs': 7} | True | 44174789 | ATTACTATAACCCATACTTCTG | [] |
| 1625 | 4 | ATP10D | {'exs': 0, 'nonexs': 7} | True | 47591380 | TATCTAGCTTTCTTTACCTTTG | [] |
| 1626 | 4 | CNGA1 | {'exs': 0, 'nonexs': 7} | True | 47936420 | TCGGTTCTATGTAGAGTCGATG | [] |
| 1627 | 4 | ZAR1.00 | {'exs': 0, 'nonexs': 7} | True | 48494243 | TTTCACCTAAATGATGTATTTG | [] |
| 1628 | 4 | FRYL | {'exs': 1, 'nonexs': 5} | False | 48548782 | TCACTGCTATGTTACACCTATG | [] |
| 1629 | 4 | DCUN1D4 | {'exs': 0, 'nonexs': 7} | True | 51913583 | AAAGTCCTAGGACATCTGTTTG | [] |
| 1630 | 4 | USP46 | {'exs': 0, 'nonexs': 7} | True | 52598600 | ATGACTACTGGAGAATAATCTG | [] |
| 1631 | 4 | FIP1L1 | {'exs': 8, 'nonexs': 1} | False | 53459448 | CATGCCTATTCTGCAGGTGTAG | [] |
| 1632 | 4 | LNX1 | {'exs': 0, 'nonexs': 7} | True | 53460906 | TGATTCTATAAAAAAGTGCCAG | [] |
| 1633 | 4 | CHIC2 | {'exs': 0, 'nonexs': 7} | True | 54010094 | TGCTAATCTGGTCGAAAAATCG | [] |
| 1634 | 4 | CHIC2 | {'exs': 0, 'nonexs': 7} | True | 54042679 | GGGGATCCAAACTACAAGAGCC | [] |
| 1635 | 4 | CLOCK | {'exs': 0, 'nonexs': 7} | True | 55435414 | GTGCTACTGTGGTTGAACCTTG | [] |
| 1636 | 4 | PDCL2 | {'exs': 0, 'nonexs': 7} | True | 55556556 | CTATTTGGTATCATTATCACTG | [] |
| 1637 | 4 | PPAT | {'exs': 7, 'nonexs': 1} | False | 56395351 | AGCTACCATTCTAATTCTACAG | [] |
| 1638 | 4 | LOC10050 | unavailable | False | 56420148 | ATCTAAATCTGAAACACTTCTG | [] |
| 1639 | 4 | SRP72 | {'exs': 4, 'nonexs': 4} | False | 56493954 | TACTCCTCTATGCCCTGTACAG | ['CCCTGTACAATCACAACAGGTG'] |
| 1640 | 4 | THEGL | {'exs': 1, 'nonexs': 4} | False | 56625866 | AACTCCGGCTATCTGGAAGCTC | [] |
| 1641 | 4 | SPINK2 | {'exs': 1, 'nonexs': 7} | True | 56809984 | TATCCATCACTACACATGGCTG | [] |
| 1642 | 4 | SPINK2 | {'exs': 1, 'nonexs': 7} | True | 56811774 | TGGTAATCTATACTGAGAGCAG | [] |
| 1643 | 4 | POLR2B | {'exs': 9, 'nonexs': 0} | False | 57030987 | ATAGCTAAACACTCATCATTCG | [] |
| 1644 | 4 | ADGRL3 | unavailable | False | 62070907 | TGTCATCTTCTATAGACTAGTG | [] |
| 1645 | 4 | STAP1 | {'exs': 0, 'nonexs': 7} | True | 67589516 | TGGCTAACACGGTGAAACCCC | [] |
| 1646 | 4 | TMPRSS11 | {'exs': 0, 'nonexs': 7} | True | 67822336 | TGTTGCACTAGATCCCAGTTTG | [] |
| 1647 | 4 | TMPRSS11 | {'exs': 0, 'nonexs': 7} | True | 68053908 | CCACACTACATACCAGTCTTTG | [] |
| 1648 | 4 | UGT2B17 | {'exs': 1, 'nonexs': 7} | True | 68537524 | -1 | -1 |
| 1649 | 4 | UGT2B15 | {'exs': 0, 'nonexs': 7} | True | 68647103 | -1 | -1 |
| 1650 | 4 | UGT2B10 | {'exs': 0, 'nonexs': 3} | True | 68830878 | -1 | -1 |
| 1651 | 4 | UGT2A3 | {'exs': 0, 'nonexs': 7} | True | 68929812 | AAGATCTATTCCCTCTTTTCTG | ['AGTTTTATTAAATTTTTGACAG', 'CTTTTCTATCTTTCTAGTTTTG'] |
| 1652 | 4 | UGT2B7 | {'exs': 0, 'nonexs': 7} | True | 69112735 | -1 | -1 |
| 1653 | 4 | UGT2B11 | {'exs': 0, 'nonexs': 7} | True | 69200439 | -1 | -1 |
| 1654 | 4 | UGT2B28 | {'exs': 0, 'nonexs': 7} | True | 69294808 | -1 | -1 |
| 1655 | 4 | UGT2A1 | {'exs': 0, 'nonexs': 5} | True | 69589371 | -1 | -1 |
| 1656 | 4 | UGT2A2 | unavailable | False | 69589371 | -1 | -1 |
| 1657 | 4 | MUC7 | {'exs': 0, 'nonexs': 7} | True | 70481877 | AATATACTATTGCTCCACCATG | [] |
| 1658 | 4 | ENAM | {'exs': 0, 'nonexs': 7} | True | 70644854 | ATAACCCCTAGGCCTGAAGTAG | [] |
| 1659 | 4 | RUFY3 | {'exs': 0, 'nonexs': 7} | True | 70791296 | AAATGCCTAATGATGTTTTGGG | [] |
| 1660 | 4 | GC | {'exs': 1, 'nonexs': 7} | True | 71749175 | CAGGACTACAGGATATTCTTCG | ['TCTTCAATTCAGCATCAATCTG'] |
| 1661 | 4 | AFM | {'exs': 0, 'nonexs': 7} | True | 73499138 | GGGAGCTATCTTCGTGAGCCT | [] |
| 1662 | 4 | AFM | {'exs': 0, 'nonexs': 7} | True | 73501802 | TCAGCTTCACTAAGTTGACAAG | [] |
| 1663 | 4 | RASSF6 | {'exs': 0, 'nonexs': 7} | True | 73576234 | TACTGCTAAACTGTTGTCTCTG | [] |
| 1664 | 4 | PF4V1 | {'exs': 0, 'nonexs': 7} | True | 73854121 | CTAGTAGCTAACTCTCCAAATG | [] |
| 1665 | 4 | PF4 | {'exs': 0, 'nonexs': 7} | True | 73981203 | CTAGTAGCTAACTCTCCAAAAG | [] |
| 1666 | 4 | MTHFD2L | {'exs': 1, 'nonexs': 5} | False | 74241473 | GTCAAGGCTACAGTGAGCCAT | [] |
| 1667 | 4 | MTHFD2L | {'exs': 1, 'nonexs': 5} | False | 74301808 | ATCTAGTAAATGATTTTTTTAG | [] |
| 1668 | 4 | THAP6 | {'exs': 0, 'nonexs': 7} | True | 75539926 | TTGCCATTCCCTATGAAATGCG | ['AAATGCATTTCTCTCTTCTTGG'] |
| 1669 | 4 | USO1 | {'exs': 1, 'nonexs': 0} | False | 75813294 | TTGAAAACTAGATATGATCTAG | [] |
| 1670 | 4 | ART3 | {'exs': 0, 'nonexs': 7} | True | 76112518 | ATCAAACTACAGAGCAACAAAG | [] |
| 1671 | 4 | FAM47E | {'exs': 0, 'nonexs': 7} | True | 76283457 | GATTCTTCTATGCTTGAGTTCG | [] |
| 1672 | 4 | CCNI | {'exs': 1, 'nonexs': 6} | True | 77048218 | TGAAACTACATGACAGAAACAG | [] |
| 1673 | 4 | CCNG2 | {'exs': 1, 'nonexs': 6} | True | 77165923 | ATTTCTAAGATGGAAAGCACAG | [] |
| 1674 | 4 | CNOT6L | {'exs': 0, 'nonexs': 5} | True | 77720430 | TACTCCACTACCTCCGATTAGG | [] |
| 1675 | 4 | FRAS1 | {'exs': 0, 'nonexs': 5} | True | 78445786 | TTTTGAGTCCCTATAAAGAAAG | [] |
| 1676 | 4 | BMP2K | {'exs': 2, 'nonexs': 6} | False | 78912032 | GAAGTATCTACTGTTTAGAAGG | [] |
| 1677 | 4 | NAA11 | {'exs': 0, 'nonexs': 7} | True | 79325187 | ATGCTCTAGGAGGTGGAATCCC | [] |
| 1678 | 4 | ENOPH1 | {'exs': 1, 'nonexs': 7} | True | 82460119 | CCCTTCTCTAGGTTGAGGAAGG | [] |
| 1679 | 4 | SLC10A6 | {'exs': 0, 'nonexs': 7} | True | 86823687 | TCCCTGCTATTCACATGAAGTG | [] |
| 1680 | 4 | NUDT9 | {'exs': 0, 'nonexs': 7} | True | 87458020 | CATCAGCTACAACGCATGGCAC | [] |
| 1681 | 4 | DSPP | {'exs': 0, 'nonexs': 5} | True | 87616567 | TTGTTCTAATCATCACTGGTTG | [] |

| | | | | | | | |
|---|---|---|---|---|---|---|---|
| 1682 | 4 | DMP1 | {'exs': 0, 'nonexs': 7} | True | 87663319 | CTGATGCTAATAGCCGTCTTGG | [] |
| 1683 | 4 | MEPE | {'exs': 0, 'nonexs': 7} | True | 87846645 | CCTGGTGGACTAGTCACCATC( | [] |
| 1684 | 4 | PYURF | unavailable | False | 88521887 | ATTATGAACTAGCGCTGCTCCG | ['AGCGCTGCTCCACTTCTTCTTG'] |
| 1685 | 4 | CCSER1 | unavailable | False | 90731867 | GATGAGTCTAGTCGACCTACAG | [] |
| 1686 | 4 | CCSER1 | unavailable | False | 90780454 | ATTCCCTAGGGTATGTGGTCTG | [] |
| 1687 | 4 | CCSER1 | unavailable | False | 90923411 | TTATCTACTCTTGAAGCAAAAG | [] |
| 1688 | 4 | CCSER1 | unavailable | False | 90938734 | ATTTAACCTAAGTTGAAATAAG | [] |
| 1689 | 4 | CCSER1 | unavailable | False | 91213330 | ACTACATCTCTGTTTTTCCCAG | [] |
| 1690 | 4 | ATOH1 | {'exs': 1, 'nonexs': 7} | True | 93829990 | CCTTCCTAACTTGCCTCATCC( | [] |
| 1691 | 4 | HPGDS | {'exs': 0, 'nonexs': 7} | True | 94299479 | GATCAGCTAGAGTTTGGTTTGG | [] |
| 1692 | 4 | TSPAN5 | {'exs': 0, 'nonexs': 7} | True | 98472521 | GGGGGTCTACCAGCTCGCCCT | [] |
| 1693 | 4 | C4orf17 | {'exs': 0, 'nonexs': 7} | True | 99542108 | AACTTCTACCTCCGGTGTGTTG | [] |
| 1694 | 4 | DAPP1 | {'exs': 0, 'nonexs': 7} | True | 99860082 | TCAGCTTCTACTCCGGTCTTTG | [] |
| 1695 | 4 | DAPP1 | {'exs': 0, 'nonexs': 7} | True | 99869184 | AAGATCTATTTAAAGATGAACG | [] |
| 1696 | 4 | DAPP1 | {'exs': 0, 'nonexs': 7} | True | 99872182 | CCTGCTGGTCTAACTTCTCTGC | [] |
| 1697 | 4 | SLC39A8 | {'exs': 0, 'nonexs': 7} | True | 1.02E+08 | AAATTACTACCTTGCCTGACAG | [] |
| 1698 | 4 | NFKB1 | {'exs': 0, 'nonexs': 7} | True | 1.03E+08 | CAGGCTAAATTTTGCCTTCTAG | [] |
| 1699 | 4 | UBE2D3 | {'exs': 1, 'nonexs': 6} | True | 1.03E+08 | CCTCTTACCCTACAACATAGCG | [] |
| 1700 | 4 | SLC9B2 | unavailable | False | 1.03E+08 | CACCTCTAAACTTGCACAGAAG | [] |
| 1701 | 4 | CENPE | {'exs': 4, 'nonexs': 3} | False | 1.03E+08 | AAGAGGAGTCTACTGAGTTTTG | [] |
| 1702 | 4 | PAPSS1 | {'exs': 1, 'nonexs': 6} | True | 1.08E+08 | AACAGCCTAAGCTTTCTCCAAG | [] |
| 1703 | 4 | OSTC | {'exs': 1, 'nonexs': 6} | True | 1.09E+08 | GGCACTCTAACCCATCAGATAG | [] |
| 1704 | 4 | CFI | {'exs': 0, 'nonexs': 7} | True | 1.1E+08 | CCCAGCTACTCGGGAGGCGGA | [] |
| 1705 | 4 | CFI | {'exs': 0, 'nonexs': 7} | True | 1.1E+08 | TGAACCTAGGAGGTGGAGGTTC | [] |
| 1706 | 4 | CFI | {'exs': 0, 'nonexs': 7} | True | 1.1E+08 | TGTTCTCTACTTTATACGACTG | [] |
| 1707 | 4 | EGF | {'exs': 1, 'nonexs': 6} | True | 1.1E+08 | ACACTGCTACAGCAAATGGGTG | [] |
| 1708 | 4 | ELOVL6 | {'exs': 0, 'nonexs': 7} | True | 1.1E+08 | CAACACTATTCAGCTTTCGTTG | [] |
| 1709 | 4 | AP1AR | {'exs': 1, 'nonexs': 7} | True | 1.12E+08 | TTTTACCCTATCGAGTCTGTTG | [] |
| 1710 | 4 | ALPK1 | {'exs': 1, 'nonexs': 4} | False | 1.12E+08 | CGTATTCTATGTGCCTGGAAGG | [] |
| 1711 | 4 | NEUROG2 | {'exs': 0, 'nonexs': 7} | True | 1.13E+08 | GCAGCTCTAGATACAATCCCTG | [] |
| 1712 | 4 | NDST4 | {'exs': 0, 'nonexs': 7} | True | 1.15E+08 | TCCAGTGCTATCTCACTTTCTG | [] |
| 1713 | 4 | NDST3 | {'exs': 1, 'nonexs': 7} | True | 1.18E+08 | TTTCTTCTATCTCATTTTCTAG | [] |
| 1714 | 4 | NDST3 | {'exs': 1, 'nonexs': 7} | True | 1.18E+08 | CTCAGTGCTATCTTACTTTCTG | [] |
| 1715 | 4 | METTL14 | {'exs': 7, 'nonexs': 2} | False | 1.19E+08 | GCCTTTGGATCTAAAGTCTTAG | [] |
| 1716 | 4 | USP53 | {'exs': 0, 'nonexs': 7} | True | 1.19E+08 | CCTTTTTCACTCTAAGATAGTG | [] |
| 1717 | 4 | PDE5A | {'exs': 0, 'nonexs': 7} | True | 1.2E+08 | AATAACTAGTCGATTCCCACTG | [] |
| 1718 | 4 | NDNF | unavailable | False | 1.21E+08 | GGTAACTAACAGAACTTTCTAG | [] |
| 1719 | 4 | TNIP3 | {'exs': 0, 'nonexs': 7} | True | 1.21E+08 | TTCTACGGATGGACTTTCTTTG | ['CTTTACTGAGGATAAACCTATG'] |
| 1720 | 4 | BBS7 | {'exs': 1, 'nonexs': 7} | True | 1.22E+08 | GAATGGTCCACTAACCATAGAG | [] |
| 1721 | 4 | BBS12 | {'exs': 0, 'nonexs': 7} | True | 1.23E+08 | GTAACACTACAAAAATAGAAAG | [] |
| 1722 | 4 | INTU | {'exs': 1, 'nonexs': 7} | True | 1.28E+08 | CAGCTACAAGGTTAACCCAAAG | [] |
| 1723 | 4 | ABHD18 | unavailable | False | 1.28E+08 | CCAACTAGTTAGCGTATTTATG | [] |
| 1724 | 4 | LARP1B | {'exs': 0, 'nonexs': 7} | True | 1.28E+08 | CCTATCAAATCTAAATAAACTG | [] |
| 1725 | 4 | LARP1B | {'exs': 0, 'nonexs': 7} | True | 1.28E+08 | TAGGAACTAGATGTTTATGATG | [] |
| 1726 | 4 | JADE1 | unavailable | False | 1.29E+08 | AAGTCACTATAAGGTGTCAGTG | [] |
| 1727 | 4 | SCLT1 | {'exs': 0, 'nonexs': 7} | True | 1.29E+08 | ATGTTCTAGAAGAAAACACCTG | [] |
| 1728 | 4 | SCLT1 | {'exs': 0, 'nonexs': 7} | True | 1.29E+08 | CCCAGCTACTGGGGAGGCTGA | [] |
| 1729 | 4 | C4orf33 | {'exs': 0, 'nonexs': 7} | True | 1.29E+08 | AGGAGAACTAGAGAGATAACTG | [] |
| 1730 | 4 | C4orf33 | {'exs': 0, 'nonexs': 7} | True | 1.29E+08 | TCCTATATATCACATTTCTCTG | ['TTCTCTATTAGCCACAGGTCTG'] |
| 1731 | 4 | PCDH10 | {'exs': 0, 'nonexs': 7} | True | 1.33E+08 | CCGGGCTAGAGGAGGCTTTCT( | [] |
| 1732 | 4 | PCDH10 | {'exs': 0, 'nonexs': 7} | True | 1.33E+08 | TTGACTAGCATATCCTTTTCCG | [] |
| 1733 | 4 | PABPC4L | unavailable | False | 1.34E+08 | TTTCTTTCCTAGTGTCTCTGGG | [] |
| 1734 | 4 | PCDH18 | {'exs': 1, 'nonexs': 7} | True | 1.38E+08 | TAAAATCTCCTAGCTCTGGCGG | [] |
| 1735 | 4 | ZNF330 | {'exs': 1, 'nonexs': 6} | True | 1.41E+08 | AGCAGCTCCCCTAGTTCTCTTC | [] |
| 1736 | 4 | HHIP | {'exs': 0, 'nonexs': 7} | True | 1.45E+08 | AGAAACTATACAATGTAACTTG | [] |
| 1737 | 4 | HHIP | {'exs': 0, 'nonexs': 7} | True | 1.45E+08 | TTTCATCTAGGTGATGTCTATG | ['TAAAATAAAATATATAAATATG', 'TCTATATTATAAAAATAAAATG'] |
| 1738 | 4 | ABCE1 | {'exs': 6, 'nonexs': 2} | False | 1.45E+08 | AGTCTAATCATCCAAGAAAAAG | [] |
| 1739 | 4 | ZNF827 | {'exs': 0, 'nonexs': 7} | True | 1.46E+08 | TTCTTCTACTTGGTCATCCTAG | [] |
| 1740 | 4 | POU4F2 | {'exs': 0, 'nonexs': 7} | True | 1.47E+08 | AGAGTCTTCTAAATGCCGGCGC | [] |
| 1741 | 4 | EDNRA | {'exs': 0, 'nonexs': 7} | True | 1.48E+08 | GTGTGGCTAGAATGATGTTTTG | [] |
| 1742 | 4 | ARHGAP10 | {'exs': 0, 'nonexs': 7} | True | 1.48E+08 | CAGGAGCTACAGCAGCTTGAC( | [] |
| 1743 | 4 | IQCM | unavailable | False | 1.49E+08 | TCTAAACAACAAACACAAATAG | [] |
| 1744 | 4 | DCLK2 | {'exs': 1, 'nonexs': 7} | True | 1.5E+08 | GCCGCCTACCTTCGGAATGGT( | [] |
| 1745 | 4 | MAB21L2 | {'exs': 0, 'nonexs': 7} | True | 1.51E+08 | AGCACCCTATAGTTTGTCCAGG | [] |
| 1746 | 4 | SH3D19 | {'exs': 0, 'nonexs': 7} | True | 1.51E+08 | TTCTCCTCTAGCTGATCTGTAG | [] |
| 1747 | 4 | SH3D19 | {'exs': 0, 'nonexs': 7} | True | 1.51E+08 | TTTCAGCCTAATTATGCTTATG | [] |
| 1748 | 4 | FAM160A1 | {'exs': 0, 'nonexs': 5} | True | 1.52E+08 | AAAAAAAAGCTAACAGCAGGAG | [] |
| 1749 | 4 | TLR2 | {'exs': 0, 'nonexs': 7} | True | 1.54E+08 | GGAACCTAGGACTTTATCGCAG | [] |
| 1750 | 4 | RNF175 | {'exs': 0, 'nonexs': 7} | True | 1.54E+08 | CTGTCCTATTCCAGCCCTAGTC | [] |
| 1751 | 4 | SFRP2 | {'exs': 0, 'nonexs': 7} | True | 1.54E+08 | CCGGGACTAGCACTGCAGCTT( | [] |
| 1752 | 4 | FGB | {'exs': 0, 'nonexs': 7} | True | 1.55E+08 | TGGGGACTATTGCTGTGGGAAC | [] |
| 1753 | 4 | FGA | {'exs': 0, 'nonexs': 7} | True | 1.55E+08 | TTCAGCCTATTGGGTCACAAGG | [] |
| 1754 | 4 | FGA | {'exs': 0, 'nonexs': 7} | True | 1.55E+08 | TTAGTCTAGGGGGACAGGGAAC | [] |
| 1755 | 4 | FGG | {'exs': 1, 'nonexs': 7} | True | 1.55E+08 | ATTTTCTACAAATCATCCTCAG | [] |
| 1756 | 4 | RBM46 | {'exs': 0, 'nonexs': 7} | True | 1.55E+08 | ATTCCACTTCTACAATGCAAAG | [] |
| 1757 | 4 | GUCY1A1 | unavailable | False | 1.56E+08 | AATTACTACCTGTAAGTTGTTG | [] |
| 1758 | 4 | GUCY1A1 | unavailable | False | 1.56E+08 | GGTTGCTAATCTATTCCTGATG | [] |
| 1759 | 4 | PDGFC | {'exs': 0, 'nonexs': 7} | True | 1.57E+08 | GTGATGCGGCTATCCTCCTGT( | [] |
| 1760 | 4 | GRIA2 | {'exs': 0, 'nonexs': 7} | True | 1.57E+08 | CCCCTAAATTTTAACACTTTCG | [] |
| 1761 | 4 | PPID | {'exs': 1, 'nonexs': 6} | True | 1.59E+08 | CCTTTCTAAGCAAACATTTTTG | [] |
| 1762 | 4 | NAF1 | {'exs': 4, 'nonexs': 4} | False | 1.63E+08 | CAATGCTATTGCATACTTAATG | ['CTTAATAAACTAAAGTAGCATG'] |
| 1763 | 4 | NAF1 | {'exs': 4, 'nonexs': 4} | False | 1.63E+08 | GCATAGTCACCTAATAGTAAGG | [] |
| 1764 | 4 | KLHL2 | {'exs': 0, 'nonexs': 7} | True | 1.65E+08 | AGGTAACACTATTTGATAAGTG | [] |
| 1765 | 4 | DDX60 | {'exs': 0, 'nonexs': 7} | True | 1.68E+08 | ACAGCTCCTAACTTGAATCTGG | [] |
| 1766 | 4 | DDX60 | {'exs': 0, 'nonexs': 7} | True | 1.68E+08 | TAGCAGTTCTATTGCTTTCCAG | [] |
| 1767 | 4 | GALNTL6 | {'exs': 0, 'nonexs': 7} | True | 1.72E+08 | TTCTTTCTAGGAGTTGGCATGG | [] |
| 1768 | 4 | GALNT7 | {'exs': 1, 'nonexs': 7} | True | 1.73E+08 | TTCTCTCTAAACACTATGGATG | [] |
| 1769 | 4 | SAP30 | {'exs': 0, 'nonexs': 7} | True | 1.73E+08 | CGTCTCCTAGTGAACACCACTC | ['CCACTATCAACCTTGAGATCTG'] |
| 1770 | 4 | CEP44 | unavailable | False | 1.74E+08 | AAATCCTACAAGTAGTGATTTG | [] |
| 1771 | 4 | HPGD | {'exs': 0, 'nonexs': 7} | True | 1.75E+08 | CTCTTTCTGCTAGAGAAAAAGG | [] |
| 1772 | 4 | ADAM29 | {'exs': 0, 'nonexs': 7} | True | 1.75E+08 | TGGCTCTAGGAGGGCGTCACA( | [] |
| 1773 | 4 | CDKN2AIP | {'exs': 2, 'nonexs': 6} | False | 1.83E+08 | CTCTTCTACCCCTTCTAAAAAG | [] |
| 1774 | 4 | ING2 | {'exs': 0, 'nonexs': 7} | True | 1.84E+08 | CCTTTACTACCTCGATCTTCTG | ['CTATCCTTTTTTGTCTTTTCAG'] |
| 1775 | 4 | RWDD4 | {'exs': 0, 'nonexs': 7} | True | 1.84E+08 | AAGTGCTACTCATCATCCTTAG | [] |
| 1776 | 4 | TRAPPC11 | unavailable | False | 1.84E+08 | TGACCTCTAGAAGCTGGAGAAG | [] |
| 1777 | 4 | TRAPPC11 | unavailable | False | 1.84E+08 | GGCAGATCTACAGAGAAAGATG | [] |
| 1778 | 4 | STOX2 | {'exs': 0, 'nonexs': 7} | True | 1.84E+08 | CTCTCCCTACCGTGGGGAGTT( | [] |
| 1779 | 4 | PRIMPOL | unavailable | False | 1.85E+08 | TGAAATTACCTACAGTTCTGTG | [] |
| 1780 | 4 | ACSL1 | {'exs': 1, 'nonexs': 6} | True | 1.85E+08 | TTCACACTAAACCTTGATAGTG | [] |
| 1781 | 4 | SNX25 | {'exs': 0, 'nonexs': 7} | True | 1.85E+08 | TCTCTACTAAAGATACAAAAAG | ['AAAAAAAATTTAGCTGGGTGTG'] |
| 1782 | 4 | LRP2BP | {'exs': 0, 'nonexs': 7} | True | 1.85E+08 | TTGTGGTCTAAATTCTTTGACG | [] |
| 1783 | 4 | C4orf47 | {'exs': 0, 'nonexs': 7} | True | 1.85E+08 | AGTTGCTAATAGGATGGTACTG | [] |
| 1784 | 4 | CCDC110 | {'exs': 1, 'nonexs': 7} | True | 1.85E+08 | GAATCCTAATGATGCTTGAGAG | [] |
| 1785 | 4 | ZFP42 | {'exs': 0, 'nonexs': 7} | True | 1.88E+08 | TGGAGGACTACTTTCCCTCTTG | [] |
| 1786 | 4 | DUX4 | unavailable | False | 1.9E+08 | CCGCGTCCTAAAGCTCCTCCA( | [] |
| 1787 | 4 | DUX4 | unavailable | False | 1.9E+08 | CGGGCCTAGACCTAGAAGGCA( | [] |
| 1788 | 5 | LRRC14B | {'exs': 0, 'nonexs': 7} | True | 195352 | GAGGAACTACTCTATTTGTTTG | [] |
| 1789 | 5 | AHRR | {'exs': 0, 'nonexs': 7} | True | 434833 | ACTGCGCTATGGCAGGAATGTC | [] |
| 1790 | 5 | EXOC3 | {'exs': 3, 'nonexs': 3} | False | 466897 | CGGAGGCTACTTGAGCAGCTTC | [] |
| 1791 | 5 | TPPP | {'exs': 0, 'nonexs': 7} | True | 665101 | GCGGGGGCTACTTGCCCCCTT | [] |
| 1792 | 5 | ZDHHC11B | unavailable | False | 732802 | GCCTCTAATTCCCCTGAGCCTC | [] |
| 1793 | 5 | BRD9 | {'exs': 3, 'nonexs': 4} | False | 875812 | AACTACAGGAAATATTTTTTTG | ['TTTTTTTAAACTCTTCAGGATG'] |
| 1794 | 5 | BRD9 | {'exs': 3, 'nonexs': 4} | False | 883438 | ACTGGGTCTAGAACGTCTCCTC | [] |
| 1795 | 5 | NKD2 | {'exs': 0, 'nonexs': 7} | True | 1038372 | GTGGCGCTAGGACGGGTGGAA | [] |
| 1796 | 5 | SLC6A3 | {'exs': 0, 'nonexs': 7} | True | 1394734 | CTCCCTCTACACCTTGAGCCAC | [] |
| 1797 | 5 | LPCAT1 | {'exs': 0, 'nonexs': 7} | True | 1463650 | TGGGTCCTAATCCAGCTTCTTG | [] |
| 1798 | 5 | MRPL36 | {'exs': 0, 'nonexs': 7} | True | 1798623 | AAAGGGTCTACATCTGTCTCTG | [] |
| 1799 | 5 | NDUFS6 | {'exs': 0, 'nonexs': 7} | True | 1815915 | ACACGCTCTAGTGGTGGTGCTC | [] |
| 1800 | 5 | IRX2 | {'exs': 0, 'nonexs': 7} | True | 2747563 | GCCCTTCTATAGGTAGGGCTGC | [] |
| 1801 | 5 | C5orf38 | {'exs': 0, 'nonexs': 7} | True | 2753358 | CGCGCCTACCGGGAGCCTGCC | [] |
| 1802 | 5 | C5orf38 | {'exs': 0, 'nonexs': 7} | True | 2755040 | CTCGTTCTAAATTCTTTGGCTG | [] |
| 1803 | 5 | IRX1 | {'exs': 0, 'nonexs': 7} | True | 3601019 | AGGGCTGCTAGGATCCGCGGC | [] |
| 1804 | 5 | CMBL | {'exs': 0, 'nonexs': 7} | True | 10280452 | CTTGCTACATGTACTTGTTCAG | [] |
| 1805 | 5 | LOC105369 | unavailable | False | 10464594 | CATTCTATTTTCTTGTGCAAAG | [] |
| 1806 | 5 | DNAH5 | {'exs': 0, 'nonexs': 7} | True | 13794048 | TAGCTCTCTATCGTCCTCTGTG | [] |
| 1807 | 5 | H3Y2 | unavailable | False | 17491324 | AGAAGTCCTCTAGAGTGCAAGG | [] |
| 1808 | 5 | CDH18 | {'exs': 1, 'nonexs': 4} | False | 19473225 | ACCCCCTAAGTTGTTCTTTCAG | [] |
| 1809 | 5 | CDH6 | {'exs': 0, 'nonexs': 7} | True | 31318033 | CTCACCTAGATCAGGCTGTATG | [] |
| 1810 | 5 | SLC45A2 | {'exs': 1, 'nonexs': 7} | True | 33944547 | CTAATCCACATATCTAACAAAG | [] |
| 1811 | 5 | SLC45A2 | {'exs': 1, 'nonexs': 7} | True | 33951651 | GTGCACTATAGGGATCCCCGC( | [] |
| 1812 | 5 | TTC23L | {'exs': 0, 'nonexs': 7} | True | 34896777 | GAACTGCTACTTGTCCTTTAAG | [] |
| 1813 | 5 | DNAJC21 | {'exs': 1, 'nonexs': 7} | True | 34954713 | AGAATCTCTATCTGTTTTTACG | [] |
| 1814 | 5 | PRLR | {'exs': 2, 'nonexs': 6} | False | 35049286 | CTTCAGCTCTACTGGACTGTGC | [] |
| 1815 | 5 | SKP2 | {'exs': 5, 'nonexs': 4} | False | 36178612 | AAGCTCCTAGGAGTTCTGGATG | [] |
| 1816 | 5 | RANBP3L | {'exs': 0, 'nonexs': 7} | True | 36248540 | TTATCCCCTAATCAGATAATTG | [] |
| 1817 | 5 | RANBP3L | {'exs': 0, 'nonexs': 7} | True | 36251304 | CAAAACAAAAGCTATTTACCTG | [] |
| 1818 | 5 | SLC1A3 | {'exs': 0, 'nonexs': 7} | True | 36686268 | AGTCTACATCTTGGTTTCACTG | [] |
| 1819 | 5 | OSMR | {'exs': 0, 'nonexs': 7} | True | 38844232 | AATCCAGCTATCTTCTATTAAG | [] |
| 1820 | 5 | FYB1 | unavailable | False | 39107442 | TGAGTGCTAGTCATTGTCATAG | [] |
| 1821 | 5 | C9 | {'exs': 0, 'nonexs': 7} | True | 39285198 | ACAGCTCTATTTTTCATTGGGG | [] |
| 1822 | 5 | C7 | {'exs': 1, 'nonexs': 7} | True | 40881572 | GGAGCCTACTGGGTTTCCGCA( | [] |
| 1823 | 5 | C6 | {'exs': 2, 'nonexs': 7} | False | 41142824 | TGCTAGGCCAAACACTTTCCAG | [] |
| 1824 | 5 | C5orf51 | {'exs': 0, 'nonexs': 7} | True | 41917298 | GAACTTACTAGTTACAGCGATG | [] |
| 1825 | 5 | GHR | {'exs': 0, 'nonexs': 7} | True | 42719423 | GAAAGGCTAAGGCATGATTTTG | [] |
| 1826 | 5 | ANXA2R | unavailable | False | 43039454 | GCTGCTATCTAAGGCTGCTTAG | [] |
| 1827 | 5 | CCL28 | {'exs': 0, 'nonexs': 7} | True | 43381859 | CTCTCTAATAAGGAGTTTTATG | [] |
| 1828 | 5 | FGF10 | {'exs': 1, 'nonexs': 6} | True | 44304994 | CTCTATGAGTGTACCACCATTG | [] |
| 1829 | 5 | MOCS2 | {'exs': 0, 'nonexs': 7} | True | 53107094 | AAAGCACTATCCTCCACTAATG | [] |
| 1830 | 5 | FST | {'exs': 1, 'nonexs': 7} | True | 53485949 | ATGGCTAGATGGGGGAATACAG | [] |
| 1831 | 5 | NDUFS4 | {'exs': 1, 'nonexs': 7} | True | 53683220 | ACCTATTTTGTGGATACTCTTG | [] |
| 1832 | 5 | SNX18 | {'exs': 0, 'nonexs': 7} | True | 54519838 | AAACTCTAAAATGATAGCACTG | [] |
| 1833 | 5 | SNX18 | {'exs': 0, 'nonexs': 7} | True | 54606406 | GGACTAACCTGGAAATACAAAG | [] |
| 1834 | 5 | CDC20B | {'exs': 0, 'nonexs': 7} | True | 55114217 | TGCTAGTAGCAATTCCATACAG | [] |
| 1835 | 5 | MTREX | unavailable | False | 55424771 | CTGACTCTACAAGTAGAGGCTG | [] |
| 1836 | 5 | MAP3K1 | {'exs': 2, 'nonexs': 6} | False | 56893679 | TAATTGGCTACCATGTAGTACG | [] |
| 1837 | 5 | GAPT | {'exs': 0, 'nonexs': 5} | True | 58495009 | TGAAAAGCTAATATGAATCTGG | [] |
| 1838 | 5 | RAB3C | {'exs': 0, 'nonexs': 7} | True | 58851350 | GGGACACTAGCAGGCACAGTT( | [] |
| 1839 | 5 | ELOVL7 | {'exs': 1, 'nonexs': 5} | False | 60836347 | CTATCTTCTTTCTTTCTTCAAG | [] |
| 1840 | 5 | DIMT1 | unavailable | False | 62389009 | TTCCAAATACCTAGGAAAAATG | [] |
| 1841 | 5 | RNF180 | {'exs': 0, 'nonexs': 7} | True | 64369813 | ATGAAATTCCTAAAACGGAAAG | [] |
| 1842 | 5 | RGS7BP | {'exs': 0, 'nonexs': 7} | True | 64609251 | GAGCCACCTAGCTTGAGATGAC | [] |
| 1843 | 5 | CWC27 | {'exs': 0, 'nonexs': 7} | True | 64844985 | TTCTGCTATTTGAGATCTGCAG | [] |
| 1844 | 5 | ADAMTS6 | {'exs': 0, 'nonexs': 7} | True | 65214900 | TATACCTAATTCCCAAATTCTG | [] |
| 1845 | 5 | ADAMTS6 | {'exs': 0, 'nonexs': 7} | True | 65246095 | CAATCTATATAAACATCTACAG | [] |
| 1846 | 5 | CENPK | {'exs': 2, 'nonexs': 4} | False | 65508120 | TCTCCACTATAAAGGTACTATG | ['GTACTATATTTCCCTGTGTAAG'] |
| 1847 | 5 | SREK1 | {'exs': 4, 'nonexs': 5} | False | 66178867 | TGTCGGTCCTATACTGCTTCTG | [] |
| 1848 | 5 | CD180 | {'exs': 0, 'nonexs': 7} | True | 67182856 | CCTTCAGCACTAAATGTGTTGG | [] |
| 1849 | 5 | CCDC125 | {'exs': 0, 'nonexs': 1} | False | 69285341 | CCAAATCTATGAGCATCTTAAG | [] |
| 1850 | 5 | RAD17 | {'exs': 6, 'nonexs': 1} | False | 69414291 | CTTCTATGTCCCATCACTCTCG | [] |
| 1851 | 5 | OCLN | {'exs': 0, 'nonexs': 7} | True | 69553670 | CATCAGCCTTCTATGTTTTCTG | [] |
| 1852 | 5 | GTF2H2C | {'exs': 3, 'nonexs': 4} | False | 69592210 | TACAATGTATACTACATGCTGG | [] |
| 1853 | 5 | SERF1B | {'exs': 2, 'nonexs': 5} | False | 70041739 | AACCTAAATCCAACAGCAAAAG | [] |
| 1854 | 5 | SMN2 | {'exs': 4, 'nonexs': 3} | False | 70077032 | TGCTCTATGCCAGCATTTCCTG | [] |
| 1855 | 5 | SERF1A | {'exs': 2, 'nonexs': 5} | False | 70917156 | AACCTAAATCCAACAGCAAAAG | [] |
| 1856 | 5 | SMN1 | {'exs': 5, 'nonexs': 3} | False | 70952452 | TGCTCTATGCCAGCATTTCCTG | [] |
| 1857 | 5 | GTF2H2 | {'exs': 3, 'nonexs': 4} | False | 71035663 | TACAATGTATACTACATGCTGG | [] |
| 1858 | 5 | GTF2H2 | {'exs': 3, 'nonexs': 4} | False | 71038429 | TACTCTAATAAGTGCTACTTTG | [] |
| 1859 | 5 | TMEM174 | {'exs': 1, 'nonexs': 7} | True | 73174164 | TAGCCTCTAGCGAGGGAGAGA( | [] |
| 1860 | 5 | ANKRA2 | {'exs': 1, 'nonexs': 7} | True | 73552796 | GTGTCTACTCCTTGATATTTTG | [] |
| 1861 | 5 | UTP15 | {'exs': 8, 'nonexs': 1} | False | 73580093 | ACACTATGATTCTATCTTCTTG | ['TTATTCTCTGGAAATCCATCAG'] |
| 1862 | 5 | GFM2 | {'exs': 2, 'nonexs': 6} | False | 74730667 | GACTAATGATGCCAACACTTAG | [] |
| 1863 | 5 | GFM2 | {'exs': 2, 'nonexs': 6} | False | 74736763 | ACTAATTAGTTGACACACTAAG | [] |
| 1864 | 5 | NSA2 | {'exs': 7, 'nonexs': 1} | False | 74772371 | AGGCTAAGGCGGGCGGATCAC | [] |
| 1865 | 5 | CERT1 | unavailable | False | 75374042 | GCTTTGCTATTGCAATATTTTG | [] |
| 1866 | 5 | CERT1 | unavailable | False | 75376345 | GTTAATACTAGAACAAAATAGG | [] |
| 1867 | 5 | F2RL2 | {'exs': 0, 'nonexs': 7} | True | 76617581 | TTTCACTATTTTGTAAGGTAAG | [] |
| 1868 | 5 | F2R | {'exs': 1, 'nonexs': 6} | True | 76733502 | TTCCTAAGTTAACAGCTTTTTG | [] |
| 1869 | 5 | ZBED3 | {'exs': 1, 'nonexs': 7} | True | 77077173 | CACCCCTACAGGAGGACCTTT( | [] |
| 1870 | 5 | WDR41 | {'exs': 0, 'nonexs': 7} | True | 77433134 | AAACTAGACAGCAAGGTATAAG | [] |
| 1871 | 5 | LHFPL2 | {'exs': 0, 'nonexs': 5} | True | 78488896 | CAAACTAAAGGAGGCAGATCAG | [] |
| 1872 | 5 | ARSB | {'exs': 0, 'nonexs': 7} | True | 78780396 | CCTGAAATCCTACATCCAAGGG | [] |
| 1873 | 5 | ARSB | {'exs': 0, 'nonexs': 7} | True | 78816085 | TGGTAGCTACAACAGCAGCGAC | [] |
| 1874 | 5 | DMGDH | {'exs': 0, 'nonexs': 7} | True | 79021606 | TTCACCCTAATAAGGCCTGATG | [] |
| 1875 | 5 | BHMT | {'exs': 0, 'nonexs': 7} | True | 79131115 | CTATCGAGGCTACTGTGATTTG | [] |
| 1876 | 5 | HOMER1 | {'exs': 1, 'nonexs': 7} | True | 79425194 | CCAAAACTAAAGATGATCAGAG | [] |
| 1877 | 5 | TENT2 | unavailable | False | 79646232 | CTAACACATATTTTCTCCATAG | [] |
| 1878 | 5 | THBS4 | {'exs': 0, 'nonexs': 7} | True | 80074404 | CAGGTGCCGCTAGGGCTGAGT( | [] |
| 1879 | 5 | SPZ1 | {'exs': 0, 'nonexs': 7} | True | 80321507 | ATTTTCTATCTTAGGCTGCTAG | [] |
| 1880 | 5 | ZFYVE16 | {'exs': 1, 'nonexs': 7} | True | 80477376 | CATTCTTTCACTAAAAAAGATG | [] |
| 1881 | 5 | ANKRD34B | {'exs': 0, 'nonexs': 7} | True | 80559474 | TCTCCTAGAAGTTTACTAATTG | [] |
| 1882 | 5 | RASGRF2 | {'exs': 1, 'nonexs': 7} | True | 81096169 | TCTAGCTATTTATTTGCCATTG | [] |
| 1883 | 5 | RASGRF2 | {'exs': 1, 'nonexs': 7} | True | 81183200 | CTGTTTCTAGAAATCGAAGTGG | [] |
| 1884 | 5 | ACOT12 | {'exs': 0, 'nonexs': 7} | True | 81310974 | TTTTCCTACTTCTCAAGGCCAG | [] |
| 1885 | 5 | ATP6AP1L | {'exs': 0, 'nonexs': 5} | True | 82312579 | CTCCTAAAGAACTTTTCCCACG | [] |
| 1886 | 5 | ATP6AP1L | {'exs': 0, 'nonexs': 5} | True | 82318299 | GTGCTGCTAAACATAGATTTTG | [] |
| 1887 | 5 | TMEM167A | {'exs': 2, 'nonexs': 5} | False | 83057083 | CCCCAGCTACTGTATGAAGAGC | [] |
| 1888 | 5 | XRCC4 | {'exs': 0, 'nonexs': 7} | True | 83210719 | TTGACTATCTTTCTTGAGAATG | [] |
| 1889 | 5 | RASA1 | {'exs': 1, 'nonexs': 6} | True | 87390882 | AGGCTGCTACCTGACATCATTG | [] |
| 1890 | 5 | TMEM161B | {'exs': 0, 'nonexs': 7} | True | 88189422 | GCCTTCTAGGTTTCAGGTGCTC | [] |
| 1891 | 5 | TMEM161B | {'exs': 0, 'nonexs': 7} | True | 88189550 | AGAGGCTACGTTCCAGTCCTTC | [] |
| 1892 | 5 | MBLAC2 | {'exs': 0, 'nonexs': 7} | True | 90461166 | AGATACTAGGGCGAGGTCCTAC | [] |
| 1893 | 5 | POLR3G | {'exs': 0, 'nonexs': 5} | True | 90512138 | AAATTTCATGCCTAATAGGTTG | [] |
| 1894 | 5 | LYSMD3 | {'exs': 0, 'nonexs': 1} | False | 90518818 | AATTGCTATGTTTCCTGTTGAG | [] |
| 1895 | 5 | ADGRV1 | unavailable | False | 91163899 | GAGGTGCTACAGGTGAGTGTC( | [] |
| 1896 | 5 | NR2F1 | {'exs': 2, 'nonexs': 6} | False | 93593841 | CAAGGTCTAGGAGCACTGGATC | [] |
| 1897 | 5 | FAM172A | {'exs': 0, 'nonexs': 7} | True | 93743513 | GCTGGTATCTATTCTCTTGAGG | [] |
| 1898 | 5 | POU5F2 | {'exs': 0, 'nonexs': 4} | True | 93740576 | TCAGCCCTAAAATCTGAGGAGG | [] |
| 1899 | 5 | KIAA0825 | unavailable | False | 94520242 | AATGTCACTAACCCAAAGCATG | [] |
| 1900 | 5 | MCTP1 | {'exs': 0, 'nonexs': 7} | True | 94707495 | GGGAGCTGGCTAGCCAAGATT( | [] |
| 1901 | 5 | GPR150 | {'exs': 0, 'nonexs': 7} | True | 95621579 | AAGCACCTAGAAGGCACTTTCG | [] |
| 1902 | 5 | ELL2 | {'exs': 0, 'nonexs': 7} | True | 95888870 | GAGTTCTAGGACCATGACTCTG | [] |
| 1903 | 5 | ELL2 | {'exs': 0, 'nonexs': 7} | True | 95890398 | ACCACTATACCACCTGTGTCCC | [] |
| 1904 | 5 | LNPEP | {'exs': 0, 'nonexs': 7} | True | 97028532 | TGCATGCTACAGCCACCATGTC | [] |
| 1905 | 5 | LIX1 | {'exs': 0, 'nonexs': 7} | True | 97094747 | CTTGCTAGTGATAGCCACACAG | [] |
| 1906 | 5 | RGMB | {'exs': 0, 'nonexs': 7} | True | 98793752 | ACCCCTACAAAAACACGATAAG | [] |
| 1907 | 5 | GIN1 | {'exs': 0, 'nonexs': 7} | True | 1.03E+08 | TTTACTAACTTAAGTATTCAAG | [] |
| 1908 | 5 | EFNA5 | {'exs': 3, 'nonexs': 5} | False | 1.07E+08 | GACTGTGCTATAATGTCAAAAG | [] |
| 1909 | 5 | FBXL17 | {'exs': 1, 'nonexs': 4} | False | 1.08E+08 | GAGCGCTAGGAGGAGGCGGCA | [] |
| 1910 | 5 | FBXL17 | {'exs': 1, 'nonexs': 4} | False | 1.08E+08 | TATACCTAAAAAAACAAAAGAG | [] |
| 1911 | 5 | FBXL17 | {'exs': 1, 'nonexs': 4} | False | 1.08E+08 | CAAATCTACCTAATCAGTAGTG | [] |
| 1912 | 5 | FER | {'exs': 1, 'nonexs': 6} | True | 1.09E+08 | CTATTTTGCATTTACAAACAAG | [] |
| 1913 | 5 | FER | {'exs': 1, 'nonexs': 6} | True | 1.09E+08 | TGACTTCTAAATGCTTCCTTTG | [] |
| 1914 | 5 | FER | {'exs': 1, 'nonexs': 6} | True | 1.09E+08 | TCACTATGTGAGTTTTCTCTTG | [] |
| 1915 | 5 | TMEM232 | {'exs': 0, 'nonexs': 7} | True | 1.11E+08 | TCCTCTACTCAGGTTTGCCTAG | [] |
| 1916 | 5 | SRP19 | {'exs': 6, 'nonexs': 2} | False | 1.13E+08 | TTACCTAGGAACCAGAAGTTAG | [] |
| 1917 | 5 | REEP5 | {'exs': 0, 'nonexs': 7} | True | 1.13E+08 | AATCGCTGCCTACCAAGGAAAG | [] |
| 1918 | 5 | YTHDC2 | {'exs': 0, 'nonexs': 7} | True | 1.14E+08 | CCACAGCTAAGAGTTCTAGAAG | [] |
| 1919 | 5 | KCNN2 | {'exs': 0, 'nonexs': 7} | True | 1.14E+08 | CTCTTCTAGCTACTCTCTGATG | [] |
| 1920 | 5 | TRIM36 | {'exs': 1, 'nonexs': 7} | True | 1.15E+08 | TTTCAACTACATGTCCTCTTGG | [] |
| 1921 | 5 | TRIM36 | {'exs': 1, 'nonexs': 7} | True | 1.15E+08 | ATTGTCCTAAGTTCTTCTTGGG | [] |
| 1922 | 5 | AP3S1 | {'exs': 0, 'nonexs': 7} | True | 1.16E+08 | CCTTGTCTACCTGTCCTGTCTC | [] |
| 1923 | 5 | LVRN | unavailable | False | 1.16E+08 | CAAGCTATGTGTTTCTCCTTAG | [] |
| 1924 | 5 | ARL14EPL | {'exs': 0, 'nonexs': 4} | True | 1.16E+08 | ACCTAGTCAGGAACATTAAACG | [] |
| 1925 | 5 | DTWD2 | {'exs': 0, 'nonexs': 7} | True | 1.19E+08 | TACTAAATTTTAACACTATTCG | ['CATTAACAATTCCATTTTCCTG'] |
| 1926 | 5 | LOX | {'exs': 0, 'nonexs': 7} | True | 1.22E+08 | GCCTTCTAATACCTAGATTAAG | [] |
| 1927 | 5 | ZNF474 | {'exs': 0, 'nonexs': 7} | True | 1.22E+08 | TTGTTCCCCTACAGGCAGAGGC | [] |
| 1928 | 5 | SNCAIP | {'exs': 1, 'nonexs': 7} | True | 1.22E+08 | TGGGCTCTAGTAACTGGATTTG | [] |
| 1929 | 5 | SNX2 | {'exs': 2, 'nonexs': 7} | False | 1.23E+08 | TTATTGCTAGGCAATGGCTTTG | [] |
| 1930 | 5 | SNX24 | {'exs': 0, 'nonexs': 7} | True | 1.23E+08 | AGGATTTCTACCTGGGCTGTAG | [] |
| 1931 | 5 | GRAMD2B | unavailable | False | 1.26E+08 | CCTGACCTAGGGCAACCAACAC | [] |
| 1932 | 5 | MARCHF3 | unavailable | False | 1.27E+08 | GTTACCTACTGGAGACAACAAG | [] |
| 1933 | 5 | CTXN3 | {'exs': 0, 'nonexs': 7} | True | 1.28E+08 | TCCCTCCTAAGCAAGGGCATGC | [] |
| 1934 | 5 | CCDC192 | unavailable | False | 1.28E+08 | AGCCAGTGCTAACTACCTAAGG | [] |
| 1935 | 5 | CCDC192 | unavailable | False | 1.28E+08 | GGAATCTACAAATTCTCATCAG | [] |
| 1936 | 5 | ISOC1 | {'exs': 0, 'nonexs': 7} | True | 1.29E+08 | AAATGTCCTATACTTTGGAAAG | [] |
| 1937 | 5 | CHSY3 | {'exs': 1, 'nonexs': 7} | True | 1.3E+08 | CCTACTAACTCTGGAGGAAAAG | [] |
| 1938 | 5 | CHSY3 | {'exs': 1, 'nonexs': 7} | True | 1.3E+08 | TGGAGAAGCTACAAAATCTTGG | [] |
| 1939 | 5 | LYRM7 | {'exs': 1, 'nonexs': 7} | True | 1.31E+08 | TCTACAAGAAGGTCTTTCCTAG | [] |
| 1940 | 5 | CDC42SE2 | {'exs': 0, 'nonexs': 7} | True | 1.31E+08 | GACCAGGGCTATCCCGCCTTC | [] |
| 1941 | 5 | RAPGEF6 | {'exs': 2, 'nonexs': 5} | False | 1.31E+08 | AAGGCTAGACTGCTGAAACTTG | [] |
| 1942 | 5 | IRF1 | {'exs': 1, 'nonexs': 6} | True | 1.32E+08 | GGCCCTGCTACGGTGCACAGG | [] |
| 1943 | 5 | CCNI2 | {'exs': 0, 'nonexs': 7} | True | 1.33E+08 | CAGAGGGGCTAGTTGGCAGGT( | [] |
| 1944 | 5 | SHROOM1 | {'exs': 0, 'nonexs': 7} | True | 1.33E+08 | TATAACTATGTAGGAGAAGAG | [] |
| 1945 | 5 | SHROOM1 | {'exs': 0, 'nonexs': 7} | True | 1.33E+08 | CCCTGGCTAGGGCAGTATTGTC | [] |
| 1946 | 5 | ZCCHC10 | {'exs': 0, 'nonexs': 7} | True | 1.33E+08 | -1 | -1 |
| 1947 | 5 | TCF7 | {'exs': 2, 'nonexs': 7} | False | 1.34E+08 | CGCAATGCCTATGGCTTCCTTC | [] |
| 1948 | 5 | TCF7 | {'exs': 2, 'nonexs': 7} | False | 1.34E+08 | TCCTCCTAGGAATAGTGAAGAG | [] |
| 1949 | 5 | TCF7 | {'exs': 2, 'nonexs': 7} | False | 1.34E+08 | GCAGCCTAGAGCACTGTCATC( | [] |
| 1950 | 5 | CDKL3 | {'exs': 3, 'nonexs': 6} | False | 1.34E+08 | ACACTACCAGAAAAAAAACCTG | [] |
| 1951 | 5 | CDKL3 | {'exs': 3, 'nonexs': 6} | False | 1.34E+08 | GCATTTTCTACTCTACATCTGG | [] |
| 1952 | 5 | DDX46 | {'exs': 8, 'nonexs': 1} | False | 1.35E+08 | TACCTACCTTTGATGGTTTAAG | [] |
| 1953 | 5 | DDX46 | {'exs': 8, 'nonexs': 1} | False | 1.35E+08 | TCCGGATGTCTATAAGACTTTG | [] |
| 1954 | 5 | TXNDC15 | {'exs': 0, 'nonexs': 7} | True | 1.35E+08 | ATCACTACTCCACATGTTCCTG | [] |
| 1955 | 5 | CATSPER3 | {'exs': 0, 'nonexs': 7} | True | 1.35E+08 | ATGCCCAGCTACTTCTCATCCC | ['CATCCACCTTTTCCAAGGACTG'] |
| 1956 | 5 | MACROH2A1 | unavailable | False | 1.35E+08 | GCTCAGCCTAGTTGGCGTCCA( | [] |
| 1957 | 5 | TIFAB | {'exs': 0, 'nonexs': 6} | True | 1.35E+08 | GATCTGCTACCCTGAACCAGG( | [] |
| 1958 | 5 | NEUROG1 | {'exs': 1, 'nonexs': 6} | True | 1.36E+08 | AAGGGCCTAGTGGTAAGGAATG | [] |
| 1959 | 5 | CXCL14 | {'exs': 0, 'nonexs': 5} | True | 1.36E+08 | ACCCTATTCTTCGTAGACCCTG | [] |
| 1960 | 5 | SLC25A48 | {'exs': 0, 'nonexs': 7} | True | 1.36E+08 | AGCACTATGTCTTCTTTCCTTG | [] |
| 1961 | 5 | TGFBI | {'exs': 0, 'nonexs': 7} | True | 1.36E+08 | AGCTAATGCTTCATCCTCTCTG | ['CCTCTCTAATAACTTTTGATAG'] |
| 1962 | 5 | SMIM32 | unavailable | False | 1.36E+08 | GTCCCAAGCTACACCGGCCGC | [] |
| 1963 | 5 | TRPC7 | {'exs': 0, 'nonexs': 1} | False | 1.36E+08 | CTGCTAAATGTCTTTGCCCTTG | [] |
| 1964 | 5 | SPOCK1 | {'exs': 1, 'nonexs': 4} | False | 1.37E+08 | TGGGCACTACCATATGTACCCC | [] |
| 1965 | 5 | PKD2L2 | {'exs': 0, 'nonexs': 7} | True | 1.38E+08 | TTGTCTCAATACTATAGAGCTG | [] |
| 1966 | 5 | WNT8A | {'exs': 0, 'nonexs': 7} | True | 1.38E+08 | CTTCTATTCTCCTGATTCCCAG | [] |
| 1967 | 5 | NME5 | {'exs': 1, 'nonexs': 6} | True | 1.38E+08 | GCAATCTACAGATTCAGTGCAG | [] |
| 1968 | 5 | CDC23 | {'exs': 8, 'nonexs': 0} | False | 1.38E+08 | GCCAACTATGGCGTGACAGAAC | [] |
| 1969 | 5 | GFRA3 | {'exs': 0, 'nonexs': 7} | True | 1.38E+08 | GTCCAGCTACCATAGGCTCAGC | [] |
| 1970 | 5 | KDM3B | {'exs': 2, 'nonexs': 6} | False | 1.38E+08 | TCCATGCCTAGGACCTTGCCA( | [] |
| 1971 | 5 | ETF1 | {'exs': 7, 'nonexs': 1} | False | 1.39E+08 | GACTACCTAGTAGTCATCAAGG | [] |
| 1972 | 5 | MZB1 | unavailable | False | 1.39E+08 | CAGGACTAGAGCTCTTCTCTTG | [] |
| 1973 | 5 | SPATA24 | {'exs': 1, 'nonexs': 7} | True | 1.39E+08 | ACAAGTCTATTCTGGACCAGAG | [] |
| 1974 | 5 | SMIM33 | unavailable | False | 1.39E+08 | GTCTCCCTCCTACAGACAAGTC | [] |
| 1975 | 5 | PSD2 | {'exs': 0, 'nonexs': 7} | True | 1.4E+08 | GTCAGCTAAGTATCAGGCCCAC | [] |
| 1976 | 5 | NRG2 | {'exs': 0, 'nonexs': 7} | True | 1.4E+08 | GGGGCCCTAGAGTGGCGCCGA | [] |
| 1977 | 5 | NRG2 | {'exs': 0, 'nonexs': 7} | True | 1.4E+08 | TTCCTAAGTTCTTTCCTTTTTG | ['TCCTTTTTATTCCATTCAGGTG'] |
| 1978 | 5 | PFDN1 | {'exs': 5, 'nonexs': 4} | False | 1.4E+08 | GCTCCCTACTGGGCCCTTCGT | [] |
| 1979 | 5 | PFDN1 | {'exs': 5, 'nonexs': 4} | False | 1.4E+08 | GCATGTGCCTAAAATATAACAG | [] |
| 1980 | 5 | SLC4A9 | {'exs': 1, 'nonexs': 7} | True | 1.4E+08 | CTCCAGCTAATTCACAGAAATG | [] |
| 1981 | 5 | ANKHD1 | {'exs': 1, 'nonexs': 6} | True | 1.4E+08 | ACAACATCTATTCTTTACGTAG | [] |
| 1982 | 5 | SLC35A4 | {'exs': 1, 'nonexs': 7} | True | 1.41E+08 | CAGGGACTAGCGGCTGCCATA( | [] |
| 1983 | 5 | LOC112267 | unavailable | False | 1.41E+08 | CAGGCCTAAGGACAGATCTCAC | [] |
| 1984 | 5 | TMCO6 | {'exs': 1, 'nonexs': 7} | True | 1.41E+08 | CAGAAACCTATACCTCTGGCTG | [] |
| 1985 | 5 | PCDHA2 | {'exs': 0, 'nonexs': 1} | False | 1.41E+08 | AACTATGAATCTAAATAACTAG | [] |
| 1986 | 5 | PCDHA5 | {'exs': 0, 'nonexs': 1} | False | 1.41E+08 | GTGTACTAAGACATAATTTTTG | ['ATTTTTAAATACAAAAAAAGTG'] |
| 1987 | 5 | PCDHA8 | {'exs': 0, 'nonexs': 1} | False | 1.41E+08 | TAACTACAATTTCCACAAATAG | [] |
| 1988 | 5 | PCDHA13 | {'exs': 0, 'nonexs': 3} | True | 1.41E+08 | TTACTAAGACAATTTTTTTAAAG | ['ATATAAGCTTACCTCTTTCAAG'] |
| 1989 | 5 | PCDHB1 | {'exs': 0, 'nonexs': 3} | True | 1.41E+08 | AGGAGCTACATATAGTCATCAG | [] |
| 1990 | 5 | PCDHB5 | {'exs': 0, 'nonexs': 3} | True | 1.41E+08 | AGATCTCTAATTTAATCCAAAG | [] |
| 1991 | 5 | PCDHB16 | {'exs': 0, 'nonexs': 3} | True | 1.41E+08 | AGTGCCCTAAGGAGAGAAGTTG | [] |
| 1992 | 5 | PCDHB11 | {'exs': 0, 'nonexs': 3} | True | 1.41E+08 | TTACTAAAAATTAAATCCAAAG | [] |
| 1993 | 5 | PCDHGA4 | unavailable | False | 1.41E+08 | CTTTGTCTATATTTCAGTGTGG | [] |
| 1994 | 5 | PCDHGB6 | {'exs': 0, 'nonexs': 1} | False | 1.41E+08 | TGAGAAACTAGTAGACTATTAG | [] |
| 1995 | 5 | DIAPH1 | {'exs': 1, 'nonexs': 4} | False | 1.42E+08 | GAAGCTAGCAGAGATGTGACTG | [] |
| 1996 | 5 | FCHSD1 | {'exs': 0, 'nonexs': 7} | True | 1.42E+08 | CTAGAAGCCCAATCTCCACCAC | [] |
| 1997 | 5 | FCHSD1 | {'exs': 0, 'nonexs': 7} | True | 1.42E+08 | GTTCTATGCTTGGAGCCTCCC( | [] |
| 1998 | 5 | RNF14 | {'exs': 2, 'nonexs': 6} | False | 1.42E+08 | AGTTAACTAGTCTTCTACCTCG | ['TACCTCATCTTCCCAAATATCG'] |
| 1999 | 5 | GNPDA1 | {'exs': 1, 'nonexs': 7} | True | 1.42E+08 | GCACAGGCTAATCGCTGTATGC | [] |
| 2000 | 5 | NR3C1 | {'exs': 1, 'nonexs': 6} | True | 1.43E+08 | GGTGTTCTACCAACCTGAAGAG | [] |
| 2001 | 5 | PRELID2 | {'exs': 0, 'nonexs': 7} | True | 1.46E+08 | ATCAAAACTCTAAGCACAGCAG | [] |
| 2002 | 5 | GRXCR2 | {'exs': 0, 'nonexs': 7} | True | 1.46E+08 | CGGGCTATTGATTGCAAATCTG | [] |
| 2003 | 5 | GPR151 | {'exs': 0, 'nonexs': 7} | True | 1.47E+08 | ATCTATTTAACACCTTCCCCTG | [] |
| 2004 | 5 | PPP2R2B | {'exs': 1, 'nonexs': 6} | True | 1.47E+08 | CCACCTAGTTAACCTTGTCCTG | [] |
| 2005 | 5 | STK32A | {'exs': 1, 'nonexs': 6} | True | 1.47E+08 | TTCTCAACCTACTGTTTCTGGG | [] |
| 2006 | 5 | SPINK14 | {'exs': 0, 'nonexs': 7} | True | 1.48E+08 | TCAGCTAACATTTTCCATCATG | [] |
| 2007 | 5 | HTR4 | {'exs': 0, 'nonexs': 7} | True | 1.48E+08 | GAGGCTATTTTCCTTCCCTAAG | ['AACATGACTTTCTCCCCAACAG'] |
| 2008 | 5 | HTR4 | {'exs': 0, 'nonexs': 7} | True | 1.48E+08 | CAGGGGCCTAAGTGTCACTGG( | [] |
| 2009 | 5 | ABLIM3 | {'exs': 0, 'nonexs': 7} | True | 1.49E+08 | CTGCCTAGAACAGCCGGGCTT( | [] |
| 2010 | 5 | AFAP1L1 | {'exs': 0, 'nonexs': 7} | True | 1.49E+08 | GGCACCTACAGTTTGGCAGTTC | [] |
| 2011 | 5 | AFAP1L1 | {'exs': 0, 'nonexs': 7} | True | 1.49E+08 | TCCTCTTCCTAGGTCTTCTTCG | ['TCTTCATTTCCCATTCCTGAAG'] |
| 2012 | 5 | ARHGEF37 | {'exs': 0, 'nonexs': 7} | True | 1.5E+08 | ACCCTAAGAGGGCAGACTCCA( | [] |
| 2013 | 5 | PPARGC1B | {'exs': 0, 'nonexs': 7} | True | 1.5E+08 | CCAAACTAGATGTTCTGCCCTG | [] |
| 2014 | 5 | PDGFRB | {'exs': 1, 'nonexs': 6} | True | 1.5E+08 | CCCTACAGGAAGCTATCCTCTC | [] |
| 2015 | 5 | CDX1 | {'exs': 0, 'nonexs': 7} | True | 1.5E+08 | GGGCATGGGGCTATGGCAGAA | ['GCAGAAACTCCTCTTTCACAGG'] |
| 2016 | 5 | NDST1 | {'exs': 1, 'nonexs': 6} | True | 1.51E+08 | CCACGGCTACCTGGTGTTCTG( | [] |
| 2017 | 5 | SYNPO | {'exs': 0, 'nonexs': 7} | True | 1.51E+08 | GGGGCTCTAGGTGGTGCAGGA | [] |
| 2018 | 5 | MYOZ3 | {'exs': 0, 'nonexs': 7} | True | 1.51E+08 | GGCTAGGGCTACAGCTCCTCG | [] |

| | | | | | | | |
|---|---|---|---|---|---|---|---|
| 2019 | 5 | RBM22 | {'ess': 5, 'noness': 2} | False | 1.51E+08 | CAAGGTGCTAGGGGCTGCTGT | [] |
| 2020 | 5 | IRGM | {'ess': 0, 'noness': 7} | True | 1.51E+08 | TGGCTAGCTGTTGAATATCCTG | [] |
| 2021 | 5 | TNIP1 | {'ess': 1, 'noness': 6} | True | 1.51E+08 | CTTCTAGTTTCTTGGCAATCTG | [] |
| 2022 | 5 | TNIP1 | {'ess': 1, 'noness': 6} | True | 1.51E+08 | CAATCTCTACCACTCATCAAAG | [] |
| 2023 | 5 | ANXA6 | {'ess': 0, 'noness': 7} | True | 1.51E+08 | GCCCTAGTCCTCACCACCACA | [] |
| 2024 | 5 | CCDC69 | {'ess': 0, 'noness': 7} | True | 1.51E+08 | GCACCCTATGTGGCGAGGAAA | [] |
| 2025 | 5 | SLC36A1 | {'ess': 0, 'noness': 7} | True | 1.51E+08 | AGATCCCTATATGAAGGCACAG | [] |
| 2026 | 5 | SLC36A1 | {'ess': 0, 'noness': 7} | True | 1.52E+08 | AGTGAACTATGATTGCACCATG | ['ACCATACACTCCAGCCTGGGTG'] |
| 2027 | 5 | SLC36A1 | {'ess': 0, 'noness': 7} | True | 1.52E+08 | TCTAGATCTGTGCTCTTTCTAG | [] |
| 2028 | 5 | FAT2 | {'ess': 0, 'noness': 7} | True | 1.52E+08 | CTAGAACATGACCTCCTCACAG | [] |
| 2029 | 5 | SPARC | {'ess': 1, 'noness': 7} | True | 1.52E+08 | TTAAATCTATGTTAGCACCTTG | [] |
| 2030 | 5 | ATOX1 | {'ess': 0, 'noness': 5} | True | 1.52E+08 | CCCCTGCTACTCAAGGCCAAG | [] |
| 2031 | 5 | GRIA1 | {'ess': 0, 'noness': 7} | True | 1.54E+08 | GGATTCACTACGGGATTTGTAG | [] |
| 2032 | 5 | PPP1R2B | unavailable | False | 1.57E+08 | CTTCTATGAACTTCGTAATTTG | [] |
| 2033 | 5 | HAVCR1 | {'ess': 0, 'noness': 7} | True | 1.57E+08 | GTTACTCTACCCAGTATCACTG | [] |
| 2034 | 5 | HAVCR2 | {'ess': 0, 'noness': 7} | True | 1.57E+08 | TTGGATCTATGGCATTGCAAAG | [] |
| 2035 | 5 | ITK | {'ess': 1, 'noness': 6} | True | 1.57E+08 | CTCTACTAAAGTCCTGATTCTG | [] |
| 2036 | 5 | ADAM19 | {'ess': 0, 'noness': 7} | True | 1.57E+08 | TGGACAGGTCTAGATTTTCGAG | [] |
| 2037 | 5 | CLINT1 | {'ess': 1, 'noness': 7} | True | 1.58E+08 | CATATCTACCAGACGAAAACAG | [] |
| 2038 | 5 | EBF1 | unavailable | False | 1.59E+08 | CCCCCTCTATTCGCCTTTATTG | [] |
| 2039 | 5 | RNF145 | {'ess': 0, 'noness': 7} | True | 1.59E+08 | CTCCTCTAGGCTGATTCAACAG | [] |
| 2040 | 5 | UBLCP1 | {'ess': 0, 'noness': 7} | True | 1.59E+08 | AACTACTGTCCTTGCTTCTTTG | [] |
| 2041 | 5 | IL12B | {'ess': 0, 'noness': 7} | True | 1.59E+08 | CTCACCTAACTGCAGGGCACA | [] |
| 2042 | 5 | ADRA1B | {'ess': 0, 'noness': 7} | True | 1.6E+08 | GGGCCCTAAAACTGCCCGGGC | [] |
| 2043 | 5 | PWWP2A | {'ess': 0, 'noness': 7} | True | 1.6E+08 | GCCTGCTAATGTCTTTGTGCTG | [] |
| 2044 | 5 | CCNJL | {'ess': 0, 'noness': 7} | True | 1.6E+08 | GGTGGCCTATCTGTCAAAGCA | [] |
| 2045 | 5 | C1QTNF2 | {'ess': 0, 'noness': 7} | True | 1.6E+08 | GCATGTCTATACCTCGTTGGG | [] |
| 2046 | 5 | SLU7 | {'ess': 5, 'noness': 4} | False | 1.6E+08 | AGTTGCTACTGTCCAAGGAAAG | [] |
| 2047 | 5 | ATP10B | {'ess': 0, 'noness': 7} | True | 1.61E+08 | TTCTAACCTTTGTAAACAAATG | [] |
| 2048 | 5 | GABRA6 | {'ess': 1, 'noness': 7} | True | 1.62E+08 | AAGCTATTCAACACTGCTACTG | [] |
| 2049 | 5 | GABRA1 | {'ess': 0, 'noness': 7} | True | 1.62E+08 | AAAGATCTATTGATGTGGTGTG | [] |
| 2050 | 5 | MAT2B | {'ess': 0, 'noness': 7} | True | 1.64E+08 | TAAACTAATGAAAGACCGTTTG | [] |
| 2051 | 5 | FBLL1 | unavailable | False | 1.69E+08 | GTGCTATTTGCTGCTGCTCTTG | [] |
| 2052 | 5 | DOCK2 | {'ess': 0, 'noness': 7} | True | 1.7E+08 | GGTTGCCTAGGCTCTCAATCC | ['AATCCATCTCCAGTTTCCACTG'] |
| 2053 | 5 | DOCK2 | {'ess': 0, 'noness': 7} | True | 1.7E+08 | GCCCTGTGCTAACCCTTGGCA | [] |
| 2054 | 5 | INSYN2B | unavailable | False | 1.7E+08 | TACTCTACACTGAAAAACACAG | [] |
| 2055 | 5 | FOXI1 | {'ess': 0, 'noness': 7} | True | 1.7E+08 | GTTCTGTACCTAGACCTCGGT | [] |
| 2056 | 5 | LCP2 | {'ess': 0, 'noness': 7} | True | 1.7E+08 | ACTTGCTATGGGTACCCTGCA | [] |
| 2057 | 5 | KCNMB1 | {'ess': 0, 'noness': 7} | True | 1.7E+08 | GGCTCTACTTCTGGGCCGCCA | [] |
| 2058 | 5 | FGF18 | {'ess': 1, 'noness': 6} | True | 1.71E+08 | GGTGGCCTAGGCAGGGTGTGT | [] |
| 2059 | 5 | EFCAB9 | {'ess': 0, 'noness': 7} | True | 1.72E+08 | ATCTACTTGATGTGACATTTTG | [] |
| 2060 | 5 | SH3PXD2B | {'ess': 2, 'noness': 7} | False | 1.72E+08 | AGTCGGCTACGGCTTCTTTCT | [] |
| 2061 | 5 | NEURL1B | {'ess': 0, 'noness': 7} | True | 1.73E+08 | CAGGCTAGGCTATGGCCTGTA | [] |
| 2062 | 5 | NKX2-5 | {'ess': 0, 'noness': 7} | True | 1.73E+08 | GTCCCTTCCCTACCAGGCTCG | [] |
| 2063 | 5 | NSG2 | unavailable | False | 1.74E+08 | GCAGGCCTCTAGTGGCCCTGG | [] |
| 2064 | 5 | MSX2 | {'ess': 0, 'noness': 7} | True | 1.75E+08 | ACCTCCTACTCCTGCCCCGGC | [] |
| 2065 | 5 | SFXN1 | {'ess': 0, 'noness': 7} | True | 1.76E+08 | GTCCTGCTAACCCATGCCAGG | [] |
| 2066 | 5 | HRH2 | {'ess': 0, 'noness': 7} | True | 1.76E+08 | CTAGGTCTAGACAGCCTCAGA | [] |
| 2067 | 5 | LOC10537 | unavailable | False | 1.76E+08 | CAATCCTCTAAGCATGGCAGG | [] |
| 2068 | 5 | FAM153B | {'ess': 0, 'noness': 7} | True | 1.76E+08 | TCTCTAAAATATATTAAAAATG | [] |
| 2069 | 5 | KIAA1191 | {'ess': 0, 'noness': 7} | True | 1.76E+08 | GGGCTCTAGAAGCCAGTGGGT | [] |
| 2070 | 5 | ARL10 | {'ess': 0, 'noness': 7} | True | 1.76E+08 | AATTCCTAGCTTTCTCCTACAG | [] |
| 2071 | 5 | ARL10 | {'ess': 0, 'noness': 7} | True | 1.76E+08 | AGCCTAGGAGAGGAGCTCCAA | [] |
| 2072 | 5 | ARL10 | {'ess': 0, 'noness': 7} | True | 1.76E+08 | GACACCTACACTGGTTCCTCA | [] |
| 2073 | 5 | NOP16 | {'ess': 8, 'noness': 1} | False | 1.76E+08 | CGTGGTTCTCTACCATGTAGC | [] |
| 2074 | 5 | CLTB | {'ess': 0, 'noness': 7} | True | 1.76E+08 | GGCACCTAGCGGGACAGTGGC | [] |
| 2075 | 5 | LOC11226 | unavailable | False | 1.77E+08 | GAGGAGCTAAACCTGGGCTAA | [] |
| 2076 | 5 | SNCB | {'ess': 1, 'noness': 7} | True | 1.77E+08 | TCCTGGGCCCCTACGCCTCTG | [] |
| 2077 | 5 | TSPAN17 | {'ess': 0, 'noness': 7} | True | 1.77E+08 | TGAAATCTACCAATTTTTAAAG | [] |
| 2078 | 5 | UNC5A | {'ess': 0, 'noness': 7} | True | 1.77E+08 | CTTGCCCTACCTCTGGGCCCA | [] |
| 2079 | 5 | ZNF346 | {'ess': 0, 'noness': 7} | True | 1.77E+08 | TCACCTCTAGTCTTCCAAGGTG | [] |
| 2080 | 5 | ZNF346 | {'ess': 0, 'noness': 7} | True | 1.77E+08 | TCTGGTCTAAGACTCAGGAAAG | [] |
| 2081 | 5 | NSD1 | {'ess': 0, 'noness': 7} | True | 1.77E+08 | GGACTACAGGCATGCACCACC | [] |
| 2082 | 5 | NSD1 | {'ess': 0, 'noness': 7} | True | 1.77E+08 | GATTGGTACTACTTCTGTTCTG | [] |
| 2083 | 5 | PRELID1 | {'ess': 7, 'noness': 1} | False | 1.77E+08 | AGACTGGCTACACAAACTGTTG | [] |
| 2084 | 5 | SLC34A1 | {'ess': 0, 'noness': 7} | True | 1.77E+08 | CCCACAGCCTAGAGGCGGGTG | [] |
| 2085 | 5 | PFN3 | {'ess': 0, 'noness': 7} | True | 1.77E+08 | GGCCGGCTAGGCGCCCTGCAT | [] |
| 2086 | 5 | GRK6 | {'ess': 1, 'noness': 6} | True | 1.77E+08 | TAGCTGCTACCGCCAACTGCT | [] |
| 2087 | 5 | PRR7 | {'ess': 0, 'noness': 7} | True | 1.77E+08 | GCGCCCCTCTATACGGCTGTA | [] |
| 2088 | 5 | DBN1 | {'ess': 0, 'noness': 7} | True | 1.77E+08 | CCACCGCTAATCACCACCCTC | [] |
| 2089 | 5 | PDLIM7 | {'ess': 0, 'noness': 7} | True | 1.77E+08 | GCCGGGCTACGACTGCACGTT | [] |
| 2090 | 5 | TMED9 | {'ess': 1, 'noness': 7} | True | 1.78E+08 | GGACAGCTACACAAGCTTCTTG | [] |
| 2091 | 5 | FAM153A | {'ess': 0, 'noness': 7} | True | 1.78E+08 | TCTCTAAAATATATTAAAAATG | [] |
| 2092 | 5 | PHYKPL | unavailable | False | 1.78E+08 | AGATCTAGCTCTTGCCACCTAG | [] |
| 2093 | 5 | LOC10272 | unavailable | False | 1.79E+08 | CTTCTTCTACATCCACCCAGTG | [] |
| 2094 | 5 | ZNF354A | {'ess': 1, 'noness': 6} | True | 1.79E+08 | TACTTTCTAGGGGTCCTCTTCG | [] |
| 2095 | 5 | GRM6 | {'ess': 0, 'noness': 5} | True | 1.79E+08 | CCCTGCTACTTGTGGGCCTCT | [] |
| 2096 | 5 | ZNF354C | {'ess': 0, 'noness': 7} | True | 1.79E+08 | GAGATGAACTAAACCTCATAGG | [] |
| 2097 | 5 | HNRNPH1 | {'ess': 6, 'noness': 2} | False | 1.8E+08 | TTACCTATGCAATGTTTGATTG | [] |
| 2098 | 5 | C5orf60 | {'ess': 0, 'noness': 3} | True | 1.8E+08 | TCTCCCTAGAAGGAAGCCTTTG | [] |
| 2099 | 5 | MAPK9 | {'ess': 1, 'noness': 6} | True | 1.8E+08 | GCAATCACTAACCTGAAGGCTG | [] |
| 2100 | 5 | CNOT6 | {'ess': 1, 'noness': 7} | True | 1.81E+08 | TGCTTGACTACCTCCTGCCAG | [] |
| 2101 | 5 | FLT4 | {'ess': 0, 'noness': 7} | True | 1.81E+08 | CTTCAGCTACCTAAGAGGGAAG | [] |
| 2102 | 5 | OR2Y1 | {'ess': 0, 'noness': 3} | True | 1.81E+08 | CACCTCCTACCCTGAGTCCCT | [] |
| 2103 | 5 | MGAT1 | {'ess': 2, 'noness': 6} | False | 1.81E+08 | GCAGGTGCTAATTCCAGCTAG | [] |
| 2104 | 5 | ZFP62 | {'ess': 0, 'noness': 7} | True | 1.81E+08 | CTACAGAGGCATCCTCATCCT | [] |
| 2105 | 5 | BTNL8 | {'ess': 0, 'noness': 7} | True | 1.81E+08 | TTCATCCTACATTTCACCCCTG | [] |
| 2106 | 5 | BTNL9 | {'ess': 0, 'noness': 7} | True | 1.81E+08 | TGAGACTATGCATTCAGTGTTG | [] |
| 2107 | 5 | RACK1 | unavailable | False | 1.81E+08 | AAACTTCTAGCGTGTGCCAATG | [] |
| 2108 | 5 | TRIM52 | {'ess': 3, 'noness': 5} | False | 1.81E+08 | TGACTAACACGGTGAAACCCTG | [] |
| 2109 | 5 | TRIM52 | {'ess': 3, 'noness': 5} | False | 1.81E+08 | CGAACTCTAACTCATGGGATGG | [] |
| 2110 | 6 | LOC10798 | unavailable | False | 1311698 | ACTCGCCTAAACTGCGTCCCC | [] |
| 2111 | 6 | GMDS | {'ess': 0, 'noness': 7} | True | 1949082 | AAACTAATTCTCTAGCTTTCAG | [] |
| 2112 | 6 | MYLK4 | {'ess': 0, 'noness': 7} | True | 2671300 | TGTAGACTATTTGGTCACAAAG | [] |
| 2113 | 6 | WRNIP1 | {'ess': 1, 'noness': 6} | True | 2772997 | AATGTTCTAGTTGCTATCTAGG | [] |
| 2114 | 6 | SERPINB1 | {'ess': 0, 'noness': 7} | True | 2833607 | GTCTCTTTCTTCTAAGGGGAAG | [] |
| 2115 | 6 | RIPK1 | {'ess': 0, 'noness': 5} | True | 3090812 | GGTTACTAGGTGCGGCAGGTA | [] |
| 2116 | 6 | TUBB2B | {'ess': 0, 'noness': 7} | True | 3224750 | GGGCATCTACGCCTCGTCCTC | [] |
| 2117 | 6 | PSMG4 | {'ess': 5, 'noness': 3} | False | 3264249 | CCACTCTACGCCACACTCACA | [] |
| 2118 | 6 | PSMG4 | {'ess': 5, 'noness': 3} | False | 3267711 | ACTCAGCTAGAACTTTTCGGGG | [] |
| 2119 | 6 | SLC22A23 | {'ess': 0, 'noness': 7} | True | 3273054 | GCCGGGCTACATGGCCTTCAT | [] |
| 2120 | 6 | FAM217A | unavailable | False | 4050541 | TAGCTGTCACTATGGTGACCTG | [] |
| 2121 | 6 | LYRM4 | {'ess': 3, 'noness': 4} | False | 5109422 | GGCTTGCTAGGTCCTGGGCAT | [] |
| 2122 | 6 | LYRM4 | {'ess': 3, 'noness': 4} | False | 5186794 | GACCAGCCTGACTAATACAGTG | [] |
| 2123 | 6 | FARS2 | {'ess': 8, 'noness': 1} | False | 5619501 | CACCTCTAGCATCTCAGCTCT | [] |
| 2124 | 6 | RIOK1 | {'ess': 9, 'noness': 0} | False | 7417440 | TCATTCTATTTGCCTTTTTTCG | [] |
| 2125 | 6 | DSP | {'ess': 0, 'noness': 7} | True | 7585877 | ACTGACTACTAGTGCCCAATAG | [] |
| 2126 | 6 | TXNDC5 | {'ess': 1, 'noness': 7} | True | 7883143 | TGTTCCTAAAGTTCGTCTTTCG | [] |
| 2127 | 6 | BLOC1S5 | {'ess': 0, 'noness': 6} | True | 8015809 | TAGCTACTAAGTGGTCACTATG | [] |
| 2128 | 6 | EEF1E1 | {'ess': 0, 'noness': 7} | True | 8079889 | GACAGCTTCTAGTGGGAATTAG | [] |
| 2129 | 6 | SLC35B3 | {'ess': 0, 'noness': 7} | True | 8413548 | CTGTCTATACAGTCTGTGCCAG | [] |
| 2130 | 6 | SLC35B3 | {'ess': 0, 'noness': 7} | True | 8417435 | CACTAGTGCATGTCAATCCCAG | [] |
| 2131 | 6 | TMEM14C | {'ess': 0, 'noness': 7} | True | 10730665 | CTTCTGCTAATGGGGTCTGTTG | [] |
| 2132 | 6 | TMEM14B | {'ess': 0, 'noness': 7} | True | 10756517 | CATGACTTCTGCTAATCAGATG | [] |
| 2133 | 6 | TMEM14B | {'ess': 0, 'noness': 7} | True | 10759100 | TACTAAAAATACAAAAAATTAG | [] |
| 2134 | 6 | MAK | {'ess': 1, 'noness': 4} | False | 10764651 | CTACCGGTGGCCTCCATACTT | [] |
| 2135 | 6 | SMIM13 | {'ess': 0, 'noness': 6} | True | 11134501 | ACAGGACTATGTCAGGGGTCT | [] |
| 2136 | 6 | PHACTR1 | {'ess': 0, 'noness': 3} | True | 13253048 | GCCACAGCTATTTTCCTTCCAG | [] |
| 2137 | 6 | GFOD1 | {'ess': 0, 'noness': 7} | True | 13364742 | AATCTGTGCTAACAGTAGAGGG | [] |
| 2138 | 6 | SIRT5 | {'ess': 1, 'noness': 6} | True | 13611907 | GGCCTAGTTCTTAGATATACTG | [] |
| 2139 | 6 | RANBP9 | {'ess': 3, 'noness': 5} | False | 13622361 | AATGCATAGCTAATGTAGGTAG | [] |
| 2140 | 6 | MCUR1 | unavailable | False | 13788672 | AGGGCAATTGCTAAGTGTGCTG | [] |
| 2141 | 6 | ATXN1 | {'ess': 0, 'noness': 7} | True | 16306328 | TGCCTCTACTTGCCTACATTAG | [] |
| 2142 | 6 | RBM24 | {'ess': 0, 'noness': 7} | True | 17292118 | CTGGTCTATTGCATTCGGTCTG | [] |
| 2143 | 6 | CDKAL1 | {'ess': 0, 'noness': 5} | True | 21231038 | TGTATTCTAATTATAGACCTTG | [] |
| 2144 | 6 | HDGFL1 | {'ess': 0, 'noness': 7} | True | 22570330 | CTGGTAACTACAGGCTGTCGC | [] |
| 2145 | 6 | DCDC2 | {'ess': 1, 'noness': 7} | True | 24174729 | AAAATGTTCTAAGCCACGGCAG | [] |
| 2146 | 6 | MRS2 | {'ess': 1, 'noness': 7} | True | 24423693 | CTGTTCCTAACGGTTTGTTAGG | [] |
| 2147 | 6 | ALDH5A1 | {'ess': 0, 'noness': 7} | True | 24533711 | CAAAGAATCCTACAAGCCCCC | [] |
| 2148 | 6 | C6orf62 | {'ess': 0, 'noness': 7} | True | 24706136 | GTACTCTACTCTGGCATATAAG | [] |
| 2149 | 6 | ARMH2 | unavailable | False | 24797409 | TTTAGAGAAACTAATTCCTATG | [] |
| 2150 | 6 | RIPOR2 | unavailable | False | 24839593 | GGATCTTCTCTACTTCTGTTTG | [] |
| 2151 | 6 | LOC10537 | unavailable | False | 25279411 | GCCGCACCTACAGCCCTAGGT | [] |
| 2152 | 6 | CARMIL1 | unavailable | False | 25577177 | TTTTGCTAGCTGAACCATCTTG | [] |
| 2153 | 6 | TRIM38 | {'ess': 0, 'noness': 7} | True | 25983166 | ACACTAACTGCAAAACAAAACG | ['AAAACAAACAAAAATAATTTTG'] |
| 2154 | 6 | H1-2 | unavailable | False | 26055786 | TTCGCCTATTTCTTCTTGGGCG | [] |
| 2155 | 6 | H1-4 | unavailable | False | 26157049 | ACTTTCTACTTTTTCTTGGCTG | [] |
| 2156 | 6 | H3C8 | unavailable | False | 26270973 | ACCTCTACGCTCTCTCCCCAC | [] |
| 2157 | 6 | BTN2A2 | {'ess': 1, 'noness': 7} | True | 26392966 | TTGATTCTATAGGCTCTGGTGG | [] |
| 2158 | 6 | BTN2A2 | {'ess': 1, 'noness': 7} | True | 26394328 | GTCTAGGTATTCCTTAATCCAG | [] |
| 2159 | 6 | BTN2A1 | {'ess': 0, 'noness': 7} | True | 26468548 | TTGATTCTATAGGCTCTGGTGG | [] |
| 2160 | 6 | POM121L2 | {'ess': 0, 'noness': 5} | True | 27309062 | CAGGCTACTTCTTGTAGGCATG | [] |
| 2161 | 6 | H2AC13 | unavailable | False | 27808501 | CCAGGCTTCTACTTGCCCTTC | [] |
| 2162 | 6 | H2AC15 | unavailable | False | 27837946 | CAGCCCCGCTACTTGCCCTTG | [] |
| 2163 | 6 | H1-5 | unavailable | False | 27866848 | GCTTCCTACTTCTTTTTGGCAG | [] |
| 2164 | 6 | ZKSCAN8 | unavailable | False | 28154016 | TGTTGTTCCTAAACGCTTATGG | [] |
| 2165 | 6 | ZSCAN9 | unavailable | False | 28233177 | GCCAAGCCCTAGACCAGCTCA | [] |
| 2166 | 6 | PGBD1 | {'ess': 0, 'noness': 7} | True | 28302283 | GTACCCTAATCTGACAGATGAG | [] |
| 2167 | 6 | ZSCAN23 | {'ess': 0, 'noness': 7} | True | 28434464 | CCAAGGAGCTAGCTTGATTCAG | [] |
| 2168 | 6 | GPX6 | {'ess': 0, 'noness': 7} | True | 28504291 | CTTCCTAGTGGGTATTGAACTG | [] |
| 2169 | 6 | GPX5 | {'ess': 0, 'noness': 7} | True | 28534166 | CTTCCTATTTGGTTTTGAATTG | [] |
| 2170 | 6 | OR2W1 | {'ess': 0, 'noness': 1} | False | 29044212 | AGTCTTTATCTATGACTTGCAG | ['TTGCAATTCCTCTTTATTTTTG'] |
| 2171 | 6 | OR12D3 | {'ess': 0, 'noness': 1} | False | 29374336 | ATTAGTCCTAGTGGTGTTGCTG | [] |
| 2172 | 6 | GABBR1 | {'ess': 0, 'noness': 1} | False | 29609466 | TGATGCTAGAAGGAAAGAACAG | [] |
| 2173 | 6 | MOG | {'ess': 0, 'noness': 1} | False | 29670878 | GCCCTCTACCTCCCAGGAGGA | [] |
| 2174 | 6 | TRIM31 | {'ess': 0, 'noness': 1} | False | 30108168 | CACTTCTAGGAAGATGTGGTTG | [] |
| 2175 | 6 | TRIM10 | {'ess': 0, 'noness': 1} | False | 30153687 | CCTTTCTATGCAGCTACTTCTG | [] |
| 2176 | 6 | PPP1R10 | {'ess': 0, 'noness': 1} | False | 30601548 | ATGGTCCCTAGGGCAGGGGGG | [] |
| 2177 | 6 | MRPS18B | {'ess': 0, 'noness': 1} | False | 30625796 | AGCTCCTACAGAGCACTCTGA | [] |
| 2178 | 6 | C6orf15 | {'ess': 0, 'noness': 1} | False | 31111380 | CTCTAGCCCCACTGCAACCTA | [] |
| 2179 | 6 | MICB | {'ess': 0, 'noness': 1} | False | 31509908 | TAGAGTCTAGGTGCCCTCAGT | [] |
| 2180 | 6 | DDX39B | unavailable | False | 31530433 | TTCTACCGTGTCTGTTCAACTG | [] |
| 2181 | 6 | ATP6V1G2 | {'ess': 0, 'noness': 1} | False | 31545407 | GTGGCCCTAGGCAGAAATCCG | [] |
| 2182 | 6 | LTA | {'ess': 1, 'noness': 0} | False | 31573692 | TCCAAGTTCTACAGAGCGAAGG | [] |
| 2183 | 6 | LTB | {'ess': 0, 'noness': 1} | False | 31581558 | TCCTTACCTATGAGGTGGGCA | [] |
| 2184 | 6 | NCR3 | {'ess': 0, 'noness': 1} | False | 31589311 | AAGAGGACTAGGGACATCTGG | [] |
| 2185 | 6 | BAG6 | unavailable | False | 31639130 | GCAAAGAGCTAAGGATCATCAG | [] |
| 2186 | 6 | LY6G5C | {'ess': 0, 'noness': 1} | False | 31676956 | AGTCACACTAAGGAGTATAGAG | [] |
| 2187 | 6 | ABHD16A | unavailable | False | 31687211 | TTGGTCCCTAGAGGTGCCAGG | [] |
| 2188 | 6 | LY6G6F-LY | unavailable | False | 31717803 | ACTCCCCCTATCCGCTCCACA | [] |
| 2189 | 6 | LY6G6D | {'ess': 0, 'noness': 7} | True | 31717803 | ACTCCCCCTATCCGCTCCACA | [] |
| 2190 | 6 | MPIG6B | unavailable | False | 31724872 | CTATCCCCTTCCCATCCCCAT | [] |
| 2191 | 6 | HSPA1A | {'ess': 0, 'noness': 1} | False | 31817581 | CCCCTAATCTACCTCCTCAATG | [] |
| 2192 | 6 | HSPA1B | {'ess': 0, 'noness': 1} | False | 31829875 | CCCCTAATCCACCTCCTCAAT | [] |
| 2193 | 6 | ZBTB12 | {'ess': 0, 'noness': 1} | False | 31899925 | GCGCGGCTAGCGGATGAGGAC | [] |
| 2194 | 6 | C2 | {'ess': 0, 'noness': 1} | False | 31934436 | CATGATTCCTACATGGCATGAG | [] |
| 2195 | 6 | C2 | {'ess': 0, 'noness': 1} | False | 31945356 | GCTAGAGGGGTAAAAAATTCAG | [] |
| 2196 | 6 | NELFE | unavailable | False | 31952300 | TTCCCTAGAAGCCATCCACAAG | [] |
| 2197 | 6 | DXO | unavailable | False | 31969876 | CATTACTATTTGGGAGAGGGAG | [] |
| 2198 | 6 | NOTCH4 | {'ess': 0, 'noness': 1} | False | 32195436 | -1 | -1 |
| 2199 | 6 | HLA-DRB5 | {'ess': 0, 'noness': 7} | True | 32517794 | -1 | -1 |
| 2200 | 6 | TAP2 | {'ess': 0, 'noness': 1} | False | 32828905 | GTCCCTCCTAGAGCTGGGCAA | [] |
| 2201 | 6 | HLA-DMB | {'ess': 0, 'noness': 7} | True | 32934970 | GATTCTGCCTCTAGGAAATGTG | [] |
| 2202 | 6 | COL11A2 | {'ess': 0, 'noness': 1} | False | 33163677 | ACGGTCCTATCCCATGAAGCA | [] |
| 2203 | 6 | COL11A2 | {'ess': 0, 'noness': 1} | False | 33186551 | GGACAACTAAGGAGGAGAGATG | [] |
| 2204 | 6 | VPS52 | {'ess': 0, 'noness': 1} | False | 33261439 | CCAGCCTAGGTGACAGAGCAA | [] |
| 2205 | 6 | WDR46 | {'ess': 0, 'noness': 1} | False | 33278965 | GCCCCTAAAAGGCAAAGCTAAG | [] |
| 2206 | 6 | RGL2 | {'ess': 0, 'noness': 1} | False | 33293306 | TGTGCTAGGAACAAACAACATG | [] |
| 2207 | 6 | DAXX | {'ess': 0, 'noness': 1} | False | 33318742 | AGGCAGCTAATCAGAGTCTGAG | [] |
| 2208 | 6 | KIFC1 | {'ess': 2, 'noness': 5} | False | 33406922 | AGAAGCCTACAACCACCCACG | [] |
| 2209 | 6 | PHF1 | {'ess': 0, 'noness': 1} | False | 33415233 | AGGCCTAGAAGAGGGGGCAAA | [] |
| 2210 | 6 | SYNGAP1 | {'ess': 0, 'noness': 1} | False | 33451905 | GGTGGGCTAGTGGTCTGCGGT | [] |
| 2211 | 6 | IP6K3 | {'ess': 1, 'noness': 7} | True | 33724306 | CTCTTGAGGACTACATTCCCG | [] |
| 2212 | 6 | GRM4 | {'ess': 0, 'noness': 7} | True | 34022820 | GACTCGCTAGATTGCATGGTTG | [] |
| 2213 | 6 | PACSIN1 | {'ess': 0, 'noness': 7} | True | 34532529 | GGGCCTCTAGATAGCCTCCAC | [] |
| 2214 | 6 | TAF11 | {'ess': 3, 'noness': 5} | False | 34878589 | GACTTTGGTCTAGAAGAAGATG | [] |
| 2215 | 6 | ANKS1A | {'ess': 0, 'noness': 7} | True | 35097946 | ACATGGCTAAATTTTGCTCCAG | ['TGCTCCAATAACTAATCCAGTG'] |
| 2216 | 6 | SCUBE3 | {'ess': 0, 'noness': 7} | True | 35248704 | GTTACTATTTGTAGGGCCTCAG | [] |
| 2217 | 6 | DEF6 | {'ess': 0, 'noness': 7} | True | 35321409 | AGAGGCTAATTTTCTGGTGCTG | [] |
| 2218 | 6 | TEAD3 | {'ess': 1, 'noness': 7} | True | 35475043 | GGGCACCCTAGTCTTTGACGA | [] |
| 2219 | 6 | CLPSL2 | unavailable | False | 35779449 | TCCTCTAAATCATATGGCACCG | [] |
| 2220 | 6 | CLPSL1 | unavailable | False | 35788009 | CACTAGAAGAACATTTTCTTAG | [] |
| 2221 | 6 | SLC26A8 | {'ess': 1, 'noness': 7} | True | 35943899 | CATCTCCTAGACATCTTCATTG | [] |
| 2222 | 6 | MAPK13 | {'ess': 1, 'noness': 6} | True | 36139372 | GATGAGTCCCTACAGCTTCATG | [] |
| 2223 | 6 | KCTD20 | {'ess': 0, 'noness': 7} | True | 36487174 | CTGGCCCTAATCCTGAAAGTC | [] |
| 2224 | 6 | STK38 | {'ess': 1, 'noness': 6} | True | 36494584 | CAATCCGCCTAAGTTGCAGGC | [] |
| 2225 | 6 | STK38 | {'ess': 1, 'noness': 6} | True | 36495783 | GTACTATTTTGCTGCTTTCATG | [] |
| 2226 | 6 | SRSF3 | {'ess': 7, 'noness': 1} | False | 36601988 | GTCTTCTATTTCCTTTCATTTG | [] |
| 2227 | 6 | CDKN1A | {'ess': 1, 'noness': 4} | False | 36685509 | TGCTCTAAGTCAAGCTGCCCA | [] |
| 2228 | 6 | RAB44 | {'ess': 0, 'noness': 1} | False | 36723215 | CAGGGCTAAGATTTGAACTCAG | [] |
| 2229 | 6 | PPIL1 | {'ess': 3, 'noness': 5} | False | 36855812 | CAAGTCTACCCAGAAGGGTATG | [] |
| 2230 | 6 | C6orf89 | {'ess': 1, 'noness': 7} | True | 36923440 | ATTTCCTACAGTTCTGAGAAAG | [] |
| 2231 | 6 | PIM1 | {'ess': 4, 'noness': 4} | False | 37174090 | GGCTGCTATTTGCTGGGCCCC | [] |
| 2232 | 6 | TMEM217 | {'ess': 0, 'noness': 7} | True | 37215208 | GACATCTATTGTGTATCTACAG | [] |
| 2233 | 6 | TMEM217 | {'ess': 0, 'noness': 7} | True | 37218421 | CACCTACCTCTGCCTCTCAAA | [] |
| 2234 | 6 | TBC1D22B | {'ess': 0, 'noness': 7} | True | 37324386 | TCACAGCTAAATTCTTGAATGG | [] |
| 2235 | 6 | TBC1D22B | {'ess': 0, 'noness': 7} | True | 37331171 | ACAGCACCTATCGGCGGTAGT | [] |
| 2236 | 6 | BTBD9 | {'ess': 1, 'noness': 7} | True | 38568582 | TTTCCTAAGTTTCATAAACAAG | [] |
| 2237 | 6 | GLO1 | {'ess': 1, 'noness': 6} | True | 38677294 | TCACAGCACTACATTAAGGTTG | [] |
| 2238 | 6 | SAYSD1 | unavailable | False | 39105431 | TGGGTCCTATCTCCCTGCCAG | [] |
| 2239 | 6 | KCNK17 | {'ess': 0, 'noness': 7} | True | 39299426 | TAACTAGCTGTCCTTGCCACAG | [] |
| 2240 | 6 | MOCS1 | {'ess': 0, 'noness': 7} | True | 39906356 | CAGGTGCTAAGCCCGATGGAA | [] |
| 2241 | 6 | LRFN2 | {'ess': 1, 'noness': 7} | True | 40391942 | CCCACCTAGACCGTGCTCTCC | ['GCTCTCCATCACCCATTCGGAG'] |
| 2242 | 6 | APOBEC2 | {'ess': 1, 'noness': 7} | True | 41061870 | TTGCCCTACTTCAGGATGTCTG | [] |
| 2243 | 6 | TREM1 | {'ess': 0, 'noness': 7} | True | 41276124 | TCGTGGGCCTAGGGTACAAAT | [] |
| 2244 | 6 | PGC | {'ess': 0, 'noness': 7} | True | 41736851 | CAGCAAGTCTAGGCGGCAGTG | [] |
| 2245 | 6 | FRS3 | {'ess': 0, 'noness': 5} | True | 41770618 | AAGTCCCTACAGAGGCAGGTC | [] |
| 2246 | 6 | PRICKLE4 | {'ess': 1, 'noness': 4} | False | 41787128 | GCGCCACTAGCACATGGTGCA | [] |
| 2247 | 6 | TOMM6 | {'ess': 0, 'noness': 7} | True | 41789327 | CTTGGCTACACCCCTGGCTGA | [] |
| 2248 | 6 | MED20 | {'ess': 7, 'noness': 1} | False | 41907071 | GCTCATCACTAACGAATCCCAG | [] |
| 2249 | 6 | MED20 | {'ess': 7, 'noness': 1} | False | 41907262 | GTCACTAGCTACCACACAGGG | [] |
| 2250 | 6 | CCND3 | {'ess': 2, 'noness': 5} | False | 41935939 | CAGGGCTACAGGTGTATGGCT | [] |
| 2251 | 6 | LOC11494 | unavailable | False | 42163248 | TCAGGCCTAGGAGCGCCACAA | [] |
| 2252 | 6 | RPL7L1 | {'ess': 7, 'noness': 1} | False | 42886463 | CTGGGTCTAGTTCAGCTGACG | [] |
| 2253 | 6 | PEX6 | {'ess': 1, 'noness': 7} | True | 42964334 | CTCCTAGCAGGCAGCAAACTT | [] |
| 2254 | 6 | KLHDC3 | {'ess': 0, 'noness': 7} | True | 43020732 | TTCCTCCTACCCATGGGAGGA | [] |
| 2255 | 6 | CUL7 | {'ess': 0, 'noness': 6} | True | 43037687 | CTAGGGCTACCGGAAGGTAGA | [] |
| 2256 | 6 | KLC4 | {'ess': 0, 'noness': 7} | True | 43067151 | GAGCTAAGAGAGGCCAAAACAG | [] |
| 2257 | 6 | KLC4 | {'ess': 0, 'noness': 7} | True | 43073948 | TGCACTAGGTTGGTTCAGATAG | [] |
| 2258 | 6 | TTBK1 | {'ess': 0, 'noness': 7} | True | 43269950 | CCCAGCCTAGGTCTTGTGGGT | [] |
| 2259 | 6 | ABCC10 | {'ess': 0, 'noness': 7} | True | 43446812 | TAGGACCTAGCCAGGAGCACA | [] |
| 2260 | 6 | TJAP1 | {'ess': 0, 'noness': 7} | True | 43505854 | GGGGCCCTAGTTGAGCAGGTT | [] |
| 2261 | 6 | POLH | {'ess': 1, 'noness': 7} | True | 43614556 | GGCAGCACTAATGTGTTAATGG | [] |
| 2262 | 6 | RSPH9 | {'ess': 0, 'noness': 7} | True | 43670948 | TCCCATTCTATAGCATGAAGGG | [] |
| 2263 | 6 | CAPN11 | {'ess': 0, 'noness': 7} | True | 44183931 | CAGCGCCTCTATCCCCACATG | [] |
| 2264 | 6 | CAPN11 | {'ess': 0, 'noness': 7} | True | 44183942 | CCAGGCTCCTACAGCGCCTCT | ['TCTATCCCCACATGGTCATCTG'] |
| 2265 | 6 | HSP90AB1 | {'ess': 3, 'noness': 4} | False | 44253597 | AACCTAATCGACTTCTTCCATG | [] |
| 2266 | 6 | SLC25A27 | {'ess': 0, 'noness': 7} | True | 46676581 | GCCCCCTATAAATCAGTCTCTG | [] |
| 2267 | 6 | PLA2G7 | {'ess': 0, 'noness': 7} | True | 46704559 | TAATCCTAATTGTATTTCTCTG | ['TCTCTATTCCTGAAGAGTTCTG'] |
| 2268 | 6 | ADGRF1 | unavailable | False | 47020451 | GTGAGCCTAGATTGTGCCATTG | [] |
| 2269 | 6 | TNFRSF2 | {'ess': 0, 'noness': 7} | True | 47232764 | CTATGTTCTACAGCAGGTCAGG | [] |
| 2270 | 6 | CD2AP | {'ess': 0, 'noness': 7} | True | 47603771 | ACAGGCTAGGAAACAAAATCTG | [] |
| 2271 | 6 | PTCHD4 | unavailable | False | 47901906 | TTCTAGCTTGAGGTAAATTCAG | [] |
| 2272 | 6 | GLYATL3 | {'ess': 0, 'noness': 7} | True | 49526913 | ACTGGGCTAGAGGTGAGGCAG | [] |
| 2273 | 6 | PGK2 | {'ess': 1, 'noness': 6} | True | 49785933 | TATTAACTACATGTTGCTGAGG | [] |
| 2274 | 6 | CRISP1 | {'ess': 0, 'noness': 7} | True | 49835315 | AAAGACCTATTTATCTCAGTG | [] |
| 2275 | 6 | TFAP2D | {'ess': 0, 'noness': 7} | True | 50772863 | TGTAGCTAGTCTGTCTTTTCTG | [] |
| 2276 | 6 | PKHD1 | {'ess': 0, 'noness': 7} | True | 51721985 | CTCTATTCTGAAATCTTCAAAG | [] |
| 2277 | 6 | FBXO9 | {'ess': 1, 'noness': 7} | True | 53097829 | AGGCTCTACAGAGGCCTTTCT | [] |
| 2278 | 6 | GCLC | {'ess': 4, 'noness': 3} | False | 53498755 | AATGTCTAGTTGGATGAGTCAG | [] |
| 2279 | 6 | LRRC1 | {'ess': 0, 'noness': 7} | True | 53822792 | GTGAAACTCTACACAGAAGTGG | [] |
| 2280 | 6 | FAM83B | {'ess': 0, 'noness': 7} | True | 54842006 | TTAATAGCTATTTATTTTTGTG | [] |
| 2281 | 6 | HCRTR2 | {'ess': 0, 'noness': 7} | True | 55282453 | ATATTCTACCAGTTTTGAAGTG | [] |
| 2282 | 6 | BMP5 | {'ess': 0, 'noness': 7} | True | 55803721 | CTTCTCTACATCCTAAGCAAAG | [] |
| 2283 | 6 | COL21A1 | {'ess': 0, 'noness': 5} | True | 56057656 | GCATCAGACACTAATAGTTTGG | [] |
| 2284 | 6 | DST | {'ess': 0, 'noness': 7} | True | 56459004 | CAATTGCACTATCTCTTTGAGG | [] |
| 2285 | 6 | BEND6 | {'ess': 0, 'noness': 7} | True | 57008202 | GAGAGTCTAAGGCAGCTCTCT | [] |
| 2286 | 6 | BEND6 | {'ess': 0, 'noness': 7} | True | 57018547 | AGGATCTATTTAATATCCTGAG | [] |
| 2287 | 6 | KIAA1586 | {'ess': 0, 'noness': 7} | True | 57054862 | TTCTATTTTAAGTTCCATATAG | [] |
| 2288 | 6 | BAG2 | {'ess': 0, 'noness': 7} | True | 57184189 | GTTTGAAGACTAATTGAATCTG | [] |
| 2289 | 6 | PRIM2 | {'ess': 0, 'noness': 1} | False | 57326062 | AACTACAAAAATATACTTACAG | [] |
| 2290 | 6 | PRIM2 | {'ess': 0, 'noness': 1} | False | 57587015 | -1 | -1 |
| 2291 | 6 | PRIM2 | {'ess': 0, 'noness': 1} | False | 57646157 | TGCCTAAGAATCTTCACTAAAG | [] |
| 2292 | 6 | KHDRBS2 | {'ess': 0, 'noness': 7} | True | 61749635 | AGATGAACTAGTAGCTTGGATG | [] |
| 2293 | 6 | KHDRBS2 | {'ess': 0, 'noness': 7} | True | 61827263 | CTTCCTCTAGGTTACAGTTGGG | [] |
| 2294 | 6 | LGSN | {'ess': 0, 'noness': 7} | True | 63280020 | ATTCTAAATAAAATACTCTAAG | [] |
| 2295 | 6 | COL19A1 | {'ess': 1, 'noness': 6} | True | 70185349 | GTGGCGCTACTGCATCTAGTG | [] |
| 2296 | 6 | FAM135A | {'ess': 0, 'noness': 7} | True | 70559920 | TTTATACTATTGGAAATATTTG | [] |
| 2297 | 6 | RIMS1 | {'ess': 1, 'noness': 7} | True | 72400713 | AGTTCACTATGATCGAATACAG | [] |
| 2298 | 6 | MTO1 | {'ess': 4, 'noness': 4} | False | 73500734 | AAAGCTATAACTCTCTCTCTTG | [] |
| 2299 | 6 | MYO6 | {'ess': 0, 'noness': 7} | True | 75841407 | -1 | -1 |
| 2300 | 6 | MYO6 | {'ess': 0, 'noness': 7} | True | 75915011 | ACATCTACTTTAACAGACTCTG | [] |
| 2301 | 6 | SH3BGRL2 | {'ess': 0, 'noness': 7} | True | 79699508 | ACTCTTCTTCTCTAAGGTTCTG | [] |
| 2302 | 6 | IBTK | {'ess': 1, 'noness': 7} | True | 82171424 | CCACAGTGAACTAGCATCCATG | [] |
| 2303 | 6 | UBE3D | unavailable | False | 82881449 | AGGGTTCTAGCCTGGGTGCCT | [] |
| 2304 | 6 | UBE3D | unavailable | False | 82944932 | TACTCCCTATGGGTTTTTGGTG | [] |
| 2305 | 6 | UBE3D | unavailable | False | 82956443 | CCAGTCTACATTTGCAGAACTG | [] |
| 2306 | 6 | ME1 | {'ess': 0, 'noness': 7} | True | 83211923 | ATTATCCTACTGGTCAACTTTG | [] |
| 2307 | 6 | MRAP2 | {'ess': 1, 'noness': 7} | True | 84145954 | TCTAATAAGAGGCATTTCTATG | [] |
| 2308 | 6 | TBX18 | {'ess': 0, 'noness': 7} | True | 84688593 | TGCTACAGTTTGGTTTCCCTTG | [] |
| 2309 | 6 | NT5E | {'ess': 0, 'noness': 7} | True | 85494003 | TGGCTATTGGTATAAAACAAAG | [] |
| 2310 | 6 | SYNCRIP | {'ess': 1, 'noness': 6} | True | 85614755 | ACTGTTTCTACTTCCACTGTTG | [] |
| 2311 | 6 | LOC10798 | unavailable | False | 86897731 | TGTTCCTATTCGGCCATCTTCG | ['ATCTTCAAAACTAGACTTTTTG'] |
| 2312 | 6 | HTR1E | {'ess': 0, 'noness': 7} | True | 87016431 | TACAGTCTAAGTATGCTCTCGG | [] |
| 2313 | 6 | SMIM8 | unavailable | False | 87340273 | CACTACTAATCCCATTTGGATG | [] |
| 2314 | 6 | SMIM8 | unavailable | False | 87341320 | GGCTAGGCTTTGTTTACACTCG | ['CTCATTTTTACCCCCATTTCTG'] |
| 2315 | 6 | C6orf163 | {'ess': 0, 'noness': 7} | True | 87365395 | GAAGACTTCTAATCAAGAGTTG | [] |
| 2316 | 6 | RARS2 | {'ess': 7, 'noness': 1} | False | 87518058 | AAAGTCTAGAGGCAGAAGGCAG | [] |
| 2317 | 6 | ORC3 | {'ess': 8, 'noness': 0} | False | 87651176 | CCAAACCTAGCCTATTATCCAG | [] |
| 2318 | 6 | ORC3 | {'ess': 8, 'noness': 0} | False | 87667031 | AGCTCTAATAAACCGAGCACTG | [] |
| 2319 | 6 | ORC3 | {'ess': 8, 'noness': 0} | False | 87667122 | GCTTTCTAGCAGCCTCCCCAT | [] |
| 2320 | 6 | RNGTT | {'ess': 7, 'noness': 1} | False | 88633366 | GGTCTCTTGCTCTAGGACTCT | [] |
| 2321 | 6 | PNRC1 | {'ess': 1, 'noness': 6} | True | 89084195 | TGAAATCTAAGTTTGAACTTTG | [] |
| 2322 | 6 | PM20D2 | {'ess': 0, 'noness': 7} | True | 89161507 | CTACTCCCTTTCTTCTCTTCTG | [] |
| 2323 | 6 | GABRR1 | {'ess': 0, 'noness': 7} | True | 89178769 | ACAAGCATCTAGGAGAAAATAG | [] |
| 2324 | 6 | GABRR2 | {'ess': 0, 'noness': 5} | True | 89257569 | TGGAGCCCCTAGGAAAACACT | [] |
| 2325 | 6 | UBE2J1 | {'ess': 0, 'noness': 1} | False | 89329857 | GTCCGTACTAGGAAATTTTAAG | [] |
| 2326 | 6 | RRAGD | {'ess': 0, 'noness': 7} | True | 89368055 | CTCACCTACAGCAGCACTCTA | [] |
| 2327 | 6 | ANKRD6 | {'ess': 0, 'noness': 7} | True | 89631003 | ATTGGTGCTAATTTTCCTTCTG | [] |
| 2328 | 6 | MDN1 | {'ess': 7, 'noness': 1} | False | 89644004 | TCTGTTCTATGGGTGGTCAGAG | [] |
| 2329 | 6 | BACH2 | {'ess': 2, 'noness': 7} | False | 89932407 | TCACTAGGTATAATCTTTCCTG | [] |
| 2330 | 6 | EPHA7 | {'ess': 1, 'noness': 6} | True | 93343794 | GCAGTCTAAAGGAGGTCAGCT | [] |
| 2331 | 6 | FHL5 | {'ess': 0, 'noness': 6} | True | 96615771 | GACTGTCTCCTAGATGTCAGTG | [] |
| 2332 | 6 | FBXL4 | {'ess': 0, 'noness': 5} | True | 98904143 | TGTATACTCTAAATATTCACAG | [] |
| 2333 | 6 | PNISR | unavailable | False | 99400539 | AAAGTATACTACCTTGATCGGG | [] |
| 2334 | 6 | USP45 | {'ess': 0, 'noness': 7} | True | 99462809 | AAAACCCTATCTCTACAAAAAG | ['AAAAAATAACAAAAATTAGCCG'] |
| 2335 | 6 | TSTD3 | unavailable | False | 99531719 | AAAACACTAGCTGTCAGATTTG | [] |
| 2336 | 6 | MCHR2 | {'ess': 1, 'noness': 7} | True | 99820939 | TTTCCTAAAAGTGTGATTTCAG | [] |
| 2337 | 6 | ASCC3 | {'ess': 5, 'noness': 4} | False | 1.01E+08 | TCTCTAAAAGCAGATAAATAAG | [] |
| 2338 | 6 | GRIK2 | {'ess': 0, 'noness': 7} | True | 1.02E+08 | GGGTGGCTAAATGTTGCTTTCG | [] |
| 2339 | 6 | GRIK2 | {'ess': 0, 'noness': 7} | True | 1.02E+08 | AAGATTACTAAACAGTGTCTGG | [] |
| 2340 | 6 | ATG5 | {'ess': 1, 'noness': 7} | True | 1.06E+08 | GATCTCCTAGTGTGTGCAACTG | [] |
| 2341 | 6 | CRYBG1 | unavailable | False | 1.07E+08 | TCTTCTGCTATCTGCTCAGTGG | [] |
| 2342 | 6 | SCML4 | {'ess': 0, 'noness': 7} | True | 1.08E+08 | TGAGGGGCTACAGGGTGGCAA | [] |
| 2343 | 6 | SCML4 | {'ess': 0, 'noness': 7} | True | 1.08E+08 | TGCAGACTAACGTAATAACTGG | [] |
| 2344 | 6 | CEP57L1 | {'ess': 0, 'noness': 7} | True | 1.09E+08 | TTACTAAAGCCTTCTTACACTG | [] |
| 2345 | 6 | MICAL1 | {'ess': 1, 'noness': 7} | True | 1.09E+08 | CTCGTCTAGCCCTGGGCCCCT | [] |
| 2346 | 6 | ZBTB24 | {'ess': 0, 'noness': 7} | True | 1.09E+08 | ATCATCACTACTCCCAATTCTG | [] |
| 2347 | 6 | AK9 | unavailable | False | 1.09E+08 | AGTAAACTACCCATTAATTGGG | [] |
| 2348 | 6 | AK9 | unavailable | False | 1.1E+08 | TTCTTATCTACTTTTGGTTGTG | [] |
| 2349 | 6 | METTL24 | unavailable | False | 1.1E+08 | CCTGCATCCTATTTCCATCTTG | [] |
| 2350 | 6 | DDO | {'ess': 0, 'noness': 7} | True | 1.1E+08 | TGTCATCTACAGGTTTGACTTG | [] |
| 2351 | 6 | GTF3C6 | {'ess': 3, 'noness': 5} | False | 1.11E+08 | GATTTCTAAGGCAGCATTTGAG | [] |
| 2352 | 6 | MFSD4B | unavailable | False | 1.11E+08 | CTCTAAACGTTAGTCCCTTTTG | [] |
| 2353 | 6 | LOC10272 | unavailable | False | 1.12E+08 | GGCCCCACCTACTGCCGTCCC | [] |
| 2354 | 6 | FAM229B | unavailable | False | 1.12E+08 | CTTAATGGTGCTATTTAGGATG | [] |
| 2355 | 6 | DSE | {'ess': 0, 'noness': 6} | True | 1.16E+08 | TCAGTGCTAACACTGTGATTGG | [] |

| | | | | | | | |
|---|---|---|---|---|---|---|---|
| 2356 | 6 | CALHM5 | unavailable | False | 1.17E+08 | ATGAGCTACATATTGTGGGCAG | [] |
| 2357 | 6 | RSPH4A | {'exs': 0, 'nonexs': 7} | True | 1.17E+08 | CCAGCCTATGTAGAAATTTTCG | ['TTTCAAACTTTCTATAAGTAAG'] |
| 2358 | 6 | VGLL2 | {'exs': 1, 'nonexs': 7} | True | 1.17E+08 | CTAAAACCAGTACACATCCTTG | [] |
| 2359 | 6 | NUS1 | {'exs': 6, 'nonexs': 1} | False | 1.18E+08 | ATGACCACTACTTTCCCAGACG | [] |
| 2360 | 6 | SLC35F1 | {'exs': 0, 'nonexs': 7} | True | 1.18E+08 | CACCCTAGGCCACACGAACAT | [] |
| 2361 | 6 | TBC1D32 | unavailable | False | 1.21E+08 | TCATGATCTATGTGCTCTGCAG | [] |
| 2362 | 6 | GJA1 | {'exs': 1, 'nonexs': 6} | True | 1.21E+08 | AGCCTGTATCTAGATCTCCAGG | [] |
| 2363 | 6 | HSF2 | {'exs': 0, 'nonexs': 7} | True | 1.22E+08 | CTAACTAAAAAATTACTTTATG | [] |
| 2364 | 6 | SMPDL3A | {'exs': 0, 'nonexs': 7} | True | 1.23E+08 | CTGTGAAATACTAGTAATTGTG | [] |
| 2365 | 6 | RNF217 | {'exs': 0, 'nonexs': 7} | True | 1.25E+08 | CATATTCCTATCTTCCAAGTTG | [] |
| 2366 | 6 | RNF217 | {'exs': 0, 'nonexs': 7} | True | 1.25E+08 | TGTCTAAAAATCCATATTCCTG | ['CATATTCCTATCTTCCAAGTTG'] |
| 2367 | 6 | TPD52L1 | {'exs': 1, 'nonexs': 6} | True | 1.25E+08 | CCGCCTACTTTCGTCTGAAATG | [] |
| 2368 | 6 | CENPW | {'exs': 5, 'nonexs': 3} | False | 1.26E+08 | TCTAACCTCTGCTCTTCTTTAG | [] |
| 2369 | 6 | RSPO3 | {'exs': 0, 'nonexs': 7} | True | 1.27E+08 | AACATTCTAGTGATTAAAACAG | [] |
| 2370 | 6 | RSPO3 | {'exs': 0, 'nonexs': 7} | True | 1.27E+08 | AACCCTCTAGTGTACAGTGCTG | [] |
| 2371 | 6 | PTPRK | {'exs': 1, 'nonexs': 6} | True | 1.28E+08 | ACCCAACTAAGATGATTCCAGG | [] |
| 2372 | 6 | PTPRK | {'exs': 1, 'nonexs': 6} | True | 1.28E+08 | CAACTCTACAAATAGGATATAG | [] |
| 2373 | 6 | ARHGAP18 | {'exs': 1, 'nonexs': 7} | True | 1.3E+08 | TAAGTCTTCTACAATGGCTTTG | [] |
| 2374 | 6 | TMEM244 | unavailable | False | 1.3E+08 | AGAATCTAAACAAGCAATTTTG | [] |
| 2375 | 6 | TAAR2 | {'exs': 0, 'nonexs': 7} | True | 1.33E+08 | AAGCCTACTCACTTTCTTTTTG | [] |
| 2376 | 6 | TAAR1 | {'exs': 0, 'nonexs': 7} | True | 1.33E+08 | TAATTCTATGAACTCAATTCCG | ['TTCCAAAAATAATTTACACCTG'] |
| 2377 | 6 | VNN1 | {'exs': 1, 'nonexs': 7} | True | 1.33E+08 | AATATTCTACCAACTTAATGAG | [] |
| 2378 | 6 | VNN2 | {'exs': 0, 'nonexs': 7} | True | 1.33E+08 | CCTATAACATTACAATATTTTG | [] |
| 2379 | 6 | SLC18B1 | unavailable | False | 1.33E+08 | CATCGGACTAGGTTTCATTAGG | [] |
| 2380 | 6 | EYA4 | {'exs': 1, 'nonexs': 7} | True | 1.34E+08 | ATCTATAAATTTAAATCATTTG | [] |
| 2381 | 6 | SLC2A12 | {'exs': 0, 'nonexs': 7} | True | 1.34E+08 | CTGACCTAGAAGCAAGACAGTG | [] |
| 2382 | 6 | HBS1L | {'exs': 5, 'nonexs': 4} | False | 1.35E+08 | TGTTACTATTCTCTAGTAAATG | [] |
| 2383 | 6 | AHI1 | {'exs': 1, 'nonexs': 6} | True | 1.35E+08 | TTTATCTAGAGGCCTTCTCTAG | [] |
| 2384 | 6 | AHI1 | {'exs': 1, 'nonexs': 6} | True | 1.35E+08 | CCTGAGTCTAGTGCTTTTTCCG | ['TTTCCACCATAATATACATGTG'] |
| 2385 | 6 | PDE7B | {'exs': 0, 'nonexs': 7} | True | 1.36E+08 | GGCCGGCCCCTAGGGGCTGT | [] |
| 2386 | 6 | MAP3K5 | {'exs': 1, 'nonexs': 6} | True | 1.37E+08 | ACAAAGCTAATTAGAGACCCAG | [] |
| 2387 | 6 | ARFGEF3 | unavailable | False | 1.38E+08 | CGGCTACACAATGATGTCATAG | [] |
| 2388 | 6 | NHSL1 | {'exs': 0, 'nonexs': 7} | True | 1.38E+08 | GGCCCTAACTCTCCTCGCTCA | [] |
| 2389 | 6 | CCDC28A | {'exs': 0, 'nonexs': 7} | True | 1.39E+08 | GGAAAATGCCTAACCTTTAAAG | [] |
| 2390 | 6 | AIG1 | {'exs': 0, 'nonexs': 7} | True | 1.43E+08 | TAGTTGCTATAGCTGCAAATTG | ['GCAAATTAAACAAATAAATGAG'] |
| 2391 | 6 | AIG1 | {'exs': 0, 'nonexs': 7} | True | 1.43E+08 | GGAACTAACTAGATGGTCCTAG | [] |
| 2392 | 6 | AIG1 | {'exs': 0, 'nonexs': 7} | True | 1.43E+08 | CCCACCTACTTTAGTTTGGCAG | [] |
| 2393 | 6 | LTV1 | {'exs': 7, 'nonexs': 1} | False | 1.44E+08 | GCTCCACTGTCTATAGCTTTAG | [] |
| 2394 | 6 | STX11 | {'exs': 0, 'nonexs': 7} | True | 1.44E+08 | CGGCCTGCTACTTGAGGCAGG | [] |
| 2395 | 6 | EPM2A | {'exs': 0, 'nonexs': 7} | True | 1.46E+08 | ACCAGCTACAGGCTACACACA | [] |
| 2396 | 6 | SAMD5 | {'exs': 0, 'nonexs': 7} | True | 1.48E+08 | GATATCTAGCGGCTCTGTGATG | [] |
| 2397 | 6 | SASH1 | {'exs': 0, 'nonexs': 7} | True | 1.49E+08 | GCCTGGCTACATGGCCTCAGG | [] |
| 2398 | 6 | UST | {'exs': 1, 'nonexs': 7} | True | 1.49E+08 | CCAATTCTATGGCTGATCCTGG | [] |
| 2399 | 6 | LATS1 | {'exs': 0, 'nonexs': 6} | True | 1.5E+08 | AAAACCCTAAAGCAACCAGAG | [] |
| 2400 | 6 | NUP43 | {'exs': 6, 'nonexs': 3} | False | 1.5E+08 | AGTACTTCTACGAAAAAAGATG | [] |
| 2401 | 6 | LRP11 | {'exs': 0, 'nonexs': 7} | True | 1.5E+08 | CTATAGATACATCCCATTTATG | [] |
| 2402 | 6 | RAET1E | {'exs': 0, 'nonexs': 7} | True | 1.5E+08 | TCACCTCTAGGTGGATCTTCTG | [] |
| 2403 | 6 | RAET1E | {'exs': 0, 'nonexs': 7} | True | 1.5E+08 | ACCAGACTAAGACGTCCTCAAG | [] |
| 2404 | 6 | IYD | {'exs': 1, 'nonexs': 7} | True | 1.5E+08 | CCCTGCCTACACTGTCACCAT | [] |
| 2405 | 6 | MTHFD1L | {'exs': 3, 'nonexs': 5} | False | 1.51E+08 | GTGAGCCTAGATCACGCGCCT | [] |
| 2406 | 6 | MTHFD1L | {'exs': 3, 'nonexs': 5} | False | 1.51E+08 | CATGCCTAGAACAAGCCTGGA | [] |
| 2407 | 6 | ESR1 | {'exs': 1, 'nonexs': 6} | True | 1.52E+08 | TGCTGCGCTAGCAGTGAGCAA | [] |
| 2408 | 6 | OPRM1 | {'exs': 0, 'nonexs': 7} | True | 1.54E+08 | AATTCTAGAGACTGCGTACCTG | [] |
| 2409 | 6 | OPRM1 | {'exs': 0, 'nonexs': 7} | True | 1.54E+08 | AATGCTCTAGACTCTTCCTGGG | [] |
| 2410 | 6 | OPRM1 | {'exs': 0, 'nonexs': 7} | True | 1.54E+08 | ATACATTATCTACCTCTTCATG | [] |
| 2411 | 6 | TIAM2 | {'exs': 0, 'nonexs': 7} | True | 1.55E+08 | ATCATACTATGATTTTCCGTGG | [] |
| 2412 | 6 | TFB1M | {'exs': 3, 'nonexs': 5} | False | 1.55E+08 | AGCTAGAGTCTGTAATTCTCTG | [] |
| 2413 | 6 | NOX3 | {'exs': 0, 'nonexs': 7} | True | 1.55E+08 | CAAAGTCTAGAAGCTCTCCTTG | [] |
| 2414 | 6 | SNX9 | {'exs': 0, 'nonexs': 7} | True | 1.58E+08 | TCTGTCCTACATCACTGGAAAG | [] |
| 2415 | 6 | SYTL3 | {'exs': 0, 'nonexs': 5} | True | 1.59E+08 | AATACCTACCCCTTAACAAATG | [] |
| 2416 | 6 | RSPH3 | {'exs': 0, 'nonexs': 5} | True | 1.59E+08 | -1 | -1 |
| 2417 | 6 | TAGAP | {'exs': 0, 'nonexs': 7} | True | 1.59E+08 | CTCCTAAATATACGATTCTTTG | [] |
| 2418 | 6 | WTAP | {'exs': 2, 'nonexs': 5} | False | 1.6E+08 | GTTCTAGAAGGGGAATTTTTCG | [] |
| 2419 | 6 | PNLDC1 | {'exs': 1, 'nonexs': 5} | False | 1.6E+08 | GCTACTTGGAGTAAATCAACAG | [] |
| 2420 | 6 | PRKN | unavailable | False | 1.61E+08 | CCCTGGCTACACGTCGAACCA | [] |
| 2421 | 6 | PACRG | {'exs': 0, 'nonexs': 7} | True | 1.63E+08 | TACTAAGAGGAACATTTCTCCG | ['TTCTCCAACAGTTTCCTATTTG'] |
| 2422 | 6 | PACRG | {'exs': 0, 'nonexs': 7} | True | 1.63E+08 | TTTCCGAGTCTCTATGCAAGGG | [] |
| 2423 | 6 | PACRG | {'exs': 0, 'nonexs': 7} | True | 1.63E+08 | GAGAACCTACGGAGGAGCCCA | [] |
| 2424 | 6 | PACRG | {'exs': 0, 'nonexs': 7} | True | 1.63E+08 | GGCTCTAGGGCCATGTGGAAA | [] |
| 2425 | 6 | QKI | {'exs': 1, 'nonexs': 7} | True | 1.64E+08 | ACTAAACTAACCTCGGTCTGCG | [] |
| 2426 | 6 | RPS6KA2 | {'exs': 1, 'nonexs': 6} | True | 1.66E+08 | ACCCGCTACAGCCGCGTGGAC | [] |
| 2427 | 6 | FGFR1OP | {'exs': 3, 'nonexs': 4} | False | 1.67E+08 | GTGTCTATGCAACATCTTCCAG | [] |
| 2428 | 6 | FRMD1 | {'exs': 1, 'nonexs': 7} | True | 1.68E+08 | GTGCCTACACCACAAACTCCT | [] |
| 2429 | 6 | WDR27 | {'exs': 0, 'nonexs': 5} | True | 1.69E+08 | ACTCAGCTAGAAAGAGCTGGAG | [] |
| 2430 | 6 | WDR27 | {'exs': 0, 'nonexs': 5} | True | 1.69E+08 | GCTACAAGCCCCTTTCCTCCA | [] |
| 2431 | 6 | WDR27 | {'exs': 0, 'nonexs': 5} | True | 1.7E+08 | TGGCAGCTAATTCCATGAAATG | [] |
| 2432 | 6 | WDR27 | {'exs': 0, 'nonexs': 5} | True | 1.7E+08 | TGTCTACATGTAACCGTATTTG | [] |
| 2433 | 6 | TCTE3 | {'exs': 1, 'nonexs': 7} | True | 1.7E+08 | AATGAGCTATTCATAATAAAGG | [] |
| 2434 | 6 | FAM120B | {'exs': 0, 'nonexs': 6} | True | 1.7E+08 | TTGACTAGTACCTTCTCCACTG | [] |
| 2435 | 7 | FAM20C | {'exs': 1, 'nonexs': 7} | True | 259979 | CGGACACTACCTCGCCGAGGC | [] |
| 2436 | 7 | ADAP1 | {'exs': 0, 'nonexs': 6} | True | 898920 | CACTCGCTAAGGTTTATGCTTG | [] |
| 2437 | 7 | C7orf50 | {'exs': 0, 'nonexs': 7} | True | 1112649 | TAATTCCTAGGACATTAATCGG | [] |
| 2438 | 7 | CYP2W1 | {'exs': 0, 'nonexs': 7} | True | 988821 | AGCTCCTAGGGCCTGGGCACC | [] |
| 2439 | 7 | GPR146 | {'exs': 0, 'nonexs': 5} | True | 1058516 | GGGCCGCCTACGCCAGCACCT | [] |
| 2440 | 7 | LOC107986 | unavailable | False | 1060059 | GCCGCCTATTGAGAGGCATCC | [] |
| 2441 | 7 | GPER1 | {'exs': 0, 'nonexs': 6} | True | 1092855 | GGCTGTCTACACGGCACTGCT | [] |
| 2442 | 7 | MICALL2 | {'exs': 2, 'nonexs': 6} | False | 1434595 | TGGCTACTACTGGGAGGGGCT | [] |
| 2443 | 7 | MAFK | {'exs': 1, 'nonexs': 7} | True | 1540374 | CCGGCACTAGGATGCAGCCGA | [] |
| 2444 | 7 | TMEM184 | {'exs': 2, 'nonexs': 7} | False | 1545375 | CTCTCTAGCCCGGACTCCACT | [] |
| 2445 | 7 | TMEM184 | {'exs': 2, 'nonexs': 7} | False | 1546925 | TGGGTCCCTACAGCACTGGCA | [] |
| 2446 | 7 | TMEM184 | {'exs': 2, 'nonexs': 7} | False | 1546951 | GCCCCCCTACAGGTCCTCCGA | [] |
| 2447 | 7 | TMEM184 | {'exs': 2, 'nonexs': 7} | False | 1547512 | CTCCGCCCTAATAGGGGGTGT | [] |
| 2448 | 7 | MAD1L1 | {'exs': 1, 'nonexs': 6} | True | 1816069 | TGCAGGCTACGCCACGGTCTG | [] |
| 2449 | 7 | NUDT1 | {'exs': 2, 'nonexs': 6} | False | 2251000 | CCGCTAGACCGTGTCCACCTC | [] |
| 2450 | 7 | SNX8 | {'exs': 0, 'nonexs': 7} | True | 2255055 | CAGGCGCTAGTGAGGACACAG | [] |
| 2451 | 7 | LFNG | {'exs': 0, 'nonexs': 7} | True | 2527211 | TGGCCACTAGAAGATGGCAGT | [] |
| 2452 | 7 | IQCE | {'exs': 0, 'nonexs': 7} | True | 2610161 | GACGGGGACCTAAACTGGAAA | [] |
| 2453 | 7 | TTYH3 | {'exs': 0, 'nonexs': 7} | True | 2660019 | GAACAGCTACCACAGCGTTCG | [] |
| 2454 | 7 | TTYH3 | {'exs': 0, 'nonexs': 7} | True | 2661738 | CGCGGTCTAGTGGCTGCCGCT | [] |
| 2455 | 7 | AMZ1 | {'exs': 1, 'nonexs': 7} | True | 2712877 | CTAACTCTCTTCCCCATCCCA | [] |
| 2456 | 7 | AMZ1 | {'exs': 1, 'nonexs': 7} | True | 2764859 | CATTCACTAAGAGGACGAGGC | [] |
| 2457 | 7 | SDK1 | {'exs': 0, 'nonexs': 7} | True | 4268749 | GATCCATGCTACCAGGACGCT | [] |
| 2458 | 7 | RADIL | {'exs': 0, 'nonexs': 7} | True | 4799377 | GCCCCCCTAGAGAGGGGGCGT | [] |
| 2459 | 7 | PAPOLB | {'exs': 0, 'nonexs': 1} | False | 4859896 | TCCAACTATAGGATTAGATATG | [] |
| 2460 | 7 | MMD2 | {'exs': 0, 'nonexs': 7} | True | 4909733 | ACCTGGCCTACTGCCTCTTTA | [] |
| 2461 | 7 | FBXL18 | {'exs': 0, 'nonexs': 7} | True | 5490406 | TTCAATTCTAGGTGCTCACCTG | [] |
| 2462 | 7 | ACTB | {'exs': 5, 'nonexs': 3} | False | 5527747 | GTCCGCCTAGAAGCATTTGCG | [] |
| 2463 | 7 | FSCN1 | {'exs': 0, 'nonexs': 7} | True | 5605473 | CGGCCCTAGTACTCCCAGAGC | [] |
| 2464 | 7 | LOC107986 | unavailable | False | 5607428 | GTGCCCCTAAGGATAGCAGCT | [] |
| 2465 | 7 | RSPH10B | {'exs': 0, 'nonexs': 5} | True | 5926367 | TCTCTACTTTTTCTTCTTCTTG | [] |
| 2466 | 7 | USP42 | {'exs': 1, 'nonexs': 7} | True | 6152968 | TTCCTAATAACTAAGACTAAAG | [] |
| 2467 | 7 | USP42 | {'exs': 1, 'nonexs': 7} | True | 6157086 | TTTCTATCTTCCTAATTCCAAG | [] |
| 2468 | 7 | CYTH3 | {'exs': 0, 'nonexs': 7} | True | 6164943 | CAGGAAAGCTATTTTTTATTGG | [] |
| 2469 | 7 | ZDHHC4 | {'exs': 0, 'nonexs': 7} | True | 6588502 | CCCACCTACTTGTGTCCAGAT | [] |
| 2470 | 7 | RSPH10B2 | {'exs': 0, 'nonexs': 5} | True | 6798542 | TCTCTACTTTTTCTTCTTCTTG | [] |
| 2471 | 7 | LOC11226 | unavailable | False | 6955568 | ATCAACCTACTGGAGCAACACG | [] |
| 2472 | 7 | DGKB | {'exs': 0, 'nonexs': 7} | True | 14176823 | ATCAACTACTCTGTACTCACTG | [] |
| 2473 | 7 | DGKB | {'exs': 0, 'nonexs': 7} | True | 14336662 | ACAGCCTATGACAATATTTGAG | [] |
| 2474 | 7 | AGMO | unavailable | False | 15158355 | ACATTCTAAAAGAGAAACTATG | [] |
| 2475 | 7 | SOSTDC1 | {'exs': 0, 'nonexs': 7} | True | 16462547 | CTGAGTTCTAACTCATGCTGTG | [] |
| 2476 | 7 | TWIST1 | {'exs': 0, 'nonexs': 1} | False | 19116712 | TCCGCCTGCTAGTGGGACGCG | [] |
| 2477 | 7 | TWISTNB | {'exs': 7, 'nonexs': 1} | False | 19698315 | TAAAATACACTAAAGAAAATTG | ['TTACTTTTCCCTTTTCTTTTTG'] |
| 2478 | 7 | TMEM196 | {'exs': 0, 'nonexs': 7} | True | 19724263 | GGTAACTCCCTAGAAATCAGCG | [] |
| 2479 | 7 | MACC1 | {'exs': 0, 'nonexs': 7} | True | 20140945 | TGTTTCTATACTTCCTCAGAAG | [] |
| 2480 | 7 | SP4 | {'exs': 1, 'nonexs': 7} | True | 21443630 | ATCACTATCTCGTTCCTATACG | [] |
| 2481 | 7 | IL6 | {'exs': 1, 'nonexs': 6} | True | 22731572 | CCATGCTACATTTGCCGAAGAG | [] |
| 2482 | 7 | FAM126A | {'exs': 1, 'nonexs': 7} | True | 22893996 | GAATTTCTATATTTTAAAAACG | ['AAACAATCTTCTGACAATTAAG'] |
| 2483 | 7 | FAM126A | {'exs': 1, 'nonexs': 7} | True | 22949378 | GCTCTGCTAAAACTGCACTCTG | [] |
| 2484 | 7 | GPNMB | {'exs': 2, 'nonexs': 6} | False | 23274089 | TTCTATTGGGTTGTATTCCTTG | [] |
| 2485 | 7 | CCDC126 | {'exs': 0, 'nonexs': 7} | True | 23643114 | AACTGCTATCTGATACTGCCCG | [] |
| 2486 | 7 | HOXA6 | {'exs': 0, 'nonexs': 7} | True | 27145657 | GGCATCTACTCGCCCGCCTTT | [] |
| 2487 | 7 | TAX1BP1 | {'exs': 0, 'nonexs': 7} | True | 27828828 | AAAGTAACTAGTCAAAATTTAG | [] |
| 2488 | 7 | TRIL | unavailable | False | 28955610 | CCTGGACCTAGTCGGCAAATC | [] |
| 2489 | 7 | PRR15 | {'exs': 0, 'nonexs': 7} | True | 29566718 | TTGGGGCTACTGCTTGTCCTC | [] |
| 2490 | 7 | FKBP14 | {'exs': 0, 'nonexs': 7} | True | 30014734 | TGTATCTCTATAACTCATCGTG | [] |
| 2491 | 7 | PLEKHA8 | {'exs': 0, 'nonexs': 7} | True | 30129270 | ACATTCCTACACCTACACCAGG | [] |
| 2492 | 7 | PLEKHA8 | {'exs': 0, 'nonexs': 7} | True | 30129284 | TCCGTCTACACCCACATTCCT | ['ACATTCCTACACCTACACCAGG'] |
| 2493 | 7 | LOC10192 | unavailable | False | 30469216 | CTGCCTATAGTGTTCTGGCATG | [] |
| 2494 | 7 | GGCT | {'exs': 0, 'nonexs': 7} | True | 30497091 | GTTATGTTCTAAAGAGTTTGTG | [] |
| 2495 | 7 | AQP1 | {'exs': 0, 'nonexs': 7} | True | 30923628 | GACCCCTTCTATTTGGGCTTC | ['GGGCTTCATCTCCACCCTGGAG'] |
| 2496 | 7 | GHRHR | {'exs': 0, 'nonexs': 7} | True | 30979243 | CAGCCTAGCACATAGATGTCAG | [] |
| 2497 | 7 | ITPRID1 | unavailable | False | 31652828 | CTGATCTAGAGGAAGACATCTG | [] |
| 2498 | 7 | PDE1C | {'exs': 0, 'nonexs': 7} | True | 31753383 | CTCGGCCTATTTTCTGTTCCAG | [] |
| 2499 | 7 | KBTBD2 | {'exs': 2, 'nonexs': 6} | False | 32869344 | CCCCACTATACAGGTGGTAGT | [] |
| 2500 | 7 | NPSR1 | {'exs': 0, 'nonexs': 7} | True | 34849654 | CTAGGGTCTAGATGAATTCTGG | [] |
| 2501 | 7 | KIAA0895 | {'exs': 0, 'nonexs': 7} | True | 36326759 | ATTACTATATAAGATCTTTCAG | [] |
| 2502 | 7 | GPR141 | {'exs': 0, 'nonexs': 5} | True | 37741310 | TTGTGGCTAACGGCACAAAACG | ['GCACAAAACACAATTCCATAAG'] |
| 2503 | 7 | AMPH | {'exs': 2, 'nonexs': 6} | False | 38384817 | GTTGCCCTAATCTAAGCGTCG | [] |
| 2504 | 7 | VPS41 | {'exs': 2, 'nonexs': 6} | False | 38726245 | ATGAGCTATTTTTTCATCTCCG | ['ATCTCCAAAATTGCACTTCCTG'] |
| 2505 | 7 | YAE1 | unavailable | False | 39609885 | CTTCTAAATGTGTCCACCTCTG | [] |
| 2506 | 7 | INHBA | {'exs': 1, 'nonexs': 7} | True | 41689649 | GGCAACTCTATGAGCACCCAC | ['CCCACACTCCTCCACGATCATG'] |
| 2507 | 7 | GLI3 | {'exs': 2, 'nonexs': 6} | False | 41964329 | AAGCCTATTGCATAACTGCAAG | [] |
| 2508 | 7 | COA1 | unavailable | False | 43639581 | GGTCGTCTCTACTCCTTTTTCG | ['CTCCTTTTTCACTTCATCACCG'] |
| 2509 | 7 | COA1 | unavailable | False | 43640469 | ATAATTCTTCCTATGCCCACAG | [] |
| 2510 | 7 | MRPS24 | {'exs': 7, 'nonexs': 1} | False | 43866698 | TTGTTCTAGAGGTACTTATACG | ['ACTTATACACAACCTTTGAGGG'] |
| 2511 | 7 | UBE2D4 | {'exs': 1, 'nonexs': 7} | True | 43955860 | TTATGGCTAGATCCAGTAGAGG | [] |
| 2512 | 7 | SPDYE1 | {'exs': 0, 'nonexs': 7} | True | 44007767 | GGAGCTCTAGGAAAGGCGAGC | [] |
| 2513 | 7 | TMED4 | {'exs': 0, 'nonexs': 7} | True | 44579478 | AGGGCACTACACCAGCTTCTT | [] |
| 2514 | 7 | OGDH | {'exs': 8, 'nonexs': 1} | False | 44707998 | CAGCATCTACGAGAAGTTCTTG | [] |
| 2515 | 7 | H2AZ2 | unavailable | False | 44830143 | -1 | -1 |
| 2516 | 7 | H2AZ2 | unavailable | False | 44834500 | CATCCCTCTAAGCAGTTTTCTG | [] |
| 2517 | 7 | CCM2 | {'exs': 1, 'nonexs': 7} | True | 45070536 | TCAATGGACTAACAGACTCCTG | [] |
| 2518 | 7 | NACAD | {'exs': 0, 'nonexs': 7} | True | 45080508 | GTGGCTACATGGTCAGTTCCT | [] |
| 2519 | 7 | ADCY1 | {'exs': 1, 'nonexs': 7} | True | 45664374 | CCTGAAGCTACGTGCACAGTC | [] |
| 2520 | 7 | ADCY1 | {'exs': 1, 'nonexs': 7} | True | 45713994 | CTCCACTAAGCCTCCTTCCCA | [] |
| 2521 | 7 | IGFBP3 | {'exs': 2, 'nonexs': 6} | False | 45914819 | AGGCGTCTACTTGCTCTGCAT | [] |
| 2522 | 7 | C7orf65 | {'exs': 0, 'nonexs': 3} | True | 47659227 | CTAGTTCCTGCCTTCCCTCTC | ['AGGGGTCCCCTTTCCTTTTTTG'] |
| 2523 | 7 | HUS1 | {'exs': 6, 'nonexs': 2} | False | 47965355 | AGGGTGCTAGGACAGCGCAGG | [] |
| 2524 | 7 | SUN3 | {'exs': 0, 'nonexs': 7} | True | 47987329 | CTCTTCTAGATGTGCTTGCCTG | [] |
| 2525 | 7 | ABCA13 | {'exs': 0, 'nonexs': 7} | True | 48292894 | CCAGGCTAGGTGCCGGACTGA | [] |
| 2526 | 7 | ABCA13 | {'exs': 0, 'nonexs': 7} | True | 48629411 | CCATCTCTAGCCATCAGCTCT | [] |
| 2527 | 7 | VWC2 | {'exs': 0, 'nonexs': 7} | True | 49912184 | GGGAAGCGTCTACATTTGCCT | [] |
| 2528 | 7 | SPATA48 | unavailable | False | 50159168 | CGCCTCCTCTAGTAGTCTATG | [] |
| 2529 | 7 | DDC | {'exs': 0, 'nonexs': 7} | True | 50463230 | ACTCCTACTCCCTCTCTGCTC | [] |
| 2530 | 7 | VSTM2A | {'exs': 0, 'nonexs': 7} | True | 54554263 | AATGCTAGGAGCCAGTAAATAG | [] |
| 2531 | 7 | VOPP1 | {'exs': 0, 'nonexs': 7} | True | 55472854 | ACTACTTGGCCTTCACTACCTG | [] |
| 2532 | 7 | SUMF2 | {'exs': 0, 'nonexs': 7} | True | 56079646 | CCCTAGAAGGCTTTTCTCCTTG | [] |
| 2533 | 7 | ZNF680 | {'exs': 1, 'nonexs': 7} | True | 64526398 | GAAGGGCTACATAGTAGCCTG | [] |
| 2534 | 7 | ZNF138 | {'exs': 0, 'nonexs': 7} | True | 64831485 | TCTGCTCTAGCCAAAGGTCTTG | [] |
| 2535 | 7 | ZNF273 | {'exs': 0, 'nonexs': 7} | True | 64929037 | TCTAGGATTTCTCACCCATATG | [] |
| 2536 | 7 | ERV3-1 | unavailable | False | 64991211 | GAGACTATCCTTTCCAAGTCTG | [] |
| 2537 | 7 | ASL | {'exs': 0, 'nonexs': 7} | True | 66092911 | GGACCTAGGCCTGCTGTGCCT | [] |
| 2538 | 7 | CRCP | {'exs': 3, 'nonexs': 5} | False | 66152356 | CTTCTATGCTGGGTCCTCTTC | [] |
| 2539 | 7 | TPST1 | {'exs': 1, 'nonexs': 7} | True | 66352590 | CCAATCTAGTATACATCAGGAG | [] |
| 2540 | 7 | TPST1 | {'exs': 1, 'nonexs': 7} | True | 66356841 | TTCTGCTACTCCACTTGCTCAG | [] |
| 2541 | 7 | AUTS2 | {'exs': 0, 'nonexs': 7} | True | 70290507 | AACGGAACTAGAATCCTTTAAG | [] |
| 2542 | 7 | GALNT17 | unavailable | False | 71578671 | GGGCTTTGGCTATTCCAGATG | [] |
| 2543 | 7 | GALNT17 | unavailable | False | 71712145 | TCCCTCCCTCTACTTGATGGA | [] |
| 2544 | 7 | CALN1 | {'exs': 0, 'nonexs': 5} | True | 71787774 | GCTGCTACTCCATGCCGCTCC | [] |
| 2545 | 7 | POM121 | {'exs': 0, 'nonexs': 7} | True | 72946233 | CAAAGGCTACTTTTTGCGGGTG | [] |
| 2546 | 7 | TRIM74 | {'exs': 1, 'nonexs': 6} | True | 72954867 | TGCCGGGCTAGGGAGTGCGAC | [] |
| 2547 | 7 | NSUN5 | {'exs': 2, 'nonexs': 6} | False | 73303414 | CTCTGCTATGTGCAAGGCGGT | [] |
| 2548 | 7 | TRIM50 | {'exs': 0, 'nonexs': 7} | True | 73312920 | GGCGGCCCTACAGCTTGGTGG | [] |
| 2549 | 7 | TRIM50 | {'exs': 0, 'nonexs': 7} | True | 73313628 | TCTCTCTAGCCCCAGGGTCTA | [] |
| 2550 | 7 | FZD9 | {'exs': 2, 'nonexs': 6} | False | 73435782 | GTGTGGCTAGAGGTGTGTGGG | [] |
| 2551 | 7 | BAZ1B | {'exs': 1, 'nonexs': 7} | True | 73442195 | CTCTCTACTTCTTCTGTCTTCG | [] |
| 2552 | 7 | BCL7B | {'exs': 1, 'nonexs': 7} | True | 73537297 | TGGTGCTAGCTTTCTGACGCC | [] |
| 2553 | 7 | MLXIPL | {'exs': 0, 'nonexs': 7} | True | 73593864 | AGGACTATAAAGGTTTGCCAAG | [] |
| 2554 | 7 | VPS37D | {'exs': 0, 'nonexs': 7} | True | 73671375 | GTGGATCCTACCGGTGGGGGG | [] |
| 2555 | 7 | STX1A | {'exs': 1, 'nonexs': 7} | True | 73700406 | GGGTGGCTTCTAGGCGAAGAT | [] |
| 2556 | 7 | LAT2 | {'exs': 0, 'nonexs': 7} | True | 74224741 | CTGCCCTAGGCTTCTGTGGCT | [] |
| 2557 | 7 | RFC2 | {'exs': 7, 'nonexs': 1} | False | 74232105 | TCTGCTCTAACTGGCCACCGG | [] |
| 2558 | 7 | GTF2IRD1 | {'exs': 0, 'nonexs': 7} | True | 74603432 | CTGAGGTCTAGTAATTAAGAGG | [] |
| 2559 | 7 | GTF2I | {'exs': 1, 'nonexs': 4} | False | 74716954 | TTTTCTCCTAACCTTCTGCTGG | [] |
| 2560 | 7 | GTF2I | {'exs': 1, 'nonexs': 4} | False | 74758837 | GAGGTCTACCACGTGGGGTCT | [] |
| 2561 | 7 | CASTOR2 | unavailable | False | 75024698 | CCTTCTAGTGCTTCTCTGCTTG | [] |
| 2562 | 7 | POM121C | {'exs': 0, 'nonexs': 4} | True | 75418795 | CAAAGGCTACTTTTTGCGGGTG | [] |
| 2563 | 7 | SPDYE5 | unavailable | False | 75501936 | GGAGCTCTAGGAAAGGTGAGC | [] |
| 2564 | 7 | HIP1 | {'exs': 0, 'nonexs': 7} | True | 75538171 | GTTTGGCTCTATTCTTTTTCGG | [] |
| 2565 | 7 | POR | {'exs': 1, 'nonexs': 7} | True | 75986480 | GCCCCTAGCTCCACACGTCCA | [] |
| 2566 | 7 | STYXL1 | {'exs': 0, 'nonexs': 7} | True | 75996291 | ATCAGCTAGGTGGCGTGAACA | [] |
| 2567 | 7 | UPK3B | {'exs': 0, 'nonexs': 4} | True | 76515203 | GCCAGGCTAGGGGCTGAGGCT | [] |
| 2568 | 7 | SPDYE16 | unavailable | False | 76533693 | GGAGCTCTAGGAAAGGCGAGC | [] |
| 2569 | 7 | SPDYE17 | unavailable | False | 77024685 | GGAGCTCTAGGAAAGGTGGGC | [] |
| 2570 | 7 | SPDYE18 | unavailable | False | 77052777 | GGAGCTCTAGGAAAGGCGAGC | [] |
| 2571 | 7 | GSAP | unavailable | False | 77311401 | GCTGCTTCCTCTACAAATTCTG | [] |
| 2572 | 7 | PHTF2 | {'exs': 0, 'nonexs': 7} | True | 77940316 | AAGTCTACATTATACCCACCTG | [] |
| 2573 | 7 | MAGI2 | {'exs': 1, 'nonexs': 6} | True | 78019919 | ACGCCTACTTCCGGCAGACCT | [] |
| 2574 | 7 | MAGI2 | {'exs': 1, 'nonexs': 6} | True | 78026084 | TTGTCAGTTCTAGCCAGTTTGG | [] |
| 2575 | 7 | HGF | {'exs': 1, 'nonexs': 6} | True | 81702580 | CTTACTTCAGCTATGACTGTGG | [] |
| 2576 | 7 | CACNA2D1 | {'exs': 0, 'nonexs': 7} | True | 81957431 | CAAAAACTAAGAAAAATTAGAG | [] |
| 2577 | 7 | SEMA3D | {'exs': 0, 'nonexs': 7} | True | 84999439 | GTAGAAAACTACGTGGCTACAG | [] |
| 2578 | 7 | GRM3 | {'exs': 0, 'nonexs': 7} | True | 86793162 | AGGCTGGACTACCCATTTTCTG | [] |
| 2579 | 7 | GRM3 | {'exs': 0, 'nonexs': 7} | True | 86864395 | TCTAACAGTCTAAAAACACAAG | [] |
| 2580 | 7 | DMTF1 | {'exs': 0, 'nonexs': 7} | True | 87195139 | TTATTCTAATGACAGTTTACCG | ['ACCAAATCTTCGACATCCTTTG'] |
| 2581 | 7 | CROT | {'exs': 0, 'nonexs': 7} | True | 87398643 | TCATCTCTAAAGATGAGTAGAG | [] |
| 2582 | 7 | ABCB4 | {'exs': 1, 'nonexs': 6} | True | 87401921 | TCTAACTCTCAAATTTCAAATG | [] |
| 2583 | 7 | SLC25A40 | {'exs': 0, 'nonexs': 7} | True | 87836248 | ATCACTAGTATTGCTGCCTTCG | [] |
| 2584 | 7 | DBF4 | {'exs': 2, 'nonexs': 5} | False | 87908162 | AAATTCTAAAAGCCAGTAAATG | [] |
| 2585 | 7 | ADAM22 | {'exs': 1, 'nonexs': 7} | True | 88196574 | ATCCTAAGCCAATGACAAAAAG | [] |
| 2586 | 7 | STEAP4 | {'exs': 0, 'nonexs': 7} | True | 88279397 | ATGCTTTTTCTAGTGTTTTGAG | [] |
| 2587 | 7 | TEX47 | unavailable | False | 88794180 | CAATCCTAGAAACGTGAAGGAG | [] |
| 2588 | 7 | STEAP1 | {'exs': 0, 'nonexs': 7} | True | 90164733 | AAACAGTAATTCTACAACTGGG | [] |
| 2589 | 7 | CFAP69 | unavailable | False | 90282091 | TGAGCTACTGCACCTGGCCTT | [] |
| 2590 | 7 | LOC10272 | unavailable | False | 90494379 | TATCTATTCATCATATACTATG | [] |
| 2591 | 7 | AKAP9 | {'exs': 0, 'nonexs': 7} | True | 92108718 | ATTATCCTAATGAGTGTGAAAG | [] |
| 2592 | 7 | GATAD1 | {'exs': 1, 'nonexs': 7} | True | 92456561 | TTTGTGAACTACAAATGGTTGG | [] |
| 2593 | 7 | ERVW-1 | unavailable | False | 92468764 | CCGCTCTAACTGCTTCCTGCT | [] |
| 2594 | 7 | RBM48 | unavailable | False | 92536936 | CACTCTATATTCTTCTTCTTTG | [] |
| 2595 | 7 | VPS50 | unavailable | False | 93276346 | AAATCTAAAGGAGAGATATATG | [] |
| 2596 | 7 | VPS50 | unavailable | False | 93320568 | CCGCTAACACGGTGAAACCCC | [] |
| 2597 | 7 | CASD1 | {'exs': 0, 'nonexs': 7} | True | 94555757 | CCTAATGTTTTGATTTATCTTG | [] |
| 2598 | 7 | PEG10 | {'exs': 0, 'nonexs': 5} | True | 94664533 | GTCCCTCTACAGCGGGGCCGG | [] |
| 2599 | 7 | PON3 | {'exs': 1, 'nonexs': 7} | True | 95359972 | GAGTCTAGAGCTCACAGTACAG | [] |
| 2600 | 7 | DYNC1I1 | {'exs': 0, 'nonexs': 7} | True | 96097592 | CACTAGGCAGATAACTCAACAG | [] |
| 2601 | 7 | SLC25A13 | {'exs': 2, 'nonexs': 7} | False | 96121190 | TCCTATGGGCCTCCACCAATA | [] |
| 2602 | 7 | SLC25A13 | {'exs': 2, 'nonexs': 7} | False | 96169916 | GCCCTAGTAGACTCTGCCCTT | [] |
| 2603 | 7 | SEM1 | unavailable | False | 96688923 | TCTGGATGCTATGAAGTCTCCG | ['AAGTCTCCATCTTATAACCATG'] |
| 2604 | 7 | DLX5 | {'exs': 0, 'nonexs': 7} | True | 97020735 | GCCCATCTAATAGAGTGTCCC | [] |
| 2605 | 7 | SDHAF3 | unavailable | False | 97181214 | GACTAAAATTTTGGTTTCATAG | [] |
| 2606 | 7 | ASNS | {'exs': 2, 'nonexs': 6} | False | 97852258 | ACCACCTAAGCTTTGACAGCTG | [] |
| 2607 | 7 | LMTK2 | {'exs': 0, 'nonexs': 7} | True | 98205491 | AGCCACCTAGTCCTTTTCTCC | [] |
| 2608 | 7 | BHLHA15 | {'exs': 1, 'nonexs': 7} | True | 98212878 | TGGGCGCTAGGTGCCCTCTCG | [] |
| 2609 | 7 | BRI3 | {'exs': 0, 'nonexs': 7} | True | 98306554 | CACATGTCTACTGCTGATAGGG | [] |
| 2610 | 7 | BRI3 | {'exs': 0, 'nonexs': 7} | True | 98307528 | CAGGCTATTTTCTGTATGTCTG | [] |
| 2611 | 7 | NPTX2 | {'exs': 0, 'nonexs': 7} | True | 98628628 | AAGGCGGCTACAAGTCAAGGA | [] |
| 2612 | 7 | KPNA7 | {'exs': 0, 'nonexs': 7} | True | 99173707 | GCTTGGCTATTTTTTTGCTAAG | [] |
| 2613 | 7 | BUD31 | {'exs': 7, 'nonexs': 1} | False | 99417548 | AGTCGACTAAGGCAGAATTTTG | [] |
| 2614 | 7 | CPSF4 | {'exs': 3, 'nonexs': 5} | False | 99456647 | GATAGGAAACTAACAGACGTGG | [] |
| 2615 | 7 | ZKSCAN5 | {'exs': 0, 'nonexs': 7} | True | 99532248 | TAAGAGCTATTCTACAACAGAG | [] |
| 2616 | 7 | TMEM225B | unavailable | False | 99610564 | CTGGGCTAGATGACTGTTTCTG | [] |
| 2617 | 7 | OR2AE1 | {'exs': 0, 'nonexs': 3} | True | 99876061 | CTATTCCACTCTACACTCGGG | [] |
| 2618 | 7 | AZGP1 | {'exs': 1, 'nonexs': 7} | True | 99967002 | TGCTTCCTAGCTGGCCTCCCA | [] |
| 2619 | 7 | MAP11 | unavailable | False | 1E+08 | TGGCACGGTGCTACTTGACGG | [] |
| 2620 | 7 | CASTOR3 | unavailable | False | 1E+08 | GGCTGCTACAGTCTAGCCACT | [] |
| 2621 | 7 | ZCWPW1 | {'exs': 1, 'nonexs': 7} | True | 1E+08 | ACCAGCTACTTCCCAAACAGC | [] |
| 2622 | 7 | PPP1R35 | unavailable | False | 1E+08 | TTGGGGTCTACGCTTCCCATC | [] |
| 2623 | 7 | C7orf61 | {'exs': 1, 'nonexs': 7} | True | 1E+08 | TTCCCCCTAGAGGGGACTTTG | [] |
| 2624 | 7 | AGFG2 | {'exs': 0, 'nonexs': 7} | True | 1.01E+08 | ACAGTGCTACAAGAAGGGGTTG | [] |
| 2625 | 7 | SAP25 | unavailable | False | 1.01E+08 | AGAAGGCTATGGACAATGGGTG | [] |
| 2626 | 7 | LRCH4 | {'exs': 0, 'nonexs': 7} | True | 1.01E+08 | TTCACCCCTAAGGCCGGTCCA | [] |
| 2627 | 7 | LRCH4 | {'exs': 0, 'nonexs': 7} | True | 1.01E+08 | TTGGGGCCTAGGAACCCAGGA | [] |
| 2628 | 7 | GIGYF1 | {'exs': 0, 'nonexs': 7} | True | 1.01E+08 | CTGGCCTACAGCCCAGGGGCT | [] |
| 2629 | 7 | AP1S1 | {'exs': 0, 'nonexs': 7} | True | 1.01E+08 | AGCAGGGGCTATGCCAAACCC | ['CCATCTCCTCCAGCACACTCCG'] |
| 2630 | 7 | MOGAT3 | {'exs': 0, 'nonexs': 7} | True | 1.01E+08 | CCAGGCCTAGATGAAGGTGAG | [] |
| 2631 | 7 | CLDN15 | {'exs': 1, 'nonexs': 7} | True | 1.01E+08 | CTGCTACACGTAGGCGTTTCT | [] |
| 2632 | 7 | IFT22 | unavailable | False | 1.01E+08 | AAGGCTGGCTAGGTCATAATTG | [] |
| 2633 | 7 | MYL10 | {'exs': 0, 'nonexs': 7} | True | 1.02E+08 | CCCCTAATCCTTCTCTTCACC | [] |
| 2634 | 7 | SH2B2 | {'exs': 1, 'nonexs': 0} | False | 1.02E+08 | CGGGCTAGTAGAAGGAGTACT | [] |
| 2635 | 7 | SPDYE6 | {'exs': 1, 'nonexs': 4} | False | 1.02E+08 | GGAGCTCTAGGAAAGGTGAGC | [] |
| 2636 | 7 | LRWD1 | {'exs': 2, 'nonexs': 6} | False | 1.02E+08 | GTGAGGCTACATCCTCCCCCA | [] |
| 2637 | 7 | POLR2J | {'exs': 5, 'nonexs': 3} | False | 1.02E+08 | GCCCCTACTCAATTCCTTCCT | [] |
| 2638 | 7 | SPDYE2 | {'exs': 1, 'nonexs': 6} | True | 1.03E+08 | GGAGCTCTAGGAAAGGTGAGC | [] |
| 2639 | 7 | SPDYE2B | unavailable | False | 1.03E+08 | GGAGCTCTAGGAAAGGTGAGC | [] |
| 2640 | 7 | SLC26A5 | {'exs': 0, 'nonexs': 7} | True | 1.03E+08 | CCCCTACGGTCTCCATCTTCT | [] |
| 2641 | 7 | SLC26A5 | {'exs': 0, 'nonexs': 7} | True | 1.03E+08 | CCTCATCTATGCCTCAGGAGT | [] |
| 2642 | 7 | ORC5 | {'exs': 3, 'nonexs': 5} | False | 1.04E+08 | AACACTATTTCCTGACTGAATG | [] |
| 2643 | 7 | SRPK2 | {'exs': 1, 'nonexs': 7} | True | 1.05E+08 | ATTTGCTAAGAATTCAACCAAG | [] |
| 2644 | 7 | NAMPT | {'exs': 6, 'nonexs': 2} | False | 1.06E+08 | CATAAAGCCTAATGATGTGCTG | [] |
| 2645 | 7 | PIK3CG | {'exs': 1, 'nonexs': 6} | True | 1.07E+08 | GGAATACTAAAATAAGTGACAG | [] |
| 2646 | 7 | COG5 | {'exs': 2, 'nonexs': 5} | False | 1.07E+08 | CTCATTTCTCTAATCTCCGACG | ['TCCTCCTTATTCTCAGTTGATG', 'ACACCTCCTTCCCTCCTCCTTG'] |
| 2647 | 7 | GPR22 | {'exs': 1, 'nonexs': 7} | True | 1.07E+08 | TTCTCTAGTCTGTGACAACCTG | [] |
| 2648 | 7 | LAMB4 | {'exs': 0, 'nonexs': 5} | True | 1.08E+08 | TCTGCCTAGCTATAGCACCTAG | [] |
| 2649 | 7 | LAMB4 | {'exs': 0, 'nonexs': 5} | True | 1.08E+08 | TTTTCTATCTTGCTGAGTTCTG | [] |
| 2650 | 7 | LAMB4 | {'exs': 0, 'nonexs': 5} | True | 1.08E+08 | AGTGCACTAGTAAGCTTTAAAG | [] |
| 2651 | 7 | NRCAM | {'exs': 0, 'nonexs': 7} | True | 1.08E+08 | CCTACTGTCTGTATAATTAAAG | [] |
| 2652 | 7 | THAP5 | {'exs': 0, 'nonexs': 7} | True | 1.09E+08 | CTAGTTATTCTATATCATAGTG | [] |
| 2653 | 7 | DNAJB9 | {'exs': 1, 'nonexs': 6} | True | 1.09E+08 | TAAGAACTACTGTCCTGAACAG | [] |
| 2654 | 7 | IMMP2L | {'exs': 0, 'nonexs': 5} | True | 1.11E+08 | CTTCTATCTATCTGTTGTCCTG | [] |
| 2655 | 7 | IFRD1 | {'exs': 1, 'nonexs': 7} | True | 1.12E+08 | GAAAATCTAGAAGAATTCTCCG | ['CAACATCTGCTCTCTTATCTCG'] |
| 2656 | 7 | FOXP2 | {'exs': 0, 'nonexs': 7} | True | 1.15E+08 | GAGCTATTTACTGTTTATAAAG | [] |
| 2657 | 7 | TES | {'exs': 0, 'nonexs': 7} | True | 1.16E+08 | CTCCTAAGACATCCTCTTCTTG | ['CTTCTTACATTCCACTGAACAG'] |
| 2658 | 7 | MET | {'exs': 1, 'nonexs': 6} | True | 1.17E+08 | CTAGCACTATGATGTCTCCCAG | [] |
| 2659 | 7 | CFTR | {'exs': 0, 'nonexs': 7} | True | 1.18E+08 | CTCTCTAAAGCCTTGTATCTTG | [] |
| 2660 | 7 | CTTNBP2 | {'exs': 1, 'nonexs': 7} | True | 1.18E+08 | GTAGGCAGGCCTATTTGTTAGG | [] |
| 2661 | 7 | ANKRD7 | {'exs': 0, 'nonexs': 7} | True | 1.18E+08 | AATAAGGTGTCTATTTAGCATG | [] |
| 2662 | 7 | CPED1 | unavailable | False | 1.21E+08 | TCAGCCTAGTGAGGTCCATGT | [] |
| 2663 | 7 | CPED1 | unavailable | False | 1.21E+08 | GCCTACATAGTCCTTTTATTTG | [] |
| 2664 | 7 | TAS2R16 | {'exs': 0, 'nonexs': 7} | True | 1.23E+08 | AGGCCTAGCACTTTCCCTTTAG | [] |
| 2665 | 7 | IQUB | {'exs': 0, 'nonexs': 7} | True | 1.23E+08 | TCACCTAATGAGGAGGCCTCT | [] |
| 2666 | 7 | C7orf77 | unavailable | False | 1.25E+08 | ATTTTCTACAACAGCAGACCTG | [] |
| 2667 | 7 | GRM8 | {'exs': 2, 'nonexs': 7} | False | 1.27E+08 | GCAACCTAAGAAGATAAGTCTG | [] |
| 2668 | 7 | FSCN3 | {'exs': 0, 'nonexs': 7} | True | 1.28E+08 | CTCCCTTACCTAAAATTCCCAG | [] |
| 2669 | 7 | PAX4 | {'exs': 1, 'nonexs': 7} | True | 1.28E+08 | TTAAGGCTAGTGTGAGAAGTGG | [] |
| 2670 | 7 | FAM71F1 | {'exs': 1, 'nonexs': 6} | True | 1.29E+08 | CAGACTCTATATTAAGAAAACG | [] |
| 2671 | 7 | ATP6V1F | {'exs': 6, 'nonexs': 2} | False | 1.29E+08 | AGTCCCCTAGCGCAGGTCTTC | [] |
| 2672 | 7 | KCP | unavailable | False | 1.29E+08 | TGTGCTACCACCTGGCTGACA | [] |
| 2673 | 7 | TNPO3 | {'exs': 7, 'nonexs': 1} | False | 1.29E+08 | TGTGAGCTATCGAAACAACCTG | [] |
| 2674 | 7 | STRIP2 | unavailable | False | 1.29E+08 | CACTTCCTATGGTGTGGATCTG | [] |
| 2675 | 7 | STRIP2 | unavailable | False | 1.29E+08 | CTCTACCTACAGCATGTGTTGG | [] |
| 2676 | 7 | UBE2H | {'exs': 3, 'nonexs': 5} | False | 1.3E+08 | TCTACTACAACTCCATATCCTG | [] |
| 2677 | 7 | TMEM209 | {'exs': 2, 'nonexs': 5} | False | 1.3E+08 | TATGACTTGCTACTCGCCAAAG | [] |
| 2678 | 7 | CPA2 | {'exs': 1, 'nonexs': 6} | True | 1.3E+08 | GGCCCTAATAGGGGTGGTCTC | [] |
| 2679 | 7 | CPA4 | {'exs': 1, 'nonexs': 7} | True | 1.3E+08 | TCGCCTAGTAGAGGTTGTCCC | [] |
| 2680 | 7 | CPA5 | {'exs': 0, 'nonexs': 7} | True | 1.3E+08 | GCTGCTAGTAGGGGTGATTCA | [] |
| 2681 | 7 | KLF14 | {'exs': 0, 'nonexs': 4} | True | 1.31E+08 | ACTGTCCTACAGGCAGGTGGT | [] |
| 2682 | 7 | PODXL | {'exs': 1, 'nonexs': 6} | True | 1.32E+08 | CAGACCGGACTAGAGGTGTGT | [] |
| 2683 | 7 | CHCHD3 | {'exs': 2, 'nonexs': 5} | False | 1.33E+08 | GGCTAAGCAGTCAACACTCCA | [] |
| 2684 | 7 | EXOC4 | {'exs': 5, 'nonexs': 3} | False | 1.33E+08 | AGTCCTATGGCCACAGTAATAG | [] |
| 2685 | 7 | EXOC4 | {'exs': 5, 'nonexs': 3} | False | 1.33E+08 | CGCCGCTACCCTGCAACGGAA | [] |
| 2686 | 7 | EXOC4 | {'exs': 5, 'nonexs': 3} | False | 1.34E+08 | AGTACGCCCTGCTAAACGGTA | [] |
| 2687 | 7 | LRGUK | {'exs': 1, 'nonexs': 6} | True | 1.34E+08 | CTGAAATACTAGCTTAGACAAG | [] |
| 2688 | 7 | LRGUK | {'exs': 1, 'nonexs': 6} | True | 1.34E+08 | CTAGTCTACCTGCGGCCCTGA | [] |
| 2689 | 7 | BPGM | {'exs': 0, 'nonexs': 7} | True | 1.35E+08 | AGAAAGACTATTTTTTAGCTTG | [] |
| 2690 | 7 | CYREN | unavailable | False | 1.35E+08 | GTTTATGCCCTAGCTGAAAAAG | [] |
| 2691 | 7 | STRA8 | {'exs': 0, 'nonexs': 7} | True | 1.35E+08 | TCAGGGCCTAGTCTGAACCAT | [] |
| 2692 | 7 | CNOT4 | {'exs': 2, 'nonexs': 6} | False | 1.35E+08 | TTTGCCTAATGGCGGTCCAGT | [] |

| | | | | | | | |
|---|---|---|---|---|---|---|---|
| 2683 | 7 | ATP6V0A4 | {'exs': 0, 'nonexs': 7} | True | 1.39E+08 | CTCAGCCTACTCCTCGGCTGT | [] |
| 2684 | 7 | ZC3HAV1 | {'exs': 1, 'nonexs': 7} | True | 1.39E+08 | TCGGTTCTAACTAATCACGCAG | [] |
| 2685 | 7 | TBXAS1 | {'exs': 0, 'nonexs': 7} | True | 1.4E+08 | AATGCTAAATCTCTGGATTCAG | [] |
| 2686 | 7 | PARP12 | {'exs': 1, 'nonexs': 7} | True | 1.4E+08 | TCTTACCCGTCTATTCACGTTG | [] |
| 2687 | 7 | MKRN1 | {'exs': 0, 'nonexs': 7} | True | 1.4E+08 | GCAAGGTTGCTATAGATCCAAG | [] |
| 2688 | 7 | DENND2A | {'exs': 0, 'nonexs': 7} | True | 1.41E+08 | TACAGACTAGAGGGTCCAAGAG | [] |
| 2689 | 7 | BRAF | {'exs': 7, 'nonexs': 4} | False | 1.41E+08 | GATGGTGGCTACTTGAAGGCTC | [] |
| 2700 | 7 | MRPS33 | {'exs': 3, 'nonexs': 4} | False | 1.41E+08 | CAACACTATTTCCTTTTTGCTG | [] |
| 2701 | 7 | DENND11 | unavailable | False | 1.42E+08 | CTCCTCCTACGGGCAACAGGG | [] |
| 2702 | 7 | SSBP1 | {'exs': 5, 'nonexs': 3} | False | 1.42E+08 | CCTTTCTACTCCTTCTCTTTCG | [] |
| 2703 | 7 | TAS2R3 | {'exs': 0, 'nonexs': 7} | True | 1.42E+08 | TGTTACTCTAAGAGAAAATGGG | [] |
| 2704 | 7 | TAS2R4 | {'exs': 0, 'nonexs': 7} | True | 1.42E+08 | TCCACTATTTTTTGAAACAAAG | [] |
| 2705 | 7 | OR9A4 | {'exs': 0, 'nonexs': 2} | False | 1.42E+08 | AGCAAGGCTAATTCCTGAATAG | [] |
| 2706 | 7 | OR9A2 | {'exs': 1, 'nonexs': 3} | False | 1.43E+08 | AGAACAGCTAATCTTTCAGGAG | [] |
| 2707 | 7 | FAM131B | {'exs': 0, 'nonexs': 7} | True | 1.43E+08 | AGGAAACTAGTTGTTGGCCTCG | [] |
| 2708 | 7 | TAS2R41 | {'exs': 0, 'nonexs': 7} | True | 1.43E+08 | TGCCATCTAGGCCACCCAGAA | [] |
| 2709 | 7 | OR6B1 | {'exs': 0, 'nonexs': 3} | True | 1.44E+08 | AATTGACTAGTCAGATCTGCTG | [] |
| 2710 | 7 | NOBOX | {'exs': 0, 'nonexs': 3} | True | 1.44E+08 | ACCCCCCAACTAGGGGACATG | [] |
| 2711 | 7 | TPK1 | {'exs': 2, 'nonexs': 6} | False | 1.45E+08 | GCTCCCTAGCCATCAGCATGT | [] |
| 2712 | 7 | EZH2 | {'exs': 5, 'nonexs': 3} | False | 1.49E+08 | ATCTCCTAAAGCCTAAAAAATG | ['AAAAAATAAAAATAAAAACTTG'] |
| 2713 | 7 | ZNF425 | {'exs': 0, 'nonexs': 7} | True | 1.49E+08 | GCCTGACCTAGAGGCTGGAGG | [] |
| 2714 | 7 | ZNF282 | {'exs': 0, 'nonexs': 7} | True | 1.49E+08 | CAGCCCTAGTCTCGCTCAGGA | [] |
| 2715 | 7 | ACTR3C | {'exs': 0, 'nonexs': 7} | True | 1.5E+08 | AACTTCCTAGGCGCGTGGAGG | [] |
| 2716 | 7 | ZNF775 | {'exs': 1, 'nonexs': 7} | True | 1.5E+08 | CCACTAGCGCGCCTCCTCCTT | [] |
| 2717 | 7 | GIMAP8 | {'exs': 1, 'nonexs': 7} | True | 1.5E+08 | GCTACCTATTGTAAAATACCTG | ['TACCTATTAAATTTTTAAGGAG'] |
| 2718 | 7 | GIMAP2 | {'exs': 1, 'nonexs': 7} | True | 1.51E+08 | ATCTGTAACTATAACCTAGGAG | [] |
| 2719 | 7 | TMEM176 | {'exs': 0, 'nonexs': 7} | True | 1.51E+08 | AGGCATGGCTAGATTCCACTCC | ['TCACTTCCAACATTTCCTTCTG'] |
| 2720 | 7 | KCNH2 | {'exs': 0, 'nonexs': 7} | True | 1.51E+08 | CCCCACTAACTGCCCGGGTCC | [] |
| 2721 | 7 | KCNH2 | {'exs': 0, 'nonexs': 7} | True | 1.51E+08 | ATTTGATCCTACTTTAAGGAAG | [] |
| 2722 | 7 | NOS3 | {'exs': 0, 'nonexs': 7} | True | 1.51E+08 | ATTAGCTAGGCATCATGGCATG | [] |
| 2723 | 7 | NOS3 | {'exs': 0, 'nonexs': 7} | True | 1.51E+08 | CGTCTATAATCGCAGCACTTTG | [] |
| 2724 | 7 | NOS3 | {'exs': 0, 'nonexs': 7} | True | 1.51E+08 | CGAAACTATTCTGTGGTGACAG | [] |
| 2725 | 7 | CDK5 | {'exs': 1, 'nonexs': 6} | True | 1.51E+08 | CGGGGCCTAGGGCGGACAGAA | [] |
| 2726 | 7 | SLC4A2 | {'exs': 0, 'nonexs': 7} | True | 1.51E+08 | GTGGCGGCTACACAGGCATGG | [] |
| 2727 | 7 | TMUB1 | {'exs': 0, 'nonexs': 7} | True | 1.51E+08 | GAGGCACTACGGGCGGTACAT | [] |
| 2728 | 7 | AGAP3 | {'exs': 0, 'nonexs': 7} | True | 1.51E+08 | TGGTCTTCTAATTTGGAAGGAG | [] |
| 2729 | 7 | ASB10 | {'exs': 1, 'nonexs': 7} | True | 1.51E+08 | GGGGCTCACCTAGTAGAGCAC | [] |
| 2730 | 7 | CHPF2 | {'exs': 0, 'nonexs': 7} | True | 1.51E+08 | CGGGCTAAGTGCTATTGGCCT | [] |
| 2731 | 7 | SMARCD3 | {'exs': 0, 'nonexs': 7} | True | 1.51E+08 | TTTTGGGCTCCTAGGTGTTGCC | [] |
| 2732 | 7 | WDR86 | {'exs': 0, 'nonexs': 7} | True | 1.51E+08 | AATCCCTACCTGTTCATCCGCC | [] |
| 2733 | 7 | WDR86 | {'exs': 0, 'nonexs': 7} | True | 1.51E+08 | TAAGACTATGAAGCGCATGCTG | [] |
| 2734 | 7 | CRYGN | {'exs': 0, 'nonexs': 7} | True | 1.51E+08 | GCTTCTAGGGCTCCATGCTCT | [] |
| 2735 | 7 | LOC64409 | unavailable | False | 1.52E+08 | GGCTGGCCTAGGGATTCACCT | [] |
| 2736 | 7 | GALNT11 | {'exs': 0, 'nonexs': 7} | True | 1.52E+08 | AGAGCAAAGCTAACGAGAAGTG | [] |
| 2737 | 7 | GALNT11 | {'exs': 0, 'nonexs': 7} | True | 1.52E+08 | CACCTCTGCTAACACACCCAG | [] |
| 2738 | 7 | ACTR3B | {'exs': 0, 'nonexs': 7} | True | 1.53E+08 | AAAAGCTACATAGTATTCCACG | [] |
| 2739 | 7 | ACTR3B | {'exs': 0, 'nonexs': 7} | True | 1.53E+08 | ACACTAGGACATGACTCCAAAG | [] |
| 2740 | 7 | EN2 | {'exs': 0, 'nonexs': 7} | True | 1.55E+08 | CCCCCGCCCTACTCGCTGTCC | [] |
| 2741 | 7 | CNPY1 | {'exs': 0, 'nonexs': 7} | True | 1.56E+08 | GGCACTCCTAGAGCTCAGTATC | [] |
| 2742 | 7 | RNF32 | {'exs': 0, 'nonexs': 7} | True | 1.57E+08 | TTTCCTAAGTATCTGTTGCCTG | [] |
| 2743 | 7 | NOM1 | {'exs': 5, 'nonexs': 3} | False | 1.57E+08 | CCTGACTACATTCTCAGGGAAG | [] |
| 2744 | 7 | MNX1 | {'exs': 0, 'nonexs': 7} | True | 1.57E+08 | GGGGCTCCTACTGGGGCGCGC | [] |
| 2745 | 7 | DNAJB6 | {'exs': 1, 'nonexs': 6} | True | 1.57E+08 | TCCGGTCTAGTGATTGCCTTTG | [] |
| 2746 | 7 | PTPRN2 | {'exs': 1, 'nonexs': 6} | True | 1.58E+08 | CAGTCGTCTACCTTCCCTGCC | [] |
| 2747 | 7 | NCAPG2 | {'exs': 4, 'nonexs': 4} | False | 1.59E+08 | CAGTCCTAGCCAATTGGGACTC | [] |
| 2748 | 7 | ESYT2 | {'exs': 0, 'nonexs': 7} | True | 1.59E+08 | TGCGGCTATGTCATCGCCTGA | [] |
| 2749 | 7 | WDR60 | {'exs': 0, 'nonexs': 7} | True | 1.59E+08 | GCCACACCCGCTACTTGTGCA | [] |
| 2750 | 7 | VIPR2 | {'exs': 0, 'nonexs': 7} | True | 1.59E+08 | GGGTGGGGCTAGATGACCGAG | [] |
| 2751 | 8 | ERICH1 | {'exs': 0, 'nonexs': 7} | True | 627098 | TATCCCTAGACACCTGCATCAG | [] |
| 2752 | 8 | ERICH1 | {'exs': 0, 'nonexs': 7} | True | 637372 | CACGTTCCTCTAGCCCCATGT | [] |
| 2753 | 8 | CLN8 | {'exs': 0, 'nonexs': 7} | True | 1780566 | GCAGCTATGGCCTCTTCTTCCC | [] |
| 2754 | 8 | KBTBD11 | {'exs': 0, 'nonexs': 7} | True | 2003063 | CGCCGGCCTACGGGGCGCCC | [] |
| 2755 | 8 | CSMD1 | {'exs': 0, 'nonexs': 5} | True | 2938584 | TGAGGGCTATACCACTGTACAG | [] |
| 2756 | 8 | MCPH1 | {'exs': 4, 'nonexs': 5} | False | 6626119 | TAATCACTAACAATTTTTATCG | [] |
| 2757 | 8 | MCPH1 | {'exs': 4, 'nonexs': 5} | False | 6637940 | CTCTAATATGCCACATCCAATG | [] |
| 2758 | 8 | AGPAT5 | {'exs': 2, 'nonexs': 5} | False | 6757387 | CTACTTGTCTATGCTTTAATAG | [] |
| 2759 | 8 | LOC10192 | unavailable | False | 7216432 | TTCCTAGCCCCAGACCCATCT | [] |
| 2760 | 8 | LOC10192 | unavailable | False | 7217234 | AAATATCTATATCTCGGCAGAG | [] |
| 2761 | 8 | LOC10192 | unavailable | False | 7219169 | CTAGTGGTCTACGCAGACAAAG | ['CAGACAAAATCTCATAAAAGGG'] |
| 2762 | 8 | SPAG11B | {'exs': 0, 'nonexs': 7} | True | 7450878 | TGGTATTTCTAATTCCCGGTGG | [] |
| 2763 | 8 | LOC10798 | unavailable | False | 7613365 | CTAGTGGTCTACGCAGACAAAG | ['CAGACAAAATCTCATAAAAGGG'] |
| 2764 | 8 | LOC10192 | unavailable | False | 7677765 | CTAGTGGTCTACGCAGACAAAG | ['CAGACAAAATCTCATAAAAGGG'] |
| 2765 | 8 | SPAG11A | {'exs': 0, 'nonexs': 7} | True | 7860666 | TGGTATTTCTAATTCCCAGTGG | [] |
| 2766 | 8 | CLDN23 | {'exs': 1, 'nonexs': 5} | False | 8703276 | AAGCGTCTAGAGGTCGGAGTC | [] |
| 2767 | 8 | PPP1R3B | {'exs': 1, 'nonexs': 6} | True | 9140793 | GTCACTAGTAGTAGGGCCCTA | [] |
| 2768 | 8 | TNKS | {'exs': 1, 'nonexs': 6} | True | 9776735 | ATTCACTAGGTCTTCTGCTCTG | [] |
| 2769 | 8 | MSRA | {'exs': 0, 'nonexs': 7} | True | 10250468 | ATTTCTAAAAGCTTTTGAATAG | [] |
| 2770 | 8 | RP1L1 | {'exs': 0, 'nonexs': 7} | True | 10606894 | TGATCTTGTCTAGAAATCTAAG | [] |
| 2771 | 8 | C8orf74 | {'exs': 0, 'nonexs': 7} | True | 10700470 | ACCTTCTACTTCCTTGCCTTCG | [] |
| 2772 | 8 | SOX7 | {'exs': 0, 'nonexs': 7} | True | 10725737 | CCAGCTCTATGACACACTGTAG | [] |
| 2773 | 8 | BLK | {'exs': 1, 'nonexs': 6} | True | 11564107 | GGCCGGCTAGGGCTGCAGCTC | [] |
| 2774 | 8 | DEFB135 | {'exs': 0, 'nonexs': 7} | True | 11984589 | CTCACAACTATTTCCCAGTCAG | [] |
| 2775 | 8 | DEFB130B | unavailable | False | 12064388 | CACCTAAGGAGATTTTCCTTTG | [] |
| 2776 | 8 | FAM86B1 | {'exs': 0, 'nonexs': 7} | True | 12183605 | AGTCCTACAGTGTGAGGTTCAG | [] |
| 2777 | 8 | FAM86B1 | {'exs': 0, 'nonexs': 7} | True | 12185375 | GGGTTCACCTAGCTCGGTGGT | [] |
| 2778 | 8 | DEFB130A | unavailable | False | 12310961 | CACCTAAGGAGATTTTCCTTTG | [] |
| 2779 | 8 | FAM86B2 | {'exs': 0, 'nonexs': 5} | True | 12425888 | AGTCCTACAGTGTGAGGTTCAG | [] |
| 2780 | 8 | DLC1 | {'exs': 0, 'nonexs': 7} | True | 13305224 | TTTATTCTAAAATAGATTGGCG | [] |
| 2781 | 8 | MICU3 | unavailable | False | 17098181 | TGCCCCTCTACTGAAACGTGT | [] |
| 2782 | 8 | MICU3 | unavailable | False | 17102729 | GTCACATACTATGTTTTAAATG | [] |
| 2783 | 8 | VPS37A | {'exs': 2, 'nonexs': 6} | False | 17286426 | AATTTACCTATAGTGGAGCATG | [] |
| 2784 | 8 | NAT1 | {'exs': 0, 'nonexs': 6} | True | 18222919 | CCTTATTCTAAATAGTAAAAAG | ['AAAAAATCTATCACCATGTTTG'] |
| 2785 | 8 | NAT2 | {'exs': 1, 'nonexs': 7} | True | 18400875 | TCCTTATTCTAAATAGTAAGGG | [] |
| 2786 | 8 | PSD3 | {'exs': 0, 'nonexs': 7} | True | 18535742 | ATGGACTCTAAGTAACTTTTTG | [] |
| 2787 | 8 | PSD3 | {'exs': 0, 'nonexs': 7} | True | 18774914 | ATCTTGGCTACGGCTGCAAAAG | [] |
| 2788 | 8 | SLC18A1 | {'exs': 0, 'nonexs': 7} | True | 20145762 | TGCTGCTACTCCTCATGGTCA | [] |
| 2789 | 8 | ATP6V1B2 | {'exs': 6, 'nonexs': 2} | False | 20220401 | CAGCAGCTAATGCTTTGCAGAG | [] |
| 2790 | 8 | GFRA2 | {'exs': 0, 'nonexs': 7} | True | 21693277 | CAGCCTACAAGGCCAGTTTCA | [] |
| 2791 | 8 | FGF17 | {'exs': 1, 'nonexs': 6} | True | 22049248 | CCCAGACTACGTGAGGGGCTG | [] |
| 2792 | 8 | NUDT18 | {'exs': 0, 'nonexs': 1} | False | 22107299 | ACCTCTCTATCTGTTCACTGGG | [] |
| 2793 | 8 | HR | {'exs': 2, 'nonexs': 6} | False | 22115699 | CATCCCTCTATTTGGCCTCCT | [] |
| 2794 | 8 | REEP4 | {'exs': 0, 'nonexs': 7} | True | 22139486 | AGACCCTAGCTGTCCACGTCT | [] |
| 2795 | 8 | LGI3 | {'exs': 0, 'nonexs': 7} | True | 22148159 | TCGGCACCCCCTAGGCACTGA | [] |
| 2796 | 8 | SFTPC | {'exs': 0, 'nonexs': 7} | True | 22164040 | GCGTCCTAGATGTAGTAGAGCC | [] |
| 2797 | 8 | PHYHIP | {'exs': 0, 'nonexs': 7} | True | 22221352 | GAGTCCCTAGCGGCCCACGCT | [] |
| 2798 | 8 | SLC39A14 | {'exs': 0, 'nonexs': 7} | True | 22419697 | AGCCCTACCCAATCTGGATCT | [] |
| 2799 | 8 | BIN3 | {'exs': 0, 'nonexs': 7} | True | 22625300 | GGGCGTCTACCAGGCAAGCCA | [] |
| 2800 | 8 | PEBP4 | {'exs': 0, 'nonexs': 7} | True | 22713369 | ATCTAGCAGGCAGCTATCTCCC | [] |
| 2801 | 8 | CHMP7 | {'exs': 1, 'nonexs': 5} | False | 23260510 | ACACTATAACAGGAAAATCTTG | [] |
| 2802 | 8 | CHMP7 | {'exs': 1, 'nonexs': 5} | False | 23260598 | GGGTCCTACAATGGCTTTAGAG | [] |
| 2803 | 8 | ADAM28 | {'exs': 0, 'nonexs': 7} | True | 24343023 | AGGCTAGCAGATGTTGTTTAAG | [] |
| 2804 | 8 | ADAM7 | {'exs': 0, 'nonexs': 7} | True | 24507538 | TAACCTACCACTTGGCACTTTG | [] |
| 2805 | 8 | PPP2R2A | {'exs': 4, 'nonexs': 4} | False | 26370412 | AACCCTAATTCACTTTGTCTTG | [] |
| 2806 | 8 | DPYSL2 | {'exs': 0, 'nonexs': 7} | True | 26655705 | GGAGCTCTAGCCCAGGCTGGT | [] |
| 2807 | 8 | ADRA1A | {'exs': 0, 'nonexs': 1} | False | 26770148 | CCTGTCCTAGACTTCCTCCCC | [] |
| 2808 | 8 | STMN4 | {'exs': 0, 'nonexs': 7} | True | 27239784 | TGGTTCCTATTTCTTGCCTCTG | [] |
| 2809 | 8 | EPHX2 | {'exs': 0, 'nonexs': 7} | True | 27544521 | CGCTGCGTTCTACATCTTTGAG | [] |
| 2810 | 8 | SCARA3 | {'exs': 0, 'nonexs': 7} | True | 27718965 | TATCCTCTATTTACGTCTTGAG | [] |
| 2811 | 8 | PBK | {'exs': 0, 'nonexs': 7} | True | 27810304 | ATCACTAGACATCTGTTTCCAG | [] |
| 2812 | 8 | ELP3 | {'exs': 5, 'nonexs': 3} | False | 28183251 | TGCCTAGAGTGAACAGCTTCTG | [] |
| 2813 | 8 | PNOC | {'exs': 1, 'nonexs': 7} | True | 28339443 | CCGGCTACACATTACCATTCTG | [] |
| 2814 | 8 | EXTL3 | {'exs': 2, 'nonexs': 6} | False | 28750865 | CTGCCCCTAGATGAACTTGAAG | [] |
| 2815 | 8 | INTS9 | {'exs': 6, 'nonexs': 2} | False | 28769606 | CCTCACTACCCTGCCCCACTC | [] |
| 2816 | 8 | HMBOX1 | {'exs': 1, 'nonexs': 7} | True | 29042701 | CAATCTCCTACCCTGCAGAAAG | [] |
| 2817 | 8 | HMBOX1 | {'exs': 1, 'nonexs': 7} | True | 29046213 | GTAGGGTCTACATTGCTAGAAG | [] |
| 2818 | 8 | DUSP4 | {'exs': 1, 'nonexs': 6} | True | 29337025 | GCGGCTCTAACAGCTGGGAGA | [] |
| 2819 | 8 | SARAF | unavailable | False | 30069334 | CCCAAACTATTTCTGTACTTTG | ['TACTTTACTCCTAGGTAAAAGG'] |
| 2820 | 8 | RBPMS | {'exs': 0, 'nonexs': 7} | True | 30545174 | TCCTTTTCTACAGGGTTTCCTG | [] |
| 2821 | 8 | GTF2E2 | {'exs': 6, 'nonexs': 2} | False | 30578920 | ACTGTTCCCTATTTGCTGGAAG | [] |
| 2822 | 8 | UBXN8 | unavailable | False | 30766393 | CCCTTCTTTCCTAGTTGGTCTG | [] |
| 2823 | 8 | LOC10537 | unavailable | False | 30743774 | GTGGGCTATAACACCTTCCTTG | ['CTTATCTCCCTAACGGTTCCTG'] |
| 2824 | 8 | PURG | {'exs': 0, 'nonexs': 7} | True | 31031738 | CACTCTAGTCGAGGCATTCTTG | [] |
| 2825 | 8 | NRG1 | {'exs': 1, 'nonexs': 6} | True | 32742767 | GCTCCTATTCAGGCAGAGACA | [] |
| 2826 | 8 | FUT10 | {'exs': 0, 'nonexs': 7} | True | 33308811 | ACAAAAAGAGCTAGGGTCCCAG | [] |
| 2827 | 8 | UNC5D | {'exs': 0, 'nonexs': 7} | True | 35790562 | AAGTGGACTAGAGTCCATTTTG | [] |
| 2828 | 8 | KCNU1 | {'exs': 0, 'nonexs': 7} | True | 36935919 | GGGCAGGTCTATAGTGGCTCT | [] |
| 2829 | 8 | GOT1L1 | {'exs': 0, 'nonexs': 7} | True | 37934292 | ACTAAAGTTTTATTCCAATCAG | [] |
| 2830 | 8 | ADRB3 | {'exs': 0, 'nonexs': 7} | True | 37964217 | CAGGCCTAAGAAACTCCCCAA | [] |
| 2831 | 8 | PLPP5 | unavailable | False | 38267198 | CCATCCCTACTAGCTTTCTAGG | [] |
| 2832 | 8 | PLPP5 | unavailable | False | 38267347 | ATGGGCTAGCCCATCAGGGAA | [] |
| 2833 | 8 | FGFR1 | {'exs': 3, 'nonexs': 5} | False | 38412388 | CGTGGCTATTCCTAGGTAGGT | [] |
| 2834 | 8 | HTRA4 | {'exs': 0, 'nonexs': 7} | True | 38979990 | TCTCATCTATAAGCAGGTCAAG | [] |
| 2835 | 8 | ADAM32 | {'exs': 0, 'nonexs': 7} | True | 39284798 | GAATCACTAGTTACTGTGGAAG | [] |
| 2836 | 8 | ADAM2 | {'exs': 0, 'nonexs': 7} | True | 39744859 | GTTGTCCAGACTACCCTTTAGG | [] |
| 2837 | 8 | IDO2 | unavailable | False | 40015601 | CAGCCTCCTAACCACGTGGGT | [] |
| 2838 | 8 | ZMAT4 | {'exs': 0, 'nonexs': 7} | True | 40532222 | CTTTCACTACTTATTCTTCAGG | [] |
| 2839 | 8 | IKBKB | {'exs': 3, 'nonexs': 5} | False | 42325324 | AAAGGCTATGGGAATTCCTGTG | [] |
| 2840 | 8 | SLC20A2 | {'exs': 0, 'nonexs': 7} | True | 42419625 | GGAGTTCTACTTCAAGGAAGAG | [] |
| 2841 | 8 | CHRNB3 | {'exs': 0, 'nonexs': 7} | True | 42736517 | TAAATTCCTAATGGTAACTATG | [] |
| 2842 | 8 | HOOK3 | {'exs': 0, 'nonexs': 7} | True | 43019497 | CTTCTCTACCTTGCTGTGGCC | [] |
| 2843 | 8 | POTEA | unavailable | False | 43358894 | ATAGCTATTTTTGTAGTACATG | [] |
| 2844 | 8 | LOC10537 | unavailable | False | 47100953 | CTGTTCCTATTCGGCCATCTTG | [] |
| 2845 | 8 | SPIDR | unavailable | False | 47732653 | CATCTCTACCCAGGCCCCTCT | [] |
| 2846 | 8 | SPIDR | unavailable | False | 47735449 | CAACCGCTAGTGTTCTGCAGA | [] |
| 2847 | 8 | PPDPFL | unavailable | False | 49074530 | GGTGCTATGATCTTCCTGACAG | [] |
| 2848 | 8 | PPDPFL | unavailable | False | 49075172 | TCAGCTACAAAGTGCTCATCTG | [] |
| 2849 | 8 | ST18 | {'exs': 0, 'nonexs': 7} | True | 52113197 | GCTGTGATCCTACACATGGATG | ['CATGGATACCCTTCACTGCCTG'] |
| 2850 | 8 | ALKAL1 | unavailable | False | 52538442 | TTTGCTAGGTCTGGGAGCACA | [] |
| 2851 | 8 | RGS20 | {'exs': 0, 'nonexs': 7} | True | 53959457 | GAAAAATCCTATGCTTCAATAG | [] |
| 2852 | 8 | LYN | {'exs': 1, 'nonexs': 6} | True | 56010109 | TGTGCTCTAAGGCTGCTGCTG | [] |
| 2853 | 8 | PLAG1 | {'exs': 0, 'nonexs': 7} | True | 56166242 | CAGAATCCTACTGAAAAGCTTG | [] |
| 2854 | 8 | CA8 | {'exs': 0, 'nonexs': 7} | True | 60208784 | CTCTTTGGCTACTGAAATGCAG | [] |
| 2855 | 8 | CA8 | {'exs': 0, 'nonexs': 7} | True | 60208915 | TTCTTCTATCTGAAAATAAAAG | [] |
| 2856 | 8 | CA8 | {'exs': 0, 'nonexs': 7} | True | 60256482 | GAACCCTACAAAGCTAAAAGAG | [] |
| 2857 | 8 | CHD7 | {'exs': 2, 'nonexs': 6} | False | 60844764 | AAACTAGCAACAATTCACACTG | [] |
| 2858 | 8 | ASPH | {'exs': 1, 'nonexs': 7} | True | 61503358 | TTCATGCTAAATTGCTGGAAGG | [] |
| 2859 | 8 | NKAIN3 | {'exs': 0, 'nonexs': 5} | True | 62965406 | TTGCTCCCTAAGTCAAGTTCAG | [] |
| 2860 | 8 | NKAIN3 | {'exs': 0, 'nonexs': 5} | True | 62999333 | TCCACTCTTGCTATGTGATGTG | [] |
| 2861 | 8 | CYP7B1 | {'exs': 0, 'nonexs': 7} | True | 64587592 | TAACTATTGCAAAATTATCAAG | [] |
| 2862 | 8 | CYP7B1 | {'exs': 0, 'nonexs': 7} | True | 64596641 | GCTTCTCTAAGATTTCACTTTG | [] |
| 2863 | 8 | MCMDC2 | unavailable | False | 66901351 | AATACCTACAGATATTTTAATG | [] |
| 2864 | 8 | TCF24 | {'exs': 1, 'nonexs': 4} | False | 66948050 | AGAGCCTAAGGCTGTGAGTCT | [] |
| 2865 | 8 | SULF1 | {'exs': 0, 'nonexs': 7} | True | 69659321 | TTCATCTAATCAGAGGAGAGTG | [] |
| 2866 | 8 | PRDM14 | {'exs': 0, 'nonexs': 7} | True | 70052076 | GGCAGGGCTAGTAGTCTTCAT | [] |
| 2867 | 8 | LACTB2 | {'exs': 0, 'nonexs': 7} | True | 70637859 | TTAATCTGAAACTAAAGATGAG | [] |
| 2868 | 8 | EYA1 | {'exs': 0, 'nonexs': 7} | True | 71201467 | GTGGGCTAAGAGACACCAGAA | [] |
| 2869 | 8 | TRPA1 | {'exs': 0, 'nonexs': 7} | True | 72022905 | GAGGAGCTAAGGCTCAAGATGC | [] |
| 2870 | 8 | SBSPON | unavailable | False | 73067340 | CATCTATATAAAAATAAAACTG | [] |
| 2871 | 8 | SBSPON | unavailable | False | 73074520 | CTCACATCTAAATATAGCACAG | [] |
| 2872 | 8 | STAU2 | {'exs': 1, 'nonexs': 7} | True | 73421371 | AGCTGCTAGACGGCCGAGTTT | [] |
| 2873 | 8 | LY96 | {'exs': 2, 'nonexs': 7} | False | 74029053 | ATTTATTCTAATTTGAATTAGG | [] |
| 2874 | 8 | GDAP1 | {'exs': 0, 'nonexs': 7} | True | 74364366 | AACAAACCTAGAAATAATTTGG | [] |
| 2875 | 8 | GDAP1 | {'exs': 0, 'nonexs': 7} | True | 74488774 | CAGCTCTGCTATTGATACACTG | [] |
| 2876 | 8 | ZFHX4 | {'exs': 0, 'nonexs': 7} | True | 76864564 | TCACTCCTACACACTGAACATG | ['CATATCCATTCGGATGCTATTG'] |
| 2877 | 8 | PEX2 | {'exs': 0, 'nonexs': 7} | True | 76983260 | TTCTAAAGAGCATTTACTTCTG | [] |
| 2878 | 8 | IL7 | {'exs': 1, 'nonexs': 6} | True | 78721350 | TTTACCTTCTAGTGGAGAAAGG | [] |
| 2879 | 8 | PAG1 | {'exs': 0, 'nonexs': 7} | True | 80976543 | CTTCTGGGTTGCTAGAGCCTGC | [] |
| 2880 | 8 | ZFAND1 | {'exs': 0, 'nonexs': 7} | True | 81702694 | TTGAATGACTATTCCAAGTAAG | [] |
| 2881 | 8 | E2F5 | {'exs': 1, 'nonexs': 6} | True | 85213861 | GAATCTAATAATTTAGTATCTG | [] |
| 2882 | 8 | LOC10192 | unavailable | False | 85737340 | CTGGGGCTAAGGACAGGCCAG | [] |
| 2883 | 8 | LOC10192 | unavailable | False | 85744688 | CTGGGGCTAAGGACAGGCCAG | [] |
| 2884 | 8 | LOC10028 | unavailable | False | 85790324 | CTGGGGCTAAGGACAGGCCAG | [] |
| 2885 | 8 | SLC7A13 | {'exs': 0, 'nonexs': 7} | True | 86214412 | ATCTATTCCTCAGACACATCAG | [] |
| 2886 | 8 | RMDN1 | unavailable | False | 86470152 | CATCTATAATGGAGAGATATAG | [] |
| 2887 | 8 | RMDN1 | unavailable | False | 86475534 | TTTACTAGATATACAAACTTTG | [] |
| 2888 | 8 | CNBD1 | {'exs': 0, 'nonexs': 5} | True | 87382526 | TAGATCTAGGCCACTGCAAAAG | [] |
| 2889 | 8 | CALB1 | {'exs': 0, 'nonexs': 7} | True | 90060172 | CACCAACTCTAGTTATCCCCAG | [] |
| 2890 | 8 | NECAB1 | {'exs': 0, 'nonexs': 7} | True | 90955511 | TAGGAACATCTAGTTGTTCAGG | [] |
| 2891 | 8 | C8orf88 | unavailable | False | 90959006 | AAAATGCTACTTAAAACTTTGG | [] |
| 2892 | 8 | LRRC69 | {'exs': 0, 'nonexs': 3} | True | 91218999 | CCTCACCTGTTCTACACCTGA | [] |
| 2893 | 8 | RUNX1T1 | {'exs': 0, 'nonexs': 7} | True | 91960241 | GTTCACGTCTAGCGAGGGGTT | [] |
| 2894 | 8 | TRIQK | unavailable | False | 92886521 | GCTTAGCTAATCTTCATCTTGG | [] |
| 2895 | 8 | RBM12B | {'exs': 1, 'nonexs': 7} | True | 93733404 | ATGCTCTCTCTCTACAGCAAAG | [] |
| 2896 | 8 | PDP1 | {'exs': 0, 'nonexs': 7} | True | 93923672 | CCACTCACTATTCTTGGTTTTG | [] |
| 2897 | 8 | GEM | {'exs': 1, 'nonexs': 7} | True | 94250309 | GGGTTCCTAGAGTACAGAGAGC | [] |
| 2898 | 8 | ESRP1 | {'exs': 0, 'nonexs': 7} | True | 94690941 | GAGGATGTTCTAACTGCTGGGC | [] |
| 2899 | 8 | GDF6 | {'exs': 0, 'nonexs': 7} | True | 96144562 | CACCGCTACCTGCAGCCGCAC | [] |
| 2900 | 8 | MTERF3 | unavailable | False | 96239490 | AAAAATCTAAAGCGTTTTTAAG | [] |
| 2901 | 8 | CPQ | unavailable | False | 97142182 | ACTGTTTCTAGGACCTAGGCAG | [] |
| 2902 | 8 | ERICH5 | unavailable | False | 98093332 | CTATCTATGTGGCTGCTGAAAG | [] |
| 2903 | 8 | NIPAL2 | {'exs': 0, 'nonexs': 7} | True | 98193050 | ATAAGCTATGTGAATCTGGTTG | [] |
| 2904 | 8 | KCNS2 | {'exs': 0, 'nonexs': 7} | True | 98429412 | AGTCCTCCCGGCTAACGTAGG | [] |
| 2905 | 8 | OSR2 | {'exs': 0, 'nonexs': 7} | True | 98951700 | CTTCTCTAGAAGTCCTGCCGC | [] |
| 2906 | 8 | RNF19A | {'exs': 1, 'nonexs': 7} | True | 1E+08 | TATGGGCCTAAATTTCAGTCTG | [] |
| 2907 | 8 | YWHAZ | {'exs': 1, 'nonexs': 6} | True | 1.01E+08 | ACAGGCTACAGCCGCTCCGCA | [] |
| 2908 | 8 | GRHL2 | {'exs': 1, 'nonexs': 7} | True | 1.01E+08 | TGAGGCTAGTCCTTCCCCTCC | ['CCTCCACCCCTGAACAGGTCCG'] |
| 2909 | 8 | GRHL2 | {'exs': 1, 'nonexs': 7} | True | 1.02E+08 | CAGGGCTAGATTTCCATGAGCC | [] |
| 2910 | 8 | UBR5 | {'exs': 3, 'nonexs': 5} | False | 1.02E+08 | CTACACAAAACCAAAATTCTTG | [] |
| 2911 | 8 | BAALC | {'exs': 0, 'nonexs': 7} | True | 1.03E+08 | GACTCTCTGCTAGTTGACACAG | [] |
| 2912 | 8 | CTHRC1 | {'exs': 0, 'nonexs': 7} | True | 1.03E+08 | ACATCCACTAATCCAGCACCAG | ['CAGCACCAATTCCTTCACAAAG'] |
| 2913 | 8 | RIMS2 | {'exs': 0, 'nonexs': 7} | True | 1.04E+08 | AGCTGCTATGAACGAGAGTAAG | [] |
| 2914 | 8 | LRP12 | {'exs': 0, 'nonexs': 7} | True | 1.04E+08 | CGTACCTAACAAAGTAACAAAG | [] |
| 2915 | 8 | SLC30A8 | {'exs': 0, 'nonexs': 7} | True | 1.17E+08 | GTGACTGAGCTAGTCACAGGG | [] |
| 2916 | 8 | ZHX2 | {'exs': 0, 'nonexs': 7} | True | 1.23E+08 | TTACCTGTCTAGGCCTGGCCA | [] |
| 2917 | 8 | TBC1D31 | unavailable | False | 1.23E+08 | ATGCATTCTATGCAGCCAAAAG | [] |
| 2918 | 8 | FAM83A | {'exs': 0, 'nonexs': 7} | True | 1.23E+08 | TTCCCTCTAACAAACCCTGCAG | [] |
| 2919 | 8 | C8orf76 | {'exs': 0, 'nonexs': 7} | True | 1.23E+08 | TAACCCACTAAGCCAAGATTTG | [] |
| 2920 | 8 | FER1L6 | {'exs': 0, 'nonexs': 5} | True | 1.24E+08 | GATCCTCTATGAGCCCACAACC | [] |
| 2921 | 8 | TRMT12 | {'exs': 1, 'nonexs': 7} | True | 1.24E+08 | ACCTCCTCTAGCCAACTGAAGC | [] |
| 2922 | 8 | NDUFB9 | {'exs': 3, 'nonexs': 5} | False | 1.25E+08 | CTTTCTACATGGGCCGCTCCC | [] |
| 2923 | 8 | MTSS1 | {'exs': 0, 'nonexs': 7} | True | 1.25E+08 | GTGAACCTAAGAAAAGCGAGGG | [] |
| 2924 | 8 | NSMCE2 | {'exs': 3, 'nonexs': 5} | False | 1.25E+08 | GCTTTTCCTACTCGGAATGACG | [] |
| 2925 | 8 | GSDMC | {'exs': 0, 'nonexs': 7} | True | 1.3E+08 | GTTCTACAGACCAGCCTTTAAG | [] |
| 2926 | 8 | ASAP1 | {'exs': 0, 'nonexs': 5} | True | 1.3E+08 | TTTGCTAGTCAGACAGGATATG | [] |
| 2927 | 8 | OC90 | {'exs': 0, 'nonexs': 1} | False | 1.32E+08 | GGCATCTATCTTCCATGAAGAG | [] |
| 2928 | 8 | SLA | {'exs': 0, 'nonexs': 7} | True | 1.33E+08 | CTTGGCTAGTCCTCAAAGTAAG | [] |
| 2929 | 8 | CCN4 | unavailable | False | 1.33E+08 | CCTGCCTAGTTGGCAATTTCTG | [] |
| 2930 | 8 | NDRG1 | {'exs': 0, 'nonexs': 7} | True | 1.33E+08 | CGCCTAGCAGGAGACCTCCAT | [] |
| 2931 | 8 | ZFAT | {'exs': 0, 'nonexs': 7} | True | 1.34E+08 | GTCCTCTAGAGTTCCTGGGCC | [] |
| 2932 | 8 | FAM135B | {'exs': 0, 'nonexs': 7} | True | 1.38E+08 | AAGCCACTACTTGAAGTAGTTG | [] |
| 2933 | 8 | KCNK9 | {'exs': 0, 'nonexs': 5} | True | 1.4E+08 | CCACACCTAAACGGACTTCGG | [] |
| 2934 | 8 | LOC10798 | unavailable | False | 1.41E+08 | GGGTGACTACTTAATCCACATG | [] |
| 2935 | 8 | SLC45A4 | {'exs': 0, 'nonexs': 7} | True | 1.41E+08 | TTCTTCTAAGAGAACCACATTG | [] |
| 2936 | 8 | GPR20 | {'exs': 1, 'nonexs': 7} | True | 1.41E+08 | GACTAAGCCTCGGGCCCATTA | [] |
| 2937 | 8 | PTP4A3 | {'exs': 1, 'nonexs': 7} | True | 1.41E+08 | TGAGCTACATAACGCAGCACCC | [] |
| 2938 | 8 | TSNARE1 | {'exs': 0, 'nonexs': 5} | True | 1.42E+08 | CAGCCCTATATCTCCTCCTCC | ['TCCACCTTCAAGATCATCCCTG'] |
| 2939 | 8 | TSNARE1 | {'exs': 0, 'nonexs': 5} | True | 1.42E+08 | GGCCTAGTTCCAGCCCCTCCA | [] |
| 2940 | 8 | ARC | {'exs': 1, 'nonexs': 6} | True | 1.43E+08 | CCCTCTACTCGGGCTGGGTCC | [] |
| 2941 | 8 | PSCA | {'exs': 0, 'nonexs': 5} | True | 1.43E+08 | CCAGAGCCTAGAGCTGGCCGG | [] |
| 2942 | 8 | LYPD2 | {'exs': 1, 'nonexs': 7} | True | 1.43E+08 | ACTCTACAGTCGGAGGCTCAA | [] |
| 2943 | 8 | CYP11B2 | {'exs': 0, 'nonexs': 7} | True | 1.43E+08 | GCAAGACTAGTTAATCGCTCTG | [] |
| 2944 | 8 | GLI4 | {'exs': 1, 'nonexs': 5} | False | 1.43E+08 | CGCCCGGCTACTCGCGGTAGT | [] |
| 2945 | 8 | MAFA | {'exs': 0, 'nonexs': 7} | True | 1.43E+08 | CGGCGCCTACAGGAAGAAGTC | [] |
| 2946 | 8 | ZC3H3 | {'exs': 6, 'nonexs': 2} | False | 1.43E+08 | TGCTGTCTAAAGAGACTCACAG | [] |
| 2947 | 8 | GSDMD | {'exs': 1, 'nonexs': 7} | True | 1.44E+08 | CACAGGCTAGTGGGGCTCCTG | [] |
| 2948 | 8 | NAPRT | unavailable | False | 1.44E+08 | CTTCCTGCTACCTCGAGCTCA | [] |
| 2949 | 8 | PYCR3 | unavailable | False | 1.44E+08 | GCCCAGCCTACTTTCTGCTGA | [] |
| 2950 | 8 | ZNF707 | {'exs': 0, 'nonexs': 7} | True | 1.44E+08 | GGCGCCCCTACACCTCCCCGT | [] |
| 2951 | 8 | IQANK1 | unavailable | False | 1.44E+08 | CAGCACTAGAGGCCTGTGCCT | [] |
| 2952 | 8 | SCRIB | {'exs': 2, 'nonexs': 6} | False | 1.44E+08 | GCTCCTCTAGGAGGGCACAGG | [] |
| 2953 | 8 | NRBP2 | {'exs': 1, 'nonexs': 0} | False | 1.44E+08 | CTGATCCTAAGGACCTGGGAG | [] |
| 2954 | 8 | GRINA | {'exs': 0, 'nonexs': 5} | True | 1.44E+08 | GCTCGGCTACTCCTTGGCGCG | [] |
| 2955 | 8 | SHARPIN | {'exs': 2, 'nonexs': 6} | False | 1.44E+08 | GGCTGCTAGGTGGAAGCTGCA | [] |
| 2956 | 8 | BOP1 | {'exs': 2, 'nonexs': 3} | False | 1.44E+08 | GAACAGCTAGGTGAAGAGGCG | [] |
| 2957 | 8 | SCX | unavailable | False | 1.44E+08 | ACCTCCTAACTGCGAATCGCTC | [] |
| 2958 | 8 | HSF1 | {'exs': 0, 'nonexs': 1} | False | 1.44E+08 | GGGCCTCTAGGAGACAGTGGG | [] |
| 2959 | 8 | SCRT1 | {'exs': 1, 'nonexs': 4} | False | 1.44E+08 | GCCGCCCTAGGCCTGCACAGG | [] |
| 2960 | 8 | FBXL6 | {'exs': 1, 'nonexs': 7} | True | 1.44E+08 | GGCTGCCTAGCTGGGTGAGGG | [] |
| 2961 | 8 | ADCK5 | {'exs': 2, 'nonexs': 6} | False | 1.44E+08 | CACCCTAGGTCTCCAGGTACTC | [] |
| 2962 | 8 | CPSF1 | {'exs': 6, 'nonexs': 2} | False | 1.44E+08 | ACGGGGCTAGAAGTGGGCGGT | [] |
| 2963 | 8 | CYHR1 | {'exs': 0, 'nonexs': 7} | True | 1.44E+08 | CCCGCCCTACTTCTGTATCTGC | [] |
| 2964 | 8 | KIFC2 | {'exs': 0, 'nonexs': 7} | True | 1.44E+08 | GACCCAGGACTAGAGGGGCAG | [] |
| 2965 | 8 | KIFC2 | {'exs': 0, 'nonexs': 7} | True | 1.44E+08 | ACTTTACTACCGCCTCTGCCAC | [] |
| 2966 | 8 | RECQL4 | {'exs': 0, 'nonexs': 1} | False | 1.45E+08 | CCCTAGCCTCTGACAACCCCA | [] |
| 2967 | 8 | LRRC24 | {'exs': 0, 'nonexs': 7} | True | 1.45E+08 | GCGGCCCTAGCAGTGGATCTC | [] |
| 2968 | 8 | ARHGAP39 | {'exs': 0, 'nonexs': 7} | True | 1.45E+08 | CCGCTACAGCACACCCTCCAT | [] |
| 2969 | 8 | RPL8 | {'exs': 6, 'nonexs': 2} | False | 1.45E+08 | AGCACTAGTTCTCTTTCTCCTG | [] |
| 2970 | 8 | ZNF7 | {'exs': 0, 'nonexs': 7} | True | 1.45E+08 | TGTAAGTGGTCTATCCCATGTG | [] |
| 2971 | 8 | ZNF16 | {'exs': 0, 'nonexs': 7} | True | 1.45E+08 | CAACAGCCTATTCCCTGGTGTC | [] |
| 2972 | 8 | C8orf33 | {'exs': 2, 'nonexs': 4} | False | 1.45E+08 | GGGGAGACGCTAAAAGAAATTG | [] |
| 2973 | 9 | WASHC1 | unavailable | False | 14806 | AGCCCCCTACGATTCCCAGTC | [] |
| 2974 | 9 | WASHC1 | unavailable | False | 16882 | CTCGGCTACCTCAGTGTGGAA | [] |
| 2975 | 9 | FOXD4 | {'exs': 0, 'nonexs': 7} | True | 116799 | CTGGCTCTAGGAGGGCCCTGC | [] |
| 2976 | 9 | CBWD1 | {'exs': 3, 'nonexs': 4} | False | 121561 | TCCTTATCTAAATTTCTGCCTG | [] |
| 2977 | 9 | DMRT2 | {'exs': 0, 'nonexs': 7} | True | 1055773 | TTTCTAAAGATGGTTCATTATG | [] |
| 2978 | 9 | KCNV2 | {'exs': 0, 'nonexs': 7} | True | 2729726 | AAATACTAATTCTCTTGTCTTG | [] |
| 2979 | 9 | PUM3 | unavailable | False | 2804330 | CTCTTTCCACCTATGTGCTCAG | [] |
| 2980 | 9 | GLIS3 | {'exs': 0, 'nonexs': 7} | True | 3855497 | AAATACTACCAGAGAAAGACAG | [] |
| 2981 | 9 | GLIS3 | {'exs': 0, 'nonexs': 7} | True | 4093248 | ATCACTATGCTATGCTGCCTGG | [] |
| 2982 | 9 | SLC1A1 | {'exs': 0, 'nonexs': 7} | True | 4585557 | GGCCCTAGAACTGTGAGGTCT | [] |
| 2983 | 9 | LOC10798 | unavailable | False | 5113849 | ACCTTCAGCATCTACTCCAGGC | [] |
| 2984 | 9 | INSL4 | {'exs': 0, 'nonexs': 7} | True | 5233876 | TACTATGTACATAATTTAACTG | [] |
| 2985 | 9 | CD274 | {'exs': 0, 'nonexs': 7} | True | 5463176 | CATGCTAGGTGCTAGGGGACA | [] |
| 2986 | 9 | RIC1 | unavailable | False | 5769329 | TATCAACTAATAGCAAAATGTG | [] |
| 2987 | 9 | IL33 | {'exs': 0, 'nonexs': 7} | True | 6256167 | CCATCAACTAAGTTTCAGAGAG | [] |
| 2988 | 9 | GLDC | {'exs': 0, 'nonexs': 7} | True | 6533016 | ACTAAGAAGACGCCCTCTTTTG | [] |
| 2989 | 9 | KDM4C | {'exs': 1, 'nonexs': 7} | True | 7076450 | GTTGCTGTGTCTAGATTCCCAG | [] |
| 2990 | 9 | KDM4C | {'exs': 1, 'nonexs': 7} | True | 7174728 | TGCAGACTACTGTCTCTTCTGG | [] |
| 2991 | 9 | PTPRD | {'exs': 1, 'nonexs': 4} | False | 8317873 | GGGTTTCTACGTTGCATAGTGG | [] |
| 2992 | 9 | LURAP1L | unavailable | False | 12777228 | ATGTACCTAAGGGTTGTCCTTG | ['AAAAAATAACAATCACTAAATG', 'CCTTAAATTTTTGTCCAACCTG'] |
| 2993 | 9 | LURAP1L | unavailable | False | 12821759 | CTGTCACTAGCCAAAGCAGTAG | [] |
| 2994 | 9 | NFIB | {'exs': 0, 'nonexs': 7} | True | 14088308 | ACCAAGCTAGCCCAGGTACCA | [] |
| 2995 | 9 | NFIB | {'exs': 0, 'nonexs': 7} | True | 14305977 | CAAGGACATCTATATCTTGATG | [] |
| 2996 | 9 | FREM1 | {'exs': 1, 'nonexs': 7} | True | 14791252 | TTGCTATTCTAGGCCTAAAGTG | [] |
| 2997 | 9 | PSIP1 | {'exs': 1, 'nonexs': 6} | True | 15465519 | TCAACCTAGTTATCTAGTGTAG | [] |
| 2998 | 9 | BNC2 | {'exs': 0, 'nonexs': 7} | True | 16418988 | CTGAGACTAATCTACTGAAGTG | [] |
| 2999 | 9 | CNTLN | {'exs': 0, 'nonexs': 7} | True | 17425416 | GTCAGCTTCTAGCAGCAGATCC | [] |
| 3000 | 9 | CNTLN | {'exs': 0, 'nonexs': 7} | True | 17509828 | GGAACTAAGACCTCTAGGATAG | [] |
| 3001 | 9 | SH3GL2 | {'exs': 0, 'nonexs': 7} | True | 17795742 | ATAACATCCTAATGGGGCAGGG | [] |
| 3002 | 9 | SLC24A2 | {'exs': 0, 'nonexs': 7} | True | 19516152 | TTCCTGCTAGATGGAGACGGG | [] |
| 3003 | 9 | KLHL9 | {'exs': 0, 'nonexs': 7} | True | 21333005 | TAGACCTAAGAATGATCTGAAG | [] |
| 3004 | 9 | IFNE | {'exs': 0, 'nonexs': 7} | True | 21481067 | CTCCACCTACCTCGGGCTTCT | ['GGCTTCTAAACTCTGTAGTAAG'] |
| 3005 | 9 | CDKN2A | {'exs': 1, 'nonexs': 6} | True | 21968724 | GTAGCTTCCCTACGCATGCCT | [] |
| 3006 | 9 | CDKN2A | {'exs': 1, 'nonexs': 6} | True | 21969731 | CGGATGCTAGCTGTAACTGGA | [] |
| 3007 | 9 | CDKN2B | {'exs': 1, 'nonexs': 6} | True | 22008716 | GGCGATCTAGGTTCCAGCCCC | [] |
| 3008 | 9 | TUSC1 | {'exs': 0, 'nonexs': 7} | True | 25677682 | CTCGGCTACAGCCAGGGCCCA | [] |
| 3009 | 9 | CAAP1 | unavailable | False | 26842300 | AATACCTAGGCTGGCTTTTTTG | ['TTATATCACCAGCTTTCATTAG'] |
| 3010 | 9 | PLAA | {'exs': 2, 'nonexs': 6} | False | 26905510 | CCACTGCTACAGCAAATTTAGG | [] |
| 3011 | 9 | TEK | {'exs': 1, 'nonexs': 6} | True | 27229231 | CTGTCCTAGGCCGCTTCTTCA | [] |
| 3012 | 9 | ACO1 | {'exs': 1, 'nonexs': 7} | True | 32450110 | CGTCTCCTACTTGGCCATCTTC | [] |
| 3013 | 9 | APTX | {'exs': 0, 'nonexs': 7} | True | 32926415 | CAGCTGCCTAAAACTTCGGATG | [] |
| 3014 | 9 | APTX | {'exs': 0, 'nonexs': 7} | True | 32973683 | GCTACTCTGGAAAACCAATATG | [] |
| 3015 | 9 | B4GALT1 | {'exs': 0, 'nonexs': 7} | True | 33113453 | AAAACGCTAGCTCGGTGTCCC | [] |
| 3016 | 9 | CHMP5 | {'exs': 2, 'nonexs': 5} | False | 33280858 | AAATCTATGAAGCAGGGATCTG | [] |
| 3017 | 9 | NFX1 | {'exs': 1, 'nonexs': 7} | True | 33347739 | TGCTGCTATAAACATGCGTGTG | [] |
| 3018 | 9 | KIF24 | {'exs': 0, 'nonexs': 5} | True | 34254379 | GTCTGGCTCTAAGACGGCACTC | [] |
| 3019 | 9 | C9orf24 | {'exs': 0, 'nonexs': 7} | True | 34379075 | CAGGCTCTACCAGCAACGATA | [] |
| 3020 | 9 | C9orf24 | {'exs': 0, 'nonexs': 7} | True | 34379645 | ACTTACCTAAGGTTCGGATATG | [] |
| 3021 | 9 | FAM219A | unavailable | False | 34400963 | CGCCCGCTACTGAATGTGGCA | [] |
| 3022 | 9 | SIGMAR1 | {'exs': 0, 'nonexs': 7} | True | 34635846 | CGTGTACTACCGTCTCCCCTG | [] |
| 3023 | 9 | IL11RA | {'exs': 0, 'nonexs': 7} | True | 34661497 | TGGGTCCTCTACAGGTTTGGA | [] |
| 3024 | 9 | CCL21 | {'exs': 0, 'nonexs': 7} | True | 34709393 | ACTGGGCTATGGCCCTTTAGG | [] |
| 3025 | 9 | FAM205A | unavailable | False | 34723231 | AGGAGCTAGGTTGGGTGCCCT | [] |
| 3026 | 9 | PHF24 | unavailable | False | 34978110 | CATGCTATCCTGGGGGCTTCA | [] |
| 3027 | 9 | DNAJB5 | {'exs': 0, 'nonexs': 7} | True | 34997258 | CAGAGCCTAGGAACAGGGTAG | [] |
| 3028 | 9 | C9orf131 | {'exs': 0, 'nonexs': 7} | True | 35045868 | TTTTCTCTACTGACTAGATTTG | [] |
| 3029 | 9 | FANCG | {'exs': 1, 'nonexs': 6} | True | 35074107 | CGTGGCAGCTACAGGTCACAA | [] |

| | | | | | | | |
|---|---|---|---|---|---|---|---|
| 3030 | 9 | PIGO | {'exs': 0, 'nonexs': 7} | True | 35089091 | TAGGCTACCTCTGCTGGGCCA | [] |
| 3031 | 9 | STOML2 | {'exs': 1, 'nonexs': 6} | True | 35100034 | TCCACTAACTCATCTTGACTCG | [] |
| 3032 | 9 | FAM166B | {'exs': 0, 'nonexs': 7} | True | 35562006 | CAGTGGCCTAAGCCAAGAGCT | [] |
| 3033 | 9 | TESK1 | {'exs': 1, 'nonexs': 6} | True | 35609741 | CACTGCTAAGAGCGTGCCCCA | [] |
| 3034 | 9 | CD72 | {'exs': 0, 'nonexs': 5} | True | 35610623 | ACTGTCCTAATCTGGAAACCTG | [] |
| 3035 | 9 | ARHGEF39 | unavailable | False | 35661986 | ATTCCTCTAGTTTTTCTGGCTG | [] |
| 3036 | 9 | CA9 | {'exs': 0, 'nonexs': 7} | True | 35681024 | CAGCCTCTAGGCTCCAGTCTC | [] |
| 3037 | 9 | TPM2 | {'exs': 1, 'nonexs': 6} | True | 35684790 | CTCTAGGTCCCCACATTTACTG | [] |
| 3038 | 9 | CREB3 | {'exs': 2, 'nonexs': 6} | False | 35730725 | TCATATCTAGCCTGAGTATCTG | [] |
| 3039 | 9 | GBA2 | {'exs': 0, 'nonexs': 6} | True | 35737475 | CATACTAACTTGGCACCCTCTG | [] |
| 3040 | 9 | FAM221B | unavailable | False | 35818845 | TCTGACTACAGACCCTGGAATG | [] |
| 3041 | 9 | RECK | {'exs': 0, 'nonexs': 7} | True | 36085926 | GTATAGACTAAATGTCAGAGAG | [] |
| 3042 | 9 | GNE | {'exs': 1, 'nonexs': 6} | True | 36217364 | GAGGTCTAGTAGATCCTGCGT | [] |
| 3043 | 9 | GRHPR | {'exs': 1, 'nonexs': 6} | True | 37430781 | GGCTACAGCTTGAGTTCACTAG | [] |
| 3044 | 9 | ZBTB5 | {'exs': 1, 'nonexs': 7} | True | 37440517 | AAACAGCTAGAGCAAAGTGCTG | [] |
| 3045 | 9 | POLR1E | {'exs': 7, 'nonexs': 1} | False | 37503201 | GCGTCTAGGTAATCTTCCTCC | [] |
| 3046 | 9 | TRMT10B | unavailable | False | 37763857 | GTTTACTAAGGAATCCCAGAAG | [] |
| 3047 | 9 | DCAF10 | {'exs': 0, 'nonexs': 7} | True | 37861507 | GTTGTGCCTAAAACTTTGGCTG | [] |
| 3048 | 9 | CNTNAP3 | {'exs': 0, 'nonexs': 7} | True | 39073889 | TCCTAGCACTCTTCTTTTTTTG | [] |
| 3049 | 9 | CBWD6 | {'exs': 0, 'nonexs': 3} | True | 41131828 | TCCTTATCTAAATTTCTGCCTG | [] |
| 3050 | 9 | CBWD6 | {'exs': 0, 'nonexs': 3} | True | 41143089 | AAAACCTAGGAAAGATTTTCTG | [] |
| 3051 | 9 | CNTNAP3 | {'exs': 0, 'nonexs': 7} | True | 41893988 | TCCTAGCACTCTTCTTTTTTTG | [] |
| 3052 | 9 | CBWD5 | {'exs': 4, 'nonexs': 3} | False | 65733520 | TCCTTATCTAAATTTCTGCCTG | [] |
| 3053 | 9 | CBWD3 | {'exs': 4, 'nonexs': 3} | False | 68299491 | TCCTTATCTAAATTTCTGCCTG | [] |
| 3054 | 9 | PGM5 | {'exs': 0, 'nonexs': 7} | True | 68405367 | AATTGGGTCCTAATTACTGCAG | [] |
| 3055 | 9 | PRKACG | {'exs': 1, 'nonexs': 6} | True | 69013036 | CAAGCACACCCCTAAAACTCAG | [] |
| 3056 | 9 | TJP2 | {'exs': 0, 'nonexs': 6} | True | 69254373 | ACATCTATAATTCTGTGTCCCG | [] |
| 3057 | 9 | PTAR1 | {'exs': 3, 'nonexs': 4} | False | 69718645 | GAAACCTACCATTTAAGTGATG | [] |
| 3058 | 9 | C9orf135 | {'exs': 1, 'nonexs': 7} | True | 69905983 | GGACTATGGAATAATCATCTTG | [] |
| 3059 | 9 | MAMDC2 | {'exs': 1, 'nonexs': 7} | True | 70216926 | GATGGGACTATTTAAACTACAG | [] |
| 3060 | 9 | TRPM3 | {'exs': 0, 'nonexs': 7} | True | 70784100 | GTTACCTACCCGCCTTCTTCT | [] |
| 3061 | 9 | CEMIP2 | unavailable | False | 71685196 | AGTCTCTAATGTGCTTTTGAAG | [] |
| 3062 | 9 | PRUNE2 | {'exs': 0, 'nonexs': 7} | True | 76614569 | CCAACTAAGGCTTTTCTTTCAG | [] |
| 3063 | 9 | FOXB2 | {'exs': 0, 'nonexs': 7} | True | 77020952 | GGGTCCCCTAGGAGTGCACTA | [] |
| 3064 | 9 | IDNK | unavailable | False | 83641048 | CGTGCTACAGTGTTTCACAAAG | [] |
| 3065 | 9 | IDNK | unavailable | False | 83643182 | TACCTAGGGTACTTGCCCATAG | [] |
| 3066 | 9 | UBQLN1 | {'exs': 0, 'nonexs': 7} | True | 83661786 | ATGCTGCTATGATGGCTGGGA | [] |
| 3067 | 9 | UBQLN1 | {'exs': 0, 'nonexs': 7} | True | 83669529 | CTCAAAATCTCTAAGGATGAGG | [] |
| 3068 | 9 | KIF27 | {'exs': 0, 'nonexs': 7} | True | 83837000 | ATTCAATGTCTAAGTTTTTAAG | [] |
| 3069 | 9 | NTRK2 | {'exs': 1, 'nonexs': 6} | True | 84810578 | TTTCAGCTACCCATCCAGTGG | [] |
| 3070 | 9 | NTRK2 | {'exs': 1, 'nonexs': 6} | True | 85021436 | GGCCCTAGCCTAGAATGTCCA | [] |
| 3071 | 9 | TUT7 | unavailable | False | 86297157 | ATCTATATTTCTCTCTCAACTG | [] |
| 3072 | 9 | GAS1 | {'exs': 2, 'nonexs': 6} | False | 86945741 | AGGGCTAAAAGAGCGGCCCAA | [] |
| 3073 | 9 | DAPK1 | {'exs': 1, 'nonexs': 6} | True | 87686949 | CTAGGCGCTGACAATACCCCA | [] |
| 3074 | 9 | SPATA31C | unavailable | False | 88129631 | AGGGCTACTGATCTCTGATTTG | [] |
| 3075 | 9 | SPIN1 | {'exs': 1, 'nonexs': 6} | True | 88475276 | CTAGGATGTTTTCACCAAATCG | [] |
| 3076 | 9 | SEMA4D | {'exs': 0, 'nonexs': 7} | True | 89362401 | TGGTGGCTACAGGGTGTCCTT | [] |
| 3077 | 9 | SPTLC1 | {'exs': 2, 'nonexs': 5} | False | 92032464 | CTGCCTAGAGCAGGACGGCCT | [] |
| 3078 | 9 | IARS1 | unavailable | False | 92210806 | ACATGCTAGAAGTCTGCTGTTG | [] |
| 3079 | 9 | CENPP | {'exs': 1, 'nonexs': 7} | True | 92613148 | TGGAACTAGTTGTTCTCCTCTG | [] |
| 3080 | 9 | OMD | {'exs': 0, 'nonexs': 7} | True | 92415151 | TTCTTGCTATTCTTGATTTTCG | ['TTCATAATAATGAAGGTCAAAG'] |
| 3081 | 9 | ECM2 | {'exs': 0, 'nonexs': 7} | True | 92494082 | TTGCCTCTACCAGAGAGGAAAG | [] |
| 3082 | 9 | BICD2 | {'exs': 2, 'nonexs': 6} | False | 92713455 | GCTACTCTACAGCTGAAAACGG | [] |
| 3083 | 9 | BICD2 | {'exs': 2, 'nonexs': 6} | False | 92715153 | CAGCCCCTAATCACAGTAAATG | [] |
| 3084 | 9 | ZNF484 | {'exs': 0, 'nonexs': 7} | True | 92840227 | ATTCTAACTAGATAGAAGAAAG | [] |
| 3085 | 9 | NINJ1 | {'exs': 0, 'nonexs': 7} | True | 93124907 | TGGGTGTCCTACTGCTGGGGT | [] |
| 3086 | 9 | WNK2 | {'exs': 0, 'nonexs': 3} | True | 93318026 | AGGAACCTAGGAGCGAACAGG | [] |
| 3087 | 9 | PTPDC1 | {'exs': 0, 'nonexs': 7} | True | 94107943 | GAAAGCTAGAGGCCAGGCTTA | [] |
| 3088 | 9 | LOC10798 | unavailable | False | 94154220 | TACACTACCTTTGTTGGCCTAG | [] |
| 3089 | 9 | NUTM2F | unavailable | False | 94318454 | GCGGCCCCTACTGGCTACAAT | [] |
| 3090 | 9 | FBP2 | {'exs': 0, 'nonexs': 7} | True | 94558937 | AAACTCGCTAGCTGCCTGCCT | [] |
| 3091 | 9 | AOPEP | unavailable | False | 95215885 | CTGTCTCTACTAAAAATAGAAG | [] |
| 3092 | 9 | FANCC | {'exs': 5, 'nonexs': 4} | False | 95301706 | GTGCCTTCTAGACTTGAGTTCG | [] |
| 3093 | 9 | HSD17B3 | {'exs': 0, 'nonexs': 7} | True | 96235459 | GCCTGGCTACCTGACCTTGGT | [] |
| 3094 | 9 | ZNF367 | {'exs': 0, 'nonexs': 7} | True | 96388236 | CCAAGCTACTGTCGGAGTGGC | [] |
| 3095 | 9 | CDC14B | {'exs': 1, 'nonexs': 6} | True | 96496286 | GGTCTAGTCCTTGTGGATCCC | [] |
| 3096 | 9 | CDC14B | {'exs': 1, 'nonexs': 6} | True | 96515677 | TCCTGATGGCTACTGTGTTCTG | [] |
| 3097 | 9 | NUTM2G | unavailable | False | 96939148 | GCGGCCCCTACTGGCTACAAT | [] |
| 3098 | 9 | MFSD14C | unavailable | False | 96946343 | AGGCTTGAGCTACCGCATCTG | [] |
| 3099 | 9 | TMOD1 | {'exs': 0, 'nonexs': 7} | True | 97599697 | CACACACTAGACACCACTCCG | [] |
| 3100 | 9 | TRIM14 | {'exs': 0, 'nonexs': 7} | True | 98087469 | TGGCCCCTAGGGCAGCCGGGG | [] |
| 3101 | 9 | TRIM14 | {'exs': 0, 'nonexs': 7} | True | 98090381 | AACTAACCTATAGTCCTAGAAG | [] |
| 3102 | 9 | ANKS6 | {'exs': 0, 'nonexs': 7} | True | 98732503 | CGGCTGCTACCTCTTGCCGGA | [] |
| 3103 | 9 | SEC61B | {'exs': 3, 'nonexs': 5} | False | 99230423 | AATCTACGAACGAGTGTACTTG | [] |
| 3104 | 9 | MSANTD3 | unavailable | False | 1E+08 | TTTCCTACCAAGTCCCAGAAAG | [] |
| 3105 | 9 | ALDOB | {'exs': 0, 'nonexs': 7} | True | 1.01E+08 | TGGACCCTAGTAGGTATAGCAG | [] |
| 3106 | 9 | PGAP4 | unavailable | False | 1.01E+08 | TGGCACCCTAGAGGAGACTGG | [] |
| 3107 | 9 | CYLC2 | {'exs': 1, 'nonexs': 7} | True | 1.03E+08 | GCCTACTTGCCCTTCTTCTTTG | [] |
| 3108 | 9 | SMC2 | {'exs': 8, 'nonexs': 1} | False | 1.04E+08 | CAAAATCTATTTTCCTTCCTTG | [] |
| 3109 | 9 | NIPSNAP3 | {'exs': 0, 'nonexs': 7} | True | 1.05E+08 | GTAGAAAACTATTTCAGTGGTG | [] |
| 3110 | 9 | NIPSNAP3 | {'exs': 0, 'nonexs': 7} | True | 1.05E+08 | GTAGAAAACTATTTCAATGGTG | [] |
| 3111 | 9 | FSD1L | {'exs': 0, 'nonexs': 7} | True | 1.06E+08 | ACACTATTGAGTAACAACATTG | [] |
| 3112 | 9 | FKTN | {'exs': 0, 'nonexs': 7} | True | 1.06E+08 | CATGTCTACAAACTCAGTCCAG | [] |
| 3113 | 9 | TAL2 | {'exs': 0, 'nonexs': 7} | True | 1.06E+08 | GCCACACTAAGGAATGTGGTG | [] |
| 3114 | 9 | TMEM38B | {'exs': 0, 'nonexs': 7} | True | 1.06E+08 | ACTACTGAAACAAAAATAAATG | [] |
| 3115 | 9 | TXNDC8 | {'exs': 0, 'nonexs': 7} | True | 1.1E+08 | TCCTACTCATTTCCATCATCTG | [] |
| 3116 | 9 | TXNDC8 | {'exs': 0, 'nonexs': 7} | True | 1.1E+08 | AAAGCTCTATTTTCCTGTATTG | [] |
| 3117 | 9 | LPAR1 | {'exs': 0, 'nonexs': 6} | True | 1.11E+08 | TCCGTTCTAAACCACAGAGTGG | [] |
| 3118 | 9 | SHOC1 | unavailable | False | 1.12E+08 | CTAGTGAGTGAAATATTCACAG | [] |
| 3119 | 9 | SUSD1 | {'exs': 0, 'nonexs': 7} | True | 1.12E+08 | TTTCTATCCGTCCACTACCCAG | [] |
| 3120 | 9 | HSDL2 | {'exs': 1, 'nonexs': 7} | True | 1.12E+08 | ACTCACCTCTATTTTTCATTCG | ['TTCATTCATCTTTTTTTTTAAG'] |
| 3121 | 9 | C9orf147 | {'exs': 0, 'nonexs': 1} | False | 1.12E+08 | AATAGGTCTAATTGGTGGCAAG | [] |
| 3122 | 9 | SNX30 | {'exs': 0, 'nonexs': 7} | True | 1.13E+08 | ACTACACTAGGAAGTTTCTTCG | ['TCTTCATCTCCCCAGATTGGTG'] |
| 3123 | 9 | SLC31A2 | {'exs': 1, 'nonexs': 7} | True | 1.13E+08 | TCACCAGCTAAGCTGTGCTGA | [] |
| 3124 | 9 | CDC26 | {'exs': 4, 'nonexs': 3} | False | 1.13E+08 | TAATCCATCTCTAAAATTCAAG | [] |
| 3125 | 9 | PRPF4 | {'exs': 7, 'nonexs': 1} | False | 1.13E+08 | TCATCTATTCAGCCATCCACAG | [] |
| 3126 | 9 | C9orf43 | {'exs': 0, 'nonexs': 7} | True | 1.13E+08 | CTTCAGTCTATTCCTCCAGAGG | [] |
| 3127 | 9 | RGS3 | {'exs': 0, 'nonexs': 7} | True | 1.14E+08 | CCAGTGGCCCCTAAAGCGGGG | [] |
| 3128 | 9 | KIF12 | {'exs': 0, 'nonexs': 7} | True | 1.14E+08 | CTGAGACTAGAAGCCTCCTGA | [] |
| 3129 | 9 | KIF12 | {'exs': 0, 'nonexs': 7} | True | 1.14E+08 | TGGTCTATTTCTGACTCCTCAG | [] |
| 3130 | 9 | KIF12 | {'exs': 0, 'nonexs': 7} | True | 1.14E+08 | CATTCTCTAGCATGAACTCCTG | [] |
| 3131 | 9 | ORM1 | {'exs': 0, 'nonexs': 7} | True | 1.14E+08 | GCTAGGATTCCCCCTCCTCCT | [] |
| 3132 | 9 | ORM2 | {'exs': 1, 'nonexs': 6} | True | 1.14E+08 | GCTAGGATTCCCCCTCCTCCT | [] |
| 3133 | 9 | WHRN | unavailable | False | 1.14E+08 | TGGCCTCTAGAGCATCACATTG | [] |
| 3134 | 9 | WHRN | unavailable | False | 1.14E+08 | TCAGGTCTATAAGCCACAGCAG | [] |
| 3135 | 9 | ATP6V1G1 | {'exs': 6, 'nonexs': 2} | False | 1.15E+08 | TCTCTTCTATCCATTTATGCGG | [] |
| 3136 | 9 | TNFSF15 | {'exs': 1, 'nonexs': 6} | True | 1.15E+08 | CTCTCCTCCTATAGTAAGAAGG | [] |
| 3137 | 9 | TRIM32 | {'exs': 0, 'nonexs': 5} | True | 1.17E+08 | TTTCTCATCCCCTATGGGGTG | [] |
| 3138 | 9 | PHF19 | {'exs': 0, 'nonexs': 7} | True | 1.21E+08 | GAGTCTACAAATCCTGGCCAAG | [] |
| 3139 | 9 | TRAF1 | {'exs': 0, 'nonexs': 7} | True | 1.21E+08 | CCCCGCCCACCCTAAGTGCTG | [] |
| 3140 | 9 | CNTRL | unavailable | False | 1.21E+08 | TTTTTGTCTAGAAAAGAAACTG | [] |
| 3141 | 9 | CNTRL | unavailable | False | 1.21E+08 | AGGTGTCTATTGCCATACCTGG | [] |
| 3142 | 9 | RAB14 | {'exs': 1, 'nonexs': 7} | True | 1.21E+08 | AAGAGGTCACTAGCAGCCACA | [] |
| 3143 | 9 | STOM | {'exs': 0, 'nonexs': 7} | True | 1.21E+08 | TCTACACTAGCCTAGATGGCTG | [] |
| 3144 | 9 | DAB2IP | {'exs': 1, 'nonexs': 7} | True | 1.22E+08 | TTTGTTTCTAATGCATACTGTG | [] |
| 3145 | 9 | MORN5 | {'exs': 0, 'nonexs': 6} | True | 1.22E+08 | CTCATCCTAGCCCTTTCGACA | [] |
| 3146 | 9 | LHX6 | {'exs': 0, 'nonexs': 7} | True | 1.22E+08 | ATCTCTTCTAGTGGCAGCCTG | [] |
| 3147 | 9 | OR1N1 | {'exs': 0, 'nonexs': 3} | True | 1.23E+08 | CTAAGAGGAAACAATACTCCTG | [] |
| 3148 | 9 | OR1L8 | {'exs': 1, 'nonexs': 3} | False | 1.23E+08 | GGGTGCTTCCTAGGATCTCTT | [] |
| 3149 | 9 | OR1L4 | {'exs': 0, 'nonexs': 3} | True | 1.23E+08 | TTCTTTCTATGAGTAAATTCTG | [] |
| 3150 | 9 | STRBP | {'exs': 0, 'nonexs': 5} | True | 1.23E+08 | CATTTGCTAAAAGAATGAGTAG | [] |
| 3151 | 9 | CRB2 | {'exs': 0, 'nonexs': 7} | True | 1.23E+08 | GCCTAGATGAGTCTCTCCTCC | [] |
| 3152 | 9 | LHX2 | {'exs': 0, 'nonexs': 7} | True | 1.24E+08 | GATAAATCTAGAGAGATGCCTG | [] |
| 3153 | 9 | GOLGA1 | {'exs': 0, 'nonexs': 7} | True | 1.25E+08 | CCCTCTAGGACCATGGTATCC | [] |
| 3154 | 9 | HSPA5 | {'exs': 7, 'nonexs': 1} | False | 1.25E+08 | GTCTACAACTCATCTTTTTCTG | [] |
| 3155 | 9 | GAPVD1 | {'exs': 0, 'nonexs': 7} | True | 1.25E+08 | ACAAAGGCTAATGAAATATTAG | [] |
| 3156 | 9 | MVB12B | unavailable | False | 1.26E+08 | CATCCTACTTGATCAAAAAACG | [] |
| 3157 | 9 | RALGPS1 | {'exs': 0, 'nonexs': 7} | True | 1.27E+08 | TGACTAAACGCTCCCTTCCCT | [] |
| 3158 | 9 | RALGPS1 | {'exs': 0, 'nonexs': 7} | True | 1.27E+08 | CCTGTTTTCTAAAGACCCTTAG | [] |
| 3159 | 9 | ANGPTL2 | {'exs': 0, 'nonexs': 7} | True | 1.27E+08 | GATGATTCCTAGAGTCCTAGAG | [] |
| 3160 | 9 | NIBAN2 | unavailable | False | 1.28E+08 | CACTGGCCTAGAACTCAGTCT | [] |
| 3161 | 9 | CFAP157 | unavailable | False | 1.28E+08 | TGTGTCCTACTTGGGACGTAG | [] |
| 3162 | 9 | CFAP157 | unavailable | False | 1.28E+08 | CTGCACTAGGGCGCCCTGCCT | [] |
| 3163 | 9 | TTC16 | {'exs': 0, 'nonexs': 7} | True | 1.28E+08 | CTGCTATGAGCTGGGCCATCT | [] |
| 3164 | 9 | FPGS | {'exs': 7, 'nonexs': 2} | False | 1.28E+08 | CCTTGGCTACTGGGACAGTGC | [] |
| 3165 | 9 | ENG | {'exs': 2, 'nonexs': 6} | False | 1.28E+08 | CCGGGGCTATGCCATGCTGCT | [] |
| 3166 | 9 | AK1 | {'exs': 1, 'nonexs': 6} | True | 1.28E+08 | GTTGCTACTTTAGGGCGTCCA | [] |
| 3167 | 9 | ST6GALNA | {'exs': 0, 'nonexs': 7} | True | 1.28E+08 | GGTGGCCTAGGTCCAGGAGGG | [] |
| 3168 | 9 | ST6GALNA | {'exs': 0, 'nonexs': 7} | True | 1.28E+08 | CGGAAGCTACTCAGTCCTCCA | [] |
| 3169 | 9 | C9orf16 | {'exs': 0, 'nonexs': 7} | True | 1.28E+08 | AGCCTAGGGGCTGGCATCACT | [] |
| 3170 | 9 | CERCAM | {'exs': 0, 'nonexs': 1} | False | 1.28E+08 | CCTCACCTAGAGCTCATCTCG | [] |
| 3171 | 9 | SET | {'exs': 0, 'nonexs': 7} | True | 1.29E+08 | CCACACTATATATATATATATG | ['ATATACACCCCAAAATAACATG'] |
| 3172 | 9 | ZDHHC12 | {'exs': 1, 'nonexs': 7} | True | 1.29E+08 | GCAACCCTAAACAGCTGGGCT | [] |
| 3173 | 9 | ZER1 | {'exs': 1, 'nonexs': 7} | True | 1.29E+08 | CCTCTATCTAGACGTGTCCATG | [] |
| 3174 | 9 | TBC1D13 | {'exs': 0, 'nonexs': 7} | True | 1.29E+08 | GCCGGGCTACTTTGAGTCTTG | [] |
| 3175 | 9 | KYAT1 | unavailable | False | 1.29E+08 | AGGGCTAGAGTTCCACCTTCC | ['CTTCCACTTCCGCAGCTTCTCG'] |
| 3176 | 9 | DOLK | {'exs': 5, 'nonexs': 3} | False | 1.29E+08 | ACTGTAACAGCTAGGCCATCAG | [] |
| 3177 | 9 | NUP188 | {'exs': 7, 'nonexs': 1} | False | 1.29E+08 | CACTGCCCTATCTTTGCATATG | [] |
| 3178 | 9 | SH3GLB2 | {'exs': 0, 'nonexs': 7} | True | 1.29E+08 | CTCACAGCTAGGTGACTAGGG | [] |
| 3179 | 9 | SH3GLB2 | {'exs': 0, 'nonexs': 7} | True | 1.29E+08 | TGCCTAGCTGAGCAGTTCCAA | [] |
| 3180 | 9 | MIGA2 | unavailable | False | 1.29E+08 | GCCTCTACTGCAGCTCCCCCA | [] |
| 3181 | 9 | PTPA | unavailable | False | 1.29E+08 | CCCTCCTAGCCCGACGTGACA | [] |
| 3182 | 9 | IER5L | {'exs': 0, 'nonexs': 7} | True | 1.29E+08 | GGGTCCCTAGAAGGCGACAAT | [] |
| 3183 | 9 | FNBP1 | {'exs': 0, 'nonexs': 7} | True | 1.3E+08 | TCTAGGAATCTACAACACAAAG | [] |
| 3184 | 9 | NCS1 | {'exs': 0, 'nonexs': 7} | True | 1.3E+08 | CTGGGACTATACCAGCCCGTC | [] |
| 3185 | 9 | ASS1 | {'exs': 0, 'nonexs': 7} | True | 1.31E+08 | GGTCTATTTGGCAGTGACCTTG | [] |
| 3186 | 9 | FUBP3 | {'exs': 1, 'nonexs': 7} | True | 1.31E+08 | GCTGTCCTACTGCTCCTGTGG | [] |
| 3187 | 9 | ABL1 | {'exs': 3, 'nonexs': 4} | False | 1.31E+08 | CTGCTGCTACCTCTGCACTAT | [] |
| 3188 | 9 | FIBCD1 | {'exs': 1, 'nonexs': 6} | True | 1.31E+08 | ACCAGTCTAGCGGTCCTCCCG | [] |
| 3189 | 9 | PLPP7 | unavailable | False | 1.31E+08 | GGTGCTAATCTGCAAGACATTG | [] |
| 3190 | 9 | POMT1 | {'exs': 0, 'nonexs': 7} | True | 1.32E+08 | GTTCTAGTGTTTTCGGATCAAG | [] |
| 3191 | 9 | RAPGEF1 | {'exs': 3, 'nonexs': 6} | False | 1.32E+08 | GCTCCTAGGTCTTCTCTTCCC | [] |
| 3192 | 9 | RAPGEF1 | {'exs': 3, 'nonexs': 6} | False | 1.32E+08 | GCTCTGCTACTGACTGGCTGT | [] |
| 3193 | 9 | MED27 | {'exs': 7, 'nonexs': 1} | False | 1.32E+08 | TGGGGGCTACTGCCGGCAGGT | [] |
| 3194 | 9 | NTNG2 | {'exs': 0, 'nonexs': 7} | True | 1.32E+08 | CCTAGAATTGTACCCCCAAAAG | [] |
| 3195 | 9 | NTNG2 | {'exs': 0, 'nonexs': 7} | True | 1.32E+08 | CTCCTAACAGGTGGGGCACCT | [] |
| 3196 | 9 | SETX | {'exs': 0, 'nonexs': 7} | True | 1.32E+08 | CTATAAAAGCTTTCTTTTCTTG | [] |
| 3197 | 9 | CFAP77 | unavailable | False | 1.33E+08 | AGCGTGCTAAATCCAGGTAGTG | [] |
| 3198 | 9 | CFAP77 | unavailable | False | 1.33E+08 | GAGGGGCTAGGGGTGGGTGTA | [] |
| 3199 | 9 | DDX31 | {'exs': 1, 'nonexs': 7} | True | 1.33E+08 | TGTTTTCTATTGCAACATACAG | [] |
| 3200 | 9 | DDX31 | {'exs': 1, 'nonexs': 7} | True | 1.33E+08 | AAGGCACTAAGATTCCTGGGG | [] |
| 3201 | 9 | AK8 | unavailable | False | 1.33E+08 | GATGCGCTAAGATGGGGCCCT | [] |
| 3202 | 9 | SPACA9 | unavailable | False | 1.33E+08 | AGAAGCTAAGTCGTCCCCTGT | [] |
| 3203 | 9 | LOC10099 | unavailable | False | 1.33E+08 | ACCTAAAGCTGCTCTAAAAATG | [] |
| 3204 | 9 | CEL | {'exs': 0, 'nonexs': 7} | True | 1.33E+08 | GGACGCTAAAACCTAATGACTG | [] |
| 3205 | 9 | OBP2B | {'exs': 0, 'nonexs': 7} | True | 1.33E+08 | CTCACCCTAGTGTTCGGGAAC | [] |
| 3206 | 9 | MED22 | {'exs': 0, 'nonexs': 1} | False | 1.33E+08 | CAGCGCTATTTATACCTGGACG | [] |
| 3207 | 9 | STKLD1 | unavailable | False | 1.33E+08 | AACATCTATTCCAGTCCCCCAG | [] |
| 3208 | 9 | REXO4 | {'exs': 0, 'nonexs': 7} | True | 1.33E+08 | GGACTGCTAGGCGTCGTCACT | [] |
| 3209 | 9 | SLC2A6 | {'exs': 0, 'nonexs': 7} | True | 1.33E+08 | ACCTTGACCTAGCGCAAGAAG | [] |
| 3210 | 9 | ADAMTSL | {'exs': 0, 'nonexs': 7} | True | 1.34E+08 | GCCGGGCCTAGGAGTGGGGG | [] |
| 3211 | 9 | RXRA | {'exs': 0, 'nonexs': 7} | True | 1.34E+08 | CAGGCCTAAGTCATTTGGTGCC | [] |
| 3212 | 9 | COL5A1 | {'exs': 1, 'nonexs': 6} | True | 1.35E+08 | GGCTCCTAGCCCATGAAGCAA | [] |
| 3213 | 9 | FCN2 | {'exs': 0, 'nonexs': 7} | True | 1.35E+08 | CCGGCCTGGGCTAGGCAGGTC | [] |
| 3214 | 9 | FCN1 | {'exs': 0, 'nonexs': 7} | True | 1.35E+08 | CCTGGCCCGTCTAGGCGGGC | [] |
| 3215 | 9 | OLFM1 | {'exs': 0, 'nonexs': 7} | True | 1.35E+08 | AGGGAGCTACAACTCGTCGGA | [] |
| 3216 | 9 | LCN1 | {'exs': 0, 'nonexs': 7} | True | 1.36E+08 | TCACCCTAATCGCTCCCTGGA | [] |
| 3217 | 9 | OBP2A | {'exs': 1, 'nonexs': 7} | True | 1.36E+08 | CTCACCCTAGTGTTCGAGAAC | [] |
| 3218 | 9 | PAEP | {'exs': 0, 'nonexs': 7} | True | 1.36E+08 | AGCTCACCTAGAAACGGCACGG | [] |
| 3219 | 9 | GLT6D1 | {'exs': 0, 'nonexs': 7} | True | 1.36E+08 | TAACAGCTAGGTGGGTTTATTG | [] |
| 3220 | 9 | SOHLH1 | {'exs': 0, 'nonexs': 7} | True | 1.36E+08 | TGGCTGCTAGCAGGCAAAGAA | [] |
| 3221 | 9 | UBAC1 | {'exs': 0, 'nonexs': 7} | True | 1.36E+08 | GCCACCTACGTGCGATTTAGT | [] |
| 3222 | 9 | LOC10192 | unavailable | False | 1.36E+08 | CCCAACTACTGGGTTCCCGCC | [] |
| 3223 | 9 | CARD9 | {'exs': 1, 'nonexs': 7} | True | 1.36E+08 | CTGCGGCTAGGAGCCCTCAGT | [] |
| 3224 | 9 | PMPCA | {'exs': 6, 'nonexs': 2} | False | 1.36E+08 | GCGGTTCTACCGGAAGAGCCG | [] |
| 3225 | 9 | SEC16A | {'exs': 4, 'nonexs': 4} | False | 1.36E+08 | GCAAGCCTAGTTCAGCACCAG | [] |
| 3226 | 9 | SEC16A | {'exs': 4, 'nonexs': 4} | False | 1.36E+08 | CCTCCCTACCCAGCCACTACA | [] |
| 3227 | 9 | AGPAT2 | {'exs': 2, 'nonexs': 7} | False | 1.37E+08 | GTCTGGGCTACTGGGCCGGCT | [] |
| 3228 | 9 | LCN6 | {'exs': 1, 'nonexs': 6} | True | 1.37E+08 | GGCCTGCTACTGTGACAGGAA | [] |
| 3229 | 9 | TMEM141 | {'exs': 0, 'nonexs': 7} | True | 1.37E+08 | CTCTCCTAGCTTCTCTGATCTG | [] |
| 3230 | 9 | CCDC183 | unavailable | False | 1.37E+08 | GGGCTACTTCTTCTTTTTCTTG | [] |
| 3231 | 9 | RABL6 | unavailable | False | 1.37E+08 | GCCGGCCTAGAGCTCCTCGTA | [] |
| 3232 | 9 | MAMDC4 | {'exs': 0, 'nonexs': 7} | True | 1.37E+08 | GGTGGTCTACGGATCACTGGT | [] |
| 3233 | 9 | PTGDS | {'exs': 0, 'nonexs': 7} | True | 1.37E+08 | GCTCACCTATTGTTCCGTCATG | [] |
| 3234 | 9 | LCNL1 | {'exs': 1, 'nonexs': 7} | True | 1.37E+08 | GTAAGCTAAAGGGAAGGGGCT | [] |
| 3235 | 9 | CLIC3 | {'exs': 0, 'nonexs': 7} | True | 1.37E+08 | GGGGCGCTAGCGGGGGTGCA | [] |
| 3236 | 9 | ENTPD2 | {'exs': 0, 'nonexs': 7} | True | 1.37E+08 | TCGGCCCCTAAATGGTGCTTG | [] |
| 3237 | 9 | SAPCD2 | unavailable | False | 1.37E+08 | CACAAGGACTAGATGAAGGTGG | [] |
| 3238 | 9 | MAN1B1 | {'exs': 0, 'nonexs': 6} | True | 1.37E+08 | TGCCTCCTAAACTGGGGGAT | [] |
| 3239 | 9 | MAN1B1 | {'exs': 0, 'nonexs': 6} | True | 1.37E+08 | TCCACCCTAGGCAGGGGTCCA | [] |
| 3240 | 9 | MAN1B1 | {'exs': 0, 'nonexs': 6} | True | 1.37E+08 | CCGGTGCTACGTGGGCCCTTG | [] |
| 3241 | 9 | LOC10192 | unavailable | False | 1.37E+08 | CCCAGCTACTTGGGAGGCTGA | [] |
| 3242 | 9 | GRIN1 | {'exs': 2, 'nonexs': 7} | False | 1.37E+08 | CCTTCCTATGGAATGGGTACCC | [] |
| 3243 | 9 | TMEM210 | {'exs': 0, 'nonexs': 2} | False | 1.37E+08 | AGCCCTCTACTCAGGTGGTAG | [] |
| 3244 | 9 | TMEM203 | {'exs': 1, 'nonexs': 5} | False | 1.37E+08 | GTGAGGCTAGTTGACCCGACA | [] |
| 3245 | 9 | RNF208 | {'exs': 0, 'nonexs': 5} | True | 1.37E+08 | GCGCTACTACATGATGGAGCA | [] |
| 3246 | 9 | CYSRT1 | unavailable | False | 1.37E+08 | GCTGGGCTAGGAGATGACACA | [] |
| 3247 | 9 | TUBB4B | unavailable | False | 1.37E+08 | CTCTAGGCCACCTCCTCCTCA | [] |
| 3248 | 9 | FAM166A | {'exs': 0, 'nonexs': 7} | True | 1.37E+08 | GGGTTTCTAAACAGGAACTGGG | [] |
| 3249 | 9 | TOR4A | unavailable | False | 1.37E+08 | CTCCACTACAGGAGGTTCACC | [] |
| 3250 | 9 | ENTPD8 | {'exs': 0, 'nonexs': 7} | True | 1.37E+08 | TTCCCACTAGTCCTGCAACCA | [] |
| 3251 | 9 | PNPLA7 | {'exs': 1, 'nonexs': 7} | True | 1.37E+08 | GGCCTCTACCCGTCCTGGTCA | [] |
| 3252 | 9 | MRPL41 | {'exs': 6, 'nonexs': 2} | False | 1.38E+08 | CAGCTAGCGCAGGAAGTTCCT | [] |
| 3253 | 9 | DPH7 | unavailable | False | 1.38E+08 | CCTCCGCTAGTCTTTCATCCA | [] |
| 3254 | 9 | DPH7 | unavailable | False | 1.38E+08 | CAGAGTCTAAGCCCTCAGAGA | [] |
| 3255 | 9 | CACNA1B | {'exs': 1, 'nonexs': 5} | False | 1.38E+08 | GCAGCTAGCACCAGTGGTCTT | [] |
| 3256 | 10 | TUBB8 | {'exs': 2, 'nonexs': 2} | False | 47056 | GTTCTAGGCCACCTCCTCCTC | [] |
| 3257 | 10 | ZMYND11 | {'exs': 0, 'nonexs': 5} | True | 249108 | ATAAGGGTCTAAAAAAGACTGG | [] |
| 3258 | 10 | DIP2C | {'exs': 0, 'nonexs': 7} | True | 277324 | AGACGAGACTACATGTTGTAGG | [] |
| 3259 | 10 | LOC10537 | unavailable | False | 954622 | CAGAACCTAAATGGGACCCGT | [] |
| 3260 | 10 | GTPBP4 | {'exs': 7, 'nonexs': 1} | False | 1017226 | ATACTATCTCCTGTCCTTTTTG | ['TTACCAGCTTTCCTCTTCCCAG'] |
| 3261 | 10 | ADARB2 | {'exs': 0, 'nonexs': 7} | True | 1183192 | CCGCAGCCTAGAGAGTCAGTA | [] |
| 3262 | 10 | AKR1C4 | {'exs': 0, 'nonexs': 5} | True | 5218759 | CCTCTATGCTAATATTCATCTG | [] |
| 3263 | 10 | UCN3 | {'exs': 0, 'nonexs': 7} | True | 5374205 | TCTACTTCTTCCTCCCAATTTG | [] |
| 3264 | 10 | NET1 | {'exs': 1, 'nonexs': 7} | True | 5456993 | TCTACACCAAAGTCTCTTTCCG | [] |
| 3265 | 10 | IL15RA | {'exs': 0, 'nonexs': 7} | True | 5963841 | GGCAGGGTCTCCTAGAGAGAG | [] |
| 3266 | 10 | IL2RA | {'exs': 1, 'nonexs': 6} | True | 6012871 | GGTTTTCTAGATTGTTCTTCTG | ['TCTTCTACTCTTCCTCCTGTAG'] |
| 3267 | 10 | GATA3 | {'exs': 0, 'nonexs': 7} | True | 8074022 | AGGGCTCTAACCCATGGCGGT | [] |
| 3268 | 10 | CAMK1D | {'exs': 1, 'nonexs': 6} | True | 12828792 | ATGAAACTACAGAGCGTGGAAG | [] |
| 3269 | 10 | BEND7 | {'exs': 0, 'nonexs': 7} | True | 13443341 | AAAGTCTAGGCAATGAAACCTG | [] |
| 3270 | 10 | BEND7 | {'exs': 0, 'nonexs': 7} | True | 13480925 | TTGGTCTAGTCCTTTCCGGTTG | [] |
| 3271 | 10 | FRMD4A | {'exs': 1, 'nonexs': 7} | True | 13651904 | ACCTCTATTCATCAGTACTTTG | [] |
| 3272 | 10 | FAM107B | {'exs': 0, 'nonexs': 7} | True | 14521189 | CAGCCTAGGACTCCTGGGCTT | [] |
| 3273 | 10 | HSPA14 | {'exs': 7, 'nonexs': 2} | False | 14843580 | GTCAAGGGCCTATCCCCACCA | [] |
| 3274 | 10 | HSPA14 | {'exs': 7, 'nonexs': 2} | False | 14871605 | CACTAAGATGCTATCTCAATAG | [] |
| 3275 | 10 | DCLRE1C | {'exs': 0, 'nonexs': 7} | True | 14911319 | ACTAAATGCTACTCTGTTCCTG | [] |
| 3276 | 10 | MEIG1 | {'exs': 0, 'nonexs': 5} | True | 14972640 | AAGACAGGCTAGTAAGCATAAG | ['AGCATAAATTTTCACTTTGTGG'] |
| 3277 | 10 | OLAH | {'exs': 0, 'nonexs': 7} | True | 15073228 | AATATCTAAAAATTGGATATCG | [] |
| 3278 | 10 | ACBD7 | {'exs': 0, 'nonexs': 7} | True | 15078529 | TTCTAAATTCCGTATTTTTCTG | ['TCTATCAGCTCCTTTGCTTTAG'] |
| 3279 | 10 | NMT2 | {'exs': 0, 'nonexs': 7} | True | 15105002 | GCCTAAATTGACTTTTTCAAAG | [] |
| 3280 | 10 | NMT2 | {'exs': 0, 'nonexs': 7} | True | 15109194 | TATCCATCTATTGTAGTACTAG | [] |
| 3281 | 10 | ITGA8 | {'exs': 0, 'nonexs': 7} | True | 15580892 | CGCTTCTATTGATCATGCACTG | [] |
| 3282 | 10 | PTER | {'exs': 0, 'nonexs': 7} | True | 16511255 | AACCATCCTATTTGAAAGTTAG | [] |
| 3283 | 10 | STAM | {'exs': 0, 'nonexs': 7} | True | 17714779 | GGTCCTATAGCAGAGCCTTCT | [] |
| 3284 | 10 | TMEM236 | unavailable | False | 17800562 | TTCCTATGCCTCATTGCAATTG | [] |
| 3285 | 10 | MRC1 | {'exs': 0, 'nonexs': 4} | True | 17910464 | TGAGGTACTAGATGACCGAGTG | [] |
| 3286 | 10 | NSUN6 | {'exs': 0, 'nonexs': 7} | True | 18545932 | ATCCCTCTCCTATGTGCTTTTG | [] |
| 3287 | 10 | MALRD1 | unavailable | False | 19734236 | TGCTGCTATTTGAGATGATGTG | [] |
| 3288 | 10 | MLLT10 | {'exs': 1, 'nonexs': 6} | True | 21556941 | ACACTACCACCTCTCCTTCTTG | [] |
| 3289 | 10 | C10orf67 | {'exs': 0, 'nonexs': 7} | True | 23265355 | CATGCTCTAAGTGAGAGAAGTG | [] |
| 3290 | 10 | C10orf67 | {'exs': 0, 'nonexs': 7} | True | 23302951 | GTCTTTCAAACTACTTTGACTG | [] |
| 3291 | 10 | C10orf67 | {'exs': 0, 'nonexs': 7} | True | 23303443 | CTTCTTCTACCTCAAAGAATTG | [] |
| 3292 | 10 | ARHGAP21 | {'exs': 0, 'nonexs': 7} | True | 24585808 | TTTTCCTAGTGATATCTTTTCG | ['CTTTTCATCTTTTGTCGTGCTG'] |
| 3293 | 10 | GPR158 | {'exs': 0, 'nonexs': 7} | True | 25599273 | CTACACTTTAAAACTATCCCAG | [] |
| 3294 | 10 | APBB1IP | {'exs': 0, 'nonexs': 7} | True | 26567487 | TGCCCGTCCCTAGGACACGTT | [] |
| 3295 | 10 | ABI1 | {'exs': 1, 'nonexs': 7} | True | 26752801 | TTTGAGAACTAGTGCTCTAGGG | [] |
| 3296 | 10 | ABI1 | {'exs': 1, 'nonexs': 7} | True | 26764586 | CATTCTTGACTAGTATGCGAGG | [] |
| 3297 | 10 | ANKRD26 | {'exs': 1, 'nonexs': 7} | True | 26992709 | GGCTATGCTACTGCCTCCTCA | [] |
| 3298 | 10 | MASTL | {'exs': 9, 'nonexs': 0} | False | 27180535 | TTGTGCTACAGACTAAATCCAG | [] |
| 3299 | 10 | RAB18 | {'exs': 2, 'nonexs': 6} | False | 27538090 | TGTCACTATCTGATCAAATATG | [] |
| 3300 | 10 | LYZL2 | {'exs': 0, 'nonexs': 7} | True | 30611914 | TGCAAACCCTAGGGCGTTGCT | [] |
| 3301 | 10 | EPC1 | {'exs': 0, 'nonexs': 7} | True | 32269062 | GGAAGCTACGTCACCTCCATC | [] |
| 3302 | 10 | CREM | {'exs': 0, 'nonexs': 7} | True | 35179328 | TATAACTAGAGAAGACTTAAAG | [] |
| 3303 | 10 | CREM | {'exs': 0, 'nonexs': 7} | True | 35211802 | ATATTTCTAGTAATCTGTTTTG | [] |
| 3304 | 10 | ZNF37A | {'exs': 0, 'nonexs': 7} | True | 38115391 | CTCTAAAACATGATTCCTAAAG | [] |
| 3305 | 10 | ZNF37A | {'exs': 0, 'nonexs': 7} | True | 38116814 | TTTTCTAGCTCTGATTCACTAG | [] |
| 3306 | 10 | ZNF37A | {'exs': 0, 'nonexs': 7} | True | 38146819 | TTCCTCTAGTGCCGCTCGGTT | [] |
| 3307 | 10 | RET | {'exs': 0, 'nonexs': 7} | True | 43126753 | CGGTGCTAGAATCTAGTAAATG | [] |
| 3308 | 10 | CSGALNA | {'exs': 0, 'nonexs': 7} | True | 43159347 | TGCATTCTAGCCTGGGTGATAG | [] |
| 3309 | 10 | HNRNPF | {'exs': 4, 'nonexs': 4} | False | 43386536 | AAACTAGTCATAGCCACCCATG | [] |
| 3310 | 10 | ZNF485 | {'exs': 1, 'nonexs': 7} | True | 43617368 | GCCACACTAACTACATTTCATG | [] |
| 3311 | 10 | CXCL12 | {'exs': 0, 'nonexs': 7} | True | 44372887 | CCTTCACTAGGCTTTGCCCAG | [] |
| 3312 | 10 | CXCL12 | {'exs': 0, 'nonexs': 7} | True | 44375938 | ATAACTAGTTTTTCCTTTTCTG | [] |
| 3313 | 10 | WASHC2C | unavailable | False | 45792399 | TGTGCTCTACTGGCCTCCAAA | [] |
| 3314 | 10 | AGAP4 | {'exs': 0, 'nonexs': 7} | True | 45825914 | CAGATACTACACACACTTGTCG | [] |
| 3315 | 10 | NCOA4 | {'exs': 1, 'nonexs': 4} | False | 46009146 | AAACTACTAGAATCCACATGGG | [] |
| 3316 | 10 | LOC10799 | unavailable | False | 46837946 | CAGATACTACACGCACTCGTC | [] |
| 3317 | 10 | SYT15 | {'exs': 2, 'nonexs': 6} | False | 46589647 | CTGGCATCTATCCGGATGTCA | [] |
| 3318 | 10 | GDF2 | {'exs': 0, 'nonexs': 4} | True | 47325783 | TACTACCTGCACCCACACTCT | [] |
| 3319 | 10 | RBP3 | {'exs': 0, 'nonexs': 4} | True | 47357456 | CCCTTCCCTACAGGTGGTCCT | [] |
| 3320 | 10 | ZNF488 | {'exs': 0, 'nonexs': 4} | True | 47367806 | GTCACCTGCTAGCTGTGAGAA | [] |
| 3321 | 10 | ANXA8 | {'exs': 0, 'nonexs': 4} | True | 47472961 | CTTCACTATTCCCTGCTGGCA | [] |
| 3322 | 10 | LOC10272 | unavailable | False | 47752172 | CTGGCATCTATCCGGATGTCA | [] |
| 3323 | 10 | ARHGAP22 | {'exs': 1, 'nonexs': 6} | True | 48532881 | TGAGGGCCTACTTTGCGCCAG | [] |
| 3324 | 10 | ARHGAP22 | {'exs': 1, 'nonexs': 6} | True | 48533523 | TTATTCTAAGTCTTCTCAATAG | [] |
| 3325 | 10 | WDFY4 | {'exs': 0, 'nonexs': 5} | True | 48808294 | CTTCATTCTATTGTCTGCCAAG | [] |
| 3326 | 10 | WDFY4 | {'exs': 0, 'nonexs': 5} | True | 48882574 | CTCTCTTCCTACCCATCTGCA | [] |
| 3327 | 10 | LRRC18 | {'exs': 0, 'nonexs': 7} | True | 48910236 | GCTACTCTAGGAAGATGTCAAG | [] |
| 3328 | 10 | VSTM4 | unavailable | False | 49019549 | GTACTACAGCTTGTTCTCCTCG | [] |
| 3329 | 10 | TMEM273 | unavailable | False | 49155884 | TCACCTACAGCTCAATCACCTG | [] |
| 3330 | 10 | TMEM273 | unavailable | False | 49157217 | CTGAGCTACCTTATGGATGAAG | [] |
| 3331 | 10 | TMEM273 | unavailable | False | 49161817 | AGTCCCTCTATTTGCAGAGGAG | [] |
| 3332 | 10 | ERCC6 | {'exs': 1, 'nonexs': 6} | True | 49515332 | GTGACACCTGCTATTCAGTGTG | [] |
| 3333 | 10 | PGBD3 | {'exs': 0, 'nonexs': 5} | True | 49515332 | GTGACACCTGCTATTCAGTGTG | [] |
| 3334 | 10 | SLC18A3 | {'exs': 0, 'nonexs': 7} | True | 49612338 | GGGATGCTAGCTGCGGGTGTA | [] |
| 3335 | 10 | OGDHL | {'exs': 0, 'nonexs': 7} | True | 49735227 | TGCCCAGCTCTAAAATGTCTTG | [] |
| 3336 | 10 | AGAP6 | {'exs': 1, 'nonexs': 6} | True | 50010185 | CAGATACTACACACACTCGTCG | [] |
| 3337 | 10 | WASHC2A | unavailable | False | 50132944 | TGTGCTCTACTGGCCTCCAAA | [] |
| 3338 | 10 | ASAH2 | {'exs': 0, 'nonexs': 7} | True | 50187314 | ACTTTTCACTAAATAGTTACAG | ['GTTACAACTTCAAAAGCCGGGG'] |
| 3339 | 10 | ASAH2B | {'exs': 1, 'nonexs': 6} | True | 50754739 | ACTTTTCACTAAATAGTTACAG | ['GTTACAACTTCAAAAGCCGGGG'] |
| 3340 | 10 | PCDH15 | {'exs': 0, 'nonexs': 7} | True | 53809479 | ACTACTTCTTCCTCCTCACTAG | [] |
| 3341 | 10 | BICC1 | {'exs': 0, 'nonexs': 7} | True | 58828890 | GTGCTGCTACCAGCGGCCACT | [] |
| 3342 | 10 | SLC16A9 | {'exs': 0, 'nonexs': 7} | True | 59852771 | ATATTCTTCTAAACATTAGAGG | [] |
| 3343 | 10 | MRLN | unavailable | False | 59737059 | AGCTTTCTAAGAAGTTATCACG | ['GTTATCACAACATATATAATAG'] |
| 3344 | 10 | CDK1 | {'exs': 7, 'nonexs': 1} | False | 60793974 | AAAGCTACATCTTCTTAATCTG | [] |
| 3345 | 10 | TMEM26 | {'exs': 0, 'nonexs': 7} | True | 61410321 | CAATAACTAAGGGGTGTGGTGG | [] |
| 3346 | 10 | TMEM26 | {'exs': 0, 'nonexs': 7} | True | 61426215 | TCTAGGAAAGAAAAATCCACAG | [] |
| 3347 | 10 | ARID5B | {'exs': 0, 'nonexs': 7} | True | 62093029 | CTGAGCCTACAGTTTTGTACTG | [] |
| 3348 | 10 | RTKN2 | {'exs': 0, 'nonexs': 7} | True | 62184321 | AAGCTATGTGAAGAGGAATTTG | [] |
| 3349 | 10 | RTKN2 | {'exs': 0, 'nonexs': 7} | True | 62197907 | GAGTTTTCTATACTTGTGCCTG | [] |
| 3350 | 10 | JMJD1C | {'exs': 0, 'nonexs': 7} | True | 63205003 | TAGTGAAACTAAGTGCTTTCTG | [] |
| 3351 | 10 | REEP3 | unavailable | False | 63620868 | GATGACTAAAAATACACTTGTG | [] |
| 3352 | 10 | CTNNA3 | {'exs': 0, 'nonexs': 5} | True | 67176900 | GACTCTTCCCTAGAGACTCCA | [] |
| 3353 | 10 | SIRT1 | {'exs': 2, 'nonexs': 5} | False | 67916592 | TTACACTATGATTTGTTTGATG | [] |
| 3354 | 10 | HERC4 | {'exs': 0, 'nonexs': 7} | True | 68070492 | CCCAGCTATTCGGGAGGCTGA | [] |
| 3355 | 10 | ATOH7 | {'exs': 0, 'nonexs': 7} | True | 68231218 | CGCGCCCTAGGTGGCCATCTG | [] |
| 3356 | 10 | PBLD | {'exs': 0, 'nonexs': 7} | True | 68284176 | CCACCTCTAGGCTGTCAGTGT | [] |
| 3357 | 10 | TET1 | {'exs': 0, 'nonexs': 7} | True | 68665851 | ATTGCTACCAAGGCATCCCTTG | [] |
| 3358 | 10 | SUPV3L1 | {'exs': 6, 'nonexs': 2} | False | 69209034 | CAGAAAACTAGTCCGAATCAGG | [] |
| 3359 | 10 | HKDC1 | {'exs': 2, 'nonexs': 6} | False | 69259179 | ACCACCACTAGACCTTATGGG | [] |
| 3360 | 10 | HKDC1 | {'exs': 2, 'nonexs': 6} | False | 69266756 | GGTTCCTAGTTCTCCTTCTGTG | [] |
| 3361 | 10 | TSPAN15 | {'exs': 1, 'nonexs': 7} | True | 69506977 | GCCCTAATTGGGGTAGCACAA | [] |
| 3362 | 10 | MACROH2 | unavailable | False | 70111682 | GCGGCGGCTACTTGGCGTCGA | [] |
| 3363 | 10 | LRRC20 | {'exs': 0, 'nonexs': 7} | True | 70301353 | AGGGTGGCCTAAGGTAGGGGG | [] |
| 3364 | 10 | PALD1 | unavailable | False | 70566732 | GGCCCCCCTACAGCAAGTCCTC | [] |
| 3365 | 10 | TBATA | unavailable | False | 70771461 | ATGTACTCTAGTGGGAGGAGAG | [] |
| 3366 | 10 | TBATA | unavailable | False | 70771687 | ATGCCTACTCTGTGCCTCTCT | [] |

| | | | | | | | |
|---|---|---|---|---|---|---|---|
| 3367 | 10 | PCBD1 | {'exs': 0, 'nonexs': 7} | True | 70883949 | AGGGTCTATGTCATGGACACTG | [] |
| 3368 | 10 | SLC29A3 | {'exs': 0, 'nonexs': 7} | True | 71362607 | CCTTCTAGATGAGGTGCACCAC | [] |
| 3369 | 10 | VSIR | unavailable | False | 71751252 | GCTGGGCTAGATGACCTCAAAC | [] |
| 3370 | 10 | PSAP | {'exs': 1, 'nonexs': 6} | True | 71817440 | CTCCTCCTAGTTCCACACATGC | [] |
| 3371 | 10 | CHST3 | {'exs': 0, 'nonexs': 7} | True | 72008470 | GCCCCCCTACGTGACCCAGAA | [] |
| 3372 | 10 | SPOCK2 | {'exs': 1, 'nonexs': 4} | False | 72062759 | CGTCTACCAGATGTAGCCCCC | [] |
| 3373 | 10 | ASCC1 | {'exs': 1, 'nonexs': 6} | True | 72102368 | CTCGATTCTAGGATTTTCCTGG | [] |
| 3374 | 10 | DNAJB12 | {'exs': 0, 'nonexs': 5} | True | 72225809 | CCAGGACTATCCATGCAGGGAC | [] |
| 3375 | 10 | OIT3 | {'exs': 2, 'nonexs': 7} | False | 72932523 | TACGAACTAGTCCTCCCAGTCC | [] |
| 3376 | 10 | NUDT13 | {'exs': 0, 'nonexs': 7} | True | 73130902 | CCGGGCTAAGCAGGCAGGGAA | [] |
| 3377 | 10 | FAM149B1 | {'exs': 2, 'nonexs': 6} | False | 73227416 | AGAAAATATGACTAAGAGCAAG | [] |
| 3378 | 10 | CFAP70 | unavailable | False | 73291208 | CTCTATTCCCTGGCTCTACCTC | [] |
| 3379 | 10 | ANXA7 | {'exs': 0, 'nonexs': 7} | True | 73376094 | CCCTCCTACTGGCCCACAATAC | [] |
| 3380 | 10 | AGAP5 | {'exs': 0, 'nonexs': 5} | True | 73674598 | CAGATACTACACGCACTCGTCC | [] |
| 3381 | 10 | NDST2 | {'exs': 0, 'nonexs': 7} | True | 73802714 | TTGCTCCGGCCTAGACAGCGAC | [] |
| 3382 | 10 | VCL | {'exs': 3, 'nonexs': 6} | False | 74118168 | GGTGCCTACTGGTACCAGGGAC | [] |
| 3383 | 10 | ADK | {'exs': 1, 'nonexs': 6} | True | 74601956 | GCTAATTTTTTTTTTTTTTCCG | ['TTTTCCATTTTTTTTTGTAGAG'] |
| 3384 | 10 | KAT6B | unavailable | False | 75031045 | CGTTGTCTACCTTCTCATGTAG | [] |
| 3385 | 10 | DUPD1 | {'exs': 1, 'nonexs': 7} | True | 75037835 | GAGTCGGGCCTACAGCTCCCT | [] |
| 3386 | 10 | COMTD1 | {'exs': 1, 'nonexs': 6} | True | 75234072 | GGCCAGCCCTAGATCTTGAAGC | [] |
| 3387 | 10 | KCNMA1 | {'exs': 0, 'nonexs': 1} | False | 77636322 | TGAGCCTAGCAACCATACATCG | [] |
| 3388 | 10 | DLG5 | {'exs': 5, 'nonexs': 4} | False | 77792439 | TTCTCCTAGAGCGGGCAGGCT | [] |
| 3389 | 10 | DLG5 | {'exs': 5, 'nonexs': 4} | False | 77800095 | TTTTCTACTACAGCATATTCAG | [] |
| 3390 | 10 | POLR3A | {'exs': 7, 'nonexs': 1} | False | 77977477 | TTTGGACTATGTGACAAGGGGG | [] |
| 3391 | 10 | NUTM2B | unavailable | False | 79712484 | GAAGCCCCTACTGGCTACAATC | [] |
| 3392 | 10 | NUTM2E | unavailable | False | 79850605 | GAAGCCCCTACTGGCTACAATC | [] |
| 3393 | 10 | PLAC9 | {'exs': 1, 'nonexs': 7} | True | 80145000 | ATGAAGCTAAGGAAGGAAGTCG | [] |
| 3394 | 10 | MAT1A | {'exs': 0, 'nonexs': 7} | True | 80273780 | GCTCTAAAATACAAGCTTCCTG | [] |
| 3395 | 10 | DYDC1 | {'exs': 0, 'nonexs': 7} | True | 80336155 | GTCCTACAAATCTTGATCAATG | [] |
| 3396 | 10 | DYDC2 | {'exs': 1, 'nonexs': 7} | True | 80366950 | TGTAACCTAAAAAGGAGATTTG | [] |
| 3397 | 10 | DYDC2 | {'exs': 1, 'nonexs': 7} | True | 80366964 | CTACCTTCTGTAACCTAAAAAG | [] |
| 3398 | 10 | C10orf99 | {'exs': 0, 'nonexs': 7} | True | 84184765 | GAGTGCTACACCTGTGGGAGTC | [] |
| 3399 | 10 | C10orf99 | {'exs': 0, 'nonexs': 7} | True | 84185088 | TGCACTCTAGCAGGATTGATGG | [] |
| 3400 | 10 | CDHR1 | {'exs': 0, 'nonexs': 7} | True | 84214620 | CATACACTAGAAGTAAGCCTTG | [] |
| 3401 | 10 | WAPL | unavailable | False | 86437542 | AAGCAGCTAGCAATGTTCCAAG | ['TCCAAATATTCAATCACTCTAG'] |
| 3402 | 10 | OPN4 | {'exs': 1, 'nonexs': 7} | True | 86665750 | CGTCCTACATCCTGGGGTCCT | [] |
| 3403 | 10 | LDB3 | {'exs': 1, 'nonexs': 7} | True | 86732975 | GGCCGCCTACAAGTTGATGGTC | [] |
| 3404 | 10 | SNCG | {'exs': 0, 'nonexs': 7} | True | 86962984 | CCCTCTAGTCTCCCCCACTCT | [] |
| 3405 | 10 | AGAP11 | unavailable | False | 87009904 | CAGATACTACACGCACTCGTCC | [] |
| 3406 | 10 | GLUD1 | {'exs': 0, 'nonexs': 7} | True | 87051750 | ATCTATGTGAAGGTCACACCAG | [] |
| 3407 | 10 | NUTM2A | unavailable | False | 87234707 | GAAGCCCCTACTGGCTACAAC | [] |
| 3408 | 10 | RNLS | {'exs': 0, 'nonexs': 7} | True | 88285353 | CACTAAATATAATTCTTTAAAG | [] |
| 3409 | 10 | LIPF | {'exs': 0, 'nonexs': 7} | True | 88678680 | CAGAACTACTTTTTATCTTCTG | [] |
| 3410 | 10 | FAS | {'exs': 0, 'nonexs': 6} | True | 89014449 | TTCACTCTAGACCAAGCTTTGG | [] |
| 3411 | 10 | IFIT1 | {'exs': 1, 'nonexs': 7} | True | 89403711 | GCCTAAGGACCTTGTCTCACAC | [] |
| 3412 | 10 | PANK1 | {'exs': 0, 'nonexs': 7} | True | 89584405 | GCTCGTCTCTACTTGTCATCAG | [] |
| 3413 | 10 | PCGF5 | {'exs': 1, 'nonexs': 7} | True | 91278315 | GTCTAACCGAAATCAATTCTTG | [] |
| 3414 | 10 | CPEB3 | {'exs': 1, 'nonexs': 7} | True | 92080031 | CCGTCTCTACTAAAAATACAAG | ['AATACAAAAAAAAAAAATTAG'] |
| 3415 | 10 | EXOC6 | {'exs': 0, 'nonexs': 7} | True | 93058354 | TGTGAGGTCTACATGTGCTGGC | [] |
| 3416 | 10 | CEP55 | {'exs': 4, 'nonexs': 4} | False | 93528152 | CTTATTTTGCTACTTTGAACAG | [] |
| 3417 | 10 | RBP4 | {'exs': 0, 'nonexs': 7} | True | 93592074 | TGCTACAAAAGGTTTCTTTCTG | [] |
| 3418 | 10 | PLCE1 | {'exs': 0, 'nonexs': 5} | True | 94283346 | GCTATTTACTTCTTTTACCTTG | [] |
| 3419 | 10 | NOC3L | {'exs': 5, 'nonexs': 3} | False | 94334176 | TTCCTCTACTAGTGTAGTGATG | [] |
| 3420 | 10 | TBC1D12 | {'exs': 0, 'nonexs': 7} | True | 94533095 | GAAGACTAGCTTTTCAAAGCAG | [] |
| 3421 | 10 | ENTPD1 | {'exs': 0, 'nonexs': 7} | True | 95866382 | TTGCTATACCATATCTTTCCAG | [] |
| 3422 | 10 | ZNF518A | unavailable | False | 96160773 | TATGGTAAACTATTCTAACATG | [] |
| 3423 | 10 | DNTT | {'exs': 1, 'nonexs': 7} | True | 96328223 | TCCTAGGCATTTCTTTCCCACG | [] |
| 3424 | 10 | TLL2 | {'exs': 0, 'nonexs': 7} | True | 96368087 | ATCAGCACTATTTCTTCATGTG | [] |
| 3425 | 10 | TM9SF3 | {'exs': 2, 'nonexs': 7} | False | 96522262 | CTCTAGTCAATTTTCACATTAG | [] |
| 3426 | 10 | TM9SF3 | {'exs': 2, 'nonexs': 7} | False | 96527235 | CTGTGCTAAATACCGCCATATG | ['GCCATATATCCAAAGTAAAATG'] |
| 3427 | 10 | LCOR | {'exs': 0, 'nonexs': 7} | True | 96955921 | CAGTATTCCTACTCGTTTTTTG | [] |
| 3428 | 10 | SLIT1 | {'exs': 0, 'nonexs': 7} | True | 97001111 | GCCCAGCGCTATGCGCAGAGG | [] |
| 3429 | 10 | C10orf62 | {'exs': 0, 'nonexs': 7} | True | 97590568 | CAGATCTAGTGTCTGCGATTAG | [] |
| 3430 | 10 | PI4K2A | {'exs': 0, 'nonexs': 7} | True | 97673741 | CTGGAGCTACCACCATGAAAAG | [] |
| 3431 | 10 | CRTAC1 | {'exs': 0, 'nonexs': 7} | True | 97878645 | GAGCATTCTATTTTCCAAGGAG | [] |
| 3432 | 10 | HPS1 | {'exs': 0, 'nonexs': 7} | True | 98417563 | TGGCCTAGAGCAGGGGGATAC | [] |
| 3433 | 10 | HPS1 | {'exs': 0, 'nonexs': 7} | True | 98429534 | AGCTAGCTATTGTTTCCTCCTG | [] |
| 3434 | 10 | NKX2-3 | {'exs': 0, 'nonexs': 7} | True | 99535720 | CCGTCCCTACCAGGCCCGGAT | [] |
| 3435 | 10 | COX15 | {'exs': 7, 'nonexs': 1} | False | 99713805 | TCTACTAAAAATACAAAATTAG | [] |
| 3436 | 10 | CUTC | {'exs': 1, 'nonexs': 7} | True | 99755738 | GAGGTCTGGCTACACCAGGATC | [] |
| 3437 | 10 | ABCC2 | {'exs': 2, 'nonexs': 6} | False | 99851630 | CTTCTGCTAGAATTTTGTGCTG | [] |
| 3438 | 10 | PKD2L1 | {'exs': 1, 'nonexs': 7} | True | 1E+08 | TGTTGTCCTAGAAGTGCAGGGC | [] |
| 3439 | 10 | HIF1AN | {'exs': 0, 'nonexs': 7} | True | 1.01E+08 | AGGCTAGTTGTATCGGCCCTTC | [] |
| 3440 | 10 | PAX2 | {'exs': 0, 'nonexs': 7} | True | 1.01E+08 | AGTCTAGGCAAGGAGCCTCCAC | [] |
| 3441 | 10 | PAX2 | {'exs': 0, 'nonexs': 7} | True | 1.01E+08 | CGGTAACTAGTGGCGGTCATAC | [] |
| 3442 | 10 | TWNK | unavailable | False | 1.01E+08 | CCGCTAAAGGCCACTCACTTTC | [] |
| 3443 | 10 | LZTS2 | {'exs': 1, 'nonexs': 7} | True | 1.01E+08 | GGGCCCTAGATCTCAGTAGCAC | [] |
| 3444 | 10 | PDZD7 | {'exs': 1, 'nonexs': 7} | True | 1.01E+08 | GGAGACTAAGGCGGGTAGATAC | [] |
| 3445 | 10 | PDZD7 | {'exs': 1, 'nonexs': 7} | True | 1.01E+08 | TTGAGACTAGCCTGGGCAACAC | [] |
| 3446 | 10 | KAZALD1 | {'exs': 1, 'nonexs': 7} | True | 1.01E+08 | GCTCTGGACCTAGTAGTAATCG | [] |
| 3447 | 10 | DPCD | unavailable | False | 1.02E+08 | GGCCAACTACTGGGTCTTGCAC | [] |
| 3448 | 10 | FBXW4 | {'exs': 0, 'nonexs': 7} | True | 1.02E+08 | TGGCCTAGAACTTGAACCTATG | [] |
| 3449 | 10 | FGF8 | {'exs': 0, 'nonexs': 7} | True | 1.02E+08 | CAGCACCTATCGGGGCTCGGG | [] |
| 3450 | 10 | NPM3 | {'exs': 1, 'nonexs': 7} | True | 1.02E+08 | AGGGCTAGGGCCTGCCCCCCT | [] |
| 3451 | 10 | KCNIP2 | {'exs': 0, 'nonexs': 7} | True | 1.02E+08 | TCTGGACTACTGAATCCCCAAG | [] |
| 3452 | 10 | KCNIP2 | {'exs': 0, 'nonexs': 7} | True | 1.02E+08 | GGGCTAGATGACATTGTCAAAG | [] |
| 3453 | 10 | ARMH3 | unavailable | False | 1.02E+08 | TCATACCTAATTGGAGAAAATG | [] |
| 3454 | 10 | MFSD13A | unavailable | False | 1.02E+08 | CATCCTCTAGAGGGCATCATTG | [] |
| 3455 | 10 | SUFU | {'exs': 0, 'nonexs': 7} | True | 1.03E+08 | CCCAGGCTAGTGTAGCGGACT | [] |
| 3456 | 10 | WBP1L | unavailable | False | 1.03E+08 | CCTGCTCTAGCTGGGGGAGCT | [] |
| 3457 | 10 | BORCS7 | unavailable | False | 1.03E+08 | TGTCATTCTATTTCAACAGATG | [] |
| 3458 | 10 | CNNM2 | {'exs': 0, 'nonexs': 7} | True | 1.03E+08 | CGCGGCCTAGATGGCGCCTTC | [] |
| 3459 | 10 | PDCD11 | {'exs': 8, 'nonexs': 1} | False | 1.03E+08 | TGCCACTAGTCCTCTAGCACTC | [] |
| 3460 | 10 | CALHM3 | {'exs': 0, 'nonexs': 7} | True | 1.03E+08 | CAGGACCCTACACGTCGGTGT | [] |
| 3461 | 10 | NEURL1 | unavailable | False | 1.04E+08 | TGGCCTATCTGTGCCCAACATC | [] |
| 3462 | 10 | NEURL1 | unavailable | False | 1.04E+08 | AACGGGCTAGGAGCTGCGGTA | [] |
| 3463 | 10 | SH3PXD2A | {'exs': 0, 'nonexs': 7} | True | 1.04E+08 | TCTGCTAGTTCTTTTTCTCAAG | [] |
| 3464 | 10 | SORCS1 | {'exs': 0, 'nonexs': 7} | True | 1.07E+08 | GGTAAAGCTACTCTCCTAAGAG | [] |
| 3465 | 10 | MXI1 | {'exs': 0, 'nonexs': 7} | True | 1.1E+08 | GGTTCTATGAAGTGAATGAAAG | [] |
| 3466 | 10 | GPAM | {'exs': 0, 'nonexs': 5} | True | 1.12E+08 | CCTACAGCACCACAAAACTCAG | [] |
| 3467 | 10 | TECTB | {'exs': 0, 'nonexs': 7} | True | 1.12E+08 | CTGGCTAACTCCTATAACACGG | [] |
| 3468 | 10 | ACSL5 | {'exs': 0, 'nonexs': 7} | True | 1.12E+08 | CCTTATCCTAATCCTGGATGTG | [] |
| 3469 | 10 | ZDHHC6 | {'exs': 0, 'nonexs': 7} | True | 1.12E+08 | -1 | -1 |
| 3470 | 10 | TCF7L2 | {'exs': 2, 'nonexs': 6} | False | 1.13E+08 | GAATCTAGTAAGCTTCCATCTG | [] |
| 3471 | 10 | TCF7L2 | {'exs': 2, 'nonexs': 6} | False | 1.13E+08 | CTAAAGCTATTCTAAAGACTTG | [] |
| 3472 | 10 | CASP7 | {'exs': 0, 'nonexs': 7} | True | 1.14E+08 | ATATGGCTATTGACTGAAGTAG | [] |
| 3473 | 10 | NHLRC2 | {'exs': 3, 'nonexs': 5} | False | 1.14E+08 | GGCTGGCTAAAATACATACCTG | [] |
| 3474 | 10 | ADRB1 | {'exs': 0, 'nonexs': 7} | True | 1.14E+08 | GGGCCCTACACCTTGGATTCC | [] |
| 3475 | 10 | AFAP1L2 | {'exs': 1, 'nonexs': 6} | True | 1.14E+08 | -1 | -1 |
| 3476 | 10 | ATRNL1 | {'exs': 1, 'nonexs': 7} | True | 1.16E+08 | GATGGCCTATTGTGGGACCTTC | [] |
| 3477 | 10 | GFRA1 | {'exs': 0, 'nonexs': 7} | True | 1.16E+08 | TAATGCAGCTATGATGTTTCTG | [] |
| 3478 | 10 | PNLIP | {'exs': 0, 'nonexs': 7} | True | 1.17E+08 | AGTCTCCTAACACGGTGTGAGC | [] |
| 3479 | 10 | C10orf82 | {'exs': 0, 'nonexs': 7} | True | 1.17E+08 | TTCTAGCCTTCTGCATACTTTG | [] |
| 3480 | 10 | ENO4 | {'exs': 0, 'nonexs': 3} | True | 1.17E+08 | GTACAGCCCTATCCAGAGGATC | [] |
| 3481 | 10 | SHTN1 | unavailable | False | 1.17E+08 | CAATGCTACTGGGAGGCCTATC | [] |
| 3482 | 10 | VAX1 | {'exs': 0, 'nonexs': 7} | True | 1.17E+08 | GGGCCCCGCACTATAGGGGCT | [] |
| 3483 | 10 | PDZD8 | {'exs': 1, 'nonexs': 7} | True | 1.17E+08 | GTCTGCTACACAGACTCGGATC | [] |
| 3484 | 10 | CACUL1 | unavailable | False | 1.19E+08 | CATTCACTATCTGTACCCCCTG | [] |
| 3485 | 10 | SFXN4 | {'exs': 0, 'nonexs': 7} | True | 1.19E+08 | CTCACGCCTACACCCCTCTGT | [] |
| 3486 | 10 | PRDX3 | {'exs': 1, 'nonexs': 6} | True | 1.19E+08 | TGATCTACTGATTTACCTTCTG | [] |
| 3487 | 10 | GRK5 | {'exs': 1, 'nonexs': 6} | True | 1.19E+08 | CCGAAACTAGCTGCTTCCGGTC | [] |
| 3488 | 10 | BAG3 | {'exs': 0, 'nonexs': 7} | True | 1.2E+08 | CAGAGGCTACGGTGCTGCTGG | [] |
| 3489 | 10 | INPP5F | {'exs': 0, 'nonexs': 7} | True | 1.2E+08 | AAAGCTAAATCTGAATTATCCG | [] |
| 3490 | 10 | FGFR2 | {'exs': 2, 'nonexs': 5} | False | 1.21E+08 | AGTCCTACTGGTCCACAGCCAC | [] |
| 3491 | 10 | PLEKHA1 | {'exs': 0, 'nonexs': 7} | True | 1.22E+08 | CTCCACTAAGAAGGCACGTTTG | [] |
| 3492 | 10 | HTRA1 | {'exs': 0, 'nonexs': 7} | True | 1.23E+08 | TGCCTATGGGTCAATTTCTTCG | [] |
| 3493 | 10 | DMBT1 | {'exs': 1, 'nonexs': 5} | False | 1.23E+08 | CCTACCGAGGCTCCTCTTCTC | [] |
| 3494 | 10 | CPXM2 | {'exs': 0, 'nonexs': 7} | True | 1.24E+08 | CATCCTAATGTCCCAAAGTCTG | [] |
| 3495 | 10 | CTBP2 | {'exs': 0, 'nonexs': 7} | True | 1.25E+08 | TCTCTGCTATTGCTCGTTGGGC | [] |
| 3496 | 10 | INSYN2A | unavailable | False | 1.27E+08 | ATAGCTTCCTAAGACATGGGTG | [] |
| 3497 | 10 | PTPRE | {'exs': 1, 'nonexs': 6} | True | 1.28E+08 | CCCCGCTAAGGGCGAACGCT | [] |
| 3498 | 10 | EBF3 | {'exs': 2, 'nonexs': 5} | False | 1.3E+08 | GGGACTACCAGCCCAGACATAC | [] |
| 3499 | 10 | C10orf143 | unavailable | False | 1.3E+08 | TCTTCTAATGATTCTTGGTATG | [] |
| 3500 | 10 | PPP2R2D | {'exs': 1, 'nonexs': 1} | False | 1.32E+08 | GTCTCTAGTTGATTTTGTCCTG | [] |
| 3501 | 10 | JAKMIP3 | {'exs': 0, 'nonexs': 7} | True | 1.32E+08 | GACGGACTATGACCAGAGAATG | [] |
| 3502 | 10 | DPYSL4 | {'exs': 1, 'nonexs': 7} | True | 1.32E+08 | GGGCGTCTAGGAGAGAGAGGT | [] |
| 3503 | 10 | STK32C | {'exs': 2, 'nonexs': 6} | False | 1.32E+08 | CCGGCCTAGCCGCTCCCGGC | [] |
| 3504 | 10 | STK32C | {'exs': 2, 'nonexs': 6} | False | 1.32E+08 | CTCCTCTATACCATGTGAGTTG | [] |
| 3505 | 10 | ADGRA1 | unavailable | False | 1.33E+08 | CCCCATCTACACAGTAGTTTCG | [] |
| 3506 | 10 | KNDC1 | {'exs': 1, 'nonexs': 7} | True | 1.33E+08 | GTCCCTATAGCCCAGTGAACC | [] |
| 3507 | 10 | KNDC1 | {'exs': 1, 'nonexs': 7} | True | 1.33E+08 | GAGCTCGGCTACTGGAATGTAC | [] |
| 3508 | 10 | ADAM8 | {'exs': 0, 'nonexs': 6} | True | 1.33E+08 | CCCCCCTAGGGTGCTGTGGGA | [] |
| 3509 | 10 | PRAP1 | {'exs': 1, 'nonexs': 7} | True | 1.33E+08 | GGAGCCCTACTGGGGGTGGTA | [] |
| 3510 | 10 | FUOM | unavailable | False | 1.33E+08 | CCAGGCCTACAGCAGGGGGTT | [] |
| 3511 | 10 | PAOX | {'exs': 1, 'nonexs': 7} | True | 1.33E+08 | GCCCAGCTAGAGCCTGGGCCT | [] |
| 3512 | 10 | SPRN | {'exs': 0, 'nonexs': 5} | True | 1.33E+08 | GCCAGGCCTAGGGCCGCAGCA | [] |
| 3513 | 11 | BET1L | {'exs': 1, 'nonexs': 7} | True | 205469 | AGTCCTAAGGGAGGAATCCCAC | [] |
| 3514 | 11 | SIRT3 | {'exs': 0, 'nonexs': 7} | True | 216697 | AGCCATCATCCTATTTGTCTGG | [] |
| 3515 | 11 | PSMD13 | {'exs': 8, 'nonexs': 0} | False | 252599 | GGCCCTAGGTGAGGATGTCATC | [] |
| 3516 | 11 | PGGHG | unavailable | False | 293592 | CCTCTCCTACAGGGAGGTGCC | [] |
| 3517 | 11 | IFITM2 | {'exs': 2, 'nonexs': 6} | False | 309164 | CCTGATCTATCGCTGGGCCTG | [] |
| 3518 | 11 | IFITM1 | {'exs': 0, 'nonexs': 7} | True | 315112 | TGGGCGGCTACTAGTAACCCC | [] |
| 3519 | 11 | IFITM3 | {'exs': 0, 'nonexs': 7} | True | 319837 | CCTGATCTATCCATAGGCCTGC | [] |
| 3520 | 11 | PKP3 | {'exs': 0, 'nonexs': 7} | True | 404568 | CTTCACCTATGGGCCCAGGAAC | [] |
| 3521 | 11 | SIGIRR | {'exs': 0, 'nonexs': 7} | True | 405895 | TGGGAGCTACATATCATCCTTG | [] |
| 3522 | 11 | ANO9 | {'exs': 0, 'nonexs': 7} | True | 418370 | GCCCTACACGTCTGTGCTCCT | [] |
| 3523 | 11 | LOC101059 | unavailable | False | 531296 | GGTTCCCTATCCCTGAGGTTGC | [] |
| 3524 | 11 | LMNTD2 | unavailable | False | 554979 | CCGCCCCTAGGCGCCGCGGC | [] |
| 3525 | 11 | IRF7 | {'exs': 0, 'nonexs': 7} | True | 612644 | GGTTCTAGGCGGGCTGCTCCA | [] |
| 3526 | 11 | DEAF1 | {'exs': 0, 'nonexs': 7} | True | 646528 | TTCTCTGGTCTACAGGTTTGAG | [] |
| 3527 | 11 | DEAF1 | {'exs': 0, 'nonexs': 7} | True | 651314 | TGCCTATAATCCTAGCACTTTG | [] |
| 3528 | 11 | DEAF1 | {'exs': 0, 'nonexs': 7} | True | 659772 | GCCGCCTAATTCCACAGGCCC | [] |
| 3529 | 11 | TMEM80 | {'exs': 0, 'nonexs': 7} | True | 703149 | CGTAGCTACCTGGTGAAGGCC | [] |
| 3530 | 11 | EPS8L2 | {'exs': 0, 'nonexs': 7} | True | 720880 | -1 | -1 |
| 3531 | 11 | TALDO1 | {'exs': 0, 'nonexs': 7} | True | 764844 | ATGCGCTACTTTCCATTCTCTG | [] |
| 3532 | 11 | GATD1 | unavailable | False | 772159 | GAGACTAGGGGCCAGAGGTTC | [] |
| 3533 | 11 | PANO1 | unavailable | False | 798272 | CCCCGCCTAGGTGCTGGCCGA | [] |
| 3534 | 11 | PIDD1 | unavailable | False | 799306 | GTGGGGCCTAGGCCTGGGCAG | [] |
| 3535 | 11 | TSPAN4 | {'exs': 1, 'nonexs': 7} | True | 866629 | GGCGGCCTACGCGCAGTAGGT | [] |
| 3536 | 11 | CHID1 | {'exs': 0, 'nonexs': 7} | True | 869857 | GCCCACCTAGAGCAGGTCGTAC | [] |
| 3537 | 11 | CHID1 | {'exs': 0, 'nonexs': 7} | True | 874780 | CCGGCTGCTAGGGCTGTCCTG | [] |
| 3538 | 11 | AP2A2 | {'exs': 0, 'nonexs': 7} | True | 1010624 | TCAGGACTAAAACTGCGCTGAG | [] |
| 3539 | 11 | MUC6 | {'exs': 0, 'nonexs': 7} | True | 1013455 | CGACCCTGCTAGTCTCCACAG | [] |
| 3540 | 11 | TOLLIP | {'exs': 0, 'nonexs': 7} | True | 1277038 | TCTATGGCTCCTCCCCCATCTC | [] |
| 3541 | 11 | BRSK2 | {'exs': 1, 'nonexs': 6} | True | 1460722 | GTGTGTCTAAGGCTGCTCGCG | [] |
| 3542 | 11 | CTSD | {'exs': 0, 'nonexs': 7} | True | 1753502 | CTTGGGAACTAGAGGCGGGCA | [] |
| 3543 | 11 | TNNI2 | {'exs': 0, 'nonexs': 7} | True | 1841550 | CAGCGAGTGGCCTAGGACTCG | [] |
| 3544 | 11 | LSP1 | {'exs': 0, 'nonexs': 7} | True | 1887562 | ATGGGACGCCTAGGGAGCCGG | [] |
| 3545 | 11 | TNNT3 | {'exs': 1, 'nonexs': 7} | True | 1938491 | GCCTCTCTACTTCCAGCGCCC | [] |
| 3546 | 11 | INS | {'exs': 2, 'nonexs': 6} | False | 2159851 | GGGCTGCGTCTAGTTGCAGTAC | [] |
| 3547 | 11 | TH | {'exs': 1, 'nonexs': 7} | True | 2164232 | GCACCTAGCCAATGGCACTCAC | [] |
| 3548 | 11 | TSPAN32 | {'exs': 0, 'nonexs': 7} | True | 2316071 | GGCTCTAGGGAAAGGCAGCAC | [] |
| 3549 | 11 | TSPAN32 | {'exs': 0, 'nonexs': 7} | True | 2317655 | GGTGAGCTACCCCTTACCCTTC | [] |
| 3550 | 11 | SLC22A18 | {'exs': 0, 'nonexs': 7} | True | 2888179 | TGTACTCTATGTTGCTGCAGAG | [] |
| 3551 | 11 | NAP1L4 | {'exs': 0, 'nonexs': 7} | True | 2945607 | CTCCGCTTCCTACTGCTGCTTC | [] |
| 3552 | 11 | OSBPL5 | {'exs': 1, 'nonexs': 7} | True | 3088204 | GGGCTCCTATTTGAGGATGTGC | [] |
| 3553 | 11 | STIM1 | {'exs': 0, 'nonexs': 7} | True | 4091797 | ATCCTGCCTACTTCTTAAGAGG | [] |
| 3554 | 11 | OR52K2 | {'exs': 0, 'nonexs': 3} | True | 4450283 | ACCCTCTACATATCCTTTCTTG | [] |
| 3555 | 11 | OR52K1 | {'exs': 0, 'nonexs': 3} | True | 4489844 | CCATCTACATGTTCTTTCTCTG | [] |
| 3556 | 11 | OR51E1 | {'exs': 0, 'nonexs': 3} | True | 4653482 | TGACACCTAGGGCTCTGAAGCC | [] |
| 3557 | 11 | OR51J1 | {'exs': 0, 'nonexs': 1} | False | 5403546 | TGGTTTTTGGCTACACTCTGCG | [] |
| 3558 | 11 | UBQLN3 | {'exs': 1, 'nonexs': 6} | True | 5507590 | AAGGCTCCTACGACTGTCTCAC | [] |
| 3559 | 11 | UBQLNL | {'exs': 0, 'nonexs': 7} | True | 5515313 | GGGTTGCCTAAGCACTAAGCAC | [] |
| 3560 | 11 | TRIM5 | {'exs': 0, 'nonexs': 7} | True | 5645855 | TTCATCTCATCTAGTCTTCTGG | [] |
| 3561 | 11 | TRIM5 | {'exs': 0, 'nonexs': 7} | True | 5664034 | TCTACTGAAAATACAAATTTAG | [] |
| 3562 | 11 | TRIM5 | {'exs': 0, 'nonexs': 7} | True | 5664087 | CCCAGCTACTTGGGAGGCTGA | [] |
| 3563 | 11 | OR56B1 | {'exs': 0, 'nonexs': 3} | True | 5737490 | CTCTAAGATCTTATTTCTTTTG | [] |
| 3564 | 11 | SMPD1 | {'exs': 0, 'nonexs': 7} | True | 6394217 | TTCCATCTATTTGGTACACACG | [] |
| 3565 | 11 | SMPD1 | {'exs': 0, 'nonexs': 7} | True | 6394506 | TGGGGCCCTAGCAAAACAGTGC | [] |
| 3566 | 11 | TRIM3 | {'exs': 1, 'nonexs': 5} | False | 6449027 | TGTACAGCTACTGGAGGTAGCC | [] |
| 3567 | 11 | ARFIP2 | {'exs': 0, 'nonexs': 7} | True | 6477322 | GGGTCTATGAGCCTGAGCCCA | [] |
| 3568 | 11 | TIMM10B | unavailable | False | 6482220 | TATGGCTAGCTGCCTGATGGAC | [] |
| 3569 | 11 | ILK | {'exs': 4, 'nonexs': 5} | False | 6610510 | CCTTCCAGTCCTACTTGTCCTC | [] |
| 3570 | 11 | DCHS1 | {'exs': 1, 'nonexs': 7} | True | 6621778 | CACAGCTAGATGTGCAGCTCCC | [] |
| 3571 | 11 | OR2AG2 | {'exs': 0, 'nonexs': 3} | True | 6768006 | CTTCCCTAGAGCGTGGAATGTC | [] |
| 3572 | 11 | OR2AG1 | {'exs': 0, 'nonexs': 3} | True | 6785987 | CTTCCCTAGAGCGTGGAGTGTC | [] |
| 3573 | 11 | OR6A2 | {'exs': 0, 'nonexs': 3} | True | 6794724 | CCCTTCTATACATTTCTGCTAG | [] |
| 3574 | 11 | OR10A5 | {'exs': 0, 'nonexs': 3} | True | 6846635 | GGTCTATGGGATACAGTTTCTG | [] |
| 3575 | 11 | OR10A2 | {'exs': 0, 'nonexs': 3} | True | 6870665 | GGTCTATGGGATACAGTTTCTG | [] |
| 3576 | 11 | ZNF215 | {'exs': 0, 'nonexs': 7} | True | 6984237 | CCCAGCTACTCAGGAGGCTGA | [] |
| 3577 | 11 | NLRP14 | {'exs': 0, 'nonexs': 7} | True | 7089011 | GCGCCTAGTCGCCGAGTCCAC | [] |
| 3578 | 11 | SYT9 | {'exs': 0, 'nonexs': 7} | True | 7314195 | TTAGGGGCTAGCTAAGAATCTG | [] |
| 3579 | 11 | SYT9 | {'exs': 0, 'nonexs': 7} | True | 7432800 | GTGCCTTCTACCCATGAGCCTC | [] |
| 3580 | 11 | OR5P2 | {'exs': 0, 'nonexs': 3} | True | 7795873 | TTCTATGTAATATCATTATTTG | [] |
| 3581 | 11 | TUB | {'exs': 0, 'nonexs': 7} | True | 8101518 | GAGGCCTCTACTCGCACGCCA | [] |
| 3582 | 11 | AKIP1 | unavailable | False | 8917465 | ACTATAGACTCAATTCAAATAG | [] |
| 3583 | 11 | C11orf16 | {'exs': 0, 'nonexs': 7} | True | 8921315 | CTTAGTCTAACGGGAATTCTTG | [] |
| 3584 | 11 | ADM | {'exs': 0, 'nonexs': 7} | True | 10306640 | AAATCCTAAAGAAAGTGGGGAG | [] |
| 3585 | 11 | AMPD3 | {'exs': 0, 'nonexs': 7} | True | 10505883 | CTGGACCTAGTTGGTCAAGGCC | [] |
| 3586 | 11 | MTRNR2L | {'exs': 0, 'nonexs': 6} | True | 10508151 | ATACTATGCCCGCCTCTTCACC | [] |
| 3587 | 11 | LYVE1 | {'exs': 0, 'nonexs': 7} | True | 10559110 | TCATCTAAACTTCAGCTTCCAG | [] |
| 3588 | 11 | MRVI1 | {'exs': 0, 'nonexs': 7} | True | 10576331 | AGGTTTCCTACTGCTCTGTAGG | [] |
| 3589 | 11 | CTR9 | {'exs': 6, 'nonexs': 2} | False | 10779104 | AAAACCTAGTCACTATCATCTG | [] |
| 3590 | 11 | GALNT18 | unavailable | False | 11256316 | CAGGAAAACTAGAATAACAATG | [] |
| 3591 | 11 | DKK3 | {'exs': 1, 'nonexs': 7} | True | 11964453 | TCTAAATCTCTTCCCCTCCCAG | [] |
| 3592 | 11 | RASSF10 | unavailable | False | 13011099 | CCACCCCTACACAAGGGATTCC | [] |
| 3593 | 11 | SPON1 | {'exs': 0, 'nonexs': 1} | False | 14265586 | TTGCTAACAAGGATGAACATTG | [] |
| 3594 | 11 | RRAS2 | {'exs': 1, 'nonexs': 5} | False | 14279336 | GATTCTAGAAAATGACACAATG | [] |
| 3595 | 11 | PDE3B | {'exs': 0, 'nonexs': 7} | True | 14869659 | TGTCGCTATTCCTCTTCATCTG | [] |
| 3596 | 11 | INSC | {'exs': 0, 'nonexs': 7} | True | 15248339 | CTACACAAAACTCTCCTCCATG | [] |
| 3597 | 11 | INSC | {'exs': 0, 'nonexs': 7} | True | 15269626 | CTCACTTACCTAGTTCAGATCG | ['CAGATCATCTTCTTTACACATG'] |
| 3598 | 11 | PLEKHA7 | {'exs': 0, 'nonexs': 7} | True | 16789086 | CCTCCTAACATACTGAGCCAAG | [] |
| 3599 | 11 | USH1C | {'exs': 0, 'nonexs': 7} | True | 17494331 | TGTTTCCTAACGGTGAATTTGG | [] |
| 3600 | 11 | USH1C | {'exs': 0, 'nonexs': 7} | True | 17495252 | AAGTACTACAGACTGGGCTTTG | [] |
| 3601 | 11 | SAA2 | {'exs': 0, 'nonexs': 7} | True | 18239947 | TTCACTCTCTACAGTTCAGCTG | [] |
| 3602 | 11 | HPS5 | {'exs': 1, 'nonexs': 7} | True | 18279881 | TTCTCCCACTAGGCCTGCTGG | [] |
| 3603 | 11 | UEVLD | {'exs': 0, 'nonexs': 7} | True | 18563965 | GTACCACTACACTCCAGCCTGC | [] |
| 3604 | 11 | SPTY2D1 | {'exs': 1, 'nonexs': 7} | True | 18609860 | GCAGCAGCTAACGCCTCTTCAC | [] |
| 3605 | 11 | MRGPRX2 | {'exs': 0, 'nonexs': 7} | True | 19055409 | TGTCCATCTCTACACCAGACTG | [] |
| 3606 | 11 | DBX1 | {'exs': 0, 'nonexs': 1} | False | 20156213 | GAGCGGCTTCTAGGACACGGT | [] |
| 3607 | 11 | SLC6A5 | {'exs': 1, 'nonexs': 7} | True | 20654867 | TGGACTAGCACTGAGTGCCCAC | [] |
| 3608 | 11 | FANCF | {'exs': 2, 'nonexs': 6} | False | 22624685 | CATTGCCTATACAGAACTGAGG | [] |
| 3609 | 11 | GAS2 | {'exs': 1, 'nonexs': 6} | True | 22766448 | AATGTCTATATGGTCTGAAATG | [] |
| 3610 | 11 | ANO3 | {'exs': 0, 'nonexs': 7} | True | 26660443 | TGTCAACTAAGGCCATTCATGG | [] |
| 3611 | 11 | MUC15 | {'exs': 0, 'nonexs': 7} | True | 26561064 | GTTAGTTCTATACAGAAGTACG | [] |
| 3612 | 11 | FIBIN | {'exs': 0, 'nonexs': 7} | True | 26995161 | ATCCTTCCACCTATACTTTAAG | [] |
| 3613 | 11 | CCDC34 | {'exs': 0, 'nonexs': 7} | True | 27338820 | CATACGCTATCTTTGTATTCTG | [] |
| 3614 | 11 | BDNF | {'exs': 0, 'nonexs': 7} | True | 27657820 | CCACTATCTTCCCCTTTTAATG | [] |
| 3615 | 11 | METTL15 | unavailable | False | 28328137 | ATTTTCCTAATAAAGAGTTTGG | [] |
| 3616 | 11 | MPPED2 | {'exs': 0, 'nonexs': 7} | True | 30388888 | CTAGACTAGGAAGTTTTTGAAG | [] |
| 3617 | 11 | DCDC1 | {'exs': 0, 'nonexs': 1} | False | 30912837 | TATACCCTAATTTACTAGCCTG | [] |
| 3618 | 11 | DCDC1 | {'exs': 0, 'nonexs': 1} | False | 30944310 | ACCAAGTCTATCTCCTTCCTTG | [] |
| 3619 | 11 | DCDC1 | {'exs': 0, 'nonexs': 1} | False | 31135596 | AGAGGCTATGGAATATATTTAG | [] |
| 3620 | 11 | IMMP1L | {'exs': 0, 'nonexs': 7} | True | 31432499 | TTACTAATCATCAGAAAATCTG | [] |
| 3621 | 11 | ELP4 | {'exs': 4, 'nonexs': 4} | False | 31783537 | GCGACTAAGGAGGAATCCCTAC | [] |
| 3622 | 11 | PAX6 | {'exs': 0, 'nonexs': 7} | True | 31789629 | GTCTACATTGTTCTTTTTTTCG | ['TTTTCATTATAACATACAAATG'] |
| 3623 | 11 | QSER1 | {'exs': 0, 'nonexs': 7} | True | 32975555 | AGCTCCTACAAGAGGTGGTGAC | [] |
| 3624 | 11 | CSTF3 | {'exs': 6, 'nonexs': 2} | False | 33085286 | AACCCTACCGAATCCGCTTCTC | [] |
| 3625 | 11 | CSTF3 | {'exs': 6, 'nonexs': 2} | False | 33141579 | CTCCTTGTCCTAATTGGCTATG | [] |
| 3626 | 11 | CSTF3 | {'exs': 6, 'nonexs': 2} | False | 33141878 | ATCACTAAACCTGTGCCTCTCG | [] |
| 3627 | 11 | FBXO3 | {'exs': 0, 'nonexs': 7} | True | 33741907 | GGCTTGCTAAAAAAGGCGTGAG | [] |
| 3628 | 11 | LMO2 | {'exs': 4, 'nonexs': 4} | False | 33859355 | TCGGGCCTATATCATCCCATTG | [] |
| 3629 | 11 | NAT10 | {'exs': 7, 'nonexs': 1} | False | 34146191 | TTCACTATTTCTTCCGCTTCAG | [] |
| 3630 | 11 | PDHX | {'exs': 0, 'nonexs': 7} | True | 34995171 | GAGGACTAGGCAAGTCGGATAC | [] |
| 3631 | 11 | FJX1 | {'exs': 1, 'nonexs': 6} | True | 35619949 | GTGACACTAAGTCCCAGACCGC | [] |
| 3632 | 11 | TRAF6 | {'exs': 1, 'nonexs': 7} | True | 36489837 | GCAAGCTATACCCCTGCATCAC | [] |
| 3633 | 11 | RAG2 | {'exs': 1, 'nonexs': 7} | True | 36592584 | ACTAATCAAACAACCTTCTAAG | [] |
| 3634 | 11 | API5 | {'exs': 2, 'nonexs': 5} | False | 43339277 | GTCTTACTAGGGTGATGTCTTG | [] |
| 3635 | 11 | API5 | {'exs': 2, 'nonexs': 5} | False | 43342454 | CCACCTCTACTTCCCCTGAAGC | [] |
| 3636 | 11 | TTC17 | {'exs': 0, 'nonexs': 7} | True | 43445964 | TTATGTCTATGGCACTTTCCTG | [] |
| 3637 | 11 | TTC17 | {'exs': 0, 'nonexs': 7} | True | 43493903 | AAGTGCACTAAGGAGAGCGCC | [] |
| 3638 | 11 | ACCSL | {'exs': 0, 'nonexs': 7} | True | 44059919 | CAGACGGCCTACTCCCTCATTC | [] |
| 3639 | 11 | EXT2 | {'exs': 0, 'nonexs': 7} | True | 44192756 | CATGTACTAGTCTTGTGACATG | [] |
| 3640 | 11 | TSPAN18 | {'exs': 0, 'nonexs': 5} | True | 44929177 | TACCCTCTACTGGATGCCCCG | [] |
| 3641 | 11 | TP53I11 | {'exs': 1, 'nonexs': 7} | True | 44934883 | CGGGCAACTAGGCCTTCTTGG | [] |
| 3642 | 11 | PRDM11 | {'exs': 0, 'nonexs': 7} | True | 45224907 | CTTTAGGCTACCCTCCACACAC | [] |
| 3643 | 11 | PRDM11 | {'exs': 0, 'nonexs': 7} | True | 45228158 | GCCAGCTCCCTAAATGTCGGG | [] |
| 3644 | 11 | LOC101060 | unavailable | False | 45213797 | GCCCTCTCACTAACTGGCTGTC | [] |
| 3645 | 11 | SLC35C1 | {'exs': 1, 'nonexs': 7} | True | 45808178 | GGGGCTAGGAGGGGAGGCCTA | [] |
| 3646 | 11 | MAPK8IP1 | {'exs': 1, 'nonexs': 7} | True | 45905720 | GCTGCACAGCTACTCCAGGTAC | [] |
| 3647 | 11 | LARGE2 | unavailable | False | 45927832 | CACAGGACTAGACCACCATGGC | [] |
| 3648 | 11 | CREB3L1 | {'exs': 1, 'nonexs': 5} | False | 46320745 | CATGGCCTAGGAGAGTTTGATG | [] |
| 3649 | 11 | DGKZ | {'exs': 0, 'nonexs': 7} | True | 46378673 | TGTGGCCTAGACAGAGATACAG | [] |
| 3650 | 11 | DGKZ | {'exs': 0, 'nonexs': 7} | True | 46379946 | CCCGCTACACAGCCGTCTCCT | [] |
| 3651 | 11 | MDK | {'exs': 1, 'nonexs': 7} | True | 46383493 | -1 | -1 |
| 3652 | 11 | CHRM4 | {'exs': 1, 'nonexs': 6} | True | 46385117 | GCCTGCCTACCTGGCAGTGCC | [] |
| 3653 | 11 | AMBRA1 | {'exs': 1, 'nonexs': 7} | True | 46397449 | TTTGTCTCTACCTGTTCCGTGG | [] |
| 3654 | 11 | AMBRA1 | {'exs': 1, 'nonexs': 7} | True | 46430988 | GCACAGCTATCCAAAAAGCTG | [] |
| 3655 | 11 | F2 | {'exs': 1, 'nonexs': 7} | True | 46729407 | TGGCCCCCTACTCTCCAAACTC | [] |
| 3656 | 11 | ACP2 | {'exs': 1, 'nonexs': 6} | True | 47240082 | ACCTCCCCTAGGAGGTGGAGG | [] |
| 3657 | 11 | NR1H3 | {'exs': 1, 'nonexs': 6} | True | 47268259 | CCGGTCTATAGGTCACCAAGAC | [] |
| 3658 | 11 | PSMC3 | {'exs': 7, 'nonexs': 1} | False | 47418834 | GTGTGCCCTAGGCGTAGTATTC | [] |
| 3659 | 11 | RAPSN | {'exs': 1, 'nonexs': 7} | True | 47437786 | TATTTTTCTATAAAGAGCAAAG | [] |
| 3660 | 11 | NDUFS3 | {'exs': 2, 'nonexs': 6} | False | 47584480 | CCTGGAGCTACTTGGCATCAGC | [] |
| 3661 | 11 | MTCH2 | {'exs': 4, 'nonexs': 4} | False | 47606155 | TTTGAATGGCTAGGTGGTTAGG | [] |
| 3662 | 11 | MTCH2 | {'exs': 4, 'nonexs': 4} | False | 47618799 | TGGGACTACAGAAATGTCACTG | [] |
| 3663 | 11 | AGBL2 | {'exs': 1, 'nonexs': 5} | False | 47659417 | GGGCTCCTAGCATGAGCTGGA | [] |
| 3664 | 11 | AGBL2 | {'exs': 1, 'nonexs': 5} | False | 47660172 | CACCTACGGGTATGTGTATATG | [] |
| 3665 | 11 | AGBL2 | {'exs': 1, 'nonexs': 5} | False | 47661475 | TTGTCACTCCTAGAGGATCTTG | [] |
| 3666 | 11 | FNBP4 | {'exs': 6, 'nonexs': 2} | False | 47717421 | CTTAAAAACTATGTGTTTGGAG | [] |
| 3667 | 11 | FNBP4 | {'exs': 6, 'nonexs': 2} | False | 47730149 | GAGCCACCTATCCAGAACTACC | [] |
| 3668 | 11 | OR4X2 | {'exs': 0, 'nonexs': 3} | True | 48246014 | TCAGCCTCTATTTCTCATTTAG | [] |
| 3669 | 11 | OR4C5 | {'exs': 0, 'nonexs': 3} | True | 48365494 | TCTTCTCTAATCACACAAATTG | [] |
| 3670 | 11 | OR4A5 | {'exs': 0, 'nonexs': 3} | True | 54707831 | CTCCTTACCTACATGAGGACGC | [] |
| 3671 | 11 | OR4S2 | {'exs': 0, 'nonexs': 3} | True | 55651838 | AAGTCCAACTATTTCCCTTTAG | [] |
| 3672 | 11 | OR5L1 | {'exs': 0, 'nonexs': 3} | True | 55812401 | TCTTCCCTAGGAGTGAATTTTG | [] |
| 3673 | 11 | OR5L2 | {'exs': 0, 'nonexs': 3} | True | 55828153 | TCTTCCCTAGGAGTGAATTTTG | [] |
| 3674 | 11 | OR8H3 | {'exs': 0, 'nonexs': 3} | True | 56123310 | ATTTTAACTACCTGGAGTCCTG | [] |
| 3675 | 11 | OR5T1 | {'exs': 0, 'nonexs': 3} | True | 56276518 | TTAAAACTATAACTTATTTATG | [] |
| 3676 | 11 | OR8U1 | {'exs': 0, 'nonexs': 3} | True | 56370552 | ACTCTAGTTTTTATTGATAATG | [] |
| 3677 | 11 | OR5M10 | {'exs': 0, 'nonexs': 3} | True | 56576773 | CTAAACTGCAATTTTACAAAAG | [] |
| 3678 | 11 | OR5M1 | {'exs': 0, 'nonexs': 3} | True | 56612554 | AAGCCTAAACTGCAATTTTATG | [] |
| 3679 | 11 | OR5AP2 | {'exs': 0, 'nonexs': 2} | False | 56641488 | TAACATGGCTCTACAAGATGTG | [] |
| 3680 | 11 | APLNR | {'exs': 0, 'nonexs': 7} | True | 57235861 | CCAGCCCTAGTCAACCACAAGC | [] |
| 3681 | 11 | SSRP1 | {'exs': 7, 'nonexs': 1} | False | 57326406 | CGTTTCTACTCATCGGATCCTG | [] |
| 3682 | 11 | P2RX3 | {'exs': 0, 'nonexs': 7} | True | 57369996 | GAGGCCCTAGTGGCCTATGGA | [] |
| 3683 | 11 | SLC43A3 | {'exs': 0, 'nonexs': 7} | True | 57407791 | TCTGAACTATGCAATTGCAGAG | [] |
| 3684 | 11 | SLC43A1 | {'exs': 0, 'nonexs': 7} | True | 57485095 | GAAGTCTATGCGGTCACCTCAC | [] |
| 3685 | 11 | CLP1 | {'exs': 7, 'nonexs': 1} | False | 57661435 | TGATCTCTACTTCAGATCCATG | [] |
| 3686 | 11 | SELENOH | unavailable | False | 57742216 | CCCTACGACAGGTACTTCTTCC | ['TCAACTCTTCCACCACCTCTTG'] |
| 3687 | 11 | BTBD18 | {'exs': 0, 'nonexs': 5} | True | 57744133 | CTCCACTATGTTAGTATATCTG | ['TCTACCTCTGTTTCTGACTCTG'] |
| 3688 | 11 | CTNND1 | {'exs': 0, 'nonexs': 7} | True | 57816307 | TGGTGCTAAATCTTCTGCTGTG | [] |
| 3689 | 11 | OR9I1 | {'exs': 0, 'nonexs': 3} | True | 58119499 | ACTTAGATCTACATGCTCAGGG | [] |
| 3690 | 11 | OR10W1 | {'exs': 0, 'nonexs': 3} | True | 58266940 | TTTCCTAGCTGTTCTGGGAAAG | [] |
| 3691 | 11 | ZFP91 | {'exs': 3, 'nonexs': 4} | False | 58617705 | TGTCCACTAAGGTCCGGCAGAC | [] |
| 3692 | 11 | CNTF | {'exs': 0, 'nonexs': 7} | True | 58624521 | TAACTGCTACATTTTCTTGTTG | [] |
| 3693 | 11 | GLYATL1 | {'exs': 0, 'nonexs': 7} | True | 58956026 | TCATTGTCTAAAATGGAACTAG | [] |
| 3694 | 11 | GLYATL1B | unavailable | False | 59094785 | TCATTGTCTACAATGGAACAAG | [] |
| 3695 | 11 | FAM111B | {'exs': 0, 'nonexs': 7} | True | 59126301 | CTCTTTTCTAACATTCCATGGG | [] |
| 3696 | 11 | OR5A1 | {'exs': 0, 'nonexs': 3} | True | 59444115 | CTACGCATGACCTAAGAAAACG | ['CTTTCCAACACCTTCCACAGGG', 'GAAAACACTTTCTTCCTTTCCG'] |
| 3697 | 11 | OR4D10 | unavailable | False | 59478364 | TTTTTCTATTTTCTATCAGAAG | [] |
| 3698 | 11 | OR4D11 | {'exs': 1, 'nonexs': 3} | False | 59504510 | GTTTTCTATTCCCTTTCAGAAG | [] |
| 3699 | 11 | OR10V1 | {'exs': 0, 'nonexs': 3} | True | 59712915 | CCTACCTAGAATTTTCTCAATG | [] |
| 3700 | 11 | MRPL16 | {'exs': 7, 'nonexs': 1} | False | 59806346 | ACACTCACTACACACGTTTGGG | [] |
| 3701 | 11 | MS4A2 | {'exs': 0, 'nonexs': 7} | True | 60093972 | TCATCACTACAATTTCCTATTG | [] |
| 3702 | 11 | MS4A4E | {'exs': 1, 'nonexs': 3} | False | 60204423 | ACTCAACTAGAGTGAAGTTCTG | [] |
| 3703 | 11 | MS4A12 | {'exs': 0, 'nonexs': 7} | True | 60507123 | TTTTACTATTTAGGGGCATTAG | [] |

| | | | | | | | |
|---|---|---|---|---|---|---|---|
| 3704 | 11 | MS4A18 | unavailable | False | 60743993 | ATCAGCTATTTCTTGGTACGAG | [] |
| 3705 | 11 | MS4A10 | {'exs': 0, 'nonexs': 7} | True | 60799867 | CAGTGACTATCCATGTGTCCGC | [] |
| 3706 | 11 | PTGDR2 | unavailable | False | 60852534 | CGGGTTCTAACTCGAGGTGCTC | [] |
| 3707 | 11 | PRPF19 | {'exs': 8, 'nonexs': 1} | False | 60891165 | GGCCTACAGGCTGTAGAACTTC | [] |
| 3708 | 11 | SLC15A3 | {'exs': 2, 'nonexs': 7} | False | 60937549 | GGTGTCTACAATTCCTCTTCAG | [] |
| 3709 | 11 | CD6 | {'exs': 0, 'nonexs': 7} | True | 61019317 | CCCGGCCTAGGCTGCGCTGAT | [] |
| 3710 | 11 | VPS37C | {'exs': 0, 'nonexs': 7} | True | 61131819 | GTGTGTCTAATACCCAGGCCAC | [] |
| 3711 | 11 | DDB1 | {'exs': 7, 'nonexs': 1} | False | 61300135 | TTGGCTAATGGATCCGAGTTAG | [] |
| 3712 | 11 | TKFC | unavailable | False | 61346502 | TCACACACCCTAGCTCTGCAAC | [] |
| 3713 | 11 | TKFC | unavailable | False | 61348206 | TGACTAGGTTGAGGTCACCCAC | [] |
| 3714 | 11 | TMEM138 | {'exs': 0, 'nonexs': 7} | True | 61368155 | AGTCTTCTAGCAAGGGCAGGGC | [] |
| 3715 | 11 | TMEM138 | {'exs': 0, 'nonexs': 7} | True | 61375222 | AAAGCTATAAATAGCTTAAAAG | [] |
| 3716 | 11 | DAGLA | {'exs': 0, 'nonexs': 7} | True | 61739524 | TCACCTCTACCTCCTCAGGCAC | [] |
| 3717 | 11 | DAGLA | {'exs': 0, 'nonexs': 7} | True | 61744688 | GGGGTGCTAGCGTGCTGAGATC | [] |
| 3718 | 11 | RAB3IL1 | {'exs': 1, 'nonexs': 7} | True | 61898277 | CGCGCCCTAAGCCTCCTGGGG | [] |
| 3719 | 11 | SCGB1D4 | {'exs': 0, 'nonexs': 7} | True | 62296409 | ATTTTTTCACTATTTCCACCTG | [] |
| 3720 | 11 | SCGB1A1 | {'exs': 1, 'nonexs': 6} | True | 62423090 | TAAATGCTAATTACACAGTGAG | [] |
| 3721 | 11 | AHNAK | {'exs': 0, 'nonexs': 7} | True | 62433883 | CCTGGCCGCTTCTACAGTCCTC | [] |
| 3722 | 11 | AHNAK | {'exs': 0, 'nonexs': 7} | True | 62516743 | GGCCTGCTACTCTTTCTTTGTG | [] |
| 3723 | 11 | EML3 | {'exs': 0, 'nonexs': 7} | True | 62602301 | CGGGTCTAAACAGTGTGTGCAC | [] |
| 3724 | 11 | ROM1 | {'exs': 0, 'nonexs': 7} | True | 62614838 | CAGGCCTCTAGGCCTCAGATAC | [] |
| 3725 | 11 | B3GAT3 | {'exs': 1, 'nonexs': 7} | True | 62616185 | CTTGCGCTAAGCCAAGATTGCC | [] |
| 3726 | 11 | B3GAT3 | {'exs': 1, 'nonexs': 7} | True | 62616201 | TGCGCTACTGCACTCCAGCCTC | [] |
| 3727 | 11 | INTS5 | {'exs': 7, 'nonexs': 1} | False | 62647019 | GAAAAGGCTACGTCCCCTGTCC | [] |
| 3728 | 11 | LBHD1 | unavailable | False | 62662128 | GTGGTTCACTAGTCCTGGCTGC | [] |
| 3729 | 11 | STX5 | {'exs': 5, 'nonexs': 4} | False | 62807608 | GGCTCCTAGCACGTTCTCGTCT | [] |
| 3730 | 11 | SLC22A25 | {'exs': 0, 'nonexs': 7} | True | 63163823 | AGCACAGACCTATAGCACAGAG | [] |
| 3731 | 11 | SLC22A10 | {'exs': 0, 'nonexs': 7} | True | 63304704 | AAATCTATTCCTGAAGCTCTTG | [] |
| 3732 | 11 | PLAAT5 | unavailable | False | 63463530 | TTTATGCTATCCACTACAGCTG | [] |
| 3733 | 11 | ATL3 | {'exs': 2, 'nonexs': 6} | False | 63629318 | CTATTGAGCTTTTTTATCCATG | [] |
| 3734 | 11 | C11orf95 | {'exs': 0, 'nonexs': 1} | False | 63763417 | GGCCGCTACGCCCGACACACA | [] |
| 3735 | 11 | RCOR2 | {'exs': 0, 'nonexs': 7} | True | 63911772 | GCCAGAGCCCTAGTCCCTTCTC | [] |
| 3736 | 11 | OTUB1 | {'exs': 2, 'nonexs': 6} | False | 63997545 | GCCCTATTTGTAGAGGATATCG | [] |
| 3737 | 11 | DNAJC4 | {'exs': 1, 'nonexs': 7} | True | 64234112 | GCCCTAGCCGTTGTCGCTCCT | [] |
| 3738 | 11 | VEGFB | {'exs': 1, 'nonexs': 6} | True | 64237632 | TGAGCTCTAAGCCCCGCCCTT | [] |
| 3739 | 11 | GPR137 | {'exs': 1, 'nonexs': 7} | True | 64289364 | CATCCTAGGGGGACGGCACAA | [] |
| 3740 | 11 | KCNK4 | {'exs': 1, 'nonexs': 5} | False | 64299725 | CCCCTACACCGGCACGCCTTT | [] |
| 3741 | 11 | SLC22A11 | {'exs': 0, 'nonexs': 7} | True | 64571041 | AGGCACAATTTCTAGAGCGAGG | [] |
| 3742 | 11 | SLC22A12 | {'exs': 0, 'nonexs': 7} | True | 64601550 | CCAGGAGGCTAAAACTGTGTGC | [] |
| 3743 | 11 | SF1 | {'exs': 7, 'nonexs': 1} | False | 64765817 | CAAGTCTAGTTCTGTGGTGGAG | [] |
| 3744 | 11 | PPP2R5B | {'exs': 2, 'nonexs': 5} | False | 64933843 | TGTCTAGCTCTGACCCCCACTC | [] |
| 3745 | 11 | GPHA2 | {'exs': 0, 'nonexs': 7} | True | 64934772 | GATGGGCTAGTAGCGAGAGAGC | [] |
| 3746 | 11 | SAC3D1 | {'exs': 1, 'nonexs': 5} | False | 65044307 | AGCATCTACCGCCAAGGCCTTC | [] |
| 3747 | 11 | CDCA5 | {'exs': 6, 'nonexs': 2} | False | 65065168 | ATTAGACCTAGATGCTGTAAAG | [] |
| 3748 | 11 | CDCA5 | {'exs': 6, 'nonexs': 2} | False | 65066325 | CTGGGATCTAGGGAGGAGCTGC | [] |
| 3749 | 11 | VPS51 | unavailable | False | 65111586 | CTGCGCCTAGCCGCGCTCGCA | [] |
| 3750 | 11 | CDC42EP2 | {'exs': 0, 'nonexs': 7} | True | 65321530 | TCGTGTCCTATGTGGGGATCTC | [] |
| 3751 | 11 | KCNK7 | {'exs': 0, 'nonexs': 7} | True | 65593342 | TTCGACTACCCCTCCCACGCC | [] |
| 3752 | 11 | KCNK7 | {'exs': 0, 'nonexs': 7} | True | 65593419 | -1 | -1 |
| 3753 | 11 | PCNX3 | unavailable | False | 65635986 | GATGACTAAAGGCCACTGTCAG | [] |
| 3754 | 11 | SNX32 | {'exs': 1, 'nonexs': 7} | True | 65853334 | TACTCTAAGGCTCCCCCTTTAG | [] |
| 3755 | 11 | C11orf68 | {'exs': 0, 'nonexs': 7} | True | 65917458 | GCCCCCCTAGGTCAGTTCCAC | [] |
| 3756 | 11 | DRAP1 | {'exs': 1, 'nonexs': 7} | True | 65921434 | CTAGGAGTCGTAATCTTCTTCG | [] |
| 3757 | 11 | SF3B2 | {'exs': 7, 'nonexs': 1} | False | 66068744 | GACCTAAAACTTGAACTCCTTG | ['GAACTCCTTATATTTCTTGCTG'] |
| 3758 | 11 | KLC2 | {'exs': 0, 'nonexs': 7} | True | 66267187 | GAACCTAGGGAAGAGGGCCCTC | [] |
| 3759 | 11 | RAB1B | {'exs': 0, 'nonexs': 7} | True | 66276237 | CCCCTCCTAGCAACAGCCACC | [] |
| 3760 | 11 | RIN1 | {'exs': 0, 'nonexs': 7} | True | 66332275 | CAAGCTACTCCTCTGCTGCCC | [] |
| 3761 | 11 | SLC29A2 | {'exs': 1, 'nonexs': 6} | True | 66364106 | CAGACAGCTAGGGTTGGGCTG | [] |
| 3762 | 11 | SLC29A2 | {'exs': 1, 'nonexs': 6} | True | 66365422 | GCATCTACAGTTTACGTACTCG | [] |
| 3763 | 11 | PELI3 | {'exs': 0, 'nonexs': 7} | True | 66476166 | GAGCCTAATCCAGCGGGCCCT | [] |
| 3764 | 11 | ZDHHC24 | {'exs': 0, 'nonexs': 7} | True | 66526956 | ACAGCCTAGGAGGTACACGTTC | [] |
| 3765 | 11 | CCDC87 | {'exs': 0, 'nonexs': 7} | True | 66590465 | AGCTACTAAAGGCTGGCTGCTC | [] |
| 3766 | 11 | RBM14 | {'exs': 6, 'nonexs': 2} | False | 66626492 | AGCTCGGCTAAACGCCGGTCG | [] |
| 3767 | 11 | RBM14 | {'exs': 6, 'nonexs': 2} | False | 66626667 | ATGGCCCTACATGCGGCGCTG | [] |
| 3768 | 11 | SPTBN2 | {'exs': 0, 'nonexs': 5} | True | 66685870 | CAACTACTTGTTCTTCTTAAAG | [] |
| 3769 | 11 | SPTBN2 | {'exs': 0, 'nonexs': 5} | True | 66697844 | CAGGGCCTAATACAATGCTTTG | [] |
| 3770 | 11 | C11orf86 | {'exs': 1, 'nonexs': 7} | True | 66976250 | GTGTGGGCTACGGGGAGGCCG | [] |
| 3771 | 11 | SYT12 | {'exs': 1, 'nonexs': 7} | True | 67048756 | GGTTGCTAGTTTCGCCGGACAC | [] |
| 3772 | 11 | POLD4 | {'exs': 0, 'nonexs': 7} | True | 67351966 | CAAGAGAGCTAAGGGCAGGAGC | [] |
| 3773 | 11 | PPP1CA | {'exs': 3, 'nonexs': 5} | False | 67398534 | CGGGGGCTATTTCTTGGCTTTC | [] |
| 3774 | 11 | CARNS1 | {'exs': 1, 'nonexs': 6} | True | 67424600 | AGTGCTATTTGAAGTGAGACAG | [] |
| 3775 | 11 | RPS6KB2 | {'exs': 1, 'nonexs': 6} | True | 67435168 | CGGCTTCCTAGCGCCCTGGAC | [] |
| 3776 | 11 | PTPRCAP | {'exs': 0, 'nonexs': 7} | True | 67435732 | GCCTCTACAGTGCGGTGACATC | [] |
| 3777 | 11 | CORO1B | {'exs': 0, 'nonexs': 7} | True | 67438075 | GCTGTGGCCCTACGCATCCCC | [] |
| 3778 | 11 | NDUFV1 | {'exs': 3, 'nonexs': 5} | False | 67612457 | TGGTGGGCTAAGAGGCAGCCTC | [] |
| 3779 | 11 | TCIRG1 | {'exs': 0, 'nonexs': 7} | True | 68050818 | GGGCCCTAGTCATCTGTGGCA | [] |
| 3780 | 11 | TESMIN | unavailable | False | 68708307 | TTATACTCTACTCCATTTTCAG | ['AATCCCTTAGATTTAAACTCAG'] |
| 3781 | 11 | FGF3 | {'exs': 2, 'nonexs': 6} | False | 69810304 | AGGCCCAGCTAGTGCGCACTG | [] |
| 3782 | 11 | ANO1 | {'exs': 0, 'nonexs': 7} | True | 70188003 | GCATAGCTACAGGACGCCCCC | [] |
| 3783 | 11 | CTTN | {'exs': 1, 'nonexs': 6} | True | 70422551 | TGCTCTACTGAGCGCTAAAAAG | [] |
| 3784 | 11 | CTTN | {'exs': 1, 'nonexs': 6} | True | 70435161 | GCTGGGGGCCCTACTGCCGCA | [] |
| 3785 | 11 | RNF121 | {'exs': 0, 'nonexs': 7} | True | 71995525 | GAGGTCTACCTTCTCTTTGCAG | [] |
| 3786 | 11 | RNF121 | {'exs': 0, 'nonexs': 7} | True | 71996314 | TCATCACTATTCCAGGCCCAGC | [] |
| 3787 | 11 | ANAPC15 | {'exs': 5, 'nonexs': 2} | False | 72107350 | AAATTCTAACAGAGACCTTTCG | ['CTTTCATAAATCTGAAAGATAG'] |
| 3788 | 11 | ANAPC15 | {'exs': 5, 'nonexs': 2} | False | 72109880 | GTCTACCTAGATCATCCACTGG | [] |
| 3789 | 11 | PHOX2A | {'exs': 0, 'nonexs': 7} | True | 72239748 | GCCGGCAGCTAGAAGAGATTGC | [] |
| 3790 | 11 | CLPB | {'exs': 3, 'nonexs': 3} | False | 72293366 | AGGTGGCTGCTAGATGGTGTTC | [] |
| 3791 | 11 | CLPB | {'exs': 3, 'nonexs': 3} | False | 72311013 | CCCGACCTAGGGCATGAGTCC | [] |
| 3792 | 11 | ATG16L2 | {'exs': 1, 'nonexs': 7} | True | 72829389 | GGTCGTGGCCCTACTGCCAGA | [] |
| 3793 | 11 | P2RY2 | {'exs': 0, 'nonexs': 7} | True | 73235292 | GTTCTGCTCCTACAGCCGAATC | [] |
| 3794 | 11 | MRPL48 | {'exs': 6, 'nonexs': 2} | False | 73864369 | TACAGTAGCTACTTCAACTTGG | [] |
| 3795 | 11 | DNAJB13 | {'exs': 0, 'nonexs': 7} | True | 73961727 | CCCTGCTAGAGAATCTGTAGAG | [] |
| 3796 | 11 | PPME1 | {'exs': 4, 'nonexs': 4} | False | 74253509 | CAGCAGGTCACTAACAGCCAGC | [] |
| 3797 | 11 | PGM2L1 | {'exs': 0, 'nonexs': 7} | True | 74336651 | GTACCCCTAAACAGAACGCCAC | [] |
| 3798 | 11 | POLD3 | {'exs': 7, 'nonexs': 2} | False | 74668855 | TCCTACCCTACCTCTCCAACCC | [] |
| 3799 | 11 | RNF169 | {'exs': 0, 'nonexs': 7} | True | 74836729 | ATTAGGTGCTACTTGGCCCCGC | [] |
| 3800 | 11 | XRRA1 | {'exs': 0, 'nonexs': 7} | True | 74843199 | GCTCTAGCCTTGTGAGTCACTC | [] |
| 3801 | 11 | SPCS2 | {'exs': 3, 'nonexs': 4} | False | 74977042 | CTACTTTATTTTTCTTTCTATG | [] |
| 3802 | 11 | NEU3 | {'exs': 0, 'nonexs': 7} | True | 74994926 | CAGTACCTACGACACAGAATGG | [] |
| 3803 | 11 | NEU3 | {'exs': 0, 'nonexs': 7} | True | 75006648 | CTAAGGCAAAATTTTCATTTTG | [] |
| 3804 | 11 | ARRB1 | {'exs': 0, 'nonexs': 7} | True | 75266162 | GGCCCGTCTATCTGTTGTTGAC | [] |
| 3805 | 11 | GDPD5 | {'exs': 1, 'nonexs': 6} | True | 75435506 | TCTTCAGCTAACGCCCACTCCC | [] |
| 3806 | 11 | SERPINH1 | {'exs': 0, 'nonexs': 7} | True | 75572082 | TGAGGCCCTATAACTCGTCTCC | [] |
| 3807 | 11 | GVQW3 | unavailable | False | 76386019 | CTCTCCTACTGACATCCCACAC | [] |
| 3808 | 11 | GVQW3 | unavailable | False | 76394993 | AGGTCCTGGGCTAGTAGAGAAC | [] |
| 3809 | 11 | EMSY | unavailable | False | 76550145 | AAACACTAGGACCGTTCAGCAG | [] |
| 3810 | 11 | MYO7A | {'exs': 1, 'nonexs': 6} | True | 77212269 | CTAAGAGGCCTTCTCACACCCC | [] |
| 3811 | 11 | GDPD4 | {'exs': 1, 'nonexs': 7} | True | 77217276 | ATCTATCTATCTATCTTCCTCG | ['CCTCATTCTTACTTCCACTCTG'] |
| 3812 | 11 | AAMDC | unavailable | False | 77900695 | CTTACTATTTCATAAAATAAAG | [] |
| 3813 | 11 | INTS4 | {'exs': 7, 'nonexs': 1} | False | 77977010 | CCACCTGGCTACAGCTTCATGC | [] |
| 3814 | 11 | NDUFC2-KCTD14 | unavailable | False | 78038663 | CATACCTAATGGGTGGTGGCAC | [] |
| 3815 | 11 | THRSP | {'exs': 2, 'nonexs': 7} | False | 78064321 | AAGGTCTACCAAACTTGTCCCG | [] |
| 3816 | 11 | NARS2 | {'exs': 7, 'nonexs': 1} | False | 78436669 | CCAGCTATAAAAGGCATGAATG | [] |
| 3817 | 11 | ANKRD42 | {'exs': 0, 'nonexs': 6} | True | 83213662 | GATCTAAGGAGGTTCTAAATAG | [] |
| 3818 | 11 | ANKRD42 | {'exs': 0, 'nonexs': 6} | True | 83249000 | GACATTCTAATTTTTCTCTACG | [] |
| 3819 | 11 | CCDC90B | {'exs': 0, 'nonexs': 7} | True | 83261910 | TTAATACTACTTCCAGAATCTG | ['CAGAATCTATAAAATCCCAATG'] |
| 3820 | 11 | CREBZF | {'exs': 1, 'nonexs': 5} | False | 85663810 | TGACCCTACATTTTAAGAGAAG | [] |
| 3821 | 11 | CCDC89 | {'exs': 0, 'nonexs': 7} | True | 85685005 | ACTTCCCTATGGAGAGAGATGG | [] |
| 3822 | 11 | RAB38 | {'exs': 0, 'nonexs': 7} | True | 88113987 | TGCCTACTAGGATTTGGCACAG | [] |
| 3823 | 11 | CTSC | {'exs': 1, 'nonexs': 6} | True | 88294005 | ATACCCTACAATTTAGGAATTG | [] |
| 3824 | 11 | CTSC | {'exs': 1, 'nonexs': 6} | True | 88326432 | TGCTCTATTTAATAACTACAAG | [] |
| 3825 | 11 | NAALAD2 | {'exs': 0, 'nonexs': 7} | True | 90191746 | CCTTCTATAATACTTCTTTCAG | [] |
| 3826 | 11 | FAT3 | {'exs': 0, 'nonexs': 1} | False | 92891112 | GTGATGTCTACACTTGAGTCTG | [] |
| 3827 | 11 | MTNR1B | {'exs': 0, 'nonexs': 7} | True | 92982311 | CAGGCTAGAGAGCATCTGCCTC | [] |
| 3828 | 11 | SLC36A4 | {'exs': 0, 'nonexs': 7} | True | 93148536 | TTTTACTATTTCAAACCAGATG | [] |
| 3829 | 11 | SLC36A4 | {'exs': 0, 'nonexs': 7} | True | 93149814 | CAATGTCTAAACGAGGAATAAG | [] |
| 3830 | 11 | DEUP1 | unavailable | False | 93434957 | AGAGGATGCTAACATGATAGAG | [] |
| 3831 | 11 | CEP295 | unavailable | False | 93730276 | CTATTTCTAGAAAGTCAGCATG | [] |
| 3832 | 11 | VSTM5 | unavailable | False | 93820568 | GCTAACACTCAACATCTTCCAG | [] |
| 3833 | 11 | GPR83 | {'exs': 0, 'nonexs': 7} | True | 94380148 | CTTCTAACTCATCGTCACAATG | [] |
| 3834 | 11 | AMOTL1 | {'exs': 1, 'nonexs': 7} | True | 94857665 | ATAGGACTACTGTGGAAGTAAG | [] |
| 3835 | 11 | CWC15 | unavailable | False | 94963384 | CTATTTAATATATTTCTCCATG | [] |
| 3836 | 11 | YAP1 | {'exs': 2, 'nonexs': 6} | False | 1.02E+08 | GAGGGCTCTATAACCATGTAAG | [] |
| 3837 | 11 | MMP7 | {'exs': 1, 'nonexs': 6} | True | 1.03E+08 | TGAAGTTTCTATTTCTTTCTTG | [] |
| 3838 | 11 | MMP8 | {'exs': 0, 'nonexs': 7} | True | 1.03E+08 | TTACTCTCTATTCCTGGAAAGG | [] |
| 3839 | 11 | MMP10 | {'exs': 0, 'nonexs': 7} | True | 1.03E+08 | TCTCGCCTAGCAATGTAACCAG | [] |
| 3840 | 11 | MMP12 | unavailable | False | 1.03E+08 | CCATTTCTAACAACCAAACCAG | [] |
| 3841 | 11 | DCUN1D5 | {'exs': 0, 'nonexs': 7} | True | 1.03E+08 | TTCTTGCTATGATGTCTGACGG | [] |
| 3842 | 11 | DYNC2H1 | {'exs': 0, 'nonexs': 7} | True | 1.03E+08 | TAGATTCTACTGATTTTTTAGG | [] |
| 3843 | 11 | CARD16 | {'exs': 0, 'nonexs': 7} | True | 1.05E+08 | GAAGACTATGATAATTTATGAG | [] |
| 3844 | 11 | CARD16 | {'exs': 0, 'nonexs': 7} | True | 1.05E+08 | CTAAGCTAATTTCCAGGTATCG | [] |
| 3845 | 11 | CARD17 | {'exs': 0, 'nonexs': 7} | True | 1.05E+08 | ACAGCATTACCTAGGAAGGAAG | [] |
| 3846 | 11 | GRIA4 | {'exs': 1, 'nonexs': 7} | True | 1.06E+08 | CCCCTAAATTTTAATACTTTCG | [] |
| 3847 | 11 | CWF19L2 | {'exs': 6, 'nonexs': 2} | False | 1.07E+08 | TATTATTCTATTTTATTTTGTG | [] |
| 3848 | 11 | ALKBH8 | {'exs': 2, 'nonexs': 6} | False | 1.08E+08 | AACACTATTCACAACAGCCAAG | [] |
| 3849 | 11 | ELMOD1 | {'exs': 0, 'nonexs': 7} | True | 1.08E+08 | GGGCAACTACATGTTGATTAAG | ['TTGATTAAACCTTCCGAGGCAG'] |
| 3850 | 11 | SLC35F2 | {'exs': 0, 'nonexs': 7} | True | 1.08E+08 | CTCCAGCTACAAGACAGCAGAC | [] |
| 3851 | 11 | ACAT1 | {'exs': 0, 'nonexs': 7} | True | 1.08E+08 | TTGTCTACAGCTTCTGAATTAG | [] |
| 3852 | 11 | ATM | {'exs': 1, 'nonexs': 5} | False | 1.08E+08 | TTTTACTCTAGTAGCAAAAAAG | [] |
| 3853 | 11 | POU2AF1 | {'exs': 2, 'nonexs': 5} | False | 1.11E+08 | CACGCCTAAAAGCCTTCCACAC | [] |
| 3854 | 11 | BTG4 | {'exs': 1, 'nonexs': 6} | True | 1.11E+08 | TCCCAGCTACTTGAAGGTGGAC | [] |
| 3855 | 11 | BTG4 | {'exs': 1, 'nonexs': 6} | True | 1.11E+08 | GACTTACCTACCATCTCTGATG | [] |
| 3856 | 11 | BTG4 | {'exs': 1, 'nonexs': 6} | True | 1.11E+08 | CTTTTAAAAATCTACAGTTTTG | [] |
| 3857 | 11 | LAYN | {'exs': 0, 'nonexs': 7} | True | 1.12E+08 | TATATGTCCTAATAACCATATG | ['ATTTCATTTTCCACCCATCCAG'] |
| 3858 | 11 | SIK2 | {'exs': 2, 'nonexs': 5} | False | 1.12E+08 | GTGCTGAGACTAATTCACCAGG | [] |
| 3859 | 11 | ALG9 | unavailable | False | 1.12E+08 | CTAACCTCCACTTTTCTTCCTG | [] |
| 3860 | 11 | FDXACB1 | {'exs': 1, 'nonexs': 6} | True | 1.12E+08 | ACTACCGAGGTATAACATATAG | [] |
| 3861 | 11 | CRYAB | {'exs': 0, 'nonexs': 7} | True | 1.12E+08 | GGCATCTATTTCTTGGGGGCTC | [] |
| 3862 | 11 | TIMM8B | {'exs': 2, 'nonexs': 5} | False | 1.12E+08 | TGGCCTACTGCCCTCCTTTCTC | [] |
| 3863 | 11 | SDHD | {'exs': 6, 'nonexs': 2} | False | 1.12E+08 | GCCAGCTACTTGGGAGGCTGA | [] |
| 3864 | 11 | IL18 | {'exs': 0, 'nonexs': 7} | True | 1.12E+08 | TTAATAGCTAGTCTTCGTTTTG | [] |
| 3865 | 11 | PTS | {'exs': 0, 'nonexs': 7} | True | 1.12E+08 | CAATAGCTATTCTCCTTTATAG | ['TTTATAAACCACAATATTATTG'] |
| 3866 | 11 | NCAM1 | {'exs': 1, 'nonexs': 1} | False | 1.13E+08 | CAAGTTCCTAACAGAGCAAAAG | [] |
| 3867 | 11 | TTC12 | {'exs': 0, 'nonexs': 7} | True | 1.13E+08 | CCAGAACTAGGGATCCCAAACC | [] |
| 3868 | 11 | TTC12 | {'exs': 0, 'nonexs': 7} | True | 1.13E+08 | CGAGACCTATCTCAGCTCCTGC | [] |
| 3869 | 11 | ANKK1 | {'exs': 0, 'nonexs': 7} | True | 1.13E+08 | AGTTGTCTAAATTTCCATCTCG | [] |
| 3870 | 11 | ZW10 | {'exs': 3, 'nonexs': 4} | False | 1.14E+08 | GGAGCTATTTAATTTTAGCAAG | [] |
| 3871 | 11 | CLDN25 | {'exs': 0, 'nonexs': 7} | True | 1.14E+08 | AAGCCAGTCCTAGATGACCAGC | [] |
| 3872 | 11 | NXPE2 | unavailable | False | 1.15E+08 | CCTCTAACAAATGTAGTTTAAG | [] |
| 3873 | 11 | CADM1 | {'exs': 0, 'nonexs': 7} | True | 1.15E+08 | AAAGGCTGATCTAGATGAAGTG | ['GAAGTACTCTTTCTTTTCTTCG'] |
| 3874 | 11 | ZPR1 | unavailable | False | 1.17E+08 | ACTGCTACCGTTGCGGAGCCA | [] |
| 3875 | 11 | SIDT2 | {'exs': 0, 'nonexs': 7} | True | 1.17E+08 | CTGCTAGAAGACATAGATCTTG | [] |
| 3876 | 11 | TAGLN | {'exs': 0, 'nonexs': 7} | True | 1.17E+08 | CTCCGCTCTAACTGATGATCTG | [] |
| 3877 | 11 | DSCAML1 | {'exs': 0, 'nonexs': 7} | True | 1.17E+08 | CGGGCCCTACACCAGGGTGTA | [] |
| 3878 | 11 | SCN2B | {'exs': 0, 'nonexs': 7} | True | 1.18E+08 | CACTACTTGGCGCCATCATCCC | [] |
| 3879 | 11 | UBE4A | {'exs': 0, 'nonexs': 7} | True | 1.18E+08 | TTCACAGTATCTATTCAAGTTG | [] |
| 3880 | 11 | TMEM25 | {'exs': 0, 'nonexs': 7} | True | 1.19E+08 | TTCCCCTAAAGCTTCCACACTG | [] |
| 3881 | 11 | CXCR5 | {'exs': 0, 'nonexs': 7} | True | 1.19E+08 | GGGACCTAGAACGTGGTGAGAC | [] |
| 3882 | 11 | BCL9L | {'exs': 1, 'nonexs': 7} | True | 1.19E+08 | CGACTTCTAGAAGGGCAGGTTC | [] |
| 3883 | 11 | TRAPPC4 | {'exs': 5, 'nonexs': 3} | False | 1.19E+08 | GCCTATGACCCAGGTCCAAAAC | [] |
| 3884 | 11 | HINFP | {'exs': 7, 'nonexs': 1} | False | 1.19E+08 | TCAGCTCTACACTCTCGTAGCC | [] |
| 3885 | 11 | ABCG4 | {'exs': 1, 'nonexs': 7} | True | 1.19E+08 | AAGCCTCTATCTCTCTGACTTG | [] |
| 3886 | 11 | PDZD3 | {'exs': 0, 'nonexs': 7} | True | 1.19E+08 | CTCCAGCTATCCTAGGCCACTC | [] |
| 3887 | 11 | PDZD3 | {'exs': 0, 'nonexs': 7} | True | 1.19E+08 | GGTGCTCTACAGTAGATCCGAC | [] |
| 3888 | 11 | CCDC153 | {'exs': 0, 'nonexs': 4} | True | 1.19E+08 | GAACACTAGAGACTAGGGGCTC | [] |
| 3889 | 11 | CBL | {'exs': 1, 'nonexs': 7} | True | 1.19E+08 | GGTGTGCTAGGTAGCTACTATGC | [] |
| 3890 | 11 | MCAM | {'exs': 0, 'nonexs': 7} | True | 1.19E+08 | TCGGGGCTAATGCCTCAGATCC | [] |
| 3891 | 11 | C1QTNF5 | {'exs': 0, 'nonexs': 1} | False | 1.19E+08 | TGGGCACTAAGCAAAGACTGGC | [] |
| 3892 | 11 | USP2 | {'exs': 0, 'nonexs': 7} | True | 1.19E+08 | TGGCGCTACATTCGGGAGGGC | [] |
| 3893 | 11 | NECTIN1 | unavailable | False | 1.2E+08 | TGGGCTAGGGGCACTCTCCTC | [] |
| 3894 | 11 | NECTIN1 | unavailable | False | 1.2E+08 | CTACACGTACCACTCCTTCTTG | [] |
| 3895 | 11 | OAF | {'exs': 1, 'nonexs': 7} | True | 1.2E+08 | TCCCACCCTAGCCTGGGTAGG | [] |
| 3896 | 11 | TLCD5 | unavailable | False | 1.2E+08 | GCCTTGGCTAGTGTATTTTGAG | [] |
| 3897 | 11 | ARHGEF12 | {'exs': 1, 'nonexs': 6} | True | 1.2E+08 | GCGGCTCTAACTTTTATCTGAG | [] |
| 3898 | 11 | GRIK4 | {'exs': 0, 'nonexs': 7} | True | 1.21E+08 | ACAAAGCTACCCAAAGAGGCTG | [] |
| 3899 | 11 | GRIK4 | {'exs': 0, 'nonexs': 7} | True | 1.21E+08 | CCGGGACTACTCGGGCTCGCT | [] |
| 3900 | 11 | SC5D | unavailable | False | 1.21E+08 | GCAATAATCTATTCAGTCTTTG | [] |
| 3901 | 11 | CRTAM | {'exs': 0, 'nonexs': 7} | True | 1.23E+08 | GAGCACTACACAATACTCTCTG | [] |
| 3902 | 11 | JHY | unavailable | False | 1.23E+08 | TGTTGTCTATACGATGTGAAGG | [] |
| 3903 | 11 | SCN3B | {'exs': 0, 'nonexs': 7} | True | 1.24E+08 | TCCTGTTCTATTCCTCCACTGG | [] |
| 3904 | 11 | ZNF202 | {'exs': 0, 'nonexs': 7} | True | 1.24E+08 | CCTAGCTAGGAGGTCTTTTCTG | [] |
| 3905 | 11 | TMEM225 | {'exs': 0, 'nonexs': 7} | True | 1.24E+08 | CTTGCACTCTAGGAGAGAGAGC | [] |
| 3906 | 11 | OR4D5 | {'exs': 0, 'nonexs': 3} | True | 1.24E+08 | CCTCTGCTAATGTAAGGGTCTG | [] |
| 3907 | 11 | OR10S1 | {'exs': 0, 'nonexs': 3} | True | 1.24E+08 | CACAGACTATGGGGGTGGGCTC | [] |
| 3908 | 11 | OR10G8 | {'exs': 0, 'nonexs': 3} | True | 1.24E+08 | GTGTCTATTTGCTCTGAGAATG | [] |
| 3909 | 11 | OR10G7 | {'exs': 0, 'nonexs': 3} | True | 1.24E+08 | TTTCTTAACTATTCACCCTGAG | [] |
| 3910 | 11 | VWA5A | {'exs': 0, 'nonexs': 7} | True | 1.24E+08 | TTTCTTCCTACCTGTGTTTCTG | [] |
| 3911 | 11 | OR10D3 | {'exs': 0, 'nonexs': 3} | True | 1.24E+08 | CTTACTAACTCTCAGGAACATG | [] |
| 3912 | 11 | PANX3 | {'exs': 0, 'nonexs': 7} | True | 1.25E+08 | AACTATGGGTGTTCATCACATG | [] |
| 3913 | 11 | NRGN | {'exs': 0, 'nonexs': 7} | True | 1.25E+08 | CCTGGCCTAGTCTCCGCTGGG | [] |
| 3914 | 11 | PKNOX2 | {'exs': 0, 'nonexs': 7} | True | 1.25E+08 | CGACTACTCCAGGGAGTCACTC | [] |
| 3915 | 11 | ACRV1 | {'exs': 1, 'nonexs': 7} | True | 1.26E+08 | AGGCTTCTAGATCTTATTGCAG | [] |
| 3916 | 11 | PATE1 | {'exs': 0, 'nonexs': 7} | True | 1.26E+08 | TAACTTCTAAAGGTCTTCATTG | [] |
| 3917 | 11 | PATE2 | {'exs': 1, 'nonexs': 7} | True | 1.26E+08 | GTAGAACTAAACTCCCTCAGGG | [] |
| 3918 | 11 | FLI1 | {'exs': 2, 'nonexs': 5} | False | 1.29E+08 | CTTCTAGTAGTAGCTGCCTAAG | [] |
| 3919 | 11 | C11orf45 | {'exs': 0, 'nonexs': 7} | True | 1.29E+08 | ACAGGGCTAGTTAGGACCACCC | [] |
| 3920 | 11 | TP53AIP1 | {'exs': 0, 'nonexs': 7} | True | 1.29E+08 | AAACTAAGGGATCTGAAACCTG | [] |
| 3921 | 11 | ARHGAP32 | {'exs': 0, 'nonexs': 7} | True | 1.29E+08 | TTCCTACTATAACCTAGTAAAG | [] |
| 3922 | 11 | APLP2 | {'exs': 0, 'nonexs': 7} | True | 1.3E+08 | CTGCCACCTAAATCTGCATCTG | [] |
| 3923 | 11 | ST14 | {'exs': 0, 'nonexs': 7} | True | 1.3E+08 | CCCCGGCCCCTATACCCCAGT | [] |
| 3924 | 11 | ZBTB44 | {'exs': 0, 'nonexs': 7} | True | 1.3E+08 | CAAAACTACTCAATAGTTATAG | [] |
| 3925 | 11 | SNX19 | {'exs': 0, 'nonexs': 7} | True | 1.31E+08 | ACCAGCTAAGAGGAGACACCCC | ['GACACCCATCCTCTTAGAGTTG'] |
| 3926 | 11 | NTM | {'exs': 0, 'nonexs': 6} | True | 1.32E+08 | TGCTCTCTATTTTACGTCTGTG | [] |
| 3927 | 11 | NTM | {'exs': 0, 'nonexs': 6} | True | 1.32E+08 | GTTAATCTAAATTGACACGGAG | [] |
| 3928 | 11 | IGSF9B | {'exs': 0, 'nonexs': 1} | False | 1.34E+08 | CCTTACCTATTGAGCTGAGGAG | [] |
| 3929 | 11 | IGSF9B | {'exs': 0, 'nonexs': 1} | False | 1.34E+08 | AGCTGCTACAGTTTAGAGAGTG | [] |
| 3930 | 11 | VPS26B | {'exs': 0, 'nonexs': 7} | True | 1.34E+08 | GGGGGCCCTACTGCCTGCAGTT | [] |
| 3931 | 11 | ACAD8 | {'exs': 0, 'nonexs': 7} | True | 1.34E+08 | CTTCTCTAAGCGTCTCTTCCTG | [] |
| 3932 | 11 | ACAD8 | {'exs': 0, 'nonexs': 7} | True | 1.34E+08 | TGGGTTCTACTCCTGAAGCAGC | [] |
| 3933 | 11 | GLB1L3 | {'exs': 0, 'nonexs': 1} | False | 1.34E+08 | CTGATCCTACTAGGAAAGGCAG | [] |
| 3934 | 12 | IQSEC3 | {'exs': 0, 'nonexs': 7} | True | 171320 | TGCTAGCAGTAACTTTTAATTG | [] |
| 3935 | 12 | IQSEC3 | {'exs': 0, 'nonexs': 7} | True | 175032 | AGAGTCTACACCAGGCTCCTTC | [] |
| 3936 | 12 | SLC6A12 | {'exs': 1, 'nonexs': 4} | False | 191067 | CTGGCCACACCCTACAAATGGC | [] |
| 3937 | 12 | SLC6A12 | {'exs': 1, 'nonexs': 4} | False | 191870 | GCCCTAATCTCTGACCTTCTTG | [] |
| 3938 | 12 | SLC6A13 | {'exs': 1, 'nonexs': 7} | True | 220947 | CCCCCTAGCAGTGAGACTCTAC | [] |
| 3939 | 12 | SLC6A13 | {'exs': 1, 'nonexs': 7} | True | 259758 | ATACATCTATGGGACTCCCCAG | [] |
| 3940 | 12 | KDM5A | {'exs': 0, 'nonexs': 7} | True | 285655 | TCTGCTAACTGGTCTCTTTAAG | [] |
| 3941 | 12 | B4GALNT3 | {'exs': 0, 'nonexs': 7} | True | 561650 | CCGGCTACAGCGTCTTCATCTC | [] |
| 3942 | 12 | WNK1 | unavailable | False | 908791 | CTAGGTCTAAGTGGTCCGCAGC | [] |
| 3943 | 12 | ERC1 | {'exs': 0, 'nonexs': 5} | True | 1410414 | TCTGATTGGCTATGCCCATATG | ['TACCCTCCTCCTCATCCTACAG'] |
| 3944 | 12 | WNT5B | {'exs': 0, 'nonexs': 7} | True | 1649251 | CGGGCTATTTACAGATGTACTG | [] |
| 3945 | 12 | DCP1B | {'exs': 0, 'nonexs': 7} | True | 1993120 | ATTTCCTAAACATTAACATTTG | [] |
| 3946 | 12 | CACNA1C | {'exs': 0, 'nonexs': 7} | True | 2691198 | CGCCCACTACAGGCTGCTGAC | [] |
| 3947 | 12 | FKBP4 | {'exs': 1, 'nonexs': 6} | True | 2803257 | GCTATGCTTCTGTCTCCACCTC | [] |
| 3948 | 12 | ITFG2 | {'exs': 0, 'nonexs': 7} | True | 2824192 | GTACAGCTAGGTGGGATCCTGC | [] |
| 3949 | 12 | TEX52 | unavailable | False | 2849230 | AGCCCTTCTAGAAGTGTCCAGC | [] |
| 3950 | 12 | FOXM1 | {'exs': 1, 'nonexs': 6} | True | 2858637 | CAGGGCTCTACTGTAGCTCAGC | [] |
| 3951 | 12 | FOXM1 | {'exs': 1, 'nonexs': 6} | True | 2864344 | CTCGCTTGCTATGGCGGGCAA | [] |
| 3952 | 12 | PRMT8 | {'exs': 0, 'nonexs': 7} | True | 3593181 | CGTGTGCTAACGCATTTTGTAG | [] |
| 3953 | 12 | FGF23 | {'exs': 1, 'nonexs': 6} | True | 4370342 | GCGACCCTAGATGAACTTGGCC | [] |
| 3954 | 12 | NDUFA9 | {'exs': 3, 'nonexs': 5} | False | 4687107 | AGGAGGCACTAAATGTTGACGG | [] |
| 3955 | 12 | CD9 | {'exs': 1, 'nonexs': 7} | True | 6237827 | CTGACTCTAGACCATCTCGCGC | [] |
| 3956 | 12 | TAPBPL | {'exs': 0, 'nonexs': 7} | True | 6461175 | ACTACTCTTTCTTTCTACTCTG | [] |
| 3957 | 12 | MRPL51 | {'exs': 7, 'nonexs': 1} | False | 6492270 | TCTTCTATCGAAACTTCCCATG | [] |
| 3958 | 12 | NCAPD2 | {'exs': 8, 'nonexs': 1} | False | 6531411 | GACTTCCTAGGATCTGTGCCTC | [] |
| 3959 | 12 | IFFO1 | {'exs': 1, 'nonexs': 7} | True | 6540482 | AGGTCTCTATCTCATGGAGCTG | [] |
| 3960 | 12 | NOP2 | {'exs': 8, 'nonexs': 0} | False | 6556992 | CAACCATCTAAGATAGCAGCAG | [] |
| 3961 | 12 | NOP2 | {'exs': 8, 'nonexs': 0} | False | 6558027 | CCTAGCTACTTGGGAGTCTGAC | [] |
| 3962 | 12 | ING4 | {'exs': 0, 'nonexs': 7} | True | 6651194 | CCCTTATCTATTTCTTCTTCCG | [] |
| 3963 | 12 | ZNF384 | {'exs': 0, 'nonexs': 7} | True | 6667713 | GGATCTCTAAGAGCTGGCCAGC | [] |
| 3964 | 12 | GNB3 | {'exs': 1, 'nonexs': 7} | True | 6845968 | ACAGTCCTAGTTGGGAACCAAG | [] |
| 3965 | 12 | CDCA3 | {'exs': 2, 'nonexs': 6} | False | 6849042 | CAGGGCCTAGCTCTCCACCAA | [] |
| 3966 | 12 | LRRC23 | {'exs': 1, 'nonexs': 7} | True | 6913002 | ACTGCTGCTAGATCAATGACTG | [] |
| 3967 | 12 | LRRC23 | {'exs': 1, 'nonexs': 7} | True | 6914070 | CAGGCCTAGGCACCCCATTCTC | [] |
| 3968 | 12 | ATN1 | {'exs': 1, 'nonexs': 7} | True | 6941779 | CAGGTTCTACAGTGGCTTGTCC | [] |
| 3969 | 12 | RBP5 | {'exs': 1, 'nonexs': 7} | True | 7124120 | CTCCGGCTATCTGACCTTCCTC | [] |
| 3970 | 12 | ACSM4 | {'exs': 1, 'nonexs': 7} | True | 7328372 | ACTATCTTCCTCTCCATTCTTG | [] |
| 3971 | 12 | GDF3 | {'exs': 0, 'nonexs': 7} | True | 7689877 | CTACCCACACCCACATTCATCC | [] |
| 3972 | 12 | FOXJ2 | {'exs': 0, 'nonexs': 7} | True | 8052849 | GCACTAAGTGATCAAGTCCCAG | [] |
| 3973 | 12 | CLEC4D | {'exs': 0, 'nonexs': 7} | True | 8521270 | CTGAGTTTCTAGTTCAATGTTG | [] |
| 3974 | 12 | RIMKLB | {'exs': 0, 'nonexs': 6} | True | 8776786 | TTTTTCTACTGTTCCAGGGTAG | [] |
| 3975 | 12 | M6PR | {'exs': 0, 'nonexs': 7} | True | 8941817 | CAATCTACATTGGTAATAAATG | [] |
| 3976 | 12 | A2M | {'exs': 1, 'nonexs': 6} | True | 9068051 | GTTTTTCTATAATAATGTGCAG | [] |
| 3977 | 12 | PZP | {'exs': 1, 'nonexs': 6} | True | 9142820 | TATTTACCTCTAATTCCAGGGG | [] |
| 3978 | 12 | LOC105378 | unavailable | False | 9503088 | TCAAAACTAATCTTCACCACTG | [] |
| 3979 | 12 | CLEC2D | {'exs': 0, 'nonexs': 7} | True | 9694873 | AACATCTAGACATGTATATCTG | [] |
| 3980 | 12 | CLECL1 | {'exs': 0, 'nonexs': 7} | True | 9722625 | CACCTCTAAATGTTAAATCTCG | ['AAATCTCACCATAGCAGTAATG'] |
| 3981 | 12 | KLRF1 | {'exs': 0, 'nonexs': 7} | True | 9844525 | CTAATACTGACAAATCCATTTG | [] |
| 3982 | 12 | CLEC2A | {'exs': 0, 'nonexs': 7} | True | 9913565 | CTGCTCTATAAAAAATATTTAG | [] |
| 3983 | 12 | CLEC12B | unavailable | False | 10018480 | GCATACTAATCCAAATCCTCAG | [] |
| 3984 | 12 | CLEC7A | {'exs': 0, 'nonexs': 7} | True | 10123353 | TTTACTATAAATCCCTGTAATG | [] |
| 3985 | 12 | CLEC7A | {'exs': 0, 'nonexs': 7} | True | 10125086 | CCTACCTACTCAGGAAGCTGAC | [] |
| 3986 | 12 | OLR1 | {'exs': 0, 'nonexs': 7} | True | 10160015 | CGGACTCTAAATCTGCAGGTAG | [] |
| 3987 | 12 | KLRD1 | {'exs': 1, 'nonexs': 7} | True | 10317947 | TGAGTTCTCACTAGATCTGATG | [] |
| 3988 | 12 | KLRC4 | {'exs': 0, 'nonexs': 7} | True | 10407652 | TTATCTATAGAAAGCAGATCAG | [] |
| 3989 | 12 | KLRC2 | {'exs': 1, 'nonexs': 7} | True | 10431116 | TGCTTTACTTCTAAAGCTTATG | [] |
| 3990 | 12 | KLRC1 | {'exs': 1, 'nonexs': 7} | True | 10446550 | CGCTTTACCTCTAAAGCTTATG | [] |
| 3991 | 12 | EIF2S3B | unavailable | False | 10522694 | ATTTCCTAGAAGTTTGCTGCAG | [] |
| 3992 | 12 | YBX3 | unavailable | False | 10710758 | AATCCAGCTATCTTCTATTAAG | ['TCTATTAAACCATACATCAGAG'] |
| 3993 | 12 | TAS2R9 | {'exs': 0, 'nonexs': 7} | True | 10809136 | GGGGTCTCTATGGAACAAAAGG | [] |
| 3994 | 12 | TAS2R10 | {'exs': 0, 'nonexs': 7} | True | 10825345 | AAGGTGTCTATGTGACTCTGAG | [] |
| 3995 | 12 | TAS2R20 | {'exs': 0, 'nonexs': 6} | True | 10998945 | GTGAATCTATGGAGTTGACTGG | [] |
| 3996 | 12 | TAS2R31 | unavailable | False | 11030405 | CATGAATCTATGGAGATGAAGG | [] |
| 3997 | 12 | TAS2R46 | {'exs': 0, 'nonexs': 5} | True | 11061364 | TGTGAATCTATGAAGATGAAGG | [] |
| 3998 | 12 | TAS2R43 | {'exs': 0, 'nonexs': 5} | True | 11091299 | TCGTGAATCTATGGAGATGAAG | [] |
| 3999 | 12 | TAS2R30 | {'exs': 0, 'nonexs': 4} | True | 11133284 | TTCTGCTAGAAGACACACAATG | [] |
| 4000 | 12 | SMIM10L1 | unavailable | False | 11171562 | CATGGCTCTAAATATGTACCTG | [] |
| 4001 | 12 | TAS2R42 | {'exs': 0, 'nonexs': 7} | True | 11185992 | GTCTGTCTACAAAGGTAAAGGG | [] |
| 4002 | 12 | BORCS5 | unavailable | False | 12465775 | CAGCAGCTACAGCCTGAGCTC | [] |
| 4003 | 12 | GSG1 | {'exs': 0, 'nonexs': 7} | True | 13084900 | TCCTAACACTGCTCTTCCTCTG | ['TCCTCTACAGATGACCTAACTG'] |
| 4004 | 12 | GSG1 | {'exs': 0, 'nonexs': 7} | True | 13085147 | GCTCTTACTATGCTTGCACTTG | [] |
| 4005 | 12 | GSG1 | {'exs': 0, 'nonexs': 7} | True | 13103084 | ACCGCTACTGGGCCTTTACCAC | [] |
| 4006 | 12 | ATF7IP | {'exs': 0, 'nonexs': 4} | True | 14469419 | CTAATTAACTACAGCTCACAAG | [] |
| 4007 | 12 | ATF7IP | {'exs': 0, 'nonexs': 4} | True | 14481225 | AGAGAACTACATGTATCTTGGG | [] |
| 4008 | 12 | C12orf60 | {'exs': 0, 'nonexs': 7} | True | 14823672 | TTGCATCCCTACTTGTCACTGG | [] |
| 4009 | 12 | RERG | {'exs': 0, 'nonexs': 7} | True | 15109109 | GGCTGCCTAACTACTGATTTTG | [] |
| 4010 | 12 | PTPRO | {'exs': 1, 'nonexs': 6} | True | 15570597 | TCCTATGCACCTGGATACAATG | [] |
| 4011 | 12 | PTPRO | {'exs': 1, 'nonexs': 6} | True | 15595040 | CTAGGACTTGCTAACATTCTCG | [] |
| 4012 | 12 | RERGL | {'exs': 0, 'nonexs': 7} | True | 18081190 | -1 | -1 |
| 4013 | 12 | PIK3C2G | {'exs': 1, 'nonexs': 4} | False | 18715855 | CATGCCTAAATAAAGCACTGTG | [] |
| 4014 | 12 | PLEKHA5 | {'exs': 0, 'nonexs': 7} | True | 19290552 | ATAGTGCCTAGGCACTATGATG | [] |
| 4015 | 12 | PLEKHA5 | {'exs': 0, 'nonexs': 7} | True | 19369786 | AAGACTACACACACATGAAATG | [] |
| 4016 | 12 | PDE3A | {'exs': 1, 'nonexs': 7} | True | 20678475 | AAAAAGGTTCTACCTTAGCATG | ['CATACCTCCTTTAAGAAACTTG'] |
| 4017 | 12 | SLCO1C1 | {'exs': 0, 'nonexs': 6} | True | 20752527 | TTTCTAAAGTTGAGTTTCCTTG | [] |
| 4018 | 12 | IAPP | {'exs': 0, 'nonexs': 6} | True | 21378425 | CATTGTCCTCTAAAGGGGCAAG | [] |
| 4019 | 12 | LDHB | {'exs': 0, 'nonexs': 7} | True | 21635520 | TTTAAATTTCTACAGCCTAGAG | [] |
| 4020 | 12 | ABCC9 | {'exs': 0, 'nonexs': 7} | True | 21805173 | ATCATGATGGTCTACTTGTTGG | [] |
| 4021 | 12 | CMAS | {'exs': 0, 'nonexs': 7} | True | 22065310 | GCTAATTTCTATTTTTGGCATG | [] |
| 4022 | 12 | ST8SIA1 | {'exs': 0, 'nonexs': 7} | True | 22201551 | TTGTTCCTAGGAAGTGGGCTGC | [] |
| 4023 | 12 | ETNK1 | {'exs': 3, 'nonexs': 5} | False | 22664303 | ACACTAACTAGAAAAAACTAAG | [] |
| 4024 | 12 | LRMP | {'exs': 0, 'nonexs': 7} | True | 25108124 | GTTACTAATTTACGAAGTTCAG | [] |
| 4025 | 12 | LMNTD1 | unavailable | False | 25526932 | CAGCTATTTCAACATCTCCAAG | [] |
| 4026 | 12 | RASSF8 | {'exs': 2, 'nonexs': 5} | False | 26079072 | ATATCTAATCTTTACACTCCTG | [] |
| 4027 | 12 | TM7SF3 | {'exs': 0, 'nonexs': 7} | True | 26973964 | GGCATCTACAGAAGCAAAGGCC | [] |
| 4028 | 12 | MED21 | {'exs': 5, 'nonexs': 1} | False | 27028460 | TGATGCTATGAGTCTGGAAGAG | [] |
| 4029 | 12 | C12orf71 | {'exs': 0, 'nonexs': 7} | True | 27081173 | AAAAATCTGTCTATATGGGATG | [] |
| 4030 | 12 | ARNTL2 | {'exs': 0, 'nonexs': 7} | True | 27420531 | CAAAGGCTAGAGGGTCCACTGC | [] |
| 4031 | 12 | MANSC4 | unavailable | False | 27762737 | TACTATGAAGAGTTCTCCTTCG | ['GTTCTCCTTCATATGGTTACGG'] |
| 4032 | 12 | KLHL42 | unavailable | False | 27797289 | TGATCCTAAGAAACATGACAAG | [] |
| 4033 | 12 | TMTC1 | {'exs': 0, 'nonexs': 6} | True | 29506845 | GCTATGTTTGATCCTTTTCTCG | [] |
| 4034 | 12 | CAPRIN2 | {'exs': 0, 'nonexs': 7} | True | 30710543 | TCGGCTACTGAAACAGACAAAG | [] |
| 4035 | 12 | RESF1 | unavailable | False | 31987211 | GAACTCTAGATATTGCTAATCG | ['TATTGCTAATCATCTCCTTTAG'] |
| 4036 | 12 | BICD1 | {'exs': 0, 'nonexs': 7} | True | 32332506 | GCACAGTGCTAGATGGACGGTC | [] |
| 4037 | 12 | BICD1 | {'exs': 0, 'nonexs': 7} | True | 32367795 | TGGGGTCTATGGATCTGACATG | [] |
| 4038 | 12 | BICD1 | {'exs': 0, 'nonexs': 7} | True | 32377626 | ATGAAGACTAGGGGTGAGGAGC | [] |
| 4039 | 12 | PPHLN1 | {'exs': 1, 'nonexs': 7} | True | 42399005 | AAGACAACTAGACTCTCACCTG | [] |
| 4040 | 12 | PPHLN1 | {'exs': 1, 'nonexs': 7} | True | 42441508 | AAAAATCTAAAAGGCTCTCCG | ['TCTCCAAAATCTTGAGTGGAAG'] |

| | | | | | | | |
|---|---|---|---|---|---|---|---|
| 4041 | 12 | TMEM117 | {'exs': 1, 'nonexs': 7} | True | 44388671 | CTCCGAGTCTAGTTCGTAGGT | [] |
| 4042 | 12 | ANO6 | {'exs': 0, 'nonexs': 7} | True | 45439937 | TAAATCTAACTTAAGAAATTCG | [] |
| 4043 | 12 | SCAF11 | unavailable | False | 45919273 | CTGAAGCTACATCTTTTGGATG | [] |
| 4044 | 12 | SLC38A1 | {'exs': 0, 'nonexs': 7} | True | 46190126 | AACAGACTAATACAGATTACTG | ['TTACTAAATCCAGAATCCAGAG'] |
| 4045 | 12 | PCED1B | unavailable | False | 47230361 | GGTCCGTCTATTGAGGCCTTG | [] |
| 4046 | 12 | SLC48A1 | {'exs': 0, 'nonexs': 7} | True | 47780944 | GGCCCTACACCTTGACCTCTT | [] |
| 4047 | 12 | ASB8 | {'exs': 0, 'nonexs': 7} | True | 48149365 | CCGGCTATTCTAAAAGTAACAG | [] |
| 4048 | 12 | OR10AD1 | {'exs': 0, 'nonexs': 3} | True | 48202338 | CTTCCTGCTAGACAGACTGTG | [] |
| 4049 | 12 | H1-7 | unavailable | False | 48330058 | GCGTTGGCTACTGGATGGTCC | [] |
| 4050 | 12 | LMBR1L | {'exs': 0, 'nonexs': 7} | True | 49101189 | GCTCGGTCTAGAGTCCCAGGA | [] |
| 4051 | 12 | FAM186B | {'exs': 1, 'nonexs': 7} | True | 49583027 | GAGCTCGCTAGAGGACGTGCA | [] |
| 4052 | 12 | FAM186B | {'exs': 1, 'nonexs': 7} | True | 49584559 | ACAAGCTAAGCGGATTCATTTG | [] |
| 4053 | 12 | FAM186B | {'exs': 1, 'nonexs': 7} | True | 49587604 | CAGGAGGACTACACGTCCAGT | [] |
| 4054 | 12 | FMNL3 | {'exs': 0, 'nonexs': 7} | True | 49649561 | GCTAACAGTTTGACTCATCATG | [] |
| 4055 | 12 | SMARCD1 | {'exs': 3, 'nonexs': 5} | False | 50098999 | GGAGAGGCCCTATGTATTCCG | [] |
| 4056 | 12 | COX14 | unavailable | False | 50120216 | CCAGTTCTACATGATTCCTGAG | [] |
| 4057 | 12 | METTL7A | {'exs': 0, 'nonexs': 5} | True | 50930149 | GCCAGCTCACACTATTTCACAG | [] |
| 4058 | 12 | HIGD1C | {'exs': 0, 'nonexs': 7} | True | 50972531 | CTGTTCCTATTCGGCCATCTTG | [] |
| 4059 | 12 | LETMD1 | {'exs': 0, 'nonexs': 7} | True | 51058357 | ACTAGCATGCATTCACTCTTAG | [] |
| 4060 | 12 | TFCP2 | {'exs': 0, 'nonexs': 7} | True | 51095240 | ACTCCTACTTCAGTATGATATG | [] |
| 4061 | 12 | POU6F1 | {'exs': 0, 'nonexs': 7} | True | 51190246 | TGAGCCCTAAGGGATCTGAAAG | [] |
| 4062 | 12 | POU6F1 | {'exs': 0, 'nonexs': 7} | True | 51190578 | GATGTCTAGCTTCTCGAACCTG | [] |
| 4063 | 12 | DAZAP2 | {'exs': 1, 'nonexs': 7} | True | 51240143 | -1 | -1 |
| 4064 | 12 | GALNT6 | {'exs': 0, 'nonexs': 6} | True | 51354378 | GATCTGGGTCCTAGACAAAGAG | [] |
| 4065 | 12 | SLC4A8 | {'exs': 0, 'nonexs': 7} | True | 51497177 | CCTTCCTATGTTCTGTAATTTG | [] |
| 4066 | 12 | SCN8A | {'exs': 1, 'nonexs': 7} | True | 51807428 | CCTCTAACACTTGGATTCTCTG | [] |
| 4067 | 12 | ANKRD33 | {'exs': 0, 'nonexs': 7} | True | 51891304 | CCCCTACTTCTGTGCCATTCT | [] |
| 4068 | 12 | ACVRL1 | {'exs': 2, 'nonexs': 6} | False | 51920892 | CTGGGCTATTGAATCACTTTAG | [] |
| 4069 | 12 | GRASP | {'exs': 0, 'nonexs': 7} | True | 52015198 | -1 | -1 |
| 4070 | 12 | SMIM41 | unavailable | False | 52080060 | GGGCGCCTAGGAGTCGTCGTC | [] |
| 4071 | 12 | KRT80 | {'exs': 0, 'nonexs': 7} | True | 52171522 | AGGCAGCTACAGGGAACATAAG | [] |
| 4072 | 12 | KRT86 | {'exs': 1, 'nonexs': 6} | True | 52308584 | AGCCTCCTAGCACCTCTTACA | [] |
| 4073 | 12 | KRT81 | {'exs': 0, 'nonexs': 7} | True | 52286254 | GGGTGCCTAACATTTCCGGCA | [] |
| 4074 | 12 | KRT85 | {'exs': 0, 'nonexs': 7} | True | 52360852 | CTCTACTAGGCAAAGCGGACC | [] |
| 4075 | 12 | KRT82 | {'exs': 0, 'nonexs': 7} | True | 52394974 | ATCATGCTAATGCTTGTGGCTG | [] |
| 4076 | 12 | KRT71 | {'exs': 0, 'nonexs': 7} | True | 52544531 | TCTCTACCGACTGGTTTTCTTG | [] |
| 4077 | 12 | KRT74 | {'exs': 0, 'nonexs': 7} | True | 52566968 | GTCTAGCGGGTGGCTTTCCTT | [] |
| 4078 | 12 | KRT77 | {'exs': 0, 'nonexs': 7} | True | 52691164 | GAGGCCTCTACTCCAAGATCC | [] |
| 4079 | 12 | KRT4 | {'exs': 0, 'nonexs': 5} | True | 52807068 | TCCTCTATCGTCTCTTGTTCAG | [] |
| 4080 | 12 | KRT79 | {'exs': 1, 'nonexs': 7} | True | 52821871 | CAGCAGCTAATACCTCTGGCT | [] |
| 4081 | 12 | EIF4B | {'exs': 5, 'nonexs': 3} | False | 53040222 | TGTAGAGGTCTATTCGGCATAG | ['TTTTCATCTTCACCATCAACAG', 'ATAATCTTCTCCCTCATTTTCG'] |
| 4082 | 12 | SPRYD3 | {'exs': 1, 'nonexs': 7} | True | 53065831 | GAGGCCCTAGCCACTCAAGGG | [] |
| 4083 | 12 | SOAT2 | {'exs': 0, 'nonexs': 7} | True | 53124122 | CCCGACCTCTAGGTATGGCAG | [] |
| 4084 | 12 | ZNF740 | {'exs': 0, 'nonexs': 7} | True | 53187589 | CTTGCGCCTATAGAGAAAACTG | [] |
| 4085 | 12 | ZNF740 | {'exs': 0, 'nonexs': 7} | True | 53188667 | CCTCCTCTACCTGAATACAGTG | [] |
| 4086 | 12 | ITGB7 | {'exs': 1, 'nonexs': 7} | True | 53191698 | TAGACCTAAGAGGCCAGCCTT | [] |
| 4087 | 12 | C12orf10 | {'exs': 1, 'nonexs': 7} | True | 53307148 | ATTAGACTAGGAGATTTGTGGG | [] |
| 4088 | 12 | PCBP2 | {'exs': 5, 'nonexs': 2} | False | 53479441 | ATTGTTCTAGCTGCTCCCCATG | [] |
| 4089 | 12 | HOXC5 | {'exs': 1, 'nonexs': 7} | True | 54034491 | CTGCCTCTAAAGAGCCTCTTTG | [] |
| 4090 | 12 | GPR84 | {'exs': 0, 'nonexs': 7} | True | 54362660 | ACAGTTCTAATGGAGCCTATGG | [] |
| 4091 | 12 | PDE1B | {'exs': 2, 'nonexs': 5} | False | 54577327 | GCTAATCCAGATTCCCATTCTG | [] |
| 4092 | 12 | DCD | {'exs': 0, 'nonexs': 7} | True | 54644712 | ACAGCTATAGTACTGAGTCAAG | [] |
| 4093 | 12 | OR9C2 | {'exs': 0, 'nonexs': 3} | True | 55452151 | ACAGCTTCTACTTCTTTGAGAG | [] |
| 4094 | 12 | ITGA7 | {'exs': 0, 'nonexs': 7} | True | 55695057 | TGGGAACCTAGGCGGTGCCTG | [] |
| 4095 | 12 | BLOC1S1 | {'exs': 1, 'nonexs': 4} | False | 55719608 | GGGGCTAGGAAGGGGCAGACT | [] |
| 4096 | 12 | CD63 | {'exs': 1, 'nonexs': 7} | True | 55725560 | AGACCCCTACATCACCTCGTA | [] |
| 4097 | 12 | IKZF4 | {'exs': 1, 'nonexs': 7} | True | 56035330 | GAGGTTGCTAGCCCACCTTAT | [] |
| 4098 | 12 | ESYT1 | {'exs': 0, 'nonexs': 7} | True | 56143861 | CAGCTCCTAGGAGCTGCCCTT | [] |
| 4099 | 12 | NABP2 | unavailable | False | 56229212 | AATGTCATGCTATCTCTTGCTG | [] |
| 4100 | 12 | ANKRD52 | {'exs': 3, 'nonexs': 5} | False | 56243141 | AGGGGGCTACTCAGAGTAGCA | [] |
| 4101 | 12 | STAT2 | {'exs': 0, 'nonexs': 7} | True | 56343388 | GTGGTTCCTAGAAGTCAGAAGG | [] |
| 4102 | 12 | MIP | {'exs': 0, 'nonexs': 7} | True | 56451279 | GAGCTTCTACAGGGCCTGGGT | [] |
| 4103 | 12 | SPRYD4 | {'exs': 1, 'nonexs': 5} | False | 56469576 | GGACATACTAGAGGCCCTCGG | [] |
| 4104 | 12 | RBMS2 | {'exs': 0, 'nonexs': 7} | True | 56589260 | ATGGCTAAGTGTTAGTCCTTAG | [] |
| 4105 | 12 | PTGES3 | {'exs': 1, 'nonexs': 5} | False | 56666155 | TTTCTTACTATTTAATAGTATG | [] |
| 4106 | 12 | TAC3 | {'exs': 0, 'nonexs': 7} | True | 57012378 | CTTACCCTATTCTGCTCTCGG | [] |
| 4107 | 12 | NEMP1 | unavailable | False | 57059878 | CACTACCTAGGTTAGAAAGTTG | [] |
| 4108 | 12 | LRP1 | {'exs': 0, 'nonexs': 7} | True | 57212554 | GGCAGGGCCCTATGCCAAGGG | [] |
| 4109 | 12 | NXPH4 | {'exs': 0, 'nonexs': 7} | True | 57225746 | GGGGCGCTATCCGAAGTAGGG | [] |
| 4110 | 12 | NXPH4 | {'exs': 0, 'nonexs': 7} | True | 57225945 | AGGGGGCTACCCCAGCCTTGC | [] |
| 4111 | 12 | STAC3 | {'exs': 0, 'nonexs': 7} | True | 57243811 | CGCACGCCTAAATTTCCTCTAG | [] |
| 4112 | 12 | INHBC | {'exs': 0, 'nonexs': 7} | True | 57450021 | CATAGACTAACTGCACCCACAG | [] |
| 4113 | 12 | INHBE | {'exs': 0, 'nonexs': 7} | True | 57456847 | CTCTTGCTAGCTGCAGCCACA | [] |
| 4114 | 12 | MBD6 | {'exs': 0, 'nonexs': 7} | True | 57529233 | GGTATGGCTGCTATGGGGCCA | [] |
| 4115 | 12 | AGAP2 | {'exs': 1, 'nonexs': 7} | True | 57726551 | GGCAACTATACCAGCGCAACC | [] |
| 4116 | 12 | CYP27B1 | {'exs': 1, 'nonexs': 7} | True | 57763141 | GGGACTATCTGTCCAAAAACTG | [] |
| 4117 | 12 | METTL1 | {'exs': 6, 'nonexs': 2} | False | 57769423 | AATCACCCTAGACAGGAGGCT | [] |
| 4118 | 12 | TSFM | {'exs': 4, 'nonexs': 4} | False | 57790582 | GGAACCTATTCAGTTTCTGCTG | [] |
| 4119 | 12 | CTDSP2 | {'exs': 2, 'nonexs': 6} | False | 57823601 | GGGCAGGCTAAGGGGCCCGC | [] |
| 4120 | 12 | LRIG3 | {'exs': 0, 'nonexs': 7} | True | 58872571 | TCATTCAGTCTATGTGTCCAAG | [] |
| 4121 | 12 | USP15 | {'exs': 0, 'nonexs': 7} | True | 62335189 | TTCACTAGCAACTCTGCTCTAG | [] |
| 4122 | 12 | SRGAP1 | {'exs': 0, 'nonexs': 7} | True | 64091931 | TAGCACTATACCTGATGTCTTG | [] |
| 4123 | 12 | C12orf56 | {'exs': 0, 'nonexs': 7} | True | 64193841 | ACCTGCTAACCTTTGGCTCCA | [] |
| 4124 | 12 | C12orf56 | {'exs': 1, 'nonexs': 7} | True | 64267182 | GTTTCTATTCAACCAATTTCAG | [] |
| 4125 | 12 | C12orf56 | {'exs': 1, 'nonexs': 7} | True | 64274937 | ATCCAGCTAATAGCAAACTTTG | [] |
| 4126 | 12 | TBK1 | {'exs': 1, 'nonexs': 6} | True | 64501380 | AGAAAGCTAAAGACAGTCAACG | [] |
| 4127 | 12 | GNS | {'exs': 2, 'nonexs': 6} | False | 64716740 | AGGTCGCTACAGAAGATGTTTG | [] |
| 4128 | 12 | MSRB3 | {'exs': 0, 'nonexs': 7} | True | 65463321 | ATTACTCTAGAGCTCCGCTTTG | [] |
| 4129 | 12 | HMGA2 | {'exs': 0, 'nonexs': 6} | True | 65842613 | GAAAGGCTTCTAAGCTGCAATG | [] |
| 4130 | 12 | HMGA2 | {'exs': 0, 'nonexs': 6} | True | 65952429 | CTCTTCTAAGTGGCATTTTTAG | [] |
| 4131 | 12 | HMGA2 | {'exs': 0, 'nonexs': 6} | True | 65963291 | GCCCCCTAGTCCTCTTCGGCA | [] |
| 4132 | 12 | LLPH | {'exs': 0, 'nonexs': 7} | True | 66123839 | GTCTACCAGGCCAAACCCTTT | [] |
| 4133 | 12 | TMBIM4 | {'exs': 0, 'nonexs': 7} | True | 66138152 | CACTATAAAAAAAAACTATAG | [] |
| 4134 | 12 | HELB | {'exs': 0, 'nonexs': 7} | True | 66338101 | AAACTAAGTTTCTTGATTATCG | [] |
| 4135 | 12 | GRIP1 | {'exs': 0, 'nonexs': 7} | True | 66349018 | GCGTTGCTATAATGTATTAGTG | [] |
| 4136 | 12 | CAND1 | {'exs': 3, 'nonexs': 4} | False | 67312829 | AACATCTAACTAGTGTCCATTG | [] |
| 4137 | 12 | MDM1 | {'exs': 0, 'nonexs': 7} | True | 68303898 | TGACTATGTATCACTGTCCTAG | [] |
| 4138 | 12 | MDM1 | {'exs': 0, 'nonexs': 7} | True | 68326485 | AACTAATTCACAATAACTAAAG | [] |
| 4139 | 12 | NUP107 | {'exs': 7, 'nonexs': 1} | False | 68742461 | ATTAAACTATAACTGAATTTCG | ['TTTCATACCCTAATGGGTCAAG'] |
| 4140 | 12 | MDM2 | {'exs': 2, 'nonexs': 6} | False | 68839848 | TCAACTAGGGGAAATAAGTTAG | [] |
| 4141 | 12 | CPSF6 | {'exs': 6, 'nonexs': 3} | False | 69262558 | CTTCTAACGATGACGATATTCG | [] |
| 4142 | 12 | BEST3 | {'exs': 1, 'nonexs': 7} | True | 69684552 | AGCAGAGGAACTAAAAGCTCTG | [] |
| 4143 | 12 | RAB3IP | {'exs': 1, 'nonexs': 7} | True | 69789251 | ATATTCTACCGATTTCTGCACG | [] |
| 4144 | 12 | MYRFL | unavailable | False | 69925944 | CCTGCTAGAAGTGAGACTCCA | [] |
| 4145 | 12 | MYRFL | unavailable | False | 69949382 | GAACCCTATGAATACAATTGAG | [] |
| 4146 | 12 | KCNMB4 | {'exs': 0, 'nonexs': 7} | True | 70409547 | CGTGTCTGCTATTTTGTTGTTG | [] |
| 4147 | 12 | PTPRB | {'exs': 1, 'nonexs': 7} | True | 70524402 | GCCTCTATGCCTGGCCAGATT | [] |
| 4148 | 12 | ATXN7L3B | {'exs': 0, 'nonexs': 7} | True | 74539405 | TGCAGCTACTGGAATTCCGGA | [] |
| 4149 | 12 | KCNC2 | {'exs': 0, 'nonexs': 7} | True | 75192217 | TCCTTGTCTAATTTCCTTGCTG | [] |
| 4150 | 12 | CAPS2 | {'exs': 0, 'nonexs': 7} | True | 75278889 | AAAGACTAAATCCCCCATGGAG | [] |
| 4151 | 12 | OSBPL8 | {'exs': 0, 'nonexs': 7} | True | 76355888 | GAACTTCTACTTGAACATGAAG | [] |
| 4152 | 12 | ZDHHC17 | {'exs': 0, 'nonexs': 7} | True | 76850984 | AAGATGTCGCTACACCAGCTG | [] |
| 4153 | 12 | NAV3 | {'exs': 1, 'nonexs': 6} | True | 78210516 | CCCTCTAGAGGGTAGATTCAAG | [] |
| 4154 | 12 | PAWR | {'exs': 0, 'nonexs': 7} | True | 79592606 | GAATCCTCTACCTGGTCAGCT | [] |
| 4155 | 12 | LIN7A | {'exs': 1, 'nonexs': 7} | True | 80797648 | TTTACTATGTGCTGGGATTCTG | [] |
| 4156 | 12 | LIN7A | {'exs': 1, 'nonexs': 7} | True | 80811464 | TCACCTATGACATGTGGTTTTG | [] |
| 4157 | 12 | LOC10536 | unavailable | False | 80937462 | TCTCCCCGCTACGTTCCCGCT | [] |
| 4158 | 12 | PPFIA2 | {'exs': 0, 'nonexs': 6} | True | 81314680 | TAAACTAGTCAATTCTGGATTG | [] |
| 4159 | 12 | SLC6A15 | {'exs': 0, 'nonexs': 7} | True | 84861631 | CCCAGCTACAAATCAGATTCTG | [] |
| 4160 | 12 | SLC6A15 | {'exs': 0, 'nonexs': 7} | True | 84883744 | CTACCTAGAATGAAGTGTTATG | [] |
| 4161 | 12 | TSPAN19 | {'exs': 0, 'nonexs': 7} | True | 85017089 | TTTGCCTATTCTTTTACTCCTG | [] |
| 4162 | 12 | LRRIQ1 | {'exs': 0, 'nonexs': 7} | True | 85244940 | GATTTCTAAATTAATTTTGAAG | [] |
| 4163 | 12 | RASSF9 | {'exs': 0, 'nonexs': 7} | True | 85804701 | ATTGGAACTATGTTGACAACAG | [] |
| 4164 | 12 | MGAT4C | {'exs': 0, 'nonexs': 5} | True | 85979288 | AATTGGCTAAGAAGTCCAAATG | [] |
| 4165 | 12 | POC1B-GA | unavailable | False | 89522812 | TGCTCTATTTCTCAAAACTCCG | ['TCCAAATTTGATTTTTATCTAG'] |
| 4166 | 12 | GALNT4 | {'exs': 1, 'nonexs': 4} | False | 89522812 | TGCTCTATTTCTCAAAACTCCG | ['TCCAAATTTGATTTTTATCTAG'] |
| 4167 | 12 | ATP2B1 | {'exs': 1, 'nonexs': 7} | True | 89591269 | AAAGAACTACGAAATGCATTCG | ['ATGCATTCACCACTCGAATCTG'] |
| 4168 | 12 | CCER1 | unavailable | False | 90953521 | ATGTCTAACAGTTAAAGTCCTG | [] |
| 4169 | 12 | PLEKHG7 | {'exs': 1, 'nonexs': 7} | True | 92770194 | TTAGGTCCCTAAATTTCAGAGG | [] |
| 4170 | 12 | NUDT4 | {'exs': 3, 'nonexs': 5} | False | 93399378 | TCTCTCTATCTTACACTAGATG | [] |
| 4171 | 12 | SOCS2 | {'exs': 1, 'nonexs': 7} | True | 93583045 | TTCTAACCAGGATAAAAAACAG | [] |
| 4172 | 12 | CRADD | {'exs': 0, 'nonexs': 7} | True | 93790707 | CTTCTAGCACAGCTGTCTCTTG | [] |
| 4173 | 12 | LOC10536 | unavailable | False | 94650276 | AGTGCTAACCCACGCGCCCCC | [] |
| 4174 | 12 | FGD6 | {'exs': 0, 'nonexs': 7} | True | 95081519 | CCAATACTGCTACAATATTGTG | [] |
| 4175 | 12 | VEZT | {'exs': 4, 'nonexs': 3} | False | 95299115 | TTCCCCCTCTAATTCCATGCG | [] |
| 4176 | 12 | NTN4 | {'exs': 0, 'nonexs': 7} | True | 95659085 | TGCTACTTGCACTCTCTTTTTG | ['CTCTCTTTTTAAAATATCCATG'] |
| 4177 | 12 | SNRPF | {'exs': 7, 'nonexs': 1} | False | 95866070 | ATGCTATTCTCTCATTTCCCCG | ['CTTCCACACCTCTGATATAAAG', 'CCATCTTCTTCCTCTTCTTCCG'] |
| 4178 | 12 | CCDC38 | {'exs': 0, 'nonexs': 7} | True | 95881507 | ATCTAAAAGGAATTCCAAACTG | [] |
| 4179 | 12 | TMPO | {'exs': 0, 'nonexs': 7} | True | 98534341 | TTTACTAGTGTTTATTTCCACG | [] |
| 4180 | 12 | SLC25A3 | {'exs': 5, 'nonexs': 3} | False | 98601527 | CTAACTACTGAGTTAACCCAAG | [] |
| 4181 | 12 | IKBIP | {'exs': 0, 'nonexs': 7} | True | 98613584 | ATGTCAAACTAATTCATATCTG | [] |
| 4182 | 12 | APAF1 | {'exs': 1, 'nonexs': 6} | True | 98665078 | TGCATCCTAGCCTGAGGAACA | [] |
| 4183 | 12 | UHRF1BP | {'exs': 0, 'nonexs': 7} | True | 1E+08 | TGCCACTCCTGCTATTCAACTG | [] |
| 4184 | 12 | UHRF1BP | {'exs': 0, 'nonexs': 7} | True | 1E+08 | TGCTTTATGGTCTAAATGACTG | [] |
| 4185 | 12 | DEPDC4 | {'exs': 0, 'nonexs': 7} | True | 1E+08 | GACTTCTATCGAAGAGGGCTTG | [] |
| 4186 | 12 | DEPDC4 | {'exs': 0, 'nonexs': 7} | True | 1E+08 | AGTATCCTAAGCTGTAGATGTG | [] |
| 4187 | 12 | SYCP3 | {'exs': 0, 'nonexs': 7} | True | 1.02E+08 | TTATACCTATTTCAGCAAATAG | ['CAGCAAATAAAATATTTCATTG'] |
| 4188 | 12 | GNPTAB | {'exs': 0, 'nonexs': 7} | True | 1.02E+08 | AGATCTTCTATACTCTGATTCG | [] |
| 4189 | 12 | GNPTAB | {'exs': 0, 'nonexs': 7} | True | 1.02E+08 | TTCCACTAGTCAGGAGGCTGA | [] |
| 4190 | 12 | PMCH | {'exs': 0, 'nonexs': 7} | True | 1.02E+08 | AAAAGACTACTCACTGTCAAAG | ['TCAAAATCTCTCCTTCCTATAG'] |
| 4191 | 12 | IGF1 | {'exs': 2, 'nonexs': 5} | False | 1.02E+08 | TTCCTACATCCTGTAGTTCTTG | [] |
| 4192 | 12 | IGF1 | {'exs': 2, 'nonexs': 5} | False | 1.02E+08 | CTCTACTTGCGTTCTTCAAATG | [] |
| 4193 | 12 | C12orf42 | {'exs': 0, 'nonexs': 5} | True | 1.03E+08 | GTGCTACGTTAGTTCTATCATG | [] |
| 4194 | 12 | STAB2 | {'exs': 0, 'nonexs': 7} | True | 1.04E+08 | CTTTGCCTATTGCACCTCTGTG | [] |
| 4195 | 12 | NT5DC3 | {'exs': 0, 'nonexs': 7} | True | 1.04E+08 | GCCCTTGGCTACTTGGCCTGG | [] |
| 4196 | 12 | C12orf45 | {'exs': 3, 'nonexs': 5} | False | 1.05E+08 | GACTATTTCTTTTTTTTACTTG | [] |
| 4197 | 12 | ALDH1L2 | {'exs': 0, 'nonexs': 7} | True | 1.05E+08 | GTGTTGCTCTAATATTCCAGTG | [] |
| 4198 | 12 | WASHC4 | unavailable | False | 1.05E+08 | CACCTAGAGTTCAAGATCCATG | [] |
| 4199 | 12 | C12orf75 | {'exs': 0, 'nonexs': 6} | True | 1.05E+08 | ATTTTCTTCTAATCTTAAAGAG | [] |
| 4200 | 12 | NUAK1 | {'exs': 1, 'nonexs': 6} | True | 1.06E+08 | GGAATGCTAGTTGAGCTTGCTG | [] |
| 4201 | 12 | RIC8B | {'exs': 0, 'nonexs': 7} | True | 1.07E+08 | CTGTTGCTAACCACCATCCTG | [] |
| 4202 | 12 | CRY1 | {'exs': 1, 'nonexs': 7} | True | 1.07E+08 | TATTTACCTAATTAGTGCTCTG | [] |
| 4203 | 12 | ASCL4 | {'exs': 0, 'nonexs': 7} | True | 1.08E+08 | TCGCTAGCTGCCCCCCTCCTC | [] |
| 4204 | 12 | SELPLG | {'exs': 0, 'nonexs': 7} | True | 1.09E+08 | AGTGAGCTAAGGGAGGAAGCT | [] |
| 4205 | 12 | SVOP | {'exs': 0, 'nonexs': 1} | False | 1.09E+08 | GGTCACTATTCCTGAGAGCCA | [] |
| 4206 | 12 | MYO1H | {'exs': 0, 'nonexs': 4} | True | 1.09E+08 | TATTCTCATCTAAAACGGCTTG | ['ACGGCTTACTTCCCTCCCAGAG'] |
| 4207 | 12 | UBE3B | {'exs': 1, 'nonexs': 7} | True | 1.09E+08 | GGTTCCTCTAGCGAGACAACT | [] |
| 4208 | 12 | UBE3B | {'exs': 1, 'nonexs': 7} | True | 1.1E+08 | CAGGAGCTAGGAGAGTTCAAAG | [] |
| 4209 | 12 | MMAB | {'exs': 0, 'nonexs': 6} | True | 1.1E+08 | AAACACTAAGGGAATGAAGGAG | [] |
| 4210 | 12 | TRPV4 | {'exs': 0, 'nonexs': 7} | True | 1.1E+08 | CTGCAGTCCCTAGAGCGGGGC | [] |
| 4211 | 12 | GLTP | {'exs': 0, 'nonexs': 7} | True | 1.1E+08 | ATGCCTACACCTTGTAGTTAAG | [] |
| 4212 | 12 | ANKRD13a | {'exs': 0, 'nonexs': 7} | True | 1.1E+08 | AAGGTCTATTTGTCAGTGAGTG | [] |
| 4213 | 12 | ATP2A2 | {'exs': 9, 'nonexs': 0} | False | 1.1E+08 | ACAACTAAAGTTCTGAGCTAAG | [] |
| 4214 | 12 | ANAPC7 | {'exs': 3, 'nonexs': 4} | False | 1.1E+08 | GAGGCTAGTGCCTACCTTAGT | [] |
| 4215 | 12 | PPTC7 | {'exs': 0, 'nonexs': 7} | True | 1.11E+08 | CTCAGCTAGTCTGTATACTCAG | [] |
| 4216 | 12 | PPTC7 | {'exs': 0, 'nonexs': 7} | True | 1.11E+08 | AGCCACTATTGAAAGAAGGACG | [] |
| 4217 | 12 | TCTN1 | {'exs': 1, 'nonexs': 6} | True | 1.11E+08 | TTAGTCTATAATATCAACTCTG | [] |
| 4218 | 12 | HVCN1 | {'exs': 1, 'nonexs': 6} | True | 1.11E+08 | GGGTCTAGTTCACTTCACCAAG | [] |
| 4219 | 12 | PPP1CC | {'exs': 3, 'nonexs': 5} | False | 1.11E+08 | CTACCTAACTTGCCAGGGAAAG | [] |
| 4220 | 12 | PPP1CC | {'exs': 3, 'nonexs': 5} | False | 1.11E+08 | ACGACATCTATTTCTTTGCTTG | [] |
| 4221 | 12 | MYL2 | {'exs': 0, 'nonexs': 7} | True | 1.11E+08 | CTCCTAGTCCTTCTCTTCTCC | [] |
| 4222 | 12 | SH2B3 | {'exs': 0, 'nonexs': 7} | True | 1.11E+08 | TGAACTCTACGAGACCAAGTTG | [] |
| 4223 | 12 | ACAD10 | {'exs': 0, 'nonexs': 7} | True | 1.12E+08 | GGCTCTAAATGCGGTGCTTCA | [] |
| 4224 | 12 | TMEM116 | {'exs': 0, 'nonexs': 5} | True | 1.12E+08 | ATTTTCTAACTCCTCCACTGTG | [] |
| 4225 | 12 | RPH3A | {'exs': 0, 'nonexs': 7} | True | 1.13E+08 | ACTAGCCTAATCACTTGACACG | [] |
| 4226 | 12 | OAS1 | {'exs': 0, 'nonexs': 7} | True | 1.13E+08 | TGTTCACTAATTATTGGTATAG | [] |
| 4227 | 12 | OAS2 | {'exs': 0, 'nonexs': 7} | True | 1.13E+08 | ATGGCCTAGAGGTTGCACAGA | [] |
| 4228 | 12 | OAS2 | {'exs': 0, 'nonexs': 7} | True | 1.13E+08 | GATCTCTAGAAGTTTCTTTTAG | [] |
| 4229 | 12 | CFAP73 | unavailable | False | 1.13E+08 | CAGGGCTAGGAGGCAGGGGCT | [] |
| 4230 | 12 | TPCN1 | {'exs': 0, 'nonexs': 6} | True | 1.13E+08 | GCTGGGCTAGGTAACGGTCTG | [] |
| 4231 | 12 | TBX3 | {'exs': 0, 'nonexs': 7} | True | 1.15E+08 | ACGGGTCTACGGGGACGCGCT | [] |
| 4232 | 12 | RNFT2 | {'exs': 0, 'nonexs': 7} | True | 1.17E+08 | TCGGTCCTAGTACACCTGGAA | [] |
| 4233 | 12 | HRK | {'exs': 0, 'nonexs': 7} | True | 1.17E+08 | GCGTTCCTACAAGTTCCGCCT | [] |
| 4234 | 12 | FBXW8 | {'exs': 1, 'nonexs': 7} | True | 1.17E+08 | ACATCCCTAAACATGGTTATAG | [] |
| 4235 | 12 | KSR2 | {'exs': 0, 'nonexs': 1} | False | 1.18E+08 | GGCTAGACAGGCTAAAACAATG | [] |
| 4236 | 12 | RFC5 | {'exs': 8, 'nonexs': 1} | False | 1.18E+08 | AAGCAGGCTACCTTCTCCACA | [] |
| 4237 | 12 | RFC5 | {'exs': 8, 'nonexs': 1} | False | 1.18E+08 | TCTAGGCCTCTGCAACAATCAG | [] |
| 4238 | 12 | PEBP1 | {'exs': 0, 'nonexs': 7} | True | 1.18E+08 | TAACCCCCTACTTCCCAGACA | [] |
| 4239 | 12 | CCDC60 | {'exs': 0, 'nonexs': 7} | True | 1.2E+08 | CCCAGCCTACCTCAGGGCGCT | [] |
| 4240 | 12 | BICDL1 | unavailable | False | 1.2E+08 | GCCACTAGATCAGAAACCAAAG | [] |
| 4241 | 12 | PXN | {'exs': 2, 'nonexs': 6} | False | 1.2E+08 | GGGCACCTAGCAGAAGAGCTT | [] |
| 4242 | 12 | OASL | {'exs': 0, 'nonexs': 7} | True | 1.21E+08 | AAAACTAACTGGCTGGAAACAG | [] |
| 4243 | 12 | CAMKK2 | {'exs': 1, 'nonexs': 6} | True | 1.21E+08 | AGGCAGCTACTCGGGCTCCAT | [] |
| 4244 | 12 | CAMKK2 | {'exs': 1, 'nonexs': 6} | True | 1.21E+08 | TCTAGGTCTAAACACATCAAAG | [] |
| 4245 | 12 | ANAPC5 | {'exs': 8, 'nonexs': 0} | False | 1.21E+08 | TCTCTAGAGATGGTTTATCAAG | [] |
| 4246 | 12 | KDM2B | {'exs': 2, 'nonexs': 6} | False | 1.21E+08 | CCTTGGACTAACTCAGTTTTTG | [] |
| 4247 | 12 | ORAI1 | {'exs': 0, 'nonexs': 4} | True | 1.22E+08 | ATGGGCCTAGGCATAGTGGCT | [] |
| 4248 | 12 | SETD1B | {'exs': 1, 'nonexs': 3} | False | 1.22E+08 | GGGCCTAGTTGAGGGTCCCCC | [] |
| 4249 | 12 | BCL7A | {'exs': 1, 'nonexs': 6} | True | 1.22E+08 | CATCGTCTACATCTCTTCGGAG | [] |
| 4250 | 12 | MLXIP | {'exs': 0, 'nonexs': 3} | True | 1.22E+08 | ACATGCCAGCTAAGCAGCTAG | [] |
| 4251 | 12 | LRRC43 | {'exs': 1, 'nonexs': 7} | True | 1.22E+08 | CCTGTCCTACCCTTGACCCAA | [] |
| 4252 | 12 | LRRC43 | {'exs': 1, 'nonexs': 7} | True | 1.22E+08 | CACGCCCTACACGGCGAACAT | [] |
| 4253 | 12 | VPS33A | {'exs': 6, 'nonexs': 3} | False | 1.22E+08 | CCTAGAAAGGTTTTTCCATCAG | [] |
| 4254 | 12 | HIP1R | {'exs': 0, 'nonexs': 7} | True | 1.23E+08 | GGGGGGCCTAGTAGTTCACGA | [] |
| 4255 | 12 | PITPNM2 | {'exs': 0, 'nonexs': 6} | True | 1.23E+08 | GGTGCCCTACTTGGGGCCCGC | [] |
| 4256 | 12 | CDK2AP1 | {'exs': 1, 'nonexs': 6} | True | 1.23E+08 | GCAGCTAGGATCTGGCATTCC | [] |
| 4257 | 12 | RILPL2 | {'exs': 0, 'nonexs': 5} | True | 1.23E+08 | TGGATCTAGGTCTGTTTCCCC | [] |
| 4258 | 12 | SNRNP35 | {'exs': 6, 'nonexs': 2} | False | 1.23E+08 | -1 | -1 |
| 4259 | 12 | RILPL1 | {'exs': 0, 'nonexs': 7} | True | 1.23E+08 | AAGAAGCTAAACCTTTGGAAGG | [] |
| 4260 | 12 | TCTN2 | {'exs': 0, 'nonexs': 7} | True | 1.24E+08 | GGTTGTCTAACTATAGGCTTTG | [] |
| 4261 | 12 | SCARB1 | {'exs': 0, 'nonexs': 7} | True | 1.25E+08 | CCACCTACAGTTTTGCTTCCTG | [] |
| 4262 | 12 | LOC10798 | unavailable | False | 1.27E+08 | CCGCAGCTAGACTCCCTCCCG | [] |
| 4263 | 12 | TMEM132 | {'exs': 0, 'nonexs': 1} | False | 1.29E+08 | AGGGGCCTAAGCCTTATCTTTG | [] |
| 4264 | 12 | STX2 | {'exs': 0, 'nonexs': 7} | True | 1.31E+08 | GGTTGCTAGGACAATGTTGTTG | [] |
| 4265 | 12 | STX2 | {'exs': 0, 'nonexs': 7} | True | 1.31E+08 | TGGGTGCTACCTAAGACTGCA | [] |
| 4266 | 12 | SFSWAP | {'exs': 3, 'nonexs': 4} | False | 1.32E+08 | GAGCTAATTTCTTACGCTAAAG | [] |
| 4267 | 12 | PUS1 | {'exs': 1, 'nonexs': 7} | True | 1.32E+08 | GAAAAGCTATCTGTTCTCTCCG | [] |
| 4268 | 12 | EP400 | {'exs': 1, 'nonexs': 4} | False | 1.32E+08 | GCCCTGACTACTGGCACGGAG | [] |
| 4269 | 12 | DDX51 | {'exs': 7, 'nonexs': 1} | False | 1.32E+08 | CCCCAGCCTAGGCCGCCCTCT | [] |
| 4270 | 12 | DDX51 | {'exs': 7, 'nonexs': 1} | False | 1.32E+08 | AAGAGCCTAGGCAGAGAGAAG | [] |
| 4271 | 12 | FBRSL1 | {'exs': 0, 'nonexs': 7} | True | 1.33E+08 | GGCTACCGCGCCTCCACCTCC | [] |
| 4272 | 12 | POLE | {'exs': 8, 'nonexs': 1} | False | 1.33E+08 | GCTGGCTAATGGCCCAGCTGT | [] |
| 4273 | 12 | POLE | {'exs': 8, 'nonexs': 1} | False | 1.33E+08 | GCCTTCCCTACACCCACCCTC | [] |
| 4274 | 12 | ANKLE2 | {'exs': 5, 'nonexs': 3} | False | 1.33E+08 | GCCAAGCCTACAGGGCGGCAA | [] |
| 4275 | 12 | ZNF84 | {'exs': 0, 'nonexs': 4} | True | 1.33E+08 | TATTCCTAGGATTTTTTTACTG | [] |
| 4276 | 13 | ZMYM5 | {'exs': 1, 'nonexs': 7} | True | 19837780 | GCTTGATTCTACTGGAAGAATG | [] |
| 4277 | 13 | GJA3 | {'exs': 0, 'nonexs': 7} | True | 20141980 | ACCCGGGCACTAGATGGCCAA | [] |
| 4278 | 13 | IFT88 | {'exs': 0, 'nonexs': 7} | True | 20671033 | GCTAGCACTTCCTTCTCTTTTG | [] |
| 4279 | 13 | LATS2 | {'exs': 0, 'nonexs': 6} | True | 20974869 | CCCCCATCTACACGTACACAG | [] |
| 4280 | 13 | LOC10192 | unavailable | False | 24659196 | ACCCTTGCTACTGAGAGTGTG | [] |
| 4281 | 13 | MTMR6 | {'exs': 0, 'nonexs': 7} | True | 25249231 | TGAGTCTAACAAGTCATTCTTG | [] |
| 4282 | 13 | GPR12 | {'exs': 0, 'nonexs': 5} | True | 26758822 | GGGTGCTACACATCACTGGGC | [] |
| 4283 | 13 | LNX2 | {'exs': 0, 'nonexs': 7} | True | 27548334 | AAAATCTATACAAGGCTGCCAG | [] |
| 4284 | 13 | POLR1D | {'exs': 1, 'nonexs': 7} | True | 27623249 | TGCATAAAGGACTAGAATGTGG | [] |
| 4285 | 13 | GSX1 | {'exs': 0, 'nonexs': 7} | True | 27793947 | GCGCCTAGGGAGTGACCGTAA | [] |
| 4286 | 13 | URAD | unavailable | False | 27976105 | CGGCAGCTACAGCTTGGCGGG | [] |
| 4287 | 13 | FLT3 | {'exs': 1, 'nonexs': 6} | True | 28004051 | TCCTCTACGAATCTTCGACCTG | [] |
| 4288 | 13 | PAN3 | {'exs': 1, 'nonexs': 6} | True | 28292521 | AAATACTACAACTGACCATTTG | [] |
| 4289 | 13 | FLT1 | {'exs': 1, 'nonexs': 6} | True | 28303166 | CAAACTCTAGATGGGTGGGGT | [] |
| 4290 | 13 | FLT1 | {'exs': 1, 'nonexs': 6} | True | 28368577 | GATAGCTATGATGATGATGATG | [] |
| 4291 | 13 | HMGB1 | {'exs': 3, 'nonexs': 4} | False | 30461682 | ATAACTAGAACTTCAATAAATG | [] |
| 4292 | 13 | HSPH1 | {'exs': 0, 'nonexs': 7} | True | 31137317 | TATCTAGTCCAAGTCCATATTG | ['TTAACAGAATTTTTCTCATTAG'] |
| 4293 | 13 | RXFP2 | {'exs': 0, 'nonexs': 7} | True | 31802404 | AAAATGATTGCTAGGAAACTGG | [] |
| 4294 | 13 | N4BP2L1 | {'exs': 0, 'nonexs': 7} | True | 32402941 | TAGGCCTCTAATATCCATGGTG | [] |
| 4295 | 13 | N4BP2L1 | {'exs': 0, 'nonexs': 7} | True | 32403667 | CAACTACTGAATGGGGTCCATG | [] |
| 4296 | 13 | KL | {'exs': 0, 'nonexs': 7} | True | 33064185 | GAACTATTTGTAACTTCTTCTG | [] |
| 4297 | 13 | RFC3 | {'exs': 8, 'nonexs': 1} | False | 33973930 | TGAGTCTTCTACTGGGATACAG | [] |
| 4298 | 13 | MAB21L1 | {'exs': 1, 'nonexs': 7} | True | 35475058 | CATCCTCTAAAGTTTTTCCAAG | ['CCAAACTTTTCGGGTTGGTCAG'] |
| 4299 | 13 | ALG5 | {'exs': 1, 'nonexs': 6} | True | 36949941 | ACAACCTAATTCATTTTCCGAG | [] |
| 4300 | 13 | ELF1 | {'exs': 0, 'nonexs': 7} | True | 40933424 | ATATTAACTAAAAAGAGTTGGG | [] |
| 4301 | 13 | WBP4 | {'exs': 3, 'nonexs': 4} | False | 41082913 | GCAACTATTGATCATCACCTCG | [] |
| 4302 | 13 | AKAP11 | {'exs': 1, 'nonexs': 6} | True | 42319227 | TTTGCTCTATGCATTTTCCAAG | [] |
| 4303 | 13 | EPSTI1 | {'exs': 0, 'nonexs': 7} | True | 42897462 | AGCCTAGGACACAGCCTTTATG | [] |
| 4304 | 13 | TSC22D1 | {'exs': 0, 'nonexs': 7} | True | 44433964 | ACACTACATACACAAATATACG | ['AATAAGCAAAACAACCTTCATG'] |
| 4305 | 13 | TSC22D1 | {'exs': 0, 'nonexs': 7} | True | 44434625 | GGCAGCTATGCGGTTGGTCCT | [] |
| 4306 | 13 | NUFIP1 | {'exs': 3, 'nonexs': 5} | False | 44941205 | GCCTATACATCTTTACTTTTCG | [] |
| 4307 | 13 | TPT1 | {'exs': 7, 'nonexs': 1} | False | 45337475 | CAGGCTAAAGATGTATTCCCCG | ['TTCCCCAATTCAATCTGGGCCG'] |
| 4308 | 13 | LRRC63 | {'exs': 0, 'nonexs': 1} | False | 46276802 | TCATTGTACTATAATTCTTTTG | [] |
| 4309 | 13 | NUDT15 | {'exs': 1, 'nonexs': 7} | True | 48045798 | CACCTAGAGATGATTTCCTTTG | [] |
| 4310 | 13 | LPAR6 | {'exs': 0, 'nonexs': 7} | True | 48401331 | GGGCTCTAGAATTCTGGATATG | [] |
| 4311 | 13 | CDADC1 | {'exs': 0, 'nonexs': 7} | True | 49292781 | ACCCCTCTCTAACCTAGGAGA | [] |
| 4312 | 13 | SETDB2-P | unavailable | False | 49528564 | AATCCCTAATAAGACAGGGATG | [] |
| 4313 | 13 | PHF11 | {'exs': 0, 'nonexs': 7} | True | 49528564 | AATCCCTAATAAGACAGGGATG | [] |
| 4314 | 13 | EBPL | {'exs': 0, 'nonexs': 7} | True | 49661902 | ATAGCAATCTATTCACCCCAGG | [] |
| 4315 | 13 | RNASEH2 | {'exs': 4, 'nonexs': 3} | False | 50950069 | ATCTAGGAGATAAACATACCAG | [] |
| 4316 | 13 | RNASEH2 | {'exs': 4, 'nonexs': 3} | False | 50961477 | GGGTCTCCCACTAGCTCTTCT | [] |
| 4317 | 13 | FAM124A | {'exs': 1, 'nonexs': 7} | True | 51252272 | ACTGGTACTATAGTGTTTGAGG | [] |
| 4318 | 13 | SERPINE3 | unavailable | False | 51361996 | ATAAAAACAGCTAGAAAGAATG | [] |
| 4319 | 13 | DHRS12 | {'exs': 0, 'nonexs': 7} | True | 51768186 | TTGGCCTATTTAAATGTCTGAG | [] |
| 4320 | 13 | CCDC70 | {'exs': 0, 'nonexs': 7} | True | 51866079 | TGCTGGCTACGCCCTGCCTCG | [] |
| 4321 | 13 | UTP14C | {'exs': 0, 'nonexs': 7} | True | 52031104 | CTACAGTTTTTCTTCTTTATTG | [] |
| 4322 | 13 | NEK5 | {'exs': 0, 'nonexs': 7} | True | 52071569 | GTGACACTAACCAAAATAAAGG | [] |
| 4323 | 13 | THSD1 | {'exs': 0, 'nonexs': 7} | True | 52377410 | GATTCTCAGTCTAGATCACCAG | [] |
| 4324 | 13 | BORA | unavailable | False | 72755215 | CATTCTATGGACTGCTGCATTG | [] |
| 4325 | 13 | DIS3 | {'exs': 8, 'nonexs': 1} | False | 72759794 | TTGAATATAGCTATTTTCCAAG | [] |
| 4326 | 13 | PIBF1 | {'exs': 0, 'nonexs': 7} | True | 72939849 | TTTCAACTACTGAAAAAGAGAG | [] |
| 4327 | 13 | PIBF1 | {'exs': 0, 'nonexs': 7} | True | 73015918 | AACACTAGGTCTTCATCTTTTG | [] |
| 4328 | 13 | UCHL3 | {'exs': 1, 'nonexs': 7} | True | 75605811 | ACAAGCTATGCTGCAGAAAGAG | [] |
| 4329 | 13 | MYCBP2 | {'exs': 0, 'nonexs': 3} | True | 77045377 | TGCGTGTTCTAAAAAGTGTGGG | [] |
| 4330 | 13 | NDFIP2 | {'exs': 0, 'nonexs': 7} | True | 79551119 | TTCTTACCTCTACAATAAGAAG | [] |
| 4331 | 13 | SPRY2 | {'exs': 1, 'nonexs': 6} | True | 80336757 | ATGCTATGTTGGTTTTTCAAAG | [] |
| 4332 | 13 | SLITRK6 | {'exs': 0, 'nonexs': 7} | True | 85793982 | CCATCTATGTTTGCTGCTCCAG | [] |
| 4333 | 13 | DCT | {'exs': 0, 'nonexs': 7} | True | 94439897 | AGCACCCTAGGCTTCTTCTGT | [] |
| 4334 | 13 | ABCC4 | {'exs': 0, 'nonexs': 7} | True | 95096134 | GAGCCTAGGAGTTTGAGACCA | [] |
| 4335 | 13 | ABCC4 | {'exs': 0, 'nonexs': 7} | True | 95157284 | ACTGCTATCTCAGTACCAATAG | [] |
| 4336 | 13 | UGGT2 | {'exs': 0, 'nonexs': 7} | True | 95801789 | CAGTGCTAGAGTTCATCATGTG | [] |
| 4337 | 13 | UGGT2 | {'exs': 0, 'nonexs': 7} | True | 95885360 | GGTTCTTCTAGTTCCCCAAGG | [] |
| 4338 | 13 | RAP2A | {'exs': 1, 'nonexs': 6} | True | 97464461 | GGATGCTATTGTATGTTACATG | [] |
| 4339 | 13 | DOCK9 | {'exs': 0, 'nonexs': 7} | True | 98860339 | TTCACCAACTATTGGCTCCAAG | [] |
| 4340 | 13 | ZIC5 | {'exs': 0, 'nonexs': 7} | True | 99965376 | ATAAATTCTAATGTATCGTCCG | [] |
| 4341 | 13 | PCCA | {'exs': 0, 'nonexs': 7} | True | 1E+08 | CATACTAGACCGCACAACTATG | [] |
| 4342 | 13 | PCCA | {'exs': 0, 'nonexs': 7} | True | 1E+08 | TTGCTCTAATTTATAATAGCTG | [] |
| 4343 | 13 | NALCN | {'exs': 0, 'nonexs': 7} | True | 1.01E+08 | ATCCACCTAAATATCCAGAAGG | [] |
| 4344 | 13 | FGF14 | {'exs': 1, 'nonexs': 6} | True | 1.02E+08 | TGGCTATGTTGTCTTACTCTTG | [] |
| 4345 | 13 | FGF14 | {'exs': 1, 'nonexs': 6} | True | 1.02E+08 | TTAACCTAAATCATTAGAGGTG | [] |
| 4346 | 13 | TEX30 | unavailable | False | 1.03E+08 | TTTTAACTAATGACATTTCTTG | [] |
| 4347 | 13 | SLC10A2 | {'exs': 0, 'nonexs': 7} | True | 1.03E+08 | TTGATGTCTACTTTTCGTCAGG | [] |
| 4348 | 13 | EFNB2 | {'exs': 1, 'nonexs': 7} | True | 1.07E+08 | CTCAACCTAAGATGTTGATCTG | [] |
| 4349 | 13 | ING1 | {'exs': 4, 'nonexs': 5} | False | 1.11E+08 | TCCACAAACTACCTGTTGTAAG | [] |
| 4350 | 13 | ANKRD10 | {'exs': 0, 'nonexs': 7} | True | 1.11E+08 | AGCGTCTCTAGGAGCCGTGGT | [] |
| 4351 | 13 | ANKRD10 | {'exs': 0, 'nonexs': 7} | True | 1.11E+08 | GAGCTACTGTTTTTCATATCTG | [] |
| 4352 | 13 | ANKRD10 | {'exs': 0, 'nonexs': 7} | True | 1.11E+08 | TCAGCAGTTTCTATATGGAATG | [] |
| 4353 | 13 | LOC10798 | unavailable | False | 1.12E+08 | GATGGCTAATGACCATGTGCAG | [] |
| 4354 | 13 | SOX1 | {'exs': 1, 'nonexs': 6} | True | 1.12E+08 | AAGGCGCTAGATGTGCGTCAG | [] |
| 4355 | 13 | TUBGCP3 | {'exs': 6, 'nonexs': 3} | False | 1.13E+08 | CGCACCCTACAAACGAGTTCT | [] |
| 4356 | 13 | ATP11A | {'exs': 0, 'nonexs': 5} | True | 1.13E+08 | CGTGGAGCTAACTGCGCCTTG | [] |
| 4357 | 13 | MCF2L | {'exs': 0, 'nonexs': 7} | True | 1.13E+08 | GCCGCGCTACCCCTGCAGGTC | [] |
| 4358 | 13 | F7 | {'exs': 0, 'nonexs': 7} | True | 1.13E+08 | TGGGCTAGGGAAATGGGGCTC | [] |
| 4359 | 13 | LOC10272 | unavailable | False | 1.13E+08 | GTTCTAGTACAAAACTCCCCG | ['TCCCCAAAAAAGTCATGAGCTG'] |
| 4360 | 13 | PCID2 | {'exs': 8, 'nonexs': 1} | False | 1.13E+08 | GTCCTAATCTCCATTTCCTATG | [] |
| 4361 | 13 | LAMP1 | {'exs': 0, 'nonexs': 7} | True | 1.13E+08 | ACCAGGCTAGATAGTCTGGTAG | [] |
| 4362 | 13 | GRTP1 | {'exs': 1, 'nonexs': 7} | True | 1.13E+08 | CTGTGCCCTATGCCCACTGGT | [] |
| 4363 | 13 | DCUN1D2 | {'exs': 0, 'nonexs': 7} | True | 1.13E+08 | TTGCTGCCTAGAAAAGGCTGC | [] |
| 4364 | 13 | TMCO3 | {'exs': 0, 'nonexs': 7} | True | 1.14E+08 | CATGACTAAAACGCTTCAAAAG | [] |
| 4365 | 13 | GRK1 | {'exs': 0, 'nonexs': 3} | True | 1.14E+08 | GGGGCGTCACCTAGGAAACCA | [] |
| 4366 | 13 | GAS6 | {'exs': 0, 'nonexs': 7} | True | 1.14E+08 | TGGGGGCCTAGGCTGCGGCGG | [] |
| 4367 | 13 | C13orf46 | unavailable | False | 1.14E+08 | GGGCGCTAGGTGCTCTCCTTT | [] |
| 4368 | 13 | C13orf46 | unavailable | False | 1.14E+08 | CTGCAGCTAAAACCTAAACAGG | [] |
| 4369 | 13 | CFAP97D2 | unavailable | False | 1.14E+08 | GCTAATCCAGTTTCACTTCCTG | [] |
| 4370 | 13 | UPF3A | {'exs': 0, 'nonexs': 7} | True | 1.14E+08 | ATCACTCTACATTTCTGTCAAG | [] |
| 4371 | 14 | OR4K2 | {'exs': 0, 'nonexs': 3} | True | 19877211 | CACAAAAACTATGAAGGCATTG | [] |
| 4372 | 14 | OR4K1 | {'exs': 1, 'nonexs': 3} | False | 19930601 | TGATCCCTAGTTTTTCCAGGAG | [] |
| 4373 | 14 | OR4L1 | {'exs': 0, 'nonexs': 3} | True | 20060982 | AACATCTAGAAATTCTGTGCAG | [] |
| 4374 | 14 | OR4K17 | {'exs': 0, 'nonexs': 3} | True | 20118437 | TTAATCTAAGTATCTTCTCTAG | [] |
| 4375 | 14 | RNASE9 | {'exs': 0, 'nonexs': 7} | True | 20556451 | CTGCTAGGGCGATATGACAAAG | [] |
| 4376 | 14 | RNASE4 | {'exs': 1, 'nonexs': 6} | True | 20699814 | GTGGCATCTAACCGTCAAAGTG | [] |
| 4377 | 14 | EDDM3A | {'exs': 0, 'nonexs': 7} | True | 20749023 | AGACTTTCTAGTTGCTGATAGG | [] |

| | | | | | | | |
|---|---|---|---|---|---|---|---|
| 4378 | 14 | EDDM3B | {'ess': 0, 'nonest': 7} | True | 20770593 | AGACTTTCTAGTTGCCGATAGG | [] |
| 4379 | 14 | TPPP2 | {'ess': 0, 'nonest': 7} | True | 21032076 | TCTCTACTTGGTCTTCTTATCG | [] |
| 4380 | 14 | TPPP2 | {'ess': 0, 'nonest': 7} | True | 21036336 | CCCCTACCCTATGAGAAACAAG | [] |
| 4381 | 14 | RNASE7 | {'ess': 0, 'nonest': 7} | True | 21043462 | AACCTAAAGGACTCTGTCCAAG | [] |
| 4382 | 14 | ZNF219 | {'ess': 0, 'nonest': 7} | True | 21090535 | TGAGGGCCCACTACCGTTCTT | [] |
| 4383 | 14 | TOX4 | {'ess': 2, 'nonest': 4} | False | 21496605 | GACTATTTCACAAACACCACTG | [] |
| 4384 | 14 | METTL3 | {'ess': 6, 'nonest': 2} | False | 21498257 | AGTGCTTCTATAAATTCTTAGG | [] |
| 4385 | 14 | SALL2 | {'ess': 0, 'nonest': 7} | True | 21522007 | AGTTTGCTACTTCTTGTCTATG | [] |
| 4386 | 14 | LOC10537 | unavailable | False | 22559516 | GGCCTAGTTCACCGTGATCCT | [] |
| 4387 | 14 | OR6J1 | unavailable | False | 22633767 | TTCTAACACTGGAGCTTTACAG | [] |
| 4388 | 14 | PRMT5 | {'ess': 7, 'nonest': 1} | False | 22920903 | ACGCAGGGCTAGAGGCCAATG | [] |
| 4389 | 14 | ACIN1 | {'ess': 6, 'nonest': 1} | False | 23059147 | CCAGCTAGCGGCGCCCACCC | [] |
| 4390 | 14 | ACIN1 | {'ess': 6, 'nonest': 1} | False | 23068260 | CCCTAGATGGGGATCCCAACA | [] |
| 4391 | 14 | RNF212B | unavailable | False | 23270625 | AAGACTCTCTAGTGTAGTTGTG | [] |
| 4392 | 14 | RNF212B | unavailable | False | 23270638 | GGAAACTAGGAAGACTCTCTAG | [] |
| 4393 | 14 | RNF212B | unavailable | False | 23272890 | AAGATGATCTATCTAGAAGTGG | [] |
| 4394 | 14 | PABPN1 | {'ess': 7, 'nonest': 1} | False | 23324298 | GCCCATCTATCCTGACCTGTA | [] |
| 4395 | 14 | IL25 | {'ess': 1, 'nonest': 7} | True | 23375879 | GTCCGGCTAGCCCATCACACG | [] |
| 4396 | 14 | MYH7 | {'ess': 0, 'nonest': 6} | True | 23412853 | GCAAAGCTACTCCTCATTCAAG | [] |
| 4397 | 14 | THTPA | {'ess': 0, 'nonest': 6} | True | 23558839 | ACACCCCTAGCCCAGGCAGTG | [] |
| 4398 | 14 | PCK2 | {'ess': 0, 'nonest': 7} | True | 24100304 | CAGTCCTAGAAGATGGTTGGAG | [] |
| 4399 | 14 | DCAF11 | {'ess': 0, 'nonest': 7} | True | 24123308 | GGTTGGATCTACTGGGGTGAG | [] |
| 4400 | 14 | RNF31 | {'ess': 2, 'nonest': 5} | False | 24160572 | CCTCAGCTACTTCCGCCTGCG | [] |
| 4401 | 14 | IRF9 | {'ess': 0, 'nonest': 7} | True | 24166195 | CCAGGCTCTACACCAGGGACA | [] |
| 4402 | 14 | IPO4 | {'ess': 1, 'nonest': 7} | True | 24180441 | CTGCAGTCTAGGAGAGGCCCA | [] |
| 4403 | 14 | TSSK4 | {'ess': 0, 'nonest': 7} | True | 24207962 | CAGGTTCTAGAGCAGGGGAGA | [] |
| 4404 | 14 | GMPR2 | {'ess': 0, 'nonest': 7} | True | 24238777 | TCAGGTCTAGCACGCCTCACT | [] |
| 4405 | 14 | TGM1 | {'ess': 0, 'nonest': 7} | True | 24249312 | CGGGGCTAAGCTCCACCTCGA | [] |
| 4406 | 14 | NOP9 | unavailable | False | 24302989 | CGAAAACAACTAAAAATGCTAG | [] |
| 4407 | 14 | LTB4R | {'ess': 0, 'nonest': 7} | True | 24316709 | CCACCAGGCCTAGTTCAGTTC | [] |
| 4408 | 14 | SCFD1 | {'ess': 6, 'nonest': 3} | False | 30676145 | TATATCTATACATCTATATCTG | [] |
| 4409 | 14 | AKAP6 | {'ess': 0, 'nonest': 7} | True | 32824772 | TCTACCTATGCATATTTCTATG | [] |
| 4410 | 14 | PRORP | unavailable | False | 35273565 | GAATCTCTATGTCTTTTGGTGG | [] |
| 4411 | 14 | RALGAPA1 | {'ess': 1, 'nonest': 7} | True | 35664468 | AAGGTTCTAAAGCAGTTTTAAG | ['TTTAAACACAAAAAAGGATATG'] |
| 4412 | 14 | MIPOL1 | {'ess': 1, 'nonest': 7} | True | 37536093 | TCCCCCAAACTATTTCGCTATG | [] |
| 4413 | 14 | FOXA1 | {'ess': 1, 'nonest': 7} | True | 37591364 | CGGGAGCTAGGAAGTGTTTAG | [] |
| 4414 | 14 | CLEC14A | {'ess': 0, 'nonest': 7} | True | 38254549 | GTTTCCCTATGCATCACTAGAG | [] |
| 4415 | 14 | TRAPPC6B | {'ess': 0, 'nonest': 7} | True | 39150349 | TCAGTATGTTCTACAGCTTCTG | [] |
| 4416 | 14 | TRAPPC6B | {'ess': 0, 'nonest': 7} | True | 39151710 | AAATTACTAAAACATTTGTAAG | [] |
| 4417 | 14 | FBXO33 | {'ess': 0, 'nonest': 7} | True | 39399515 | AAAAAGCTAAATGTCTTCACTG | [] |
| 4418 | 14 | FSCB | {'ess': 0, 'nonest': 7} | True | 44504509 | AACCTGACTACAGTTCAGGAGG | [] |
| 4419 | 14 | KLHL28 | {'ess': 1, 'nonest': 7} | True | 44937754 | ATGGTGCTAGCATCTGGGAAG | [] |
| 4420 | 14 | MIS18BP1 | unavailable | False | 45204108 | ATTTTACTATGCAGAATCAGAG | [] |
| 4421 | 14 | MDGA2 | {'ess': 0, 'nonest': 7} | True | 46994680 | CACTATAAATGGATTTTAATAG | [] |
| 4422 | 14 | POLE2 | {'ess': 8, 'nonest': 1} | False | 49650252 | GATTAACTACATTCTTACAGGG | [] |
| 4423 | 14 | NEMF | unavailable | False | 49784635 | TTTCAGCTATTTCCTTTTTACG | [] |
| 4424 | 14 | VCPKMT | unavailable | False | 50111941 | CAAACCCTAGGTGTCACTGAG | [] |
| 4425 | 14 | DMAC2L | unavailable | False | 50324119 | TGATCATCTACCTTAAATGACG | [] |
| 4426 | 14 | DMAC2L | unavailable | False | 50324552 | GTGTTCTAGGCTCTAGAGACAG | [] |
| 4427 | 14 | NIN | {'ess': 0, 'nonest': 7} | True | 50723462 | TTTGGCTATGACCTCAAAGGAG | [] |
| 4428 | 14 | TMX1 | {'ess': 0, 'nonest': 7} | True | 51254518 | ATTTAACTAGGATTTATCTGTG | [] |
| 4429 | 14 | TXNDC16 | {'ess': 0, 'nonest': 7} | True | 52472901 | CTAAGGACTAGACGTTTCCAAG | [] |
| 4430 | 14 | CDKN3 | {'ess': 2, 'nonest': 5} | False | 54418272 | TCTCATTGTCCTAGGTGAGAGG | [] |
| 4431 | 14 | CNIH1 | unavailable | False | 54427813 | CTAAGAGCTCACCAAAACATAG | [] |
| 4432 | 14 | SAMD4A | unavailable | False | 54788943 | GCTTCTAGATGGTGGAGGTCC | [] |
| 4433 | 14 | GCH1 | {'ess': 1, 'nonest': 7} | True | 54843038 | CGTGGAAGCTATGGTTCTGCA | [] |
| 4434 | 14 | SOCS4 | {'ess': 0, 'nonest': 7} | True | 55044363 | TGTTACTAGCATTGCTGTTCTG | [] |
| 4435 | 14 | TMEM260 | unavailable | False | 56650112 | AGACTACAGGAAACCCCTTTTG | ['CCTTTTATCAGATTTTATGGAG'] |
| 4436 | 14 | SLC35F4 | {'ess': 0, 'nonest': 1} | False | 57564134 | CCTCTCTAAGCCAGTGGTATAG | [] |
| 4437 | 14 | SLC35F4 | {'ess': 0, 'nonest': 1} | False | 57581066 | GGGCTTTCTAATGCGTCTTCAG | [] |
| 4438 | 14 | ACTR10 | {'ess': 6, 'nonest': 3} | False | 58234550 | TCAAACTTCTATTTCTCAGTGG | [] |
| 4439 | 14 | TIMM9 | {'ess': 2, 'nonest': 5} | False | 58409033 | TTCTCTATCGTGGTTGGCCAAG | [] |
| 4440 | 14 | LRRC9 | {'ess': 0, 'nonest': 2} | False | 60000974 | CTTTTCCTAAGTTAATTGGCTG | [] |
| 4441 | 14 | PCNX4 | unavailable | False | 60134220 | TGAGCTCTAATACAAATGTATG | [] |
| 4442 | 14 | PRKCH | {'ess': 1, 'nonest': 6} | True | 61549830 | TAAGGCTATGGTTGCAATTCTG | [] |
| 4443 | 14 | HIF1A | {'ess': 0, 'nonest': 6} | True | 61746938 | TGCTAAATCTGAAAAACAAAAG | [] |
| 4444 | 14 | SYT16 | {'ess': 0, 'nonest': 7} | True | 62100706 | CAAACCTAGGATTCCAGCAAAG | [] |
| 4445 | 14 | SGPP1 | {'ess': 0, 'nonest': 7} | True | 63698596 | AAATTCTACTTAGGCAAACTAG | [] |
| 4446 | 14 | SYNE2 | {'ess': 0, 'nonest': 7} | True | 64225525 | ATGCCCTCTATGTGGGGGGTG | [] |
| 4447 | 14 | ESR2 | {'ess': 2, 'nonest': 6} | False | 64227889 | GGCACTAAGATAACTTCAAATG | [] |
| 4448 | 14 | ZBTB25 | {'ess': 1, 'nonest': 5} | False | 64486922 | AGCTGCTACTCAACTAGGATAG | [] |
| 4449 | 14 | SPTB | {'ess': 0, 'nonest': 7} | True | 64749305 | CACCTGCTACTTCTTTTTGGGG | [] |
| 4450 | 14 | SPTB | {'ess': 0, 'nonest': 7} | True | 64766656 | CTGAGCCTAGTAGGGGTGAGA | [] |
| 4451 | 14 | CHURC1-FNTB | unavailable | False | 65061311 | AGGTCCTCTAGTCGGTTGCAG | [] |
| 4452 | 14 | GPX2 | {'ess': 0, 'nonest': 7} | True | 64939487 | ACATCTATATGGCAACTTTAAG | [] |
| 4453 | 14 | FNTB | {'ess': 7, 'nonest': 0} | False | 65061311 | AGGTCCTCTAGTCGGTTGCAG | [] |
| 4454 | 14 | MAX | {'ess': 6, 'nonest': 2} | False | 65077476 | CATGGGCCTAGCCCATAGGCT | [] |
| 4455 | 14 | CCDC196 | unavailable | False | 66491000 | AGAGAGCTAAAGGAATAAGAGG | [] |
| 4456 | 14 | CCDC196 | unavailable | False | 66495668 | GTGCCACCTACTGGGCTAAAT | [] |
| 4457 | 14 | CCDC196 | unavailable | False | 66498471 | AAAGCCTGCTAATTCTTCAGTG | [] |
| 4458 | 14 | GPHN | {'ess': 0, 'nonest': 6} | True | 67735239 | AGCAGACTAGATGGTCCTTTTG | [] |
| 4459 | 14 | ARG2 | {'ess': 2, 'nonest': 6} | False | 67650919 | GTCTCCTAAATTCTCACACGTG | [] |
| 4460 | 14 | RDH12 | {'ess': 0, 'nonest': 5} | True | 67733847 | TTCCACCAGCTACTCCCACCG | [] |
| 4461 | 14 | RAD51B | unavailable | False | 68477563 | GTATCCCTATGGCTTCTCTTGG | [] |
| 4462 | 14 | RAD51B | unavailable | False | 68497139 | CTGTCTCTAGGAATTTCCATAG | [] |
| 4463 | 14 | SLC39A9 | {'ess': 1, 'nonest': 7} | True | 69461701 | TTAGTTCTAATCCTTGAAAAAG | [] |
| 4464 | 14 | SLC10A1 | {'ess': 0, 'nonest': 7} | True | 69776281 | GAAGGGCTAGGCTGTGCAAGG | [] |
| 4465 | 14 | MAP3K9 | {'ess': 2, 'nonest': 6} | False | 70730379 | CGTGCTAAGACCAGAACTCCT | [] |
| 4466 | 14 | LOC10537 | unavailable | False | 71190250 | AGCTGCCTAGTGCTGTCCAGT | [] |
| 4467 | 14 | SIPA1L1 | {'ess': 0, 'nonest': 7} | True | 71739160 | CCTTCCCTAGCTCATGTCTATG | [] |
| 4468 | 14 | DPF3 | {'ess': 0, 'nonest': 7} | True | 72619296 | GCCCTAGGCCTGGCAGCCAAA | [] |
| 4469 | 14 | PSEN1 | {'ess': 1, 'nonest': 7} | True | 73219288 | AAATATGCTAGATATAAAATTG | [] |
| 4470 | 14 | PAPLN | {'ess': 0, 'nonest': 7} | True | 73272663 | TTCATCCCTACTGCCAGATGG | [] |
| 4471 | 14 | RIOX1 | unavailable | False | 73492942 | AGAAACTAATTTAGGGCTAGAG | [] |
| 4472 | 14 | BBOF1 | unavailable | False | 74078239 | GGTTTGCTAGTAGGAAGTTGAG | [] |
| 4473 | 14 | ALDH6A1 | {'ess': 0, 'nonest': 7} | True | 74060641 | TGTTTCTAACGGCCCATGGTAG | [] |
| 4474 | 14 | LIN52 | {'ess': 6, 'nonest': 2} | False | 74198976 | CAAGCAGCTGCTACTTCTTGG | [] |
| 4475 | 14 | VSX2 | {'ess': 0, 'nonest': 7} | True | 74260918 | TGACCTAAGCCATGTCCTCCA | [] |
| 4476 | 14 | SYNDIG1L | unavailable | False | 74407534 | GCTACTAGCCATGACCGTTCT | [] |
| 4477 | 14 | LTBP2 | {'ess': 3, 'nonest': 6} | False | 74500883 | TGACTGCTACTCCTTGGCAGT | [] |
| 4478 | 14 | PROX2 | {'ess': 0, 'nonest': 7} | True | 74855131 | AAACAGCTACTGGGGATAGCTG | [] |
| 4479 | 14 | DLST | {'ess': 7, 'nonest': 1} | False | 74902329 | TTCCTCCTAAAGATCCAGGAGG | [] |
| 4480 | 14 | RPS6KL1 | {'ess': 1, 'nonest': 4} | False | 74904840 | GGGTAGCTAACCTAGACTGCA | [] |
| 4481 | 14 | ZC2HC1C | unavailable | False | 75073619 | AGTACTCTAGAGGAATGACAGG | [] |
| 4482 | 14 | ZC2HC1C | unavailable | False | 75077563 | TTCATGCTTCTATCTCCAGTTG | [] |
| 4483 | 14 | NEK9 | {'ess': 1, 'nonest': 6} | True | 75084563 | AGACTAGAGGCTGGGTCTACA | [] |
| 4484 | 14 | GPATCH2L | unavailable | False | 76178143 | TGAACCTACAAAGAGGAATTAG | [] |
| 4485 | 14 | GPATCH2L | unavailable | False | 76201850 | TGCTCTCTAGCTGTATCCAGG | [] |
| 4486 | 14 | LRRC74A | unavailable | False | 76870148 | TTGTTGCTATGGCTTCATCGTG | [] |
| 4487 | 14 | LRRC74A | unavailable | False | 76870165 | TCTTTCTAGACCAGACTTGTTG | [] |
| 4488 | 14 | CIPC | unavailable | False | 77114317 | CTCTGTGCTATATGGCTGGGT | [] |
| 4489 | 14 | ZDHHC22 | {'ess': 0, 'nonest': 7} | True | 77133682 | GTGTCTACTACTTATCCTGCTG | [] |
| 4490 | 14 | GSTZ1 | {'ess': 0, 'nonest': 7} | True | 77331194 | GGGAGCTAGGCCCTCAGCTCA | [] |
| 4491 | 14 | SAMD15 | {'ess': 0, 'nonest': 7} | True | 77391243 | TTCCCCTATGGTTCAGTGCAAG | [] |
| 4492 | 14 | AHSA1 | {'ess': 0, 'nonest': 7} | True | 77469248 | CTGGCCCTAAAATAAGCGTGCC | [] |
| 4493 | 14 | ISM2 | {'ess': 0, 'nonest': 7} | True | 77475594 | CCCCGTCACTAGTACTCCTTG | [] |
| 4494 | 14 | SNW1 | {'ess': 7, 'nonest': 3} | False | 77718087 | CTATTCCTTCCTCCTCTTCTTG | [] |
| 4495 | 14 | SEL1L | {'ess': 4, 'nonest': 4} | False | 81499443 | TTCCACTAAAGTCTACTTACCG | ['CTACTTACCAAAACCATGTGGG'] |
| 4496 | 14 | GPR65 | {'ess': 0, 'nonest': 7} | True | 88011860 | TGGTTCTACTCAAGGACCTCTG | ['TCTAATTCCATAGTATCTTTTG'] |
| 4497 | 14 | ZC3H14 | {'ess': 0, 'nonest': 7} | True | 88611750 | ACTGGGTGCTATTCGCTGAATG | [] |
| 4498 | 14 | EML5 | {'ess': 0, 'nonest': 7} | True | 88616044 | TGTTGACTAGCTGATTTCATAG | ['TCATAAACCAAAGCTGTAGGAG'] |
| 4499 | 14 | KCNK13 | {'ess': 0, 'nonest': 7} | True | 90185002 | TGGCTTCTACCTGTCCCCACT | [] |
| 4500 | 14 | NRDE2 | unavailable | False | 90275269 | AGGACCTGCTAGGAGTACTGC | [] |
| 4501 | 14 | NRDE2 | unavailable | False | 90278335 | CTGCTCTCTAATCCTCCAGCA | [] |
| 4502 | 14 | RPS6KA5 | {'ess': 1, 'nonest': 6} | True | 90872073 | CATGCCTAAGCTACTGAGTCC | [] |
| 4503 | 14 | GPR68 | {'ess': 0, 'nonest': 5} | True | 91233952 | CCAGGCTAGGCCAACCTGCCC | [] |
| 4504 | 14 | TRIP11 | {'ess': 0, 'nonest': 7} | True | 91969672 | AATCATCTATTGCTTTAAAAGG | [] |
| 4505 | 14 | SLC24A4 | {'ess': 0, 'nonest': 7} | True | 92483807 | TTAACCACTAGGGTGTAAGGGG | [] |
| 4506 | 14 | SLC24A4 | {'ess': 0, 'nonest': 7} | True | 92492906 | CCTTCTATCTTTCCTTCCACTG | [] |
| 4507 | 14 | SLC24A4 | {'ess': 0, 'nonest': 7} | True | 92493627 | TCAGCGCTAATCGTCTTCCCG | [] |
| 4508 | 14 | ITPK1 | {'ess': 6, 'nonest': 2} | False | 92941560 | CTCCGTGGCTACTGGGAGGAG | [] |
| 4509 | 14 | UBR7 | {'ess': 0, 'nonest': 7} | True | 93227224 | CACTCTAGCTGCAGTAATACTG | [] |
| 4510 | 14 | COX8C | {'ess': 1, 'nonest': 7} | True | 93348119 | CCATCTAATTCCTTCTGAACTG | [] |
| 4511 | 14 | PRIMA1 | {'ess': 0, 'nonest': 5} | True | 93733642 | ATTCCTAGGACTCGAGGCTATG | [] |
| 4512 | 14 | FAM181A | {'ess': 0, 'nonest': 7} | True | 93926163 | GCGTGGCTAGAGGTAGCCAAA | [] |
| 4513 | 14 | IFI27 | {'ess': 0, 'nonest': 7} | True | 94116526 | GCAGGGAGCTAGTAGAACCTC | [] |
| 4514 | 14 | SERPINA9 | {'ess': 0, 'nonest': 7} | True | 94463092 | TTCCCACCTAGGATTTAGTGGG | [] |
| 4515 | 14 | SERPINA4 | {'ess': 0, 'nonest': 7} | True | 94569594 | GGAGGGCTATGGTTTCGTGGG | [] |
| 4516 | 14 | SERPINA3 | {'ess': 0, 'nonest': 5} | True | 94623813 | GCAAGCTCTAGGCTTGCTTGG | [] |
| 4517 | 14 | DICER1 | {'ess': 3, 'nonest': 6} | False | 95090613 | CTTTCCTACTACTTCCACAGTG | [] |
| 4518 | 14 | CLMN | {'ess': 1, 'nonest': 7} | True | 95194413 | TGTCTTCTATCCAAAAGTATAG | [] |
| 4519 | 14 | LOC11226 | unavailable | False | 96417531 | GAGATGCTAGGAGATGTCCTTG | [] |
| 4520 | 14 | PAPOLA | {'ess': 1, 'nonest': 7} | True | 96531695 | CACTACTGTGTTAACAATATAG | [] |
| 4521 | 14 | SETD3 | {'ess': 0, 'nonest': 7} | True | 99398678 | TCGAGCTACTCCTTAACTCCAG | [] |
| 4522 | 14 | CCDC85C | {'ess': 0, 'nonest': 7} | True | 99515245 | GTGGCCCTACAAAGGCCCCTT | [] |
| 4523 | 14 | HHIPL1 | {'ess': 1, 'nonest': 7} | True | 99675625 | TTGCCTACAGGTCGGGGTTCT | [] |
| 4524 | 14 | EML1 | {'ess': 0, 'nonest': 6} | True | 99840111 | GGTACTAAATGACGCGCCACT | [] |
| 4525 | 14 | EVL | {'ess': 0, 'nonest': 7} | True | 1E+08 | ACCTGACTAAGAAAGGCCAGCC | [] |
| 4526 | 14 | SLC25A47 | {'ess': 0, 'nonest': 7} | True | 1E+08 | CCGGCTATGTGAGCAGACCCC | [] |
| 4527 | 14 | WARS1 | unavailable | False | 1E+08 | CGAGTGCTACTGAAAGTCGAAG | [] |
| 4528 | 14 | PPP2R5C | {'ess': 1, 'nonest': 6} | True | 1.02E+08 | CTCTCTACACCTTCCTTACTTG | [] |
| 4529 | 14 | PPP2R5C | {'ess': 1, 'nonest': 6} | True | 1.02E+08 | CGGAGGCTAGCGGCCGTCCTG | [] |
| 4530 | 14 | PPP2R5C | {'ess': 1, 'nonest': 6} | True | 1.02E+08 | TTTTCTACAGAATCCGGAAAAG | [] |
| 4531 | 14 | WDR20 | {'ess': 3, 'nonest': 5} | False | 1.02E+08 | CGCTACACTACAGTTCCACCT | [] |
| 4532 | 14 | ZNF839 | {'ess': 0, 'nonest': 7} | True | 1.02E+08 | CTCCTTCTAGCGGGCCCATCC | [] |
| 4533 | 14 | TECPR2 | {'ess': 0, 'nonest': 7} | True | 1.02E+08 | CCAGCTAACTTCTGCTTACCTG | [] |
| 4534 | 14 | ANKRD9 | {'ess': 0, 'nonest': 7} | True | 1.03E+08 | CCGGGCCTAGCCTTTGCCAGT | [] |
| 4535 | 14 | LBHD2 | unavailable | False | 1.03E+08 | CTTGTCCTATCCCCGGGCCGG | [] |
| 4536 | 14 | EXOC3L4 | unavailable | False | 1.03E+08 | AGAACTAGACGCAGGTGATCAG | [] |
| 4537 | 14 | TNFAIP2 | {'ess': 1, 'nonest': 4} | False | 1.03E+08 | GGGCCTCTAGGCAAGTCGAGG | [] |
| 4538 | 14 | TNFAIP2 | {'ess': 1, 'nonest': 4} | False | 1.03E+08 | AAAGCTAACCAACCTTTATAAG | [] |
| 4539 | 14 | TRMT61A | {'ess': 6, 'nonest': 2} | False | 1.04E+08 | GGCCCCCTAGCCTGGGGTCTT | [] |
| 4540 | 14 | COA8 | unavailable | False | 1.04E+08 | CCTAGTTGCTCCTCTTCTTTTG | [] |
| 4541 | 14 | TMEM179 | {'ess': 0, 'nonest': 6} | True | 1.05E+08 | GCACTGTGCCTAAATGACAGC | [] |
| 4542 | 14 | TMEM179 | {'ess': 0, 'nonest': 6} | True | 1.05E+08 | TGTCGGCTCTAAGGAGGGCGG | [] |
| 4543 | 14 | ADSS1 | unavailable | False | 1.05E+08 | AGTCTGTGACTAAAACAGCTGG | [] |
| 4544 | 14 | SIVA1 | {'ess': 1, 'nonest': 7} | True | 1.05E+08 | GGCCCTAAGGCAACACGACCA | [] |
| 4545 | 14 | CEP170B | unavailable | False | 1.05E+08 | CCTAGGCCCTCCCCATCCCCT | [] |
| 4546 | 14 | CEP170B | unavailable | False | 1.05E+08 | GGCCTAGATCAGGAACCTCTC | [] |
| 4547 | 14 | JAG2 | {'ess': 0, 'nonest': 7} | True | 1.05E+08 | CCGCCCCTACTCCTTGCCGGC | [] |
| 4548 | 14 | PACS2 | {'ess': 0, 'nonest': 7} | True | 1.05E+08 | GTGGGGCTAGAAGGTGGCCTT | [] |
| 4549 | 14 | MTA1 | {'ess': 1, 'nonest': 7} | True | 1.05E+08 | CGGCCCCTAGTCCTCGATGAC | [] |
| 4550 | 14 | CRIP2 | {'ess': 1, 'nonest': 6} | True | 1.05E+08 | TGTAGCCTAGGGCTGGACCTT | [] |
| 4551 | 15 | GOLGA6L | {'ess': 1, 'nonest': 3} | False | 20534258 | GCAGCCTCACCTAGTGTTCCT | [] |
| 4552 | 15 | GOLGA6L | unavailable | False | 22469824 | GCAGCCTCACCTAGTGTTCCT | [] |
| 4553 | 15 | NIPA1 | {'ess': 1, 'nonest': 7} | True | 22824238 | ATCTAGTCTGTTTTCATATTAG | [] |
| 4554 | 15 | MKRN3 | {'ess': 0, 'nonest': 7} | True | 23567305 | ACGATGCTACAGAATCAAATTG | [] |
| 4555 | 15 | NDN | {'ess': 2, 'nonest': 6} | False | 23686251 | TTGCTAGTCCTCAGAGACACTG | [] |
| 4556 | 15 | SNRPN | {'ess': 1, 'nonest': 7} | True | 24978443 | CAGTATGCTAAGGTCTTGGTGG | [] |
| 4557 | 15 | ATP10A | {'ess': 1, 'nonest': 7} | True | 25673403 | TCAGCTACATCAGCTGCAAAAG | [] |
| 4558 | 15 | LOC64520 | unavailable | False | 28350250 | GCAGCCTCACCTAGTGTTCCT | [] |
| 4559 | 15 | LOC10013 | unavailable | False | 28561378 | GCAGCCTCACCTAGTGTTCCT | [] |
| 4560 | 15 | APBA2 | {'ess': 0, 'nonest': 5} | True | 29117132 | GGTGGCCTAGATGTACAGCGG | [] |
| 4561 | 15 | ARHGAP11B | {'ess': 0, 'nonest': 7} | True | 30635629 | -1 | -1 |
| 4562 | 15 | FAN1 | {'ess': 0, 'nonest': 7} | True | 30916100 | ATGACCACATCCTAGCCACTAG | [] |
| 4563 | 15 | TRPM1 | {'ess': 0, 'nonest': 6} | True | 31001821 | ACAGACTAGCATTCAGTTTCTG | [] |
| 4564 | 15 | OTUD7A | {'ess': 0, 'nonest': 7} | True | 31501582 | CAGGTGTGCTTCTAAGGGGGG | [] |
| 4565 | 15 | ARHGAP11A | {'ess': 0, 'nonest': 7} | True | 32635937 | GTAAAACTCTAACAGTAGTATG | [] |
| 4566 | 15 | FMN1 | {'ess': 1, 'nonest': 7} | True | 32810944 | TGATGCTAACGAGGAAACAAAG | [] |
| 4567 | 15 | EMC7 | unavailable | False | 34084333 | CCTGACTACCTCCTTTTGCCA | [] |
| 4568 | 15 | NUTM1 | {'ess': 0, 'nonest': 6} | True | 34357490 | CTGCTCCCTACTGGCTACGAC | [] |
| 4569 | 15 | AQR | {'ess': 7, 'nonest': 0} | False | 34856791 | TTTGGCTACTTGGTCTCTTCCG | [] |
| 4570 | 15 | DPH6 | unavailable | False | 35220516 | ACTATAAAGATGCCACCATCTG | [] |
| 4571 | 15 | DPH6 | unavailable | False | 35373400 | AAACCACTAATGATAAAAATCG | ['AAAATCAATTTTCTTTTTCCTG'] |
| 4572 | 15 | MEIS2 | {'ess': 1, 'nonest': 7} | True | 36892172 | CTTATACTATTGGGCATGAATG | [] |
| 4573 | 15 | TMCO5A | {'ess': 1, 'nonest': 7} | True | 37966642 | TCTTGCCACTAAAAGGAGGAGG | [] |
| 4574 | 15 | RASGRP1 | {'ess': 0, 'nonest': 7} | True | 38490553 | TGGGCTAAGAACAGTCACCCT | [] |
| 4575 | 15 | FSIP1 | {'ess': 0, 'nonest': 7} | True | 39639092 | AAGCAATCTAGGAAACCACAAG | [] |
| 4576 | 15 | GPR176 | {'ess': 0, 'nonest': 7} | True | 39801131 | CCTTGCTAGGAATCCACCTTTG | [] |
| 4577 | 15 | PLCB2 | {'ess': 1, 'nonest': 7} | True | 40288056 | ACTTCTCTATGGGCTCAGTATG | [] |
| 4578 | 15 | KNSTRN | unavailable | False | 40393230 | ATGAAATCCTCTACATTTGGTG | [] |
| 4579 | 15 | IVD | {'ess': 0, 'nonest': 7} | True | 40418262 | TCTCAGGACTAGTGAAAGTCTG | [] |
| 4580 | 15 | IVD | {'ess': 0, 'nonest': 7} | True | 40419201 | TGCTAGGAGCTGATACCTCTAG | [] |
| 4581 | 15 | IVD | {'ess': 0, 'nonest': 7} | True | 40427207 | GAGCGTAGCCCTAGTCTTCTT | [] |
| 4582 | 15 | BAHD1 | {'ess': 3, 'nonest': 5} | False | 40466129 | GAGGCTACTGGGGGTTCTTAA | [] |
| 4583 | 15 | KNL1 | unavailable | False | 40662187 | AAGGGTCTAGTGGTAGAAATGG | [] |
| 4584 | 15 | DNAJC17 | {'ess': 6, 'nonest': 2} | False | 40774967 | GAGACCTAGGGTGTGGACTGA | [] |
| 4585 | 15 | ZFYVE19 | {'ess': 1, 'nonest': 7} | True | 40810469 | CCCTACCCTATCGTGCTGTTT | [] |
| 4586 | 15 | VPS18 | {'ess': 5, 'nonest': 3} | False | 40903340 | CACCCTCCTACAGCCAACTGA | [] |
| 4587 | 15 | EXD1 | {'ess': 1, 'nonest': 7} | True | 41183930 | CCCATCTCTAGGGCAGATTTAG | [] |
| 4588 | 15 | CHP1 | unavailable | False | 41264186 | -1 | -1 |
| 4589 | 15 | RPAP1 | {'ess': 7, 'nonest': 1} | False | 41517541 | TATCAACTATCCTAGGTCTCTG | [] |
| 4590 | 15 | TYRO3 | {'ess': 2, 'nonest': 5} | False | 41574230 | TTAGACTAGCTGGGGTCTCCA | [] |
| 4591 | 15 | TYRO3 | {'ess': 2, 'nonest': 5} | False | 41578275 | GTGGGCTAACAGCTACTGTGT | [] |
| 4592 | 15 | PLA2G4E | {'ess': 0, 'nonest': 5} | True | 41993753 | TGGGGCCTAGGAGGGACACTG | [] |
| 4593 | 15 | GANC | {'ess': 0, 'nonest': 7} | True | 42314095 | TGAGAGCTATTTTTCTCCAATG | [] |
| 4594 | 15 | SNAP23 | {'ess': 3, 'nonest': 5} | False | 42529890 | CCCACAGTATCTAGCACAGATG | [] |
| 4595 | 15 | LRRC57 | {'ess': 0, 'nonest': 7} | True | 42539587 | AAGACTAGCCCAGGCAACACA | [] |
| 4596 | 15 | HAUS2 | {'ess': 5, 'nonest': 3} | False | 42566815 | GATGACTACTTGGCATTAATAG | [] |
| 4597 | 15 | STARD9 | {'ess': 0, 'nonest': 7} | True | 42719573 | GAGATGCTACCTACCAAGGAAG | [] |
| 4598 | 15 | CDAN1 | {'ess': 4, 'nonest': 4} | False | 42724690 | CTCAGCCCTAGCTCTGGGCCA | [] |
| 4599 | 15 | TTBK2 | {'ess': 1, 'nonest': 7} | True | 42745794 | CTCCTATCTGCTGAGTTTACTG | [] |
| 4600 | 15 | PPIP5K1 | {'ess': 0, 'nonest': 7} | True | 43534673 | GACTTCTAATTTATCTCCTCAG | [] |
| 4601 | 15 | SERF2 | {'ess': 1, 'nonest': 7} | True | 43793772 | -1 | -1 |
| 4602 | 15 | MFAP1 | {'ess': 7, 'nonest': 1} | False | 43805093 | TTGGACCCTAGGTAGTTTTCCG | [] |
| 4603 | 15 | FRMD5 | {'ess': 0, 'nonest': 7} | True | 43873147 | AGGCTCTAGACTAAGGGGACA | [] |
| 4604 | 15 | SPG11 | {'ess': 0, 'nonest': 7} | True | 44563120 | AATCATCTAACCTGCTAGCATG | [] |
| 4605 | 15 | TERB2 | unavailable | False | 44978627 | TTACTATTTCCTCTTCAATTTG | [] |
| 4606 | 15 | DUOXA1 | {'ess': 0, 'nonest': 7} | True | 45120360 | CTTCTACCTGCTGGTGAATTTG | [] |
| 4607 | 15 | DUOX1 | {'ess': 0, 'nonest': 7} | True | 45164900 | GGGCCTAGAAGTTCTCATAATG | [] |
| 4608 | 15 | SHF | {'ess': 1, 'nonest': 7} | True | 45167946 | TCACATCTAAAGAGTCCGGATG | [] |
| 4609 | 15 | SHF | {'ess': 1, 'nonest': 7} | True | 45170468 | GACTCCTAGAGTATGACAGCTG | [] |
| 4610 | 15 | LOC10537 | unavailable | False | 45476542 | AAGGCTATGGGGCAATTCCTTG | [] |
| 4611 | 15 | SLC30A4 | {'ess': 0, 'nonest': 7} | True | 45502940 | GTGAGCTATGATGGTGTCATTG | [] |
| 4612 | 15 | SEMA6D | {'ess': 0, 'nonest': 6} | True | 47765059 | ATATACCTACTTTGCATGGTTG | [] |
| 4613 | 15 | SEMA6D | {'ess': 0, 'nonest': 6} | True | 47771784 | TGAGGCCTAGTATGTGTATTTG | [] |
| 4614 | 15 | SLC24A5 | {'ess': 0, 'nonest': 7} | True | 48141224 | ACCTAGTTCTTACCTTTTCTTG | [] |
| 4615 | 15 | EID1 | {'ess': 0, 'nonest': 5} | True | 48878739 | ATCTAACTACTCTCTATCAATG | ['ATCAATAATCTCATCACAGCCG'] |
| 4616 | 15 | DTWD1 | {'ess': 0, 'nonest': 7} | True | 49643577 | AACTAATGTGTAAGTTTTCCTG | [] |
| 4617 | 15 | DTWD1 | {'ess': 0, 'nonest': 7} | True | 49653890 | TCCATCTATTTTTCCTTTTCTG | ['TTTTCTATAAAAAAATGACAAG'] |
| 4618 | 15 | ATP8B4 | {'ess': 0, 'nonest': 5} | True | 49867127 | GGTCTGCTACTGGAGAATCAAG | [] |
| 4619 | 15 | HDC | {'ess': 0, 'nonest': 7} | True | 50242259 | CCTGTGTCTAAACCATGGCCT | [] |
| 4620 | 15 | AP4E1 | {'ess': 1, 'nonest': 7} | True | 51002661 | TTCTGCTAGGATCCCTCCATCC | ['CCTCCATCACCTTTTGACACTG'] |
| 4621 | 15 | CYP19A1 | {'ess': 1, 'nonest': 6} | True | 51210807 | TCTCTAGTGTTCCAGACACCTG | [] |
| 4622 | 15 | LYSMD2 | {'ess': 0, 'nonest': 7} | True | 51723606 | AATCACCTAACTGTGATAGAGG | [] |
| 4623 | 15 | MYO5C | {'ess': 0, 'nonest': 7} | True | 52193901 | TCTCCTGCTATAACCTATTCAG | [] |
| 4624 | 15 | NEDD4 | {'ess': 0, 'nonest': 7} | True | 55829896 | AATCTAATCAACTCCATCAAAG | [] |
| 4625 | 15 | GCOM1 | {'ess': 0, 'nonest': 3} | True | 57714568 | TTCAGCTAATTTTTGCGCTGCG | [] |
| 4626 | 15 | MYZAP | {'ess': 1, 'nonest': 4} | False | 57684497 | CCTCTGAGTGCCTAAGTCAGA | [] |
| 4627 | 15 | AQP9 | {'ess': 0, 'nonest': 6} | True | 58184134 | ATGCCACTACATGATGACACTG | [] |
| 4628 | 15 | MINDY2 | unavailable | False | 58831781 | TGCTCAGCTACAAAGCCTATAG | [] |
| 4629 | 15 | CCNB2 | {'ess': 1, 'nonest': 6} | True | 59124876 | CAGCCTAGGACCTTCCTATCA | [] |
| 4630 | 15 | RAB8B | {'ess': 0, 'nonest': 7} | True | 63251295 | TTATCCCTAAAATTCCTTCCTG | [] |
| 4631 | 15 | FBXL22 | {'ess': 0, 'nonest': 5} | True | 63599192 | AGTACCTACTATGTGCCAGGTG | [] |
| 4632 | 15 | FBXL22 | {'ess': 0, 'nonest': 5} | True | 63601038 | GCGGCGTCTAGCGCTGCAGCA | [] |
| 4633 | 15 | DAPK2 | {'ess': 0, 'nonest': 5} | True | 63911105 | ATCCCAGCTACTCAGGAGGCT | [] |
| 4634 | 15 | SNX22 | {'ess': 1, 'nonest': 7} | True | 64154150 | GTGTCTACAAAGCAGGCTCCA | [] |
| 4635 | 15 | PPIB | {'ess': 1, 'nonest': 6} | True | 64156022 | TGTGCCCTACTCCTTGGCGAT | [] |
| 4636 | 15 | PCLAF | unavailable | False | 64369329 | TTCTATTCTTTTTCATCATTTG | [] |
| 4637 | 15 | RBPMS2 | {'ess': 0, 'nonest': 7} | True | 64741179 | GAAAAACTAACAGAACTGACGG | [] |
| 4638 | 15 | PLEKHO2 | {'ess': 0, 'nonest': 7} | True | 64869887 | GCCCTAGGGTGCACTTCTCCT | [] |
| 4639 | 15 | ANKDD1A | {'ess': 2, 'nonest': 6} | False | 64957187 | CCCAGCTACTCGGGAGGCTGA | [] |
| 4640 | 15 | SPG21 | {'ess': 0, 'nonest': 7} | True | 64963619 | GACACACTACTGCTCCTCCTG | [] |
| 4641 | 15 | MTFMT | {'ess': 2, 'nonest': 6} | False | 65003061 | TTCCTAACTACTCAATGCATTG | [] |
| 4642 | 15 | RASL12 | {'ess': 0, 'nonest': 7} | True | 65051989 | GTCAGAATCTCTAGGCAGACAG | [] |
| 4643 | 15 | SLC51B | unavailable | False | 65052163 | CCTCACTAGCTCTCAGTTTCTG | [] |
| 4644 | 15 | UBAP1L | {'ess': 0, 'nonest': 4} | True | 65093062 | GGGCCTAGATGCCCAGGCATC | [] |
| 4645 | 15 | PDCD7 | {'ess': 2, 'nonest': 6} | False | 65118716 | CTTTACTAATGCAGCTTAACAG | [] |
| 4646 | 15 | IGDCC3 | {'ess': 1, 'nonest': 7} | True | 65328908 | ACTGGCTACTGTTCCGAGTGA | [] |
| 4647 | 15 | IGDCC4 | {'ess': 0, 'nonest': 7} | True | 65384008 | GAAGAGCTAGGCAGAGGAGGA | [] |
| 4648 | 15 | SMAD6 | {'ess': 0, 'nonest': 7} | True | 66781534 | CCGCCACTATCTGGGGTTGTT | [] |
| 4649 | 15 | SMAD3 | {'ess': 2, 'nonest': 6} | False | 67190535 | TGATGTCTCTAAGACACACTGG | [] |
| 4650 | 15 | AAGAB | {'ess': 0, 'nonest': 7} | True | 67226900 | TTTTAATTACCTAGATTTTCAG | [] |
| 4651 | 15 | IQCH | {'ess': 0, 'nonest': 7} | True | 67306827 | TGAGGACCTACTAGAAAAAATG | [] |
| 4652 | 15 | SKOR1 | {'ess': 0, 'nonest': 7} | True | 67833835 | CGGCCCTAGGGCAACAGCGGT | [] |
| 4653 | 15 | CORO2B | {'ess': 0, 'nonest': 5} | True | 68725973 | GGGGAGCTAACAGTTCTTGGG | [] |
| 4654 | 15 | NOX5 | {'ess': 0, 'nonest': 1} | False | 69056695 | GTGAGGCTAGAAATTCTCTTGG | [] |
| 4655 | 15 | GLCE | {'ess': 1, 'nonest': 5} | False | 69269243 | TGTGAGCTCTAGTTGTGCTTTG | [] |
| 4656 | 15 | UACA | {'ess': 0, 'nonest': 7} | True | 70657055 | ACGGCTAGCACACAAGCCCCT | [] |
| 4657 | 15 | LARP6 | {'ess': 0, 'nonest': 7} | True | 70851638 | ATGTCCTATGTGCTGGGTGCT | [] |
| 4658 | 15 | CT62 | {'ess': 1, 'nonest': 7} | True | 71111150 | CCTTCCTATCTCCTCTGACCA | [] |
| 4659 | 15 | NR2E3 | {'ess': 1, 'nonest': 0} | False | 71817683 | CCCCCACTAGTTTTTGAACATG | ['TTTTGAACATATCACAAAGGAG'] |
| 4660 | 15 | SENP8 | {'ess': 0, 'nonest': 7} | True | 72140261 | AATAGCTACTTTTTAGCAAGTG | [] |
| 4661 | 15 | GOLGA6B | {'ess': 0, 'nonest': 7} | True | 72666341 | GTGCTCCTAGGTGTCCTGCAT | [] |
| 4662 | 15 | ADPGK | {'ess': 0, 'nonest': 7} | True | 72752340 | AATCTTCCTAATAGTGAGGGTG | [] |
| 4663 | 15 | INSYN1 | unavailable | False | 73739916 | CGGCCCCTAGTTTTTCCCCCT | [] |
| 4664 | 15 | TBC1D21 | {'ess': 1, 'nonest': 7} | True | 73889004 | TCTAAGCAGCGACATTCTCCTG | [] |
| 4665 | 15 | TBC1D21 | {'ess': 1, 'nonest': 7} | True | 73901030 | CCTGATCTATTGAGGAATTCTG | [] |
| 4666 | 15 | STOML1 | {'ess': 0, 'nonest': 7} | True | 73983936 | AGGCTGCTACTTCAAGGCCCT | [] |
| 4667 | 15 | PML | {'ess': 1, 'nonest': 6} | True | 74043179 | CAGAGACTAAATTAGAAAGGGG | [] |
| 4668 | 15 | GOLGA6A | {'ess': 1, 'nonest': 6} | True | 74070909 | GTGCTCCTAGGTGTCCTGCAT | [] |
| 4669 | 15 | ISLR2 | {'ess': 0, 'nonest': 7} | True | 74138324 | CACATCCCTAGTAATGCCTATG | [] |
| 4670 | 15 | ISLR | {'ess': 0, 'nonest': 7} | True | 74176144 | GTGGGGCTAGAAGGAGGTGAG | [] |
| 4671 | 15 | CCDC33 | {'ess': 1, 'nonest': 6} | True | 74297090 | GCCTTCCTATTCGTCTGATGG | [] |
| 4672 | 15 | CCDC33 | {'ess': 1, 'nonest': 6} | True | 74335510 | GAGCCTACATCGTGGCCATCT | [] |
| 4673 | 15 | SEMA7A | {'ess': 1, 'nonest': 7} | True | 74410623 | GAGGCCCTAGTGGACCAGCAA | [] |
| 4674 | 15 | EDC3 | {'ess': 0, 'nonest': 7} | True | 74632611 | ACCCTCTAAGCAGAGTGCAGT | [] |
| 4675 | 15 | CYP1A1 | {'ess': 0, 'nonest': 7} | True | 74720488 | CAAGCACCTAAGAGCGCAGCT | [] |
| 4676 | 15 | SCAMP2 | {'ess': 0, 'nonest': 7} | True | 74845082 | GGAGGACTAATTCCCCTGGAA | [] |
| 4677 | 15 | FAM219B | unavailable | False | 74902618 | TAATGTCTACTGGAGGGTACAG | [] |
| 4678 | 15 | C15orf39 | {'ess': 1, 'nonest': 6} | True | 75211115 | AGTGCTACAGGTGATGGCTCT | [] |
| 4679 | 15 | LOC10537 | unavailable | False | 75226780 | TCACAGCTCTACCAAAGTCTTG | [] |
| 4680 | 15 | GOLGA6C | {'ess': 0, 'nonest': 7} | True | 75270198 | GTGCTCCTAGGTGTCCTGCAT | [] |
| 4681 | 15 | GOLGA6D | {'ess': 0, 'nonest': 7} | True | 75294474 | GTGCTCCTAGGTGTCCTGCAT | [] |
| 4682 | 15 | NEIL1 | {'ess': 0, 'nonest': 7} | True | 75352296 | CCATGCTAGGCAGTGGGGAAT | [] |
| 4683 | 15 | NEIL1 | {'ess': 0, 'nonest': 7} | True | 75355033 | GCCTCCTGCTAAGAGGCTGAG | [] |
| 4684 | 15 | IMP3 | {'ess': 5, 'nonest': 1} | False | 75639613 | GATCCGCTAGGCTTCCAGATC | [] |
| 4685 | 15 | TMEM266 | unavailable | False | 76204314 | GCTCTAGGCCTCAGGGACCCT | [] |
| 4686 | 15 | SCAPER | {'ess': 0, 'nonest': 7} | True | 76458082 | AGCACTCTAAAGATGGTCACTG | [] |
| 4687 | 15 | TSPAN3 | {'ess': 0, 'nonest': 7} | True | 77049834 | TGTCAACTATGCATAGGTTCCG | [] |
| 4688 | 15 | HMG20A | {'ess': 0, 'nonest': 7} | True | 77479314 | CATTCCCTAACGATCGAGTCTG | [] |
| 4689 | 15 | SH2D7 | {'ess': 0, 'nonest': 7} | True | 78101614 | GGTTGTCTCCTACCACCAGGT | [] |
| 4690 | 15 | ACSBG1 | {'ess': 0, 'nonest': 7} | True | 78168375 | TCCAGCTATTCAGGACGCTGA | [] |
| 4691 | 15 | IREB2 | {'ess': 3, 'nonest': 6} | False | 78498142 | AGATACTATGAGAATTTTCGTG | [] |
| 4692 | 15 | PSMA4 | {'ess': 5, 'nonest': 4} | False | 78548943 | TCTATTTATCCTTTTCTTTCTG | [] |
| 4693 | 15 | CHRNA5 | {'ess': 0, 'nonest': 7} | True | 78593115 | AACCTGGGCTATGAATTTCCAG | ['ATTTCCAATCTTCAACAACCTG'] |
| 4694 | 15 | ANKRD34C | {'ess': 0, 'nonest': 5} | True | 79294891 | CTTCTAGGTCATGATTTCTTCG | ['TTTCTTCAAAATCAGACAGCTG'] |
| 4695 | 15 | TMED3 | {'ess': 0, 'nonest': 7} | True | 79314563 | CAGCTACAGGCAGAAGACCCT | [] |
| 4696 | 15 | TMED3 | {'ess': 0, 'nonest': 7} | True | 79322213 | TGCCGGGGCTAGGAGTGGACT | [] |
| 4697 | 15 | ARNT2 | {'ess': 0, 'nonest': 7} | True | 80593697 | CGCAGCTACTCAGAAAACGGT | [] |
| 4698 | 15 | TLNRD1 | unavailable | False | 81003359 | GGGGTGCTAATTCACAGAATTG | [] |
| 4699 | 15 | IL16 | {'ess': 1, 'nonest': 6} | True | 81308797 | CCTGCCTAGGAGTCTCCAGCA | [] |
| 4700 | 15 | TMC3 | {'ess': 0, 'nonest': 7} | True | 81332418 | CTGGAGGAAGCCTAGACATCT | [] |
| 4701 | 15 | EFL1 | unavailable | False | 82130372 | AGGTAGCTACTTATTTTTGCTG | [] |
| 4702 | 15 | CPEB1 | {'ess': 1, 'nonest': 6} | True | 82544591 | GCTCCTCTAGCTGGAATCTCG | [] |
| 4703 | 15 | AP3B2 | {'ess': 0, 'nonest': 6} | True | 82689432 | GCTAGAGACTCTCCACTCTCC | [] |
| 4704 | 15 | FSD2 | {'ess': 0, 'nonest': 7} | True | 82759347 | TGTTCTAATAGAAAGTGACATG | [] |
| 4705 | 15 | WHAMM | {'ess': 1, 'nonest': 6} | True | 82833535 | TTGAGCCTAACCATCCCACTG | [] |
| 4706 | 15 | HOMER2 | {'ess': 0, 'nonest': 7} | True | 82849714 | AGCCCTAGTTATCGGTGCCCA | [] |
| 4707 | 15 | SH3GL3 | {'ess': 0, 'nonest': 7} | True | 83618155 | TTTAAATCCTAATTCTCCTTGG | [] |
| 4708 | 15 | ADAMTSL3 | {'ess': 0, 'nonest': 7} | True | 84016138 | GGTTTCTAGCCTAAATTGCAAG | [] |
| 4709 | 15 | UBE2Q2L | unavailable | False | 84185518 | GTCCCTAGCCAGCCTGAGCCA | [] |
| 4710 | 15 | LOC10537 | unavailable | False | 84290878 | GGCTCCTAGCCCCTTTCTTCA | [] |
| 4711 | 15 | ZNF592 | {'ess': 1, 'nonest': 5} | False | 84801573 | GGTATCCTATGCCTATACAAAG | ['CTATACAAAACAAAACTATTG'] |
| 4712 | 15 | ALPK3 | {'ess': 1, 'nonest': 7} | True | 84869455 | CGGAGGCTACCGCATGCCCTG | [] |
| 4713 | 15 | PDE8A | {'ess': 0, 'nonest': 7} | True | 85137902 | TCTCCCACTATTCAGGAGGTG | [] |
| 4714 | 15 | AGBL1 | {'ess': 0, 'nonest': 5} | True | 86907293 | GAACCTCGGGGCTATGGGATG | [] |

| | | | | | | | |
|---|---|---|---|---|---|---|---|
| 4715 | 15 | NTRK3 | {'exs': 1, 'nonexs': 6} | True | 87876934 | ACCACTAGCCAAGAATGTCCAG | [] |
| 4716 | 15 | MRPS11 | {'exs': 3, 'nonexs': 5} | False | 88472397 | GGACTTTCCGCTATGAGAAGGC | [] |
| 4717 | 15 | DET1 | {'exs': 2, 'nonexs': 7} | False | 88512950 | GAGGCACCTACGTGCAGCAGT | [] |
| 4718 | 15 | AEN | {'exs': 1, 'nonexs': 7} | True | 88630056 | CTGGCCCACCTACTGACGAGA | [] |
| 4719 | 15 | HAPLN3 | {'exs': 0, 'nonexs': 7} | True | 88877969 | CAGGTCCTAGTGCTGGCGGTA | [] |
| 4720 | 15 | MFGE8 | {'exs': 1, 'nonexs': 6} | True | 88899394 | GTGGCCACTAACAGCCCAGCA | [] |
| 4721 | 15 | POLG | {'exs': 5, 'nonexs': 5} | False | 89316750 | GCAGTGCTATGGTCCAGGCTG | [] |
| 4722 | 15 | RHCG | {'exs': 0, 'nonexs': 7} | True | 89472734 | TGGGAGCCTAGGGTACCAAGG | [] |
| 4723 | 15 | TICRR | unavailable | False | 89627085 | TTTGTGGCTATAAGTCCTCCAG | [] |
| 4724 | 15 | PEX11A | {'exs': 0, 'nonexs': 7} | True | 89683376 | AAACACCCTAACGGGTCTTCAG | [] |
| 4725 | 15 | WDR93 | {'exs': 0, 'nonexs': 7} | True | 89703167 | AATGAGGCTAAAAAGGAAGATG | [] |
| 4726 | 15 | ANPEP | {'exs': 0, 'nonexs': 7} | True | 89785348 | GGGGACTATTTGCTGTTTTCTG | [] |
| 4727 | 15 | ZNF710 | {'exs': 0, 'nonexs': 7} | True | 90074549 | ATCTAAGTTCTCTGTTCTTCAG | [] |
| 4728 | 15 | ZNF710 | {'exs': 0, 'nonexs': 7} | True | 90079828 | CTTGCGCTATAGCACATTGTAG | [] |
| 4729 | 15 | IDH2 | {'exs': 1, 'nonexs': 7} | True | 90084265 | CTCCCCCTACTGCCTGCCCAG | [] |
| 4730 | 15 | CIB1 | {'exs': 0, 'nonexs': 7} | True | 90230692 | TTCTAAACCTTTATTACTAATG | [] |
| 4731 | 15 | ZNF774 | {'exs': 0, 'nonexs': 7} | True | 90361282 | TACTTCCTAAATCAAATGGGTG | [] |
| 4732 | 15 | MAN2A2 | {'exs': 0, 'nonexs': 5} | True | 90919786 | GAAGCCCTAACCCAAGCGGAG | [] |
| 4733 | 15 | HDDC3 | {'exs': 1, 'nonexs': 7} | True | 90931542 | CCCAGCTATTGTATTTTTACTG | [] |
| 4734 | 15 | SLCO3A1 | {'exs': 0, 'nonexs': 7} | True | 92163134 | TCACTATAAAACGGACTCCATG | [] |
| 4735 | 15 | ST8SIA2 | {'exs': 0, 'nonexs': 7} | True | 92464384 | CCCACCCTACGTGGCCCCATC | [] |
| 4736 | 15 | CHD2 | {'exs': 1, 'nonexs': 6} | True | 92949079 | CTACCTACTTGCACCAGGAATG | [] |
| 4737 | 15 | RGMA | {'exs': 0, 'nonexs': 6} | True | 93044997 | ACGCGTCTAGCAGAACACAGG | [] |
| 4738 | 15 | MCTP2 | {'exs': 0, 'nonexs': 7} | True | 94479033 | GTGTGCCCTAGAGAGCGCTGC | [] |
| 4739 | 15 | TTC23 | {'exs': 0, 'nonexs': 5} | True | 99137960 | CACCCTAAATGACAGTGCCCAC | [] |
| 4740 | 15 | LRRC28 | {'exs': 0, 'nonexs': 7} | True | 99355453 | CTCCTAGTGGTCACTCTCCATC | [] |
| 4741 | 15 | MEF2A | {'exs': 0, 'nonexs': 7} | True | 99694122 | GGAGGCTAAAGAGACATGATTG | [] |
| 4742 | 15 | LYSMD4 | {'exs': 0, 'nonexs': 7} | True | 99717799 | CCTGCTTTCTAGCTAACCAAAG | [] |
| 4743 | 15 | LYSMD4 | {'exs': 0, 'nonexs': 7} | True | 99720955 | TCTCCTAGATCTTTCCTGTCAG | [] |
| 4744 | 15 | ADAMTS1 | {'exs': 0, 'nonexs': 7} | True | 1E+08 | GCCACTAGGGCCCCTGGAGCCT | [] |
| 4745 | 15 | CERS3 | unavailable | False | 1E+08 | TTCCAGCTAATGGCCATGCTGC | [] |
| 4746 | 15 | PCSK6 | {'exs': 0, 'nonexs': 1} | False | 1.01E+08 | AACTTACTACAAAGCTACAGTG | [] |
| 4747 | 15 | PCSK6 | {'exs': 0, 'nonexs': 1} | False | 1.01E+08 | CCCCTTCCTACACATTCCAGG | [] |
| 4748 | 15 | TM2D3 | {'exs': 0, 'nonexs': 7} | True | 1.02E+08 | CCACAGCTAAATGTACAAAGAG | [] |
| 4749 | 15 | TARS3 | unavailable | False | 1.02E+08 | TGTGCTACGTGAAAGAAGCCAG | [] |
| 4750 | 15 | OR4F4 | {'exs': 0, 'nonexs': 3} | True | 1.02E+08 | AAGATCTAAAACTTTACACTAG | [] |
| 4751 | 16 | POLR3K | {'exs': 6, 'nonexs': 2} | False | 47629 | CTGGCCCTAATCCCTCCAGCG | [] |
| 4752 | 16 | FAM234A | unavailable | False | 265021 | GTGCCTCTACGCCTCACTCTG | [] |
| 4753 | 16 | RGS11 | {'exs': 0, 'nonexs': 7} | True | 269268 | CCACTAGGCCACCCCATCTCC | ['TCTCCACCCCCAGGGCCACAAG'] |
| 4754 | 16 | RGS11 | {'exs': 0, 'nonexs': 7} | True | 272376 | GGTTCTAGGTGCATCTTTCCAG | [] |
| 4755 | 16 | PDIA2 | {'exs': 0, 'nonexs': 7} | True | 287112 | GGGCAGCTACAGTTCCTCCTT | [] |
| 4756 | 16 | DECR2 | {'exs': 0, 'nonexs': 7} | True | 411577 | CCTCACCTAGAGCTTAGCAGA | [] |
| 4757 | 16 | RAB11FIP | {'exs': 0, 'nonexs': 7} | True | 497300 | CAACAGATCTATACGAAGCTGG | [] |
| 4758 | 16 | RAB11FIP | {'exs': 0, 'nonexs': 7} | True | 520838 | CTGCCTCTACTTGACCTCCAG | [] |
| 4759 | 16 | RAB40C | {'exs': 0, 'nonexs': 7} | True | 627521 | TCCCCGCTAGGAGATCTTGCA | [] |
| 4760 | 16 | WFIKKN1 | {'exs': 0, 'nonexs': 7} | True | 634056 | CGGGGGCTAGTCCTGGAAGCG | [] |
| 4761 | 16 | METTL26 | {'exs': 0, 'nonexs': 2} | False | 634525 | CGGCCTAGGGAGGCAGGGTTC | [] |
| 4762 | 16 | MCRIP2 | {'exs': 0, 'nonexs': 2} | False | 648189 | CAGCCACTAGGAACAGCTGGT | [] |
| 4763 | 16 | JMJD8 | {'exs': 1, 'nonexs': 7} | True | 682793 | TTTTGGCTAGCCGAGGAAGGT | [] |
| 4764 | 16 | FBXL16 | {'exs': 0, 'nonexs': 7} | True | 694274 | CTCGCGCTACTCAATGACGAG | [] |
| 4765 | 16 | CCDC78 | {'exs': 0, 'nonexs': 7} | True | 722677 | GAGACGGCCTAGTGGCTGCCG | [] |
| 4766 | 16 | CIAO3 | unavailable | False | 730416 | GCCCCTACCACCGGATGCCCA | [] |
| 4767 | 16 | CHTF18 | {'exs': 2, 'nonexs': 6} | False | 797974 | AGAGAACTAGAGCAAGTCCCTG | [] |
| 4768 | 16 | LMF1 | {'exs': 0, 'nonexs': 7} | True | 854531 | TGCACGTCTAGAGGGGCCCGG | [] |
| 4769 | 16 | C1QTNF8 | {'exs': 0, 'nonexs': 7} | True | 1093500 | CCCGGCTACAGCTCGGCGGC | [] |
| 4770 | 16 | CACNA1H | {'exs': 0, 'nonexs': 7} | True | 1208690 | CCTCCCCTGCTAGGGCGTCTG | [] |
| 4771 | 16 | CACNA1H | {'exs': 0, 'nonexs': 7} | True | 1220993 | CCGAGCTACACGGGGTCATCT | [] |
| 4772 | 16 | UNKL | {'exs': 1, 'nonexs': 7} | True | 1371599 | TGCAGGTCTAACGCAGAACCTC | [] |
| 4773 | 16 | PERCC1 | unavailable | False | 1433396 | CGCTTCGCTAGGCCTCGGCTG | [] |
| 4774 | 16 | TELO2 | {'exs': 5, 'nonexs': 3} | False | 1509935 | AGGGACTAGGGAGACGCGGGT | [] |
| 4775 | 16 | IFT140 | {'exs': 0, 'nonexs': 7} | True | 1556163 | TGGGCTACAAAATTCCTGCATG | [] |
| 4776 | 16 | NME3 | {'exs': 0, 'nonexs': 7} | True | 1770648 | GCCGGGCTACTCATACAGCCA | [] |
| 4777 | 16 | MRPS34 | {'exs': 4, 'nonexs': 4} | False | 1772220 | ATTCTAGACGGGGGTGCCCTT | [] |
| 4778 | 16 | NUBP2 | {'exs': 4, 'nonexs': 4} | False | 1789012 | CTCCGCTAACTGTGTGGACAC | [] |
| 4779 | 16 | MSRB1 | unavailable | False | 1939111 | GCCCGCCTAGTGACCCTGGGA | [] |
| 4780 | 16 | RPL3L | {'exs': 1, 'nonexs': 7} | True | 1944936 | CACAGCCTACAAGTCTCCCGA | [] |
| 4781 | 16 | RPS2 | {'exs': 7, 'nonexs': 1} | False | 1962097 | AAACCCTATGTTGTAGCCACAG | [] |
| 4782 | 16 | RNF151 | {'exs': 0, 'nonexs': 5} | True | 1968924 | ATTTACCTATTTACAAGGTATG | [] |
| 4783 | 16 | TBL3 | {'exs': 6, 'nonexs': 2} | False | 1978684 | CCGGACCGGCTAGGGCAGTGC | [] |
| 4784 | 16 | GFER | {'exs': 7, 'nonexs': 1} | False | 1986027 | GACCACCCTCTAGTCACAGGA | [] |
| 4785 | 16 | SYNGR3 | {'exs': 1, 'nonexs': 7} | True | 1993071 | GCCGCTAGTAGGCGGGCACCT | [] |
| 4786 | 16 | ZNF598 | {'exs': 1, 'nonexs': 1} | False | 1998231 | CGGGAGCTACGTGATGATCCT | [] |
| 4787 | 16 | SLC9A3R2 | {'exs': 1, 'nonexs': 7} | True | 2037785 | GAGCCCTAGTCCCACACCCCA | [] |
| 4788 | 16 | PKD1 | {'exs': 0, 'nonexs': 7} | True | 2089726 | GGAGGACTAAGTGCTGCTGGG | [] |
| 4789 | 16 | RAB26 | {'exs': 1, 'nonexs': 6} | True | 2153217 | TACTTTGCTATGGCTGTGAAGG | [] |
| 4790 | 16 | MLST8 | {'exs': 3, 'nonexs': 5} | False | 2208876 | CACAGGCTAGCCCAGCACACT | [] |
| 4791 | 16 | E4F1 | {'exs': 5, 'nonexs': 3} | False | 2235571 | CATGCTAGACGATGACCGTCTC | [] |
| 4792 | 16 | RNPS1 | {'exs': 3, 'nonexs': 5} | False | 2253896 | GGCTAGAAAAGTGACAAAACTG | [] |
| 4793 | 16 | ATP6V0C | {'exs': 7, 'nonexs': 1} | False | 2519744 | GAGGGTCTACTTTGTGGAGAG | [] |
| 4794 | 16 | PDPK1 | {'exs': 4, 'nonexs': 4} | False | 2597421 | GAGCCACTAGTGTGGGCGTCT | [] |
| 4795 | 16 | SRRM2 | {'exs': 6, 'nonexs': 2} | False | 2759788 | AGCTCCTAGAGGGCAGGGACC | [] |
| 4796 | 16 | PRSS21 | {'exs': 0, 'nonexs': 7} | True | 2821290 | GAAGGGCTATGGGGCTGGGTA | [] |
| 4797 | 16 | ZG16B | {'exs': 0, 'nonexs': 7} | True | 2832158 | CCACCCTAGCGACCCACGGGT | [] |
| 4798 | 16 | PRSS22 | {'exs': 0, 'nonexs': 7} | True | 2853092 | TGCGCCCTAGGAGCGCGCGGC | [] |
| 4799 | 16 | ZSCAN10 | {'exs': 1, 'nonexs': 7} | True | 3089090 | GCCAAGCTAGTACAGCGTCTC | [] |
| 4800 | 16 | ZNF205 | {'exs': 1, 'nonexs': 7} | True | 3120324 | GCCTCCTAGGTGGGAGCGGGT | [] |
| 4801 | 16 | ZNF200 | {'exs': 0, 'nonexs': 7} | True | 3226749 | TGTTGCTATATCTGTAGGTCAG | [] |
| 4802 | 16 | ZNF263 | {'exs': 1, 'nonexs': 7} | True | 3290557 | CACTGTTACTAACCTGTGTGAG | [] |
| 4803 | 16 | TIGD7 | {'exs': 0, 'nonexs': 5} | True | 3298964 | AAAATCTAATGATTAGAACCAG | [] |
| 4804 | 16 | ZNF75A | {'exs': 0, 'nonexs': 7} | True | 3322918 | TCTTCTATCCTAGGGTTTAATG | [] |
| 4805 | 16 | ZNF75A | {'exs': 0, 'nonexs': 7} | True | 3322970 | TTCTACATTGGCTGCCTCCTTG | [] |
| 4806 | 16 | C16orf90 | {'exs': 0, 'nonexs': 7} | True | 3493838 | GGATCCCTATGGCCTCTCCAG | [] |
| 4807 | 16 | NLRC3 | {'exs': 0, 'nonexs': 1} | False | 3547120 | CCGTTCCTAACAGCCAAGTAGC | [] |
| 4808 | 16 | CREBBP | {'exs': 4, 'nonexs': 3} | False | 3727717 | ACAATGCTACAAGCCCTCCACC | ['CTCCACAAACTTCTCTAGCGTG'] |
| 4809 | 16 | SRL | {'exs': 0, 'nonexs': 7} | True | 4192152 | CACCACCTAGTGCTTCCTGTAC | [] |
| 4810 | 16 | CORO7 | {'exs': 1, 'nonexs': 6} | True | 4355157 | GGCGCAGGCTAGTCCTGGGGC | [] |
| 4811 | 16 | DNAJA3 | {'exs': 7, 'nonexs': 1} | False | 4455567 | TTCCCGGCCTAGTTTCCAGTG | [] |
| 4812 | 16 | NMRAL1 | {'exs': 0, 'nonexs': 7} | True | 4463476 | CTAAGCCAGAAATTTCACTCTG | [] |
| 4813 | 16 | ANKS3 | {'exs': 1, 'nonexs': 6} | True | 4697027 | GGCCCGCAGGCTAGGTCTCCC | [] |
| 4814 | 16 | C16orf71 | {'exs': 1, 'nonexs': 6} | True | 4747524 | GGCAGCTATAGTTGCTCCAGAC | [] |
| 4815 | 16 | UBN1 | {'exs': 1, 'nonexs': 7} | True | 4871984 | TCAGCCCTGCTACCTGCAAATC | [] |
| 4816 | 16 | PPL | {'exs': 1, 'nonexs': 7} | True | 4893383 | AGCTGTGCCTACTTCTGCCCA | [] |
| 4817 | 16 | SEC14L5 | {'exs': 0, 'nonexs': 7} | True | 5014969 | GCCCGGCTATCTGGAGACCAG | [] |
| 4818 | 16 | C16orf89 | {'exs': 0, 'nonexs': 7} | True | 5044582 | ACACCTATAATCCCACACTTTG | [] |
| 4819 | 16 | EEF2KMT | unavailable | False | 5085631 | AGTCCTACAGGGTGAGATTCAG | [] |
| 4820 | 16 | RBFOX1 | {'exs': 1, 'nonexs': 6} | True | 7709640 | CCCTCTCTATTTTCTTTCCTGG | [] |
| 4821 | 16 | METTL22 | unavailable | False | 8673549 | TACACAAGCGCTATAGTCCTTG | [] |
| 4822 | 16 | CARHSP1 | {'exs': 2, 'nonexs': 7} | False | 8855163 | TCTCCTAGGAGCTGATGACATG | [] |
| 4823 | 16 | USP7 | {'exs': 2, 'nonexs': 6} | False | 8894809 | TATGGAATGCTATTGCTCCTGG | [] |
| 4824 | 16 | ATF7IP2 | {'exs': 0, 'nonexs': 7} | True | 10481898 | TGTCTACTAATTCAGGTAGTGG | [] |
| 4825 | 16 | EMP2 | {'exs': 0, 'nonexs': 7} | True | 10532904 | AACTCTATTTGCGCTTCCTCAG | [] |
| 4826 | 16 | NUBP1 | {'exs': 7, 'nonexs': 1} | False | 10768010 | CTGTAGGCTAACGTGGCTGGG | [] |
| 4827 | 16 | TVP23A | unavailable | False | 10770271 | CATTTCTCACCTAATGCTGGTG | [] |
| 4828 | 16 | DEXI | {'exs': 1, 'nonexs': 6} | True | 10941717 | TCGAGACCCTACTCCAAGTACC | [] |
| 4829 | 16 | CLEC16A | {'exs': 0, 'nonexs': 6} | True | 11126279 | AAATTCTAAGAAGCTGGTGCTG | [] |
| 4830 | 16 | CLEC16A | {'exs': 0, 'nonexs': 6} | True | 11136175 | GCCACCTAACAATTTTCCCGTG | [] |
| 4831 | 16 | CLEC16A | {'exs': 0, 'nonexs': 6} | True | 11174229 | GGAGCACTAAACCCAGTGGCC | [] |
| 4832 | 16 | LOC10799 | unavailable | False | 11213733 | CTACCCCTCATCTCTCCCCAT | [] |
| 4833 | 16 | LOC10799 | unavailable | False | 11224606 | CCAGCCTACCAAGAACCTCGA | [] |
| 4834 | 16 | PRM2 | {'exs': 0, 'nonexs': 7} | True | 11275829 | GAATTGCTATGGCCTCACTTGG | [] |
| 4835 | 16 | RMI2 | unavailable | False | 11350789 | CATACTCTAAGGAATATTCCTG | [] |
| 4836 | 16 | LOC40049 | unavailable | False | 11389442 | GGAATCCTAGTTAGAGGCAGAG | [] |
| 4837 | 16 | LITAF | {'exs': 0, 'nonexs': 7} | True | 11549536 | TGAGTCCTACAAACGCTTGTAG | [] |
| 4838 | 16 | NPIPB2 | unavailable | False | 11929351 | CTGAACTATCCTGCAACTGCAG | [] |
| 4839 | 16 | SNX29 | {'exs': 0, 'nonexs': 5} | True | 12526612 | GGGATGCCGCTAAGAGGATGA | [] |
| 4840 | 16 | SNX29 | {'exs': 0, 'nonexs': 5} | True | 12561100 | CGTGAAATCTAATTCCACTGTG | [] |
| 4841 | 16 | SHISA9 | {'exs': 0, 'nonexs': 5} | True | 13474247 | GCTTTCTATGTTTTGAGGAGTG | [] |
| 4842 | 16 | PARN | {'exs': 3, 'nonexs': 5} | False | 14461206 | CGTGGTCTAAAGAAGCCTATGG | [] |
| 4843 | 16 | PARN | {'exs': 3, 'nonexs': 5} | False | 14576257 | GGAAGGCCTAGATTTCCAGCG | [] |
| 4844 | 16 | LOC10537 | unavailable | False | 14702996 | CATACTCTAAGCCCCGCCAGC | [] |
| 4845 | 16 | LOC10537 | unavailable | False | 14742118 | CATACTCTAAGCCCCGCCAGC | [] |
| 4846 | 16 | PDXDC1 | {'exs': 0, 'nonexs': 7} | True | 15036014 | TTCTGGCCTACAGAAAGGGCAC | [] |
| 4847 | 16 | BMERB1 | unavailable | False | 15596928 | GGGGCTACATGATGTTGCACT | [] |
| 4848 | 16 | NDE1 | {'exs': 1, 'nonexs': 7} | True | 15699821 | GTTTTCCTAAAGCAACATCTTG | [] |
| 4849 | 16 | NDE1 | {'exs': 1, 'nonexs': 7} | True | 15699892 | AAAAACCTAGTGTAAAAAGACG | [] |
| 4850 | 16 | ABCC6 | {'exs': 0, 'nonexs': 7} | True | 16161369 | CCCACTAGCAGGGGTCCGACA | [] |
| 4851 | 16 | ABCC6 | {'exs': 0, 'nonexs': 7} | True | 16175919 | GCGTCTAAGCTCGGGCCTCCT | [] |
| 4852 | 16 | NOMO3 | {'exs': 0, 'nonexs': 7} | True | 16294764 | TACCTCCTAATAGTGGATACAG | [] |
| 4853 | 16 | XYLT1 | {'exs': 0, 'nonexs': 7} | True | 17108694 | TGCCCAGTGCTACCTGAGCCG | [] |
| 4854 | 16 | XYLT1 | {'exs': 0, 'nonexs': 7} | True | 17117426 | CAAATAGCTACCTAATACCTAG | [] |
| 4855 | 16 | NOMO2 | {'exs': 0, 'nonexs': 7} | True | 18499969 | TACCTCCTAATAGTGGATACAG | [] |
| 4856 | 16 | RPS15A | {'exs': 7, 'nonexs': 1} | False | 18783008 | ATCCCTAGAAAAAGAATCCCAG | [] |
| 4857 | 16 | GDE1 | {'exs': 0, 'nonexs': 7} | True | 19503459 | GAAAGTCTAGAAGTGAGGTTCG | [] |
| 4858 | 16 | PDILT | {'exs': 0, 'nonexs': 7} | True | 20359318 | GAGAAGCTAAAGTTCTTCCTTG | [] |
| 4859 | 16 | PDILT | {'exs': 0, 'nonexs': 7} | True | 20365458 | GTCAAAGACGACTACGTTGAAG | [] |
| 4860 | 16 | ACSM5 | {'exs': 0, 'nonexs': 7} | True | 20440451 | GGAGGTCTACGGGGCCTTCCT | [] |
| 4861 | 16 | ACSM2A | {'exs': 0, 'nonexs': 7} | True | 20480938 | CCAGGCTACTGCTTCCCCATC | ['CCCATCACCTCTCCTCGGACGG'] |
| 4862 | 16 | ACSM1 | {'exs': 0, 'nonexs': 7} | True | 20627075 | CACGGCTCTATTCAGATTTAGG | [] |
| 4863 | 16 | THUMPD1 | {'exs': 0, 'nonexs': 7} | True | 20736879 | TTCCTATGAGAAGTCATTTTCG | ['TCATTTGACTTGGATCCTTCTG'] |
| 4864 | 16 | DCUN1D3 | {'exs': 0, 'nonexs': 7} | True | 20859885 | GAGCCACTAAGTCTGCTCCTC | [] |
| 4865 | 16 | IGSF6 | {'exs': 0, 'nonexs': 7} | True | 21641533 | ACGTTTCTATGGCCTTTCATAG | [] |
| 4866 | 16 | PDZD9 | {'exs': 0, 'nonexs': 7} | True | 21984266 | CAGATTTGCTAACCAACCTTTG | [] |
| 4867 | 16 | VWA3A | {'exs': 0, 'nonexs': 7} | True | 22155901 | AACACTTCTCTAGGAAAGAGTG | [] |
| 4868 | 16 | SDR42E2 | {'exs': 0, 'nonexs': 1} | False | 22170906 | TCTATGGGCTTCCCTCCAAATG | [] |
| 4869 | 16 | SDR42E2 | {'exs': 0, 'nonexs': 1} | False | 22181947 | GGACTACTGTGAAGATTAAATG | [] |
| 4870 | 16 | POLR3E | {'exs': 7, 'nonexs': 1} | False | 22326759 | ACTCCAGGCCACTATACTGAGC | [] |
| 4871 | 16 | EARS2 | {'exs': 5, 'nonexs': 1} | False | 23524370 | CCTAGCTGGAAACCACCTTCTC | [] |
| 4872 | 16 | UBFD1 | {'exs': 0, 'nonexs': 7} | True | 23564562 | CAGAACTAAGCAAAATCACCTG | [] |
| 4873 | 16 | DCTN5 | {'exs': 7, 'nonexs': 1} | False | 23667023 | CCCAGCTCTAAGGAGCTTGAAC | [] |
| 4874 | 16 | DCTN5 | {'exs': 7, 'nonexs': 1} | False | 23667143 | GAGATGCTAGACTTGCGTCAGC | [] |
| 4875 | 16 | ERN2 | {'exs': 1, 'nonexs': 7} | True | 23695111 | GGGTTTTCTACGTACTGAGCAG | [] |
| 4876 | 16 | ERN2 | {'exs': 1, 'nonexs': 7} | True | 23695960 | CCCACTACGGTGAGTTGCTCA | [] |
| 4877 | 16 | PRKCB | {'exs': 1, 'nonexs': 6} | True | 24220112 | TGCATTCACCTACACATTAATG | [] |
| 4878 | 16 | SLC5A11 | {'exs': 0, 'nonexs': 7} | True | 24911532 | ACACTAAGCAAAATAGCCCCAG | [] |
| 4879 | 16 | KDM8 | unavailable | False | 27220729 | TCCTGGCTACGACCACCAGAA | [] |
| 4880 | 16 | NSMCE1 | {'exs': 5, 'nonexs': 3} | False | 27225156 | ATGGCTAATGCTGCCTGGACC | [] |
| 4881 | 16 | IL4R | {'exs': 0, 'nonexs': 7} | True | 27363829 | ATGCACCTAAGAGACCCTCATG | [] |
| 4882 | 16 | GTF3C1 | {'exs': 7, 'nonexs': 1} | False | 27461349 | GTCCTAGAGGTGGATCCACTTC | [] |
| 4883 | 16 | GSG1L | {'exs': 0, 'nonexs': 7} | True | 27806044 | CAACCTCTATCCCTGCTGTGTC | [] |
| 4884 | 16 | XPO6 | {'exs': 1, 'nonexs': 7} | True | 28098537 | GCAGGCCTAGAGCTTCACAGT | [] |
| 4885 | 16 | NPIPB6 | unavailable | False | 28342606 | ACCTACGGATTTATTCTTATTG | [] |
| 4886 | 16 | NPIPB8 | unavailable | False | 28658629 | ACCTACGGATTTATTCTTATTG | [] |
| 4887 | 16 | NPIPB9 | unavailable | False | 28772754 | ACCTACGGATTTATTCTTATTG | [] |
| 4888 | 16 | ATXN2L | {'exs': 3, 'nonexs': 5} | False | 28836479 | CCTCGGCTAAGTCAGGAGACA | [] |
| 4889 | 16 | CD19 | {'exs': 2, 'nonexs': 4} | False | 28935442 | CCATAGTACTAGGGGGAAGGG | [] |
| 4890 | 16 | QPRT | {'exs': 0, 'nonexs': 7} | True | 29697410 | TTTAGGACTAGTGGATTTTGGG | [] |
| 4891 | 16 | PRRT2 | {'exs': 0, 'nonexs': 7} | True | 29813953 | CCTGGGCTAGGGTCCCATGGG | [] |
| 4892 | 16 | KCTD13 | {'exs': 0, 'nonexs': 7} | True | 29921853 | GTAATTCTAGCTACTTGGGAGG | [] |
| 4893 | 16 | TAOK2 | {'exs': 1, 'nonexs': 6} | True | 29987979 | GAGTCAGCTACCTCCAGGGGG | [] |
| 4894 | 16 | INO80E | {'exs': 3, 'nonexs': 6} | False | 30004635 | AGCCTCCTAACTGGACTCCCT | [] |
| 4895 | 16 | MAPK3 | {'exs': 2, 'nonexs': 6} | False | 30116667 | GTCTGGGCTAGGGGGCCTCCA | [] |
| 4896 | 16 | MAPK3 | {'exs': 2, 'nonexs': 6} | False | 30116936 | GGGATGCCTACGTGCCCCCCT | [] |
| 4897 | 16 | CORO1A | {'exs': 1, 'nonexs': 7} | True | 30188963 | GGGCTCTACTTGGCCTGGACT | [] |
| 4898 | 16 | MYLPF | {'exs': 1, 'nonexs': 7} | True | 30377899 | GTGCCCCTACTCCTGGTCCTT | [] |
| 4899 | 16 | SEPTIN1 | unavailable | False | 30378282 | GCGGGGCTACGGACTCAGGCT | [] |
| 4900 | 16 | DCTPP1 | {'exs': 0, 'nonexs': 7} | True | 30424232 | ATCTTTCTAGGTTGAGGTCTGG | [] |
| 4901 | 16 | ZNF689 | {'exs': 0, 'nonexs': 7} | True | 30604263 | CTTCCCCTTCTAATGGGCGTTC | [] |
| 4902 | 16 | PRR14 | {'exs': 0, 'nonexs': 7} | True | 30656310 | GGGGCACCTAGGTCCAGTGGG | [] |
| 4903 | 16 | PHKG2 | {'exs': 1, 'nonexs': 6} | True | 30757396 | TGAGGTCCTAGCCCAGCACAA | [] |
| 4904 | 16 | FBXL19 | {'exs': 1, 'nonexs': 5} | False | 30947229 | CGCCCAACTAGCTGTCCTTGA | [] |
| 4905 | 16 | STX1B | {'exs': 0, 'nonexs': 7} | True | 30992820 | GGGGGGGCCTACAAGCCCAGC | [] |
| 4906 | 16 | ITGAM | {'exs': 0, 'nonexs': 6} | True | 31331706 | AGGAGCCGCTACTGGGGTTCG | [] |
| 4907 | 16 | ARMC5 | {'exs': 1, 'nonexs': 7} | True | 31466888 | ACAGTCTAGGCAGGGACCCTT | [] |
| 4908 | 16 | AHSP | {'exs': 0, 'nonexs': 7} | True | 31528690 | CTGAGCTAGGAGGAGGGCGGT | [] |
| 4909 | 16 | LOC10798 | unavailable | False | 31793876 | GTCTCCTAGAGACTGAGATCTG | [] |
| 4910 | 16 | VN1R3 | unavailable | False | 31807925 | GAGAACTATCTGACCAGCTTAG | [] |
| 4911 | 16 | LOC10050 | unavailable | False | 32264028 | GCCTCTAGCCTGCGGCTGCTT | [] |
| 4912 | 16 | LOC10537 | unavailable | False | 33385982 | GCCGGCTCTACCGGCGTCCTT | [] |
| 4913 | 16 | LOC10537 | unavailable | False | 33484764 | GCCGGCTCTACCGGCGTCCTT | [] |
| 4914 | 16 | PHKB | {'exs': 1, 'nonexs': 6} | True | 47699365 | CCCCACTAGCTAATCAGACACC | [] |
| 4915 | 16 | ABCC12 | {'exs': 0, 'nonexs': 7} | True | 48083714 | GACCTCTACAATCTGACTTCTG | [] |
| 4916 | 16 | LONP2 | {'exs': 0, 'nonexs': 7} | True | 48336403 | TCTAAGATGCTATCTATAAAAG | [] |
| 4917 | 16 | LONP2 | {'exs': 0, 'nonexs': 7} | True | 48351801 | CCTACAGTTTGCTATTTAACAG | [] |
| 4918 | 16 | LONP2 | {'exs': 0, 'nonexs': 7} | True | 48361595 | ATGTAACTATTTCCATGTGTTG | [] |
| 4919 | 16 | TENT4B | unavailable | False | 50229327 | GCTGACTATCTACAGAGGTCTG | [] |
| 4920 | 16 | ADCY7 | {'exs': 0, 'nonexs': 5} | True | 50311703 | GGAACAGCTAGGGGTGCCACT | [] |
| 4921 | 16 | NKD1 | {'exs': 0, 'nonexs': 7} | True | 50633780 | GGGGCTCTATGTCTGGTAGAAC | [] |
| 4922 | 16 | NOD2 | {'exs': 0, 'nonexs': 7} | True | 50712373 | CACTCACTACAGAGCCTTGCAC | [] |
| 4923 | 16 | CYLD | {'exs': 2, 'nonexs': 5} | False | 50788853 | CCATATCTAAGTTAGTAAAAAG | ['AAAAAACTATTAATTGATATTG'] |
| 4924 | 16 | LOC10537 | unavailable | False | 51134200 | CTTTTCCTAAACTCTTCTGATG | [] |
| 4925 | 16 | RPGRIP1L | {'exs': 0, 'nonexs': 7} | True | 53607981 | TACAGTCTAGTCTGAGGACCTG | [] |
| 4926 | 16 | RPGRIP1L | {'exs': 0, 'nonexs': 7} | True | 53633336 | TTCCTACCAAAATGCATAACTG | [] |
| 4927 | 16 | FTO | {'exs': 0, 'nonexs': 7} | True | 53965732 | CTGCCCTATGTCGTTGCTCTAC | [] |
| 4928 | 16 | FTO | {'exs': 0, 'nonexs': 7} | True | 54111914 | CCTTCTAGGGTTTTGCTTCCAG | [] |
| 4929 | 16 | IRX3 | {'exs': 0, 'nonexs': 7} | True | 54283685 | AAGAACTAGGATGAGGAGAGAG | [] |
| 4930 | 16 | IRX3 | {'exs': 0, 'nonexs': 7} | True | 54283797 | AGAGGCTGCCTAGGGCGGGGA | [] |
| 4931 | 16 | IRX6 | {'exs': 0, 'nonexs': 7} | True | 55330304 | TTGCGCTAACCTGCTGAGAGAC | [] |
| 4932 | 16 | AMFR | {'exs': 2, 'nonexs': 7} | False | 56362908 | GGGAGCGCTAGGAGGTCTGCT | [] |
| 4933 | 16 | BBS2 | {'exs': 0, 'nonexs': 7} | True | 56494760 | TTCCTCACCTAGGAAGAAGCTG | [] |
| 4934 | 16 | MT1E | {'exs': 0, 'nonexs': 7} | True | 56626820 | CCTGCCCTAATAGCAAGCCTGC | [] |
| 4935 | 16 | CETP | {'exs': 0, 'nonexs': 7} | True | 56983665 | AGACTTCTAGCTCAAGCTCTGG | [] |
| 4936 | 16 | PSME3IP1 | unavailable | False | 57154289 | GAGAAACTAGGGGGCCTCGAG | [] |
| 4937 | 16 | ARL2BP | {'exs': 0, 'nonexs': 7} | True | 57252266 | TAGGACCTAGTGCCGCAGATTC | [] |
| 4938 | 16 | CIAPIN1 | {'exs': 4, 'nonexs': 4} | False | 57429169 | GAACCTCCTAGGCATCATGAAG | [] |
| 4939 | 16 | CCDC102 | {'exs': 0, 'nonexs': 7} | True | 57512740 | AGCTCTAGGCCACCTGGATTTC | [] |
| 4940 | 16 | ADGRG5 | unavailable | False | 57575537 | CCCGGACTACTGTGTTGTTTGC | [] |
| 4941 | 16 | ADGRG1 | unavailable | False | 57663581 | GAGGCCTAGATGCGGCTGGAC | [] |
| 4942 | 16 | ADGRG3 | unavailable | False | 57689460 | GCCTTCCTATTCTTGAGATGCG | [] |
| 4943 | 16 | ADGRG3 | unavailable | False | 57698686 | CAGGACTATAGGAACTTTAGAG | [] |
| 4944 | 16 | CCDC113 | {'exs': 0, 'nonexs': 7} | True | 58279776 | CTCTGGCTACTTCCCAGCAAG | [] |
| 4945 | 16 | PRSS54 | {'exs': 0, 'nonexs': 7} | True | 58290223 | GGACTAGATACTGCTGCAAAAG | [] |
| 4946 | 16 | SETD6 | {'exs': 1, 'nonexs': 7} | True | 58519028 | AGGGAAACTGCTAACTTGTCAG | [] |
| 4947 | 16 | CNOT1 | {'exs': 7, 'nonexs': 2} | False | 58520957 | AGTTTCGTCTAACTGGCACCTG | [] |
| 4948 | 16 | SLC38A7 | {'exs': 0, 'nonexs': 7} | True | 58667305 | TGGACTAAGAATGGCCCCATAG | [] |
| 4949 | 16 | CDH11 | {'exs': 0, 'nonexs': 7} | True | 64948090 | ACAATACTACAATGACTGGAGG | [] |
| 4950 | 16 | CDH5 | {'exs': 1, 'nonexs': 7} | True | 66403168 | CCTCGGCCGCCTAATACAGCA | [] |
| 4951 | 16 | BEAN1 | {'exs': 1, 'nonexs': 6} | True | 66493440 | GGGGCCCTAGAGGACTGGAGA | [] |
| 4952 | 16 | TK2 | {'exs': 2, 'nonexs': 6} | False | 66511967 | CCTCCTATGGGCAATGCTTCC | [] |
| 4953 | 16 | CMTM4 | {'exs': 0, 'nonexs': 7} | True | 66617276 | ACTTTTCTAGGCAAGTTGCCAG | [] |
| 4954 | 16 | NAE1 | {'exs': 5, 'nonexs': 3} | False | 66803008 | TTACTCTACAACTGGAAAGTTG | [] |
| 4955 | 16 | RRAD | {'exs': 0, 'nonexs': 7} | True | 66922075 | TGGGACCTAGAGAACCGAGAG | [] |
| 4956 | 16 | CES2 | {'exs': 0, 'nonexs': 7} | True | 66944024 | CAGGGAGCTACAGCTCTGTGT | [] |
| 4957 | 16 | CES4A | {'exs': 0, 'nonexs': 5} | True | 67006685 | CACCCTACTCCTGCTGCTTCT | [] |
| 4958 | 16 | CBFB | {'exs': 2, 'nonexs': 5} | False | 67098731 | AACCAGGACTAGGGTCTTGTTG | [] |
| 4959 | 16 | C16orf70 | {'exs': 0, 'nonexs': 7} | True | 67147130 | TGTCCCTAGGGGAGTTCCGCT | [] |
| 4960 | 16 | B3GNT9 | {'exs': 0, 'nonexs': 7} | True | 67149276 | GGGGAGCTAGGAGTCCCATTG | [] |
| 4961 | 16 | TRADD | {'exs': 0, 'nonexs': 7} | True | 67154548 | GGTCTAGGCCAGGCCGCCATT | [] |
| 4962 | 16 | NOL3 | {'exs': 2, 'nonexs': 4} | False | 67175096 | CAGGTCCTATCCAGCATGGGC | [] |
| 4963 | 16 | NOL3 | {'exs': 2, 'nonexs': 4} | False | 67175322 | GGACTCCTAAGGGCAGATTAGC | [] |
| 4964 | 16 | TMEM208 | {'exs': 0, 'nonexs': 7} | True | 67229112 | ATGGCTATAACCGCTTCATCTG | [] |
| 4965 | 16 | SLC9A5 | {'exs': 0, 'nonexs': 7} | True | 67271209 | GAGCTACAGCCGGCTGCCTCT | [] |
| 4966 | 16 | KCTD19 | {'exs': 0, 'nonexs': 7} | True | 67289568 | GGCACCCTAGTCCTCTTGTAG | [] |
| 4967 | 16 | LRRC36 | {'exs': 0, 'nonexs': 7} | True | 67385088 | TAGCTCTATAAGCCAGGCTCTG | [] |
| 4968 | 16 | ZDHHC1 | {'exs': 0, 'nonexs': 7} | True | 67394609 | CGGCCCAGCTAAGCCAGACCA | [] |
| 4969 | 16 | ATP6V0D1 | {'exs': 5, 'nonexs': 2} | False | 67438527 | AGGACGCTAGAAGATAGGGATG | [] |
| 4970 | 16 | AGRP | {'exs': 0, 'nonexs': 7} | True | 67482535 | GGCCAGCTAGGTGCGGCTGCA | [] |
| 4971 | 16 | ENKD1 | unavailable | False | 67663655 | GACCTACTAATGTGGCACATGG | [] |
| 4972 | 16 | RANBP10 | {'exs': 0, 'nonexs': 7} | True | 67726427 | CACAGTCAGCTAGTGCAAGTAG | [] |
| 4973 | 16 | TSNAXIP1 | {'exs': 0, 'nonexs': 7} | True | 67827992 | AGTTCCTAGCTTGCAGGCTCT | [] |
| 4974 | 16 | CENPT | {'exs': 2, 'nonexs': 4} | False | 67828266 | TGGCCACTACTGGGCAGGGAA | [] |
| 4975 | 16 | EDC4 | {'exs': 3, 'nonexs': 4} | False | 67884147 | TAGCAGCTAAGGGAGGCTGGG | [] |
| 4976 | 16 | NRN1L | {'exs': 0, 'nonexs': 7} | True | 67886258 | CAAGCTAGGCCAGAGGCCTCA | [] |
| 4977 | 16 | NFATC3 | {'exs': 0, 'nonexs': 7} | True | 68214428 | ACCCTCTAGATTCTCCAGAAAG | [] |
| 4978 | 16 | ESRP2 | {'exs': 2, 'nonexs': 6} | False | 68230225 | CTACAAACACACCCATTCCTTG | [] |
| 4979 | 16 | SLC7A6 | {'exs': 0, 'nonexs': 7} | True | 68297327 | CCTCTAGTCAGTTTTCCTCTCG | ['CCTCTCATCCTTTTTTTCTTCG'] |
| 4980 | 16 | PRMT7 | {'exs': 1, 'nonexs': 7} | True | 68352495 | GCCGCTCTACAGGCACCCAAG | [] |
| 4981 | 16 | SMPD3 | {'exs': 0, 'nonexs': 7} | True | 68361205 | GTCTATGCCTCCTCCTCCCCC | [] |
| 4982 | 16 | ZFP90 | {'exs': 0, 'nonexs': 5} | True | 68563013 | AGTCCCCTATTCCTTCCTGTTG | [] |
| 4983 | 16 | ZFP90 | {'exs': 0, 'nonexs': 5} | True | 68564697 | GGTTCCTAGAGTTTATTTCGAG | [] |
| 4984 | 16 | CDH3 | {'exs': 1, 'nonexs': 6} | True | 68698399 | GGCCGCCTAGTCGTCCTCCCC | [] |
| 4985 | 16 | CDH1 | {'exs': 1, 'nonexs': 6} | True | 68833498 | AGTCCCCTAGTCGTCCTCGCC | [] |
| 4986 | 16 | TANGO6 | unavailable | False | 69083660 | CCAGGTCTACGGCAGGACCAT | [] |
| 4987 | 16 | DERPC | unavailable | False | 69118468 | CACGTTTCTAGAGAGCAGTGAG | [] |
| 4988 | 16 | COG8 | {'exs': 4, 'nonexs': 4} | False | [illegible] | [illegible] | [] |
| [illegible] | [illegible] | [illegible] | [illegible] | [illegible] | [illegible] | [illegible] | [illegible] |
| 4990 | 16 | CLEC18A | {'exs': 0, 'nonexs': 7} | True | 69966055 | CTCACTACAGGGATCCCCTTT | [] |
| 4991 | 16 | PDPR | {'exs': 2, 'nonexs': 6} | False | 70163379 | GGTTGACTATTCCTGGAACTGG | [] |
| 4992 | 16 | COG4 | {'exs': 7, 'nonexs': 1} | False | 70481009 | CAGCTACAGGCGCAGCCTCTT | [] |
| 4993 | 16 | VAC14 | {'exs': 0, 'nonexs': 7} | True | 70712906 | GGTGCTACCTCCCCAGGGCAT | [] |
| 4994 | 16 | VAC14 | {'exs': 0, 'nonexs': 7} | True | 70728761 | GCTATAGATTCCTCCTCCAAAG | [] |
| 4995 | 16 | HYDIN | {'exs': 0, 'nonexs': 7} | True | 70807579 | ACTAAAGGGTGATCCCCTTCAG | [] |
| 4996 | 16 | HYDIN | {'exs': 0, 'nonexs': 7} | True | 71027589 | AATGTGCTATCCCCTTACCCTG | [] |
| 4997 | 16 | TAT | {'exs': 0, 'nonexs': 7} | True | 71568143 | CCTATTTATCACACTCCTCCTG | [] |
| 4998 | 16 | MARVELD | {'exs': 1, 'nonexs': 7} | True | 71640403 | CCACTATCTGCACCACACCTAC | [] |
| 4999 | 16 | IST1 | {'exs': 0, 'nonexs': 6} | True | 71927812 | AGACCTATGTTTTCTTTTTCAG | [] |
| 5000 | 16 | PKD1L3 | {'exs': 1, 'nonexs': 0} | False | 71944113 | TGGGGCTACATAGACGGGGTT | [] |
| 5001 | 16 | TXNL4B | {'exs': 1, 'nonexs': 7} | True | 72086636 | AATGTACTAAATGTCTTGATAG | [] |
| 5002 | 16 | PMFBP1 | {'exs': 0, 'nonexs': 6} | True | 72119337 | TGTGGATTCTAGCAGTATGAGG | [] |
| 5003 | 16 | GLG1 | {'exs': 0, 'nonexs': 7} | True | 74453166 | GGTGGCTCTACCTGTCCTTGA | [] |
| 5004 | 16 | MLKL | {'exs': 2, 'nonexs': 6} | False | 74672503 | GATACACTACTTAGAAAAGGTG | [] |
| 5005 | 16 | WDR59 | {'exs': 3, 'nonexs': 5} | False | 74912146 | ATCCATCTAGACAATGATGCAG | [] |
| 5006 | 16 | CFDP1 | {'exs': 4, 'nonexs': 4} | False | 75372546 | AGATCAATCTCTAGGCATCAAG | ['TCAAATTTCTTCTTCCTGAAAG'] |
| 5007 | 16 | TMEM170 | {'exs': 0, 'nonexs': 7} | True | 75447557 | GCTATAGAGTAGCTAAAATCCG | [] |
| 5008 | 16 | CHST6 | {'exs': 0, 'nonexs': 7} | True | 75478640 | CTCCACTAATTTCGGGGGTGCC | [] |
| 5009 | 16 | TMEM231 | {'exs': 0, 'nonexs': 7} | True | 75539993 | AATGGCCTTTCTAGGATAAGTG | [] |
| 5010 | 16 | KARS1 | unavailable | False | 75627894 | TATTATTTTCTAGACAGAAGTG | [] |
| 5011 | 16 | SYCE1L | {'exs': 0, 'nonexs': 7} | True | 77212930 | GCTGGCCTAGAGGGCGTCAGG | [] |
| 5012 | 16 | CLEC3A | {'exs': 0, 'nonexs': 5} | True | 78030840 | AAAGACCTATTGAGGGATGGTG | [] |
| 5013 | 16 | WWOX | {'exs': 1, 'nonexs': 6} | True | 78278655 | CTAGTGGAAAATTTTTATTCTG | [] |
| 5014 | 16 | DYNLRB2 | {'exs': 0, 'nonexs': 7} | True | 80550057 | AGGTCTATTCACATGGATTCTG | [] |
| 5015 | 16 | CENPN | {'exs': 4, 'nonexs': 4} | False | 81022837 | TTTCATGTTCACTAATTGCAAG | [] |
| 5016 | 16 | C16orf46 | {'exs': 0, 'nonexs': 7} | True | 81054070 | CTACTTTATCTTCTTTTTCCTG | [] |
| 5017 | 16 | C16orf46 | {'exs': 0, 'nonexs': 7} | True | 81057916 | CCCAGCTACTCAGGGGGCTGA | [] |
| 5018 | 16 | PLCG2 | {'exs': 0, 'nonexs': 7} | True | 81957997 | CAGCTTCTATGAGTAAAACTTG | [] |
| 5019 | 16 | HSD17B2 | {'exs': 0, 'nonexs': 7} | True | 82098435 | TGCCTAGGTGGCCTTTTTCTTG | [] |
| 5020 | 16 | DNAAF1 | unavailable | False | 84177840 | GAAAACTATGATGCTTTCGGTG | [] |
| 5021 | 16 | ATP2C2 | {'exs': 2, 'nonexs': 5} | False | 84463731 | GGTCCACTACACATCTTCAGGC | [] |
| 5022 | 16 | ZDHHC7 | {'exs': 0, 'nonexs': 7} | True | 84988546 | CACCTTCTACCCTGACTTCCTC | [] |
| 5023 | 16 | KIAA0513 | {'exs': 0, 'nonexs': 5} | True | 85088324 | GGGCCTACTCAGTGGCCATTTC | [] |
| 5024 | 16 | FAM92B | {'exs': 2, 'nonexs': 4} | False | 85298534 | GCCAGCTAAGTTCTTCTGACTG | [] |
| 5025 | 16 | GINS2 | {'exs': 7, 'nonexs': 1} | False | 85678211 | TTCTCTAGAAGTCCTGAGACTG | [] |
| 5026 | 16 | EMC8 | unavailable | False | 85779707 | GCCTGCCTAGCACAAGTGTAG | [] |
| 5027 | 16 | COX4I1 | {'exs': 6, 'nonexs': 2} | False | 85806501 | AGGTCCTCTCTAAAGGCTATGG | [] |
| 5028 | 16 | SLC7A5 | {'exs': 4, 'nonexs': 4} | False | 87832969 | TCCTGGCTATGTCTCCTGGGG | [] |
| 5029 | 16 | CA5A | {'exs': 0, 'nonexs': 7} | True | 87888128 | CTAATGTCTCTAGGACCTTGTG | [] |
| 5030 | 16 | CA5A | {'exs': 0, 'nonexs': 7} | True | 87923914 | AGGAACTATTTTTGGTTTTTCG | ['GTTTTTCAATATTTTGAAGTGG'] |
| 5031 | 16 | LOC10799 | unavailable | False | 88263090 | TGAGAACTAGGAGTCTCCAGGC | [] |
| 5032 | 16 | ZC3H18 | {'exs': 7, 'nonexs': 2} | False | 88631298 | CACGGCCTATGCTTTCCCGGG | [] |
| 5033 | 16 | CYBA | {'exs': 0, 'nonexs': 7} | True | 88645165 | AGCCACTAGCTGTGCGTGGCA | [] |
| 5034 | 16 | RNF166 | {'exs': 2, 'nonexs': 7} | False | 88697935 | ACCTCTAACAATTTCCATCCTG | [] |
| 5035 | 16 | RNF166 | {'exs': 2, 'nonexs': 7} | False | 88700619 | CCTCTAGCTGCTCAGTCCTCC | [] |
| 5036 | 16 | PIEZO1 | unavailable | False | 88715604 | AGCTCCTACTCCTTCTCACGAC | [] |
| 5037 | 16 | GALNS | {'exs': 0, 'nonexs': 7} | True | 88814438 | CAGGTGCTAGTGGGACCAGAG | [] |
| 5038 | 16 | GALNS | {'exs': 0, 'nonexs': 7} | True | 88817171 | GCCACCCTATGGGACCCCCAA | [] |
| 5039 | 16 | LOC10798 | unavailable | False | 88824721 | CCCCATCTCTATCCAGACGTGC | [] |
| 5040 | 16 | CBFA2T3 | {'exs': 3, 'nonexs': 5} | False | 88878276 | TGGTGGTCTAGCTTCTCCCTAC | [] |
| 5041 | 16 | LOC10537 | unavailable | False | 88930532 | CGCACTGGAGCCTAGTGGGGA | [] |
| 5042 | 16 | ACSF3 | {'exs': 0, 'nonexs': 7} | True | 89159010 | AGGCTCTAGAAGGGCCACCAA | [] |
| 5043 | 16 | SLC22A31 | {'exs': 1, 'nonexs': 0} | False | 89195998 | GCAGGACTAGTGCTGCTCGGG | [] |
| 5044 | 16 | ANKRD11 | {'exs': 3, 'nonexs': 5} | False | 89270555 | TCAAGCCTAGGGTCCTGTTGA | [] |
| 5045 | 16 | SPG7 | {'exs': 0, 'nonexs': 7} | True | 89541010 | AGCTCTTCTAGTCGCTGTGTAG | [] |
| 5046 | 16 | SPG7 | {'exs': 0, 'nonexs': 7} | True | 89549619 | CCCGTTCCTACTGACGGTCGG | [] |
| 5047 | 16 | SPG7 | {'exs': 0, 'nonexs': 7} | True | 89557092 | CCCAACTACTTGGGCCAAGTC | [] |
| 5048 | 16 | CPNE7 | {'exs': 2, 'nonexs': 6} | False | 89596537 | CTACGCCCTCCACATCTTCAC | [] |
| 5049 | 16 | CHMP1A | {'exs': 0, 'nonexs': 7} | True | 89645913 | AACAACCCTAAGGCCACGCAG | [] |
| 5050 | 16 | CHMP1A | {'exs': 0, 'nonexs': 7} | True | 89646065 | GCACGGCTAGTTCCTCAAGGC | [] |
| 5051 | 16 | CHMP1A | {'exs': 0, 'nonexs': 7} | True | 89647483 | CTATGGAAAGACCACCAACTCG | [] |

| | | | | | | | |
|---|---|---|---|---|---|---|---|
| 5052 | 16 | SPATA2L | {'ess': 0, 'noness': 7} | True | 89697233 | GCTGGCCTAGGGCCGGGCCC( | [] |
| 5053 | 16 | VPS9D1 | unavailable | False | 89707860 | CTAGCTGTACTACTTGGCCAG( | [] |
| 5054 | 16 | ZNF276 | {'ess': 0, 'noness': 7} | True | 89729966 | CCAACTACCTGGGAGTGTTCT( | [] |
| 5055 | 16 | DEF8 | {'ess': 0, 'noness': 7} | True | 89963820 | TCCCTACTTGAAAGGCACTAAG | [] |
| 5056 | 16 | DEF8 | {'ess': 0, 'noness': 7} | True | 89965962 | TCGGCGCTAGGCCTCCACATC | [] |
| 5057 | 16 | CENPBD1 | {'ess': 0, 'noness': 7} | True | 89971358 | TCATCTACGTTAAAGACACCTG | [] |
| 5058 | 16 | DBNDD1 | {'ess': 0, 'noness': 7} | True | 90006334 | AGATGGTCTAGTCCTCCTGGG( | [] |
| 5059 | 16 | GAS8 | {'ess': 1, 'noness': 6} | True | 90043344 | GGGGGGCAGCTACGTCGGGGT | [] |
| 5060 | 17 | DOC2B | {'ess': 0, 'noness': 1} | False | 160689 | CAGCCCTATGAACCAGGTGAA( | [] |
| 5061 | 17 | VPS53 | {'ess': 5, 'noness': 4} | False | 519127 | CTGCTACAGTCTCTTTTTAATG | [] |
| 5062 | 17 | VPS53 | {'ess': 5, 'noness': 4} | False | 532826 | GATACTACGTCCATCTCACCTG | [] |
| 5063 | 17 | NXN | {'ess': 0, 'noness': 7} | True | 800948 | AGCCACGCTAGATGGGCTCCG | [] |
| 5064 | 17 | ABR | {'ess': 1, 'noness': 7} | True | 1006079 | CTCGGGCTACACGTCGGTGGA | [] |
| 5065 | 17 | ABR | {'ess': 1, 'noness': 7} | True | 1033431 | TGAGCTACGTGTCCAAGGCCA( | [] |
| 5066 | 17 | ABR | {'ess': 1, 'noness': 7} | True | 1228146 | GATGGGCCTAAGCCTCCCCTT( | [] |
| 5067 | 17 | SLC43A2 | {'ess': 0, 'noness': 7} | True | 1575603 | GCCACTACACGAAGGCCTCCT( | [] |
| 5068 | 17 | SCARF1 | {'ess': 2, 'noness': 6} | False | 1635503 | CCCGAGCTAGGCTGGAGGTGC | [] |
| 5069 | 17 | TLCD2 | {'ess': 0, 'noness': 7} | True | 1707769 | TTCTCTCTAGTCTTTCAGGCTG | [] |
| 5070 | 17 | WDR81 | {'ess': 0, 'noness': 7} | True | 1737584 | CTCAGTCTATGCCAGGAGGCG( | [] |
| 5071 | 17 | SMYD4 | {'ess': 0, 'noness': 7} | True | 1781285 | CCCTACAATGCAGGCCCTACA( | [] |
| 5072 | 17 | SMYD4 | {'ess': 0, 'noness': 7} | True | 1782503 | CTCACTATAGACAGAAGCCCAG | [] |
| 5073 | 17 | SMYD4 | {'ess': 0, 'noness': 7} | True | 1784325 | CTCACCTAGTTCACCATCTCTG | [] |
| 5074 | 17 | HIC1 | {'ess': 0, 'noness': 7} | True | 2058834 | CGCTCTAGGTGGGAGAGAAAC( | [] |
| 5075 | 17 | SMG6 | {'ess': 7, 'noness': 1} | False | 2274369 | GTCTTCTACTCCAAAAACTCAG | [] |
| 5076 | 17 | METTL16 | unavailable | False | 2419969 | GTTTCTAGTTAACTGCAACAAG | [] |
| 5077 | 17 | CLUH | unavailable | False | 2690593 | CCTCTCTATCCCTGCACGCTC( | [] |
| 5078 | 17 | CCDC92B | unavailable | False | 2724410 | TGGCGCCTACTCCGGGTCCCC | [] |
| 5079 | 17 | OR1G1 | {'ess': 0, 'noness': 3} | True | 3126609 | GTCACTACCACTAAGGGGAATG | [] |
| 5080 | 17 | OR1A2 | {'ess': 0, 'noness': 3} | True | 3198447 | CGGCTATGAGGAGATTCTCTTG | [] |
| 5081 | 17 | ASPA | {'ess': 0, 'noness': 7} | True | 3499087 | TGATTTCTAATGTAAACAGCAG | [] |
| 5082 | 17 | TRPV3 | {'ess': 0, 'noness': 7} | True | 3513916 | CGCTTCTACACCGAGGTTTCC( | [] |
| 5083 | 17 | SHPK | {'ess': 0, 'noness': 7} | True | 3610559 | TGCTGTCTAAGATTCCTTCTGG | [] |
| 5084 | 17 | CTNS | {'ess': 1, 'noness': 7} | True | 3660368 | CCCTGGGTGCTAGTTCAGCTG( | [] |
| 5085 | 17 | TAX1BP3 | {'ess': 0, 'noness': 7} | True | 3663747 | GCTGCTAGGACAGCATGGACT( | [] |
| 5086 | 17 | NCBP3 | unavailable | False | 3812545 | CCCACACTAGGGTCCCGGAAG | [] |
| 5087 | 17 | ATP2A3 | {'ess': 2, 'noness': 7} | False | 3927730 | GTGAGCCCCTACATGTGTCTG( | [] |
| 5088 | 17 | ZZEF1 | {'ess': 0, 'noness': 1} | False | 4006889 | GGGCTAACACTCCACATTCCA( | [] |
| 5089 | 17 | ANKFY1 | {'ess': 0, 'noness': 7} | True | 4167778 | CTCACTAAGAAACCCCACCCA( | [] |
| 5090 | 17 | PELP1 | {'ess': 4, 'noness': 2} | False | 4671438 | GATGGCTAGGAGTCAGGCTCT( | [] |
| 5091 | 17 | ARRB2 | {'ess': 0, 'noness': 7} | True | 4721038 | ACCCCGCTTCCTAGCAGAGTT( | [] |
| 5092 | 17 | MED11 | {'ess': 5, 'noness': 3} | False | 4733186 | TGGCCTAGTTCTCCAGCATCT( | [] |
| 5093 | 17 | PLD2 | {'ess': 0, 'noness': 7} | True | 4822863 | TCAACTATGTCCACACTTCTAG | [] |
| 5094 | 17 | CHRNE | {'ess': 0, 'noness': 7} | True | 4898735 | GCGAGCTAAGGCTGGATACA( | [] |
| 5095 | 17 | C17orf107 | {'ess': 0, 'noness': 7} | True | 4900532 | CTGGCCCCTAACTCACCCCGT | [] |
| 5096 | 17 | SPAG7 | {'ess': 1, 'noness': 6} | True | 4959533 | GGGGCGCCTAGGAGGTTGGCG | [] |
| 5097 | 17 | ZNF594 | {'ess': 0, 'noness': 7} | True | 5181832 | GAATTCTATGATGTCTCAGAAG | [] |
| 5098 | 17 | SCIMP | unavailable | False | 5214726 | CCCAGCTACTCCGGAGGCTGA | [] |
| 5099 | 17 | RABEP1 | {'ess': 0, 'noness': 6} | True | 5336531 | TCTAAGGAGACACAACTAAATG | [] |
| 5100 | 17 | RPAIN | {'ess': 5, 'noness': 3} | False | 5432570 | AGCTGGCTCTAGAGGATCACA( | [] |
| 5101 | 17 | C1QBP | {'ess': 5, 'noness': 3} | False | 5433014 | CTGTCTGCTCTACTGGCTCTT( | [] |
| 5102 | 17 | NLRP1 | {'ess': 0, 'noness': 7} | True | 5501813 | TGGGCACTAGTATCTCCTGGC( | [] |
| 5103 | 17 | PITPNM3 | {'ess': 0, 'noness': 7} | True | 6463539 | GCTGAGGTCTAGAAAGCAGGA( | [] |
| 5104 | 17 | SLC13A5 | {'ess': 0, 'noness': 7} | True | 6686206 | TCTTCCTAAGTCTCAATATGTG | [] |
| 5105 | 17 | XAF1 | {'ess': 0, 'noness': 7} | True | 6772168 | TCTAGCTGAAATTTCTCACTTG | [] |
| 5106 | 17 | RNASEK | {'ess': 1, 'noness': 5} | False | 7014285 | GCCGGGGCCCTAGCGCACCAT | [] |
| 5107 | 17 | BCL6B | {'ess': 0, 'noness': 7} | True | 7027618 | CGCTCAGCTAGGGCCCCCCGA | [] |
| 5108 | 17 | DVL2 | {'ess': 0, 'noness': 7} | True | 7225864 | GGGCTACATAACATCCACAAAG | [] |
| 5109 | 17 | ELP5 | unavailable | False | 7258787 | TGCCCCATCCCTAGAATCCCT( | [] |
| 5110 | 17 | CLDN7 | {'ess': 0, 'noness': 7} | True | 7260486 | AGGATGACTAGGGCAGACCCT( | [] |
| 5111 | 17 | TMEM95 | {'ess': 0, 'noness': 7} | True | 7356720 | CTGGGACTAAGGGGCTGACTG( | [] |
| 5112 | 17 | PLSCR3 | unavailable | False | 7390384 | GGCCTCTAACTGGTGACGGCA( | [] |
| 5113 | 17 | NLGN2 | {'ess': 0, 'noness': 7} | True | 7417798 | CACCCCCTATACCCGAGTGGT( | [] |
| 5114 | 17 | MPDU1 | {'ess': 2, 'noness': 5} | False | 7587550 | GGCTCTACTGCGCCTTTTTCT( | [] |
| 5115 | 17 | DNAH2 | {'ess': 0, 'noness': 5} | True | 7818214 | CTGTCCTAAGGCCCCAGGCCA | [] |
| 5116 | 17 | DNAH2 | {'ess': 0, 'noness': 5} | True | 7820700 | ATTCCCCTACTGTCTTGCCAG( | [] |
| 5117 | 17 | TMEM88 | {'ess': 0, 'noness': 7} | True | 7855940 | CCAGCCTAGATGTTCAGGATTG | [] |
| 5118 | 17 | CYB5D1 | {'ess': 0, 'noness': 7} | True | 7859611 | TCCTTGCCTACAACTCCGTGA( | [] |
| 5119 | 17 | CHD3 | {'ess': 0, 'noness': 7} | True | 7902876 | TGCTCCTAGATGTTCTCTGATG | [] |
| 5120 | 17 | CHD3 | {'ess': 0, 'noness': 7} | True | 7911703 | GGGGTTCCTAGCCCAGTGATG( | [] |
| 5121 | 17 | KCNAB3 | {'ess': 0, 'noness': 7} | True | 7923101 | ACAGACTACTTCTTGGAATGCG | [] |
| 5122 | 17 | ALOX12B | {'ess': 0, 'noness': 7} | True | 8072770 | GCGCTCCTAAATAGAAATGCTG | [] |
| 5123 | 17 | PER1 | {'ess': 0, 'noness': 7} | True | 8141067 | GAATGGAGTCTAGCTGGTGCA( | [] |
| 5124 | 17 | PER1 | {'ess': 0, 'noness': 7} | True | 8144583 | CATCCTAGTGGGGAGAAGCTA( | [] |
| 5125 | 17 | SLC25A35 | {'ess': 0, 'noness': 7} | True | 8289913 | TTCACGATCCTAACATCTTTGG | [] |
| 5126 | 17 | SPDYE4 | {'ess': 0, 'noness': 7} | True | 8753137 | GGAGCTCTAGGAAATGAGGGT( | [] |
| 5127 | 17 | PIK3R5 | {'ess': 0, 'noness': 7} | True | 8880638 | ACACTAGGGCAGAGCTCCACT( | [] |
| 5128 | 17 | NTN1 | {'ess': 0, 'noness': 5} | True | 9239967 | GCGCTAGGCCTTCTTGCACTT( | [] |
| 5129 | 17 | GSG1L2 | unavailable | False | 9802385 | TGGCTAGCATATGGACACCTTG | [] |
| 5130 | 17 | GLP2R | {'ess': 0, 'noness': 7} | True | 9889704 | CCTAGATCTCACTCTCTTCCAG | [] |
| 5131 | 17 | GAS7 | {'ess': 1, 'noness': 6} | True | 9917227 | CAGGCCCATCTAGATCTCCAT( | [] |
| 5132 | 17 | SCO1 | {'ess': 4, 'noness': 4} | False | 10681118 | CTTTGGCTAGCTCTTTTTTCTG | [] |
| 5133 | 17 | ADPRM | unavailable | False | 10711143 | AAATTAGACTAACAATGGAAGG | [] |
| 5134 | 17 | DNAH9 | {'ess': 0, 'noness': 7} | True | 11969526 | CAGGATGCTAAATCTGGAGAAG | [] |
| 5135 | 17 | ZNF18 | {'ess': 0, 'noness': 7} | True | 11977956 | TGGTTACTGGCTATTGAAAGGG | [] |
| 5136 | 17 | MYOCD | {'ess': 0, 'noness': 7} | True | 12763643 | GGCATTCTACCACTGCTGCAA( | [] |
| 5137 | 17 | TEKT3 | {'ess': 0, 'noness': 5} | True | 15303835 | GTCCCTAGCAGAAGCCGACCA( | [] |
| 5138 | 17 | CDRT1 | {'ess': 0, 'noness': 1} | False | 15588974 | GCTGCTACACACCACTGAAAAG | [] |
| 5139 | 17 | TRIM16 | {'ess': 1, 'noness': 7} | True | 15628614 | GCTCCTGGAGTCTAGGGAGCA( | [] |
| 5140 | 17 | TTC19 | {'ess': 1, 'noness': 5} | False | 16009005 | GTCTCTATCCTTCTTTGATATG | [] |
| 5141 | 17 | TTC19 | {'ess': 1, 'noness': 5} | False | 16044857 | ATCTACTAGTCTCCTGATCCTG | [] |
| 5142 | 17 | NCOR1 | {'ess': 1, 'noness': 6} | True | 16091854 | CTACCTACCTTCCCACTCTCT( | [] |
| 5143 | 17 | PIGL | {'ess': 0, 'noness': 7} | True | 16317083 | AGTTCCTAACTCAGGCAGATTG | [] |
| 5144 | 17 | TRPV2 | {'ess': 0, 'noness': 7} | True | 16434976 | GTCACCAGGCTACTCACGAGG( | [] |
| 5145 | 17 | LRRC75A | unavailable | False | 16446922 | CATCTCCTAGTTGTGGGTGGG( | [] |
| 5146 | 17 | CCDC144 | {'ess': 0, 'noness': 6} | True | 16764243 | AAGTAACTATTTTACCAAATAG | ['TAACATACAAGAATTCTATCAG'] |
| 5147 | 17 | CCDC144 | {'ess': 0, 'noness': 6} | True | 16772867 | CATATGCTATAACCTGAAAAAG | [] |
| 5148 | 17 | CCDC144 | {'ess': 0, 'noness': 6} | True | 16772528 | GAGAAGCTATATAGGATCCAGG | [] |
| 5149 | 17 | FAM106C | unavailable | False | 16789409 | ATATTACACCTAACGTTCTGTG | [] |
| 5150 | 17 | MPRIP | {'ess': 1, 'noness': 6} | True | 17184893 | ACACACCTAGTCTTTCTTCAAG | [] |
| 5151 | 17 | FLCN | {'ess': 2, 'noness': 6} | False | 17207047 | GAGCAGCTAGGCTGGGCTGGG | [] |
| 5152 | 17 | MED9 | {'ess': 4, 'noness': 4} | False | 17491494 | ACTCTACTCCTTGGGGATTTCG | [] |
| 5153 | 17 | RASD1 | {'ess': 0, 'noness': 7} | True | 17495124 | GGGCTCCTAGCTGATGACGCA( | [] |
| 5154 | 17 | RAI1 | {'ess': 0, 'noness': 7} | True | 17804093 | ATGCTTCTATCTTCCCTCACTG | [] |
| 5155 | 17 | RAI1 | {'ess': 0, 'noness': 7} | True | 17809880 | TGGATTACTACGGCAGCCTCT( | [] |
| 5156 | 17 | SREBF1 | {'ess': 2, 'noness': 5} | False | 17812036 | CACTAGTCAGCACATCCATCAG | [] |
| 5157 | 17 | SREBF1 | {'ess': 2, 'noness': 5} | False | 17812621 | GGGGTCTAGCTGGAAGTGACA( | [] |
| 5158 | 17 | DRC3 | unavailable | False | 18016670 | ATTCATCTAGTCTAGGATGTCG | [] |
| 5159 | 17 | ATPAF2 | {'ess': 2, 'noness': 5} | False | 18020959 | GGCCTACTTGACAATACTTCTG | [] |
| 5160 | 17 | GID4 | unavailable | False | 18057025 | TCTCTGATTCTAGTTTATAATG | [] |
| 5161 | 17 | MIEF2 | unavailable | False | 18264763 | ACCCACCTAGAGCAGCCCCTC | [] |
| 5162 | 17 | EVPLL | {'ess': 0, 'noness': 7} | True | 18387084 | CCGTCTCTACTAAAAATACAAG | ['ACAAAAAAAAAATTAGCCATG'] |
| 5163 | 17 | LGALS9C | {'ess': 0, 'noness': 7} | True | 18494366 | AGCCGCCTATGTCTGCACGTG( | [] |
| 5164 | 17 | FAM106A | {'ess': 0, 'noness': 2} | False | 18526293 | ATATTACACCTAACGTTCTGTG | [] |
| 5165 | 17 | FOXO3B | unavailable | False | 18672308 | GTTCTTCTAGCCGGCAGAGCT( | [] |
| 5166 | 17 | TRIM16L | {'ess': 0, 'noness': 7} | True | 18735459 | GCTCCTGGAGTCTAGGGAGCA( | [] |
| 5167 | 17 | FBXW10 | {'ess': 1, 'noness': 7} | True | 18771674 | ATTACTATTGCTGCTGGGAATG | [] |
| 5168 | 17 | FAM83G | {'ess': 0, 'noness': 5} | True | 18971358 | ATGCTGCTAGGGGTCTTTGCG( | [] |
| 5169 | 17 | GRAPL | {'ess': 0, 'noness': 6} | True | 19158375 | CCGGCTCTACCCCAATCTGCT( | [] |
| 5170 | 17 | EPN2 | {'ess': 0, 'noness': 7} | True | 19334253 | CAGGCACTAGAGAAGGAAAGG( | [] |
| 5171 | 17 | B9D1 | {'ess': 0, 'noness': 7} | True | 19337719 | CACTGCTATCTTTGACAGACAG | [] |
| 5172 | 17 | B9D1 | {'ess': 0, 'noness': 7} | True | 19347138 | CTGTTTCTATTTGTCCTCAGTG | [] |
| 5173 | 17 | SLC47A2 | {'ess': 0, 'noness': 7} | True | 19678685 | TTTGCTAGTGCCTGGTGGCTA( | [] |
| 5174 | 17 | SLC47A2 | {'ess': 0, 'noness': 7} | True | 19703087 | CTGATTACCTACTCATCATTGG | [] |
| 5175 | 17 | SLC47A2 | {'ess': 0, 'noness': 7} | True | 19705192 | ATGATACTAATTAATTTTTAAG | ['TTTTAAAATTCCAACATCTGTG'] |
| 5176 | 17 | SPECC1 | {'ess': 1, 'noness': 7} | True | 20236880 | TCTGCTAGCTGACAGAGCCCA( | [] |
| 5177 | 17 | SPECC1 | {'ess': 1, 'noness': 7} | True | 20312828 | GGTGATGCTAAAAAATTATCAG | [] |
| 5178 | 17 | FAM106B | {'ess': 0, 'noness': 1} | False | 20417958 | ATATTACACCTAACGTTCTGTG | [] |
| 5179 | 17 | LGALS9B | {'ess': 0, 'noness': 7} | True | 20449972 | AGCCGCCTATGTCTGCACGTG( | [] |
| 5180 | 17 | USP22 | {'ess': 1, 'noness': 6} | True | 21003030 | ATAAGGCTACTCGTATTCCAGG | [] |
| 5181 | 17 | DHRS7B | {'ess': 0, 'noness': 7} | True | 21191152 | TCAGAGTACTAGGAGTTCTTGG | [] |
| 5182 | 17 | MAP2K3 | {'ess': 2, 'noness': 6} | False | 21314229 | GCCCCTATGAGTCTTCTCCCA( | [] |
| 5183 | 17 | WSB1 | {'ess': 1, 'noness': 6} | True | 27312368 | AATCTTCTAAATACGATACGAG | [] |
| 5184 | 17 | KSR1 | {'ess': 0, 'noness': 5} | True | 27621934 | CCAGGCCTACAACCTACAGAG( | [] |
| 5185 | 17 | KSR1 | {'ess': 0, 'noness': 5} | True | 27623391 | TGGCTACATCTTTGGATTACTG | [] |
| 5186 | 17 | LGALS9 | {'ess': 1, 'noness': 6} | True | 27648981 | AGCCGCCTATGTCTGCACATG( | [] |
| 5187 | 17 | IFT20 | {'ess': 0, 'noness': 7} | True | 28326629 | ACTATAAAAGCGAAATTTTCAG | [] |
| 5188 | 17 | TNFAIP1 | {'ess': 0, 'noness': 7} | True | 28344702 | GGGTGCTAGAGCAGAGACCCA( | [] |
| 5189 | 17 | POLDIP2 | unavailable | False | 28348116 | CTGGCCTACCAGTGAAGGCCT( | [] |
| 5190 | 17 | SEBOX | {'ess': 0, 'noness': 4} | True | 28364267 | AGACTAGGAGTGGTCCACATTG | [] |
| 5191 | 17 | VTN | {'ess': 0, 'noness': 1} | False | 28367368 | GACTCCTACAGATGGCCAGGA( | [] |
| 5192 | 17 | SLC13A2 | {'ess': 1, 'noness': 7} | True | 28497268 | CCCAGCCTAGGGGCTTGGTGT | [] |
| 5193 | 17 | SUPT6H | {'ess': 7, 'noness': 1} | False | 28701624 | GGCCCCCTACCGATCCATCTC | [] |
| 5194 | 17 | FAM222B | unavailable | False | 28758269 | GGCTACCTATCTATACCCTGG( | [] |
| 5195 | 17 | PHF12 | {'ess': 1, 'noness': 7} | True | 28906182 | CCAGTCCTAAGGAACAGAGTTG | [] |
| 5196 | 17 | SEZ6 | {'ess': 0, 'noness': 7} | True | 28955848 | CCCCACCTAGATGGAGACTTC( | ['CTTCATATTCTCTCGTCTCCTG'] |
| 5197 | 17 | MYO18A | {'ess': 1, 'noness': 7} | True | 29074769 | CTGGGCTATGCGTTAGTCTCC( | [] |
| 5198 | 17 | CRYBA1 | {'ess': 0, 'noness': 7} | True | 29254348 | AATCAGCTACTGTTGGATTCGG | [] |
| 5199 | 17 | ANKRD13B | {'ess': 1, 'noness': 7} | True | 29613581 | GGGCGCTACTGCTCGGTCAGT | [] |
| 5200 | 17 | CORO6 | {'ess': 0, 'noness': 5} | True | 29615731 | CGCGCGGGGCTAGTCCGTGCC | [] |
| 5201 | 17 | EFCAB5 | {'ess': 0, 'noness': 7} | True | 30093628 | AATAAGCTAGGATAGTTCCCCG | [] |
| 5202 | 17 | CRLF3 | {'ess': 0, 'noness': 7} | True | 30784186 | CCAAACATCTAAAACACTAACG | ['TAACACTTTCCATCCAGGATAG'] |
| 5203 | 17 | TEFM | unavailable | False | 30899168 | CTCAGAATTCTAAGGCTGAGAG | [] |
| 5204 | 17 | LOC10799 | unavailable | False | 31035502 | CAGATTCAACTAGATGAGGCTG | [] |
| 5205 | 17 | EVI2A | {'ess': 0, 'noness': 7} | True | 31318302 | TCTTCACTAACCTATCTGTTTG | [] |
| 5206 | 17 | RHOT1 | {'ess': 0, 'noness': 7} | True | 32212735 | CGTATGTTGTTCTAAAGGGCAG | [] |
| 5207 | 17 | PSMD11 | {'ess': 8, 'noness': 1} | False | 32480630 | ACTCACCTATGTCAGTTTCTTG | [] |
| 5208 | 17 | TMEM98 | {'ess': 0, 'noness': 7} | True | 32940992 | GTAGGCACTAAATTGCAGACTG | [] |
| 5209 | 17 | SPACA3 | {'ess': 0, 'noness': 7} | True | 32997777 | CCGTCCATCCTAGAAGTCACA( | [] |
| 5210 | 17 | TMEM132E | {'ess': 0, 'noness': 7} | True | 34638231 | GCGCCTCTATGCAATCTCTTTG | [] |
| 5211 | 17 | ZNF830 | {'ess': 5, 'noness': 3} | False | 34962684 | TCTTTAAAACCCTATAACAATG | [] |
| 5212 | 17 | LIG3 | {'ess': 3, 'noness': 5} | False | 35003047 | CTGTTCTATCTCCTGCCTGCT( | [] |
| 5213 | 17 | LIG3 | {'ess': 3, 'noness': 5} | False | 35004505 | AAACCTAGCAGGGAGCTACCA( | [] |
| 5214 | 17 | UNC45B | {'ess': 0, 'noness': 7} | True | 35186558 | TGTCTAAGACACTGGTTTAATG | [] |
| 5215 | 17 | SLFN11 | {'ess': 0, 'noness': 7} | True | 35362355 | TCTTCCTAATGGCCACCCCAC( | [] |
| 5216 | 17 | LOC10537 | unavailable | False | 35374436 | CGGGTGCTAAAGACTGCAGCG( | [] |
| 5217 | 17 | RDM1 | {'ess': 0, 'noness': 4} | True | 35918341 | AGAAATGCTAGTCAAGTTCTGG | [] |
| 5218 | 17 | TBC1D3B | {'ess': 4, 'noness': 0} | False | 36165994 | ATGCTTCTAGAAGCCTGGAGG( | [] |
| 5219 | 17 | TBC1D3I | unavailable | False | 36253953 | ATGCTTCTAGAAGCCTGGAGG( | [] |
| 5220 | 17 | TBC1D3G | {'ess': 3, 'noness': 1} | False | 36334443 | ATGCTTCTAGAAGCCTGGAGG( | [] |
| 5221 | 17 | TBC1D3H | {'ess': 2, 'noness': 2} | False | 36377845 | ATGCTTCTAGAAGCCTGGAGG( | [] |
| 5222 | 17 | TBC1D3F | {'ess': 4, 'noness': 0} | False | 36428926 | ATGCTTCTAGAAGCCTGGAGG( | [] |
| 5223 | 17 | ZNHIT3 | {'ess': 2, 'noness': 4} | False | 36497765 | TCTACTAAAAATACAAAAATTG | [] |
| 5224 | 17 | MYO19 | {'ess': 0, 'noness': 4} | True | 36513368 | AAAAGCTCTATGCAAAGGGCTG | [] |
| 5225 | 17 | DHRS11 | {'ess': 0, 'noness': 4} | True | 36600202 | AGTCACTAGGTCACCTGCTCC( | [] |
| 5226 | 17 | LHX1 | {'ess': 0, 'noness': 4} | True | 36943130 | AGACCCCGCTACCACACGGCC | [] |
| 5227 | 17 | ACACA | {'ess': 3, 'noness': 4} | False | 37087315 | AGCTCTTCCTACGTGGAAGGG( | [] |
| 5228 | 17 | DUSP14 | {'ess': 0, 'noness': 4} | True | 37512868 | TGGCACTAAATCCCCCAGTAAG | [] |
| 5229 | 17 | HNF1B | {'ess': 0, 'noness': 4} | True | 37721284 | GAAGGCTTCTAACAGCACACAG | [] |
| 5230 | 17 | TBC1D3K | unavailable | False | 37835055 | ATGCTTCTAGAAGCCTGGAGG( | [] |
| 5231 | 17 | TBC1D3L | unavailable | False | 37976459 | ATGCTTCTAGAAGCCTGGAGG( | [] |
| 5232 | 17 | TBC1D3D | unavailable | False | 38014592 | ATGCTTCTAGAAGCCTGGAGG( | [] |
| 5233 | 17 | TBC1D3C | {'ess': 4, 'noness': 0} | False | 38058007 | ATGCTTCTAGAAGCCTGGAGG( | [] |
| 5234 | 17 | TBC1D3E | unavailable | False | 38138552 | ATGCTTCTAGAAGCCTGGAGG( | [] |
| 5235 | 17 | TBC1D3 | {'ess': 4, 'noness': 0} | False | 38181973 | ATGCTTCTAGAAGCCTGGAGG( | [] |
| 5236 | 17 | SOCS7 | {'ess': 0, 'noness': 4} | True | 38395967 | GAGCCCCTCGCTACGTGGAGG | [] |
| 5237 | 17 | SRCIN1 | {'ess': 0, 'noness': 4} | True | 38533296 | GGCTTCTAGAAGGAGATGGAAG | [] |
| 5238 | 17 | PIP4K2B | {'ess': 0, 'noness': 4} | True | 38769690 | AGAGAACTACGTCAGGATGTTG | [] |
| 5239 | 17 | FBXO47 | {'ess': 0, 'noness': 7} | True | 38938753 | AAGCGCTATAAACCTTCATGTG | [] |
| 5240 | 17 | CACNB1 | {'ess': 2, 'noness': 7} | False | 39177244 | CCCACTACATGGCATGTTCCT( | [] |
| 5241 | 17 | STAC2 | {'ess': 1, 'noness': 6} | True | 39214406 | GTGTGCTACCCTGGGGGCTGT | [] |
| 5242 | 17 | MED1 | {'ess': 6, 'noness': 2} | False | 39407474 | AGGTTCCTAATTCCCAATCAGG | [] |
| 5243 | 17 | CDK12 | {'ess': 4, 'noness': 4} | False | 39564251 | CTCCTACCTGTAAAGGAAAATG | [] |
| 5244 | 17 | PPP1R1B | {'ess': 0, 'noness': 7} | True | 39635864 | GGTGCCTATGTGCCAGGCTCA( | [] |
| 5245 | 17 | ZPBP2 | {'ess': 0, 'noness': 7} | True | 39876808 | TTCACCTCTAATCTTCATATTG | [] |
| 5246 | 17 | MSL1 | {'ess': 3, 'noness': 5} | False | 40129637 | AGTGTTCTCTATGCCTTACTTG | [] |
| 5247 | 17 | MSL1 | {'ess': 3, 'noness': 5} | False | 40134368 | ACAGCTATTTCCTACACGTCCG | [] |
| 5248 | 17 | GJD3 | {'ess': 0, 'noness': 5} | True | 40362930 | GCCCCTAGATGGCCAGATCTC( | [] |
| 5249 | 17 | TNS4 | {'ess': 0, 'noness': 7} | True | 40477587 | TCTCCCCTACATCCTTTCTGC( | [] |
| 5250 | 17 | CCR7 | {'ess': 0, 'noness': 7} | True | 40554741 | AGTCGCCTATGGGGAGAAGGT( | [] |
| 5251 | 17 | KRT28 | {'ess': 0, 'noness': 7} | True | 40792426 | ACAGCTAGAAAGGAACCCTTTG | [] |
| 5252 | 17 | KRTAP4-4 | {'ess': 0, 'noness': 7} | True | 41160190 | TGGGCTAGCAGCATAGAGACT( | [] |
| 5253 | 17 | KRT33A | {'ess': 0, 'noness': 7} | True | 41348118 | GGGCCTCTAGTACCCAAATGT( | [] |
| 5254 | 17 | KRT33B | {'ess': 0, 'noness': 1} | False | 41363835 | GGGTATCTAGTACCCAAAGGTG | [] |
| 5255 | 17 | KRT31 | {'ess': 0, 'noness': 1} | False | 41394015 | CCTAGGTTCTAGCGCACGAAG( | [] |
| 5256 | 17 | KRT13 | {'ess': 1, 'noness': 6} | True | 41501343 | AGAGGTGCTACGGGGGCTGAC | [] |
| 5257 | 17 | KRT15 | {'ess': 0, 'noness': 7} | True | 41514120 | AAGAGGCTAGAGAGAGAAGGAG | [] |
| 5258 | 17 | KRT9 | {'ess': 0, 'noness': 7} | True | 41567272 | GAAGAAGACTAGGAATGGGATG | [] |
| 5259 | 17 | KRT16 | {'ess': 0, 'noness': 7} | True | 41609834 | AGTTCTAGGAGCTCTGGCCCT( | [] |
| 5260 | 17 | JUP | {'ess': 3, 'noness': 6} | False | 41755743 | GGCCGCCTAGGCCAGCATGTG | [] |
| 5261 | 17 | KLHL11 | {'ess': 1, 'noness': 7} | True | 41853739 | AGAACCTAGCATTCAATCTGAG | [] |
| 5262 | 17 | ZNF385C | {'ess': 0, 'noness': 1} | False | 42026896 | TGAGGGCTAATAAGGGGCAAG( | [] |
| 5263 | 17 | KAT2A | {'ess': 2, 'noness': 6} | False | 42113648 | ATGGGCCTACTTGTCAATGAGG | [] |
| 5264 | 17 | STAT5B | {'ess': 2, 'noness': 5} | False | 42214581 | TTGAACTAACAGTCAAATCCTG | ['GTCAAATCCTATATTCTAACAG'] |
| 5265 | 17 | CAVIN1 | unavailable | False | 42412043 | GAGACCCTAAACACTCCCACC( | [] |
| 5266 | 17 | TUBG2 | {'ess': 0, 'noness': 7} | True | 42666830 | TACCATCTAGAAGGAGAAGGAG | [] |
| 5267 | 17 | CCR10 | {'ess': 0, 'noness': 7} | True | 42679552 | GCAGCCCTAGTTGTCCCAGGA( | [] |
| 5268 | 17 | EZH1 | {'ess': 0, 'noness': 7} | True | 42702531 | TGGGAGGGCTAAAGGACGTCG( | [] |
| 5269 | 17 | RAMP2 | {'ess': 1, 'noness': 7} | True | 42762851 | CCCCTAGGCCTGGGCCTCACT | [] |
| 5270 | 17 | VPS25 | {'ess': 8, 'noness': 1} | False | 42779068 | TCCCTGCTAGAAGAACTTGACG | [] |
| 5271 | 17 | BECN1 | {'ess': 4, 'noness': 4} | False | 42812230 | AATACCTAATTAGCCGGGCGTG | [] |
| 5272 | 17 | AOC2 | {'ess': 0, 'noness': 7} | True | 42850347 | CAGGACTAGAAGCCGTGGTAA( | [] |
| 5273 | 17 | AOC3 | {'ess': 0, 'noness': 7} | True | 42856549 | GACCGCCTAGTTGTGAGAGAA( | [] |
| 5274 | 17 | RPL27 | {'ess': 7, 'noness': 1} | False | 43002919 | GCATCTAAAACCGCAGTTTCTG | [] |
| 5275 | 17 | IFI35 | {'ess': 2, 'noness': 6} | False | 43014298 | AGGCCCCTAGCCTGACTCAGA( | [] |
| 5276 | 17 | [illegible] | [illegible] | True | [illegible] | [illegible] | [] |
| 5277 | 17 | RND2 | {'ess': 0, 'noness': 6} | True | 43029113 | TGTTGCCTAGAGGTTAGATGTG | [] |
| 5278 | 17 | ETV4 | {'ess': 0, 'noness': 7} | True | 43528518 | CGCTGGGGGCTAGTAAGAGTA( | [] |
| 5279 | 17 | SOST | {'ess': 0, 'noness': 7} | True | 43755341 | GGCTCTAGTAGGCGTTCTCCA( | [] |
| 5280 | 17 | DUSP3 | {'ess': 0, 'noness': 7} | True | 43769608 | TGGGGGTGCCCTAGGGTTTCA( | ['GGTTTCAACTTCCCCTCCTTGG'] |
| 5281 | 17 | CFAP97D1 | unavailable | False | 43783892 | ACATACCTATTCATTTTGGGAG | [] |
| 5282 | 17 | CD300LG | {'ess': 0, 'noness': 7} | True | 43861910 | CCCTACGCTGAGACAAACTTC( | [] |
| 5283 | 17 | RUNDC3A | {'ess': 0, 'noness': 7} | True | 44317503 | GAGCCACTAATTTGGCTCTGAG | [] |
| 5284 | 17 | FAM171A2 | {'ess': 0, 'noness': 7} | True | 44353732 | GCGCGGGCGCTACTTGACGTT | [] |
| 5285 | 17 | MEIOC | unavailable | False | 44676854 | TCTCAAACTAATGCTTAAACAG | [] |
| 5286 | 17 | DBF4B | {'ess': 2, 'noness': 6} | False | 44751252 | CACCTCTAACCTGAGTCTACAG | [] |
| 5287 | 17 | HIGD1B | {'ess': 0, 'noness': 7} | True | 44850395 | TAGGAGTCCTACTTCTCTCCAG | [] |
| 5288 | 17 | FAM187A | {'ess': 0, 'noness': 1} | False | 44905070 | GAGCTCCTAGGGGGAGAAGCT( | [] |
| 5289 | 17 | GFAP | {'ess': 0, 'noness': 7} | True | 44910135 | GGCAGCTAACCGCGAGCCGGC | [] |
| 5290 | 17 | GFAP | {'ess': 0, 'noness': 7} | True | 44910458 | TCTGGCTAGGAGCGCTGCAGT( | [] |
| 5291 | 17 | KIF18B | {'ess': 2, 'noness': 4} | False | 44926399 | TGTCCCTAGTGCCAGTCACCT( | [] |
| 5292 | 17 | KIF18B | {'ess': 2, 'noness': 4} | False | 44932507 | CAGCACTCTAGGTGACAGTCA( | [] |
| 5293 | 17 | DCAKD | {'ess': 0, 'noness': 7} | True | 45024432 | TGCCCCACTAGGCGTAGGCA( | [] |
| 5294 | 17 | ACBD4 | {'ess': 3, 'noness': 5} | False | 45138364 | GGAGTTCTAGAAGGGAAGAGG( | [] |
| 5295 | 17 | ACBD4 | {'ess': 3, 'noness': 5} | False | 45143640 | GTTCTTCTAAACCCTAGGAAGG | [] |
| 5296 | 17 | HEXIM1 | {'ess': 2, 'noness': 6} | False | 45150269 | TTCAGTCTAGTCTCCAAACTTG | [] |
| 5297 | 17 | HEXIM2 | {'ess': 0, 'noness': 7} | True | 45169808 | GGAGGCACCCCTAGGTACCCG | [] |
| 5298 | 17 | FMNL1 | {'ess': 0, 'noness': 7} | True | 45246595 | TACCTGAGGGGCTAGAGGGGC | ['GAGGGGCATCTCTTCTCCCAGG'] |
| 5299 | 17 | FMNL1 | {'ess': 0, 'noness': 7} | True | 45246969 | TAGCGGCTACAGCGAGAGGTC( | [] |
| 5300 | 17 | ARHGAP27 | {'ess': 0, 'noness': 7} | True | 45397642 | ATTCCTACCTCTATTAATACAG | [] |
| 5301 | 17 | ARHGAP27 | {'ess': 0, 'noness': 7} | True | 45401647 | GCTTCTAGATCCCATGTTCATG | [] |
| 5302 | 17 | STH | {'ess': 0, 'noness': 4} | True | 45999665 | CCCTTTCTACACTTGGCATGAG | [] |
| 5303 | 17 | LRRC37A | {'ess': 0, 'noness': 7} | True | 46337722 | GTGTTCTCCTCCTATGGCAGG( | [] |
| 5304 | 17 | ARL17B | {'ess': 0, 'noness': 7} | True | 46339499 | TCAGGCTACTTTACCACAGCTG | [] |
| 5305 | 17 | LOC10798 | unavailable | False | 46341691 | AACCATCCTAAACGGCGGCTG( | [] |
| 5306 | 17 | LRRC37A2 | {'ess': 0, 'noness': 5} | True | 46555578 | GTGTTCTCCTCCTATGGCAGG( | [] |
| 5307 | 17 | LRRC37A2 | {'ess': 0, 'noness': 5} | True | 47049060 | CCAATCCTAAACCTATGAAATG | [] |
| 5308 | 17 | ARL17A | {'ess': 1, 'noness': 6} | True | 46526768 | GATTCTCTAGATTTTGCCCCTG | [] |
| 5309 | 17 | ARL17A | {'ess': 1, 'noness': 6} | True | 46557355 | TCAGGCTACTTTACCACAGCTG | [] |
| 5310 | 17 | WNT3 | {'ess': 0, 'noness': 7} | True | 46768319 | TGGTGCCCTACTTGCAGGTGT( | [] |
| 5311 | 17 | WNT9B | {'ess': 0, 'noness': 7} | True | 46876717 | GTAGGCCTAGTGCTTGCAGGT( | [] |
| 5312 | 17 | EFCAB13 | unavailable | False | 47429965 | GAAACTACTAACCATGTTCAGG | [] |
| 5313 | 17 | EFCAB13 | unavailable | False | 47440713 | ACTACCTAAAATTTTGAGTTTG | [] |
| 5314 | 17 | TBKBP1 | {'ess': 0, 'noness': 7} | True | 47710625 | GCTGGTGCCCTAGATCTTGCT( | [] |
| 5315 | 17 | OSBPL7 | {'ess': 1, 'noness': 6} | True | 47808290 | GGCCAGGGCTACCAGAGCACG | [] |
| 5316 | 17 | MRPL10 | {'ess': 7, 'noness': 1} | False | 47824204 | GCTGGCTACGAGTCCGGAACA( | [] |
| 5317 | 17 | SCRN2 | {'ess': 1, 'noness': 6} | True | 47838251 | GGTAGCCCCTACTCCCTGGGG | [] |
| 5318 | 17 | CDK5RAP3 | {'ess': 0, 'noness': 6} | True | 47981733 | TACTAACTAGAAGGAGAATCTG | [] |
| 5319 | 17 | HOXB4 | {'ess': 1, 'noness': 6} | True | 48576721 | GGGGCACTAGAGCGCGCGGG( | [] |
| 5320 | 17 | HOXB8 | {'ess': 0, 'noness': 7} | True | 48613201 | AGCTGAAGCCTACTTCTTGTCG | [] |
| 5321 | 17 | TTLL6 | {'ess': 1, 'noness': 7} | True | 48769888 | GGCCTACCTAGCTCCTCTCAC( | ['CTCACATCCTTTTTGGCCAGAG'] |
| 5322 | 17 | UBE2Z | {'ess': 2, 'noness': 7} | False | 48927133 | GAGCAGGGTCTAAACCCTCAG( | [] |
| 5323 | 17 | SLC35B1 | {'ess': 6, 'noness': 2} | False | 49701457 | TCCTAGTGGGATGTCTTCTTAG | [] |
| 5324 | 17 | PDK2 | {'ess': 2, 'noness': 6} | False | 50100151 | GTGGTCTAGGTCCCCTCAGCA( | [] |
| 5325 | 17 | SAMD14 | {'ess': 0, 'noness': 7} | True | 50112892 | ACCCTCCCCTAGCTCTTCTTG( | [] |
| 5326 | 17 | XYLT2 | {'ess': 2, 'noness': 6} | False | 50360290 | GCCCTGCTACCTGAGTCGCCC | [] |
| 5327 | 17 | ANKRD40 | unavailable | False | 50768922 | CACCTCTAGTCCGTCAGATGT( | [] |
| 5328 | 17 | UTP18 | {'ess': 3, 'noness': 5} | False | 51286758 | ATCTGCTACTCATGAATTGCTG | [] |
| 5329 | 17 | CA10 | {'ess': 0, 'noness': 7} | True | 51631583 | GCTTTGTTCCCTACTTGAGGAG | [] |
| 5330 | 17 | COX11 | {'ess': 6, 'noness': 2} | False | 54961281 | ACTCTCTAAAACGTAGACTCTG | [] |
| 5331 | 17 | STXBP4 | {'ess': 0, 'noness': 7} | True | 55043628 | TCTTCATCTAATTCTGCCCTGG | [] |
| 5332 | 17 | HLF | {'ess': 0, 'noness': 7} | True | 55320878 | TGCCATCCTACAGGGGCCCGT | [] |
| 5333 | 17 | PCTP | {'ess': 0, 'noness': 7} | True | 55775342 | CTAGTGATACAAACTACCACTG | [] |
| 5334 | 17 | PCTP | {'ess': 0, 'noness': 7} | True | 55822840 | TCAATCTATTGCAGAATGTTTG | [] |
| 5335 | 17 | ANKFN1 | {'ess': 1, 'noness': 7} | True | 56482546 | TAATATTTCTAGGTGAAACAAG | [] |
| 5336 | 17 | ANKFN1 | {'ess': 1, 'noness': 7} | True | 56511268 | GCCTCCCTAAAGCATGCTGCT( | [] |
| 5337 | 17 | NOG | {'ess': 1, 'noness': 6} | True | 56594921 | CGAGTTCTAGCACGAGCACTT( | [] |
| 5338 | 17 | DGKE | {'ess': 1, 'noness': 6} | True | 56862790 | CCCTCATCCATCTATTCAGTCG | [] |
| 5339 | 17 | TRIM25 | {'ess': 0, 'noness': 7} | True | 56891699 | GCCTGCCTACTTGGGGGAGCA | [] |
| 5340 | 17 | SCPEP1 | {'ess': 0, 'noness': 7} | True | 57006234 | CATCCATCCTATTCTTGCTGAG | [] |
| 5341 | 17 | MSI2 | {'ess': 1, 'noness': 7} | True | 57631830 | GAGCTAGTTGATAAAAATTATG | [] |
| 5342 | 17 | MSI2 | {'ess': 1, 'noness': 7} | True | 57655791 | GCTGGTACTAACAGGAAGCTG( | [] |
| 5343 | 17 | DYNLL2 | {'ess': 1, 'noness': 4} | False | 58089278 | GGCCACCTAGCCTGACTTGAA( | [] |
| 5344 | 17 | MKS1 | {'ess': 0, 'noness': 7} | True | 58205889 | GTGGTCTCTAATCAGAAAGACG | [] |
| 5345 | 17 | MKS1 | {'ess': 0, 'noness': 7} | True | 58206078 | GCTGTGAGCTAGGAGACCAGG( | [] |
| 5346 | 17 | MKS1 | {'ess': 0, 'noness': 7} | True | 58206282 | CATACCTAGCACATTGTGAATG | [] |
| 5347 | 17 | LPO | {'ess': 0, 'noness': 7} | True | 58267993 | GGGCCCCTAATTCTTCACTGA( | [] |
| 5348 | 17 | MPO | {'ess': 1, 'noness': 7} | True | 58270655 | CCTCTAGGAGGCTTCCCTCCA( | [] |
| 5349 | 17 | TSPOAP1 | unavailable | False | 58304369 | TCTATCTCCATCTAGCACTGGG | ['CACTGGACTCTTCTCCTCCTCG'] |
| 5350 | 17 | SUPT4H1 | {'ess': 4, 'noness': 4} | False | 58349245 | TCTTGCTAGGTCTTTATAGCTG | [] |
| 5351 | 17 | SEPTIN4 | unavailable | False | 58521881 | CCACCTGCTAGTGGCAGCCCT | [] |
| 5352 | 17 | GDPD1 | {'ess': 0, 'noness': 7} | True | 59273772 | TTTTCTATGCTGAAAAGTTATG | [] |
| 5353 | 17 | GDPD1 | {'ess': 0, 'noness': 7} | True | 59275197 | AGGTACACCTAGAAATAACAAG | [] |
| 5354 | 17 | PTRH2 | {'ess': 1, 'noness': 7} | True | 59897438 | GTCCACCTAGTAAAGTTTTAGG | [] |
| 5355 | 17 | TUBD1 | {'ess': 5, 'noness': 3} | False | 59860371 | GTCTAAAAAGTCCTCTTCTTCG | [] |
| 5356 | 17 | BCAS3 | {'ess': 0, 'noness': 7} | True | 61235536 | TCTGTACTACGTAGTCACCTCG | ['ACCTCACAATCTGTTTGGAATG'] |
| 5357 | 17 | BCAS3 | {'ess': 0, 'noness': 7} | True | 61392124 | GGTACCTACGGGAAGCCAGTC( | [] |
| 5358 | 17 | TBX4 | {'ess': 0, 'noness': 7} | True | 61481367 | TTGCAACTACGAAACTCTGCCG | [] |
| 5359 | 17 | MRC2 | {'ess': 1, 'noness': 7} | True | 62692450 | GCGCCTGGCTCTATTCTTGTT( | [] |
| 5360 | 17 | MARCHF10 | unavailable | False | 62701702 | CTTGCTAGACGACCTGGCTTT( | [] |
| 5361 | 17 | KCNH6 | {'ess': 0, 'noness': 7} | True | 63542708 | ATTTATTGTCTCTAATCCTCAG | [] |
| 5362 | 17 | DCAF7 | {'ess': 3, 'noness': 1} | False | 63589171 | GCCACCAACACTACACTCTGA( | [] |
| 5363 | 17 | DDX42 | {'ess': 3, 'noness': 4} | False | 63818397 | ATCCCCTCTAACTGTCCCATC( | [] |
| 5364 | 17 | CSH2 | {'ess': 0, 'noness': 7} | True | 63872125 | CGCGGGCACCTAGAAGCCACA | [] |
| 5365 | 17 | CSH2 | {'ess': 0, 'noness': 7} | True | 63872528 | CCAGCCCTCTAAGCCAGTGGG | [] |
| 5366 | 17 | CSH1 | {'ess': 0, 'noness': 7} | True | 63895021 | GGGCACCTAGAAGCCACAGCT( | [] |
| 5367 | 17 | CSHL1 | {'ess': 1, 'noness': 6} | True | 63909710 | GGGCCCCTAGAAGCCACAGCT | [] |
| 5368 | 17 | GH1 | {'ess': 0, 'noness': 6} | True | 63917308 | GGGCAGCTAGAAGCCACAGCT( | [] |
| 5369 | 17 | SCN4A | {'ess': 1, 'noness': 5} | False | 63940770 | CTGCTAGACAAGAGACTCCTTG | [] |
| 5370 | 17 | ICAM2 | {'ess': 0, 'noness': 7} | True | 64002746 | GTTGCTATGGCCGGAAGGCCT( | [] |
| 5371 | 17 | PECAM1 | unavailable | False | 64323815 | TGCTGTCTAAGTTCCATCAAGG | [] |
| 5372 | 17 | POLG2 | {'ess': 6, 'noness': 3} | False | 64477822 | AAAATCTATACATTCTTAGCTG | [] |
| 5373 | 17 | LRRC37A3 | {'ess': 0, 'noness': 7} | True | 64854598 | GGGTTCTCCTCCTATGGCAGG( | [] |
| 5374 | 17 | KPNA2 | {'ess': 4, 'noness': 4} | False | 68040595 | TACATGATCTAAAAGTTAAAGG | [] |
| 5375 | 17 | FAM20A | {'ess': 0, 'noness': 7} | True | 68541556 | GTGCTAGGTCCTGGGGATACA( | [] |
| 5376 | 17 | KCNJ16 | {'ess': 0, 'noness': 5} | True | 70133343 | GACTACATTTGGGATTCTACAG | [] |
| 5377 | 17 | SLC39A11 | {'ess': 0, 'noness': 6} | True | 72647583 | TCAGCCCTAGCCCAGGCCAAC | [] |
| 5378 | 17 | SLC39A11 | {'ess': 0, 'noness': 6} | True | 72725637 | AACCTATGCTACCAAGCTACCG | [] |
| 5379 | 17 | C17orf80 | {'ess': 0, 'noness': 7} | True | 73245189 | GTCAACACTACTTACGCCACC( | [] |
| 5380 | 17 | C17orf80 | {'ess': 0, 'noness': 7} | True | 73247340 | CGTGCTTCCTAATCCAAACATG | [] |
| 5381 | 17 | DNAI2 | {'ess': 0, 'noness': 7} | True | 74314215 | -1 | -1 |
| 5382 | 17 | BTBD17 | {'ess': 0, 'noness': 7} | True | 74356656 | CCGAGGCTACTTGGGGGTCCG | [] |
| 5383 | 17 | GPR142 | unavailable | False | 74372599 | CTCCTCTACACTTCTGCCCCT( | [] |
| 5384 | 17 | GPRC5C | {'ess': 1, 'noness': 7} | True | 74449366 | TCTACCATCTCGTTTTACTTTG | [] |
| 5385 | 17 | CD300A | {'ess': 0, 'noness': 7} | True | 74484125 | CTATGTCTTCCTTATCACACTG | [] |
| 5386 | 17 | CD300LB | {'ess': 0, 'noness': 7} | True | 74522737 | TCCATCTCTCTAAGTGGCCATG | [] |
| 5387 | 17 | CD300C | {'ess': 1, 'noness': 7} | True | 74534628 | AAGGAAGGAGACTAAAATAATG | [] |
| 5388 | 17 | CD300C | {'ess': 1, 'noness': 7} | True | 74541588 | ATGCTACTGGTTCTCACCCTTG | [] |

| | | | | | | | |
|---|---|---|---|---|---|---|---|
| 5389 | 17 | CD300C | {'exs': 1, 'nonexs': 7} | True | 74543172 | CCCATCCCTAGGTACTGCAGA | [] |
| 5390 | 17 | CD300E | {'exs': 0, 'nonexs': 5} | True | 74612652 | TGGCTCTATCTTCCAGGAGGA | [] |
| 5391 | 17 | CD300LF | {'exs': 0, 'nonexs': 7} | True | 74695095 | TGCAGGCTAAGGCCTGCTGAT | [] |
| 5392 | 17 | NT5C | {'exs': 0, 'nonexs': 7} | True | 75130827 | GTTCTACGAACTGGGGCCCCA | [] |
| 5393 | 17 | JPT1 | unavailable | False | 75136101 | TCAGAGCTAACCCAAGACGAGG | [] |
| 5394 | 17 | MRPS7 | {'exs': 5, 'nonexs': 3} | False | 75265922 | GAGACTCTACCACCAGCGGTA | [] |
| 5395 | 17 | MIF4GD | {'exs': 0, 'nonexs': 7} | True | 75266739 | GGAGGCCTAGTCGGAGACTTC | [] |
| 5396 | 17 | MIF4GD | {'exs': 0, 'nonexs': 7} | True | 75267267 | ACCCACAGCTAGCCATGGCTT | [] |
| 5397 | 17 | LLGL2 | {'exs': 0, 'nonexs': 7} | True | 75564541 | GCCTCCCACACCTAACCCTGG | [] |
| 5398 | 17 | UNC13D | {'exs': 0, 'nonexs': 7} | True | 75827964 | TCTACGGCTACGGTGCCGGCC | [] |
| 5399 | 17 | WBP2 | {'exs': 0, 'nonexs': 7} | True | 75846733 | GAGGGCCTACTGGGTCTTCTT | ['CTTCTTATCTTCCGGTGGGTAG'] |
| 5400 | 17 | TRIM65 | {'exs': 1, 'nonexs': 7} | True | 75882676 | CTATCCTAAAGGATGGCAACTG | [] |
| 5401 | 17 | TEN1 | {'exs': 1, 'nonexs': 4} | False | 76000261 | TTTCCTACTGGCTGCCGCCCC | [] |
| 5402 | 17 | RNF157 | {'exs': 0, 'nonexs': 7} | True | 76147268 | TACCTATTCCTCAATGTTTATG | [] |
| 5403 | 17 | PRPSAP1 | {'exs': 0, 'nonexs': 7} | True | 76311541 | TGAAAGCTAGTCATCCACAGTG | [] |
| 5404 | 17 | UBE2O | {'exs': 1, 'nonexs': 7} | True | 76390942 | TGGCAGCTACTTGTCCTCTGT | [] |
| 5405 | 17 | CYGB | {'exs': 0, 'nonexs': 7} | True | 76528577 | GGGGTCCTACGGCCCCGAAGA | [] |
| 5406 | 17 | MXRA7 | {'exs': 0, 'nonexs': 7} | True | 76677587 | GCGCAGTCTACATGTCGTAGA | [] |
| 5407 | 17 | MXRA7 | {'exs': 0, 'nonexs': 7} | True | 76683893 | AACTTGCTACAGGGAAGTGAGG | [] |
| 5408 | 17 | MFSD11 | {'exs': 0, 'nonexs': 7} | True | 76785567 | GAGTCTAGGTGTTCAGGACCA | [] |
| 5409 | 17 | MGAT5B | {'exs': 1, 'nonexs': 7} | True | 76944173 | TAGGTCTAATTACAGGTGTGCG | [] |
| 5410 | 17 | SEC14L1 | {'exs': 0, 'nonexs': 7} | True | 77214022 | GCGGCACTACCTGGAGATCAT | [] |
| 5411 | 17 | SEPTIN9 | unavailable | False | 77498657 | GGTGGCGTCTACATCTCCGGG | [] |
| 5412 | 17 | TNRC6C | unavailable | False | 78104944 | CAGAGCCTACAGGGACTCCCC | [] |
| 5413 | 17 | TMC6 | {'exs': 0, 'nonexs': 7} | True | 78112147 | CCCTAGGCATCCTGTTCATCT | [] |
| 5414 | 17 | TMC8 | {'exs': 1, 'nonexs': 7} | True | 78139314 | CAGACGTGTTCTAGGCCCTCG | [] |
| 5415 | 17 | C17orf99 | {'exs': 0, 'nonexs': 7} | True | 78164665 | CATTTCCTACCTGGCCCACGT | [] |
| 5416 | 17 | C17orf99 | {'exs': 0, 'nonexs': 7} | True | 78166045 | GGTTCATTCTACATGGCTGCTG | [] |
| 5417 | 17 | AFMID | {'exs': 0, 'nonexs': 7} | True | 78206936 | CGTCAGAACTACTGGAAGATTG | [] |
| 5418 | 17 | BIRC5 | {'exs': 8, 'nonexs': 0} | False | 78223656 | CCTTTCCTAAGACATTGCTAAG | [] |
| 5419 | 17 | PGS1 | {'exs': 6, 'nonexs': 2} | False | 78419687 | GTGATGCTAGAGACAGCACCTC | [] |
| 5420 | 17 | PGS1 | {'exs': 6, 'nonexs': 2} | False | 78420228 | TCCAGAGTCTAGAAGGGCGCA | [] |
| 5421 | 17 | DNAH17 | {'exs': 0, 'nonexs': 1} | False | 78423905 | GAGCGAGCTAAACCTGTAGGA | [] |
| 5422 | 17 | CYTH1 | {'exs': 1, 'nonexs': 6} | True | 78677022 | CAGCTGCTATGGCTCTCCTGG | [] |
| 5423 | 17 | LGALS3BP | {'exs': 0, 'nonexs': 7} | True | 78971575 | CGCCGTCTAGTCCACACCTGA | [] |
| 5424 | 17 | C1QTNF1 | {'exs': 0, 'nonexs': 7} | True | 79049087 | GGCCAGCTAGGGCTCGGTGGC | [] |
| 5425 | 17 | RBFOX3 | {'exs': 0, 'nonexs': 5} | True | 79092596 | CTCTTGCTAGTAGGGGGTGAA | [] |
| 5426 | 17 | CBX2 | {'exs': 0, 'nonexs': 7} | True | 79782148 | ATGAGCACCTACTGTACGCATG | [] |
| 5427 | 17 | CBX4 | {'exs': 1, 'nonexs': 7} | True | 79833958 | GCCCTCCGGCTACACCGTCAC | [] |
| 5428 | 17 | TBC1D16 | {'exs': 1, 'nonexs': 7} | True | 79940858 | CCCGACCTATCTGCGGAAGCC | [] |
| 5429 | 17 | CCDC40 | {'exs': 0, 'nonexs': 7} | True | 80099774 | GCTCCCTAGGAGGGCCCTGGT | [] |
| 5430 | 17 | GAA | {'exs': 0, 'nonexs': 7} | True | 80119330 | GCCCGGCTAACACCAGCTGAC | [] |
| 5431 | 17 | CARD14 | {'exs': 0, 'nonexs': 5} | True | 80198755 | TCATCTATAAATGAGCAGAACG | [] |
| 5432 | 17 | CARD14 | {'exs': 0, 'nonexs': 5} | True | 80207088 | TGCACCTACTTGAGCTTCTTTG | [] |
| 5433 | 17 | RNF213 | {'exs': 0, 'nonexs': 7} | True | 80319479 | GAACTAGATTTCAGCGCCATTG | [] |
| 5434 | 17 | RNF213 | {'exs': 0, 'nonexs': 7} | True | 80370210 | AAGCTACCATCTGAAAAATCAG | [] |
| 5435 | 17 | RNF213 | {'exs': 0, 'nonexs': 7} | True | 80393497 | AATAATTCTATCTCATTTCTCG | [] |
| 5436 | 17 | ENDOV | unavailable | False | 80429922 | TCCACCCTACTGCCAGTCCTT | [] |
| 5437 | 17 | RPTOR | {'exs': 9, 'nonexs': 1} | False | 80964329 | GGTCACGCCGCTATCTGACAC | [] |
| 5438 | 17 | LOC10798 | unavailable | False | 80727594 | CTCTCTTCTAAAGCAGTTGTAG | [] |
| 5439 | 17 | PVALEF | unavailable | False | 81183010 | CATGGTGCTACTTCTTCTTTGG | [] |
| 5440 | 17 | NDUFAF8 | unavailable | False | 81241015 | CCTCCTAACAGCCTCCCTCCA | [] |
| 5441 | 17 | NDUFAF8 | unavailable | False | 81241152 | CAGGATCTAACACCGAGGAATG | [] |
| 5442 | 17 | SLC38A10 | {'exs': 2, 'nonexs': 5} | False | 81245555 | AGGTGCTAGGACTCCTCTGGA | [] |
| 5443 | 17 | LOC11226 | unavailable | False | 81346461 | CCGGCTCTAGCTGGGCGACCT | [] |
| 5444 | 17 | NPLOC4 | {'exs': 6, 'nonexs': 2} | False | 81559258 | GGCGCCCTAGGTCCTGGGGAG | [] |
| 5445 | 17 | PDE6G | {'exs': 0, 'nonexs': 5} | True | 81651073 | CGTGCTAGATGATGCCATATTG | [] |
| 5446 | 17 | OXLD1 | unavailable | False | 81666298 | AGGCTAAGCGGCCCCTCCACA | [] |
| 5447 | 17 | CCDC137 | {'exs': 1, 'nonexs': 7} | True | 81670548 | TGGCTATGAAACAGGGTACCTG | [] |
| 5448 | 17 | MRPL12 | {'exs': 8, 'nonexs': 0} | False | 81707239 | TGGAGGCTACTCCAGAACCAC | [] |
| 5449 | 17 | SLC25A10 | {'exs': 1, 'nonexs': 6} | True | 81720516 | GCAGAGCTAGCACTTGGGTGA | [] |
| 5450 | 17 | MCRIP1 | unavailable | False | 81823246 | CGGGCGCTCTAGGACTTCTTG | [] |
| 5451 | 17 | MAFG | {'exs': 1, 'nonexs': 6} | True | 81922604 | CGTCCCTACGATCGGGCATCC | [] |
| 5452 | 17 | MYADML2 | {'exs': 0, 'nonexs': 5} | True | 81940817 | TGTGGGCTACAGGCTGGGCAC | [] |
| 5453 | 17 | NOTUM | {'exs': 1, 'nonexs': 6} | True | 81952960 | CTGCCTAGCTTCCGTTGCTCA | [] |
| 5454 | 17 | CENPX | unavailable | False | 82019204 | GAGATCCCTAGAAGTCCAGGA | [] |
| 5455 | 17 | RAC3 | {'exs': 0, 'nonexs': 7} | True | 82033828 | GCCAGGGCTCTAGAAGACGGT | [] |
| 5456 | 17 | FASN | {'exs': 5, 'nonexs': 3} | False | 82079142 | ACGGGCCTAGCCCTCCCCGCAC | [] |
| 5457 | 17 | CCDC57 | {'exs': 0, 'nonexs': 3} | True | 82157732 | AGGAGGAGCTAGCGGGCACCC | ['GCACCCACCTCTCTCCCAAGGG'] |
| 5458 | 17 | CCDC57 | {'exs': 0, 'nonexs': 3} | True | 82175663 | CTCCCTACAATGTGTAAAACCG | [] |
| 5459 | 17 | CSNK1D | {'exs': 0, 'nonexs': 7} | True | 82249006 | CGAGCAGCTACTTGCCGTGGT | [] |
| 5460 | 17 | SECTM1 | {'exs': 1, 'nonexs': 7} | True | 82322160 | GCAGGCGGCTATGGGTCTGCG | [] |
| 5461 | 17 | TEX19 | {'exs': 2, 'nonexs': 6} | False | 82362644 | CCCCTGCTATGAAGGAAAGAAG | [] |
| 5462 | 17 | OGFOD3 | unavailable | False | 82392397 | TGCTGGCTACGGGAACGCTGG | [] |
| 5463 | 17 | FOXK2 | {'exs': 1, 'nonexs': 7} | True | 82601498 | TCCCGGTCGCTAGTTCTGGAC | ['CTGGACACCCTTTTCCCTTACG'] |
| 5464 | 17 | LOC10537 | unavailable | False | 82702480 | TCTATTTCAGCATTTTCCCACG | [] |
| 5465 | 17 | FN3K | {'exs': 1, 'nonexs': 6} | True | 82750754 | GCCGCTACTTGAGCAGCCTTC | [] |
| 5466 | 18 | TUBB8B | unavailable | False | 47389 | GTTCTAGGCCACCTCCTCCTC | [] |
| 5467 | 18 | THOC1 | {'exs': 6, 'nonexs': 3} | False | 214625 | AAGCTAACTATTTGTCTCATTG | [] |
| 5468 | 18 | TYMS | {'exs': 7, 'nonexs': 1} | False | 672996 | GCACCCTAAACAGCCATTTCCC | ['TTTCCATTTTAATAGTTGGATG'] |
| 5469 | 18 | ADCYAP1 | {'exs': 0, 'nonexs': 7} | True | 909634 | ATCGCTACAAATAAGCTATTCG | [] |
| 5470 | 18 | METTL4 | {'exs': 2, 'nonexs': 6} | False | 2539066 | CCCAACTAGTCCAACCTGGCT | [] |
| 5471 | 18 | DLGAP1 | {'exs': 0, 'nonexs': 7} | True | 3502426 | GAAGCCTATTTAGACTGTCCAG | [] |
| 5472 | 18 | DLGAP1 | {'exs': 0, 'nonexs': 7} | True | 3709460 | GTCTTCCTACCCTAGCGCCCA | [] |
| 5473 | 18 | ZBTB14 | unavailable | False | 5290857 | CGCCTCTAGCTACAGGCTATC | [] |
| 5474 | 18 | TMEM200 | {'exs': 0, 'nonexs': 5} | True | 5890197 | CCCCTTTCTCTAAATGCCTGTG | [] |
| 5475 | 18 | L3MBTL4 | {'exs': 0, 'nonexs': 7} | True | 6032114 | TAGGTGGGACTATAGGCACCC | [] |
| 5476 | 18 | L3MBTL4 | {'exs': 0, 'nonexs': 7} | True | 6078862 | TGCAACTAGACAGTCCCATCTG | [] |
| 5477 | 18 | L3MBTL4 | {'exs': 0, 'nonexs': 7} | True | 6080737 | AGGGCGGGAACTAGAACACAA | [] |
| 5478 | 18 | L3MBTL4 | {'exs': 0, 'nonexs': 7} | True | 6161502 | TGCACTCTACCTTGGCTCAATG | [] |
| 5479 | 18 | ARHGAP2 | {'exs': 0, 'nonexs': 7} | True | 6898573 | CTGCCTATGTGGAGATGTCCA | [] |
| 5480 | 18 | PTPRM | {'exs': 1, 'nonexs': 6} | True | 8212931 | GGCTAGGAAGAATGTTCATAAG | [] |
| 5481 | 18 | MTCL1 | unavailable | False | 8828945 | GCAGGGCTAGAGTCCCCAGGG | [] |
| 5482 | 18 | MTCL1 | unavailable | False | 8831627 | ATGGAGCTAGTGCAGAACAAGG | [] |
| 5483 | 18 | ANKRD12 | {'exs': 0, 'nonexs': 7} | True | 9281125 | AGTACTGACTGCTATATAGGAG | [] |
| 5484 | 18 | PPP4R1 | {'exs': 2, 'nonexs': 6} | False | 9547788 | GCCTTCTAGTAGGTTGAGGAC | [] |
| 5485 | 18 | VAPA | {'exs': 1, 'nonexs': 6} | True | 9954210 | TCTACAAGATGAATTTCCCTAG | [] |
| 5486 | 18 | APCDD1 | {'exs': 0, 'nonexs': 7} | True | 10488037 | CTAAAACTTCTATCTGCGGATG | [] |
| 5487 | 18 | NAPG | {'exs': 3, 'nonexs': 5} | False | 10550219 | AAATACTAGCATAGTCCTCCTG | [] |
| 5488 | 18 | MPPE1 | {'exs': 0, 'nonexs': 7} | True | 11885028 | CAACTAAGCATCTGTTCCCTAG | [] |
| 5489 | 18 | ANKRD62 | {'exs': 0, 'nonexs': 3} | True | 12127938 | ACCAGCTACAGAGTAACTGTTG | [] |
| 5490 | 18 | CIDEA | {'exs': 0, 'nonexs': 7} | True | 12277269 | TCCCTATCCACACGTGAACCT | [] |
| 5491 | 18 | TUBB6 | {'exs': 0, 'nonexs': 7} | True | 12326129 | CTCTATTCCGACTATCCATCGG | ['TCCTCTTCCTCATCCTCAAAAG', 'TCCATCGATCTCCTCTTCCTCG'] |
| 5492 | 18 | AFG3L2 | {'exs': 7, 'nonexs': 0} | False | 12329564 | CCTCTGGGCCTCTAGTTGGCA | ['TTGGCAACTTTCTCACCCGGGG'] |
| 5493 | 18 | SPIRE1 | {'exs': 0, 'nonexs': 7} | True | 12475053 | GAGTCACTAGATGCCAGACATG | [] |
| 5494 | 18 | CEP76 | {'exs': 0, 'nonexs': 7} | True | 12662152 | CCTAAGAGGCACATTCACCTAG | [] |
| 5495 | 18 | CEP76 | {'exs': 0, 'nonexs': 7} | True | 12669833 | GTCTCTACTTAAAAAAAATTAG | [] |
| 5496 | 18 | CEP76 | {'exs': 0, 'nonexs': 7} | True | 12673364 | ATTGGCCCTATAATACCGAGCG | [] |
| 5497 | 18 | MC2R | {'exs': 0, 'nonexs': 7} | True | 13884624 | GCCATTCTACCAGTACCTGCT | [] |
| 5498 | 18 | ZNF519 | {'exs': 0, 'nonexs': 7} | True | 14104916 | TGCAGGGTTTCTACCTGGTATG | [] |
| 5499 | 18 | POTEC | {'exs': 0, 'nonexs': 7} | True | 14511897 | TTCATCTAGTTCCAGTCTCCAG | [] |
| 5500 | 18 | ANKRD30B | {'exs': 0, 'nonexs': 7} | True | 14853909 | AGCCTCTATTGATGCCTCAGAG | [] |
| 5501 | 18 | RBBP8 | {'exs': 5, 'nonexs': 3} | False | 23026239 | CGTCTATGTCTTCTGCTCCTTG | [] |
| 5502 | 18 | CABLES1 | {'exs': 0, 'nonexs': 7} | True | 23257366 | AGTGCTAGGAACTCTGGACCA | [] |
| 5503 | 18 | RIOK3 | {'exs': 2, 'nonexs': 6} | False | 23481278 | ATTAGTGCTATTCATCATATAG | [] |
| 5504 | 18 | NPC1 | {'exs': 1, 'nonexs': 7} | True | 23532201 | GAGAGGGCTAGAAATTTAGAAG | [] |
| 5505 | 18 | OSBPL1A | {'exs': 0, 'nonexs': 1} | False | 24265878 | CACCACCTACTGCTTCCCATC | [] |
| 5506 | 18 | ZNF521 | {'exs': 0, 'nonexs': 6} | True | 25062711 | CTTGCACTAACTGCTGTGTTGG | [] |
| 5507 | 18 | CHST9 | {'exs': 0, 'nonexs': 7} | True | 26916258 | GCAAACTACAAAAATGGAGTTG | [] |
| 5508 | 18 | DSG1 | {'exs': 0, 'nonexs': 7} | True | 31355345 | GTCCTGACTACTTGCTATATTG | [] |
| 5509 | 18 | GAREM1 | unavailable | False | 32267870 | TGGCTATATTTTGGGCCTCCAG | [] |
| 5510 | 18 | DTNA | {'exs': 0, 'nonexs': 7} | True | 34829070 | ATCTAGACGCATCCACCAAAAG | [] |
| 5511 | 18 | SLC39A6 | {'exs': 1, 'nonexs': 6} | True | 36109592 | ACCTTAACTAGAAATTTATACG | [] |
| 5512 | 18 | TPGS2 | unavailable | False | 36783041 | AGCTCTAGAACCTTCTGCTCCC | [] |
| 5513 | 18 | SYT4 | {'exs': 1, 'nonexs': 7} | True | 43270340 | TAGGATGCTAACCATCACAGAG | [] |
| 5514 | 18 | SETBP1 | {'exs': 0, 'nonexs': 7} | True | 44959737 | CTGCTATGAAGAGTATTTCCAG | [] |
| 5515 | 18 | SETBP1 | {'exs': 0, 'nonexs': 7} | True | 45063697 | CCCGCCCCTAGGGAAGGACCTC | [] |
| 5516 | 18 | EPG5 | unavailable | False | 45852466 | CAGTGTTAACTATCGTATGTGG | [] |
| 5517 | 18 | EPG5 | unavailable | False | 45853205 | AGTTGCTAAGTGCTGCCAAGAG | [] |
| 5518 | 18 | PIAS2 | {'exs': 0, 'nonexs': 7} | True | 46827942 | CTATAAATTAACAAACCCTTTG | [] |
| 5519 | 18 | ZBTB7C | {'exs': 0, 'nonexs': 6} | True | 48029259 | ACCAGCCTAGTTGTTGGCTTC | [] |
| 5520 | 18 | SMAD7 | {'exs': 1, 'nonexs': 6} | True | 48921371 | ACGCGGCTACCGGCTGTTGAA | [] |
| 5521 | 18 | CFAP53 | unavailable | False | 50227380 | GAACTACGGTGGAAGCTTACTG | [] |
| 5522 | 18 | MBD1 | {'exs': 0, 'nonexs': 7} | True | 50270010 | ATGAGTCTAAATGTCAATCAAG | ['CAATCAAATAAATTTCATATAG'] |
| 5523 | 18 | MBD1 | {'exs': 0, 'nonexs': 7} | True | 50271500 | CCTCCAGTCTACTGCTTTCTAG | [] |
| 5524 | 18 | ME2 | {'exs': 0, 'nonexs': 7} | True | 50947183 | TGCTTCTATTCTGTTATCACAG | [] |
| 5525 | 18 | RAB27B | {'exs': 0, 'nonexs': 7} | True | 54889412 | TATGTAGAGTCTAGCAGATACG | ['AGATACATTTCTTCTCTGGTGG'] |
| 5526 | 18 | RAB27B | {'exs': 0, 'nonexs': 7} | True | 54893072 | CTCTTTCTAACTGCTGCACTCG | ['CACTCACTCCACTGGTGAAGAG'] |
| 5527 | 18 | ZNF532 | {'exs': 1, 'nonexs': 7} | True | 58884465 | CTGTGGCTATTTCTCGGCTGA | [] |
| 5528 | 18 | RAX | {'exs': 0, 'nonexs': 7} | True | 59269003 | TGCCCCTAGAGGGCCTGCCAC | [] |
| 5529 | 18 | CCBE1 | {'exs': 0, 'nonexs': 7} | True | 59435907 | GTGCTATGGGTAGAAGTCTCTG | [] |
| 5530 | 18 | RELCH | unavailable | False | 62305533 | CCTGCTTCTACTTCTTTCTCTG | [] |
| 5531 | 18 | SERPINB4 | {'exs': 0, 'nonexs': 7} | True | 63637718 | CATCTATGGGGATGAGAATCTG | [] |
| 5532 | 18 | SERPINB3 | {'exs': 0, 'nonexs': 7} | True | 63655656 | CATCTACGGGGATGAGAATCTG | [] |
| 5533 | 18 | CDH7 | {'exs': 1, 'nonexs': 6} | True | 65880893 | GCTATGAGTACAAACTCTCTTG | [] |
| 5534 | 18 | CDH19 | {'exs': 0, 'nonexs': 7} | True | 66504811 | AAAAGCCCTAATTATTTGACTG | [] |
| 5535 | 18 | NETO1 | {'exs': 0, 'nonexs': 7} | True | 72749027 | TTCTTTCTAGACCCTAGTTGTG | [] |
| 5536 | 18 | NETO1 | {'exs': 0, 'nonexs': 7} | True | 72857212 | CTTAACCTGTCCTACTTCATGG | [] |
| 5537 | 18 | FBXO15 | {'exs': 0, 'nonexs': 7} | True | 74073460 | GCCACCTACTGCTAATATTCAG | [] |
| 5538 | 18 | FBXO15 | {'exs': 0, 'nonexs': 7} | True | 74126652 | TGCTATGCATGCCATAATCCAG | [] |
| 5539 | 18 | C18orf63 | unavailable | False | 74354512 | CATGTTCTAAGCATTATGAATG | [] |
| 5540 | 18 | DIPK1C | unavailable | False | 74436500 | GTGGCTACTTCTCTGCCTCCT | [] |
| 5541 | 18 | CNDP2 | {'exs': 0, 'nonexs': 7} | True | 74520067 | CTTGGCCTAGTCCTTCAGCTG | [] |
| 5542 | 18 | TSHZ1 | {'exs': 0, 'nonexs': 7} | True | 75288640 | CGCTACTGTTTCTCCAACTCAG | [] |
| 5543 | 18 | LOC10537 | unavailable | False | 76055028 | GCTGCACTATCCTCTGTTGGG | [] |
| 5544 | 18 | MBP | {'exs': 1, 'nonexs': 7} | True | 77016813 | AAAACTCCTCTACTCCTCAGAG | [] |
| 5545 | 18 | GALR1 | {'exs': 0, 'nonexs': 7} | True | 77265531 | CATTACCTAGTGGATCTGTGAG | [] |
| 5546 | 18 | SALL3 | {'exs': 0, 'nonexs': 6} | True | 78997321 | CTAGTTGATACCAATCTCCTTG | [] |
| 5547 | 18 | ATP9B | {'exs': 0, 'nonexs': 7} | True | 79359879 | AAATAGTCTATGGACCCTAGGG | [] |
| 5548 | 18 | ATP9B | {'exs': 0, 'nonexs': 7} | True | 79360146 | ACGTTCTATGCTATAAAATATG | [] |
| 5549 | 18 | ATP9B | {'exs': 0, 'nonexs': 7} | True | 79361889 | TGTCCTACATCGCAGGAGCTT | [] |
| 5550 | 18 | ATP9B | {'exs': 0, 'nonexs': 7} | True | 79376976 | ACTGCTAATGACTAGCTACTTG | [] |
| 5551 | 18 | LOC10798 | unavailable | False | 79346513 | TGTGCTAAATGTGTGCCGAATG | [] |
| 5552 | 18 | NFATC1 | {'exs': 0, 'nonexs': 7} | True | 79527576 | GGCAACTAGGAGTGGGTGCTC | [] |
| 5553 | 18 | RBFA | {'exs': 2, 'nonexs': 6} | False | 80046154 | CTCTCCATCTACTCCCTGCTT | [] |
| 5554 | 18 | PARD6G | {'exs': 0, 'nonexs': 7} | True | 80159770 | GGGAGTCTAGAGCGTGACCGC | [] |
| 5555 | 19 | OR4F17 | {'exs': 0, 'nonexs': 1} | False | 111595 | AAGATCTAAAACTTTACACTAG | [] |
| 5556 | 19 | C2CD4C | {'exs': 0, 'nonexs': 7} | True | 407095 | CCCCGCTCTACAGGAAGGGGA | [] |
| 5557 | 19 | SHC2 | {'exs': 0, 'nonexs': 7} | True | 421938 | ATTCTACAGTAGCTGGTCAAAG | [] |
| 5558 | 19 | ODF3L2 | {'exs': 0, 'nonexs': 7} | True | 463843 | CCTGGCCTAGCAGCAGCGGCC | [] |
| 5559 | 19 | PTBP1 | {'exs': 2, 'nonexs': 6} | False | 810825 | GTGCCCCTAGATGGTGGACTT | [] |
| 5560 | 19 | PLPPR3 | unavailable | False | 812569 | GCCGCGCTAGTCGGGGAAGCG | [] |
| 5561 | 19 | AZU1 | {'exs': 0, 'nonexs': 7} | True | 831876 | GGCCCCCCTAGGCTGGCCCCC | [] |
| 5562 | 19 | CFD | {'exs': 1, 'nonexs': 6} | True | 863237 | GGCACCCTAGGCCAGGACGCT | [] |
| 5563 | 19 | POLR2E | {'exs': 6, 'nonexs': 2} | False | 1089485 | CGGTAGCTACTGCACCAGCCG | [] |
| 5564 | 19 | GPX4 | {'exs': 2, 'nonexs': 7} | False | 1106571 | TGTGGAGCTAGAAATAGTGGGG | [] |
| 5565 | 19 | ATP5F1D | unavailable | False | 1244436 | CCTACTCCAGGGCCTTCACCA | [] |
| 5566 | 19 | MIDN | {'exs': 0, 'nonexs': 5} | True | 1257271 | CCGAAGATCCTAAGCCACCAC | [] |
| 5567 | 19 | CIRBP | {'exs': 0, 'nonexs': 7} | True | 1272599 | TAGGCTAGCAGAGAATGCTCAG | [] |
| 5568 | 19 | EFNA2 | {'exs': 3, 'nonexs': 5} | False | 1299944 | CTGGGACTAGGAACCCAGGAG | [] |
| 5569 | 19 | PWWP3A | unavailable | False | 1371446 | CTGGGTACTAAATGCGATTCTG | [] |
| 5570 | 19 | PWWP3A | unavailable | False | 1376156 | ACAGAGCTAAATTCTGTCTCTG | ['TAAAAAAAAAAAAAAAAAAAAG'] |
| 5571 | 19 | NDUFS7 | unavailable | False | 1395487 | GCGGCGCTACCTGCGGTACCA | [] |
| 5572 | 19 | DAZAP1 | {'exs': 1, 'nonexs': 7} | True | 1433826 | ACCTACCTAGGCCACATCACT | [] |
| 5573 | 19 | DAZAP1 | {'exs': 1, 'nonexs': 7} | True | 1434911 | CGCGGGCTAGCGTCGGTAGGG | [] |
| 5574 | 19 | APC2 | {'exs': 0, 'nonexs': 7} | True | 1470212 | AGGCCACTATTCCAGGAGGGT | [] |
| 5575 | 19 | PLK5 | {'exs': 0, 'nonexs': 6} | True | 1535249 | AGGGGCACTAGATACTCTGCA | [] |
| 5576 | 19 | MEX3D | {'exs': 0, 'nonexs': 7} | True | 1555562 | CCGCGCTCTAGGAAAAGATATG | [] |
| 5577 | 19 | MBD3 | {'exs': 1, 'nonexs': 6} | True | 1578339 | CCCTAGACGTGCTCCATCTCC | [] |
| 5578 | 19 | ATP8B3 | {'exs': 0, 'nonexs': 6} | True | 1789536 | TCGGAGCTACGGCGGGCTCTG | [] |
| 5579 | 19 | KLF16 | {'exs': 4, 'nonexs': 4} | False | 1854458 | AGGGCCCTACAGGCCTGCGGG | [] |
| 5580 | 19 | ABHD17A | unavailable | False | 1877199 | CCGCCGCTAGGCGCGCTGGCT | [] |
| 5581 | 19 | ADAT3 | {'exs': 3, 'nonexs': 3} | False | 1912150 | GCGCCTACGTGTCGGGGTCCA | [] |
| 5582 | 19 | SCAMP4 | {'exs': 3, 'nonexs': 5} | False | 1924283 | CCTCTAAGGCCACTGGCCACT | [] |
| 5583 | 19 | BTBD2 | {'exs': 0, 'nonexs': 7} | True | 1986487 | GGCAGCCTAGGTGTAGAAGAT | [] |
| 5584 | 19 | DOT1L | {'exs': 2, 'nonexs': 6} | False | 2229791 | AAATCCTAGTTACCTGCAGAAG | [] |
| 5585 | 19 | PLEKHJ1 | {'exs': 1, 'nonexs': 5} | False | 2230162 | CCGGTGCTATCTCGTCCCCAG | [] |
| 5586 | 19 | OAZ1 | {'exs': 2, 'nonexs': 4} | False | 2272107 | CCCTACTCCTCCTCCTCTCCC | [] |
| 5587 | 19 | LSM7 | {'exs': 6, 'nonexs': 2} | False | 2321679 | GGCCAGGCTAGGCGTCCTGCT | [] |
| 5588 | 19 | SPPL2B | unavailable | False | 2353208 | CTCCCCTAGGCCGAGGCGCCA | [] |
| 5589 | 19 | ZNF554 | {'exs': 0, 'nonexs': 7} | True | 2834851 | GCAATTGCTAATATCCAATAAG | [] |
| 5590 | 19 | TLE5 | unavailable | False | 3053818 | CCCCTGCTAATCCGACTTCTC | [] |
| 5591 | 19 | NFIC | {'exs': 0, 'nonexs': 7} | True | 3450599 | CTTGGTCCTAATCCCACAAAGG | [] |
| 5592 | 19 | NFIC | {'exs': 0, 'nonexs': 7} | True | 3462768 | CCTTTGCTATCCCAGATACCAG | [] |
| 5593 | 19 | NFIC | {'exs': 0, 'nonexs': 7} | True | 3462863 | TGTTCACTCTAATTTTCCACCG | [] |
| 5594 | 19 | DOHH | {'exs': 7, 'nonexs': 1} | False | 3491491 | GGGGCCCTAGGAGGGGGCCCC | [] |
| 5595 | 19 | GIPC3 | {'exs': 0, 'nonexs': 5} | True | 3590189 | AAACTAGCCACAGGCCTCTCT | [] |
| 5596 | 19 | GIPC3 | {'exs': 0, 'nonexs': 5} | True | 3593287 | GCCAAGCTACGGCCCCCCGCC | [] |
| 5597 | 19 | TBXA2R | {'exs': 1, 'nonexs': 7} | True | 3595687 | CACTTCCTACTGCAGCCCGGA | [] |
| 5598 | 19 | MRPL54 | {'exs': 4, 'nonexs': 4} | False | 3767392 | ATGCTACAACCTCTTGTTCTTG | [] |
| 5599 | 19 | DAPK3 | {'exs': 0, 'nonexs': 7} | True | 3959100 | CCACTGCGCCTAGCGCAGCCC | [] |
| 5600 | 19 | EEF2 | {'exs': 7, 'nonexs': 1} | False | 3976553 | GGCCGCCTACAATTTGTCCAG | [] |
| 5601 | 19 | EBI3 | {'exs': 0, 'nonexs': 7} | True | 4237087 | AAGCCCTTGCTACTTGCCCAG | [] |
| 5602 | 19 | TMIGD2 | {'exs': 0, 'nonexs': 7} | True | 4294386 | CTTGCTAGGTTGCACAGGCTA | [] |
| 5603 | 19 | FSD1 | {'exs': 0, 'nonexs': 7} | True | 4323642 | GGGGGCCTAGGTGAGGCTGGT | [] |
| 5604 | 19 | PLIN4 | {'exs': 0, 'nonexs': 7} | True | 4504458 | CTACAGCTACTGCCCGCCAGC | [] |
| 5605 | 19 | SEMA6B | {'exs': 0, 'nonexs': 5} | True | 4543600 | CCGGCCTAGGGCACGGGGGGG | [] |
| 5606 | 19 | SEMA6B | {'exs': 0, 'nonexs': 5} | True | 4546227 | GTCCCCTAAGCCTGAGGTGCT | [] |
| 5607 | 19 | DPP9 | {'exs': 0, 'nonexs': 7} | True | 4694519 | ACGTGACTAACGCGATGAGTC | [] |
| 5608 | 19 | PLIN3 | {'exs': 0, 'nonexs': 7} | True | 4839191 | TCCCCCTACTTCTTCTCCTCCC | [] |
| 5609 | 19 | KDM4B | {'exs': 1, 'nonexs': 6} | True | 5088297 | GAGCCGCTAAGAGCTGGATGT | [] |
| 5610 | 19 | KDM4B | {'exs': 1, 'nonexs': 6} | True | 5151510 | GCTGTCCTAGAAGGGGGCTCC | [] |
| 5611 | 19 | MICOS13 | unavailable | False | 5678550 | CTGACTCGCTACTTGGTGCGC | [] |
| 5612 | 19 | DUS3L | {'exs': 0, 'nonexs': 7} | True | 5785202 | CTGAGGCTACTTGTACGCGTT | [] |
| 5613 | 19 | RANBP3 | {'exs': 6, 'nonexs': 2} | False | 5917609 | GGCCGCTATGTGCTCCCGGTC | [] |
| 5614 | 19 | RFX2 | {'exs': 0, 'nonexs': 7} | True | 5994834 | GGGCTGCTAGATGCCCTGCAG | [] |
| 5615 | 19 | SLC25A41 | {'exs': 1, 'nonexs': 7} | True | 6426388 | CTCACAGCCTATATGCCCAGG | [] |
| 5616 | 19 | SLC25A23 | {'exs': 0, 'nonexs': 7} | True | 6436319 | GACTATCCAGGGAATTCAACTG | [] |
| 5617 | 19 | CRB3 | {'exs': 0, 'nonexs': 7} | True | 6467021 | GGGGACCTAGATGGGCAGGCA | [] |
| 5618 | 19 | TUBB4A | unavailable | False | 6495163 | GCCTAGGCCACCTCCTCCTCC | [] |
| 5619 | 19 | TNFSF14 | {'exs': 0, 'nonexs': 7} | True | 6661133 | TCCCCTATAAGCTCGCCGTCT | [] |
| 5620 | 19 | CAMSAP3 | unavailable | False | 7618056 | GTGGGGCTATTTGGGGGTGCC | [] |
| 5621 | 19 | CD209 | {'exs': 0, 'nonexs': 7} | True | 7743038 | GTTCTGCTACGCAGGAGGGGG | [] |
| 5622 | 19 | CLEC4M | {'exs': 0, 'nonexs': 7} | True | 7768987 | AAACAACTATTCGTCTCTGAAG | [] |
| 5623 | 19 | MAP2K7 | {'exs': 0, 'nonexs': 6} | True | 7912430 | CAAGCAGCTACCTGAAGAAGG | [] |
| 5624 | 19 | TGFBR3L | {'exs': 0, 'nonexs': 4} | True | 7918877 | GGAGGGGACGCTACGCGGGCC | ['GGGCCACCTCCCTTCCCAGCAG'] |
| 5625 | 19 | FBN3 | {'exs': 0, 'nonexs': 7} | True | 8065918 | CTCCCAACTAAAGCAACTGCAG | [] |
| 5626 | 19 | CERS4 | unavailable | False | 8262108 | GCCCGGCTATGTGGCTGTTGT | [] |
| 5627 | 19 | ANGPTL4 | {'exs': 0, 'nonexs': 7} | True | 8373885 | GGACGCTAGGAGGCTGCCTCT | [] |
| 5628 | 19 | ADAMTS10 | {'exs': 0, 'nonexs': 5} | True | 8580892 | CGCCCCCCTAGTGGCCATGGCA | [] |
| 5629 | 19 | OR7G1 | {'exs': 0, 'nonexs': 1} | False | 9114827 | AAGATCGCTAAAAGGAAACAG | [] |
| 5630 | 19 | OR7E24 | {'exs': 0, 'nonexs': 3} | True | 9252062 | TTCTATCCCATATAACAAAAAG | [] |
| 5631 | 19 | ZNF560 | {'exs': 0, 'nonexs': 7} | True | 9466573 | CTCTCTAGTGTGTTTTCAAATG | [] |
| 5632 | 19 | ZNF426 | {'exs': 0, 'nonexs': 7} | True | 9528379 | CTCACTAGTGAATTTGTTCATG | [] |
| 5633 | 19 | ZNF561 | {'exs': 0, 'nonexs': 7} | True | 9610199 | ATGGGCTAAATGGTAACACTCG | ['ACTCATATCCTTGCATTCATAG'] |
| 5634 | 19 | UBL5 | {'exs': 7, 'nonexs': 1} | False | 9830007 | TTCTCATCTATTGATAATAAAG | [] |
| 5635 | 19 | DNMT1 | {'exs': 8, 'nonexs': 0} | False | 10133666 | GGGCAGAACTAGTCCTTAGCA | [] |
| 5636 | 19 | MRPL4 | {'exs': 7, 'nonexs': 1} | False | 10258737 | GTGATTCTAACAGCGGAGCCT | [] |
| 5637 | 19 | MRPL4 | {'exs': 7, 'nonexs': 1} | False | 10259505 | TGTGGTCTAGTGTGACTTTGTG | [] |
| 5638 | 19 | AP1M2 | {'exs': 0, 'nonexs': 7} | True | 10573065 | CTCCCTTCTAGCTGGTACGAA | [] |
| 5639 | 19 | SLC44A2 | {'exs': 0, 'nonexs': 7} | True | 10643017 | ACTCTCTACTCCTCGGCTTCT | [] |
| 5640 | 19 | ILF3 | {'exs': 6, 'nonexs': 2} | False | 10684464 | TCTAGGAAGACCCAAAATCATG | [] |
| 5641 | 19 | DNM2 | {'exs': 7, 'nonexs': 1} | False | 10831046 | GAGGCCTAGTCGAGCAGGGAT | [] |
| 5642 | 19 | TMED1 | {'exs': 0, 'nonexs': 7} | True | 10832994 | CAGGGGCTACGTGGGCACCGG | [] |
| 5643 | 19 | CARM1 | {'exs': 1, 'nonexs': 7} | True | 10921756 | CGGGCCCCTAGCTCCCGTAGT | [] |
| 5644 | 19 | YIPF2 | {'exs': 1, 'nonexs': 7} | True | 10923290 | GCCTTCCTAGGAGGGGGCCAG | [] |
| 5645 | 19 | SPC24 | {'exs': 7, 'nonexs': 1} | False | 11147182 | GGCTCCTGGCTACCACTCGGT | [] |
| 5646 | 19 | KANK2 | {'exs': 0, 'nonexs': 5} | True | 11166557 | TCACGGCTACTCTTCTGCCGA | [] |
| 5647 | 19 | RAB3D | {'exs': 0, 'nonexs': 7} | True | 11325397 | CCATCTCTAGCAGCTGCAGCT | [] |
| 5648 | 19 | TMEM205 | {'exs': 0, 'nonexs': 7} | True | 11342814 | GCTAGAGGCTCCTTATTTCCAG | [] |
| 5649 | 19 | EPOR | {'exs': 1, 'nonexs': 6} | True | 11377983 | GTCCTAAGAGCAAGCCACATAG | [] |
| 5650 | 19 | CCDC151 | {'exs': 1, 'nonexs': 7} | True | 11420834 | ACGAGTCTAGGACCTCCGAGA | [] |
| 5651 | 19 | PRKCSH | {'exs': 1, 'nonexs': 5} | False | 11449411 | ATCCAGCTAGAGCTCGTCATGG | [] |
| 5652 | 19 | ECSIT | {'exs': 1, 'nonexs': 7} | True | 11506183 | GCTCAGACTAGCTCTGGCCCT | [] |
| 5653 | 19 | CNN1 | {'exs': 0, 'nonexs': 7} | True | 11549794 | GTGGCCCTAGGCGGAATTGTA | [] |
| 5654 | 19 | ELOF1 | {'exs': 3, 'nonexs': 3} | False | 11553745 | CTGTGTCGCTACTGATTGGCC | [] |
| 5655 | 19 | ZNF441 | {'exs': 0, 'nonexs': 7} | True | 11781905 | TTCTCTAGTGAGTCATTTCATG | [] |
| 5656 | 19 | ZNF700 | {'exs': 1, 'nonexs': 7} | True | 11950252 | AAGGAACCAGGCTAGCTGAATG | [] |
| 5657 | 19 | ZNF878 | {'exs': 0, 'nonexs': 4} | True | 12043804 | TAGAGTTTCTAGCCAGCGTGAG | [] |
| 5658 | 19 | ZNF799 | {'exs': 0, 'nonexs': 7} | True | 12390465 | AATGCTAGTGAGTCTTTTTATG | [] |
| 5659 | 19 | ZNF443 | {'exs': 0, 'nonexs': 7} | True | 12430155 | AATGCTAGTGAGTCTTTTTATG | [] |
| 5660 | 19 | ZNF564 | {'exs': 0, 'nonexs': 7} | True | 12526445 | CCTCCACTATTCATTTTCACAG | [] |
| 5661 | 19 | ZNF791 | {'exs': 0, 'nonexs': 7} | True | 12629259 | TATAGAGTTTCTATCGATTGTG | [] |
| 5662 | 19 | MAN2B1 | {'exs': 0, 'nonexs': 7} | True | 12646619 | CCTAACCATCCACCTCCTTCC | ['CCTTCCATTGAACTGAGGCCAG'] |
| 5663 | 19 | TNPO2 | {'exs': 0, 'nonexs': 6} | True | 12701345 | ATCACCTAGACCCCATAGAAAG | [] |
| 5664 | 19 | GET3 | unavailable | False | 12748103 | GCTGTGCTACTGGGCACTGGG | [] |
| 5665 | 19 | PRDX2 | {'exs': 4, 'nonexs': 5} | False | 12797080 | GCCAGCCTAATTGTGTTTGGAG | [] |
| 5666 | 19 | RNASEH2A | {'exs': 5, 'nonexs': 4} | False | 12813465 | AGCTGCTAGAGGCTGGTTGCT | [] |
| 5667 | 19 | RTBDN | {'exs': 1, 'nonexs': 7} | True | 12825705 | CGTCCGCTAGGGGCCGCTGCC | [] |
| 5668 | 19 | MAST1 | {'exs': 1, 'nonexs': 7} | True | 12874869 | CTTGGCTATGGTCCCGGGGGT | [] |
| 5669 | 19 | GCDH | {'exs': 0, 'nonexs': 7} | True | 12899742 | AGACGCTCTACAGAGAATTCCG | ['GAATTCCATCTTCTTTTTAAGG'] |
| 5670 | 19 | CALR | {'exs': 4, 'nonexs': 5} | False | 12943812 | GCCTCTCTACAGCTCGTCCTT | [] |
| 5671 | 19 | CACNA1A | {'exs': 0, 'nonexs': 7} | True | 13209045 | TACGGAACTACTGCCCTACACC | [] |
| 5672 | 19 | CCDC130 | {'exs': 6, 'nonexs': 2} | False | 13761561 | CTTGTGCTAGGCTTTGTTCTGG | [] |
| 5673 | 19 | NANOS3 | {'exs': 0, 'nonexs': 7} | True | 13880502 | AGCCTCCTAGGTGGACATGGA | [] |
| 5674 | 19 | C19orf57 | {'exs': 0, 'nonexs': 7} | True | 13882801 | AGTGGCCTACAACTCCCTCCA | [] |
| 5675 | 19 | C19orf57 | {'exs': 0, 'nonexs': 7} | True | 13885083 | TGTGGCTAAGGGAGGTGGCCA | [] |
| 5676 | 19 | RLN3 | {'exs': 0, 'nonexs': 7} | True | 14030947 | CTCAAACTAGCAAAGGCTACTG | [] |
| 5677 | 19 | PRKACA | {'exs': 1, 'nonexs': 6} | True | 14093111 | CAGGCATGCCCCTAAAACTCA | [] |
| 5678 | 19 | PKN1 | {'exs': 1, 'nonexs': 6} | True | 14471783 | GGAGGGGGCTAGCAGCCCCCC | [] |
| 5679 | 19 | GIPC1 | {'exs': 0, 'nonexs': 7} | True | 14478415 | GTCCTAGTAGCGGCCGACCTT | [] |
| 5680 | 19 | DNAJB1 | {'exs': 0, 'nonexs': 7} | True | 14515939 | ATAGCTATATTGGAAGAACCTG | [] |
| 5681 | 19 | NDUFB7 | {'exs': 4, 'nonexs': 4} | False | 14566132 | CACCCCCTACAGGGCCACCTT | [] |
| 5682 | 19 | CLEC17A | {'exs': 0, 'nonexs': 7} | True | 14599690 | GGCTTGAGGGCTACAGACTGA | [] |
| 5683 | 19 | ADGRE2 | unavailable | False | 14736235 | GAAGATTTTTCTAGTTAACCTG | [] |
| 5684 | 19 | OR7A10 | {'exs': 1, 'nonexs': 3} | False | 14840947 | GCCTTTCTATTGCTTTCCTCTG | [] |
| 5685 | 19 | CCDC105 | {'exs': 0, 'nonexs': 7} | True | 15023118 | GGGTCACTAGGGGTCCGCGCT | [] |
| 5686 | 19 | CASP14 | {'exs': 0, 'nonexs': 7} | True | 15056088 | TCTACTTCTACTGCAGATACAG | [] |
| 5687 | 19 | ILVBL | {'exs': 1, 'nonexs': 7} | True | 15115251 | AAGGCCCTATACAGCAATGGAG | [] |
| 5688 | 19 | EPHX3 | {'exs': 0, 'nonexs': 7} | True | 15227436 | ACCACTAGTCCAGCAGGTCTT | [] |
| 5689 | 19 | OR10H1 | {'exs': 0, 'nonexs': 3} | True | 15807080 | GAATTTCTCCTACATCATTACG | ['CATTACATTTTTTTCTGGGTAG'] |
| 5690 | 19 | HSH2D | {'exs': 0, 'nonexs': 1} | False | 16157793 | TGTTCTCTAGCAGTACCCAGG | [] |
| 5691 | 19 | FAM32A | {'exs': 4, 'nonexs': 4} | False | 16190954 | AGGCGGCTACTTCGTCCAGCT | [] |
| 5692 | 19 | FAM32A | {'exs': 4, 'nonexs': 4} | False | 16191148 | AGAAGCCTAAACTGAGTACAAG | [] |
| 5693 | 19 | KLF2 | {'exs': 0, 'nonexs': 7} | True | 16327030 | CGGCTACATGTGCCGTTTCAT | [] |
| 5694 | 19 | EPS15L1 | {'exs': 1, 'nonexs': 7} | True | 16355704 | CCTCGCCTAGGCGGCAGGCAT | [] |
| 5695 | 19 | CALR3 | {'exs': 0, 'nonexs': 6} | True | 16479130 | TCACTAAAGTTCATTCCTTCTG | [] |
| 5696 | 19 | CHERP | {'exs': 6, 'nonexs': 2} | False | 16519158 | CGCCTACTTACACTCGTCCCT | [] |
| 5697 | 19 | SLC35E1 | {'exs': 0, 'nonexs': 7} | True | 16553678 | TGGACTCTACACATCATAGCGG | [] |
| 5698 | 19 | SMIM7 | unavailable | False | 16652861 | CTTTTCCTAAATCCATTTACAG | [] |
| 5699 | 19 | TMEM38A | {'exs': 0, 'nonexs': 7} | True | 16688370 | CACCCCCTAATCCGCCTTCTT | [] |
| 5700 | 19 | CPAMD8 | {'exs': 0, 'nonexs': 7} | True | 16894162 | GCTGTAGCTAGGAAGGTGGGG | [] |
| 5701 | 19 | USHBP1 | {'exs': 1, 'nonexs': 7} | True | 17250224 | GTAAGGGGCCTACAGAAAGGT | [] |
| 5702 | 19 | USHBP1 | {'exs': 1, 'nonexs': 7} | True | 17258595 | TCCAGCTACTCAGGAGGCTGA | [] |
| 5703 | 19 | ANKLE1 | {'exs': 2, 'nonexs': 6} | False | 17286714 | GCTCCCTACACACACACACAC | ['ACACACACACACACCCCTTCTG', 'CACACACACACACACACACACG', 'CACACACACAAACACACACACG'] |
| 5704 | 19 | ABHD8 | {'exs': 1, 'nonexs': 7} | True | 17292660 | AGCGGCTACTTCTTGTCTTCTG | [] |
| 5705 | 19 | ANO8 | {'exs': 0, 'nonexs': 7} | True | 17323516 | GTAGAGCTAATGCCAGCCGCT | [] |
| 5706 | 19 | MVB12A | unavailable | False | 17424992 | GAGGGACTATGAGACGCTGGG | [] |
| 5707 | 19 | PGLS | {'exs': 1, 'nonexs': 7} | True | 17521080 | TGGCCAGCTACAAAGTGGAATG | [] |
| 5708 | 19 | NIBAN3 | unavailable | False | 17553683 | TAACATCTACATCCCGTCCGG | [] |
| 5709 | 19 | NIBAN3 | unavailable | False | 17555158 | GCTTGCTATATGTGAGCCTCAG | [] |
| 5710 | 19 | UNC13A | {'exs': 0, 'nonexs': 5} | True | 17606053 | CCCGCGCTAAGGCGCAGGCGC | [] |
| 5711 | 19 | MAP1S | {'exs': 0, 'nonexs': 7} | True | 17734427 | ATGGGGCTAGAACTCCACCTT | [] |
| 5712 | 19 | JAK3 | {'exs': 0, 'nonexs': 5} | True | 17826742 | CAGGAGCTATGAAAAGGACAGG | [] |
| 5713 | 19 | RPL18A | {'exs': 7, 'nonexs': 1} | False | 17863262 | CTGCACCTAGAAGAAGGTGTTG | [] |
| 5714 | 19 | IL12RB1 | {'exs': 1, 'nonexs': 7} | True | 18071208 | TCTACTAAAAATACAAAATTAG | [] |
| 5715 | 19 | PDE4C | {'exs': 1, 'nonexs': 6} | True | 18210928 | CTGGCCCTAAGTCCTCTGGTT | [] |
| 5716 | 19 | IQCN | unavailable | False | 18257179 | GGTCCTAGATGCCAGGCCAAT | [] |
| 5717 | 19 | PGPEP1 | {'exs': 1, 'nonexs': 7} | True | 18369897 | GTCTCCTAGAGCCCAGCCCCA | [] |
| 5718 | 19 | SSBP4 | {'exs': 2, 'nonexs': 6} | False | 18434280 | AGAAGCCTAGGCCCGCAGAGA | [] |
| 5719 | 19 | ELL | {'exs': 8, 'nonexs': 1} | False | 18444751 | AGGGCGGCTAGGGCCAAGCCT | [] |
| 5720 | 19 | KXD1 | {'exs': 0, 'nonexs': 6} | True | 18568630 | GGGCAGCAGGGCTATTCGCCC | [] |
| 5721 | 19 | TMEM59L | {'exs': 0, 'nonexs': 5} | True | 18620535 | GGAGGCCTACAGCTTGGTCAG | [] |
| 5722 | 19 | KLHL26 | {'exs': 0, 'nonexs': 7} | True | 18669244 | TGGGGGCTACCTCCTGGTCCC | [] |
| 5723 | 19 | COMP | {'exs': 0, 'nonexs': 7} | True | 18792914 | CTGGTCCCTAGGCTTGCCGCA | [] |
| 5724 | 19 | SUGP2 | {'exs': 1, 'nonexs': 7} | True | 18894365 | AACATACCTATTTGTTGGCCCG | [] |
| 5725 | 19 | SLC25A42 | {'exs': 1, 'nonexs': 7} | True | 19110875 | GGGTCCCCTAGCTCTGCAGGT | [] |

| | | | | | | | |
|---|---|---|---|---|---|---|---|
| 5726 | 19 | TMEM161 | {'ess': 0, 'noness': 7} | True | 19119929 | AGGCAGCTAGGAGCCTGCCAA | [] |
| 5727 | 19 | MEF2B | {'ess': 2, 'noness': 5} | False | 19145796 | GATCTCCTACCGGGGCCAGCC | [] |
| 5728 | 19 | BORCS8 | unavailable | False | 19182568 | CAGGAGCTACCTGTGGCCCTG | [] |
| 5729 | 19 | HAPLN4 | {'ess': 0, 'noness': 7} | True | 19257816 | CCCAGCCTAGACGTGCAGAGG | [] |
| 5730 | 19 | GATAD2A | {'ess': 3, 'noness': 4} | False | 19505473 | TCGCACTATTTCCACGTGGCTC | [] |
| 5731 | 19 | TSSK6 | {'ess': 1, 'noness': 4} | False | 19514605 | CGGCTTCTAGCCGGAGTCCCC | [] |
| 5732 | 19 | NDUFA13 | {'ess': 5, 'noness': 1} | False | 19528125 | CAGGGCCTACGTGTACCACATC | [] |
| 5733 | 19 | LPAR2 | {'ess': 1, 'noness': 7} | True | 19624255 | TCAAGGTAGCTAAAGGGTGGAG | [] |
| 5734 | 19 | ZNF93 | unavailable | False | 19934817 | CACTTCTAGGGTTTCTCCCCAC | [] |
| 5735 | 19 | ZNF66 | unavailable | False | 20807321 | TAGAGGAGTGTCTAAGGCTTTG | [] |
| 5736 | 19 | ZNF85 | {'ess': 2, 'noness': 6} | False | 20942834 | CTTGCTACTGCATTCCAGACTG | [] |
| 5737 | 19 | ZNF738 | {'ess': 0, 'noness': 3} | True | 21377482 | CTAAGGTTCCAAATATATAAAG | [] |
| 5738 | 19 | ZNF208 | {'ess': 0, 'noness': 7} | True | 21971190 | CATTTCTAGAGTTTCTCTCCAG | [] |
| 5739 | 19 | ZNF681 | {'ess': 0, 'noness': 7} | True | 23743611 | CATTTCTAAGATTTCTCACCAG | [] |
| 5740 | 19 | C19orf12 | {'ess': 0, 'noness': 7} | True | 29702711 | TGCGGCCTAGTCATCATACTGC | [] |
| 5741 | 19 | URI1 | {'ess': 4, 'noness': 4} | False | 30015068 | ACAGGGCCTAGTCTTTCTGTTG | [] |
| 5742 | 19 | TSHZ3 | {'ess': 0, 'noness': 7} | True | 31276546 | TGCTACTGCTTCTCTAACTCAG | [] |
| 5743 | 19 | THEG5 | unavailable | False | 31593038 | CGCAGTCCTAAAACCGTGTAGC | [] |
| 5744 | 19 | ZNF507 | {'ess': 0, 'noness': 6} | True | 32383082 | TCCACCTAATTTGTGTTTAGAG | [] |
| 5745 | 19 | DPY19L3 | {'ess': 0, 'noness': 7} | True | 32482239 | TGCGCTACTTGTTTCTGGACAG | [] |
| 5746 | 19 | PDCD5 | {'ess': 3, 'noness': 6} | False | 32580038 | GTATGACTATTCTCTCCAACAG | ['AACAATTTCTTCTGAGAGTCAG'] |
| 5747 | 19 | NUDT19 | {'ess': 0, 'noness': 7} | True | 32711956 | CCCTACAAATGGCTTTTTCTTG | ['TTTCTTACTACAGAGTTCTTAG'] |
| 5748 | 19 | TDRD12 | {'ess': 0, 'noness': 7} | True | 32827165 | AATTCACTACTAAAGGAGTAAG | [] |
| 5749 | 19 | CEP89 | unavailable | False | 32879161 | GATTCTAGCAGGTGGGGGCATC | [] |
| 5750 | 19 | GPATCH1 | {'ess': 4, 'noness': 5} | False | 33113776 | AGCCAACTAAAGTTGAACTGGG | [] |
| 5751 | 19 | SLC7A10 | {'ess': 0, 'noness': 7} | True | 33208879 | AGTCTCTACAAAAATCTCATTG | [] |
| 5752 | 19 | CEBPG | {'ess': 1, 'noness': 6} | True | 33379591 | TGAGGTCTACTGTCCTGCATTG | [] |
| 5753 | 19 | PEPD | {'ess': 0, 'noness': 7} | True | 33387343 | CTGGCTCTACTTGGGGCCAGA | [] |
| 5754 | 19 | KCTD15 | {'ess': 0, 'noness': 7} | True | 33812947 | GGGCCTAGTCCAGGGGTTCCT | [] |
| 5755 | 19 | LSM14A | {'ess': 0, 'noness': 7} | True | 34227387 | TGTAGACTATGCAGCAACTTTG | [] |
| 5756 | 19 | PDCD2L | {'ess': 1, 'noness': 7} | True | 34426119 | AAATGCTCTACTTAAACAATAG | ['ATAATTCATCTGGGTCTTCTTG'] |
| 5757 | 19 | ZNF302 | {'ess': 0, 'noness': 7} | True | 34685236 | TGCATTTCTAAACTTCAAACGG | [] |
| 5758 | 19 | ZNF792 | {'ess': 1, 'noness': 7} | True | 34957955 | CATTCCTAGGGAACCTCCCCAC | [] |
| 5759 | 19 | SCN1B | {'ess': 0, 'noness': 7} | True | 35039700 | CCAGGGCTATTCGGCCACCTG | [] |
| 5760 | 19 | FXYD1 | {'ess': 0, 'noness': 6} | True | 35142741 | GTGTTTCTACCGCCTGCGGGT | [] |
| 5761 | 19 | HAMP | {'ess': 0, 'noness': 7} | True | 35285041 | AGGTAGGTTCTACGTCTTGCAG | [] |
| 5762 | 19 | FFAR3 | {'ess': 1, 'noness': 7} | True | 35359930 | GAGGACCTAGCTTTCAGCACAC | [] |
| 5763 | 19 | GPR42 | {'ess': 0, 'noness': 1} | False | 35372399 | GAGGACCTAGTTTTCAGCACAG | [] |
| 5764 | 19 | FFAR2 | {'ess': 0, 'noness': 5} | True | 35450706 | AAACTGCTACTCTGTAGTGAAG | [] |
| 5765 | 19 | DMKN | {'ess': 0, 'noness': 7} | True | 35498715 | CCTACCAAAACTTCACCCACTG | [] |
| 5766 | 19 | TMEM147 | {'ess': 2, 'noness': 6} | False | 35547446 | CCAAGCCTAGGAGTGCACATTC | [] |
| 5767 | 19 | ATP4A | {'ess': 0, 'noness': 7} | True | 35550614 | CCCTCTAATAGTAGAGTTCCTG | [] |
| 5768 | 19 | RBM42 | {'ess': 3, 'noness': 5} | False | 35637553 | ACAGACCCTATCTCAGGCCCAC | [] |
| 5769 | 19 | HSPB6 | {'ess': 0, 'noness': 5} | True | 35755521 | CCCTCCTACTTGGCTGCGGCT | [] |
| 5770 | 19 | ARHGAP33 | {'ess': 0, 'noness': 7} | True | 35785915 | ATGGCTAGAACTGAGCAGTATG | [] |
| 5771 | 19 | PRODH2 | {'ess': 0, 'noness': 7} | True | 35800037 | GGTGCTAGTGGGGTATCCTTCC | [] |
| 5772 | 19 | NFKBID | {'ess': 0, 'noness': 7} | True | 35888558 | GAGTCCTAAGAGGACAGGCCTC | [] |
| 5773 | 19 | LRFN3 | {'ess': 0, 'noness': 5} | True | 35945018 | CGCCTGGCTAGGGTCCCACAG | [] |
| 5774 | 19 | CLIP3 | {'ess': 0, 'noness': 7} | True | 36016157 | GGCCTCTAAGACTGCATCTCCC | [] |
| 5775 | 19 | ZNF565 | {'ess': 0, 'noness': 7} | True | 36182465 | TTTCTAACCAGTATGAATTCTG | [] |
| 5776 | 19 | ZNF567 | {'ess': 0, 'noness': 6} | True | 36723222 | TGACGCTAGGATAAACATCCAG | [] |
| 5777 | 19 | ZNF875 | unavailable | False | 37363774 | TAAAGTTTCTATCCTGAGTGTG | [] |
| 5778 | 19 | ZNF569 | {'ess': 0, 'noness': 7} | True | 37412596 | CATAGGGTTTCTAATGAGTATG | [] |
| 5779 | 19 | ZNF570 | {'ess': 1, 'noness': 7} | True | 37485232 | TCAGGATCTATAGGACTTGGTG | [] |
| 5780 | 19 | DPF1 | {'ess': 0, 'noness': 7} | True | 38212062 | GCCGGCCTAGGTGAGGGTGAT | [] |
| 5781 | 19 | KCNK6 | {'ess': 0, 'noness': 7} | True | 38325175 | TGTGCTACTGCATTCCAGCCTC | [] |
| 5782 | 19 | KCNK6 | {'ess': 0, 'noness': 7} | True | 38327402 | CCCAGCTACCTGGGGATGGAA | [] |
| 5783 | 19 | RASGRP4 | {'ess': 0, 'noness': 7} | True | 38410039 | AGATGTCTAGGAATCCAGCTTG | [] |
| 5784 | 19 | MAP4K1 | {'ess': 0, 'noness': 7} | True | 38587809 | CACCACTACAGGCCTGTGGAAC | [] |
| 5785 | 19 | CAPN12 | {'ess': 0, 'noness': 7} | True | 38730851 | CGGAGATCCTAGGAGAAGGTGC | [] |
| 5786 | 19 | RINL | {'ess': 0, 'noness': 5} | True | 38869103 | TGAAACCCCTAGTTGTCACTGG | [] |
| 5787 | 19 | NFKBIB | {'ess': 0, 'noness': 6} | True | 38907706 | CCGGTCTATCCTGACCCCCCA | [] |
| 5788 | 19 | CCER2 | unavailable | False | 38910304 | ACTTGCTAGTGATGGTGGCTTG | [] |
| 5789 | 19 | FBXO17 | {'ess': 1, 'noness': 6} | True | 38942607 | TCGCTAGGACAGACGGATCCTC | [] |
| 5790 | 19 | FBXO27 | {'ess': 0, 'noness': 7} | True | 39025410 | TGGACTAGGACAGACGGACTCC | [] |
| 5791 | 19 | ACP7 | unavailable | False | 39110067 | CATCTACGATCTTCCCATCCTG | [] |
| 5792 | 19 | ACP7 | unavailable | False | 39110117 | ATCCCTAGAGGTACATCCTCCC | [] |
| 5793 | 19 | RPS16 | {'ess': 7, 'noness': 1} | False | 39433349 | GGTCAGCTACCAGCAGGGTCC | [] |
| 5794 | 19 | SELENOV | unavailable | False | 39519147 | CCTTGCCGGCTACCTTTTCTTC | [] |
| 5795 | 19 | EID2 | {'ess': 0, 'noness': 7} | True | 39539368 | ACCACATAACTATTCTCTATTG | [] |
| 5796 | 19 | C19orf47 | {'ess': 0, 'noness': 7} | True | 40295318 | GGCCTAAACAGCTGTCTTATTG | [] |
| 5797 | 19 | C19orf47 | {'ess': 0, 'noness': 7} | True | 40321881 | GGTGGGCTAGAAGGTCCTGCG | [] |
| 5798 | 19 | BLVRB | {'ess': 0, 'noness': 7} | True | 40447888 | AGAGTGCTACTGGTACTGGTGC | [] |
| 5799 | 19 | SPTBN4 | {'ess': 0, 'noness': 7} | True | 40524615 | TGGCCTCTAGCTGCTGGATCTC | [] |
| 5800 | 19 | NUMBL | {'ess': 0, 'noness': 7} | True | 40657457 | CTCGGGCTACAGTTCAATCTCC | [] |
| 5801 | 19 | C19orf54 | {'ess': 0, 'noness': 7} | True | 40741704 | TGTGCTAGTCATGTGACCTCAG | [] |
| 5802 | 19 | C19orf54 | {'ess': 0, 'noness': 7} | True | 40742165 | AATGTCTAGTGAGATGACAGAG | [] |
| 5803 | 19 | C19orf54 | {'ess': 0, 'noness': 7} | True | 40742432 | GCAGCTAATGGCTGCAGTCCTC | [] |
| 5804 | 19 | SNRPA | {'ess': 5, 'noness': 3} | False | 40765166 | AAGGTGCTACTTCTTGGCAAAG | [] |
| 5805 | 19 | EGLN2 | {'ess': 2, 'noness': 4} | False | 40807953 | GGCCACTAGGTGGGCGTAGGC | [] |
| 5806 | 19 | HNRNPUL1 | {'ess': 0, 'noness': 7} | True | 41306564 | CACTGGCTACTGTGTACTTGTG | [] |
| 5807 | 19 | TMEM91 | {'ess': 0, 'noness': 7} | True | 41383872 | GGGCAACTAGGGCGGGTCTCG | [] |
| 5808 | 19 | DMAC2 | unavailable | False | 41432230 | TAAAGGCTAGGCAGGGACAGGC | [] |
| 5809 | 19 | CEACAM21 | {'ess': 0, 'noness': 7} | True | 41585870 | TCTTACCTAGGAGATGGAGCTG | [] |
| 5810 | 19 | CEACAM4 | {'ess': 1, 'noness': 6} | True | 41618099 | GTGAACCTATATGAGATCATCG | ['GAGATCATCACCCTTCCAGGGG'] |
| 5811 | 19 | CEACAM4 | {'ess': 1, 'noness': 6} | True | 41619329 | GGAACCTAAGAGACCACATCTG | [] |
| 5812 | 19 | CEACAM7 | {'ess': 0, 'noness': 7} | True | 41677411 | TGCTATATCAGAGCCATCCCAG | [] |
| 5813 | 19 | CEACAM5 | {'ess': 0, 'noness': 7} | True | 41727315 | CACCAGGGCTGCTATATCAGAC | [] |
| 5814 | 19 | CEACAM6 | {'ess': 0, 'noness': 7} | True | 41769258 | CTGCTATATCAGAGCCACCCTC | [] |
| 5815 | 19 | CEACAM3 | {'ess': 0, 'noness': 7} | True | 41811236 | GGAAGCTAAGAGCCACTTCTG | [] |
| 5816 | 19 | LYPD4 | {'ess': 0, 'noness': 4} | True | 41837285 | AATCTGCTAAAAGCTGGTTATG | [] |
| 5817 | 19 | DMRTC2 | {'ess': 1, 'noness': 7} | True | 41851695 | GGGTTTCTAGCTCAGGAGGGAC | [] |
| 5818 | 19 | RPS19 | {'ess': 7, 'noness': 1} | False | 41871376 | TTTGTTCTAATGCTTCTTGTTG | [] |
| 5819 | 19 | CXCL17 | {'ess': 1, 'noness': 7} | True | 42428883 | CAGAGCTCCTACAAAGGCAGAC | [] |
| 5820 | 19 | CEACAM8 | {'ess': 1, 'noness': 6} | True | 42583245 | CTACTATATCAGAGCCACCCTG | [] |
| 5821 | 19 | PSG3 | {'ess': 0, 'noness': 7} | True | 42723881 | CGGCTGCTATAATGGATTAAGG | [] |
| 5822 | 19 | PSG8 | {'ess': 0, 'noness': 7} | True | 42754294 | GACTCCACCTAAATCCCTATTG | [] |
| 5823 | 19 | PSG1 | {'ess': 1, 'noness': 6} | True | 42868062 | GACTCCACCTAAAACCCTATTG | [] |
| 5824 | 19 | PSG11 | {'ess': 0, 'noness': 7} | True | 43009997 | CGGCTGCTACGTTGGATTATTG | [] |
| 5825 | 19 | PSG2 | {'ess': 0, 'noness': 7} | True | 43066556 | CAGCTGCTATGTTGGATTAAGG | [] |
| 5826 | 19 | PSG5 | {'ess': 0, 'noness': 7} | True | 43170094 | CGGCTGCTATATTGGATTAAGG | [] |
| 5827 | 19 | PSG4 | {'ess': 1, 'noness': 6} | True | 43194301 | GACTCCACCTAAATCCCTATTG | [] |
| 5828 | 19 | ZNF428 | {'ess': 1, 'noness': 7} | True | 43607616 | CTCTGCCTACACCTCACCCCG | [] |
| 5829 | 19 | SRRM5 | {'ess': 0, 'noness': 7} | True | 43614268 | GGGGGCTACGCCAGCTTGGAA | [] |
| 5830 | 19 | SMG9 | unavailable | False | 43728967 | CAGCCCATCCTAGAGCAGAAGC | [] |
| 5831 | 19 | KCNN4 | {'ess': 0, 'noness': 7} | True | 43767542 | CACCAGCTACTTGGACTGCTGC | [] |
| 5832 | 19 | LYPD5 | {'ess': 0, 'noness': 7} | True | 43797590 | GAGGGGCGGTCTATGCTGAGA | [] |
| 5833 | 19 | ZNF224 | {'ess': 0, 'noness': 6} | True | 44108283 | ACTAAGGTTTTTCTCCAACATG | [] |
| 5834 | 19 | ZNF227 | {'ess': 0, 'noness': 7} | True | 44236829 | CATTTCTAAAGCTTTTTACCAG | [] |
| 5835 | 19 | ZNF235 | {'ess': 0, 'noness': 7} | True | 44287217 | CATTTCTACAGATTCCCTCCAG | [] |
| 5836 | 19 | ZNF229 | {'ess': 0, 'noness': 4} | True | 44428302 | CATCTACTTATAAGGGTTCTCG | [] |
| 5837 | 19 | ZNF180 | {'ess': 0, 'noness': 7} | True | 44476401 | TACAAACTAGTTACATTCATAG | [] |
| 5838 | 19 | IGSF23 | unavailable | False | 44635419 | ATTCCTATTCTCTGCCTGTCAG | [] |
| 5839 | 19 | IGSF23 | unavailable | False | 44636469 | GACTTGGAAGCTATGGGATGGC | [] |
| 5840 | 19 | CLPTM1 | {'ess': 1, 'noness': 6} | True | 44992896 | TCTCGACTAATCCTTTTTCTTG | [] |
| 5841 | 19 | RELB | {'ess': 2, 'noness': 5} | False | 45037789 | CGCGGGGCTACGTGGCTTCAG | [] |
| 5842 | 19 | CLASRP | {'ess': 7, 'noness': 1} | False | 45070844 | TTCTGCCTAATGTCGGTAATGG | [] |
| 5843 | 19 | BLOC1S3 | {'ess': 2, 'noness': 6} | False | 45179904 | CATGGCTAGGCCCGCGGCCCC | [] |
| 5844 | 19 | CKM | {'ess': 1, 'noness': 7} | True | 45306749 | AGGCGCCTACTTCTGGGCGGG | [] |
| 5845 | 19 | PPP1R13L | {'ess': 0, 'noness': 7} | True | 45380189 | CCTGCTAGACTTTACTCCTTTG | [] |
| 5846 | 19 | CD3EAP | {'ess': 7, 'noness': 2} | False | 45409500 | GGGCAGACTACACAGGCTGCT | [] |
| 5847 | 19 | OPA3 | {'ess': 0, 'noness': 7} | True | 45529055 | ACCTCCTATTTCTCGGACGCC | [] |
| 5848 | 19 | OPA3 | {'ess': 0, 'noness': 7} | True | 45553513 | AGCTCCTATTTCTTGGACGCAG | [] |
| 5849 | 19 | GIPR | {'ess': 0, 'noness': 7} | True | 45681934 | GCCCCCTAGCAGTAACTTTCC | ['TAACTTTCCAACTCCCGGCTGG'] |
| 5850 | 19 | SNRPD2 | {'ess': 7, 'noness': 1} | False | 45687552 | CGGCCCCTACTTGCCGGCGAT | [] |
| 5851 | 19 | QPCTL | {'ess': 1, 'noness': 7} | True | 45698786 | AGAAAGGCTAGGACTGGAGGGC | [] |
| 5852 | 19 | QPCTL | {'ess': 1, 'noness': 7} | True | 45703048 | CCAAGCACGCTAGAGCCCCAG | [] |
| 5853 | 19 | DMPK | {'ess': 2, 'noness': 6} | False | 45770480 | CAGTTCTAGGGTTCAGGGAGCC | [] |
| 5854 | 19 | FOXA3 | {'ess': 2, 'noness': 6} | False | 45873057 | ACCCCTGCTAGGATGCATTAAG | [] |
| 5855 | 19 | IRF2BP1 | {'ess': 1, 'noness': 6} | True | 45884019 | CAGTGGTAGCCTAGGGGTCCC | [] |
| 5856 | 19 | IGFL4 | {'ess': 0, 'noness': 7} | True | 46039891 | AGAACAGTCTAGATGAGGTCAG | [] |
| 5857 | 19 | PPP5D1 | unavailable | False | 46519813 | TCTATAAAAATAAAAAAATTAG | [] |
| 5858 | 19 | PNMA8B | unavailable | False | 46493557 | GGGCCACTAGCGGCATTTAGG | [] |
| 5859 | 19 | ZC3H4 | {'ess': 7, 'noness': 1} | False | 47066355 | TGGACACTACTGGCAAAAGGGC | [] |
| 5860 | 19 | BBC3 | {'ess': 0, 'noness': 7} | True | 47221700 | CAGTATGCTACATGGTGCAGAG | [] |
| 5861 | 19 | BBC3 | {'ess': 0, 'noness': 7} | True | 47221801 | CACCTAATTGGGCTCCATCTCC | [] |
| 5862 | 19 | C5AR1 | {'ess': 0, 'noness': 7} | True | 47320829 | GCTGTCGCCTACACTGCCTGG | [] |
| 5863 | 19 | C5AR2 | unavailable | False | 47341812 | CCAGCCTACACCTCCATCTCC | [] |
| 5864 | 19 | MEIS3 | {'ess': 1, 'noness': 7} | True | 47406476 | GCATCAGCCTCTATAGATAATG | [] |
| 5865 | 19 | SLC8A2 | {'ess': 0, 'noness': 7} | True | 47430088 | GAGGCCCCTAGAAGCCCCGGAT | [] |
| 5866 | 19 | NOP53 | unavailable | False | 47757004 | GGCAGCTACAACCTGGGGACAC | [] |
| 5867 | 19 | TPRX1 | {'ess': 0, 'noness': 6} | True | 47801774 | CACAGGCCCCCTATAAATCCAC | [] |
| 5868 | 19 | CRX | {'ess': 0, 'noness': 6} | True | 47839966 | ACTGCGTCCTCTACAAGATCTG | [] |
| 5869 | 19 | PLA2G4C | {'ess': 0, 'noness': 7} | True | 48048342 | GAGGCTCATCTATGCCAAGCAC | [] |
| 5870 | 19 | PLA2G4C | {'ess': 0, 'noness': 7} | True | 48052992 | GTCTCCCACCTACCCGGCCAC | [] |
| 5871 | 19 | PLA2G4C | {'ess': 0, 'noness': 7} | True | 48064983 | GCAGACTAAGACAGTACCTCTG | [] |
| 5872 | 19 | CARD8 | {'ess': 0, 'noness': 7} | True | 48215366 | CTGATGCAGCTACAAGCTGGAC | [] |
| 5873 | 19 | KDELR1 | {'ess': 0, 'noness': 7} | True | 48383292 | CGGGGCTATGCCGGCAAACTC | ['AACTCAACTTCTTCCCCTTTAG'] |
| 5874 | 19 | NTN5 | {'ess': 0, 'noness': 7} | True | 48661676 | TCACGTCTAGTGCTCCGGCCT | [] |
| 5875 | 19 | NTN5 | {'ess': 0, 'noness': 7} | True | 48663176 | CCCTATCTCTTCCATCCCTCAC | [] |
| 5876 | 19 | TULP2 | {'ess': 0, 'noness': 7} | True | 48881010 | GCCAGCTTCTAATTGAAACTGG | [] |
| 5877 | 19 | CGB1 | {'ess': 0, 'noness': 6} | True | 49035609 | GGGTGTCCTAGGGCCCCCGGGA | [] |
| 5878 | 19 | C19orf73 | {'ess': 0, 'noness': 7} | True | 49118632 | AGAGACTAGTCCGAGGGCGGA | [] |
| 5879 | 19 | PPFIA3 | {'ess': 0, 'noness': 7} | True | 49150137 | TGCACTAGCAGGAATAGGTCCC | [] |
| 5880 | 19 | CD37 | {'ess': 0, 'noness': 7} | True | 49340327 | CGGGGCCTAACGGTATCGAGC | [] |
| 5881 | 19 | DKKL1 | {'ess': 0, 'noness': 7} | True | 49375027 | CCACCCCTACAGCTGCCGAGA | [] |
| 5882 | 19 | FLT3LG | {'ess': 1, 'noness': 7} | True | 49486064 | GTCCAGGCTATGCATCCTCTGC | [] |
| 5883 | 19 | RRAS | {'ess': 0, 'noness': 7} | True | 49635575 | CCTGGGCTACAGGAGGACGCA | [] |
| 5884 | 19 | ADM5 | unavailable | False | 49690492 | TCCAACCTAGTCGCAGCCCGG | [] |
| 5885 | 19 | MED25 | {'ess': 2, 'noness': 6} | False | 49838823 | ACAAACCCTATTGACATTTGAG | [] |
| 5886 | 19 | PTOV1 | {'ess': 0, 'noness': 5} | True | 49860266 | CCCATCTACGGCGAGAAATCAC | [] |
| 5887 | 19 | PTOV1 | {'ess': 0, 'noness': 5} | True | 49860278 | AACCACTACCCCCCCATCTACC | [] |
| 5888 | 19 | ATF5 | {'ess': 1, 'noness': 6} | True | 49933091 | CTGCCCTTCTAGCAGCTACGGC | [] |
| 5889 | 19 | VRK3 | {'ess': 1, 'noness': 6} | True | 49979093 | TTCCACCTAGGGCACCATCGGC | [] |
| 5890 | 19 | ZNF473 | {'ess': 0, 'noness': 7} | True | 50047058 | CCAATGGCTACAGGCAGTATTG | [] |
| 5891 | 19 | IZUMO2 | {'ess': 0, 'noness': 7} | True | 50153688 | CCCAGCTACTCAGGAGGCTGA | [] |
| 5892 | 19 | KCNC3 | {'ess': 0, 'noness': 7} | True | 50320245 | CGTCCACTAGGGGGATATCCAC | [] |
| 5893 | 19 | FAM71E1 | {'ess': 1, 'noness': 7} | True | 50466657 | CCGCCTCTAGGCCTTGGCAGC | [] |
| 5894 | 19 | CLEC11A | {'ess': 0, 'noness': 7} | True | 50725466 | GCCCCGCTAGAAGGGGAACTC | [] |
| 5895 | 19 | KLK3 | {'ess': 1, 'noness': 6} | True | 50859615 | TGACTAGACACCTCCTCTCCAC | [] |
| 5896 | 19 | KLK11 | {'ess': 0, 'noness': 7} | True | 51022544 | CCAGTCTAATTGTTCTTCATCG | [] |
| 5897 | 19 | CTU1 | {'ess': 3, 'noness': 5} | False | 51098600 | AAGTCGCTAGAAGGTGGGGACC | [] |
| 5898 | 19 | IGLON5 | {'ess': 0, 'noness': 7} | True | 51328758 | GCGCCTACATTCTCCACCACAC | [] |
| 5899 | 19 | VSIG10L | {'ess': 1, 'noness': 7} | True | 51332610 | CCTCCGCCTACAGAGACAACTC | [] |
| 5900 | 19 | CLDND2 | {'ess': 0, 'noness': 7} | True | 51367004 | CCACGTTCCCTATGAGTTCATG | [] |
| 5901 | 19 | C19orf84 | unavailable | False | 51388983 | GTCCTAGTACTCTGTCTCCCAC | [] |
| 5902 | 19 | SIGLEC6 | {'ess': 0, 'noness': 7} | True | 51529579 | TGGACTCTAAGCCTCCCAGAGC | [] |
| 5903 | 19 | ZNF577 | {'ess': 0, 'noness': 7} | True | 51855923 | TCATGGCTACTGTGGTTCTGAG | [] |
| 5904 | 19 | ZNF350 | unavailable | False | 51964853 | TTTTCTTCCTATGGGTTTTCTG | [] |
| 5905 | 19 | ZNF615 | {'ess': 0, 'noness': 7} | True | 51992879 | TGACTACCTGTGAATTCTCCTG | [] |
| 5906 | 19 | ZNF841 | {'ess': 0, 'noness': 7} | True | 52059157 | TGGCTAACATGGTGAAACCCTG | [] |
| 5907 | 19 | ZNF616 | {'ess': 0, 'noness': 7} | True | 52114817 | CGTCTAAGGCCTTGTCACATTG | [] |
| 5908 | 19 | ZNF480 | {'ess': 0, 'noness': 7} | True | 52322857 | AGTTTCTATCCCATATGAATTG | [] |
| 5909 | 19 | ZNF578 | {'ess': 0, 'noness': 6} | True | 52512153 | GAAGTCTATGATGACATGCAAG | [] |
| 5910 | 19 | ZNF701 | {'ess': 1, 'noness': 7} | True | 52583456 | CACTTCTAAGATTTCTTTCCAG | [] |
| 5911 | 19 | ZNF611 | {'ess': 0, 'noness': 7} | True | 52704936 | CATTTCTAAGGTTTCTCTCCAG | [] |
| 5912 | 19 | ZNF347 | {'ess': 1, 'noness': 7} | True | 53140307 | CTCCACTATGAATTCTGTGATG | [] |
| 5913 | 19 | ZNF665 | {'ess': 0, 'noness': 7} | True | 53164452 | TGTAAGGTTTCTATCCACTATG | [] |
| 5914 | 19 | ZNF677 | {'ess': 0, 'noness': 5} | True | 53236971 | CCCTGATGCTAGGTACAGCTTC | [] |
| 5915 | 19 | ZNF845 | {'ess': 0, 'noness': 5} | True | 53353587 | TATTTCTAATGTTTCTTTCCAG | [] |
| 5916 | 19 | ZNF761 | unavailable | False | 53449952 | CTCAGCTACTTGGGAGACTGAC | [] |
| 5917 | 19 | PRKCG | {'ess': 0, 'noness': 6} | True | 53907035 | GGGCGTCTAGAACCCGCGGGG | [] |
| 5918 | 19 | CACNG8 | {'ess': 1, 'noness': 7} | True | 53982848 | CCGCGCCCCTACACAGGCGTG | [] |
| 5919 | 19 | CACNG6 | {'ess': 0, 'noness': 7} | True | 54011809 | ACACCTACTATATAGCAGGCAG | [] |
| 5920 | 19 | CACNG6 | {'ess': 0, 'noness': 7} | True | 54012188 | CGTGGCTCTAGGTGGCCCGGT | [] |
| 5921 | 19 | VSTM1 | {'ess': 0, 'noness': 7} | True | 54040960 | TCTTGCTACACTTTCAGTGCCG | [] |
| 5922 | 19 | OSCAR | {'ess': 0, 'noness': 7} | True | 54095677 | GAGGACTAGACTCCTGGATCTC | [] |
| 5923 | 19 | CNOT3 | {'ess': 5, 'noness': 3} | False | 54154050 | TGAGACCTACAGGAGGCCTAGC | [] |
| 5924 | 19 | MBOAT7 | {'ess': 1, 'noness': 7} | True | 54178221 | TCTGCTTCTAAAGGCTGCACAG | [] |
| 5925 | 19 | LILRB3 | {'ess': 0, 'noness': 7} | True | 54217092 | CCGGGCTAGTGGATGGCCAGA | [] |
| 5926 | 19 | LILRB5 | {'ess': 1, 'noness': 7} | True | 54250785 | CGTGGGCTAGTGGATGGCCAG | [] |
| 5927 | 19 | LILRB2 | {'ess': 0, 'noness': 7} | True | 54274692 | TCCGGGCTAGTGGATGGCCAG | [] |
| 5928 | 19 | LENG8 | {'ess': 7, 'noness': 1} | False | 54458905 | GGTCTCTATGGTGGGAGCACAC | [] |
| 5929 | 19 | CDC42EP5 | {'ess': 0, 'noness': 7} | True | 54465100 | GGGAACCTAGAGGCCGATGACC | [] |
| 5930 | 19 | LILRB1 | {'ess': 0, 'noness': 7} | True | 54636877 | CTGGGCTAGTGGATGGCCAGA | [] |
| 5931 | 19 | KIR2DL1 | {'ess': 0, 'noness': 7} | True | 54776338 | GAGCTAAAAGGTGGATCTCATG | [] |
| 5932 | 19 | KIR3DL2 | {'ess': 0, 'noness': 7} | True | 54866730 | CTCCCTAGAAAACCCCCTCAAC | [] |
| 5933 | 19 | PTPRH | {'ess': 3, 'noness': 5} | False | 55194242 | GCCTATGGCCCATCTTCATCAC | [] |
| 5934 | 19 | PPP6R1 | {'ess': 0, 'noness': 7} | True | 55230527 | AGGCAGCTATCTGGAAACAGAG | [] |
| 5935 | 19 | TMEM150B | {'ess': 0, 'noness': 7} | True | 55310224 | TGCAGGCTAACAGGGTGCCTG | [] |
| 5936 | 19 | TMEM150B | {'ess': 0, 'noness': 7} | True | 55312858 | GTGCCTGCTACAGCTGGACCG | [] |
| 5937 | 19 | TMEM190 | {'ess': 0, 'noness': 7} | True | 55378202 | CCCCTAATCCTCCTCTTCCTCC | [] |
| 5938 | 19 | UBE2S | {'ess': 4, 'noness': 3} | False | 55401435 | GCCCACTACAGCCGCCGCAGC | [] |
| 5939 | 19 | UBE2S | {'ess': 4, 'noness': 3} | False | 55404000 | CCTACCCTATCCTTGTTTTTTG | ['TTTTTTAAAAAAAAGGGAGAAG'] |
| 5940 | 19 | SSC5D | unavailable | False | 55504185 | GCTATGGGCTAGGGAGGCGCT | [] |
| 5941 | 19 | ZNF579 | {'ess': 2, 'noness': 6} | False | 55577326 | GATCTATCTTCTCACATTCCTG | [] |
| 5942 | 19 | ZNF784 | {'ess': 1, 'noness': 7} | True | 55621750 | TGGTCTGCAGCCTACTGGTCGC | [] |
| 5943 | 19 | LOC10798 | unavailable | False | 55640769 | TTCGCCCTAAGGCCCCACAAGG | [] |
| 5944 | 19 | LOC10798 | unavailable | False | 55640792 | ACCGAGACTAAGCAAAGTTACG | [] |
| 5945 | 19 | U2AF2 | {'ess': 7, 'noness': 1} | False | 55674067 | CCGCCTCTACCAGAAGTCCCG | [] |
| 5946 | 19 | NLRP13 | {'ess': 0, 'noness': 7} | True | 55892095 | CTCTCTCACTACATTGTGCAGG | [] |
| 5947 | 19 | NLRP8 | {'ess': 0, 'noness': 7} | True | 55987912 | GACGGCCTAAGGATTAATCTGG | [] |
| 5948 | 19 | ZNF787 | {'ess': 0, 'noness': 7} | True | 56088022 | CCCCTACCGGCCCTCCCCACC | [] |
| 5949 | 19 | ZNF444 | {'ess': 0, 'noness': 7} | True | 56160200 | AGGCGGCTAACCCAAGGGCCA | [] |
| 5950 | 19 | GALP | {'ess': 1, 'noness': 7} | True | 56182233 | CACAGGTCTAGGATCTCAAGGC | [] |
| 5951 | 19 | GALP | {'ess': 1, 'noness': 7} | True | 56185269 | TGAAGACATCTATGACTTCAGG | [] |
| 5952 | 19 | EDDM13 | unavailable | False | 56310147 | GGCTCACTAGAAAGAACAGCAG | [] |
| 5953 | 19 | ZNF582 | {'ess': 0, 'noness': 7} | True | 56383862 | ACAATTAGCCTAGGCTAATGGG | [] |
| 5954 | 19 | ZFP28 | {'ess': 0, 'noness': 7} | True | 56555391 | TCGAGGCTATGGTGATGGGAGC | [] |
| 5955 | 19 | ZNF470 | {'ess': 0, 'noness': 7} | True | 56578582 | TTGAATCTATAGGACTTGGTGG | [] |
| 5956 | 19 | ZIM3 | {'ess': 0, 'noness': 7} | True | 57134917 | CATGGGGCTCCTATCTGGAGTC | [] |
| 5957 | 19 | ZNF749 | {'ess': 0, 'noness': 7} | True | 57445484 | CTCCACTAAGGCCTTTTTCCAG | [] |
| 5958 | 19 | ZNF772 | {'ess': 0, 'noness': 7} | True | 57473273 | CCATCCTAAGGCCTTTCTCCAC | [] |
| 5959 | 19 | ZNF419 | {'ess': 1, 'noness': 7} | True | 57492908 | GCAGTCCTACAGAAGCAATCTG | [] |
| 5960 | 19 | ZNF549 | {'ess': 0, 'noness': 7} | True | 57538926 | AACAATTGCTAGGGCTCTTCTG | [] |
| 5961 | 19 | ZIK1 | {'ess': 0, 'noness': 7} | True | 57591274 | CATGAGGCCTCTATGTGTTATG | [] |
| 5962 | 19 | ZNF134 | unavailable | False | 57621402 | CACTCCTAAAGCCTGCCTGCAC | [] |
| 5963 | 19 | ZNF211 | {'ess': 0, 'noness': 7} | True | 57642180 | CAGCTAAGGCTTTTCTCCAATG | [] |
| 5964 | 19 | LOC10537 | unavailable | False | 58019954 | TCTGAAGTGCTAGCTGTTAAGG | [] |
| 5965 | 19 | ZNF274 | {'ess': 0, 'noness': 1} | False | 58213142 | TTGAGAGCTATGAGGTAGGCTG | [] |
| 5966 | 19 | ZNF544 | {'ess': 0, 'noness': 5} | True | 58259292 | CCAATTCTACCATTTCTTCAGG | [] |
| 5967 | 19 | ZNF544 | {'ess': 0, 'noness': 5} | True | 58262753 | CACTCCTAAGGTTTCTCTCCAG | [] |
| 5968 | 19 | ZNF8 | {'ess': 0, 'noness': 4} | True | 58295535 | TCTCTCCTATGTGGATTCTCTG | [] |
| 5969 | 19 | ZNF584 | {'ess': 1, 'noness': 7} | True | 58417783 | AACACGGTGCTAGCAGCTAACC | [] |
| 5970 | 19 | SLC27A5 | {'ess': 1, 'noness': 7} | True | 58500987 | TACCTAGAACTTTAATTTAATG | [] |
| 5971 | 19 | UBE2M | {'ess': 8, 'noness': 1} | False | 58556088 | GCCAACCCTATTTCAGGCAGCC | [] |
| 5972 | 19 | MZF1 | {'ess': 1, 'noness': 7} | True | 58562071 | CTGGAGCTACTCGGCGCTGTG | [] |
| 5973 | 20 | C20orf96 | {'ess': 0, 'noness': 7} | True | 271206 | CTGCCATCTAGAAGGGTAGTGC | [] |
| 5974 | 20 | TRIB3 | {'ess': 1, 'noness': 6} | True | 396589 | TAGGGTGGTCCTAGCCATACAC | [] |
| 5975 | 20 | RBCK1 | {'ess': 1, 'noness': 6} | True | 420075 | GGTCATGGCTAGGGTGCTGGG | [] |
| 5976 | 20 | SRXN1 | {'ess': 1, 'noness': 6} | True | 648713 | GGCTGCTACTGCAAGTCTGGTC | [] |
| 5977 | 20 | SLC52A3 | unavailable | False | 761025 | GGCCTGCCTAGGCTGGACAGT | [] |
| 5978 | 20 | FAM110A | {'ess': 0, 'noness': 7} | True | 845691 | GGCGCCTAGCCTTCAGCTGCT | [] |
| 5979 | 20 | RSPO4 | {'ess': 1, 'noness': 7} | True | 963728 | TTTCCTAATGTATTTGGGACTG | [] |
| 5980 | 20 | SIRPB1 | {'ess': 0, 'noness': 7} | True | 1568298 | TCACTAGGCTCCCATGCCACTC | [] |
| 5981 | 20 | SIRPG | {'ess': 0, 'noness': 7} | True | 1635976 | AACTCTCTAGCGCAGAGCTTGC | [] |
| 5982 | 20 | SNRPB | {'ess': 5, 'noness': 3} | False | 2461755 | ATGAGTCTAGGGCCTTGGTGGC | [] |
| 5983 | 20 | NOP56 | {'ess': 7, 'noness': 2} | False | 2658293 | GCATTCTAATCTTCCTGGGATG | [] |
| 5984 | 20 | IDH3B | {'ess': 0, 'noness': 7} | True | 2658441 | CAGGGCCTATGGGTGGGGCAA | [] |
| 5985 | 20 | IDH3B | {'ess': 0, 'noness': 7} | True | 2658750 | GGCTCTAGCTCCCTTTAGTCTC | [] |
| 5986 | 20 | TMEM239 | unavailable | False | 2817012 | CCTTCTATTGATCCAAATCCTG | [] |
| 5987 | 20 | PCED1A | unavailable | False | 2835461 | GATCCAGTCTACCCAGGCCATC | [] |
| 5988 | 20 | MRPS26 | {'ess': 3, 'noness': 5} | False | 3047868 | TGGGCCCCTAGGAGTCCCTGC | [] |
| 5989 | 20 | ITPA | {'ess': 1, 'noness': 6} | True | 3226687 | TCTAAAACTGAAAATATATCTG | [] |
| 5990 | 20 | C20orf194 | {'ess': 0, 'noness': 7} | True | 3252571 | CCTCTAGGGCTTCAGCTCAAAC | [] |
| 5991 | 20 | ADAM33 | {'ess': 0, 'noness': 7} | True | 3671584 | ACCATTCTAGGTCCACACTGGC | [] |
| 5992 | 20 | HSPA12B | {'ess': 0, 'noness': 7} | True | 3751392 | CCTGTACCTACCTGGCGACTAC | [] |
| 5993 | 20 | MAVS | {'ess': 1, 'noness': 6} | True | 3866146 | GGCTTCACTAGTGCAGACGCC | [] |
| 5994 | 20 | PANK2 | {'ess': 0, 'noness': 7} | True | 3908292 | ATAAGCTAAAAAATGCATACTG | [] |
| 5995 | 20 | RNF24 | {'ess': 0, 'noness': 6} | True | 3934062 | GTAAGCTATACAATGTTCTCTG | [] |
| 5996 | 20 | SLC23A2 | {'ess': 0, 'noness': 7} | True | 4856971 | CAAAGGCTATCCCGTGGCCTGC | [] |
| 5997 | 20 | TMEM230 | unavailable | False | 5100790 | GGGTGCTAGTCATCAAAGTCTG | [] |
| 5998 | 20 | PCNA | {'ess': 8, 'noness': 0} | False | 5115282 | CTAAGATCCTTCTTCATCCTCG | [] |
| 5999 | 20 | CDS2 | {'ess': 3, 'noness': 5} | False | 5190233 | TGGCCCCTACTCGTCCTCTGT | [] |
| 6000 | 20 | GPCPD1 | {'ess': 0, 'noness': 7} | True | 5547660 | TAAAAACTAAGCATTCTCCACG | [] |
| 6001 | 20 | CRLS1 | {'ess': 6, 'noness': 2} | False | 6023386 | AGCATTCCTAGGCACCTGGTAC | [] |
| 6002 | 20 | LRRN4 | {'ess': 0, 'noness': 7} | True | 6041021 | GCTGGGCTAACTGACGGTCTGC | [] |
| 6003 | 20 | BMP2 | {'ess': 0, 'noness': 7} | True | 6779088 | GCTGTACTAGCGACACCCACAC | ['CCCACAACCCTCCACAACCATG'] |
| 6004 | 20 | TMX4 | {'ess': 0, 'noness': 7} | True | 7982250 | TAAATCTACAGTCCCTTGTCAG | [] |
| 6005 | 20 | LAMP5 | unavailable | False | 9529819 | CTAACGGCCTCTAGCCCATGTC | [] |
| 6006 | 20 | JAG1 | {'ess': 2, 'noness': 4} | False | 10639497 | GCTATACGATGTACTCCATTCG | [] |
| 6007 | 20 | SPTLC3 | {'ess': 0, 'noness': 7} | True | 13080046 | CCTTGTTTTCTATGTACAGAAG | [] |
| 6008 | 20 | NDUFAF5 | unavailable | False | 13801843 | CAAACTAAACATTGTCAATCAG | [] |
| 6009 | 20 | SEL1L2 | {'ess': 0, 'noness': 7} | True | 13849484 | GATCCGCATCCTACCCATGGTC | [] |
| 6010 | 20 | KIF16B | {'ess': 0, 'noness': 7} | True | 16273252 | CTGGCTCTACCCCGTCCCGTG | [] |
| 6011 | 20 | PCSK2 | {'ess': 1, 'noness': 7} | True | 17482069 | GCAGCGCTAGTTCTTGTTAAGG | [] |
| 6012 | 20 | DSTN | {'ess': 0, 'noness': 7} | True | 17607145 | CTACACAGGGCATCCTTCAAAG | [] |
| 6013 | 20 | BANF2 | {'ess': 0, 'noness': 6} | True | 17735810 | GTTTGTGTCTACAGGAAGCAGG | [] |
| 6014 | 20 | MGME1 | unavailable | False | 17990108 | GCTCCCTATTCTGAATATTCTG | [] |
| 6015 | 20 | DZANK1 | unavailable | False | 18384398 | TCAGGGCTAACAGTCATCCAGC | [] |
| 6016 | 20 | RBBP9 | {'ess': 0, 'noness': 7} | True | 18489763 | CAGTCTATGCTGGTACTTTCAG | [] |
| 6017 | 20 | DTD1 | {'ess': 0, 'noness': 7} | True | 18630544 | TCTCTACTAAAAAAATACAAAG | ['ACAAAAAAATTAGCCGGGCATG'] |
| 6018 | 20 | DTD1 | {'ess': 0, 'noness': 7} | True | 18744251 | CTGAGCTACGGCTCCCGTTCA | [] |
| 6019 | 20 | RIN2 | {'ess': 0, 'noness': 7} | True | 20000935 | TGTCTTCTAGGAGGTGGTGAGC | [] |
| 6020 | 20 | CFAP61 | unavailable | False | 20171807 | GATGCTATGAACTCCTTCCCAG | [] |
| 6021 | 20 | CFAP61 | unavailable | False | 20322751 | GACCCTCTAATGATCTCTTCTG | [] |
| 6022 | 20 | CFAP61 | unavailable | False | 20360409 | CCTACAACTAAACGATGCCTGG | [] |
| 6023 | 20 | INSM1 | {'ess': 0, 'noness': 7} | True | 20369799 | GCGCGCTCTAGCAGGCCGGGC | [] |
| 6024 | 20 | RALGAPA2 | {'ess': 0, 'noness': 7} | True | 20393098 | AGTCCACTAATGGGAAGACCTG | [] |
| 6025 | 20 | XRN2 | {'ess': 6, 'noness': 2} | False | 21339042 | AAACTGTCCTAAAAGTATAAAG | ['AGTATAAAATCCACATATTCTG'] |
| 6026 | 20 | XRN2 | {'ess': 6, 'noness': 2} | False | 21345881 | ACTCTCTCTATCCTTGCTTGGG | [] |
| 6027 | 20 | PAX1 | {'ess': 0, 'noness': 7} | True | 21714561 | TGCCCCTAGGAGGTCGAGGCC | [] |
| 6028 | 20 | PAX1 | {'ess': 0, 'noness': 7} | True | 21714802 | GCAGTCTAAAACCCAAGTTCAG | [] |
| 6029 | 20 | NXT1 | {'ess': 1, 'noness': 7} | True | 23354463 | CCCACTAGCTGGCCCAGTCCT | [] |
| 6030 | 20 | CST11 | {'ess': 0, 'noness': 7} | True | 23450505 | GGACACCTAGTCACTGCTGCAC | [] |
| 6031 | 20 | CST3 | {'ess': 0, 'noness': 7} | True | 23633915 | AGACCCCTAGGCGTCCTGACA | [] |
| 6032 | 20 | CST4 | {'ess': 0, 'noness': 7} | True | 23685893 | CCCCTAGGCTTCTTGACACCTC | [] |
| 6033 | 20 | CST1 | {'ess': 0, 'noness': 7} | True | 23747915 | TCCCTAGGATTCTTGACACCTG | [] |
| 6034 | 20 | CST2 | {'ess': 1, 'noness': 7} | True | 23824019 | TCCCTAGGCTTCTTGACACCTC | [] |
| 6035 | 20 | CST5 | {'ess': 0, 'noness': 7} | True | 23876187 | CCCCTAGACTTTCCGGCACTTC | [] |
| 6036 | 20 | SYNDIG1 | unavailable | False | 24660032 | TATTATCTACAGCACAAAACAG | [] |
| 6037 | 20 | APMAP | unavailable | False | 24963812 | GGAGGGCTAAACAGCCTGGAG | [] |
| 6038 | 20 | VSX1 | {'ess': 0, 'noness': 7} | True | 25078735 | GTTGGCTATAGAGAAGGGACTG | [] |
| 6039 | 20 | ENTPD6 | {'ess': 0, 'noness': 7} | True | 25225596 | GGCTCGGCCACTATGAGGCTG | [] |
| 6040 | 20 | PYGB | {'ess': 0, 'noness': 7} | True | 25296521 | GTGTGCCTAGTCCCGGGGGAT | [] |
| 6041 | 20 | ABHD12 | {'ess': 0, 'noness': 7} | True | 25319019 | GGCTCTGCCTATAAAATGTTTG | [] |
| 6042 | 20 | DEFB115 | {'ess': 0, 'noness': 7} | True | 31259631 | AACTAGATATATAGCTCACTTG | [] |
| 6043 | 20 | DEFB119 | {'ess': 0, 'noness': 7} | True | 31389024 | TTCTATCTATTAGTTATATTTG | [] |
| 6044 | 20 | DEFB124 | {'ess': 0, 'noness': 7} | True | 31465505 | GAACCAGCTACTCATATTCATG | [] |
| 6045 | 20 | ID1 | {'ess': 2, 'noness': 6} | False | 31605836 | GATGATCTAGTGGTCGGATCTG | [] |
| 6046 | 20 | DUSP15 | {'ess': 0, 'noness': 7} | True | 31848586 | TGTGGTCTACTGCCTTCTCCTC | [] |
| 6047 | 20 | XKR7 | {'ess': 0, 'noness': 4} | True | 31997456 | GACACTGTAGCCTACACTGTGC | [] |
| 6048 | 20 | CCM2L | unavailable | False | 32031313 | GGGCGGTGGCTACAGGTAGTT | [] |
| 6049 | 20 | PLAGL2 | {'ess': 0, 'noness': 7} | True | 32196451 | GTGGGGGCTACTGGAATGCTTC | [] |
| 6050 | 20 | C20orf203 | unavailable | False | 32650431 | TCGGCTAATTAAACAGCGACCG | [] |
| 6051 | 20 | DNMT3B | {'ess': 1, 'noness': 6} | True | 32807902 | CTGGAACTATTCACATGCAAAG | [] |
| 6052 | 20 | EFCAB8 | {'ess': 0, 'noness': 1} | False | 32946521 | AGCTTCTAAAAGATAACACTAG | [] |
| 6053 | 20 | LOC10537 | unavailable | False | 33145950 | CTCAGCTATGGTGGCAATGGTC | [] |
| 6054 | 20 | SNTA1 | {'ess': 0, 'noness': 7} | True | 33408506 | CGACTTCTAGGCCAACAGCCC | [] |
| 6055 | 20 | SNTA1 | {'ess': 0, 'noness': 7} | True | 33408565 | AAGACTATGGTTTTGGGACACG | [] |
| 6056 | 20 | CBFA2T2 | {'ess': 0, 'noness': 7} | True | 33638960 | TAAGACTAGGTGCACAAACAAG | [] |
| 6057 | 20 | NECAB3 | {'ess': 1, 'noness': 7} | True | 33660003 | ATCTGACCTAGAGGAAGTGAGG | [] |
| 6058 | 20 | LOC10798 | unavailable | False | 34363270 | CTCGCTCTAGTAGCTGTGGTCC | [] |
| 6059 | 20 | PIGU | {'ess': 3, 'noness': 5} | False | 34560865 | GCCAGGCCTACTTGAGCACGA | [] |
| 6060 | 20 | GGT7 | {'ess': 1, 'noness': 7} | True | 34845327 | CCGCTGCTCTACAGGATGGTG | [] |
| 6061 | 20 | GGT7 | {'ess': 1, 'noness': 7} | True | 34845428 | AGACATCTACTTTCTCCACGTG | [] |
| 6062 | 20 | FAM83C | {'ess': 0, 'noness': 7} | True | 35286534 | CTTTGGCTAGGACTCAAAGCGC | [] |

| | | | | | | | |
|---|---|---|---|---|---|---|---|
| 6063 | 20 | GDF5 | {'ess': 0, 'nonext': 7} | True | 35433906 | TGCTGCTACCTGCAGCCACAC | [] |
| 6064 | 20 | CEP250 | {'ess': 1, 'nonext': 6} | True | 35511625 | GCTGCTACCTGGAGGCGGCTT | [] |
| 6065 | 20 | CEP250 | {'ess': 1, 'nonext': 6} | True | 35514028 | TCCTGCTACTGGGCAAGGAAA | [] |
| 6066 | 20 | C20orf173 | {'ess': 0, 'nonext': 5} | True | 35527220 | CTATTCTGATCCAAATCCAAAG | [] |
| 6067 | 20 | C20orf173 | {'ess': 0, 'nonext': 5} | True | 35528257 | TTCCACTCCACTAGGGAACCA | [] |
| 6068 | 20 | ERGIC3 | {'ess': 1, 'nonext': 7} | True | 35557503 | GTGACTACGTTGTCTTCCCTAG | [] |
| 6069 | 20 | CPNE1 | {'ess': 1, 'nonext': 7} | True | 35626240 | GAACCTAGGCCTGGGGGGCCT | [] |
| 6070 | 20 | RBM12 | {'ess': 4, 'nonext': 4} | False | 35652523 | TGAATGGCTACCCTAATACAAG | [] |
| 6071 | 20 | NFS1 | {'ess': 7, 'nonext': 1} | False | 35669621 | CTTCTAGTGTTGGGTCCACTTG | [] |
| 6072 | 20 | CNBD2 | unavailable | False | 36030647 | TACACTAAGCCAAGAGTTCTCG | [] |
| 6073 | 20 | MYL9 | {'ess': 0, 'nonext': 7} | True | 36549248 | GGCCTAGTCGTCTTTATCCTTG | [] |
| 6074 | 20 | TGIF2 | {'ess': 2, 'nonext': 6} | False | 36591430 | GATGCCTACTGGGGATTTTCAG | [] |
| 6075 | 20 | RAB5IF | unavailable | False | 36612008 | AATGACCTAGTGAGGAGACAAG | [] |
| 6076 | 20 | SLA2 | {'ess': 0, 'nonext': 7} | True | 36613865 | TGGGCCTAGGCATCATCCAAA | [] |
| 6077 | 20 | MROH8 | unavailable | False | 37102686 | GTGTTTCTAGGCAGAGGACCT | [] |
| 6078 | 20 | RPN2 | {'ess': 7, 'nonext': 2} | False | 37241314 | CTGGAACTAATGTGCTGTTCTG | [] |
| 6079 | 20 | MANBAL | {'ess': 0, 'nonext': 7} | True | 37316414 | CTCTTCTACCGCTTCTTCTTAG | [] |
| 6080 | 20 | SRC | {'ess': 0, 'nonext': 5} | True | 37403378 | CTGTGCCTAGAGGTTCTCCCC | [] |
| 6081 | 20 | CTNNBL1 | {'ess': 6, 'nonext': 2} | False | 37872012 | GGTGCCTCTAGAAGTTCTCCA | [] |
| 6082 | 20 | VSTM2L | {'ess': 0, 'nonext': 7} | True | 37944252 | GCATCAGTCTAGAGGCTGCAG | [] |
| 6083 | 20 | TTI1 | {'ess': 7, 'nonext': 1} | False | 37999197 | AAGGCTCTAAGCACTGCCAGG | [] |
| 6084 | 20 | RPRD1B | {'ess': 4, 'nonext': 4} | False | 38089874 | CATCCTAGTCAGTTGAAAACAG | [] |
| 6085 | 20 | KIAA1755 | {'ess': 0, 'nonext': 7} | True | 38213041 | TCTCAGCTAGAGGGAGGCAAG | [] |
| 6086 | 20 | RALGAPB | {'ess': 1, 'nonext': 6} | True | 38574966 | GGTCTAAGAACTGCAGTTCTTG | [] |
| 6087 | 20 | SLC32A1 | {'ess': 0, 'nonext': 7} | True | 38728638 | TGCGCCCTAGTCCTCCGCGTT | [] |
| 6088 | 20 | ACTR5 | {'ess': 3, 'nonext': 5} | False | 38771815 | CTCTGCTATGCCTGCTCACCA | [] |
| 6089 | 20 | PPP1R16B | {'ess': 0, 'nonext': 7} | True | 38918665 | AGACTAGGAGATACGGCAACAG | [] |
| 6090 | 20 | TOP1 | {'ess': 7, 'nonext': 1} | False | 41123296 | TGAGACTGGCTAAAACTCATAG | [] |
| 6091 | 20 | PLCG1 | {'ess': 1, 'nonext': 7} | True | 41174508 | GGGGTACAACTAGAGGCGGTT | [] |
| 6092 | 20 | LPIN3 | {'ess': 1, 'nonext': 7} | True | 41357635 | TGATCCTAGAACACAGCCCCA | [] |
| 6093 | 20 | LPIN3 | {'ess': 1, 'nonext': 7} | True | 41357976 | GGGGAGGCTACAGCCCCCCTC | [] |
| 6094 | 20 | EMILIN3 | {'ess': 0, 'nonext': 7} | True | 41361267 | CCTGTCTAGTTGGCTTGCCCT | [] |
| 6095 | 20 | CHD6 | {'ess': 0, 'nonext': 7} | True | 41404592 | AAAGTTCTAATTGGTGTCGTTG | [] |
| 6096 | 20 | PTPRT | {'ess': 0, 'nonext': 7} | True | 42080878 | ATTGAGCTAAAGGAGCTTAAG | ['TAAATATTCCAGTGCCACCTCG'] |
| 6097 | 20 | SGK2 | {'ess': 1, 'nonext': 6} | True | 43585015 | TCTCTTCTAGCAATCCAAGATG | [] |
| 6098 | 20 | GDAP1L1 | {'ess': 0, 'nonext': 7} | True | 44279299 | GCCCTAGATGTATTTTTTCTTG | [] |
| 6099 | 20 | HNF4A | {'ess': 2, 'nonext': 7} | False | 44429664 | CGGCTTGCTAGATAACTTCCTG | [] |
| 6100 | 20 | TTPAL | {'ess': 1, 'nonext': 7} | True | 44480690 | TGGTTCTAGGCCCTAGTAAAAG | [] |
| 6101 | 20 | TTPAL | {'ess': 1, 'nonext': 7} | True | 44489540 | CGGGCTAGTAGCAGGAGTACA | [] |
| 6102 | 20 | CCN5 | unavailable | False | 44727306 | GGCTCTAGAAGGCACTGTTTTG | [] |
| 6103 | 20 | RIMS4 | {'ess': 0, 'nonext': 6} | True | 44756133 | CCAGCACTAAGATCGTTCTCC | [] |
| 6104 | 20 | SEMG2 | unavailable | False | 45223380 | AACAGGGCTATGTAGATATTGG | [] |
| 6105 | 20 | RBPJL | {'ess': 1, 'nonext': 7} | True | 45316810 | GGCACTAGGGTAGAAGAGCCC | [] |
| 6106 | 20 | RBPJL | {'ess': 1, 'nonext': 7} | True | 45316958 | GCGCGCCTAAGTCTGGATGAA | [] |
| 6107 | 20 | SYS1 | {'ess': 6, 'nonext': 2} | False | 45367114 | TGATTCTAGACATTGGATTTAG | [] |
| 6108 | 20 | DBNDD2 | {'ess': 0, 'nonext': 7} | True | 45410139 | CACTGCTAGCTGCAGGCTCCA | [] |
| 6109 | 20 | WFDC9 | {'ess': 0, 'nonext': 7} | True | 45508109 | AGAATCTAGGGGTTTAGCATTG | [] |
| 6110 | 20 | WFDC3 | {'ess': 1, 'nonext': 7} | True | 45774427 | TCAGCACAGCTAGGGCACCGG | [] |
| 6111 | 20 | ACOT8 | {'ess': 0, 'nonext': 7} | True | 45841837 | TCTGGCTACAGCTTGCTCTCT | [] |
| 6112 | 20 | SPATA25 | unavailable | False | 45886516 | TCCATCTACAAGCAGGAGCCA | [] |
| 6113 | 20 | NEURL2 | {'ess': 0, 'nonext': 7} | True | 45888843 | TGTATCACTAGGCGGCACAGA | [] |
| 6114 | 20 | ZNF335 | {'ess': 7, 'nonext': 1} | False | 45952729 | AAGGTTCTAGGAGAAGATGGAG | [] |
| 6115 | 20 | MMP9 | {'ess': 0, 'nonext': 7} | True | 46016367 | AGCCCTAGTCCTCAGGGCACT | [] |
| 6116 | 20 | CD40 | {'ess': 0, 'nonext': 7} | True | 46126736 | ACCTTCCTAGGCTTTCTCCAG | [] |
| 6117 | 20 | CDH22 | {'ess': 0, 'nonext': 7} | True | 46174505 | GCGAGGGGCTAGGAGGCCTGG | [] |
| 6118 | 20 | TP53RK | {'ess': 7, 'nonext': 1} | False | 46686752 | ACATTCTTCTACCCAACCATGG | [] |
| 6119 | 20 | EYA2 | {'ess': 0, 'nonext': 7} | True | 47188132 | GATCCTGCTATAAATACTCCAG | [] |
| 6120 | 20 | ZMYND8 | {'ess': 4, 'nonext': 3} | False | 47210760 | TCACTGCTAGTCCCAGAAGGT | [] |
| 6121 | 20 | ARFGEF2 | {'ess': 0, 'nonext': 7} | True | 49033198 | AGGGCTACCACACTGGTGACA | [] |
| 6122 | 20 | DDX27 | {'ess': 6, 'nonext': 2} | False | 49243833 | CGACAGCTACTTCCTCCTCTT | [] |
| 6123 | 20 | ZNFX1 | {'ess': 0, 'nonext': 7} | True | 49247266 | ATCTTCCTACATCATCCCCTGG | [] |
| 6124 | 20 | SLC9A8 | {'ess': 0, 'nonext': 7} | True | 49885622 | TTTCTGCTAGTGCTGTGCTTCG | [] |
| 6125 | 20 | SLC9A8 | {'ess': 0, 'nonext': 7} | True | 49886649 | TCCCCAACCCTAACTCCACTT | [] |
| 6126 | 20 | SPATA2 | {'ess': 1, 'nonext': 7} | True | 49905618 | GGCCGGTCTATCTGTACACGA | [] |
| 6127 | 20 | UBE2V1 | {'ess': 0, 'nonext': 7} | True | 50096246 | GGAACTAGGATACAGGTGCAC | [] |
| 6128 | 20 | CEBPB | {'ess': 1, 'nonext': 7} | True | 50192070 | GCCGCGCTAGCAGTGGCCGGA | [] |
| 6129 | 20 | PTPN1 | {'ess': 3, 'nonext': 6} | False | 50582714 | GTCAGGCTATGTGTTGCTGTTG | [] |
| 6130 | 20 | BCAS4 | {'ess': 1, 'nonext': 7} | True | 50876534 | CCGTCCTATACAGTGTGGGCA | [] |
| 6131 | 20 | ATP9A | {'ess': 2, 'nonext': 7} | False | 51601210 | ACGGCCTATGATGTGAGCTTTG | [] |
| 6132 | 20 | ZFP64 | {'ess': 0, 'nonext': 7} | True | 52084556 | GCTGAGCTAGAGGTCTTGGCT | [] |
| 6133 | 20 | ZFP64 | {'ess': 0, 'nonext': 7} | True | 52152145 | TAAGCACTAATCTGGAATGGAG | [] |
| 6134 | 20 | TSHZ2 | {'ess': 0, 'nonext': 7} | True | 53258562 | GCAGAGCTATTCTTCATCCACG | [] |
| 6135 | 20 | CBLN4 | {'ess': 1, 'nonext': 7} | True | 55998556 | AATCCTATAGGGGGAACACCAG | [] |
| 6136 | 20 | MC3R | {'ess': 0, 'nonext': 7} | True | 56249814 | TGCATCCTATCCCAAGTTCATG | [] |
| 6137 | 20 | AURKA | {'ess': 7, 'nonext': 0} | False | 56370157 | GATTCCTAAGACTGTTTGCTAG | [] |
| 6138 | 20 | BMP7 | {'ess': 0, 'nonext': 7} | True | 57170958 | GAGGAGCTAGTGGCAGCCACA | [] |
| 6139 | 20 | RAE1 | {'ess': 7, 'nonext': 1} | False | 57378098 | CAGCCACTACTTCTTATTCCTG | [] |
| 6140 | 20 | CTCFL | {'ess': 0, 'nonext': 7} | True | 57507864 | TAACTAATACAACACCTAAAAG | [] |
| 6141 | 20 | ZBP1 | {'ess': 0, 'nonext': 7} | True | 57604572 | CTAAATCCCACCTCCCCACCA | [] |
| 6142 | 20 | ZBP1 | {'ess': 0, 'nonext': 7} | True | 57613085 | CCTACCCTTTGCCCCCACCCA | [] |
| 6143 | 20 | PMEPA1 | {'ess': 0, 'nonext': 7} | True | 57652052 | CCTGGGGACCCTAGAGAGGGT | [] |
| 6144 | 20 | ANKRD60 | {'ess': 1, 'nonext': 1} | False | 58218494 | AGGAGCTAATGAGGGGTCATC | [] |
| 6145 | 20 | VAPB | {'ess': 0, 'nonext': 7} | True | 58444164 | CTAAGGCCTTCTTCCTTCCCA | [] |
| 6146 | 20 | VAPB | {'ess': 0, 'nonext': 7} | True | 58444234 | CCTCTACAAGGCAATCTTCCC | ['CTTCCCAATAATTACACCAACG'] |
| 6147 | 20 | EDN3 | {'ess': 0, 'nonext': 7} | True | 59324658 | CCTCCTAAGGGGCTCCTTCCT | [] |
| 6148 | 20 | PHACTR3 | {'ess': 0, 'nonext': 7} | True | 59847129 | AGAAAATCTCTATGGCCTGTGG | [] |
| 6149 | 20 | MTG2 | unavailable | False | 62201076 | GGCGTGGCTACCACCTGAGCG | [] |
| 6150 | 20 | LAMA5 | {'ess': 0, 'nonext': 7} | True | 62309325 | TGTGTCCTAGGCGGCTGGGCA | [] |
| 6151 | 20 | LAMA5 | {'ess': 0, 'nonext': 7} | True | 62319537 | CCTGCCCTACTGTTCACTCCA | [] |
| 6152 | 20 | RPS21 | {'ess': 7, 'nonext': 1} | False | 62388000 | CAGATCTACTCAGGCACCATC | ['CACCATCACACACTCCAATGCG'] |
| 6153 | 20 | CABLES2 | {'ess': 0, 'nonext': 7} | True | 62390970 | CCTCTGCTAGAACTGCTGGGT | [] |
| 6154 | 20 | RBBP8NL | unavailable | False | 62410877 | GGCTGGCTAGGTCTCCTCCCA | [] |
| 6155 | 20 | GATA5 | {'ess': 0, 'nonext': 6} | True | 62464835 | GGGACCTAGGCCAAGGCCAGC | [] |
| 6156 | 20 | SLCO4A1 | {'ess': 0, 'nonext': 5} | True | 62672736 | TAATCATCTCCTAAAGGCCCTG | [] |
| 6157 | 20 | NTSR1 | {'ess': 2, 'nonext': 6} | False | 62760266 | CACAGCCTAGTACAGCGTCTC | [] |
| 6158 | 20 | MRGBP | unavailable | False | 62799642 | TGAGGGTCTACGTGCGGCGCC | [] |
| 6159 | 20 | DIDO1 | {'ess': 5, 'nonext': 4} | False | 62879232 | GGGCGTCTAGGCCTGCGAGGC | [] |
| 6160 | 20 | GID8 | unavailable | False | 62944911 | CAGGCGCTACTTGGGCTCCTC | ['CTCCTCAATCACACCCTTGCTG'] |
| 6161 | 20 | SLC17A9 | {'ess': 0, 'nonext': 7} | True | 62967499 | TGGGAGCTAGAGGTCCTCATG | [] |
| 6162 | 20 | BIRC7 | {'ess': 0, 'nonext': 7} | True | 63239604 | CCTGGCCTAGGACAGGAAGGT | [] |
| 6163 | 20 | ARFGAP1 | {'ess': 0, 'nonext': 7} | True | 63287624 | TCTGGTCTATGTTGCTGTTTTG | [] |
| 6164 | 20 | ARFGAP1 | {'ess': 0, 'nonext': 7} | True | 63287872 | TGGGCCCTACCAGTTCTGGTT | [] |
| 6165 | 20 | COL20A1 | {'ess': 0, 'nonext': 7} | True | 63331097 | AGAGCTACTGAGACGGCCACA | [] |
| 6166 | 20 | CHRNA4 | {'ess': 0, 'nonext': 7} | True | 63346727 | CTTCCTAGATCATGCCAGCCA | [] |
| 6167 | 20 | GMEB2 | {'ess': 0, 'nonext': 7} | True | 63590088 | TGTCGGCTACTTCCGCTCGTG | [] |
| 6168 | 20 | RTEL1 | {'ess': 6, 'nonext': 2} | False | 63695784 | GCACATCCTAGAGGCCTGTGA | [] |
| 6169 | 20 | ARFRP1 | {'ess': 2, 'nonext': 6} | False | 63700442 | GCGCCTACGTGATGTCCCTCT | [] |
| 6170 | 20 | ZGPAT | {'ess': 1, 'nonext': 7} | True | 63735918 | TTGTGGGGTCTCTAGAACTCAG | [] |
| 6171 | 20 | ZBTB46 | {'ess': 0, 'nonext': 5} | True | 63746929 | GCGGGCCTAGGAGAGCCAGGC | [] |
| 6172 | 20 | TPD52L2 | {'ess': 0, 'nonext': 7} | True | 63889416 | CTGCACTACAGGATTTTCCAAG | [] |
| 6173 | 20 | UCKL1 | {'ess': 0, 'nonext': 7} | True | 63939975 | CTGGGCAGCTAACCCGTGTAG | [] |
| 6174 | 20 | UCKL1 | {'ess': 0, 'nonext': 7} | True | 63941762 | AGCAGGTCTAGGAGCGGGCAG | [] |
| 6175 | 20 | SAMD10 | {'ess': 1, 'nonext': 7} | True | 63975509 | CAGCAGCTAGGACATTTTCCC | [] |
| 6176 | 20 | C20orf204 | unavailable | False | 64038758 | GCCGCGCTAGGAGTCCCTGGA | [] |
| 6177 | 20 | SOX18 | {'ess': 1, 'nonext': 7} | True | 64049165 | GCGGCCTAGCCGGAGATGCAC | [] |
| 6178 | 20 | RGS19 | {'ess': 0, 'nonext': 7} | True | 64073852 | GGCGGCCTAGGCCTCGGAGGA | [] |
| 6179 | 20 | OPRL1 | {'ess': 2, 'nonext': 6} | False | 64098885 | CCGCCTAGAGAGCAGTGACCT | [] |
| 6180 | 20 | MYT1 | {'ess': 1, 'nonext': 6} | True | 64240447 | CCACACGGCCTAGACCTGGAT | [] |
| 6181 | 21 | LOC10272 | unavailable | False | 5034095 | TCCACATCTATGTTGCTCCTGG | [] |
| 6182 | 21 | LOC10272 | unavailable | False | 5045605 | CAAAGCTTCTAGGAAATAGATG | [] |
| 6183 | 21 | LOC10272 | unavailable | False | 5131280 | ACCGGCTGGCTCTAGGCCAGC | ['GGCCAGCATCTCTTCTTCTGAG'] |
| 6184 | 21 | LOC11226 | unavailable | False | 8399220 | TGAAGCCTAGGGCGCGGGCCC | [] |
| 6185 | 21 | BAGE2 | unavailable | False | 10452885 | ACTGATCTAAGAGGTCTCCCG | [] |
| 6186 | 21 | LOC10537 | unavailable | False | 10499509 | TGCTACAAGTGAAGATCCACAG | [] |
| 6187 | 21 | RBM11 | {'ess': 0, 'nonext': 7} | True | 14227292 | CTAGTATCTTTTCTTCTTCTTG | ['TCTTAGACTTTCGGAACTTTTG'] |
| 6188 | 21 | SAMSN1 | {'ess': 2, 'nonext': 5} | False | 14496292 | TACCTAGATGTGTGCTATTATG | [] |
| 6189 | 21 | CXADR | {'ess': 1, 'nonext': 7} | True | 17565614 | GGGCAGCTACCTTAGCAGGTG | [] |
| 6190 | 21 | CXADR | {'ess': 1, 'nonext': 7} | True | 17565691 | ATATGGAGGCTCTATACTATAG | [] |
| 6191 | 21 | TMPRSS15 | {'ess': 0, 'nonext': 7} | True | 18269968 | TGCGCTAATGTAGAAAACTTTG | [] |
| 6192 | 21 | NCAM2 | {'ess': 0, 'nonext': 7} | True | 21480043 | TCTTCTCTACCAGACTGTGGAG | [] |
| 6193 | 21 | JAM2 | {'ess': 1, 'nonext': 6} | True | 25714304 | TGGTCTAGAACTCCTGACCTC | [] |
| 6194 | 21 | APP | {'ess': 1, 'nonext': 6} | True | 25881669 | CGGGGGTCTAGTTCTGCATCT | [] |
| 6195 | 21 | ADAMTS5 | {'ess': 0, 'nonext': 7} | True | 26924052 | AGGCTAACATTTCTTCAACAAG | [] |
| 6196 | 21 | N6AMT1 | {'ess': 6, 'nonext': 2} | False | 28876384 | ATGCTAAGACTTGGTGAACTTG | [] |
| 6197 | 21 | CLDN8 | {'ess': 0, 'nonext': 4} | True | 30215247 | CAACTACACATACTGACTTCTG | [] |
| 6198 | 21 | KRTAP23- | {'ess': 0, 'nonext': 7} | True | 30349408 | CTGGCAGCTAGGGGTTTCCCA | [] |
| 6199 | 21 | TIAM1 | {'ess': 0, 'nonext': 7} | True | 31124658 | GGAGTTCTACTGCGGAGTGGG | [] |
| 6200 | 21 | LOC15005 | unavailable | False | 31560115 | GAAGTCTAGAGCTCTTCAGGAG | [] |
| 6201 | 21 | SCAF4 | unavailable | False | 31671398 | AGTCTCTAACGAGGAGCCTCT | [] |
| 6202 | 21 | CFAP298 | unavailable | False | 32002223 | ACAGGCTATGTGTGGGCGCCA | [] |
| 6203 | 21 | PAXBP1 | unavailable | False | 32734949 | CTCATAGATCTATTTTCCTTCG | [] |
| 6204 | 21 | C21orf62 | {'ess': 0, 'nonext': 7} | True | 32792761 | ATTATGCTAGTAAATAAATGTG | [] |
| 6205 | 21 | IL10RB | {'ess': 0, 'nonext': 7} | True | 33290356 | GAGCCTAGCTTTGGGGCCCCT | [] |
| 6206 | 21 | IFNAR1 | {'ess': 0, 'nonext': 7} | True | 33348573 | ATCCACCTACCTCGGCCTCCT | ['GGCCTCCTAAAATACAAACTGG'] |
| 6207 | 21 | SON | {'ess': 7, 'nonext': 1} | False | 33557550 | TTTGGCTTCTAAAACTGGAAAG | [] |
| 6208 | 21 | SON | {'ess': 7, 'nonext': 1} | False | 33560130 | ATTTGCTACCACCAAAATCTAG | [] |
| 6209 | 21 | ITSN1 | {'ess': 0, 'nonext': 7} | True | 33888299 | CCTACGGCTCATCAAACAACTG | [] |
| 6210 | 21 | RCAN1 | {'ess': 0, 'nonext': 7} | True | 34521301 | GTCTCTCTAACACTGCAGGTG | [] |
| 6211 | 21 | CLIC6 | {'ess': 0, 'nonext': 7} | True | 34709846 | ATTCTCAATTGCTAACACCAGG | [] |
| 6212 | 21 | SIM2 | {'ess': 0, 'nonext': 7} | True | 36745272 | AGCCCTACTTAGAAGCAGAAAG | [] |
| 6213 | 21 | TTC3 | {'ess': 0, 'nonext': 7} | True | 37201573 | GTGTGACTACCTAGAAGAGCAG | [] |
| 6214 | 21 | VPS26C | unavailable | False | 37225543 | CTCCGGGCTATATCCTGCAGA | [] |
| 6215 | 21 | KCNJ6 | {'ess': 0, 'nonext': 5} | True | 37625158 | CAGCTAGGGCACTAAACTTTGG | [] |
| 6216 | 21 | ERG | {'ess': 2, 'nonext': 5} | False | 38400361 | TAAAGGGCTAATCTGCTCTTAG | ['GCTCTTAATTCCTTTAGTGTGG'] |
| 6217 | 21 | BRWD1 | {'ess': 0, 'nonext': 7} | True | 39187025 | CTATTGTGCATTTTCTTTCCAG | [] |
| 6218 | 21 | GET1-SH3 | unavailable | False | 39511774 | TGAAAAGGCCTAGGAATCTTCG | [] |
| 6219 | 21 | SH3BGR | {'ess': 0, 'nonext': 7} | True | 39511774 | TGAAAAGGCCTAGGAATCTTCG | [] |
| 6220 | 21 | IGSF5 | {'ess': 0, 'nonext': 7} | True | 39801356 | AGGCTTTGCTATACTACAGTTG | [] |
| 6221 | 21 | PCP4 | {'ess': 0, 'nonext': 7} | True | 39929110 | TCCCACTAGGACTGAGACCCA | [] |
| 6222 | 21 | MX1 | {'ess': 2, 'nonext': 6} | False | 41447291 | AGGTCACTAGGGTGGGCCCTC | ['CCCTCATCCCATCTGCCTAGTG'] |
| 6223 | 21 | MX1 | {'ess': 2, 'nonext': 6} | False | 41458780 | GCACGTCTACGGGGCTGGACA | [] |
| 6224 | 21 | RIPK4 | {'ess': 0, 'nonext': 7} | True | 41740837 | CAGCCAGCTAGGTCTTGCTTC | [] |
| 6225 | 21 | C2CD2 | {'ess': 0, 'nonext': 7} | True | 41889123 | GCCCTACGTGCAGGGCTCCAC | [] |
| 6226 | 21 | UMODL1 | {'ess': 0, 'nonext': 7} | True | 42137619 | CCTCCTATTCGTAGAACACCTG | [] |
| 6227 | 21 | TFF1 | {'ess': 1, 'nonext': 6} | True | 42362478 | GTCTAAAATTCACACTCCTCTG | ['CCTCTACAGGGGTGAGGGGGAG'] |
| 6228 | 21 | LOC10192 | unavailable | False | 42569119 | CACAGCTAATAAGGAGGACCTG | [] |
| 6229 | 21 | PKNOX1 | {'ess': 1, 'nonext': 7} | True | 43030100 | CCCCTACTGCAGGGAGTCACT | [] |
| 6230 | 21 | HSF2BP | {'ess': 0, 'nonext': 7} | True | 43529753 | ACACCTACACATTATGCTCCAG | [] |
| 6231 | 21 | AGPAT3 | {'ess': 0, 'nonext': 7} | True | 43960644 | TTGGGTTGCTTCTAGTTTTGTG | [] |
| 6232 | 21 | PWP2 | {'ess': 7, 'nonext': 1} | False | 44130770 | ACCGGCTGGCTCTAGGCCAGC | ['GGCCAGCATCTCTTCTTCTGAG'] |
| 6233 | 21 | ICOSLG | {'ess': 0, 'nonext': 7} | True | 44218056 | CAAAGCTTCTAGGAAATAGATG | [] |
| 6234 | 21 | ICOSLG | {'ess': 0, 'nonext': 7} | True | 44229529 | TCCACATCTATGTTGCTCCTGG | [] |
| 6235 | 21 | LRRC3 | {'ess': 0, 'nonext': 7} | True | 44457417 | CAGGCGCTAGGGCCCCCGGGC | [] |
| 6236 | 21 | SUMO3 | {'ess': 0, 'nonext': 7} | True | 44806950 | GCCCTCTAGAAACTGTGCCCT | [] |
| 6237 | 21 | PTTG1IP | {'ess': 1, 'nonext': 7} | True | 44851366 | ACTAAATCTAGAAACAGAGATG | [] |
| 6238 | 21 | ITGB2 | {'ess': 0, 'nonext': 7} | True | 44886367 | CTCCTAACTCTCAGCAAACTTG | [] |
| 6239 | 21 | COL18A1 | {'ess': 0, 'nonext': 4} | True | 45512397 | GGCGGTGGCTACTTGGAGGCA | [] |
| 6240 | 21 | PCBP3 | {'ess': 0, 'nonext': 7} | True | 45936353 | AGAAGGCCTAGAGTGCACATTG | [] |
| 6241 | 21 | COL6A1 | {'ess': 0, 'nonext': 7} | True | 46004012 | TGCAGGGTGGGCTAGCCCAGC | [] |
| 6242 | 21 | COL6A2 | {'ess': 0, 'nonext': 7} | True | 46132551 | GCGGCGCTAGCAGATCCAGCG | [] |
| 6243 | 21 | FTCD | {'ess': 1, 'nonext': 7} | True | 46136453 | CGAGGCTATCCTTCCAGCGTC | [] |
| 6244 | 21 | FTCD | {'ess': 1, 'nonext': 7} | True | 46136813 | ACATGGGACTAGGGGCCTTCT | [] |
| 6245 | 21 | FTCD-AS1 | unavailable | False | 46152550 | CCCCTAGGCTGCACCAACCCA | [] |
| 6246 | 21 | DIP2A | {'ess': 0, 'nonext': 7} | True | 46546005 | CACCCCCTACCTTCTCTGACT | [] |
| 6247 | 21 | DIP2A | {'ess': 0, 'nonext': 7} | True | 46560881 | GTCCTCTAATGCCTGGGTCCC | [] |
| 6248 | 21 | DIP2A | {'ess': 0, 'nonext': 7} | True | 46562993 | GGTGAGCTAAACAATTACAGTG | [] |
| 6249 | 21 | PRMT2 | {'ess': 1, 'nonext': 7} | True | 46653053 | TACACTAACAACAGTAAGACTG | [] |
| 6250 | 22 | CCT8L2 | {'ess': 1, 'nonext': 6} | True | 16590876 | ATCACTAGTTATTCAATCCAAG | [] |
| 6251 | 22 | CECR2 | {'ess': 0, 'nonext': 1} | False | 17552839 | CTCCTTGGACTAGCTCTGAAAG | [] |
| 6252 | 22 | SLC25A18 | {'ess': 0, 'nonext': 7} | True | 17590235 | TGTCTAGTCAAAACACTTTAAG | [] |
| 6253 | 22 | BCL2L13 | {'ess': 1, 'nonext': 7} | True | 17726589 | AAGACTAAACACAGTGCCCTAG | [] |
| 6254 | 22 | BCL2L13 | {'ess': 1, 'nonext': 7} | True | 17727533 | GCCTCCTATTTCTTTCTCAGAG | [] |
| 6255 | 22 | DGCR6 | {'ess': 0, 'nonext': 7} | True | 18911688 | CTGTGGCTATGGTGGGACAGG | [] |
| 6256 | 22 | PRODH | {'ess': 0, 'nonext': 7} | True | 18913174 | CGGGTGCTAGGCAGGGCGATG | [] |
| 6257 | 22 | DGCR2 | {'ess': 1, 'nonext': 6} | True | 19038864 | GGCCGTCTACACCACAGTATT | [] |
| 6258 | 22 | TSSK2 | {'ess': 2, 'nonext': 5} | False | 19132475 | CCATTGTCATGCTAGGTGCTTG | [] |
| 6259 | 22 | ESS2 | unavailable | False | 19134195 | TGGCTCTAAAGAAGTCCGAAG | [] |
| 6260 | 22 | CLTCL1 | {'ess': 1, 'nonext': 5} | False | 19201437 | ACATTGCTATAGAGCAGCTTGG | [] |
| 6261 | 22 | HIRA | {'ess': 4, 'nonext': 4} | False | 19321439 | GGCAGGCTACTTGTCCCTCAG | [] |
| 6262 | 22 | MRPL40 | {'ess': 4, 'nonext': 4} | False | 19435961 | TCTATCTCTTAAACTCCACTTG | [] |
| 6263 | 22 | CDC45 | {'ess': 9, 'nonext': 0} | False | 19518907 | AACTCACCCTAGGACAGGAGG | [] |
| 6264 | 22 | ARVCF | {'ess': 1, 'nonext': 6} | True | 19970716 | AAGCCCTAAGAACAAGAGGCTG | [] |
| 6265 | 22 | ARVCF | {'ess': 1, 'nonext': 6} | True | 19971227 | AAGCTAGACCCAGGAATCAACG | [] |
| 6266 | 22 | TANGO2 | unavailable | False | 20064563 | CGCATCTACCAGGATGATAGTG | [] |
| 6267 | 22 | TRMT2A | {'ess': 1, 'nonext': 6} | True | 20112562 | CTGGGCCTAGGATGAGGGGAA | [] |
| 6268 | 22 | ZDHHC8 | {'ess': 2, 'nonext': 6} | False | 20147230 | ATCCTGCTAGGGCCCCGAGAG | [] |
| 6269 | 22 | DGCR6L | {'ess': 1, 'nonext': 6} | True | 20314574 | CTGTGGCTATGGTGGGACAGG | [] |
| 6270 | 22 | SCARF2 | {'ess': 1, 'nonext': 6} | True | 20425374 | AGCCTGCTACAGGGTGGGTGC | [] |
| 6271 | 22 | KLHL22 | {'ess': 1, 'nonext': 6} | True | 20442072 | GGGAGCCTAGTCCTCACTGGA | [] |
| 6272 | 22 | MED15 | {'ess': 5, 'nonext': 3} | False | 20569390 | CTGACTCTTTCTAAAGCCAACG | [] |
| 6273 | 22 | MED15 | {'ess': 5, 'nonext': 3} | False | 20586703 | TCTTGGCTAGGCGGCTGAGAG | [] |
| 6274 | 22 | SERPIND1 | {'ess': 1, 'nonext': 6} | True | 20787065 | CCACCTCTAGGACCTGCTGGG | [] |
| 6275 | 22 | P2RX6 | {'ess': 1, 'nonext': 6} | True | 21026616 | ACGGCTACAGGCTCCCGGAAT | [] |
| 6276 | 22 | SLC7A4 | {'ess': 1, 'nonext': 6} | True | 21029054 | ATCAGCTACTCCATATGGCCAG | [] |
| 6277 | 22 | HIC2 | {'ess': 2, 'nonext': 6} | False | 21446742 | TGGCTTCTAGGAGGGGGAGGT | [] |
| 6278 | 22 | CCDC116 | {'ess': 1, 'nonext': 6} | True | 21635643 | AGTGGCTGCTAGGAGTGGTGG | [] |
| 6279 | 22 | CCDC116 | {'ess': 1, 'nonext': 6} | True | 21637069 | CTAGACTCCATCCTCATCCTTG | [] |
| 6280 | 22 | PPIL2 | {'ess': 7, 'nonext': 1} | False | 21695489 | CTGCTGCTACCAGGAGCTGAA | [] |
| 6281 | 22 | PPIL2 | {'ess': 7, 'nonext': 1} | False | 21698802 | CATACCTCTAGTGGTCATCAGG | [] |
| 6282 | 22 | PPM1F | {'ess': 3, 'nonext': 5} | False | 21923091 | ACCACCTAGCTTCTTGGTGGA | [] |
| 6283 | 22 | PRAME | {'ess': 1, 'nonext': 6} | True | 22549066 | CAGCTAATTAGGCATGAAACAG | [] |
| 6284 | 22 | IGLV3-10 | unavailable | False | 22812280 | CACTGTGCTATGATTACCACTG | [] |
| 6285 | 22 | IGLL5 | unavailable | False | 22895693 | GGAACCTATGAACATTCTGTAG | [] |
| 6286 | 22 | IGLJ1 | unavailable | False | 22893817 | CTTACCTAGGACGGTGACCTT | [] |
| 6287 | 22 | IGLC1 | {'ess': 0, 'nonext': 1} | False | 22895693 | GGAACCTATGAACATTCTGTAG | [] |
| 6288 | 22 | IGLJ2 | unavailable | False | 22899654 | CTCACCTAGGACGGTCAGCTT | [] |
| 6289 | 22 | IGLC2 | {'ess': 0, 'nonext': 1} | False | 22901294 | AGAACCTATGAACATTCTGTAG | [] |
| 6290 | 22 | IGLJ3 | unavailable | False | 22905024 | CTCACCTAGGACGGTCAGCTT | [] |
| 6291 | 22 | IGLC3 | {'ess': 0, 'nonext': 1} | False | 22906660 | AGAACCTATGAACATTCTGTAG | [] |
| 6292 | 22 | IGLJ4 | unavailable | False | 22910606 | ACTCATCTAAAATGATCAGCTG | [] |
| 6293 | 22 | IGLJ5 | unavailable | False | 22914308 | CTCATCTAGGACGGTCAGCTC | [] |
| 6294 | 22 | IGLC7 | {'ess': 0, 'nonext': 1} | False | 22922912 | GGGGCCTAAGAGCATTCTGCA | [] |
| 6295 | 22 | RAB36 | {'ess': 1, 'nonext': 6} | True | 23168912 | GGCTAGGGGTGCCAACCACCT | [] |
| 6296 | 22 | LOC10798 | unavailable | False | 23394791 | TCCGGCCTAGAAGGACAGGAA | [] |
| 6297 | 22 | IGLL1 | {'ess': 1, 'nonext': 7} | True | 23573265 | GGAACCTATGAACATTCTGCAG | [] |
| 6298 | 22 | DRICH1 | unavailable | False | 23583170 | AAGATCTACTTCTTGCTGGAAG | [] |
| 6299 | 22 | RGL4 | {'ess': 0, 'nonext': 7} | True | 23699112 | TATACACTAAAGAAAATAAACG | ['ATAAACATAAAATGGTCTCCCG'] |
| 6300 | 22 | ZNF70 | {'ess': 0, 'nonext': 7} | True | 23743799 | TCTTTCTATAGCTTCTCCCCAG | [] |
| 6301 | 22 | VPREB3 | {'ess': 0, 'nonext': 7} | True | 23752875 | CCACCCCTAGGGACTAAAGCC | [] |
| 6302 | 22 | C22orf15 | {'ess': 0, 'nonext': 7} | True | 23765753 | TCTCACTACAGTGTGCAATCCG | ['CAATCCATCCCCTTAATCAGGG'] |
| 6303 | 22 | SLC2A11 | {'ess': 0, 'nonext': 7} | True | 23884848 | GGGACTAGAGTTCTGTTGACTG | [] |
| 6304 | 22 | UPB1 | {'ess': 0, 'nonext': 7} | True | 24525793 | AGCCGGCTACTCTTTCACGAT | [] |
| 6305 | 22 | CRYBB2 | {'ess': 0, 'nonext': 7} | True | 25231771 | AGGGCACTAGTTGGAGGGGTG | [] |
| 6306 | 22 | GRK3 | unavailable | False | 25722449 | TGGGTGCTAGAGGCCGTTGCT | [] |
| 6307 | 22 | MYO18B | {'ess': 0, 'nonext': 7} | True | 26027677 | AACTGGTTCCTACTTCTGGAGG | [] |
| 6308 | 22 | HPS4 | {'ess': 0, 'nonext': 7} | True | 26444581 | CAGCTTCTAAGAGGTACAACTG | [] |
| 6309 | 22 | HPS4 | {'ess': 0, 'nonext': 7} | True | 26453884 | GTCTAGTCTGCTCTCCACACT | [] |
| 6310 | 22 | PITPNB | {'ess': 0, 'nonext': 7} | True | 27854891 | CTCATCTAGACATCAGCAGCC | [] |
| 6311 | 22 | TTC28 | {'ess': 0, 'nonext': 7} | True | 28020591 | AGGGCTCTATTCCTGAGGCAG | [] |
| 6312 | 22 | TTC28 | {'ess': 0, 'nonext': 7} | True | 28028886 | ACACTCTAGTTTATCACCACAG | [] |
| 6313 | 22 | CCDC117 | {'ess': 1, 'nonext': 7} | True | 28786325 | GAAATTGGTTTCTAGAGTTCCG | ['GAGTTCCATCTCTTCTTCTGTG'] |
| 6314 | 22 | C22orf31 | {'ess': 0, 'nonext': 7} | True | 29058741 | CTATTTTTTGCTCTTTAACTTG | [] |
| 6315 | 22 | RHBDD3 | {'ess': 1, 'nonext': 7} | True | 29260059 | GCCTGGGCTAGGGAGGCCCAG | [] |
| 6316 | 22 | EWSR1 | {'ess': 3, 'nonext': 5} | False | 29290502 | TCTAGTCCCACTTTTCATTATG | [] |
| 6317 | 22 | EWSR1 | {'ess': 3, 'nonext': 5} | False | 29300160 | CATCTAGTAGGGCCGATCTCT | [] |
| 6318 | 22 | NF2 | {'ess': 2, 'nonext': 6} | False | 29694801 | CACCTGCTAGAGCTCTTCAAAG | [] |
| 6319 | 22 | CABP7 | {'ess': 1, 'nonext': 7} | True | 29729568 | TGGCGTCTACTTCATGCCACT | [] |
| 6320 | 22 | UQCR10 | {'ess': 3, 'nonext': 6} | False | 29769718 | GAACTACTTGTTCTCATACTTG | [] |
| 6321 | 22 | HORMAD2 | {'ess': 0, 'nonext': 7} | True | 30124438 | GTTCTACATCTTAGCCTAAATG | [] |
| 6322 | 22 | LIF | {'ess': 1, 'nonext': 6} | True | 30243650 | ACCTCCTGCTAGAAGGCCTGG | [] |
| 6323 | 22 | OSM | {'ess': 1, 'nonext': 6} | True | 30263882 | CTCGAGGCTACCGGGGCAGCT | [] |
| 6324 | 22 | SF3A1 | {'ess': 8, 'nonext': 1} | False | 30334593 | TCTTGTCTACTTCTTCCTCCCG | [] |
| 6325 | 22 | CCDC157 | {'ess': 0, 'nonext': 5} | True | 30373882 | GTGAGCTACATGGTACTCAGC | [] |
| 6326 | 22 | CCDC157 | {'ess': 0, 'nonext': 5} | True | 30376458 | GCTGCTAGGAAGGTGCTGTCT | [] |
| 6327 | 22 | CCDC157 | {'ess': 0, 'nonext': 5} | True | 30376744 | CAGGCTACATGGGCCGCTCCT | [] |
| 6328 | 22 | RNF215 | {'ess': 0, 'nonext': 7} | True | 30379699 | GGGCAGCTAATCATCGGAGTA | [] |
| 6329 | 22 | SEC14L2 | {'ess': 0, 'nonext': 7} | True | 30416500 | CTTCTAGTTCTAACACTCAGAG | [] |
| 6330 | 22 | MTFP1 | {'ess': 0, 'nonext': 7} | True | 30428673 | TTGTCTCTAAGCTTCTGTCCAG | [] |
| 6331 | 22 | SEC14L3 | {'ess': 1, 'nonext': 7} | True | 30460020 | GAGCCACCTAGACAGGGGTGA | [] |
| 6332 | 22 | SEC14L6 | unavailable | False | 30524996 | AGGTTCACCTAGAATTTCTCCG | ['TTCTCCATCTTCTCCATGAAGG'] |
| 6333 | 22 | TCN2 | {'ess': 0, 'nonext': 7} | True | 30626520 | TCAGGGGCTACCAGCTAACCA | [] |
| 6334 | 22 | SMTN | {'ess': 0, 'nonext': 1} | False | 31104465 | GGCAGGCTAGACATTCTTGCC | [] |
| 6335 | 22 | SELENOM | unavailable | False | 31104969 | CCGGACCTACAGGTCAGCGTG | [] |
| 6336 | 22 | LIMK2 | {'ess': 1, 'nonext': 6} | True | 31278440 | AGGGCTAGGGAGGTGAGTCCC | [] |
| 6337 | 22 | PATZ1 | {'ess': 2, 'nonext': 7} | False | 31341404 | CAGGCTAGCCAACAGGCCACT | [] |
| 6338 | 22 | SFI1 | {'ess': 0, 'nonext': 7} | True | 31619417 | GAACACGCTAGCACAGGGCCT | [] |
| 6339 | 22 | PISD | {'ess': 3, 'nonext': 3} | False | 31619611 | GAGACTCTAGAGCGAGCCCAG | [] |
| 6340 | 22 | C22orf42 | {'ess': 0, 'nonext': 7} | True | 32149539 | TGGAACACTAAAGCCGGAGAAG | [] |
| 6341 | 22 | C22orf42 | {'ess': 0, 'nonext': 7} | True | 32150853 | TGTCAACTACACGTAAATAAAG | [] |
| 6342 | 22 | RTCB | unavailable | False | 32387991 | AGGTTCTATCCTTTGATCACAG | [] |
| 6343 | 22 | LARGE1 | unavailable | False | 33274426 | TTGGTGCTAGCTGTTGTTCTCG | [] |
| 6344 | 22 | ISX | {'ess': 1, 'nonext': 7} | True | 35085692 | TCCAATCTCTATGTTGAAGTAG | [] |
| 6345 | 22 | MB | {'ess': 1, 'nonext': 7} | True | 35607296 | GGGCCTAGCCCTGGAAGCCCA | [] |
| 6346 | 22 | RBFOX2 | {'ess': 1, 'nonext': 7} | True | 35752628 | TAGCGCTAATGATTGGTTCCTG | [] |
| 6347 | 22 | APOL4 | {'ess': 0, 'nonext': 1} | False | 36193074 | ATTGGGCCTAGCCTGCTTTTAG | [] |
| 6348 | 22 | TEX33 | unavailable | False | 36991178 | AAACTAGTACTTGGAGCTCTTG | [] |
| 6349 | 22 | KCTD17 | {'ess': 0, 'nonext': 7} | True | 37061791 | CCTAACTTTCTTCTCCTCTCAG | [] |
| 6350 | 22 | IL2RB | {'ess': 1, 'nonext': 7} | True | 37128095 | TCTGTCTACACCAAGTGAGTTG | [] |
| 6351 | 22 | SSTR3 | {'ess': 0, 'nonext': 4} | True | 37206546 | AGGCCCCTACAGGTAGCTGAT | [] |
| 6352 | 22 | RAC2 | {'ess': 1, 'nonext': 6} | True | 37226672 | TTACCCCTAGAGGAGGCTGCA | [] |
| 6353 | 22 | GGA1 | {'ess': 0, 'nonext': 7} | True | 37632710 | TCTGTTCTAGAGGCTACCCCA | [] |
| 6354 | 22 | TRIOBP | {'ess': 0, 'nonext': 7} | True | 37772761 | CCACCTCTACTCAGCCAGGCT | [] |
| 6355 | 22 | ANKRD54 | {'ess': 0, 'nonext': 7} | True | 37831942 | CTCTTGCTACCTCTTCTCCATG | [] |
| 6356 | 22 | MICALL1 | {'ess': 1, 'nonext': 7} | True | 37926196 | ACGATACCTACTCACAGGTTTG | [] |
| 6357 | 22 | C22orf23 | {'ess': 0, 'nonext': 7} | True | 37944314 | GTGAGCTAGAGCCTGACCCCT | [] |
| 6358 | 22 | MAFF | {'ess': 0, 'nonext': 7} | True | 38214877 | GGGCACTAGGAGCAGGAGGCC | [] |
| 6359 | 22 | TMEM184 | {'ess': 0, 'nonext': 7} | True | 38221468 | CGCACCTAGAATTCATCATCAG | [] |
| 6360 | 22 | DMC1 | {'ess': 0, 'nonext': 7} | True | 38509292 | CCGTTAGCTATTCTTCTCGATG | [] |
| 6361 | 22 | DMC1 | {'ess': 0, 'nonext': 7} | True | 38520019 | CAATTCACCACCTACTCCTTGG | [] |
| 6362 | 22 | TOMM22 | {'ess': 6, 'nonext': 2} | False | 38683840 | TAACAATCTAGATCTTTCCAGG | [] |
| 6363 | 22 | GTPBP1 | {'ess': 0, 'nonext': 7} | True | 38735314 | CACATCTAACCAGGCCCTGAT | [] |
| 6364 | 22 | SUN2 | {'ess': 0, 'nonext': 7} | True | 38736266 | GTAAGCAGGGCTAGTGGGCGG | [] |
| 6365 | 22 | APOBEC3 | {'ess': 0, 'nonext': 7} | True | 39044216 | TGGCCCTCTAGGGTCTGCCTT | [] |
| 6366 | 22 | PDGFB | {'ess': 1, 'nonext': 6} | True | 39225722 | CGATGCCCCTAGGCTCCAAGG | [] |
| 6367 | 22 | SYNGR1 | {'ess': 0, 'nonext': 7} | True | 39377689 | GGAGGCCTACGGCTGTGCCGA | [] |
| 6368 | 22 | TAB1 | {'ess': 2, 'nonext': 5} | False | 39430221 | GCCCTACGGTGCTGTCACCAC | [] |
| 6369 | 22 | TAB1 | {'ess': 2, 'nonext': 5} | False | 39436570 | TCCTTCCTACCCTGGGGTCAG | [] |
| 6370 | 22 | MGAT3 | {'ess': 2, 'nonext': 6} | False | 39488948 | GCAGCTCTAGACTTCCGCCTC | [] |
| 6371 | 22 | MIEF1 | unavailable | False | 39514322 | ACCTGCCCTACGTCTGCAGCA | [] |
| 6372 | 22 | MIEF1 | unavailable | False | 39515372 | CATCCTAGAAAAGCAGGGCCT | [] |
| 6373 | 22 | ATF4 | {'ess': 3, 'nonext': 4} | False | 39522601 | -1 | -1 |
| 6374 | 22 | RPS19BP1 | {'ess': 4, 'nonext': 4} | False | 39529491 | CCAGGGAGCCTAGCTGCCGAA | [] |
| 6375 | 22 | ENTHD1 | {'ess': 0, 'nonext': 7} | True | 39816173 | CCAAACTAGCCTACATGGAAAG | [] |
| 6376 | 22 | ADSL | {'ess': 9, 'nonext': 1} | False | 40366521 | CAACTCTACAGACATAATTCTG | [] |
| 6377 | 22 | MRTFA | unavailable | False | 40411389 | GAGCTACAAGCAGGAATCCCA | [] |
| 6378 | 22 | XPNPEP3 | {'ess': 2, 'nonext': 6} | False | 40860760 | TCCCAGCTACTCGGGAAGCTG | [] |
| 6379 | 22 | RBX1 | {'ess': 8, 'nonext': 0} | False | 40972497 | CTTTTCCTAGTGCCCATACCTG | [] |
| 6380 | 22 | EP300 | {'ess': 7, 'nonext': 1} | False | 41178955 | TGTCTCTAGTGTATGTCTAGTG | [] |
| 6381 | 22 | RANGAP1 | {'ess': 7, 'nonext': 1} | False | 41246602 | TGAGTCTAGACCTTGTACAGCG | [] |
| 6382 | 22 | ZC3H7B | {'ess': 0, 'nonext': 7} | True | 41357428 | CTGGCCCTACTCCCCAGTGGT | [] |
| 6383 | 22 | POLR3H | {'ess': 7, 'nonext': 1} | False | 41529292 | GGGCTAGTTGCTGGTCCACCA | [] |
| 6384 | 22 | CSDC2 | {'ess': 0, 'nonext': 7} | True | 41574894 | CAGCCTAGGAGCCCACGACCT | [] |
| 6385 | 22 | CCDC134 | {'ess': 0, 'nonext': 7} | True | 41825822 | CCAGGGCTATAACTCAGACTGC | [] |
| 6386 | 22 | TNFRSF13 | {'ess': 0, 'nonext': 7} | True | 41925366 | TCCCTGCTATTGTTGCTCAGG | [] |
| 6387 | 22 | CYP2D6 | {'ess': 0, 'nonext': 7} | True | 42126573 | ACCCCATTCTAGCGGGGCACA | [] |
| 6388 | 22 | CYP2D7 | unavailable | False | 42140277 | ACCCCATTCTAGCGGGGCACA | [] |
| 6389 | 22 | NFAM1 | {'ess': 0, 'nonext': 7} | True | 42385160 | CCCATCCTAGAGATTTTCATAG | [] |
| 6390 | 22 | SERHL2 | {'ess': 1, 'nonext': 7} | True | 42574054 | CCAGAGCTACAGCTGGGCTGG | [] |
| 6391 | 22 | POLDIP3 | {'ess': 1, 'nonext': 7} | True | 42597713 | AGCAGTCTAATTACTGGAAAGG | [] |
| 6392 | 22 | ATP5MGL | unavailable | False | 42639971 | CATTCTAGCCAATGATGCCACG | [] |
| 6393 | 22 | TTLL12 | {'ess': 0, 'nonext': 1} | False | 43169007 | GCGAGTGCCTAGACAAGGCAG | [] |
| 6394 | 22 | EFCAB6 | {'ess': 0, 'nonext': 7} | True | 43528852 | GTGTCTACTGGAGGAATGCCC | [] |
| 6395 | 22 | SAMM50 | {'ess': 7, 'nonext': 1} | False | 43996382 | GGCTACAGGAACCTTATCCCA | [] |
| 6396 | 22 | PRR5-ARH | {'ess': 0, 'nonext': 1} | False | 44862594 | GCAACACTAGAGACGTCTTCTG | [] |
| 6397 | 22 | ARHGAP8 | {'ess': 0, 'nonext': 7} | True | 44862594 | GCAACACTAGAGACGTCTTCTG | [] |
| 6398 | 22 | KIAA0930 | unavailable | False | 45197175 | ACGCGGCTAGGTCATCAGGAT | [] |
| 6399 | 22 | FAM118A | {'ess': 0, 'nonext': 7} | True | 45328485 | CCATGCCTGGCTACTTTATTAG | [] |

| | | | | | | | |
|---|---|---|---|---|---|---|---|
| 6400 | 22 | FAM118A | {'exs': 0, 'nonexs': 7} | True | 45336362 | TTCTCTTCTAACTTCCTCTTTG | [] |
| 6401 | 22 | SMC1B | {'exs': 0, 'nonexs': 7} | True | 45344555 | CTGCCCCTAGCGGGACTCTCC | [] |
| 6402 | 22 | FBLN1 | {'exs': 0, 'nonexs': 7} | True | 45564991 | TCCTCACTAGGCATCAAGGTG | [] |
| 6403 | 22 | LOC10798 | unavailable | False | 45877313 | TTTTCCTATTTGTTCCCCGCAG | [] |
| 6404 | 22 | CDPF1 | unavailable | False | 46240676 | ACCTGACTAAGGGGCAGGACT | [] |
| 6405 | 22 | TTC38 | {'exs': 0, 'nonexs': 7} | True | 46280376 | CCCAGGCTACAAGGCCCAGCA | [] |
| 6406 | 22 | GTSE1 | {'exs': 1, 'nonexs': 6} | True | 46329369 | TGATATCTACAAGAAGAGCCTG | [] |
| 6407 | 22 | GRAMD4 | {'exs': 0, 'nonexs': 5} | True | 46677250 | CAATACTAGCTGTCCCCGCCA | [] |
| 6408 | 22 | TBC1D22 | {'exs': 0, 'nonexs': 7} | True | 46978587 | CCAACTATGTATCTATGTAATG | [] |
| 6409 | 22 | TBC1D22 | {'exs': 0, 'nonexs': 7} | True | 47113682 | GTCCCTATTTGGGAGCAGACA | [] |
| 6410 | 22 | C22orf34 | {'exs': 0, 'nonexs': 3} | True | 49623156 | ACAGAAACCTAGACACAGTGAG | [] |
| 6411 | 22 | IL17REL | {'exs': 0, 'nonexs': 5} | True | 49997037 | GGAAGGTCTAGGAAGGCAAAAG | [] |
| 6412 | 22 | MOV10L1 | {'exs': 0, 'nonexs': 7} | True | 50158605 | GGGGACGCTAACTCTCCTATA | ['CTCCTATAATCACATCACAGAG'] |
| 6413 | 22 | TRABD | {'exs': 0, 'nonexs': 7} | True | 50198518 | GCAGTCTCCTACTTGTGCCGG | [] |
| 6414 | 22 | DENND6B | unavailable | False | 50312138 | TTTGTTCTAGGGAGGGCACAAG | [] |
| 6415 | 22 | LOC10798 | unavailable | False | 50432133 | AGGTGCTACTCAAACAGAAGTG | [] |
| 6416 | 22 | LMF2 | {'exs': 0, 'nonexs': 7} | True | 50503390 | ACACAGCTACTTCTTTCGCCGC | [] |
| 6417 | 22 | SYCE3 | unavailable | False | 50551244 | TGGGGCCTACAGCCTTTGCTT | [] |
| 6418 | 22 | MAPK8IP2 | {'exs': 0, 'nonexs': 1} | False | 50610778 | GGCAGGCTACTCCAGGTAGAT | [] |
| 6419 | 22 | RABL2B | {'exs': 0, 'nonexs': 7} | True | 50773562 | CTAAGCTAATGACAAACATCAG | [] |
| 6420 | X | LOC10537 | unavailable | False | 1396391 | CAAGGCTAAGTCCCACGTGCA | [] |
| 6421 | X | ZBED1 | {'exs': 0, 'nonexs': 7} | True | 2488634 | TCCTCGCTACAGGAAGCTGCT | [] |
| 6422 | X | CD99 | {'exs': 0, 'nonexs': 7} | True | 2740803 | ATCTTCTATTTCTCTAAAAGAG | [] |
| 6423 | X | ARSD | {'exs': 0, 'nonexs': 7} | True | 2907976 | GATCTACTACCTTTCTACACAG | [] |
| 6424 | X | ARSD | {'exs': 0, 'nonexs': 7} | True | 2914524 | TTAATGTCTATGCTCTTTTAAG | [] |
| 6425 | X | NLGN4X | {'exs': 0, 'nonexs': 7} | True | 5892816 | GCAAAGCTATACTCTAGTGGTG | [] |
| 6426 | X | VCX3A | {'exs': 0, 'nonexs': 7} | True | 6533744 | TGGCCGTCTACACACTCGGTA | [] |
| 6427 | X | STS | {'exs': 1, 'nonexs': 6} | True | 7350260 | CCAGGCGCTGCTAGCGGCTCA | [] |
| 6428 | X | VCX | {'exs': 0, 'nonexs': 7} | True | 7844015 | TGGCCGTCTACACACTCGGTA | [] |
| 6429 | X | VCX3B | {'exs': 0, 'nonexs': 5} | True | 8466382 | TGGCCGTCTACACACTCGGTA | [] |
| 6430 | X | FAM9A | {'exs': 0, 'nonexs': 7} | True | 8791310 | AAAAACACGGCTAGTTATCAAG | [] |
| 6431 | X | FAM9B | {'exs': 1, 'nonexs': 7} | True | 9025514 | TGTCTAGTTATCAAGTTCACTG | [] |
| 6432 | X | CLCN4 | {'exs': 0, 'nonexs': 7} | True | 10233583 | TTGTTGCTAATTAAACATGATG | [] |
| 6433 | X | FRMPD4 | {'exs': 0, 'nonexs': 7} | True | 12720532 | GTCAAAGCTACACTAGGGGTAG | [] |
| 6434 | X | TLR7 | {'exs': 0, 'nonexs': 7} | True | 12888657 | GAAGGGCTAGACCGTTTCCTT | [] |
| 6435 | X | LOC10537 | unavailable | False | 13085906 | ACAAAGACTAAAACTCTTTCTG | [] |
| 6436 | X | LOC10798 | unavailable | False | 13852997 | CTCCTCTAGGAAGCCGCGCGA | [] |
| 6437 | X | GLRA2 | {'exs': 0, 'nonexs': 7} | True | 14730484 | AGGGCACATCTATTTCTTGTGG | [] |
| 6438 | X | ASB11 | {'exs': 0, 'nonexs': 7} | True | 15283504 | TTAGGACTATTGGTATAGGAGG | [] |
| 6439 | X | PIGA | {'exs': 1, 'nonexs': 7} | True | 15321505 | GCTTCCTTCTACCTGGTTTCAG | [] |
| 6440 | X | PIR | {'exs': 0, 'nonexs': 7} | True | 15385003 | TTCCACTAGTTCCCAATCTTTG | [] |
| 6441 | X | ACE2 | {'exs': 1, 'nonexs': 5} | False | 15561904 | GATTTTTCTAAAAGGAGGTCTG | [] |
| 6442 | X | AP1S2 | {'exs': 1, 'nonexs': 6} | True | 15832950 | CGTATTCTAGTACGCTTTCTTG | ['CTTAAACCCAATGAAACTCTTG'] |
| 6443 | X | GRPR | {'exs': 0, 'nonexs': 7} | True | 16152644 | GTCAATCTAGACATACCGCTCG | [] |
| 6444 | X | MAGEB17 | {'exs': 0, 'nonexs': 5} | True | 16171392 | AGACTTCACTAGGGCTGAAAGG | [] |
| 6445 | X | SYAP1 | {'exs': 0, 'nonexs': 7} | True | 16760358 | GGAACAGCTAATTTTCCTCTTG | [] |
| 6446 | X | REPS2 | {'exs': 0, 'nonexs': 7} | True | 17187390 | GCTTCTAAGGAGGGCCCATTC | [] |
| 6447 | X | NHS | {'exs': 0, 'nonexs': 7} | True | 17732463 | GGCAGTCTATGTTGAACTCTGG | [] |
| 6448 | X | PHKA2 | {'exs': 1, 'nonexs': 6} | True | 18893484 | GAGACCCTATTGCATCTGGCA | [] |
| 6449 | X | SH3KBP1 | {'exs': 0, 'nonexs': 7} | True | 19591076 | CAAGTTCTAACTGCTCACTGTG | [] |
| 6450 | X | CNKSR2 | {'exs': 0, 'nonexs': 7} | True | 21609642 | TACACAGGCTAAGAATCTCCAG | [] |
| 6451 | X | SMPX | {'exs': 0, 'nonexs': 7} | True | 21737562 | TTCCTACTACTGTTCAGCTTTG | [] |
| 6452 | X | PHEX | {'exs': 0, 'nonexs': 7} | True | 22247867 | AAAGTTACTAATTGCACCATTG | [] |
| 6453 | X | PHEX | {'exs': 0, 'nonexs': 7} | True | 22247952 | TCCCAGCTACCAGAGTCGGCA | [] |
| 6454 | X | APOO | {'exs': 0, 'nonexs': 7} | True | 23840341 | GTTTTCTACTTAGTTCCAGGTG | [] |
| 6455 | X | PDK3 | {'exs': 0, 'nonexs': 7} | True | 24536163 | GTCCTCTAGAAAGTTCTATTAG | [] |
| 6456 | X | PCYT1B | {'exs': 0, 'nonexs': 7} | True | 24562084 | GTTGTCCAGCTAGAAGTCTCTG | [] |
| 6457 | X | PCYT1B | {'exs': 0, 'nonexs': 7} | True | 24562292 | CAGCCTCTACTTTTCATCCTCG | ['ATCCTCATCCCCCTCGCTCATG'] |
| 6458 | X | POLA1 | {'exs': 8, 'nonexs': 1} | False | 24845872 | GTGCTTCTAAGACTAAAATCAG | ['CTAAAATCAAAACAATATTGAG'] |
| 6459 | X | MAGEB3 | {'exs': 0, 'nonexs': 7} | True | 30236964 | ACTTCACTAGGCGTGGGAAGA | [] |
| 6460 | X | CXorf21 | {'exs': 0, 'nonexs': 7} | True | 30559449 | TCCTCTCTATGGATTTACATTG | [] |
| 6461 | X | TAB3 | {'exs': 0, 'nonexs': 5} | True | 30834557 | TGGGCACTATCCATACAGTAAG | [] |
| 6462 | X | TAB3 | {'exs': 0, 'nonexs': 5} | True | 30846014 | CCATCTCTAATCTGGTGAGTTG | [] |
| 6463 | X | DMD | {'exs': 0, 'nonexs': 7} | True | 31121918 | GACTTCCTACATTGTGTCCTGG | [] |
| 6464 | X | DMD | {'exs': 0, 'nonexs': 7} | True | 31195564 | CCTACAATCCTTTAAAATCCAG | [] |
| 6465 | X | FAM47A | {'exs': 0, 'nonexs': 7} | True | 34129902 | ACTGCCTAATCTTCTTCCGATG | [] |
| 6466 | X | TMEM47 | {'exs': 0, 'nonexs': 7} | True | 34630312 | GTTCTAGTAGTAGTCTTCATAG | [] |
| 6467 | X | FAM47B | {'exs': 0, 'nonexs': 7} | True | 34944768 | ACCATCTAATCTTCTTTTGATG | [] |
| 6468 | X | CFAP47 | unavailable | False | 35889535 | TACAGTTAATTCTAGATATAGG | [] |
| 6469 | X | FAM47C | {'exs': 0, 'nonexs': 7} | True | 37011517 | ACCATCTAATCTTCTTCCGATG | [] |
| 6470 | X | LANCL3 | {'exs': 0, 'nonexs': 7} | True | 37675812 | AGCCTTCTAAACAAAGACGCTG | [] |
| 6471 | X | OTC | {'exs': 0, 'nonexs': 7} | True | 38411860 | TTAGCAGTCTAAAGAGATGGGG | [] |
| 6472 | X | MED14 | {'exs': 5, 'nonexs': 3} | False | 40651805 | AGTGTCTATGGACGCCCACCA | [] |
| 6473 | X | CASK | {'exs': 2, 'nonexs': 6} | False | 41520419 | AGAGGCCTAATAGACCCAGGA | [] |
| 6474 | X | PPP1R2C | unavailable | False | 42777470 | TCTTCTAAGGGTCGCATGACTG | [] |
| 6475 | X | KRBOX4 | unavailable | False | 46471184 | TGTACCTAATAACAGCACTTCG | ['CAGCACTTCAAAATACATGAGG'] |
| 6476 | X | CHST7 | {'exs': 0, 'nonexs': 7} | True | 46575391 | GGGAGGCTACGTGGCGCCGTC | [] |
| 6477 | X | ARAF | {'exs': 2, 'nonexs': 6} | False | 47565353 | GCATCCCTACCTGGGACCCTG | [] |
| 6478 | X | ARAF | {'exs': 2, 'nonexs': 6} | False | 47571456 | CGGGGCCTAAGGCACAAGGCG | [] |
| 6479 | X | ZNF182 | {'exs': 0, 'nonexs': 7} | True | 47976166 | ATTAGCTCTCTAATGTGCCATG | [] |
| 6480 | X | RBM3 | {'exs': 0, 'nonexs': 7} | True | 48577558 | TGATATACTATAAAATAAGATG | [] |
| 6481 | X | WDR13 | {'exs': 1, 'nonexs': 6} | True | 48605031 | CAGGACCCTACTTCTGCTCCC | [] |
| 6482 | X | SUV39H1 | {'exs': 1, 'nonexs': 6} | True | 48707569 | TAAGGGCTAGAAGAGGTATTTG | [] |
| 6483 | X | SLC35A2 | {'exs': 1, 'nonexs': 6} | True | 48903437 | CGGCTAGGAACCCCTTCACCTT | [] |
| 6484 | X | PRAF2 | {'exs': 0, 'nonexs': 7} | True | 49071868 | GGGCCTAGGATCCAGCCTCCT | [] |
| 6485 | X | PLP2 | {'exs': 0, 'nonexs': 7} | True | 49174693 | TTCGCCTACACCGGGCCATCT | [] |
| 6486 | X | SYP | {'exs': 0, 'nonexs': 7} | True | 49191436 | ACCAGACTACATCTGATTGGAG | [] |
| 6487 | X | PPP1R3F | {'exs': 0, 'nonexs': 7} | True | 49287089 | GAGCCTATGAGAACCAAGCCA | [] |
| 6488 | X | CLCN5 | {'exs': 0, 'nonexs': 7} | True | 50092218 | ATGATTCTAGTTGAAGAGAATG | [] |
| 6489 | X | CCNB3 | {'exs': 0, 'nonexs': 7} | True | 50351702 | GCTGCTAGAGTACCAGGCCCT | [] |
| 6490 | X | DGKK | unavailable | False | 50368939 | AAGGGCTACAGTTGAGATCTCG | [] |
| 6491 | X | KDM5C | {'exs': 2, 'nonexs': 6} | False | 53192743 | TAGAGGCTACCAGGGAGGGAA | [] |
| 6492 | X | IQSEC2 | {'exs': 0, 'nonexs': 7} | True | 53227823 | TCTTCCTACAGCCTTCTTTGAG | [] |
| 6493 | X | IQSEC2 | {'exs': 0, 'nonexs': 7} | True | 53231732 | GAAAACTAAAAAATGTTGATTG | [] |
| 6494 | X | SMC1A | {'exs': 7, 'nonexs': 1} | False | 53380102 | ATACTGCTACTGCTCATTGGGG | [] |
| 6495 | X | FGD1 | {'exs': 1, 'nonexs': 7} | True | 54446108 | AACCCTCTAGGTCTTGTCTCGC | [] |
| 6496 | X | TRO | {'exs': 0, 'nonexs': 7} | True | 54931019 | ACCTCACTAGCCATACGAGAAG | [] |
| 6497 | X | PFKFB1 | {'exs': 1, 'nonexs': 6} | True | 54936268 | ATTGGACTAGATGTACTCTGAG | [] |
| 6498 | X | PAGE2B | {'exs': 0, 'nonexs': 7} | True | 55078797 | TTGCTTGAAACCTATGGTTGCG | [] |
| 6499 | X | PAGE2 | {'exs': 0, 'nonexs': 7} | True | 55092730 | TTGCTTGAAACCTATGGTTGCG | [] |
| 6500 | X | PAGE5 | {'exs': 0, 'nonexs': 7} | True | 55224002 | ACCTATAGTTGCCCTTCACCTG | ['TATAATATAAATATAAAATTAG'] |
| 6501 | X | USP51 | {'exs': 0, 'nonexs': 7} | True | 55486803 | GGTAAGACTAGTCTTTCTCTAG | [] |
| 6502 | X | NBDY | unavailable | False | 56729559 | TTTCTCTACTTCTCCGGAGGAG | [] |
| 6503 | X | SPIN3 | {'exs': 0, 'nonexs': 7} | True | 56994170 | CTAAGATGTTTTTACCAATCG | [] |
| 6504 | X | FAAH2 | {'exs': 0, 'nonexs': 7} | True | 57488931 | GGTCCTCCTAAAACTTTCCTGG | [] |
| 6505 | X | NLRP2B | unavailable | False | 57680122 | CTTGTCTACCTATGTCTGGGGC | [] |
| 6506 | X | AMER1 | unavailable | False | 64189878 | TGATAATAACTACTTGGCTAGG | [] |
| 6507 | X | OPHN1 | {'exs': 0, 'nonexs': 7} | True | 68081782 | GGCTATGGCAATTTCTTAAAAG | [] |
| 6508 | X | EDA | {'exs': 0, 'nonexs': 7} | True | 70035608 | GGGGGAATCTAGGATGCAGGG | [] |
| 6509 | X | OTUD6A | {'exs': 0, 'nonexs': 7} | True | 70063390 | TGGGGCCTACAGGAGACGCGG | [] |
| 6510 | X | PDZD11 | {'exs': 1, 'nonexs': 7} | True | 70287081 | GCTGCAACTTTCTAGTGCACAG | [] |
| 6511 | X | TEX11 | {'exs': 0, 'nonexs': 7} | True | 70529094 | TGCCTAATCTGACTTGCTCCAG | [] |
| 6512 | X | SNX12 | {'exs': 0, 'nonexs': 7} | True | 71061015 | TCCTACTGGCGCACCTTCCCC | [] |
| 6513 | X | NLGN3 | {'exs': 0, 'nonexs': 7} | True | 71170096 | TTGGAGCTATACCCGGGTAGT | [] |
| 6514 | X | ZMYM3 | {'exs': 1, 'nonexs': 6} | True | 71249442 | CTCTAATGCACTACTCCCTCGC | [] |
| 6515 | X | ITGB1BP2 | {'exs': 1, 'nonexs': 7} | True | 71305191 | TGGTGTCACTATTCCCCCATTG | [] |
| 6516 | X | HDAC8 | {'exs': 2, 'nonexs': 5} | False | 72230053 | TCAACTAGACCACATGCTTCAG | [] |
| 6517 | X | HDAC8 | {'exs': 2, 'nonexs': 5} | False | 72474695 | TGCTTCTACAGGCCTGGATTTG | [] |
| 6518 | X | DMRTC1B | {'exs': 0, 'nonexs': 5} | True | 72848374 | GGGTTCCTAGCTCAAGAGGGA | [] |
| 6519 | X | DMRTC1 | {'exs': 0, 'nonexs': 4} | True | 72873451 | GGGTTCCTAGCTCAAGAGGGA | [] |
| 6520 | X | NAP1L2 | {'exs': 1, 'nonexs': 7} | True | 73213109 | TACTCTGCTCTAACGATCAATG | [] |
| 6521 | X | CHIC1 | {'exs': 0, 'nonexs': 7} | True | 73578711 | ATCTCTCTAGAATTTCTTTATG | [] |
| 6522 | X | ZDHHC15 | {'exs': 0, 'nonexs': 7} | True | 75379151 | TGCTATGTTTCCGTTTCCACAG | [] |
| 6523 | X | ZDHHC15 | {'exs': 0, 'nonexs': 7} | True | 75450721 | GGCTAGTTTGAGAACATATATG | [] |
| 6524 | X | MAGEE1 | {'exs': 0, 'nonexs': 7} | True | 76430803 | CATTCCTCTACCTGAAGTTGTG | [] |
| 6525 | X | PGK1 | {'exs': 7, 'nonexs': 1} | False | 78125829 | AAGTACTAAATATTGCTGAGAG | [] |
| 6526 | X | CYSLTR1 | {'exs': 0, 'nonexs': 7} | True | 78272732 | GGTTTAAACTATACTTTACATG | ['CATATTTCTTCTCCTTTTTCTG'] |
| 6527 | X | LPAR4 | {'exs': 0, 'nonexs': 7} | True | 78755981 | ATACCTAAAAGGTGGATTCTAG | [] |
| 6528 | X | TBX22 | {'exs': 0, 'nonexs': 7} | True | 80031110 | ATTGTCTTACTAAAGGTAATGG | [] |
| 6529 | X | POF1B | {'exs': 0, 'nonexs': 7} | True | 85282178 | TAGGGCTAAAAATGCCCAAGTG | [] |
| 6530 | X | TSPAN6 | {'exs': 0, 'nonexs': 7} | True | 1.01E+08 | CCTAAATGTCCTAAAATTAAAG | [] |
| 6531 | X | ARL13A | {'exs': 1, 'nonexs': 7} | True | 1.01E+08 | TCCTTCAACTATGGGCTGATGG | [] |
| 6532 | X | TMEM35A | unavailable | False | 1.01E+08 | TTTCTATGACACCTTCACTTTG | [] |
| 6533 | X | TIMM8A | {'exs': 0, 'nonexs': 7} | True | 1.01E+08 | CCGGCTAGGCCACAGGAACCT | [] |
| 6534 | X | HNRNPH2 | {'exs': 1, 'nonexs': 7} | True | 1.01E+08 | CTACCTAAGCAAGGTTTGACTG | [] |
| 6535 | X | TCEAL2 | {'exs': 0, 'nonexs': 5} | True | 1.02E+08 | GGGACACTACACATAAGGAATG | [] |
| 6536 | X | GPRASP1 | {'exs': 0, 'nonexs': 7} | True | 1.03E+08 | TAACCATTACTAATTTTCTTGG | [] |
| 6537 | X | TCEAL9 | unavailable | False | 1.03E+08 | GGTAAACTACATTAAATAGGGG | [] |
| 6538 | X | TCEAL4 | {'exs': 0, 'nonexs': 7} | True | 1.04E+08 | GGGACACTACACATAAGGAATG | [] |
| 6539 | X | SLC25A53 | {'exs': 0, 'nonexs': 4} | True | 1.04E+08 | TCTGCAGCTAGTCAGTCTTCAG | [] |
| 6540 | X | IL1RAPL2 | {'exs': 0, 'nonexs': 5} | True | 1.06E+08 | TTTTCACTACCAAATATCACTG | [] |
| 6541 | X | SERPINA7 | {'exs': 0, 'nonexs': 7} | True | 1.06E+08 | TCCCAACTACGCTTCCGTTGG | [] |
| 6542 | X | TBC1D8B | {'exs': 0, 'nonexs': 7} | True | 1.07E+08 | CTTTTTCTACTTCTCCCCAATG | ['CCCAATAACACATTGTCCTTGG'] |
| 6543 | X | RIPPLY1 | {'exs': 0, 'nonexs': 7} | True | 1.07E+08 | GGCCTCCTACTTCTCTTCTTCC | ['TTCATCTTCCTCTTCTTCTTCG', 'TTCTTCATCCTCCTGCTCTTCG'] |
| 6544 | X | RBM41 | {'exs': 0, 'nonexs': 7} | True | 1.07E+08 | TTCTAGCTACCACTAATTTCTG | [] |
| 6545 | X | FRMPD3 | {'exs': 0, 'nonexs': 1} | False | 1.08E+08 | CAGATGCCCTACTTAATAAGGG | [] |
| 6546 | X | NCBP2L | {'exs': 0, 'nonexs': 1} | False | 1.08E+08 | GATTGTTCTAGATCTGAGTCTG | [] |
| 6547 | X | VSIG1 | {'exs': 0, 'nonexs': 7} | True | 1.08E+08 | CCACCAGCCTATGCCTTAACC | ['CCTTTTCATCTTCACTTAAGGG', 'TGCCTTAACCACTCCCTTTTCG'] |
| 6548 | X | PSMD10 | {'exs': 0, 'nonexs': 7} | True | 1.08E+08 | CAGGCTAAGTGTCTGAAATCAG | [] |
| 6549 | X | ATG4A | {'exs': 1, 'nonexs': 7} | True | 1.08E+08 | TCCCAGGATTCTACACACTCAG | [] |
| 6550 | X | COL4A6 | {'exs': 0, 'nonexs': 7} | True | 1.08E+08 | GCCACCCTACAGGCTTTTCAT | ['TTTTCATACACACCTGGCAGCG'] |
| 6551 | X | CHRDL1 | {'exs': 0, 'nonexs': 7} | True | 1.11E+08 | GCCTAACAGTGGCCCTTTTCA | [] |
| 6552 | X | PAK3 | {'exs': 1, 'nonexs': 6} | True | 1.11E+08 | GATACTACCATATAGTTACTTG | [] |
| 6553 | X | ALG13 | {'exs': 3, 'nonexs': 5} | False | 1.12E+08 | AGCTTCTAGTGTCTCAAGGAAG | [] |
| 6554 | X | LOC10537 | unavailable | False | 1.15E+08 | TTCCCCTAGCAGTAGAGGCCC | [] |
| 6555 | X | LONRF3 | {'exs': 1, 'nonexs': 7} | True | 1.19E+08 | CACTCACTAGTTTTGGTTTCGG | [] |
| 6556 | X | SLC25A43 | {'exs': 0, 'nonexs': 7} | True | 1.19E+08 | CTGTTCTAGCTCTCAGGAAGAG | [] |
| 6557 | X | SEPTIN6 | unavailable | False | 1.2E+08 | GGGAATACTATTCAGTACACAG | [] |
| 6558 | X | SEPTIN6 | unavailable | False | 1.2E+08 | AGGTTTCTATAAACCTTGGCAG | [] |
| 6559 | X | UPF3B | {'exs': 0, 'nonexs': 7} | True | 1.2E+08 | CTTTCTACTGCTGAATCTCCAG | [] |
| 6560 | X | RNF113A | {'exs': 4, 'nonexs': 4} | False | 1.2E+08 | GAAACCTAAGTAATGGGAATTG | [] |
| 6561 | X | AKAP14 | {'exs': 0, 'nonexs': 7} | True | 1.2E+08 | GAGTGGTCGGCCTAACTTACC | [] |
| 6562 | X | RHOXF1 | {'exs': 0, 'nonexs': 7} | True | 1.2E+08 | CTAGGGCTAGTCCACGACGAT | [] |
| 6563 | X | ATP1B4 | {'exs': 0, 'nonexs': 7} | True | 1.2E+08 | GCCCTCCTATCTAGTTCATTCG | [] |
| 6564 | X | LAMP2 | {'exs': 0, 'nonexs': 7} | True | 1.2E+08 | TTCTAAAATTGCTCATATCCAG | [] |
| 6565 | X | CUL4B | {'exs': 1, 'nonexs': 6} | True | 1.21E+08 | AACATTCTATGCAATATAGTTG | [] |
| 6566 | X | CT47B1 | {'exs': 0, 'nonexs': 7} | True | 1.21E+08 | CTGCGTCTAGGTCCCTTTGGA | [] |
| 6567 | X | CT47A12 | unavailable | False | 1.21E+08 | CTGCGTCTAGGTCCCTTTGGA | [] |
| 6568 | X | CT47A12 | unavailable | False | 1.21E+08 | TTCTACGAATCTTTACACTTTG | [] |
| 6569 | X | CT47A11 | {'exs': 0, 'nonexs': 3} | True | 1.21E+08 | CTGCGTCTAGGTCCCTTTGGA | [] |
| 6570 | X | CT47A11 | {'exs': 0, 'nonexs': 3} | True | 1.21E+08 | TTCTACGAATCTTTACACTTTG | [] |
| 6571 | X | CT47A10 | {'exs': 0, 'nonexs': 3} | True | 1.21E+08 | CTGCGTCTAGGTCCCTTTGGA | [] |
| 6572 | X | CT47A10 | {'exs': 0, 'nonexs': 3} | True | 1.21E+08 | TTCTACGAATCTTTACACTTTG | [] |
| 6573 | X | CT47A9 | {'exs': 0, 'nonexs': 3} | True | 1.21E+08 | CTGCGTCTAGGTCCCTTTGGA | [] |
| 6574 | X | CT47A9 | {'exs': 0, 'nonexs': 3} | True | 1.21E+08 | TTCTACGAATCTTTACACTTTG | [] |
| 6575 | X | CT47A8 | {'exs': 0, 'nonexs': 3} | True | 1.21E+08 | CTGCGTCTAGGTCCCTTTGGA | [] |
| 6576 | X | CT47A8 | {'exs': 0, 'nonexs': 3} | True | 1.21E+08 | TTCTACGAATCTTTACACTTTG | [] |
| 6577 | X | CT47A7 | {'exs': 0, 'nonexs': 3} | True | 1.21E+08 | CTGCGTCTAGGTCCCTTTGGA | [] |
| 6578 | X | CT47A7 | {'exs': 0, 'nonexs': 3} | True | 1.21E+08 | TTCTACGAATCTTTACACTTTG | [] |
| 6579 | X | CT47A6 | {'exs': 0, 'nonexs': 3} | True | 1.21E+08 | CTGCGTCTAGGTCCCTTTGGA | [] |
| 6580 | X | CT47A6 | {'exs': 0, 'nonexs': 3} | True | 1.21E+08 | TTCTACGAATCTTTACACTTTG | [] |
| 6581 | X | CT47A5 | {'exs': 0, 'nonexs': 3} | True | 1.21E+08 | CTGCGTCTAGGTCCCTTTGGA | [] |
| 6582 | X | CT47A5 | {'exs': 0, 'nonexs': 3} | True | 1.21E+08 | TTCTACGAATCTTTACACTTTG | [] |
| 6583 | X | CT47A4 | {'exs': 0, 'nonexs': 3} | True | 1.21E+08 | CTGCGTCTAGGTCCCTTTGGA | [] |
| 6584 | X | CT47A4 | {'exs': 0, 'nonexs': 3} | True | 1.21E+08 | TTCTACGAATCTTTACACTTTG | [] |
| 6585 | X | CT47A3 | {'exs': 0, 'nonexs': 3} | True | 1.21E+08 | CTGCGTCTAGGTCCCTTTGGA | [] |
| 6586 | X | CT47A3 | {'exs': 0, 'nonexs': 3} | True | 1.21E+08 | TTCTACGAATCTTTACACTTTG | [] |
| 6587 | X | CT47A2 | {'exs': 0, 'nonexs': 3} | True | 1.21E+08 | CTGCGTCTAGGTCCCTTTGGA | [] |
| 6588 | X | CT47A2 | {'exs': 0, 'nonexs': 3} | True | 1.21E+08 | TTCTACGAATCTTTACACTTTG | [] |
| 6589 | X | CT47A1 | {'exs': 0, 'nonexs': 3} | True | 1.21E+08 | CTGCGTCTAGGTCCCTTTGGA | [] |
| 6590 | X | CT47A1 | {'exs': 0, 'nonexs': 3} | True | 1.21E+08 | TTCTACGAATCTTTACACTTTG | [] |
| 6591 | X | GLUD2 | {'exs': 1, 'nonexs': 7} | True | 1.21E+08 | ATCTATGTGAAGGTCACACCAG | [] |
| 6592 | X | GRIA3 | {'exs': 1, 'nonexs': 6} | True | 1.23E+08 | CCCTAGATCTTAACACTCTCTG | [] |
| 6593 | X | XPNPEP2 | {'exs': 1, 'nonexs': 7} | True | 1.3E+08 | GCCTCTAGACACTCCAGCCAA | [] |
| 6594 | X | ZDHHC9 | {'exs': 0, 'nonexs': 7} | True | 1.3E+08 | ATAGGCTACTTCTCAGCTTCAG | [] |
| 6595 | X | ZNF280C | {'exs': 0, 'nonexs': 7} | True | 1.3E+08 | AGGAATCTATTTCAATGAGCAG | [] |
| 6596 | X | SLC25A14 | {'exs': 0, 'nonexs': 7} | True | 1.3E+08 | ACTAGCCTAAAAGGCTTTAATG | ['AATCAAATTTTCCTTCCCATAG'] |
| 6597 | X | ARHGAP3 | {'exs': 1, 'nonexs': 7} | True | 1.31E+08 | AAACATCTAAGGAAAGAAGTAG | [] |
| 6598 | X | MBNL3 | {'exs': 0, 'nonexs': 7} | True | 1.32E+08 | ACTGAGAACTTCTACATCTGTG | [] |
| 6599 | X | HS6ST2 | {'exs': 0, 'nonexs': 7} | True | 1.33E+08 | CTCCCTACTGGGTATCAAACTG | [] |
| 6600 | X | PHF6 | {'exs': 0, 'nonexs': 7} | True | 1.34E+08 | ACGAACTACTTTAAATTTTCTG | [] |
| 6601 | X | PHF6 | {'exs': 0, 'nonexs': 7} | True | 1.34E+08 | TTCATACCTAGTTTCCATTAAG | [] |
| 6602 | X | FAM122B | {'exs': 2, 'nonexs': 6} | False | 1.35E+08 | TAAATAACTATTCATTCATTTG | [] |
| 6603 | X | FAM122C | {'exs': 0, 'nonexs': 7} | True | 1.35E+08 | TGGCTAACACGGTGAAACCCC | [] |
| 6604 | X | FAM122C | {'exs': 0, 'nonexs': 7} | True | 1.35E+08 | TTTGTCTACTAGAACAAATGAG | [] |
| 6605 | X | SMIM10 | unavailable | False | 1.35E+08 | AGTGGCTCTAAATATATACCTG | [] |
| 6606 | X | RTL8B | unavailable | False | 1.35E+08 | TCTCCCGGCCTAGAAGTCCTC | [] |
| 6607 | X | RTL8C | unavailable | False | 1.35E+08 | TCTCCCGGCCTAGAAGTCCTC | [] |
| 6608 | X | RTL8A | unavailable | False | 1.35E+08 | CATCACTAGGAGCCACAGTCT | [] |
| 6609 | X | RTL8A | unavailable | False | 1.35E+08 | TCTCCCGGCCTAGAAGTCCTC | [] |
| 6610 | X | INTS6L | unavailable | False | 1.36E+08 | TTTGCACTAACATGATGATCTG | [] |
| 6611 | X | BRS3 | {'exs': 0, 'nonexs': 7} | True | 1.36E+08 | AAAAGCTAGAATCTGTCCTCTG | [] |
| 6612 | X | VGLL1 | {'exs': 1, 'nonexs': 7} | True | 1.37E+08 | ACTTGACTAAAGATGCTGCAGG | [] |
| 6613 | X | RBMX | {'exs': 7, 'nonexs': 1} | False | 1.37E+08 | TTATCTACTGTGAATCAATCAG | [] |
| 6614 | X | RBMX | {'exs': 7, 'nonexs': 1} | False | 1.37E+08 | TTGTTTCTAGTATCTGCTTCTG | [] |
| 6615 | X | FGF13 | {'exs': 0, 'nonexs': 7} | True | 1.39E+08 | CACTGGCTACGTTGATTCATTG | [] |
| 6616 | X | MCF2 | {'exs': 0, 'nonexs': 7} | True | 1.4E+08 | GGAAAGAGCTACTTGCCATTTG | [] |
| 6617 | X | CXorf66 | {'exs': 0, 'nonexs': 7} | True | 1.4E+08 | TTGAGCTCCTAATCATTTAGGG | [] |
| 6618 | X | CDR1 | {'exs': 0, 'nonexs': 7} | True | 1.41E+08 | CAGAAAAATCTAGATCTTCCAG | [] |
| 6619 | X | SPANXB1 | {'exs': 0, 'nonexs': 3} | True | 1.41E+08 | TGCTACTTTTTAGGTCTTTCAG | [] |
| 6620 | X | LDOC1 | {'exs': 0, 'nonexs': 7} | True | 1.41E+08 | -1 | -1 |
| 6621 | X | SPANXC | {'exs': 0, 'nonexs': 7} | True | 1.41E+08 | TAGCTTCTTGCTACTTTGCAGG | [] |
| 6622 | X | SPANXA1 | {'exs': 2, 'nonexs': 5} | False | 1.42E+08 | TAGCTTCTTGCTACTTTGCAGG | [] |
| 6623 | X | SPANXA2 | {'exs': 2, 'nonexs': 5} | False | 1.42E+08 | TAGCTTCTTGCTACTTTGCAGG | [] |
| 6624 | X | SPANXD | {'exs': 0, 'nonexs': 7} | True | 1.42E+08 | TAGCTTCTTGCTACTTTGCAGG | [] |
| 6625 | X | MAGEC3 | {'exs': 1, 'nonexs': 7} | True | 1.42E+08 | AAGGCACTAGGGATGATACTGG | [] |
| 6626 | X | SPANXN4 | {'exs': 0, 'nonexs': 7} | True | 1.43E+08 | ACTAATTCTGCCCACCATCCTG | [] |
| 6627 | X | SPANXN3 | {'exs': 0, 'nonexs': 7} | True | 1.44E+08 | GTGTGACTAATCCTCCCCACTC | [] |
| 6628 | X | SLITRK4 | {'exs': 0, 'nonexs': 7} | True | 1.44E+08 | ATGACCTAGATCTTGTTCAAAG | [] |
| 6629 | X | SPANXN2 | {'exs': 0, 'nonexs': 4} | True | 1.44E+08 | ACTAGTCCTCCCCACCCTCCT | [] |
| 6630 | X | FMR1NB | {'exs': 1, 'nonexs': 7} | True | 1.48E+08 | TCTCTTGCTACTCGTCACCATG | [] |
| 6631 | X | AFF2 | {'exs': 0, 'nonexs': 7} | True | 1.49E+08 | AACACCCACTACAACAAGTGGG | [] |
| 6632 | X | EOLA1 | unavailable | False | 1.5E+08 | AGCTGAACCTCTATGGATTCTG | [] |
| 6633 | X | HSFX3 | unavailable | False | 1.5E+08 | GCCTTCCTACTCCTCCTCATC | [] |
| 6634 | X | HSFX2 | {'exs': 0, 'nonexs': 4} | True | 1.5E+08 | GCCCTCTATGTGCTCTGGTCT | [] |
| 6635 | X | TMEM185 | {'exs': 0, 'nonexs': 4} | True | 1.5E+08 | TGGCATCTAATCTGGCATTTCG | [] |
| 6636 | X | HSFX1 | {'exs': 1, 'nonexs': 7} | True | 1.5E+08 | GCCCTCTATGTGCTCTGGTCT | [] |
| 6637 | X | HSFX4 | unavailable | False | 1.5E+08 | GCCTTCCTACTCCTCCTCATC | [] |
| 6638 | X | EOLA2 | unavailable | False | 1.5E+08 | AGCTGAACCTCTATGGATTCTG | [] |
| 6639 | X | PRRG3 | {'exs': 0, 'nonexs': 5} | True | 1.52E+08 | TCCCACTACTTGTCAGCGCCA | [] |
| 6640 | X | GABRE | {'exs': 0, 'nonexs': 7} | True | 1.52E+08 | AGCTGGTACCTACAAGTTAAGG | [] |
| 6641 | X | GABRA3 | {'exs': 1, 'nonexs': 7} | True | 1.52E+08 | TATCTACTGTTTGCGGATCATG | [] |
| 6642 | X | GABRQ | {'exs': 0, 'nonexs': 4} | True | 1.53E+08 | ACTGGGGGACTAATACATATGG | [] |
| 6643 | X | ZNF185 | {'exs': 0, 'nonexs': 6} | True | 1.53E+08 | CACCTAGAAGAGCTTCTCATAG | [] |
| 6644 | X | PNMA5 | {'exs': 0, 'nonexs': 7} | True | 1.53E+08 | TCTGAGCCTATGTTTCCCAGGC | [] |
| 6645 | X | PNMA3 | {'exs': 1, 'nonexs': 6} | True | 1.53E+08 | TCCCGAGCCTACTTGGCCTTG | [] |
| 6646 | X | PNMA6A | {'exs': 0, 'nonexs': 3} | True | 1.53E+08 | CCTAGTTATCACATTCCTCCAG | [] |
| 6647 | X | PNMA6F | unavailable | False | 1.53E+08 | CAGAGCCTATTTGCCCGGGGG | [] |
| 6648 | X | ZNF275 | {'exs': 0, 'nonexs': 3} | True | 1.53E+08 | CGTTTCTACTCGTGGTGGGTT | [] |
| 6649 | X | PNMA6E | unavailable | False | 1.53E+08 | AGGAGCCTATTTGCCCTGGGA | [] |
| 6650 | X | HAUS7 | {'exs': 6, 'nonexs': 2} | False | 1.53E+08 | AGACTTTCTATTTCTCCATTAG | [] |
| 6651 | X | LOC11226 | unavailable | False | 1.53E+08 | ACTAGGAGCCCTCCTCCCCCC | [] |
| 6652 | X | BGN | {'exs': 0, 'nonexs': 7} | True | 1.54E+08 | CTGCCTCTACTTTTTGTAGTTG | [] |
| 6653 | X | ATP2B3 | {'exs': 0, 'nonexs': 7} | True | 1.54E+08 | AGCGAGCTACGGAACGCTTTC | ['CTTTCACCACCCGGATCTGGAG'] |
| 6654 | X | DUSP9 | {'exs': 0, 'nonexs': 7} | True | 1.54E+08 | CGGGGCCCTAGGTGGGGGCC | [] |
| 6655 | X | SRPK3 | {'exs': 1, 'nonexs': 7} | True | 1.54E+08 | CCGGGCCTAGGGGTTGAGCCA | [] |
| 6656 | X | IDH3G | {'exs': 0, 'nonexs': 7} | True | 1.54E+08 | CAGCCTAGGCCTCCACGGCCC | [] |
| 6657 | X | IDH3G | {'exs': 0, 'nonexs': 7} | True | 1.54E+08 | GGCGGGCTACAGGAGGCTGCA | [] |
| 6658 | X | L1CAM | {'exs': 1, 'nonexs': 6} | True | 1.54E+08 | ACTCCACTATTCTAGGGCCACC | [] |
| 6659 | X | AVPR2 | {'exs': 1, 'nonexs': 7} | True | 1.54E+08 | GTCAGCCCTAGCCACGGCTAC | ['GGCTACACCCACCTTCCAGAGG'] |
| 6660 | X | NAA10 | {'exs': 7, 'nonexs': 2} | False | 1.54E+08 | CAGGCTCTAGGAGGCTGAGTC | [] |
| 6661 | X | EMD | {'exs': 0, 'nonexs': 7} | True | 1.54E+08 | TGGCTCCCTCTAGAAGGGGTT | [] |
| 6662 | X | TAZ | {'exs': 5, 'nonexs': 3} | False | 1.54E+08 | CAAGGCCTATCTCCCAGGCTG | [] |
| 6663 | X | UBL4A | {'exs': 0, 'nonexs': 7} | True | 1.54E+08 | CCGAGAATTCTATTTGGAGAAG | [] |
| 6664 | X | FAM3A | {'exs': 0, 'nonexs': 7} | True | 1.55E+08 | CTGGCCGTGCTAGCTGGCCGT | [] |
| 6665 | X | IKBKG | {'exs': 2, 'nonexs': 3} | False | 1.55E+08 | GCCCTACTCAATGCACTCCATC | [] |
| 6666 | X | CTAG2 | {'exs': 0, 'nonexs': 7} | True | 1.55E+08 | GTCGGCTAAATGTGAGGGGCA | [] |
| 6667 | X | DKC1 | {'exs': 9, 'nonexs': 0} | False | 1.55E+08 | GCCTTCACTACTCAGAAACCAG | ['GAAACCAATTCTACCTCTTTTG'] |
| 6668 | X | H2AB1 | unavailable | False | 1.55E+08 | CAGAAGCTAGTCCTCGCCAGG | [] |
| 6669 | X | RAB39B | {'exs': 0, 'nonexs': 7} | True | 1.55E+08 | AAAGAACTGACTAGCACAAACG | ['CAAACATCTCCTCTCTGATTTG'] |
| 6670 | X | CLIC2 | {'exs': 0, 'nonexs': 7} | True | 1.55E+08 | CTCTCCTAACTCTTCTGTTTAG | [] |
| 6671 | X | H2AB2 | unavailable | False | 1.55E+08 | CAGAAGCTAGTCCTCGCCAGG | [] |
| 6672 | X | H2AB3 | unavailable | False | 1.55E+08 | CAGAAGCTAGTCCTCGCCAGG | [] |
| 6673 | X | VAMP7 | {'exs': 0, 'nonexs': 7} | True | 1.56E+08 | TCTTTCCTATTTCTTCACACAG | [] |
| 6674 | X | IL9R | {'exs': 1, 'nonexs': 6} | True | 1.56E+08 | GGACCTAGAATGTCCAGGACC | [] |
| 6675 | Y | LOC10537 | unavailable | False | 1396391 | CAAGGCTAAGTCCCACGTGCA | [] |
| 6676 | Y | ZBED1 | {'exs': 0, 'nonexs': 7} | True | 2488634 | TCCTCGCTACAGGAAGCTGCT | [] |
| 6677 | Y | CD99 | {'exs': 0, 'nonexs': 7} | True | 2740803 | ATCTTCTATTTCTCTAAAAGAG | [] |
| 6678 | Y | SRY | {'exs': 0, 'nonexs': 5} | True | 2786988 | ATTGTCCTACAGCTTTGTCCAG | [] |
| 6679 | Y | TGIF2LY | {'exs': 0, 'nonexs': 5} | True | 3579801 | GGGCTATTCCGGTGAGCTTCT | [] |
| 6680 | Y | UTY | {'exs': 0, 'nonexs': 5} | True | 13249785 | AACAGCTACAAAACCAGTGGTG | [] |
| 6681 | Y | NLGN4Y | {'exs': 0, 'nonexs': 5} | True | 14841261 | GAAAAGCTATACTCTAGTGGTG | [] |
| 6682 | Y | RBMY1B | {'exs': 0, 'nonexs': 5} | True | 21526052 | CTATTCCTCTAAAAAATAAACG | [] |
| 6683 | Y | RBMY1D | {'exs': 0, 'nonexs': 5} | True | 21879808 | CTATTCCTCTAAAAAATAAACG | [] |
| 6684 | Y | RBMY1E | {'exs': 0, 'nonexs': 5} | True | 21903350 | CTATTCCTCTAAAAAATAAACG | [] |
| 6685 | Y | RBMY1F | {'exs': 1, 'nonexs': 4} | False | 22169045 | GTATTACTATATGCGTGGCAGG | [] |
| 6686 | Y | VAMP7 | {'exs': 0, 'nonexs': 7} | True | 57128670 | TCTTTCCTATTTCTTCACACAG | [] |
| 6687 | Y | IL9R | {'exs': 1, 'nonexs': 6} | True | 57190928 | GGACCTAGAATGTCCAGGACC | [] |
